# Unsupervised clustering of track-weighted dynamic functional connectivity reveals white matter substrates of functional connectivity dynamics

**DOI:** 10.1101/2021.12.04.471233

**Authors:** Gianpaolo Antonio Basile, Salvatore Bertino, Victor Nozais, Alessia Bramanti, Rosella Ciurleo, Giuseppe Pio Anastasi, Demetrio Milardi, Alberto Cacciola

**Author notes:** **Correspondence** Alberto Cacciola, MD Brain Mapping Lab, Department of Biomedical, Dental Sciences and Morphological and Functional Images, University of Messina, Messina, Italy, Gianpaolo Antonio Basile, MD Brain Mapping Lab, Department of Biomedical, Dental Sciences and Morphological and Functional Images, University of Messina, Messina, Italy.

## Abstract

The contribution of structural connectivity to functional connectivity dynamics is still far from being fully elucidated. Herein, we applied track-weighted dynamic functional connectivity (tw-dFC), a model integrating structural, functional, and dynamic connectivity, on high quality diffusion weighted imaging and resting-state fMRI data from two independent repositories. The tw-dFC maps were analyzed using independent component analysis, aiming at identifying spatially independent white matter components which support dynamic changes in functional connectivity. Each component consisted of a spatial map of white matter bundles that show consistent fluctuations in functional connectivity at their endpoints, and a time course representative of such functional activity. These components show high intra-subject, inter-subject, and inter-cohort reproducibility. We provided also converging evidence that functional information about white matter activity derived by this method can capture biologically meaningful features of brain connectivity organization, as well as predict higher-order cognitive performance.

## Introduction

Functional co-activation of brain regions, as measured by resting-state functional MRI (rsfMRI), has long been employed to identify spatially segregated patterns of brain activity ^1^. During the last decades, neuroscience has seen a paradigm shift from a traditional, localizationist view of functional brain organization to a network-based perspective, in which different brain regions, which are frequently engaged together during the execution of complex tasks, tend to show correlated intrinsic activity in awake rest, the so-called resting state ^2, 3^. Inter-individual differences in activity and configuration of such intrinsic connectivity networks have been shown to reflect differences in perception, cognition and behavior ^4, 5^.

Recently, this paradigm has been further expanded by incorporating evidence for time-varying fluctuations in functional connectivity strength across brain regions ^6^. This dynamic functional connectivity approach has been employed to parcellate brain into brain networks akin to those identified by static functional connectivity ^7^, as well as to identify transitions from different states of brain activity in the resting state ^8^, and to predict inter-individual variability in age and cognition ^9, 10^. Structural connectivity, resulting either from direct or indirect axonal connections between brain regions, is thought to represent the anatomical substrate of such functional organization. At the current state-of-art, diffusion weighted imaging (DWI) and tractography are instruments of choice for the study of structural connectivity in the human brain in-vivo and non-invasively ^11–14^. Multi-modal approaches integrating tractography with resting-state fMRI have demonstrated a general agreement between structural and functional connectivity and between structural and functional brain networks ^15–17^. Notwithstanding, the contribution of structural connectivity to functional connectivity dynamics is still far from being fully elucidated.

The peculiar spatial organization of functional connectivity is thought to stem in part from the underlying anatomy of white matter circuits, so that some of the system-level properties of functional networks can be explained by the underlying structural connectivity ^15^. At the same time, the intrinsic organization of structural and functional connectivity are expected to diverge, as functional connectivity investigates neuronal activity at a very different time scale compared to synaptic activity and is not constrained by the implicit anatomy of long and short-range neuronal connections. As a clear example of this mismatch, a recent work correlating structural brain networks to functional brain networks found that multiple white matter components accounted for the spatial distribution of each functional brain network ^17^. In this perspective, while most of the existing works analyze functional and structural connectivity data separately, a joint decomposition of both structural and functional information may provide the opportunity of investigating structure-function relationships in a more intuitive fashion.

In addition, evidence suggests that white matter connections have a crucial role in driving and modulating synchronization between brain regions ^18, 19^ which is probably reflected by dynamic fluctuations in brain connectivity ^6^. Hence, incorporating the dynamic connectivity paradigm into this framework provides a simple and natural model to investigate the contribution of structural connectivity in shaping context-dependent fluctuations in functional connectivity in the human brain. As a member of the track-weighted imaging “family” ^20–22^, track-weighted dynamic functional connectivity (tw-dFC) has been recently developed to allow for a joint analysis of structural and dynamic functional connectivity data ^23^. By integrating tractography and dynamic functional connectivity information into a unified framework, tw-dFC represents a powerful tool to investigate the relationship between structure and function, as the structural constraints imposed by mapping dynamic functional connectivity on tractography-derived priors provide a solution to the high dimensionality of functional connectivity data.

In the present work, we applied this framework on high spatial and temporal resolution DWI and resting state fMRI (rs-fMRI) data from the Human Connectome Project (HCP) repository ^24^. The resulting tw-dFC maps were analyzed using an independent component analysis (ICA) at different dimensionality levels, aiming at identifying consistent, spatially independent white matter components which support dynamic changes in functional connectivity. We demonstrated that such components are stable and conserved across different dataset both using a test-retest, a split-half approach and data from an independent repository (Leipzig Study for Mind-Body-Emotion Interactions, LEMON) ^25^. In addition, we provide converging evidence that functional information about white matter activity derived by this method can be employed to capture biologically meaningful features of brain connectivity organization, as well as to predict higher-order cognitive performance.

## Results

For each subject, whole-brain tractograms derived from tractography and preprocessed rs-fMRI time series were combined to generate a 4-dimensional tw-dFC dataset with the same spatial and temporal resolution of the original fMRI time series. In this framework, each white matter voxel’s time series reflects the dynamic changes in functional connectivity occurring at the endpoints of the white matter pathways traversing that voxel.

### ICA-based parcellation of tw-dFC reveals white matter networks, sub-networks, and functional units

The obtained tw-dFC volumes were analyzed using a spatial group ICA that resulted in a series of well-recognizable, anatomically meaningful patterns of white matter connectivity. Each component consisted of a white matter spatial map, which represents the spatial distribution of white matter bundles which show consistent fluctuations in functional connectivity at their endpoints, and a time course that is representative of the dynamic connectivity fluctuations occurring along these tracts. ICA decomposition was performed at three different dimensionality levels, by selecting a different number of components (n) for each run: a first run with n=10 (ICA_10_), to reveal large-scale networks; a second run with n=20 (ICA_20_) which has been commonly used empirically to identify consistent resting-state networks ^7, 26^ and a third run with n=100 (ICA_100_) to obtain a more fine-grained parcellation. For each of these three group ICA runs, and both for the principal dataset (HCP) and the validation dataset (LEMON), peak activation for each independent component (IC) was localized in the white matter; all the components’ mean power spectra show higher low-frequency spectral power and for all the reconstructed components there was no or minimal overlap with known vascular, ventricular, and meningeal sources of artifacts (Supplementary File 1). To ensure stability of estimation, each ICA algorithm was repeated 20 times in ICASSO ^27^. On the principal datasets, the cluster stability/quality index (I_q_) over 20 ICASSO runs was very high (> 0.9) for all the components. Similar results were obtained on the validation dataset, except for the n=100 run, where nearly all components obtained moderate-to-high I_q_ values (>0.8) (Supplementary Figure 1).

For the lower-dimensionality ICA_10_, the resulting components mostly consisted of white matter pathways linking nodes of well-known gray matter functional networks derived from rs-fMRI literature: default mode network (IC3), lateral (IC5) and medial (IC1) sensorimotor network, right (IC6) and left (IC7) frontoparietal network, lateral (IC9) and medial (IC9) visual network and auditory network (IC4) (Figure 2).

**Figure 1.**
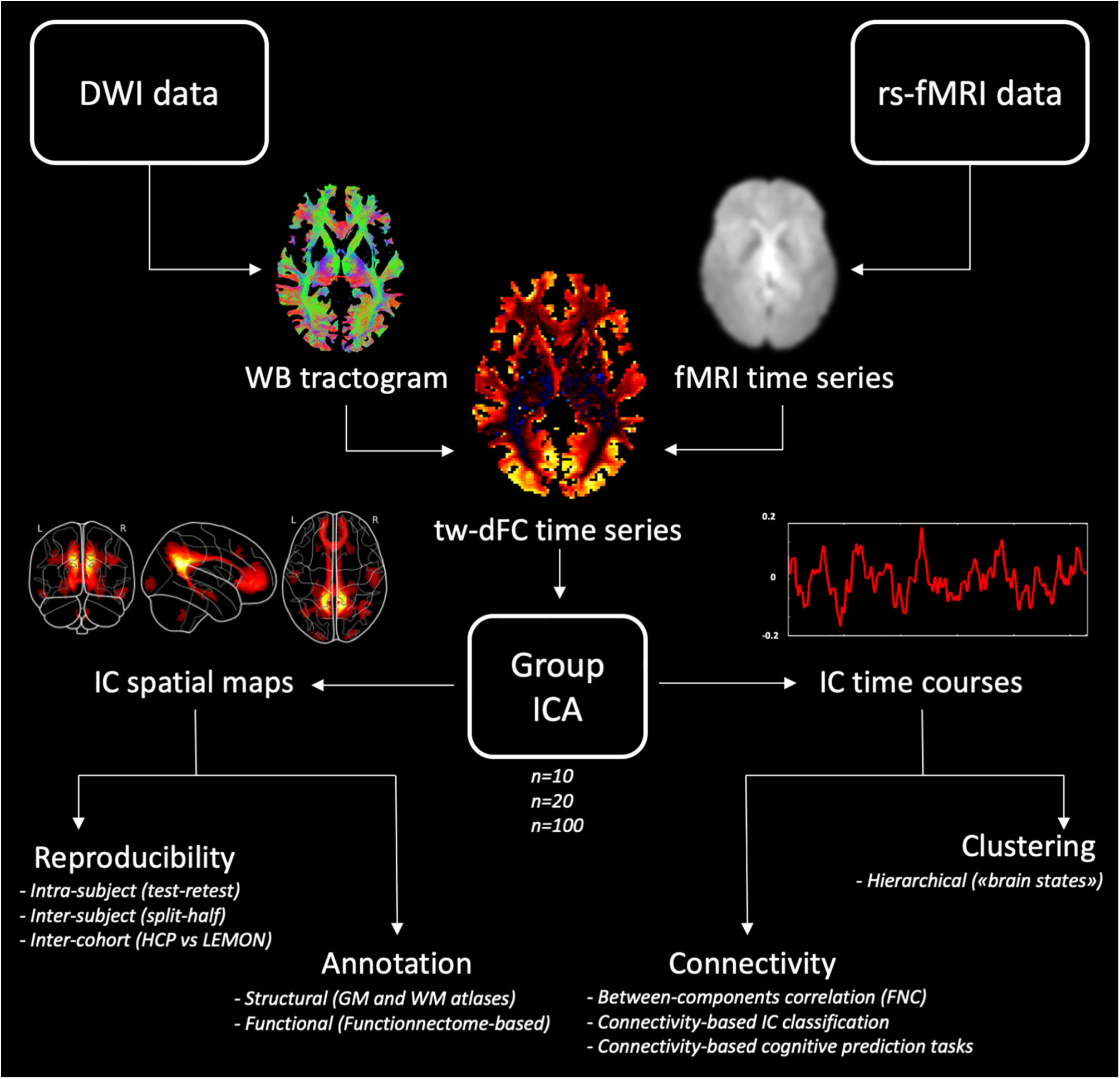
Overview of the workflow. After diffusion-MRI and fMRI preprocessing steps, whole-brain tractography and resting-state fMRI are merged to generate subject-specific track-weighted dynamic functional connectivity (tw-dFC) time series. Independent component analysis (ICA) is then applied at the group level to classify the tw-dFC signal into spatially independent component (IC) maps and their associated time courses, which are the basis for following analyses. In particular, reproducibility of IC spatial maps was evaluated at three different levels on the primary dataset: intra-subject reproducibility (test-retest), inter-subject reproducibility (split-half) and inter-cohort reproducibility (comparison with the validation dataset). IC spatial maps were also annotated by calculating percentage overlap with regions of interest from known GM (Rolls et al., 2020) and WM atlases (Hua et al., 2008; Yeh et al., 2018). IC time courses, instead, were employed to compute between-components correlation (functional network connectivity, FNC), perform connectivity-based IC classification and predict cognitive tasks. Finally, the time series of IC underwent a further clustering analysis aimed at identifying stable or quasi-stable patterns of component activity weights which tend to reoccur over time and across subjects, an analogy to “brain states” described in the dynamic functional connectivity literature.

**Figure 2.**
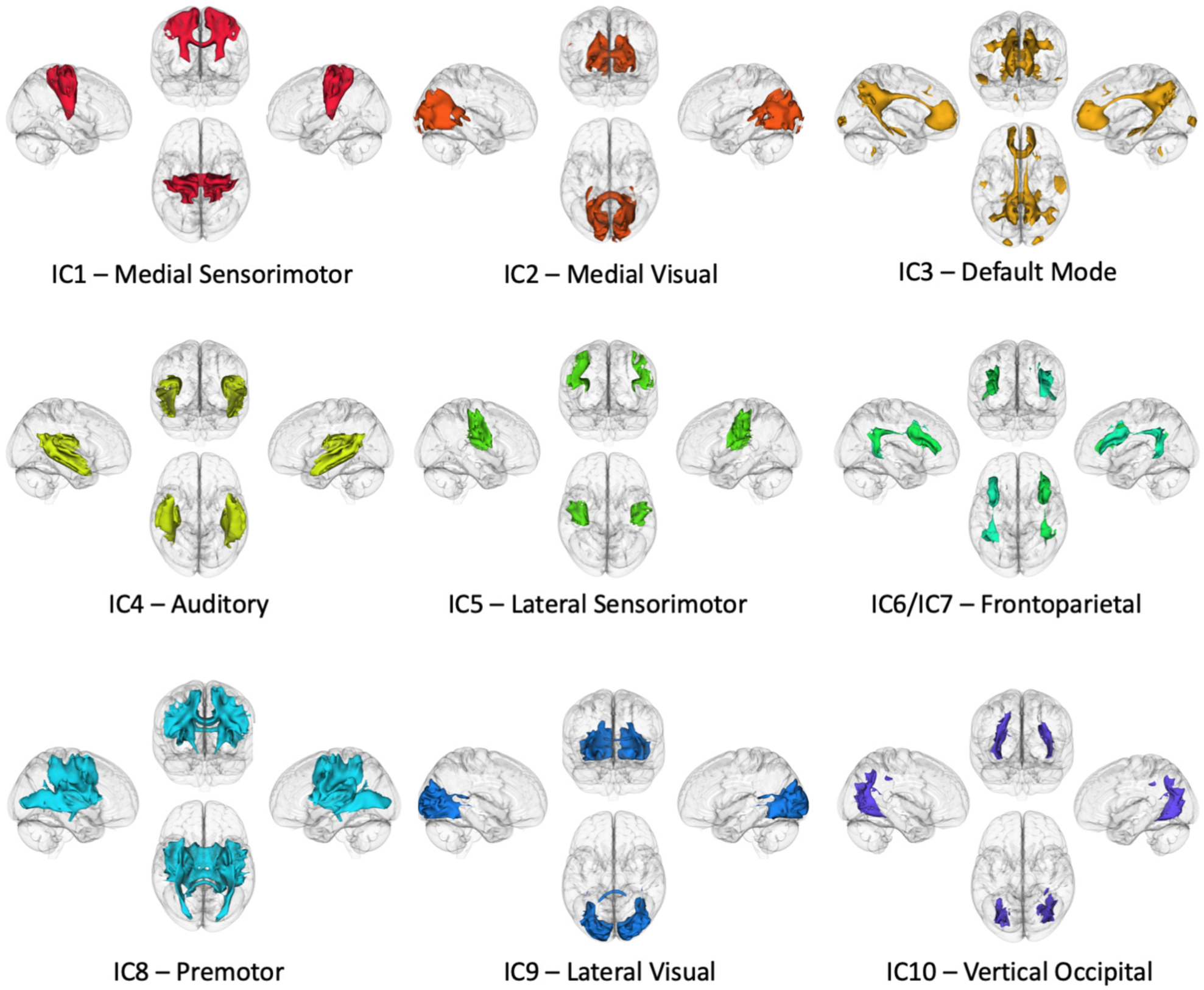
Large-scale networks as identified by low-dimensionality ICA of tw-dFC data. Independent components (ICs) identified by the ICA_10_ on the main dataset reveal white matter structures corresponding to large-scale brain connectivity networks. Group spatial maps for each component are thresholded at z > 1, binarized and volume-rendered on a glass-brain underlay for visualization purposes. Each render shows left, right, superior, and anterior 3D views, along with the putative large-scale network name attributed to each component.

The ICA_20_ parcellation retrieved a more detailed representation of the networks featured in the lower dimensionality ICA run, and some of the networks were split into distinct sub-networks. In addition, along with cortical connectivity networks, a cerebello-cerebellar connectivity component was also identified (IC5). By contrast, the higher-dimensionality ICA_100_ mostly identifies individual white matter bundles (Figure 3A). While most components are bilateral and symmetric, especially from the low-dimensionality ICA runs, some of the ICs from ICA_100_ are lateralized, and many of them show roughly symmetrical, contralateral counterparts (Supplementary Figure 2). In addition, the patterns of connectivity revealed by the higher-dimensionality ICA also highlighted “functional units” corresponding to the somatotopic subdivision of the sensorimotor cortex, to parallel cortico-basal ganglia circuits, as well as to segregated, intrinsic cerebellar connectivity patterns (Figure 3B-C-D). Component spatial maps underwent a group statistical analysis (one sample t-test) and the resulting statistical maps were hard-thresholded at z=1to obtain a white matter parcellation for each ICA run. Percentage overlap between white matter parcellations derived from the ICA_10_, ICA_20_ and ICA_100_ and known white and gray matter components is reported in Supplementary File 2.

**Figure 3.**
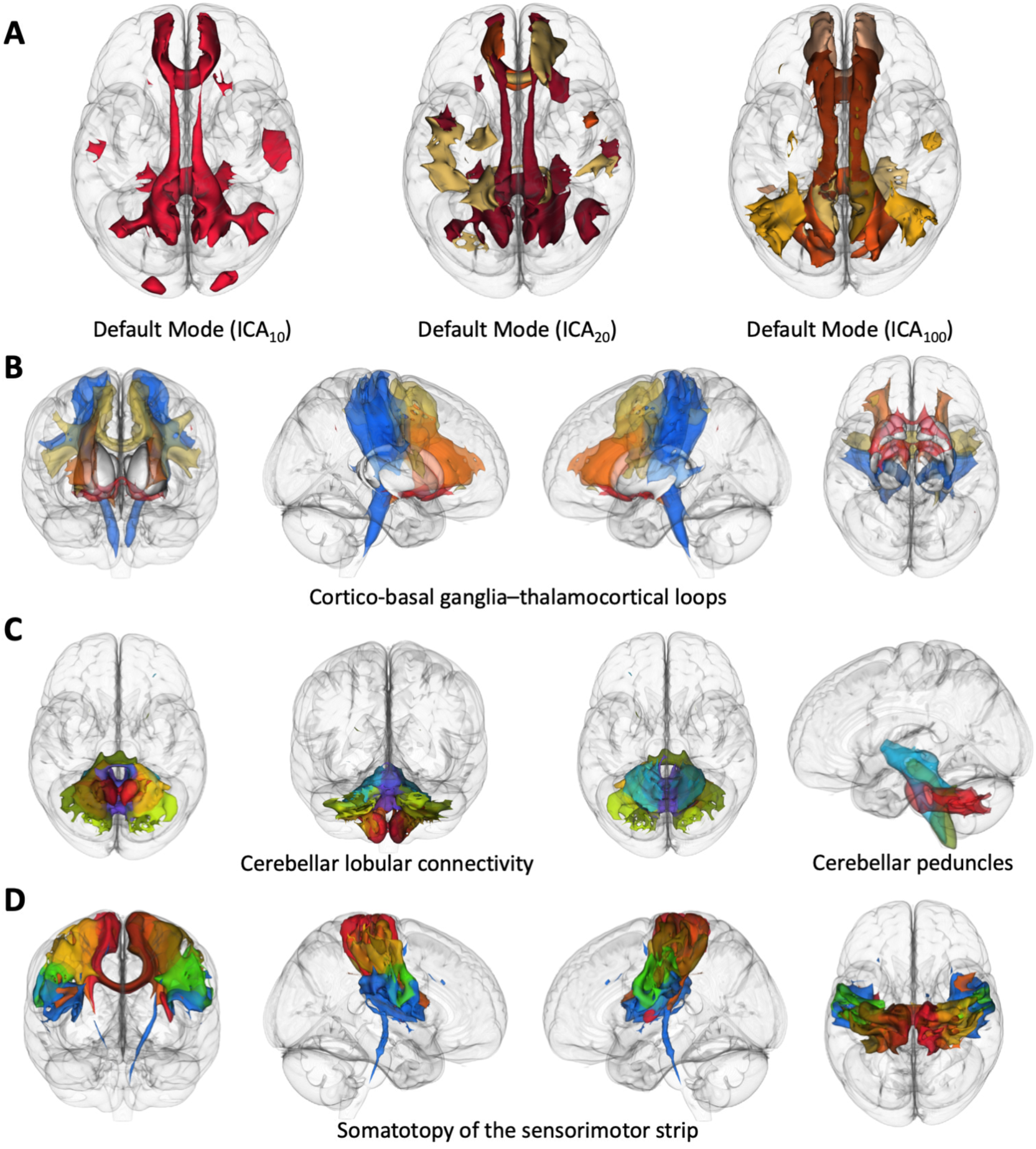
Anatomical details of white matter organization. Group spatial maps for each component are thresholded at z > 1, binarized and volume-rendered on a glass-brain for visualization purposes. **A)** Increasing ICA dimensionality splits large scale white matter networks (e.g. the default mode network white matter component, IC3, n=10) into smaller sub-networks (left anterior, IC1; right anterior; IC18; posterior, IC20, n=20) and increasingly detailed white matter sub-units (ICs 12, 30, 49, 50, 56, 68, 81, n=100). **B)** Cortico-basal ganglia-thalamocortical loops as revealed by ICA_100_ (ICs 72, 84, 38, 42, 87). Fine-grained ICA unsupervised decomposition identifies a ventromedial-orbitofrontal component (red) which extends to the basal forebrain, a ventrolateral component (orange) and dorsolateral (yellow) component involving prefrontal white matter, and two lateralized sensorimotor components (light blue) which also include part of the pyramidal tract. **C)** Intra-cerebellar connectivity networks (ICs 21, 29, 39, 80, 2, 40, 19, 24, 1, 14, 15, 16, 37) are mostly lobule-specific and include distinct cerebellar white matter regions (likely corresponding to cortico-deep nuclear connectivity); superior, middle and inferior cerebellar peduncles are roughly circumscribed by specific components. **D)** Components spanning between the sensorimotor strip (precentral and postcentral gyrus) (ICs 5, 11, 13, 31, 33, 35, 44, 75) roughly reflect the somatotopic organization of primary motor and primary somatosensory cortex.

### White matter components show high intra-subject, inter-subject and inter-cohort reproducibility

The reproducibility of the results was evaluated at three different levels on the primary dataset: intra-subject reproducibility (test-retest), inter-subject reproducibility (split-half) and inter-cohort reproducibility (comparison with the validation dataset).

All the components derived from ICA_10_, ICA_20_ and ICA_100_ were successfully replicated in the test-retest reproducibility analysis, showing very high or moderate-to-high intra-subject similarity. Pearson’s correlation was employed to quantify similarity between group spatial maps. Specifically, the ICA_10_ showed the highest test-retest reproducibility (all components with Pearson r = 1, meaning absolute identity between the paired components from the two datasets), while the ICA_20_ and ICA_100_ showed a decreasing trend (ICA_20_: median Pearson r = 0.97, IQR = 0.98 – 0.83; ICA_100_: median Pearson r = 0.95, IQR = 0.97 – 0.91). Reproducibility was evaluated for the ICA-derived white matter hard parcellations as well, using Dice similarity coefficient (DSC) as a reproducibility measure. We found a similar trend for the corresponding white matter parcellations: for the ICA_10_ a DSC value > 0.99 was reached for all components, while lower values were obtained by the ICA_20_ (median DSC = 0.78, IQR = 0.83 – 0.55) and by the ICA_100_ (median DSC = 0.64, IQR = 0.67 – 0.58) (Figure 4A) High reproducibility was shown also for the split-half replicate analysis, with slightly lower values of between-subject similarity; the ICA_10_ obtained the highest correlation between corresponding components (median Pearson r = 0.97, IQR = 0.99 – 0.96), followed by the ICA_20_ (median Pearson r = 0.95, IQR = 0.98 – 0.85) and the ICA_100_ (median Pearson r = 0.92, IQR = 0.95 – 0.87). For the white matter parcellation, a different trend was observed; while the ICA_10_ obtained the highest DSC values between corresponding components (median DSC= 0.88, IQR = 0.90 – 0.85), the ICA_100_ run showed higher spatial overlap (median DSC = 0.81, IQR = 0.86 – 0.75) than the ICA_20_ (median DSC = 0.71, IQR = 0.85 – 0.59) (Figure 4B).

**Figure 4.**
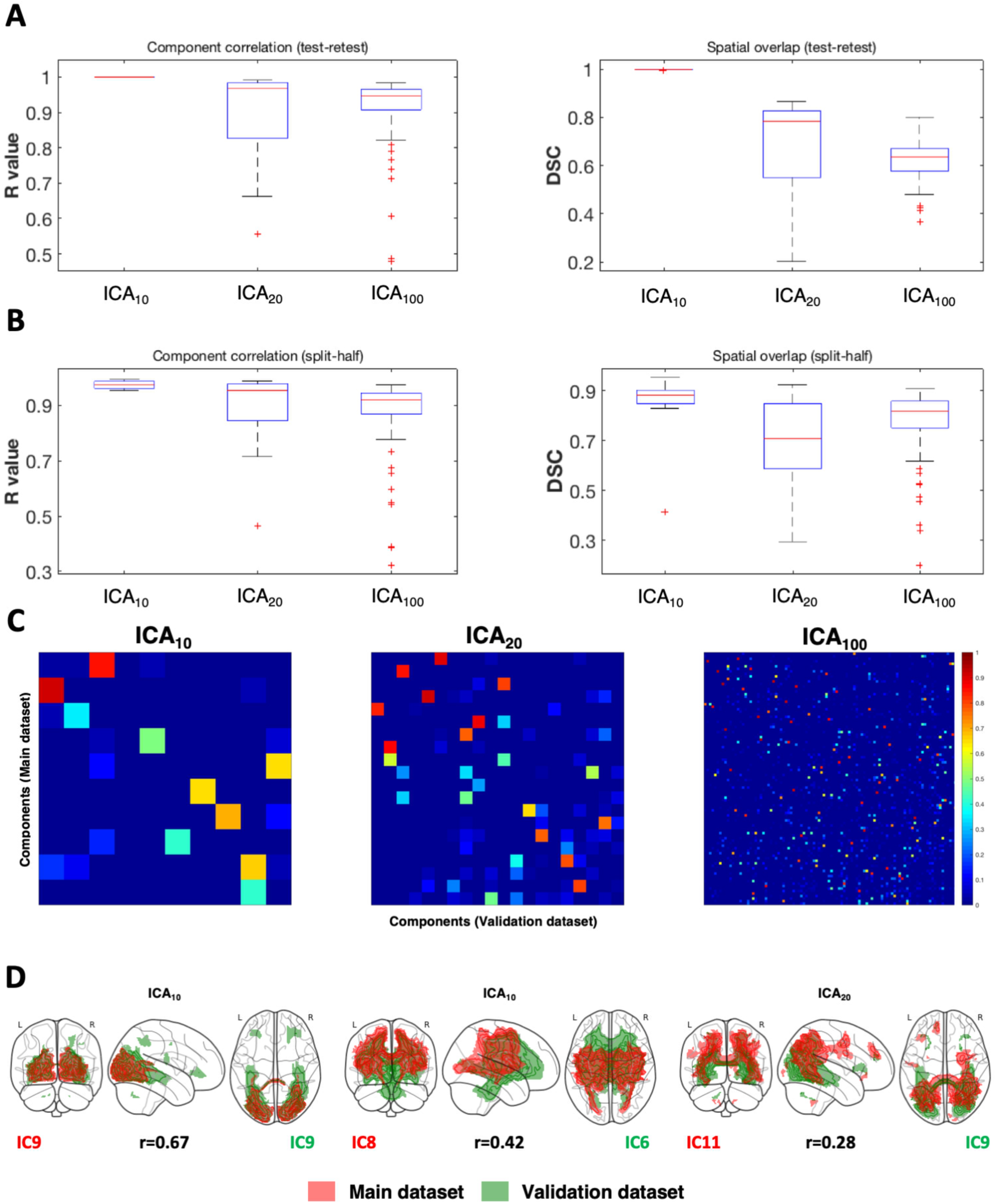
Internal and external reproducibility analysis. Box plots showing the distribution of Pearson’s correlation values (left) and Dice similarity coefficients (right) between corresponding components from the test-retest **(A)** and split-half internal reproducibility analysis **(B). C)** Matrix plots of the pairwise Pearson’s correlation coefficient between the main (HCP) and the validation dataset (LEMON). The colormap allows distinguishing between pairs of components with high correlation (red/yellow), intermediate correlation (green/turquoise) and low correlation (blue/light blue). **D)** A visual example of overlaps between spatial maps of the two independent datasets at different levels of correlation (high, left; moderate, center; low, right). Group spatial maps are thresholded at z > 1, binarized and shown in form of 2D maximum intensity projections on a glass brain in axial, sagittal and coronal sections; L=left; R=right.

Finally, many of the components resulting from the primary dataset (HCP) were totally or partially replicated in the validation dataset (LEMON). To ease with interpretation of the results, we subdivided correlation values into three ranks: high correlation (r > 0.66), moderate correlation (0.33 < r > 0.66) low or absent correlation (r < 0.33). For the ICA_10_, 4 components showed high correlation (40%), 5 components (50%) showed moderate correlation and 1 component (10%) showed low or absent correlation. For the ICA_20_, 12 components (60%) showed high correlation, 4 components (20%) showed moderate correlation and 4 components showed low or absent correlation. Finally, for the ICA_100_, 74 components (74%) showed high correlation, 21 components (21%) showed moderate correlation and 5 components (5%) showed low or absent correlation (Figure 4C). Due to the heterogeneity of the results between the two datasets, DSC between paired components from the two datasets was not calculated. However, corresponding components were visually inspected to check for similar coverage of cortical, subcortical, and white matter regions. Visual inspection confirmed that highly correlated components covered roughly the same cortical, subcortical, and white matter regions (i.e., they have substantially the same anatomical meaning). Components showing moderate correlation were anatomically distinct, but shared some degrees of overlap, while components with low or absent correlation were completely distinct (Figure 4D).

### Resting-state and task-based white matter networks are correlated together

Given that resting-state patterns of brain activity are often related to regional coactivation during behavioral tasks ^2^, we sought to investigate the involvement of resting-state white matter dynamic connectivity components in task-modulated activity by correlating them with distinct, task-dependent white matter networks obtained from track-weighted, task-based fMRI data (“Functionnectome”). In brief, the Functionnectome algorithm maps the function signal from fMRI to tractography-derived priors of white matter anatomy. It has been applied to task-based fMRI data from the HCP dataset to obtain group task-based activation maps for the HCP motor, working memory and semantic language tasks ^28^. As expected, we found high correlation between resting-state and task-based white matter networks. As an example, for the lowest dimensionality ICA run (ICA_10_), components covering sensorimotor regions (IC1, IC5 and IC8) showed relatively high correlation (r > 0.30) with the motor task-based functionnectome maps (corresponding to left and right finger tapping and left and right toe clenching). The medial, lateral and vertical occipital visual network components (IC2, IC9, IC10) and default mode network component (IC3) were not strongly correlated to any of the task-based functionnectome maps. The auditory network component IC4 was weakly correlated to the language semantic task-based functionnectome map (r = 0.24). Left and right frontoparietal components (IC6 and IC7) showed higher correlation to the working memory task-based functionnectome map (r = 0.40 both). Generally lower correlation to task-based maps were found for ICA_20_ or ICA_100_, in line with the finding that increasing ICA dimensionality leads to spatially circumscribed components, in contrast with the widespread task-activation maps. However, some components still showed high correlation to task-based maps, suggesting their possible involvement in language, working memory or motor tasks (Supplementary Figure 3).

### Connectivity-based clustering of independent components uncovers their intrinsic functional organization

Within each different dimensionality ICA run, temporal activity of the extracted ICs, which is a measure of component-specific fluctuations in functional connectivity, showed correlation to that of the other components. Pairwise correlations between components of each run were quantified to obtain ICA-specific “functional network connectivity” (FNC) matrices. Note that the interpretation of connectivity values slightly differ from the classical functional network connectivity as obtained in previous works ^8^. Since tw-dFC volumes are already derived from windowed, dynamic functional connectivity, the time series of tw-dFC derived components are a measure of mean functional connectivity fluctuations at the endpoints of the white matter tracts involved in the IC (i.e., *within* each component). Consequently, FNC can be interpreted as a measure of “co-fluctuation” of the mean functional connectivity *between* pairs of components. As expected, functionally correlated components showed anatomical and functional commonalities that were captured by the clustering analysis of FNC. For the ICA_10_, the elbow method suggested an optimal number of clusters of k=3; the following k-means clustering revealed an intrinsic functional organization of ICs into an associative cluster including the default mode, left and right frontoparietal networks (IC3, IC6 and IC7), a sensorimotor cluster covering somatomotor, somatosensory, auditory and premotor regions (IC1, IC4, IC5 and IC8) and a visual cluster which includes lateral and medial visual networks as well as a component covering the vertical occipital fasciculus (IC2, IC9 and IC10) (Figure 5). Clustering analysis from the ICA_20_ (k=5) and ICA_100_ (k=24) highlighted a roughly similar functional organization into associative, sensorimotor and visual clusters (Supplementary File 2; Supplementary Figures 4-6). In particular, clustering obtained from the higher-dimensionality ICA run showed multiple associative, sensorimotor, visual and cerebellar “sub-clusters”, each with distinctive anatomical features, that are likely to represent fine-grained levels of organization of white matter functional activity.

**Figure 5.**
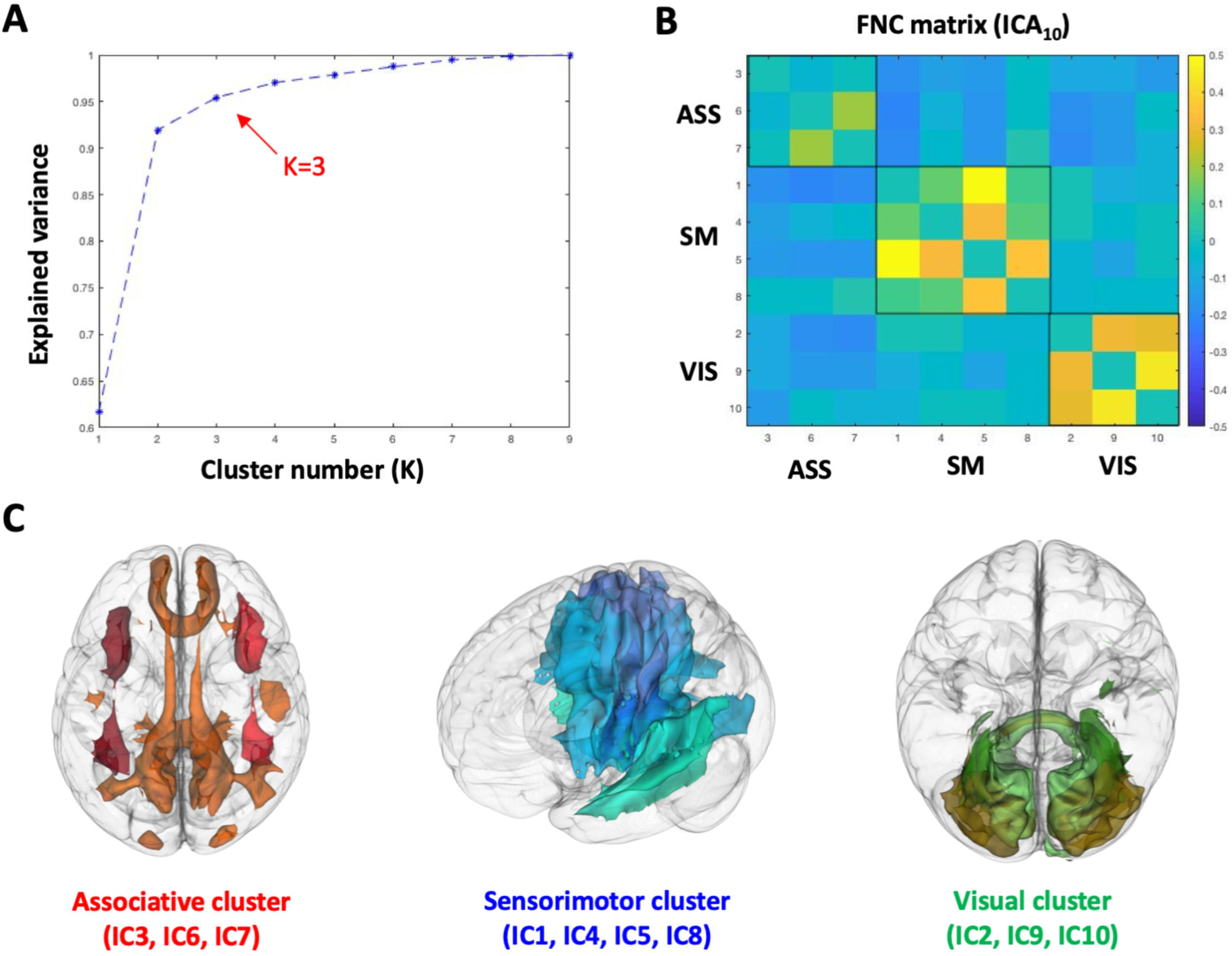
Functional network connectivity (FNC) clustering analysis (ICA10). **A)** Plot of the explained variance for different numbers of clusters; the elbow method (red arrow) suggests an optimal number of k=3. **B)** The FNC matrix, ordered according to the clustering results. Black squares delimitate the three distinct connectivity clusters; ASS = associative, SM = sensorimotor, VIS = visual. **C)** Visualization of the three FNC-derived clusters; group spatial maps for each component are thresholded at z > 1, binarized and volume-rendered on a glass-brain underlay.

### Co-fluctuations of functional connectivity between tw-dFC components predict individual cognitive performance

In order to quantify the role of correlated activity between brain functional units in predicting cognitive performance, we built predictive models based on linear regression using features extracted from individual FNC matrices. In more detail, each feature is constituted by the connectivity strength (Pearson’s correlation coefficient) between a pair of ICs from the high dimensional ICA_100_. Feature selection (based on correlation between individual connectivity and each behavioral score) resulted in 56 features being correlated to fluid intelligence scores (p < 0.01), 44 features being correlated to cognitive flexibility (p < 0.01) and 19 features being correlated to sustained attention (p < 0.01). The following LOOCV-regression model revealed that each behavioral measure could be effectively predicted by the FNC-derived features (Figure 6). In particular, the best prediction resulted for fluid intelligence scores (R^2^ = 0.45, p < 0.0001), where almost 50% of the inter-individual variance in the behavioral scale was explained by the cross-validated regression model. Cognitive flexibility and sustained attention scores were also successfully predicted by FNC-derived features (cognitive flexibility: R^2^ = 0.39, p < 0.001; sustained attention: R^2^ = 0.25, p <0.001).

**Figure 6.**
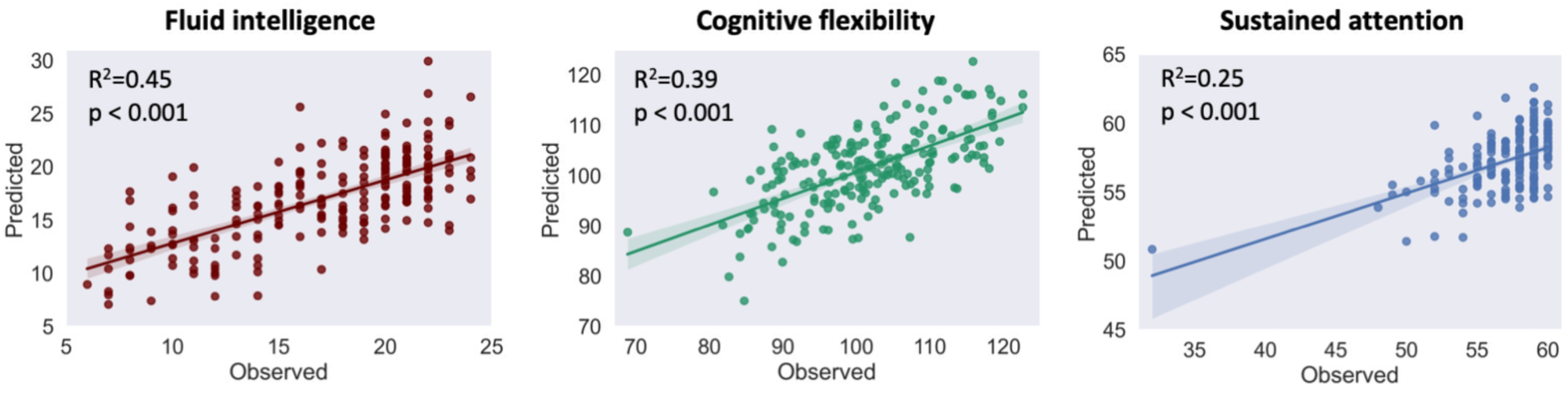
Cognitive performance prediction tasks. Observed vs predicted plots for cognitive performance scores of fluid intelligence (Penn Progressive Matrices, age adjusted), cognitive flexibility (Dimensional Change Card Sort), and sustained attention (Short Penn Continuous Performance Test). Each subject’s score is represented by a dot and the shaded area around the least square lines is the 95% confidence interval.

### Hierarchical clustering of tw-dFC time windows identifies transient “brain states”

Dynamic connectivity shows temporally organized patterns of activity between pairs of nodes or connectivity units, often described as transient “brain states” (Allen et al., 2014; Fan et al., 2021). By applying a hierarchical clustering algorithm to tw-dFC time series, we investigated whether a similar information could be obtained from track-weighted connectivity data. For hierarchical clustering of tw-dFC time windows an optimal cluster number of k=11 was identified according to the elbow method of cluster validity index (CVI). The clustering of time series identified 11 recurring temporal patterns of component activity (“brain states”). Note that, due to the dynamic and windowed nature of the tw-dFC signal, the term “activity” in this case refers to high or low functional connectivity *within* components, while the FNC previously analyzed referred to correlated variations of functional connectivity *between* components. In time windows belonging to state 1 connectivity is low in the cerebellar component (IC5), in cingulum (IC1, IC20) and in prefrontal white matter (IC15) and very high in the bilateral temporal white matter (IC19). State 2 shows high connectivity in the visual components, in particular in the medial occipital component (IC3), and low connectivity in associative and sensorimotor components. In state 3, connectivity is high in the sensorimotor components (IC2, IC4, IC7, IC10, IC12), and in temporal white matter (IC19), and low in cingulum, prefrontal and parietal white matter. State 4 is characterized by generalized low connectivity, especially in the occipital (IC3, IC6, IC8 and IC9), temporal (IC19) and prefrontal components, while in state 5, connectivity is high in the medial sensorimotor component IC2 and low in occipital and cerebellar components, and state 6 shows high connectivity in the cerebellum (IC5), prefrontal and lateral motor components. States 7 and 8 show both high connectivity in the default mode components IC1 and IC20, while in state 9 connectivity is high in parietal and prefrontal components (IC1, IC15 and IC18) and low in the temporal component IC19 and in the cerebellum. State 10 and 11 show specular patterns of connectivity (components with high connectivity in state 10 show low connectivity in state 11 and vice-versa) and are both characterized by extremely high or extremely low connectivity throughout almost all the components (Figure 7).

**Figure 7.**
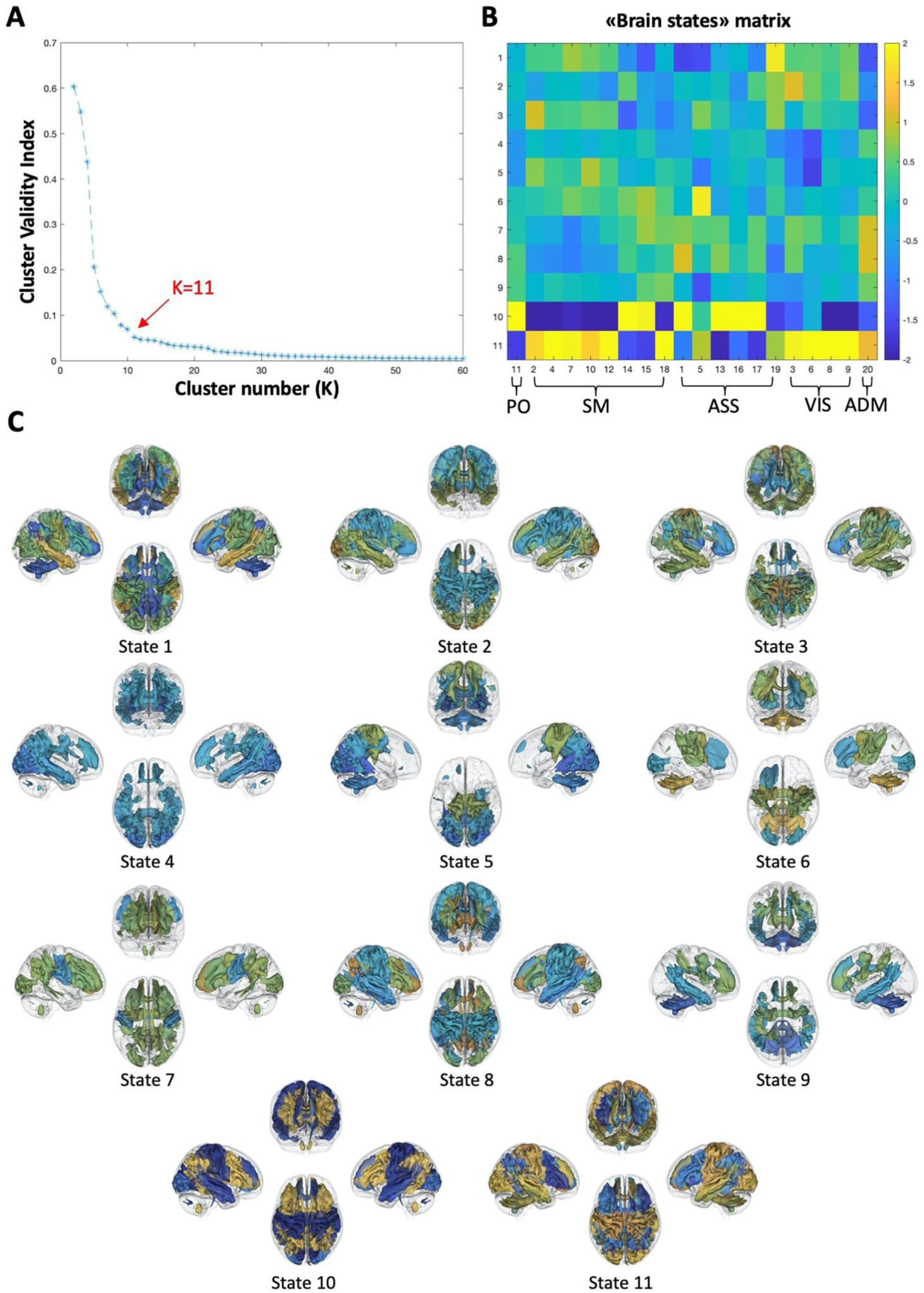
“Brain states” hierarchical clustering. **A)** The elbow criterion of cluster validity index suggests an optimal cluster number k=11 for hierarchical clustering of tw-dFC time windows. **B)** Cluster centroids of transient brain state vectors. Colors are assigned according to the average within-component connectivity value in the time windows corresponding to each state, and for each of the n=20 components. Components are sorted according to FNC-based clustering: PO = parieto-occipital, ASS = associative, SM = sensorimotor, VIS = visual, ADM = anterior default mode. **C)** Clusters visualization. For each state, group spatial maps of independent components are thresholded at z > 1, binarized and volume-rendered on a glass-brain in left, right, anterior, and superior 3D views. For simplicity, only components with average within-component connectivity r > ± 0.5 are displayed. Colormap is the same as in panel B.

## Discussion

In the present work we employed tw-dFC to incorporate structural and dynamic functional connectivity information from a large dataset of human subjects into a unified analysis framework ^23^. In such a framework, the components identified by the ICA process can be interpreted as spatially independent functional units of white matter, composed by fiber bundles sharing coordinated fluctuations of functional connectivity at their endpoints.

On a coarse ICA dimensionality scale, our findings complement and expand current knowledge on resting-state brain networks, as tw-dFC-derived components could be associated with well-known grey matter resting-state networks. Indeed, these ICs mostly consisted of white matter pathways linking nodes of gray matter functional networks as described in rs-fMRI literature during the last decades ^1, 2, 7^. Our results clarify the contribution of anatomical white matter structures to known resting state networks, such as, for instance, distinct anatomical portions of the corpus callosum in the default mode, medial sensorimotor, medial visual and lateral visual networks ^29^, the cingulum bundle in the default mode network ^30^, or the arcuate fasciculi in the left and right frontoparietal networks ^31^. While many other works combined tractography and rs-fMRI to identify the white matter correlates of intrinsic brain connectivity networks ^32^, directly decomposing the tw-dFC signal offers the advantage of identifying joint structural/functional connectivity networks in an unsupervised way. In addition, and in line with the existing literature, we confirmed that there is no one-by-one correspondence between white matter bundles and functional brain networks ^17^, as each tw-dFC large scale component included multiple association, commissural and U-fiber tracts.

As expected from previous works ^7, 17, 33^, increasing ICA dimensionality has led to a more detailed classification of white matter sub-units. At the most detailed dimensionality scale (ICA_100_), ICA decomposition of tw-dFC data reveals a fine-grained, anatomically meaningful functional parcellation of the white matter into long- or short-range connectivity patterns, adding further insights on the functional organization of white matter circuits at the macroscale. As an example, fine-grained ICA was able to reveal subtle anatomical details of white matter connectivity such as the dense, tightly-organized U-fibers system connecting somatotopically analogous regions of precentral and postcentral gyri, the contribution of the cerebellar white matter to intrinsic lobule-specific cerebellar circuitry ^34, 35^, or again, parallel white matter components corresponding to topographically organized cortico-basal ganglia-thalamic circuitry ^36–39^ (Figure 3).

Similarly to well-known gray matter resting-state intrinsic connectivity networks ^2^, white matter dynamic connectivity networks show correspondence to task-based activation networks. To identify task-based activation networks in the white matter, we employed the results obtained using a recently developed method, the “Functionnectome” ^28^. Despite substantial differences, this method shows remarkable analogies to the tw-dFC pipeline as they both involve the resampling of functional information derived from BOLD fMRI on anatomical priors derived by tractography, and the representation of outputs in terms of spatial statistical maps (z-maps), thus enabling direct comparison of results. Taken together, the high correlation values obtained between task-based and resting-state dynamic connectivity provide complementary evidence to the hypothesis that regions intrinsically connected in the resting state are more easily recruited together during tasks ^2, 40, 41^.

In addition, spatial tw-dFC components are highly stable both across subjects of the same sample and when compared to those obtained from a validation dataset (Figure 4). This last result is of utmost importance, as reproducibility is one of the key issues of modern-days neuroimaging research ^42^. Noteworthy, the validation dataset showed several demographical (larger age range, different gender proportion) and technical differences both in DWI (single shell, low b-value, no filtering) and rs-fMRI (lower temporal resolution, different denoising pipeline). Although a proper formal evaluation of the effects of these variables on the results is warranted in future studies, this finding suggests that tw-dFC-based components may be robust to experimental conditions, and, by consequence, able to capture actual features of brain activity and connectivity regardless of technical differences in data processing. While test-retest and split-half reproducibility showed a progressively decreasing trend with the increase of ICA dimensionality, external reproducibility measures seemed to benefit from high dimensionality; this observation would suggest that, while coarse-scale decomposition may be more influenced by group-specific features of the tw-dFC signal, fine-grained ICA would be more sensitive to inter-individual differences, and then less affected by technical and demographic group differences.

Aside from providing an unsupervised and reliable functional parcellation of the human white matter, ICA of tw-dFC data has also the advantage of assigning to each white matter component a time course, which allows for direct investigation of time-varying activity of white matter pathways – a difficult task for state-of-art, conventional neuroimaging ^23, 28^. Although a growing body of works employed direct white matter BOLD signal analysis to accomplish such task ^43–45^, sampling BOLD signal from white matter is challenging due to the much smaller number of blood vessels compared to the gray matter, which implies lower signal-to-noise ratio and lower correlations between seed regions ^43^. In addition, the biological mechanisms behind BOLD signal fluctuations in the white matter are still unclear ^46^. In comparison to existing functional networks obtained by clustering of white matter signal ^44, 45^, components detected from ICA of tw-dFC show some topographical similarities both at coarse and fine-grained scale, but are more adherent to the known anatomy of white matter bundles, as a consequence of incorporating white matter priors from multi-fiber tractography into the processing pipeline. This also allows the tw-dFC signal to account for complex fiber configurations such as crossing, kinking and fanning fibers ^47^, while this is not possible for white matter BOLD-fMRI ^44^.

In addition, our work provided concurrent evidence that the profiles of white matter activity identified by tw-dFC are functionally meaningful.

First, time profiles of white matter connectivity, similarly to what observed after dynamic connectivity analysis in the gray matter ^7, 8^, show peculiar patterns of activity which are recurrent in time (“brain states”). The eleven white matter brain states identified in our work are in line with the corresponding gray matter brain states identified by hierarchical clustering of dynamic functional connectivity ^7^, by showing generally opposite patterns of activity between sensorimotor (visual, auditory, and somatomotor) white matter components (e.g. in states 1, 2, 3, 5 and 11) and associative (default mode, frontoparietal, prefrontal and cerebellar) components (as in states 6, 7, 8, 9 and 10). These distinct sets of states are in line with the notion of “metastates”, which involve preferentially sensorimotor or associative connectivity patterns, and which has been suggested to represent the basis for hierarchical organization of brain functional activity over time ^48^. Further investigation of the temporal structure (e.g. transition probability or fractional time occupancy) of these brain states, could give interesting clues about their relation to inter-individual differences in brain function.

Second, correlation between time series components, as measured by FNC, revealed that dynamic activity in the white matter is coordinated across functionally homogeneous clusters, reflecting similarities in information processing. In the low-level representation of tw-dFC organization derived by ICA_10_, FNC revealed a tripartite segregation into visual unimodal, somatosensory-auditory unimodal and associative transmodal brain regions. Along with reflecting the temporal activity patterns revealed by “brain states” hierarchical clustering, such a segregation is in line with the hierarchical organization model theorized previously by Mesulam ^49^ and recently confirmed by diffusion embedding of functional connectivity ^50^. Moreover, this hierarchical organization is maintained also for increasingly fine-grained decompositions of the tw-dFC signal, where the tripartite model “breaks up” into multiple associative, sensorimotor, and visual clusters, each capturing distinct facets of cortico-cortical and cortico-subcortical information processing. In addition, while the cerebellar involvement is minimal in the ICA_10_, and limited to a single component in the ICA_20_ (IC5, which covers cerebello-cerebellar connections via the middle cerebellar peduncle and is part of the “transmodal” cluster), FNC clustering of the higher-dimensionality ICA_100_ reveals distinct segregated cerebellar connectivity clusters, consisting of components covering intrinsic cerebellar connectivity or the cerebellar peduncles, either alone or in group with cerebral connectivity components. This result fits well with recent investigations postulating the existence of a hierarchical organization of cerebellum-cerebellar and cortical-cerebellar connectivity, similar to that observed for cerebral cortex ^34^.

Last but not least, FNC between time series components can be successfully employed to predict behavior. This finding further confirms that the patterns of co-fluctuation between white matter components may be functionally relevant, by encoding inter-individual differences in behavioral traits. We choose to limit our analysis to higher-level cognition measures, which are frequently used in connectome-based prediction tasks ^51^ and, specifically, have been shown to be predictable by dynamic connectivity measures ^7, 10^. Although we did not directly test the accuracy of predictions derived from tw-dFC data versus other static or dynamic connectivity methods, we suggest that tw-dFC-based behavioral prediction may show its usefulness by allowing a stronger link between functional measures and the underlying white matter anatomy, even considering that correlation between behavior and structural connectivity measures have been shown to be generally weaker than those obtained by functional connectivity ^17, 52, 53^.

Taken together, these results confirm ICA-analysis of tw-dFC data as a powerful and versatile tool to investigate the relations between structural connectivity, functional activity, and behavior. Mapping behaviorally and clinically relevant functional information to the underlying white matter structures is of major relevance for a better understanding of brain functional anatomy in health and disease (Fox, 2018). Indeed, ICA analysis of tw-dFC data has potential translational application in diseases that involve direct damage to white matter bundles, such as stroke ^54^, traumatic brain injury ^55^or multiple sclerosis ^56^. In addition, such a method may provide new insights into the pathophysiology of some neuropsychiatric conditions in which there is evidence of subtle white matter connectivity alterations such as epilepsy, schizophrenia, bipolar disorder, major depressive disorder or autism spectrum disorders ^57^.

The present work does not come without limitations. The first concern regards the choice of dimensionality for the ICA decomposition of tw-dFC signal. In order to provide a compact representation of white matter independent components, we maintained our ICA dimensionality generally low, i.e., not more than 100 components, if compared to similar applications ^17^, and component stability was evaluated using ICASSO to avoid overfitting ^27^. Another possible limitation is related to which dimension to impose maximized independence during ICA processing. Our choice towards spatial ICA (i.e., maximizing the independence of components in space instead of time) was substantially aimed at keeping our work consistent with most of the existing literature on ICA analysis of static and dynamic fMRI data ^2, 7, 58^. In addition, the choice of minimizing between-component correlation in space has permitted the conservation of higher degrees of correlation between component time series, which, as suggested by our findings, may encode biologically relevant information. However, imposing the constraint of spatial independence to white matter components could lead to oversimplification as multiple fiber populations, each with potentially different temporal profiles, may share the same spatial localization. Additional investigation using temporal ICA of tw-dFC data could add further insights on the temporal organization of white matter activity.

In addition, it is important to remark the role of non-neuronal sources (such as CSF pulsations, motion-related noise or cerebrovascular reactivity) as a potential confound when investigating dynamic fluctuations in functional connectivity ^6^. Although our results were obtained from nuisance-corrected fMRI data, preprocessed independently using two state-of-art denoising algorithms ^59^, disentangling neuronal from non-neuronal signal in dynamic connectivity may be challenging even with the existing tools. On the other hand, constraining the dynamic functional connectivity analysis to voxels which are structurally connected by streamlines, as in tw-dFC, could help mitigate such drawback ^23^. Nevertheless, it is generally well-known that tractography-derived priors may not entirely reflect the ground-truth white matter anatomy, being affected by relatively high false positive rates ^47^. In the present work we employed a probabilistic tractography algorithm paired with track-filtering in the main dataset (multi-shell data), and the same algorithm without track-filtering in the validation dataset (single shell data). Albeit finding generally good external reproducibility, it remains difficult to rule out how much of the differences observed between the two datasets were due to tractography. It is very likely that optimizing the tw-dFC map generation by integrating a tractography pipeline with high robustness to noise could help raising the external reproducibility of the resulting tw-dFC components, and the impact of different signal modeling and tracking algorithms (such as deterministic or global tractography) is yet to be evaluated. Finally, since the quality of tw-dFC maps depends on the correct alignment of tractography and BOLD-fMRI data, non-linear registration of tractograms, as well as proper distortion correction of both DWI and BOLD data, are critical steps that need accurate control to ensure the good quality of the results.

## Conclusions

In the present work, we demonstrated that ICA decomposition of tw-dFC data may reliably identify spatial patterns of white matter connectivity, each showing distinct temporal profiles of activity. These spatial patterns, at the coarse scale, show similarity to well-known functional networks, and at increasing dimensionality are able to capture subtle anatomical details of white matter organization. Temporal patterns of activity in the white matter showed evidence of hierarchical organization at different levels, and their pattern of correlation may encode differences in behavioral traits.

In summary, we showed that tw-dFC is a powerful and versatile tool to investigate the relationships between brain structure, function, and behavior and it shows promise as a tool to deepen our knowledge about brain connectivity in health and disease.

## Materials and methods

### 1. Subjects and data acquisition

#### 1.1. Primary dataset (HCP)

Structural, diffusion and resting-state functional MRI data of 210 healthy subjects (males=92, females=118, age range 22-36 years) were retrieved from the HCP repository (https://humanconnectome.org). Data have been acquired by the Washington University, University of Minnesota and Oxford University (WU-Minn) HCP consortium. The Washington University in St. Louis Institutional Review Board (IRB) approved subject recruitment procedures, informed consent and sharing of de-identified data.

MRI data were acquired on a custom-made Siemens 3T “Connectome Skyra” (Siemens, Erlangen, Germany), provided with a Siemens SC72 gradient coil and maximum gradient amplitude (Gmax) of 100 mT/m (initially 70 mT/m and 84 mT/m in the pilot phase), to improve acquisitions of diffusion-weighted imaging (DWI).

High resolution T1-weighted MPRAGE images were collected using the following parameters: voxel size = 0.7 mm, TR = 2400 ms, TE = 2.14 ms .

DWI data were acquired using a single-shot 2D spin-echo multiband Echo Planar Imaging (EPI) sequence and equally distributed over 3 shells (b-values 1000, 2000, 3000 mm/s^2^), 90 directions per shell, spatial isotropic resolution of 1.25 mm.

For rs-fMRI, a gradient-echo EPI resolution was acquired with the following parameters: voxel size = 2mm isotropic, TR = 720 ms, TE = 33.1 ms, 1200 frames, ∼15 min/run. Scans were acquired along two different sessions on different days, with each session consisting of a left-to-right (LR) and a right-to-left (RL) phase encoding acquisition; in the present work, we employ left-to-right and right-to-left acquisitions from a single session only (first session) ^24, 60–62^.

#### 1.2. Validation dataset (LEMON)

We obtained structural, diffusion and rs-fMRI data of 213 healthy subjects (males=138, females=75, age range 20-70 years) from the Leipzig Study for Mind-Body-Emotion Interactions (LEMON) dataset (http://fcon_1000.projects.nitrc.org/indi/retro/MPI_LEMON.html). The study was carried out in accordance with the Declaration of Helsinki and the study protocol was approved by the ethics committee at the medical faculty of the University of Leipzig.

MRI was performed on a 3T scanner (MAGNETOM Verio, Siemens Healthcare GmbH, Erlangen, Germany) equipped with a 32-channel head coil.

High resolution structural MRI scans were acquired using a MP2RAGE sequence and with the following parameters: voxel size =1 m, TR = 5000 ms, TE = 2.92 ms.

Single-shell DWI data were acquired using a multi-band accelerated sequence with spatial isotropic resolution= 1.7 mm, b-value=1000, 60 diffusion-encoding directions.

For rs-fMRI data, a gradient-echo EPI was performed with the following parameters: phase encoding = AP, voxel size=2.3 mm isotropic, TR = 1400 ms, TE = 30 ms, 15.30 min/run ^25^.

### 2. Data preprocessing

#### 2.1. Structural preprocessing

All T1-weighted images were obtained in skull-stripped version ^25, 60^ and were subsequently segmented into cortical and subcortical gray matter (GM), white matter (WM) and cerebrospinal fluid (CSF) using FAST and FIRST FSL’s tools (https://fsl.fmrib.ox.ac.uk/fsl/fslwiki/). The segmentation outputs were collapsed into a 5-tissue-type (5TT) image that was required later in the tractography pipeline. T1-weighted volumes were also non-linearly registered to the 1-mm resolution MNI 152 asymmetric template using FLIRT and FNIRT from the FSL toolbox. The quality check of the registered T1 images was performed by visual inspection in specific axial, sagittal and coronal sections.

#### 2.2. DWI preprocessing

For the HCP dataset, DWI scans were retrieved in a minimally preprocessed form which includes eddy currents, EPI susceptibility-induced distortion and motion correction, as well as linear registration of structural and DWI images ^60^.

In contrast, the LEMON DWI scans were obtained in raw format and underwent preprocessing through the dedicated pipeline included in the MRtrix3 software (https://www.mrtrix.org/). It features denoising using Marchenko-Pastur principal component analysis (MP-PCA), removal of Gibbs ringing artifacts, eddy currents, distortion (by exploiting the available reverse-phase encoding scans) and motion correction using EDDY and TOPUP FSL’s tools, as well as bias field correction using the N4 algorithm ^63, 64^.

#### 2.3. Resting state fMRI preprocessing

Both HCP and LEMON rs-fMRI data were obtained in preprocessed and denoised form.

HCP data minimal preprocessing included the following steps: field inhomogeneity-related artifact correction, motion correction, registration to standard space (MNI152, 2mm resolution), high pass temporal filtering (> 2000 s full width at half maximum) for removal of slow drifts ^60^, artifact components identification using ICA-FIX ^65^ and regression of artifacts and motion-related parameters ^62^. Minimally preprocessed data were additionally band-pass filtered (0.01-0.09 Hz) and the global WM and CSF signal was regressed out to further improve ICA-based denoising ^66^.

The LEMON dataset processing pipeline included removal of the first 5 volumes to allow for signal equilibration, motion and distortion correction, artifact detection (rapidart) and denoising using component-based noise correction (aCompCor), mean-centering and variance normalization of the time series as well as spatial normalization to MNI 152, 2mm resolution template ^25^.

Finally, all rs-fMRI volumes were smoothed through convolution with a Gaussian kernel of 6mm full width at half maximum. All the additional preprocessing described above was carried out in the CONN toolbox ^67^.

### 3. Tractography and track-weighted dynamic functional connectivity (tw-dFC)

Whole-brain tractograms were obtained for each subject using the following pipeline: first, diffusion signal modeling was performed on the preprocessed DWI data within the constrained-spherical deconvolution (CSD) framework, which estimates white matter Fiber Orientation Distribution (FOD) function from the diffusion-weighted deconvolution signal using a single fiber response function (RF) as reference ^68^. Specifically, multi-shell HCP DWI data underwent multi-shell multi-tissue (MSMT) CSD signal modeling, an optimized version of the CSD approach which allows for separate response function calculation in WM, GM and CSF, reducing the presence of spurious FOD in voxels containing GM and/or CSF ^69^. To achieve a similar result on the single-shell LEMON data, the diffusion signal was modeled by using single-shell 3-tissue CSD, a variant of the MSMT model optimized for RF estimation in single-shell datasets. SS3T-CSD signal modeling was performed using MRtrix3Tissue ^70^, a fork of MRtrix3 software. After signal modeling, whole-brain tractography was performed using the IFOD2 algorithm with default parameters, and by applying the anatomically constrained tractography (ACT) frameworks, which makes use of the 5TT map previously obtained from segmentation to improve the biological plausibility of the resulting streamlines ^71, 72^.

For the high b-value, multi-shell HCP data, the spherical-deconvolution informed filtering of tractograms (SIFT) ^73^ algorithm was applied to further improve the fit between the reconstructed streamlines and the underlying DWI data, starting from a generated tractogram of 10 million streamlines and filtering it to a final whole-brain tractogram of 1 million streamlines. Since the SIFT algorithm is best suited for high b-value datasets, it was not applied on the single-shell, low b-value LEMON dataset, where instead a whole brain tractogram of 5 million streamlines was generated (note that the number of streamlines is uninfluential to the tw-dFC contrast generation). Whole-brain tractograms for each subject of both datasets were transformed to the MNI 152 standard space by applying the non-linear transformations obtained from structural images. Since each subject’s rs-fMRI volumes were already registered to the MNI 152 template, all the following analyses took place in standard space.

For each subject, whole-brain tractograms derived from tractography and preprocessed rs-fMRI time series were combined to generate a 4-dimensional tw-dFC dataset with the same spatial and temporal resolution of the original fMRI time series. In this framework, each white matter voxel’s time series reflects the dynamic changes in functional connectivity occurring at the endpoints of the white matter pathways traversing that voxel ^23^. For sliding-window functional connectivity analysis of rs-fMRI time series, a rectangular sliding window with ∼40 s length (55 time points for the HCP data, TR = 0.72 s; 29 time points for the LEMON data, TR = 1.4 s) was employed, as suggested by previous works ^6, 74^. For the HCP data, tw-dFC derived from LR and RL phase encoding volumes were temporally concatenated for each subject.

### 4. Group independent component analysis (ICA)

The obtained tw-dFC volumes were analyzed using a spatial group ICA framework as implemented in the Group ICA of FMRI Toolbox (GIFT) ^75, 76^. Group analysis was performed separately for the primary dataset (HCP) and the validation dataset (LEMON). Briefly, the pipeline for group ICA analysis involves a first step, in which a subject-level principal component analysis (PCA) is performed for dimensionality reduction purposes, and a second step in which dimensionality-reduced data are temporally concatenated and undergo a secondary PCA dimensionality reduction along directions of maximal group variability. Finally, the group PCA-reduced matrix is decomposed into a given number of independent components (ICs) using the Infomax algorithm ^77^.

For the first data reduction step, 120 subject-specific PCA components were chosen, as in previous works ^76^. The second data reduction step and subsequent ICA decomposition were performed at three different dimensionality levels (ICA_10_, ICA_20_ and ICA_100_). For each run, the ICA algorithm was repeated 20 times in ICASSO and the n most reliable components were identified as the final group-level components, to ensure stability of estimation. ICASSO returns a stability (quality) index (Iq) for each estimate-cluster. This provides a rank for the corresponding ICA estimate. In the ideal case of *m* one-dimensional independent components, the estimates are concentrated in *m* compact and close-to-orthogonal clusters. In this case, the index to all estimate-clusters is very close to one, while it drops when the clusters grow wider and less homogeneous ^27^.

Finally, the resulting components for each run were visually inspected to ensure that: 1) the peak activation of each network was localized in the white matter; 2) there was only minimal overlap to vascular, meningeal, ventricular sources of artifacts; 3) the mean power spectra of each network showed prominence of low frequency spectral power.

For each component, subject-specific spatial maps and time courses were obtained using group-information guided ICA (GIG-ICA) back-reconstruction ^78^. A one sample t-test was run to generate group statistical maps for each component’s SM, and a hard parcellation of the white matter was obtained by thresholding the obtained t-maps at z=1. To facilitate the interpretation of the results, spatial maps were annotated by calculating percentage overlap with regions of interest from known GM ^79^and WM atlases ^80, 81^.

### 5. Reproducibility analysis

The reproducibility of results was evaluated at three different levels on the primary dataset: intra-subject reproducibility (test-retest), inter-subject reproducibility (split-half) and inter-cohort reproducibility (comparison with the validation dataset).

For test-retest reproducibility analysis, the LR- and RL-phase encoding acquisitions for each subject were employed as the test and retest data respectively; in the split-half reproducibility analysis, the HCP sample was split in random halves (105 subject each) and the ICA was run separately on each of them; finally, in the external reproducibility analysis the results from ICA on the primary dataset were directly compared to the results obtained on the validation dataset. In all cases, the comparison metrics were: i) the pairwise Pearson’s correlation coefficient between each spatial component and its corresponding component (i.e., the component which scored the highest correlation coefficient); ii) the Dice similarity coefficient (DSC) ^82^ between the obtained white matter parcellations, i.e. between each component’s binarized, z-thresholded group statistical map and the corresponding component map.

reproducibility analysis was run separately for each ICA dimensionality level (n=10, n=20, n=100).

### 6. Task-based functional network annotation

To provide insights on the functional relevance of the identified white matter components, i.e. to give a measure of how large-scale white matter networks may be involved in the execution of complex tasks, we compared the resting-state spatial maps resulting from the main dataset to the task-based white matter activation maps derived with a recently developed method, namely “Functionnectome”^28^. To compare these results to the ICs derived from distinct ICA runs, pairwise Pearson’s correlation between the z-weighted functionnectome maps and each component’s z-map were computed after a thresholding of z > 0. .We considered correlations significant if the proportion of shared variance between tw-dFC and functionnectome maps was above 5% (e.g., a spatial correlation of r > 0.22).

### 7. Connectivity-based component classification

Functional network connectivity (FNC), defined as the pairwise correlation between each pair of IC time courses, was measured on the primary dataset for each ICA dimensionality level. For each run of ICA, we sought to classify components in an unsupervised way based on the similarity of their activity profiles, by performing k-means clustering on the group-average FNC matrix. The optimal number of clusters (k) was determined by plotting the ratio of between-group variance to total variance for increasing values of k and identifying the elbow point (elbow method). To obtain robust cluster centroids, k-means clustering was performed on bootstrap resamples, by iterating clustering 100 times on randomly drawn samples of 168 subjects (80% of the total subjects); the resulting centroids were then employed to perform clustering on the whole dataset.

### 8. FNC-based cognitive performance prediction tasks

To evaluate the role of tw-dFC-derived FNC in predicting individual cognitive performance, we applied a linear regression model with leave-one-out cross validation (LOOCV) to behavioral measures of cognitive performance available in the HCP database. Behavioral measures of fluid intelligence (Penn Progressive Matrices, HCP_ID: PMAT24_A_CR), cognitive flexibility (Dimensional Change Card Sort, HCP_ID: CardSort_AgeAdj) and sustained attention (Short Penn Continuous Performance Test, HCP_ID: SPCPT_TP) were selected as independent variables. FNC measures from ICA_100_ were used as features to predict, independently and separately, each behavioral variable. More specifically, FNC upper triangular matrices were vectorized to obtain a single feature vector per subject, and underwent a first feature selection step, in which only features with the highest correlation coefficient (p < 0.01) to behavioral scores were retained. The retained features were then fed into a predictive linear model and predicted scores were generated for each subject using a LOOCV approach, i.e., for each iteration, data from one subject were set aside as test sample and the remaining subjects were used as training set; such step was iterated for all subjects. Finally, the difference between observed and predicted behavioral scores was computed and the residuals were used to obtain a R^2^ for each model, as a measure of goodness of fit. The statistical significance of this value was tested using a permutational approach (10,000 permutations), i.e. by iteratively calculating the R^2^ after random permutations of the behavioral scores; results were considered significant with p < 0.01.

### 9. “Brain state” vectors identification

The time series of components underwent further clustering analysis aimed at identifying stable or quasi-stable patterns of component activity weights which tend to reoccur over time and across subjects, an analogy to “brain states” described in the dynamic functional connectivity literature ^7, 8^. For tw-dFC components, we sought to replicate the procedure described in Fan et al., 2021^7^ for brain state vector identification from dynamic functional connectivity-derived independent components. For simplicity, and in analogy with this work, we employed the subject-specific time series obtained from ICA_20_. A pairwise distance matrix based on L1 distance function (Manhattan distance) was computed from timepoints*subject data matrices for each component time series. A distance matrix between component’s activity vectors was obtained by summing all 20 component distance matrices. Then, Ward’s method of agglomerative hierarchical clustering ^83^ was applied to the resulting distance matrix. To determine the optimal number of clusters (k), a cluster validity index (CVI) was computed as the ratio of within-cluster to between-clusters distance. To obtain robust cluster centroids, clustering was performed on bootstrap resamples, by iterating the algorithm 100 times on randomly drawn samples of 168 subjects (80% of the total subjects). Consequently, all the time points were classified into k clusters, each representing a vector of activity of the 20 tw-dFC components; brain states were obtained by averaging each component’s activity weights at the time points falling within the same cluster.

A complete representation of the whole processing pipeline is provided in Figure 1.

## Acknowledgments

The primary dataset (HCP) was provided by the Human Connectome Project, WU-Minn Consortium (Principal Investigators: David Van Essen and Kamil Ugurbil; 1U54MH091657), funded by the 16 NIH institutes and centers that support the NIH Blueprint for Neuroscience Research; and by the McDonnell Center for Systems Neuroscience at Washington University.

We also gratefully acknowledge the Mind-Body-Emotion group at the Max Planck Institute for Human Cognitive and Brain Sciences for the data of the “Leipzig Study for Mind-Body-Emotion Interactions” (LEMON) that have been used as validation dataset.

## Data and code availability

The code used for this article will be available after article publication at https://github.com/BrainMappingLab

