## Supplementary Information for "Unsupervised clustering of track-weighted dynamic functional connectivity reveals white matter substrates of functional connectivity dynamics"

### Supplementary figures

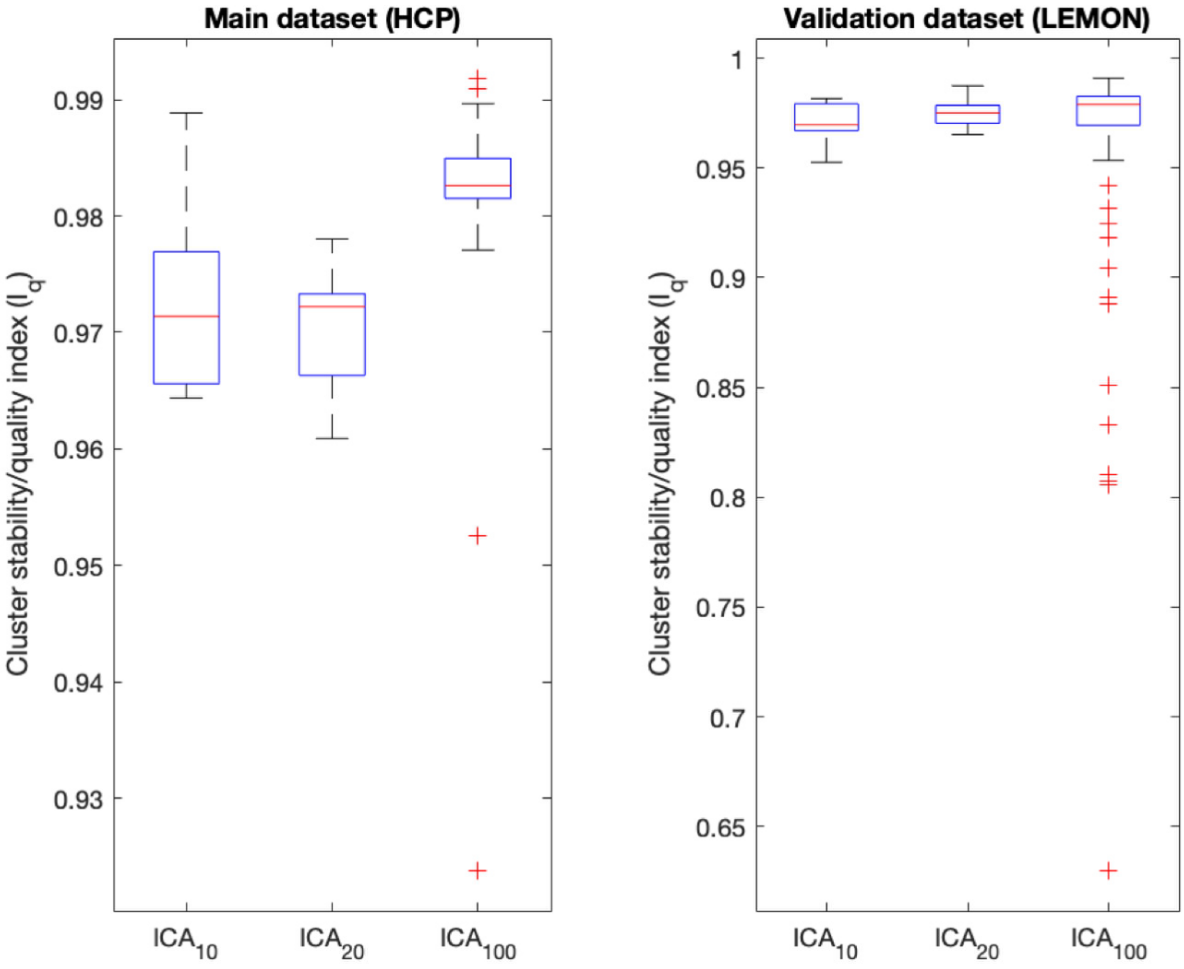


**Supplementary Figure 1. Cluster stability/quality index ($I_q$) of ICA runs.** For each ICA dimensionality, the aggregated cluster stability/quality indices of all components are displayed in box-plots. In both main and validation datasets, $I_q$ reached high values (close to 1) for most of the components. Median $I_q$ values show also a progressively increasing trend with higher ICA dimensionality.

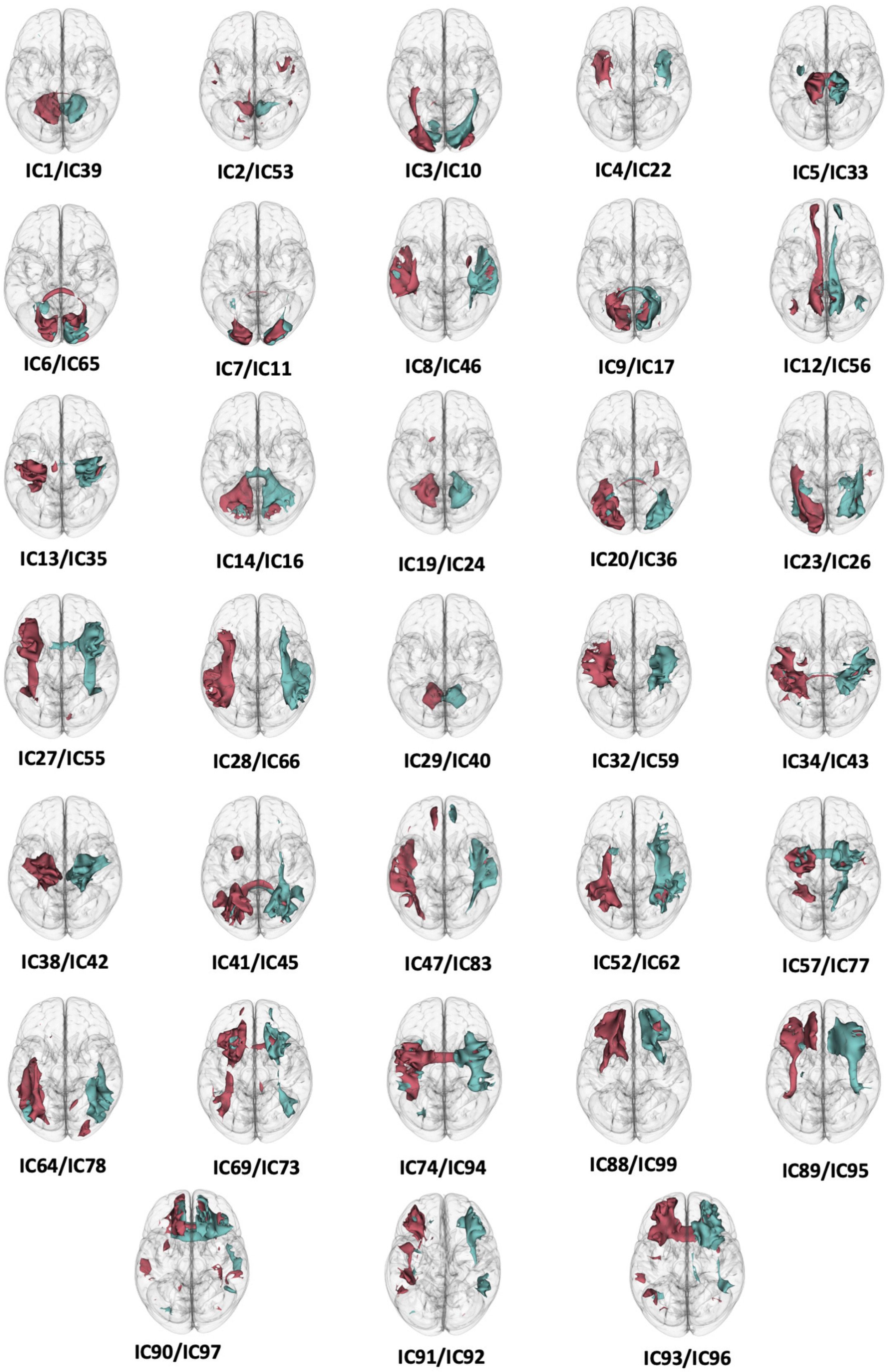


**Supplementary Figure 2. Pairs of symmetrical, lateralized components from $ICA_{100}$ run.** 66 components showed left or right lateralization and resulted to have a roughly symmetrical counterpart, which shares a similar spatial organization on the contralateral hemisphere.

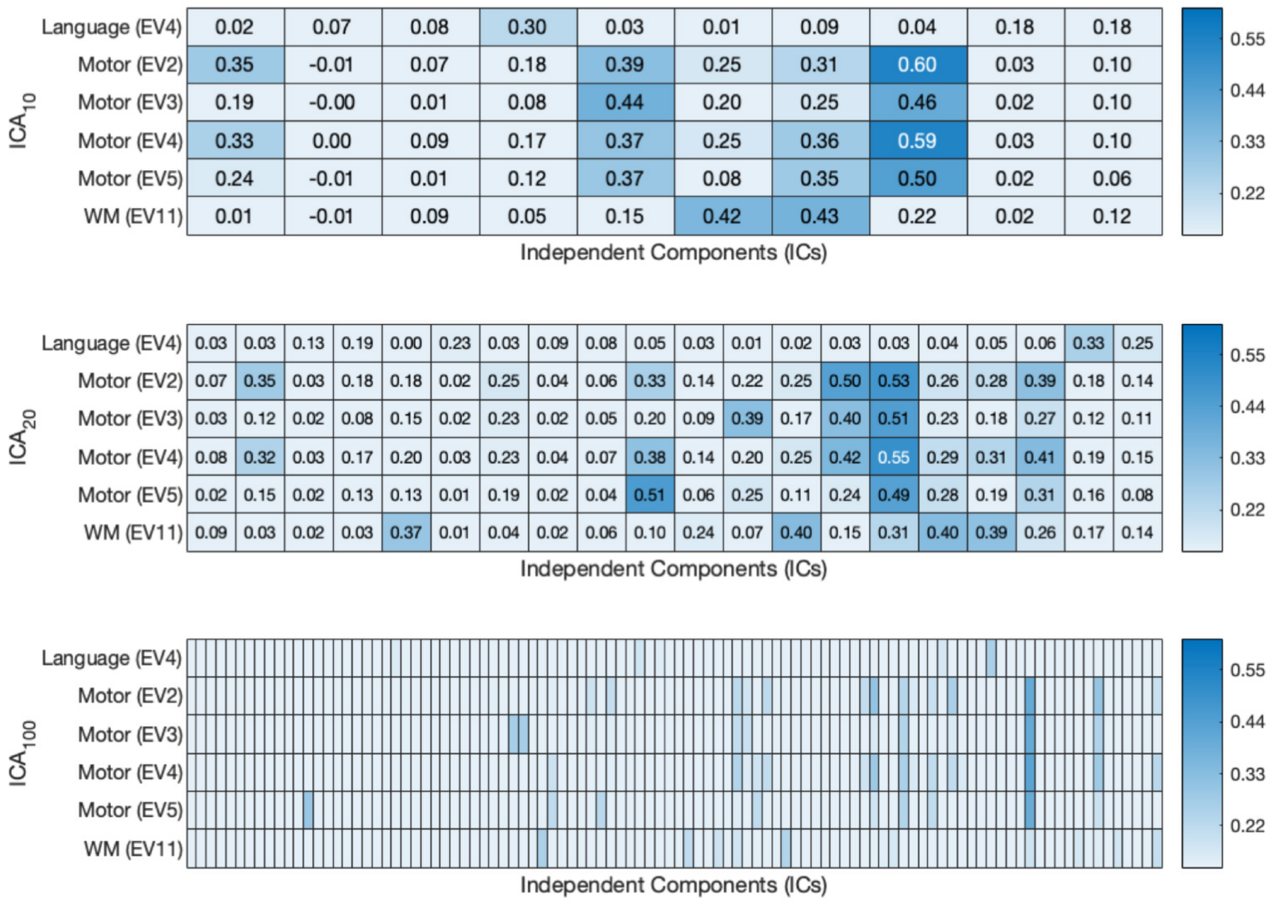


**Supplementary Figure 3. Correlation between resting-state track-weighted dynamic functional connectivity components (tw-dFC) and task-based white matter activation maps (Functionnectome).** The lower threshold was set to 0.22 (i.e. spatial maps which share 5% or above of their variance were deemed to be significantly correlated). Explanatory variables (EV) follow the same nomenclature of the HCP release.

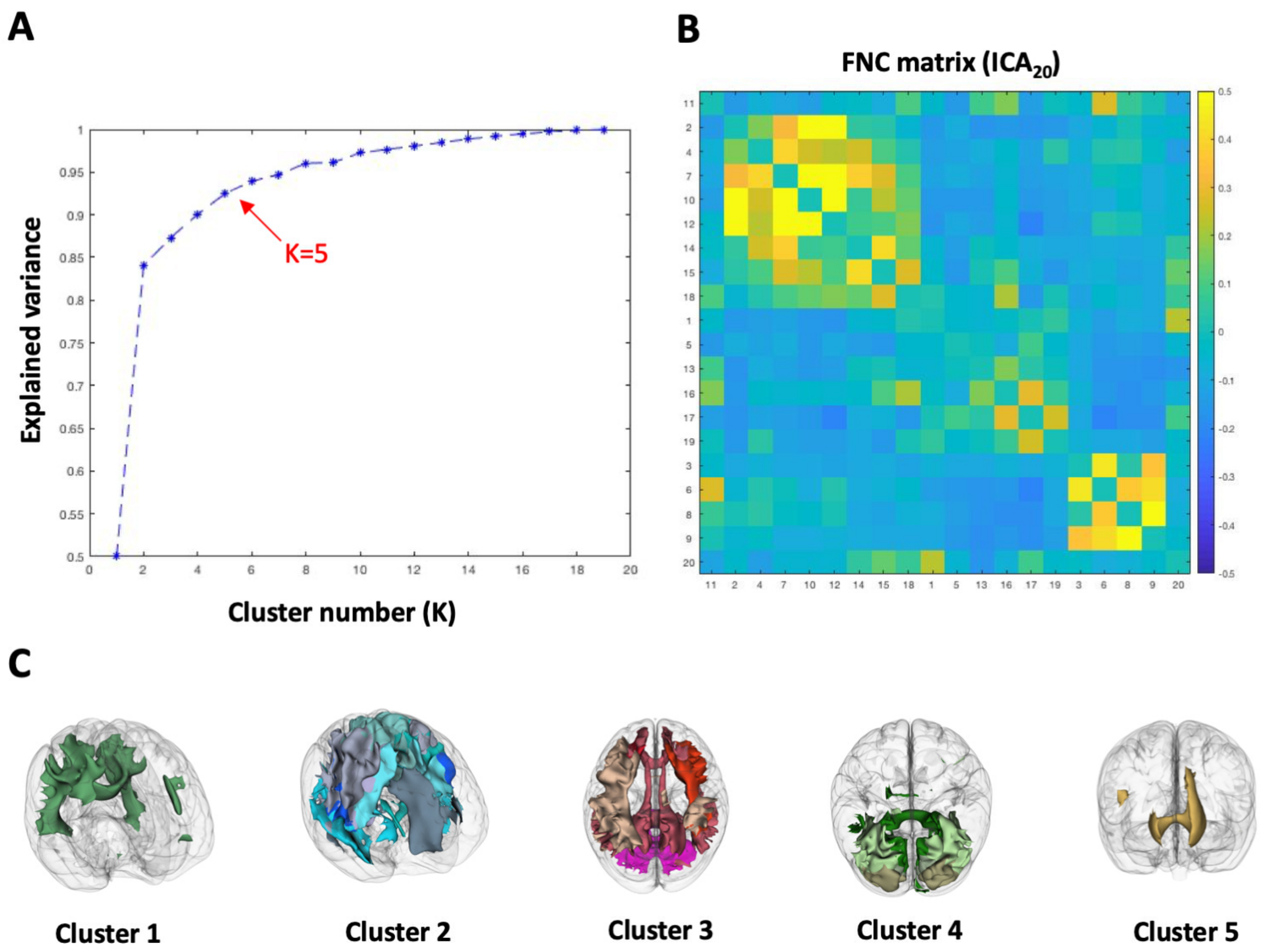


**Supplementary Figure 4. Functional network connectivity (FNC) clustering analysis for $ICA_{20}$. A)** Plot of the explained variance for different numbers of clusters. The elbow method (red arrow) suggests an optimal number of k=5. **B)** The FNC matrix, ordered according to the clustering results. **C)** Visualization of FNC-derived clusters. Group spatial maps for each component are thresholded at $z > 1$, binarized and volume-rendered on a glass-brain. For each component, red-orange shaded colors were assigned to associative/cerebellar clusters, blue-light blue colors to sensorimotor clusters, and green/brown colors to visual clusters.

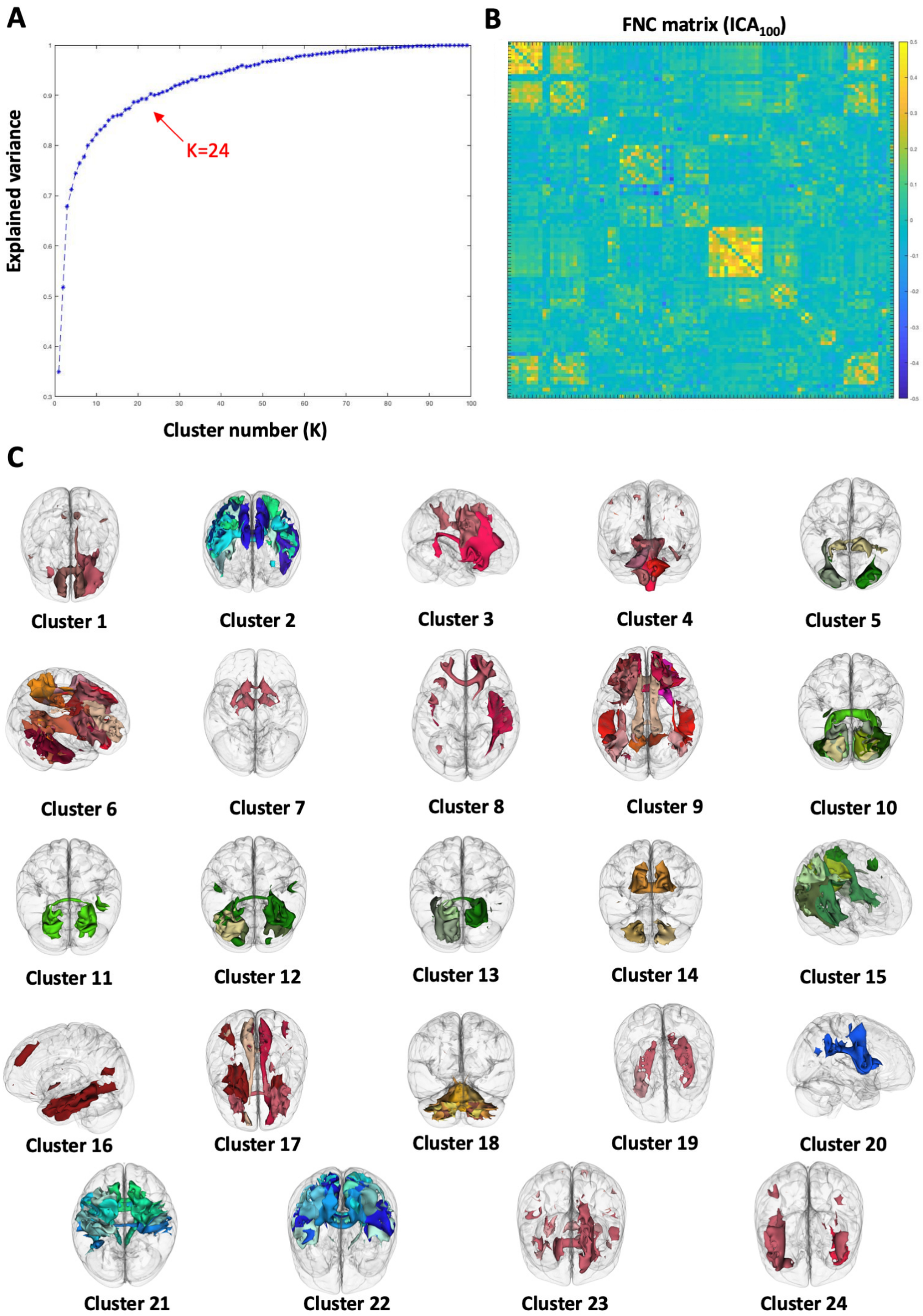


**Supplementary Figure 5. Functional network connectivity (FNC) clustering analysis for ICA$_{100}$.** **A)** Plot of the explained variance for different numbers of clusters. The elbow method (red arrow) suggests an optimal number of k=24. **B)** The FNC matrix, ordered according to the clustering results. **C)** Visualization of FNC-derived clusters. Group spatial maps for each component are thresholded at z > 1, binarized and volume-rendered on a glass-brain. For each component, red-orange shaded colors were assigned to associative/cerebellar clusters, blue-light blue colors to sensorimotor clusters, and green/brown colors to visual clusters.

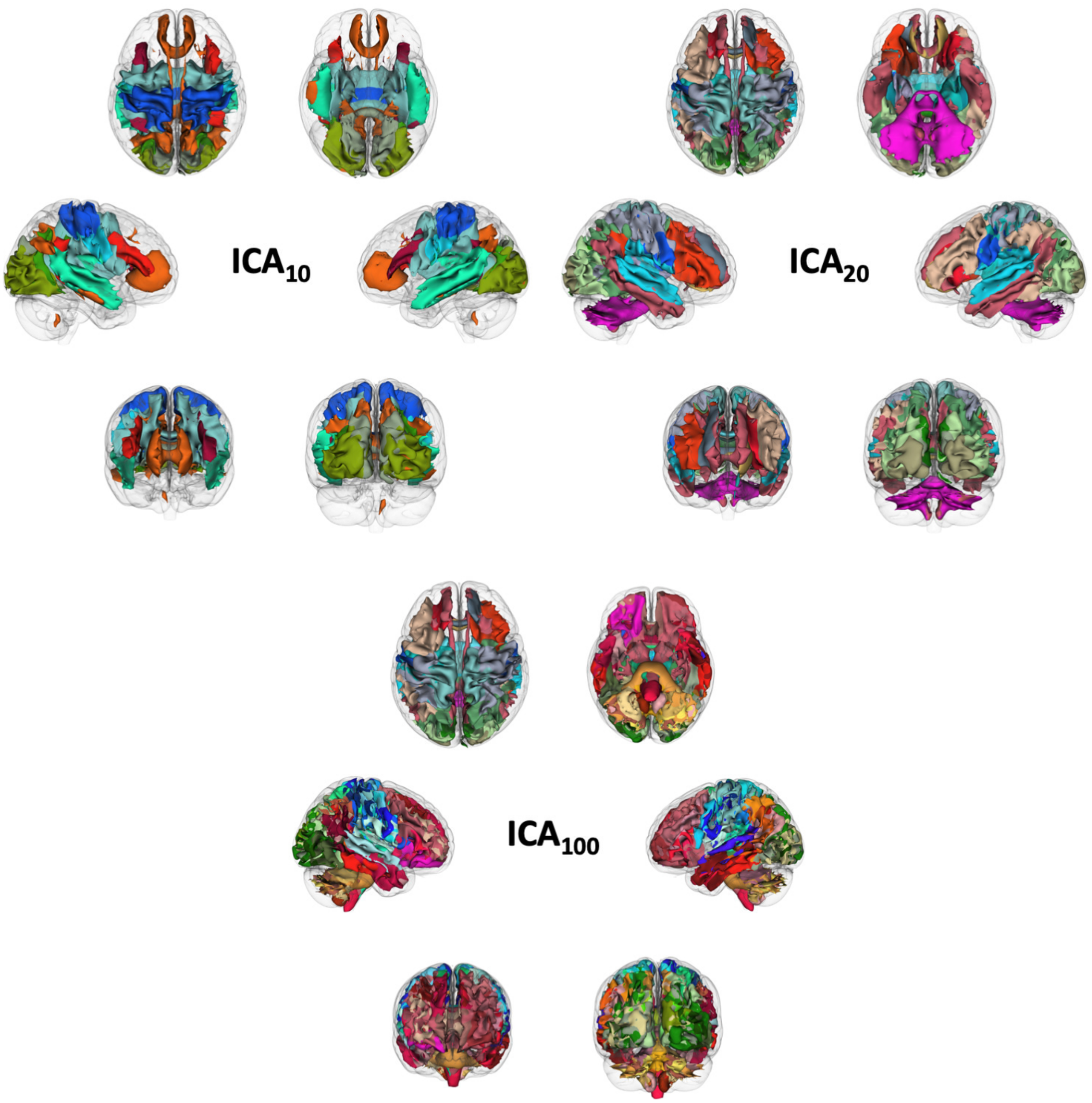


**Supplementary Figure 6. Overview of the topographical organization of white matter functional network connectivity (FNC) clusters.** Group spatial maps for each component are thresholded at $z > 1$, binarized and volume-rendered on a glass-brain. For each component, red-orange shaded colors were assigned to associative/cerebellar clusters, blue-light blue colors to sensorimotor clusters, and green/brown colors to visual clusters. The tripartite organization of white matter components into sensorimotor (unimodal), visual (unimodal) and associative (heteromodal) clusters, based on their functional correlation is maintained across all the ICA run, regardless of the number of clusters.
