## Supplementary File 1 for "Unsupervised clustering of track-weighted dynamic functional connectivity reveals white matter substrates of functional connectivity dynamics"

Summary of track-weighted dynamic functional connectivity components from each ICA run for the main and validation datasets. Component t-maps are overlaid on the MNI152 standard template (2mm resolution), thresholded at z =1 and showed in sagittal, coronal and axial slices centered on the peak t values. Mean power spectra of component time series are plotted below.

***INDEX:***


- **Pages 1-10:** *HCP dataset, n=10 run, components 1-10*
- **Pages 11-30:** *HCP dataset, n=20 run, components 1-20*
- **Pages 31-130:** *HCP dataset, n=100 run, components 1-100*
- **Pages 131-140:** *LEMON dataset, n=10 run, components 1-10*
- **Pages 141-160:** *LEMON dataset, n=20 run, components 1-20*
- **Pages 161-260:** *LEMON dataset, n=100 run, components 1-100*

### HCP dataset

*n=10 run*

3-1 10n_tmap_component_ica_s1 001

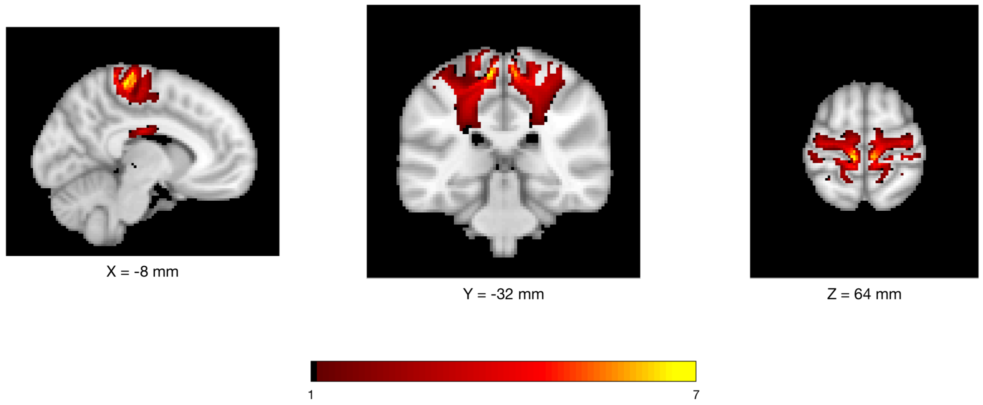


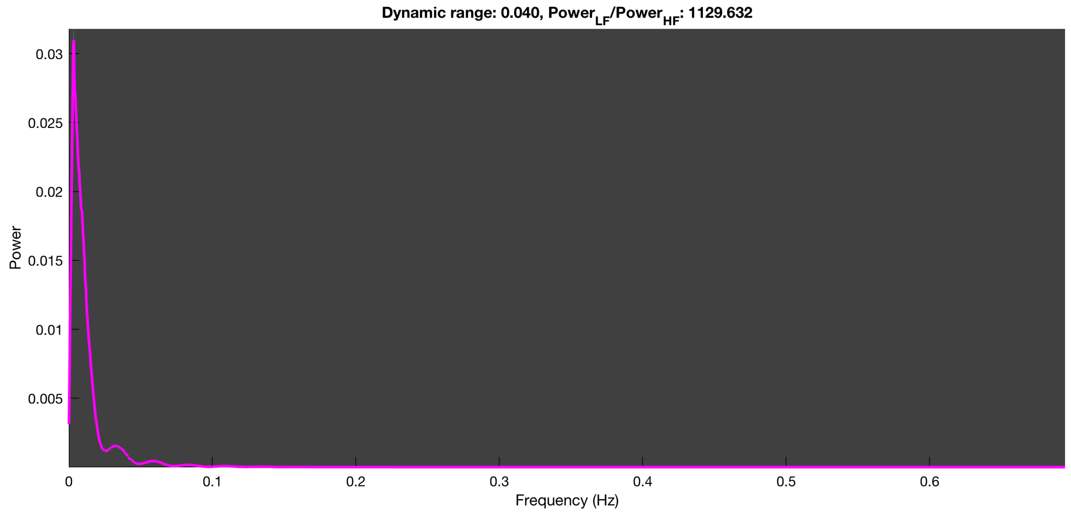

3-1 10n_tmap_component_ica_s1 002

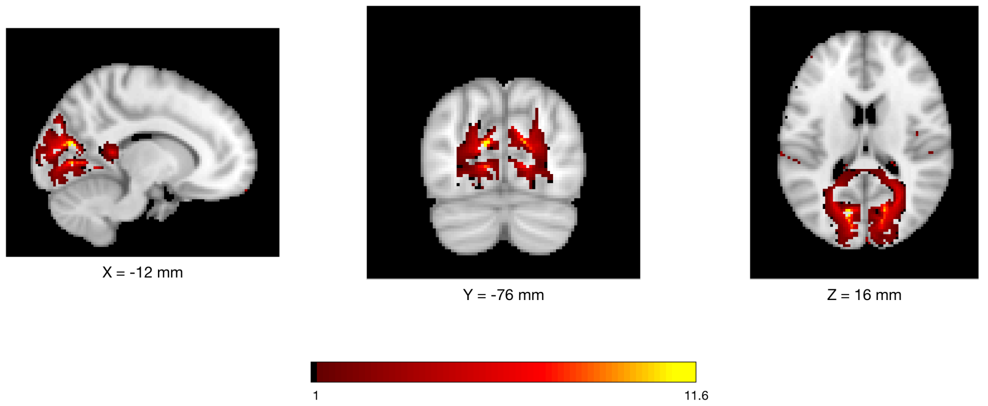


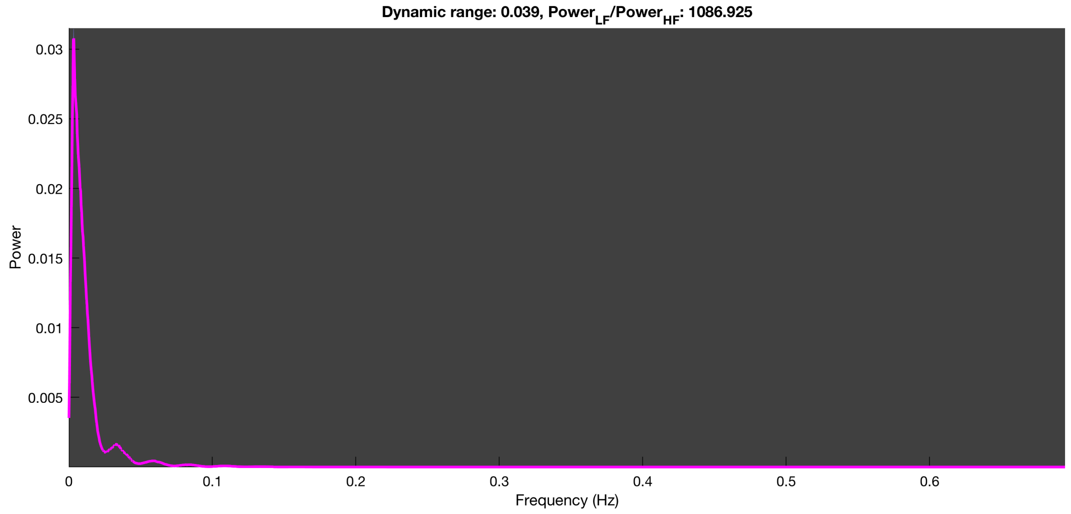

3-1 10n_tmap_component_ica_s1 003

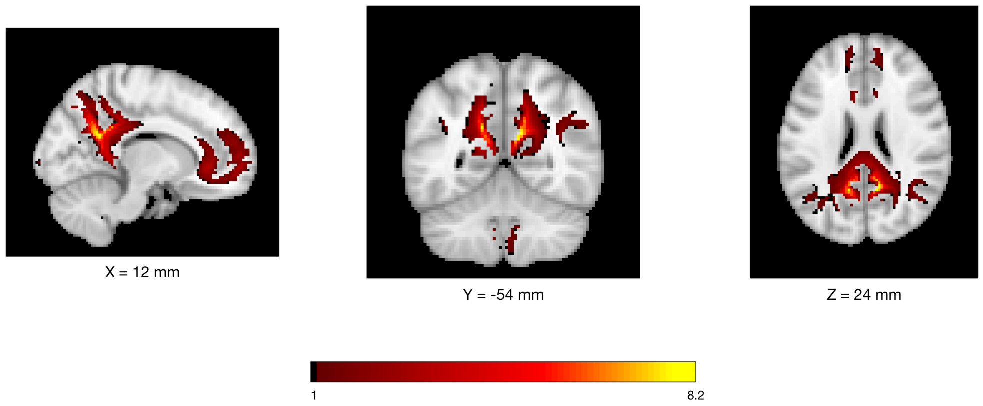


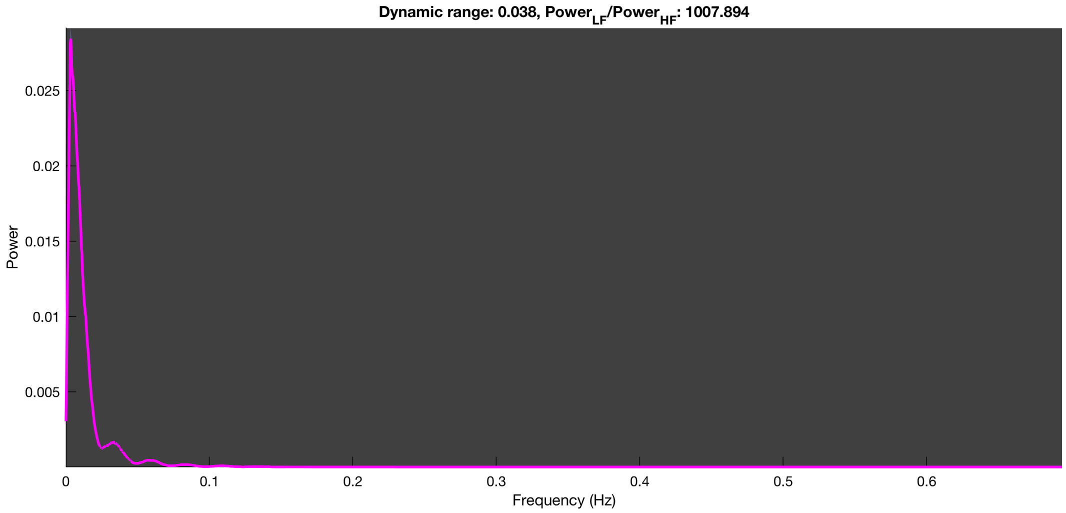

3-1 10n_tmap_component_ica_s1 004

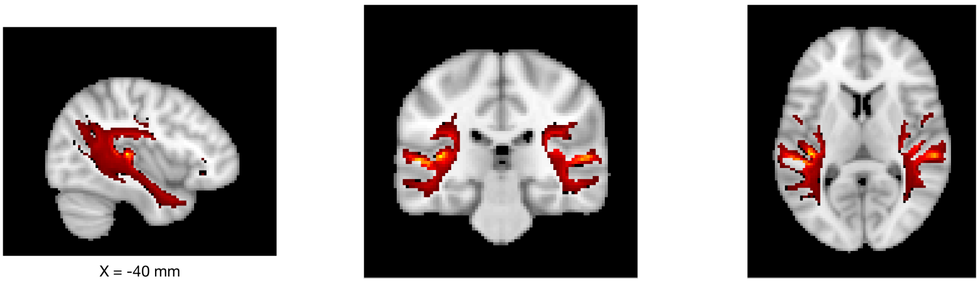

X = -40 mm Y = -28 mm Z = 8 mm

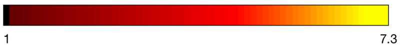


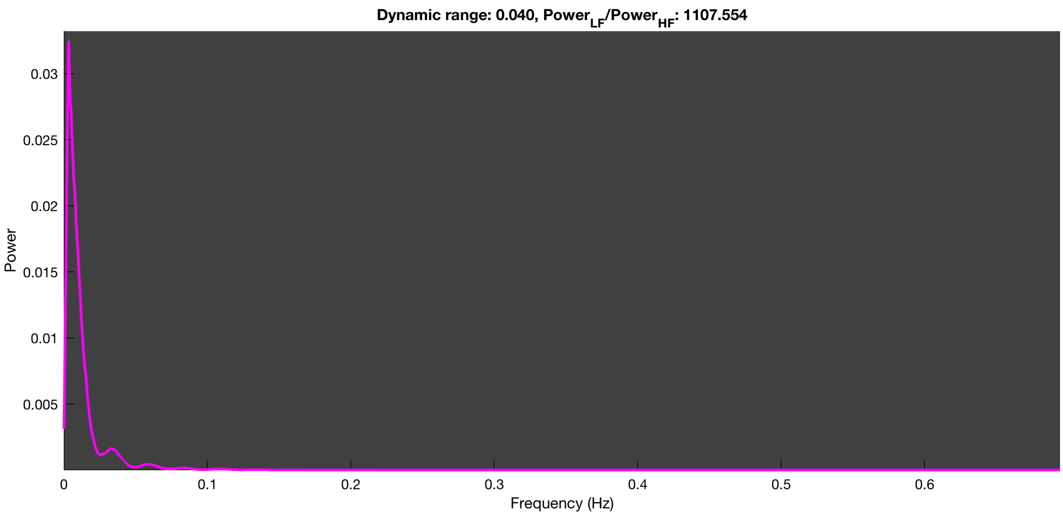

3-1 10n_tmap_component_ica_s1 005

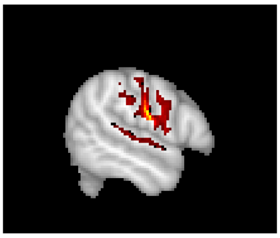

X = -54 mm

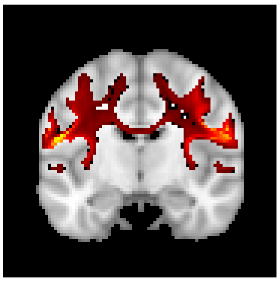

Y = -10 mm

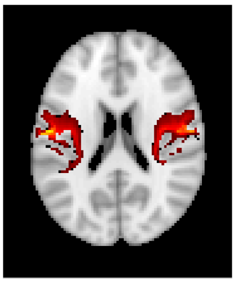

Z = 20 mm

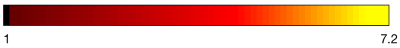


Dynamic range: 0.040, $Power_{LF}/Power_{HF}$: 1176.222

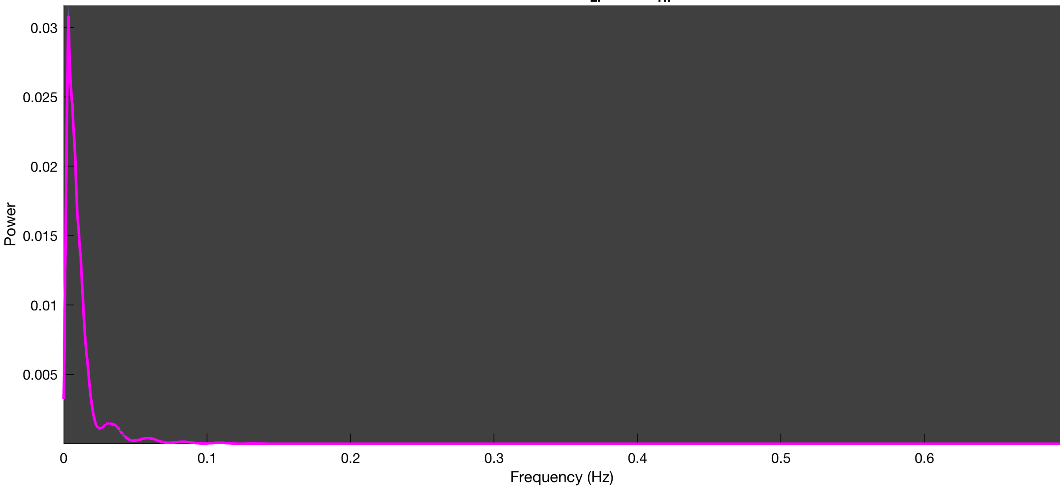

3-1 10n_tmap_component_ica_s1 006

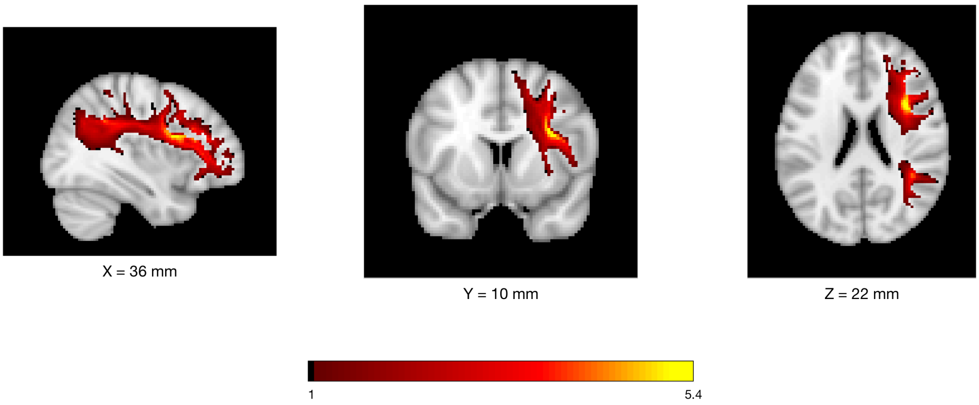


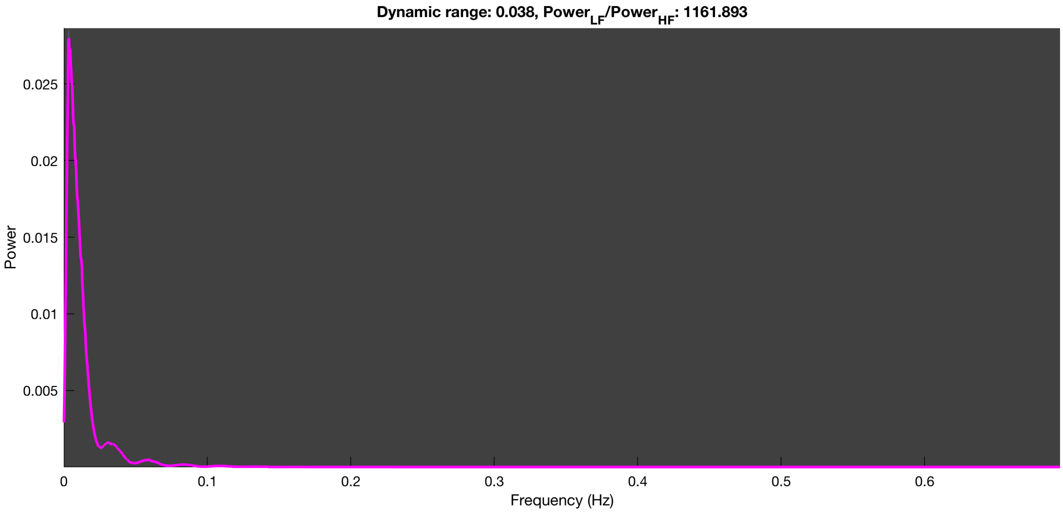

#### 3-1 10n_tmap_component_ica_s1 007

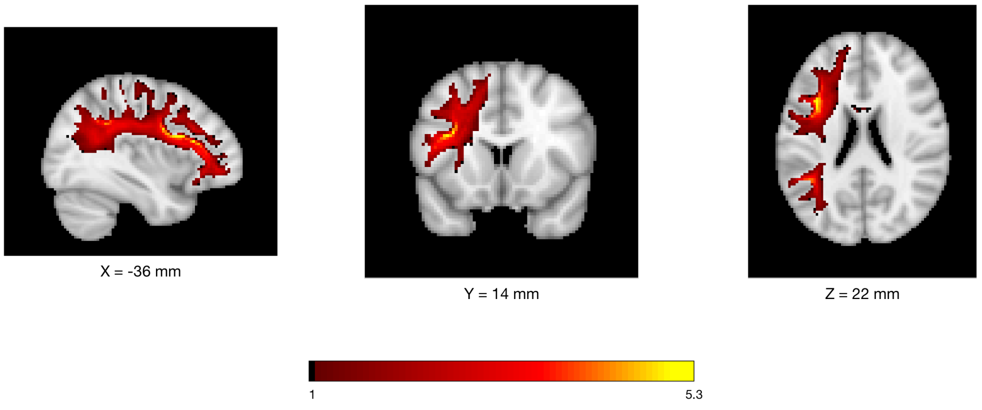


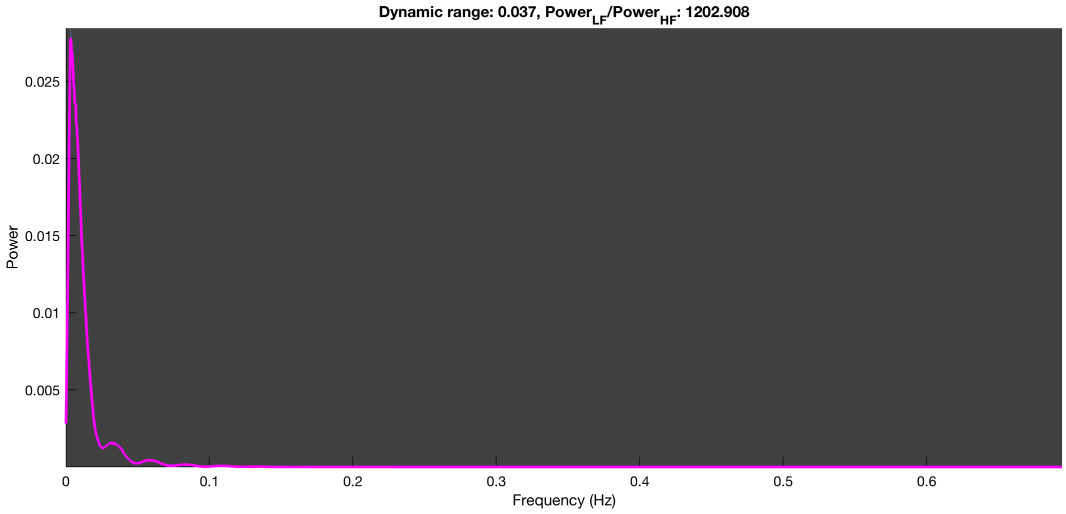

3-1 10n_tmap_component_ica_s1 008

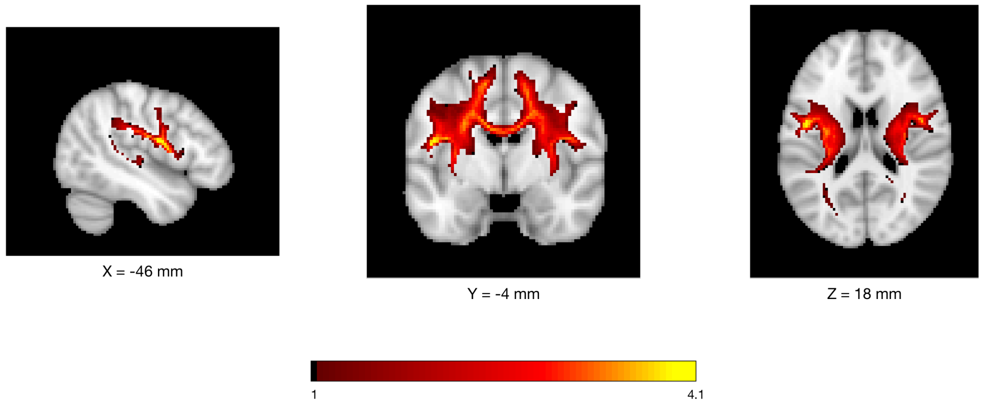


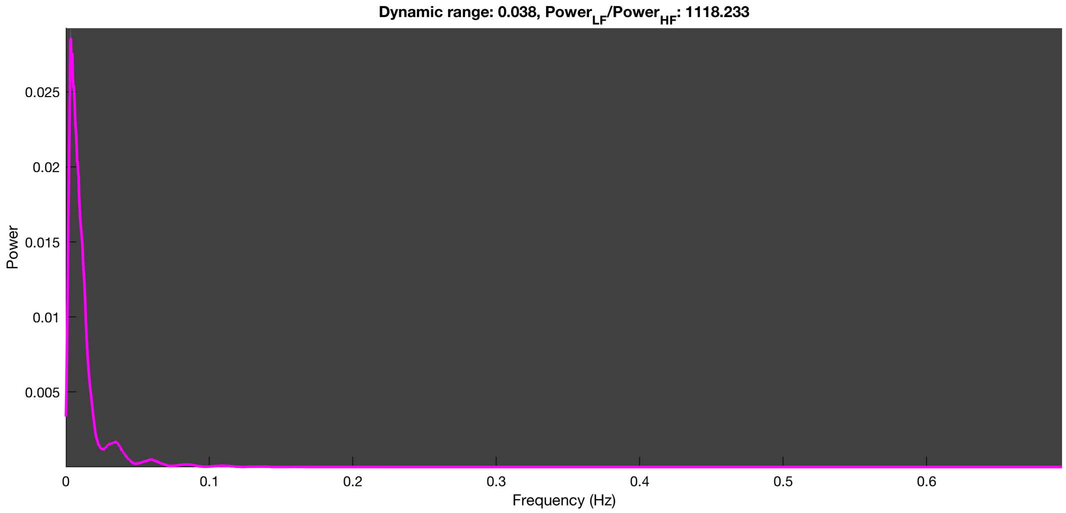

3-1 10n_tmap_component_ica_s1 009

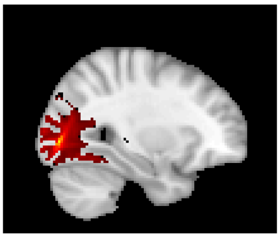

X = 30 mm

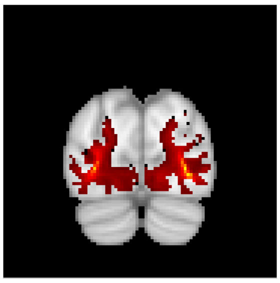

Y = -82 mm

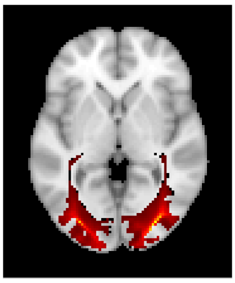

Z = 0 mm

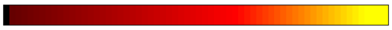


Dynamic range: 0.039, $Power_{LF}/Power_{HF}$: 1253.849

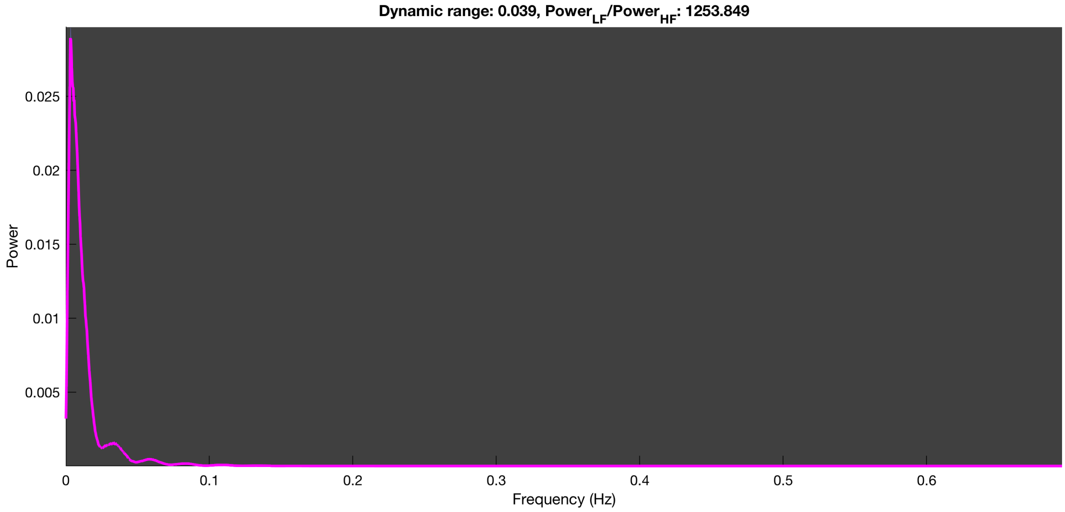

3-1 10n_tmap_component_ica_s1 010

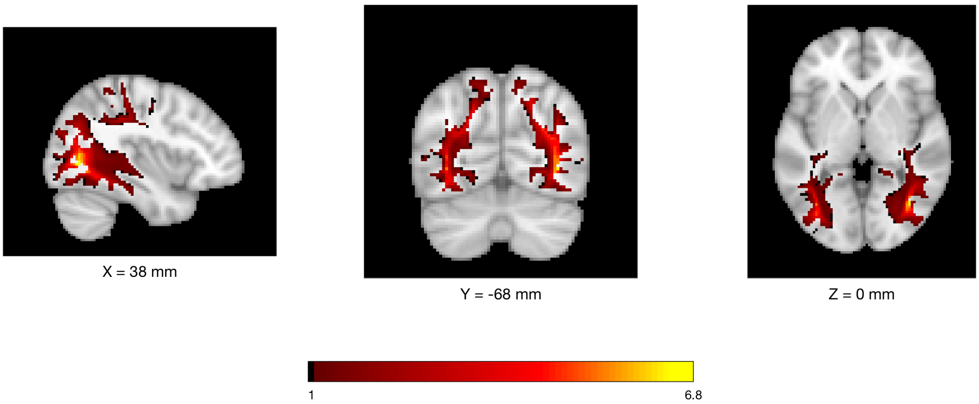


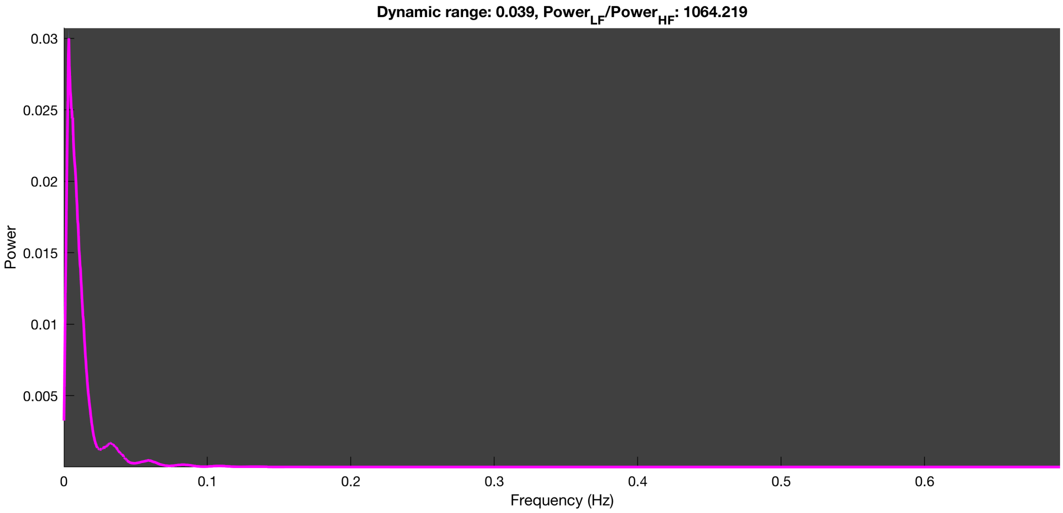

### HCP dataset

*n=20 run*

3-1 20n_tmap_component_ica_s1 001

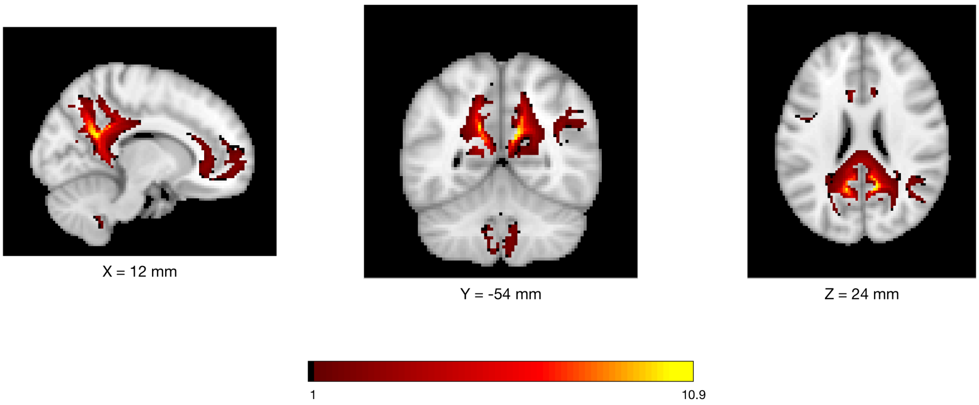


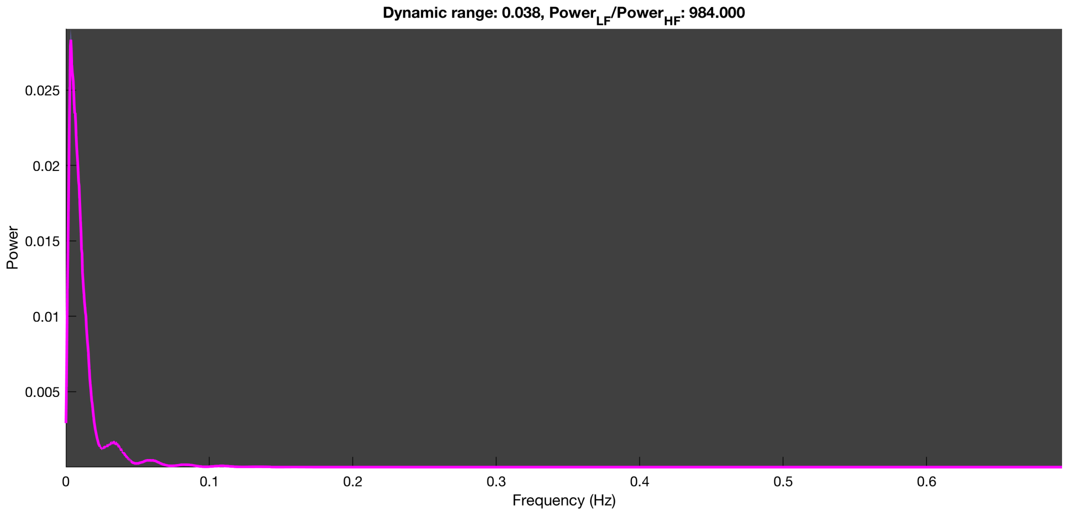

3-1 20n_tmap_component_ica_s1 002

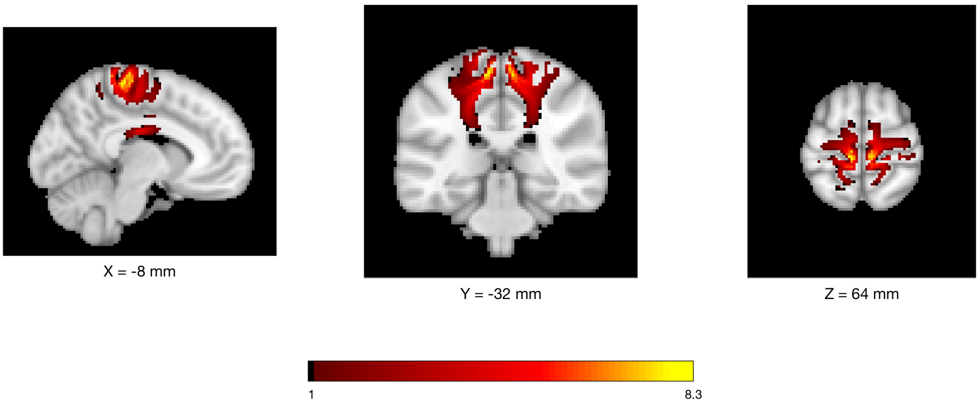


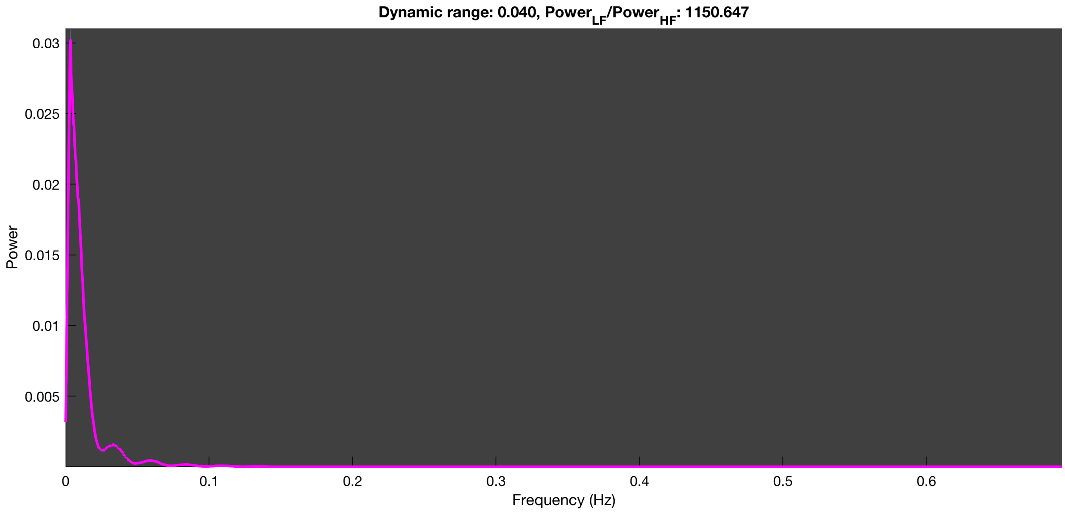

##### 3-1 20n_tmap_component_ica_s1 003

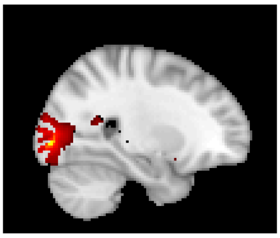

X = 26 mm

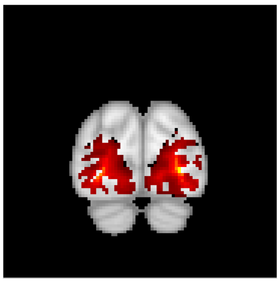

Y = -88 mm

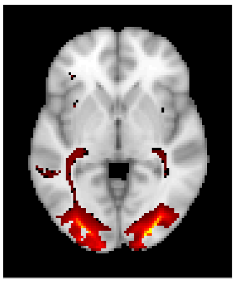

Z = -2 mm

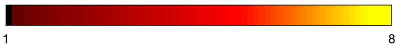


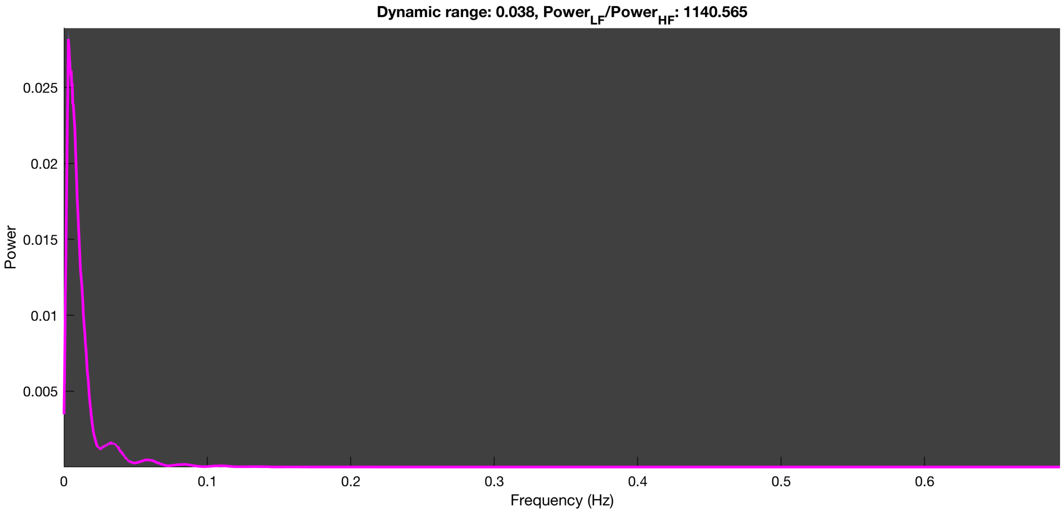

3-1 20n_tmap_component_ica_s1 004

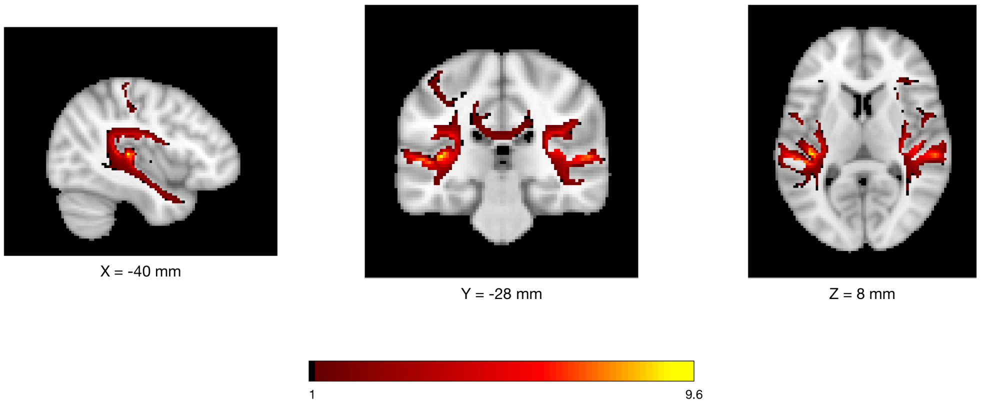


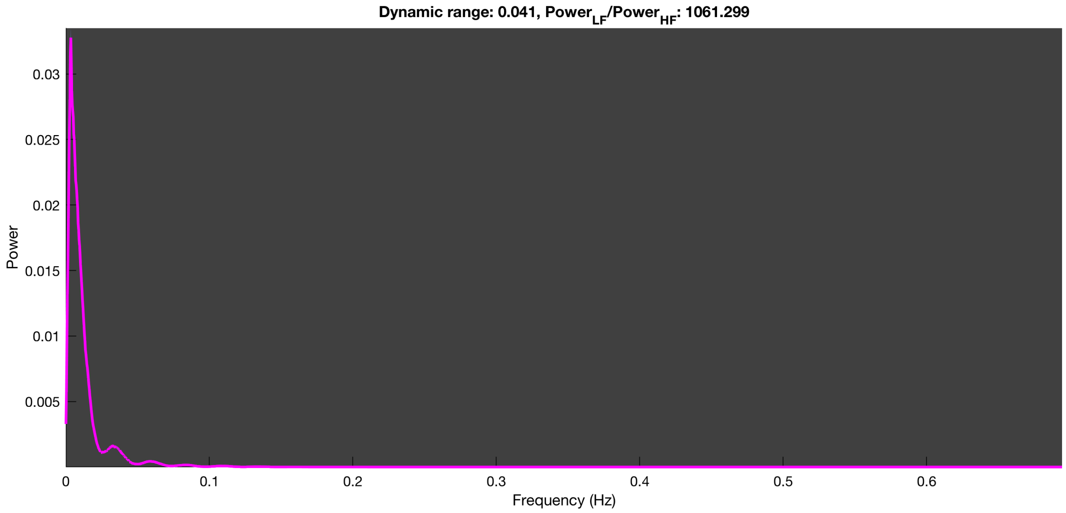

3-1 20n_tmap_component_ica_s1 005

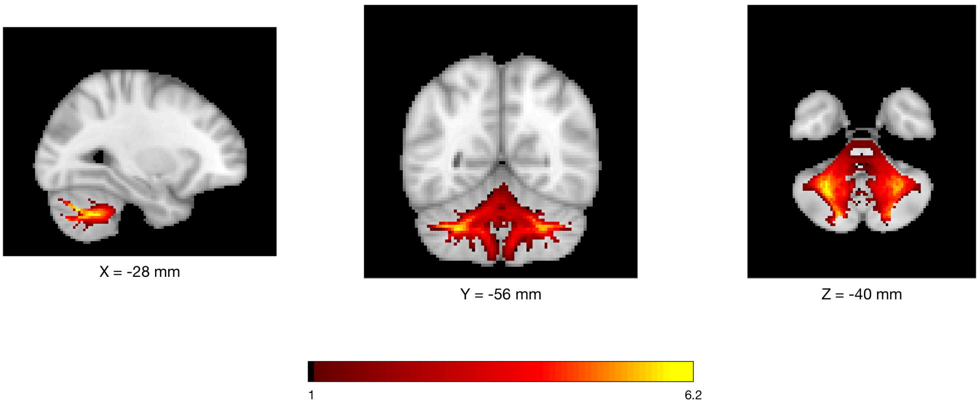


Dynamic range: 0.038, $Power_{LF}/Power_{HF}$: 1089.710

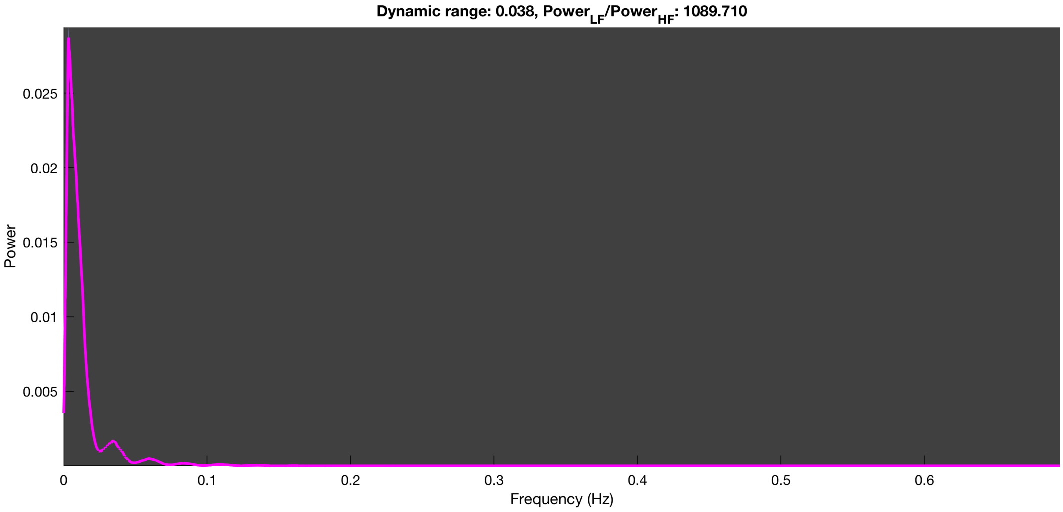

##### 3-1 20n_tmap_component_ica_s1 006

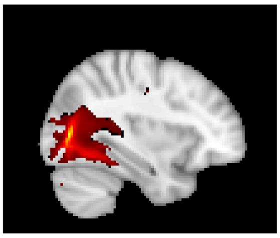

X = 34 mm

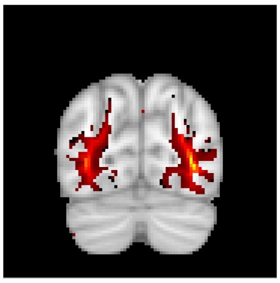

Y = -74 mm

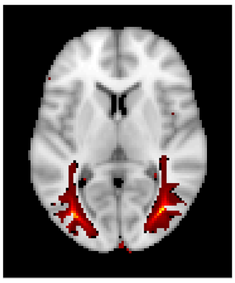

Z = 6 mm

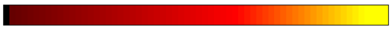


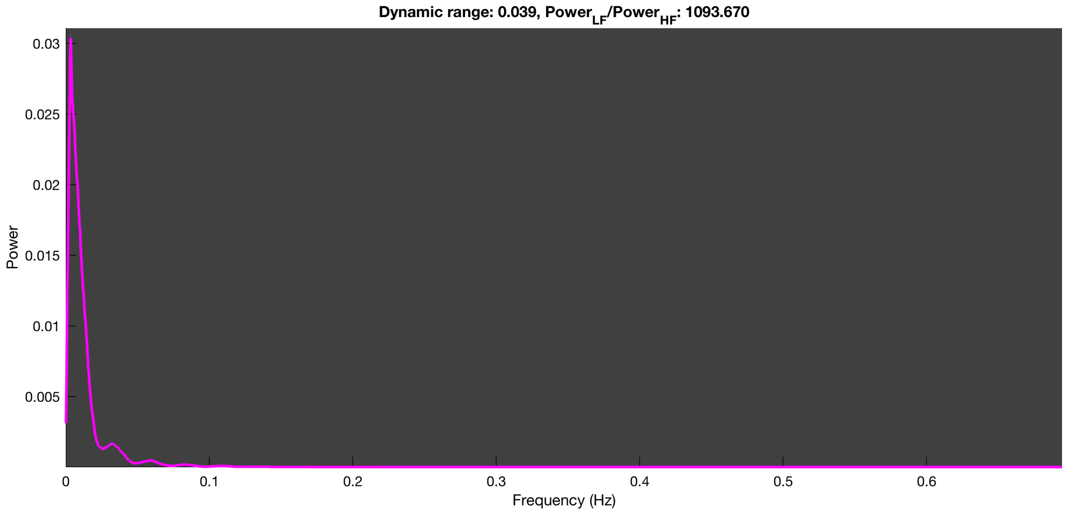

3-1 20n_tmap_component_ica_s1 007

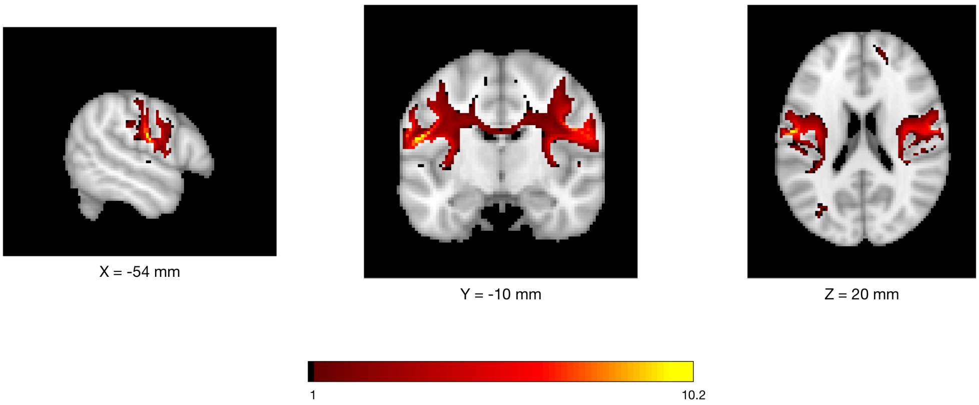


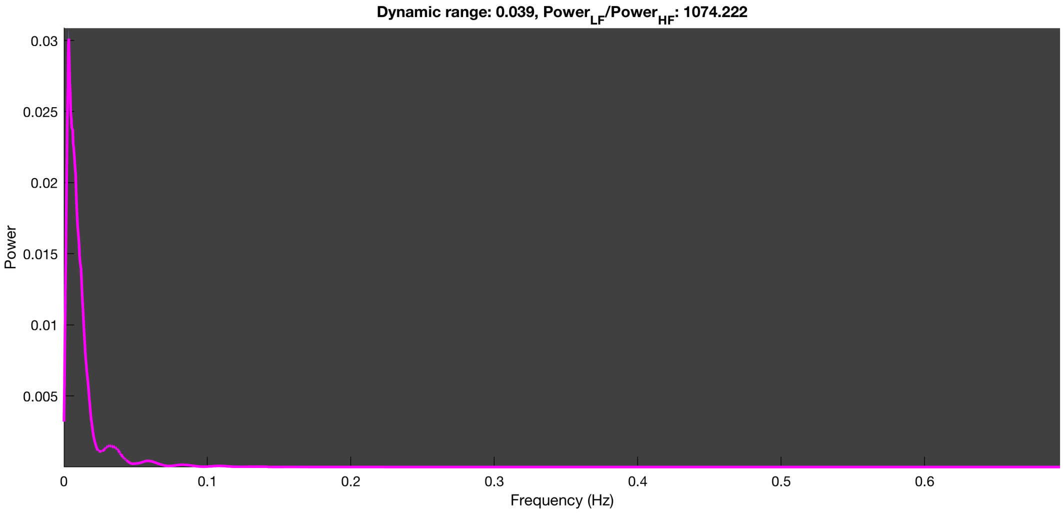

3-1 20n_tmap_component_ica_s1 008

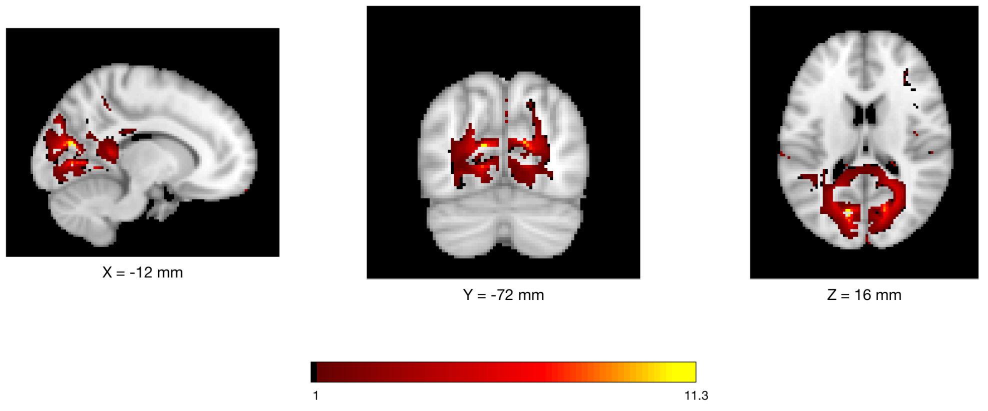


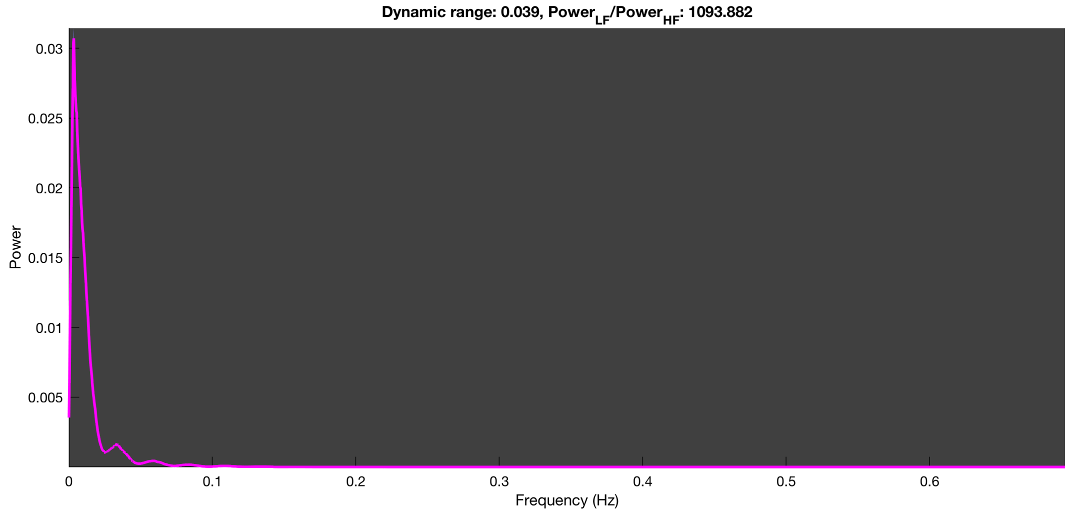

3-1 20n_tmap_component_ica_s1 009

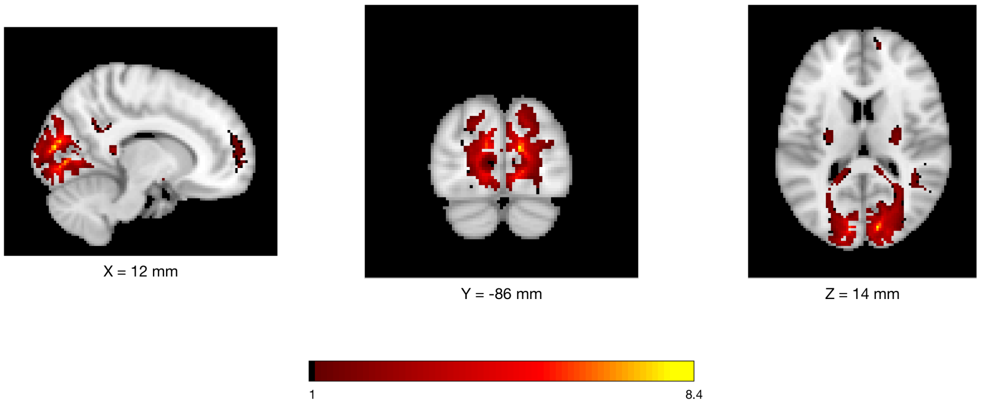


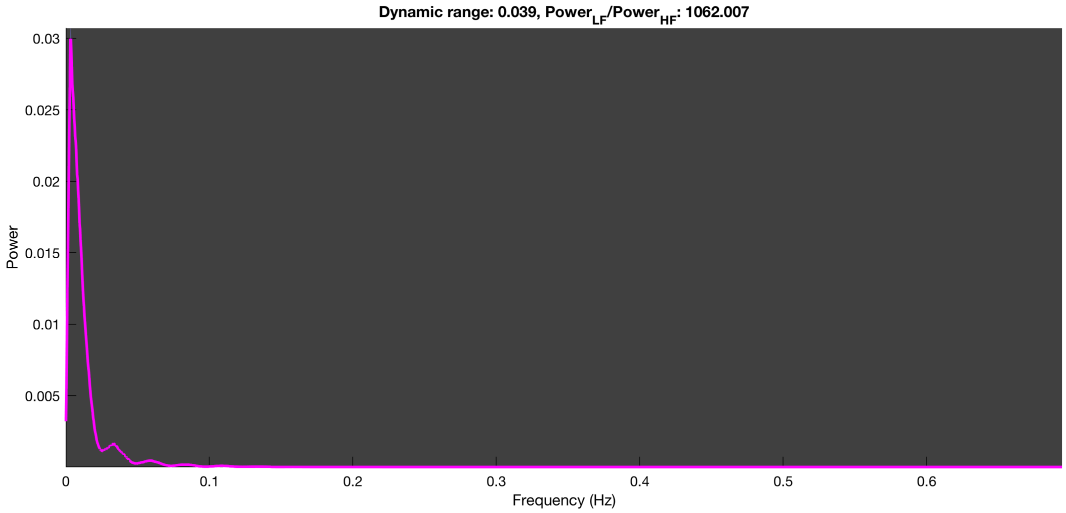

3-1 20n_tmap_component_ica_s1 010

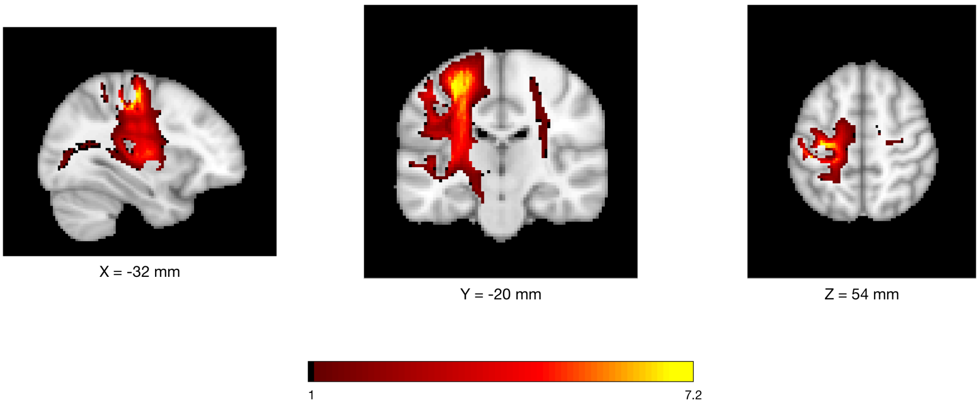


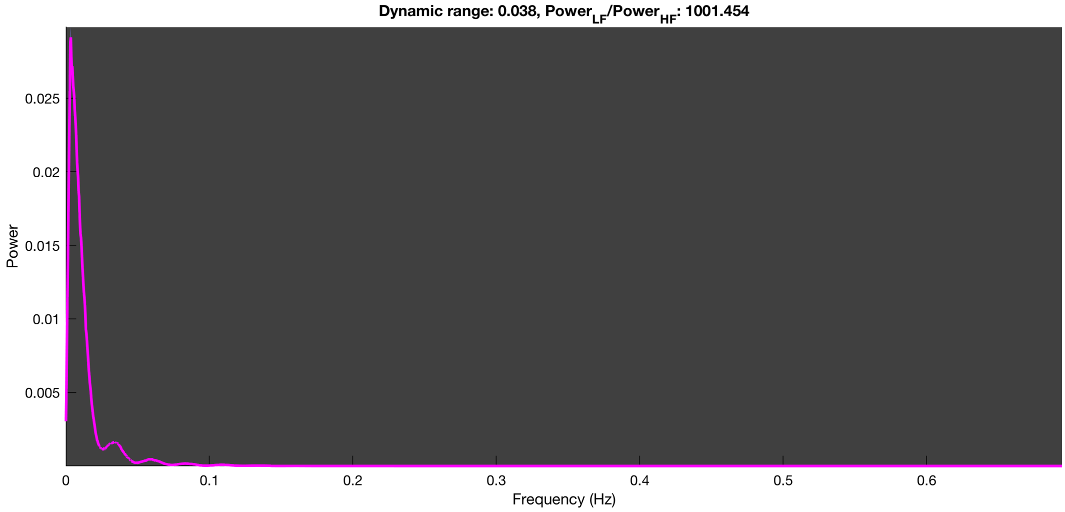

#### 3-1 20n_tmap_component_ica_s1 011

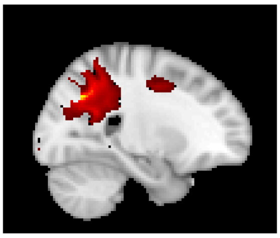

X = 24 mm

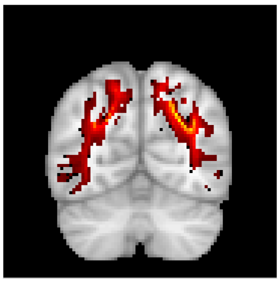

Y = -64 mm

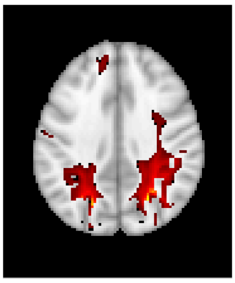

Z = 36 mm

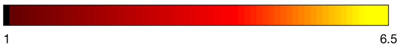


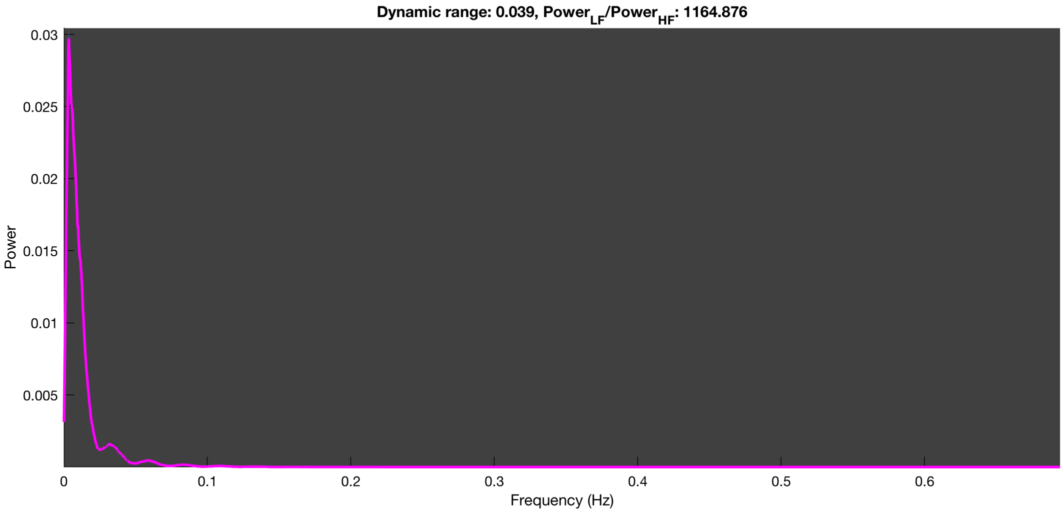

#### 3-1 20n_tmap_component_ica_s1 012

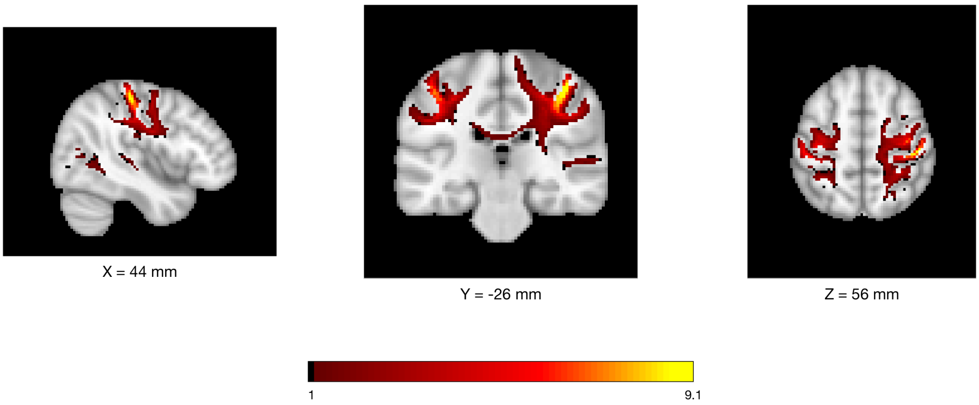


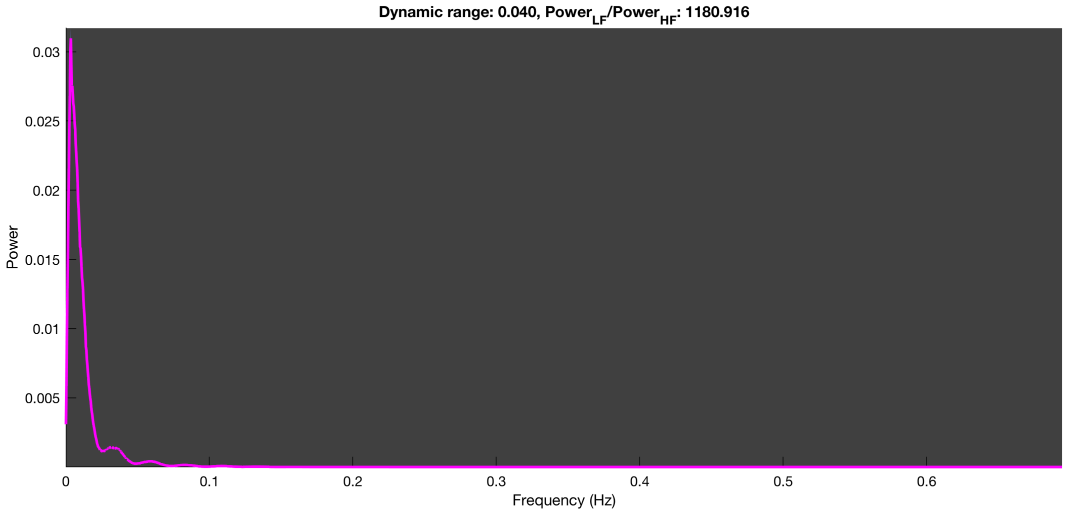

3-1 20n_tmap_component_ica_s1 013

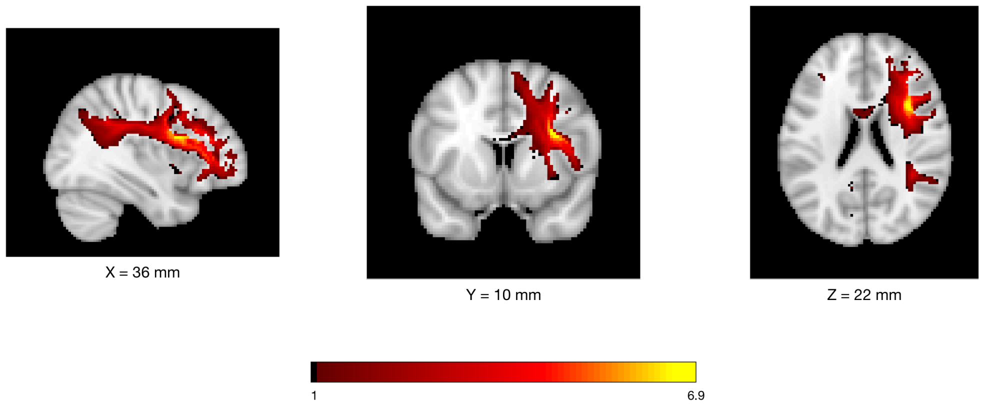


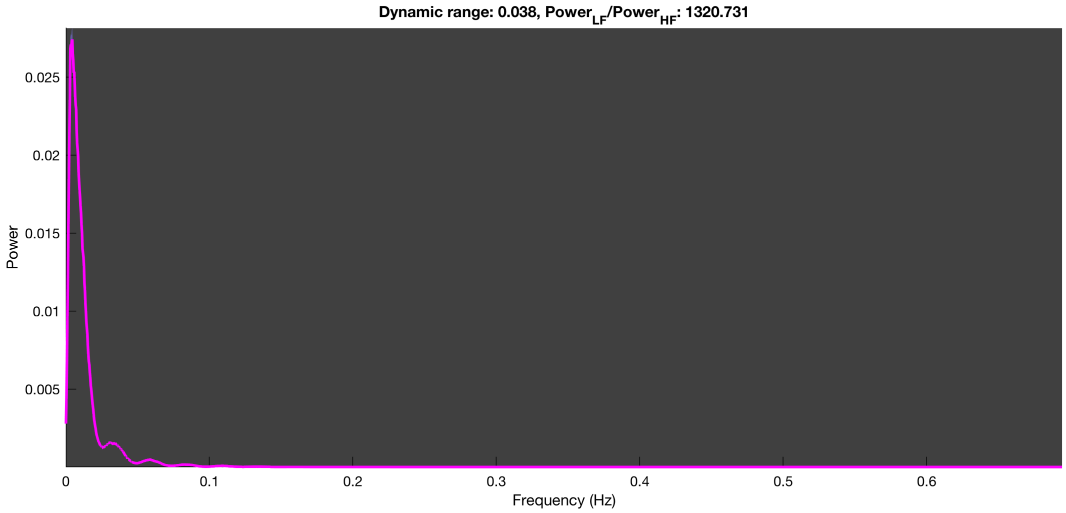

3-1 20n_tmap_component_ica_s1 014

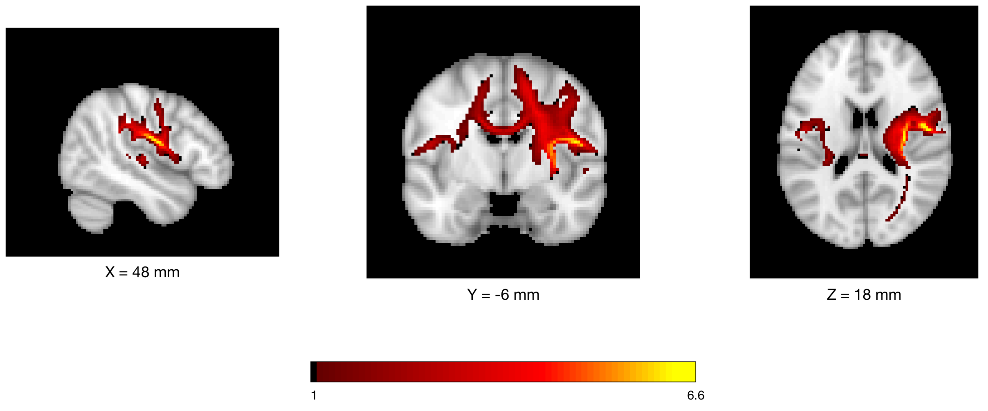


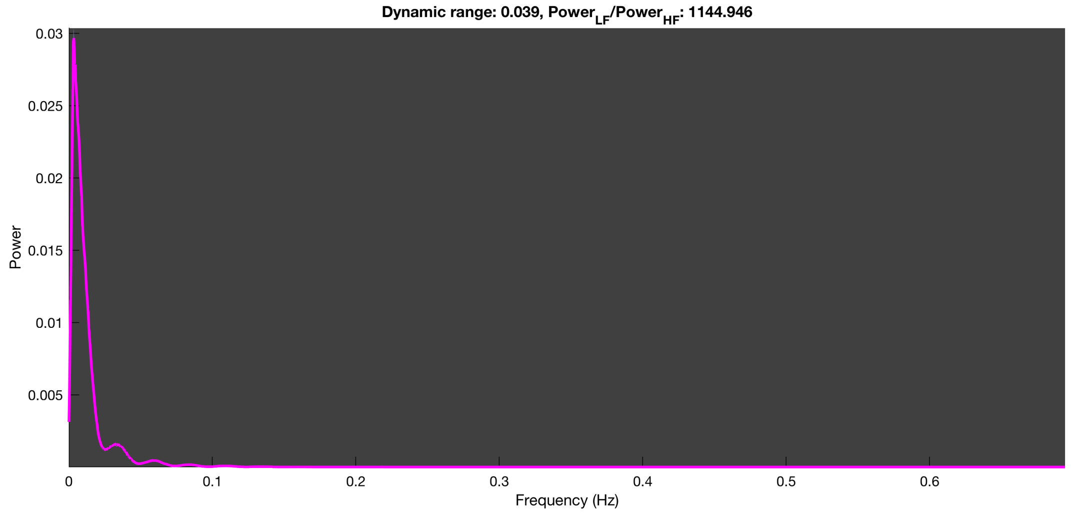

3-1 20n_tmap_component_ica_s1 015

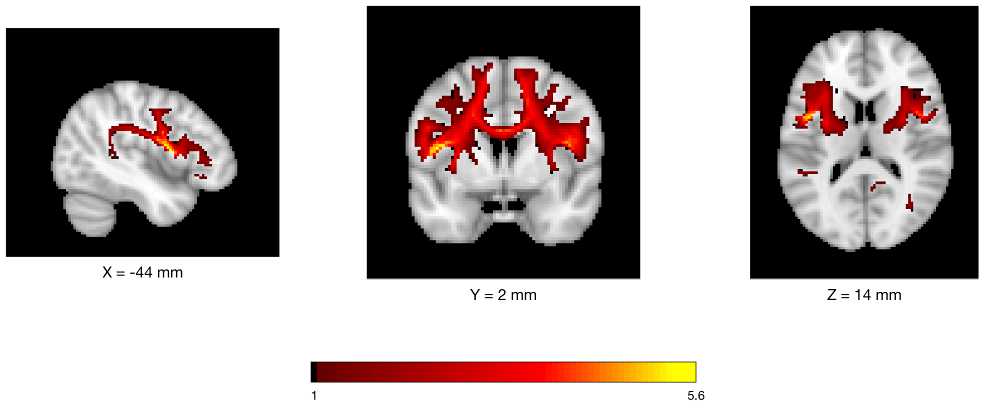


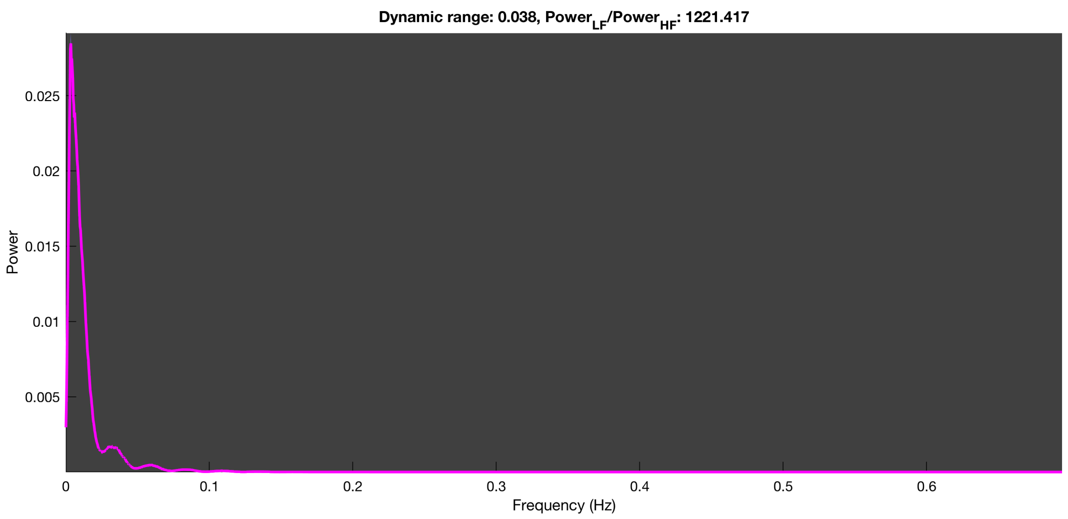

3-1 20n_tmap_component_ica_s1 016

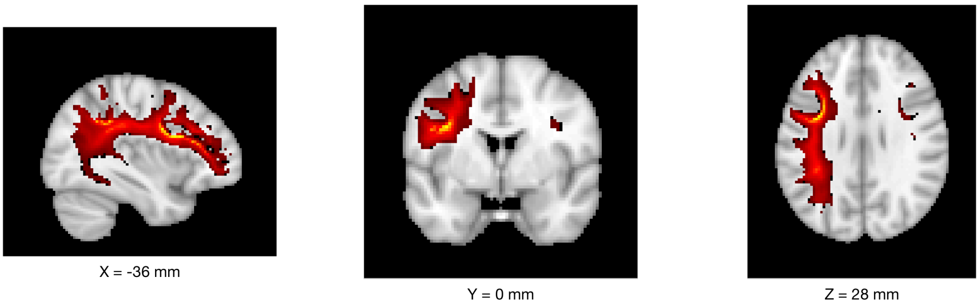


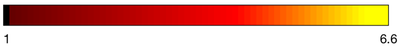


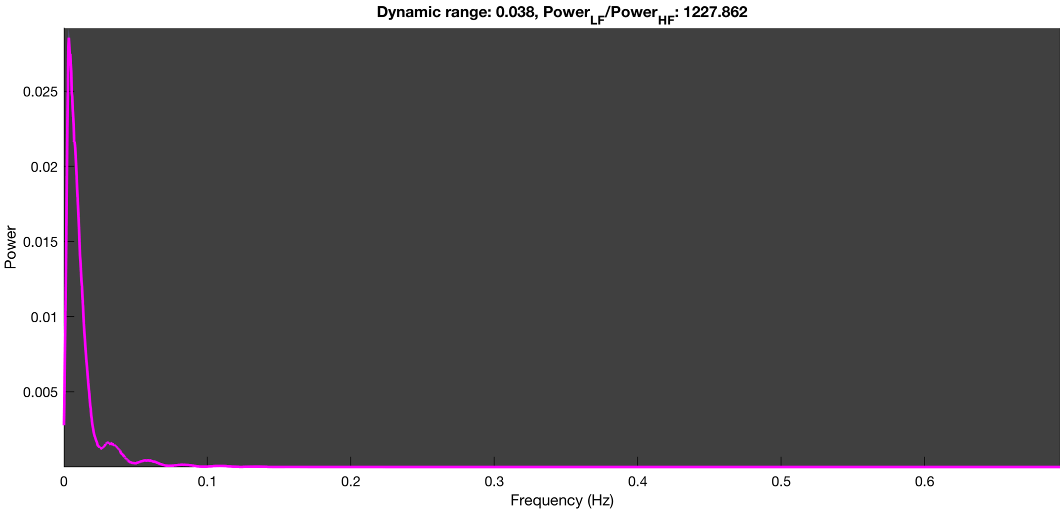

3-1 20n_tmap_component_ica_s1 017

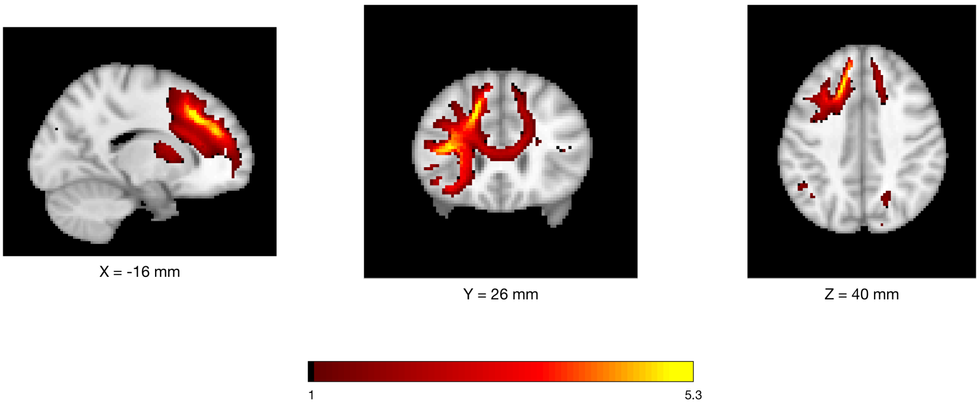


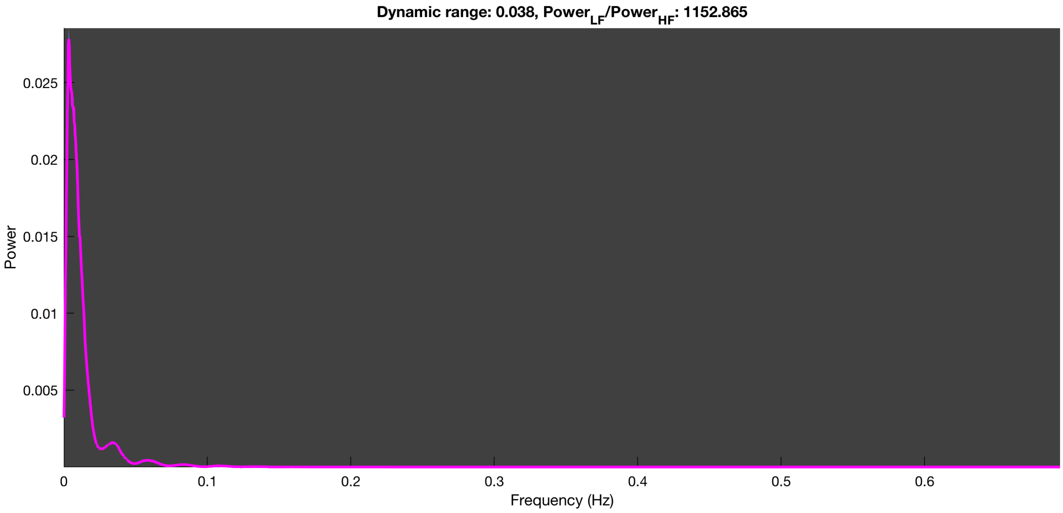

3-1 20n_tmap_component_ica_s1 018

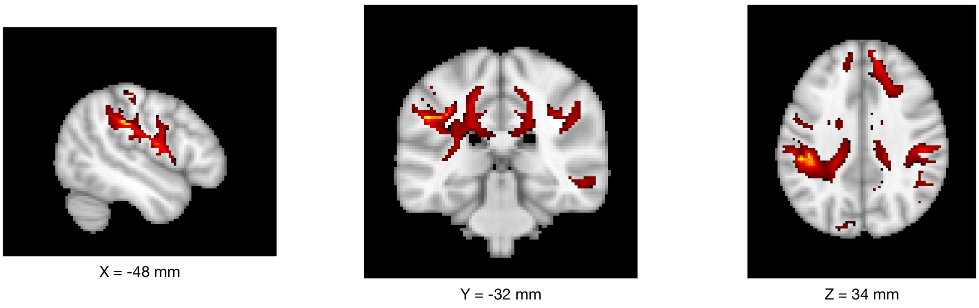


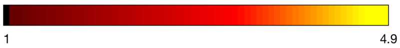


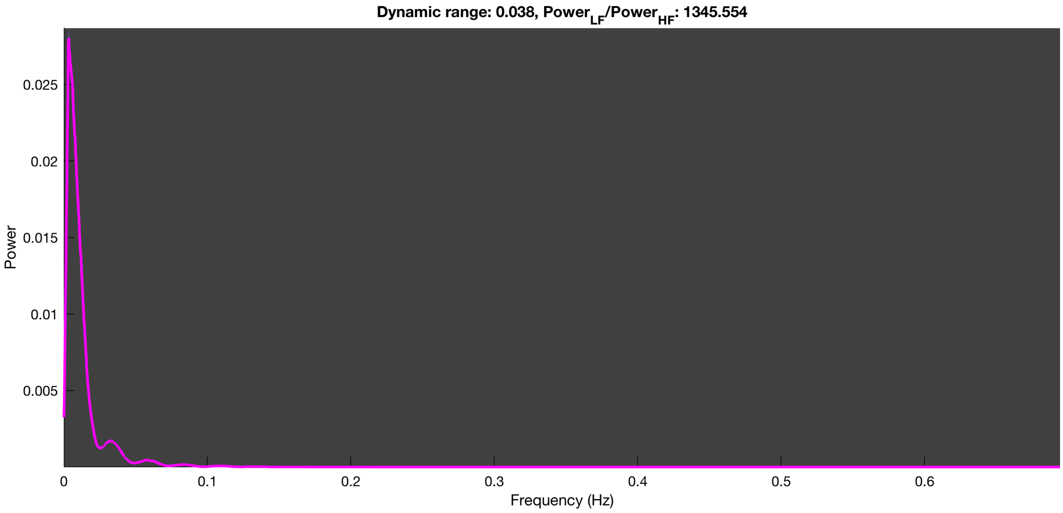

3-1 20n_tmap_component_ica_s1 019

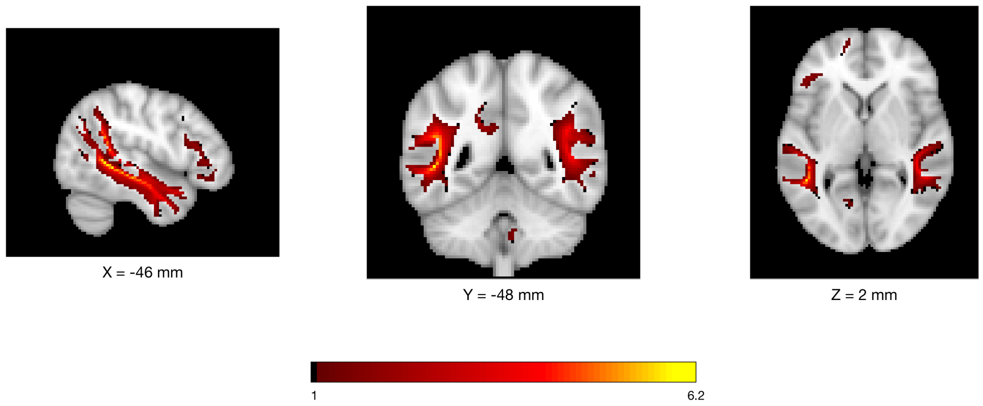


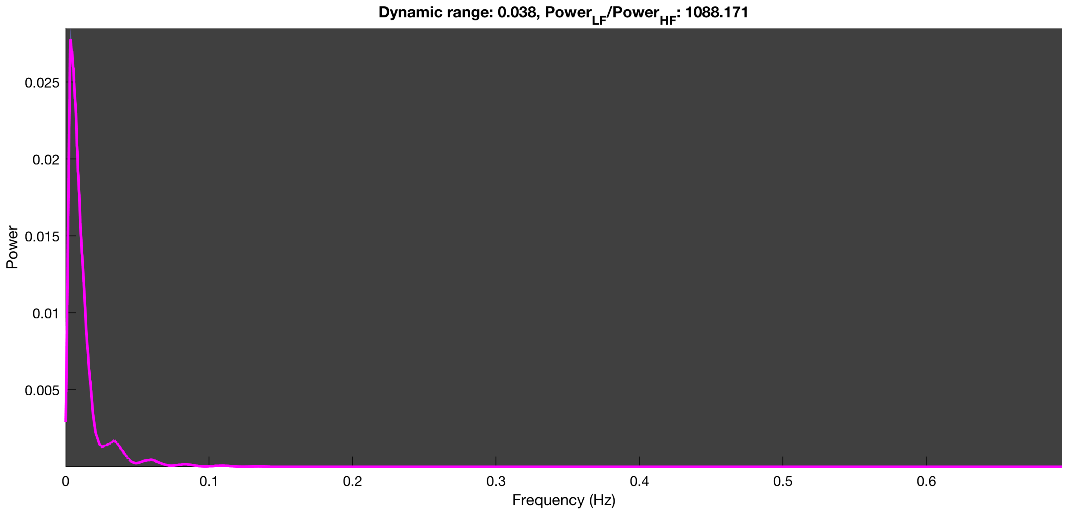

3-1 20n_tmap_component_ica_s1 020

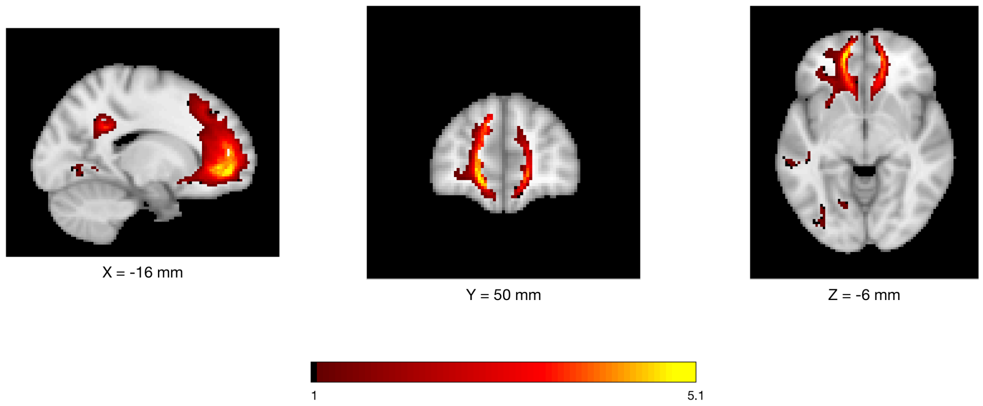


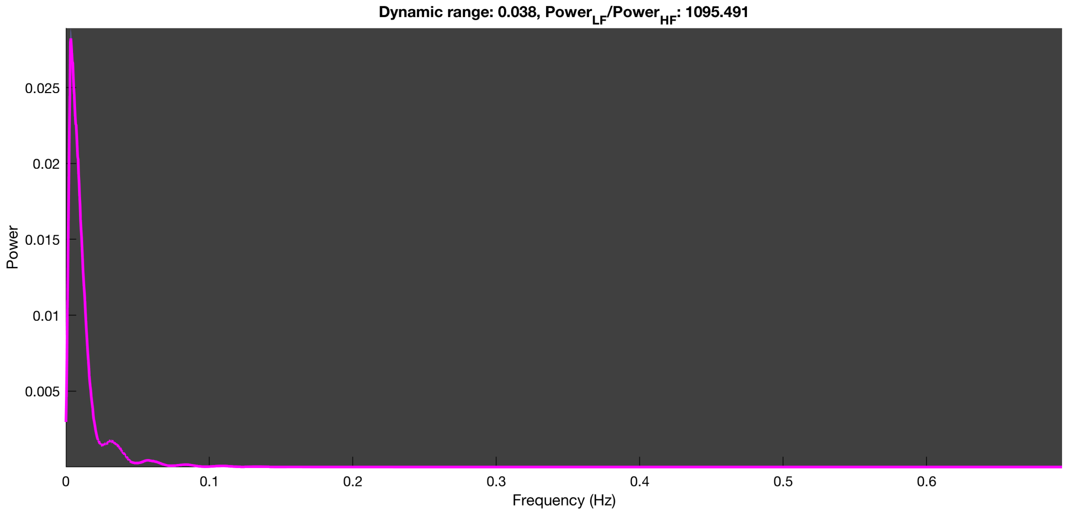

### HCP dataset

*n=100 run*

3-1 100n_tmap_component_ica_s1 001

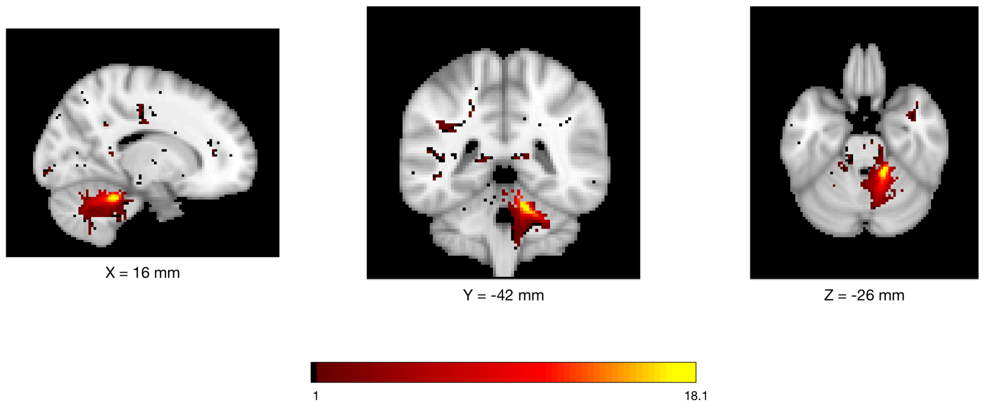


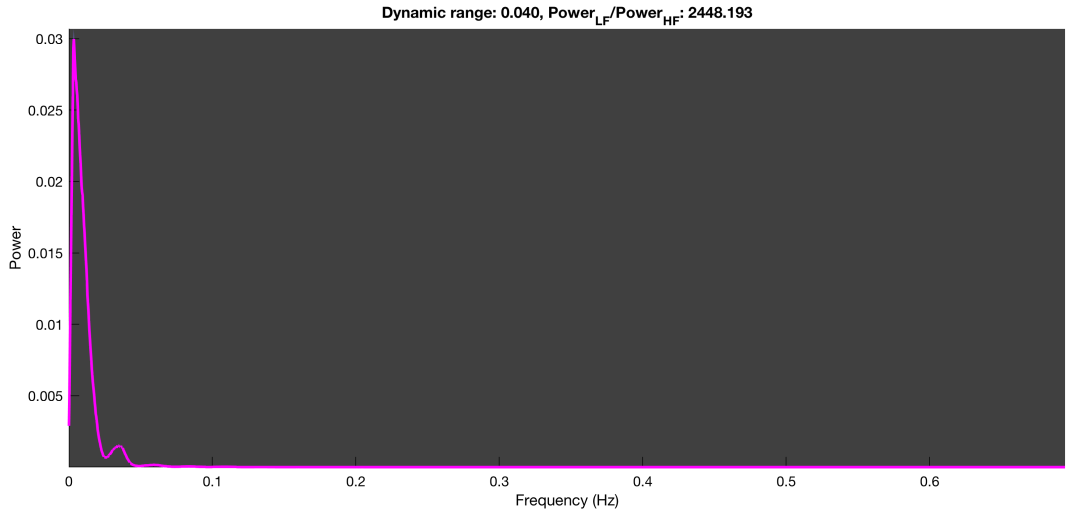

3-1 100n_tmap_component_ica_s1 002

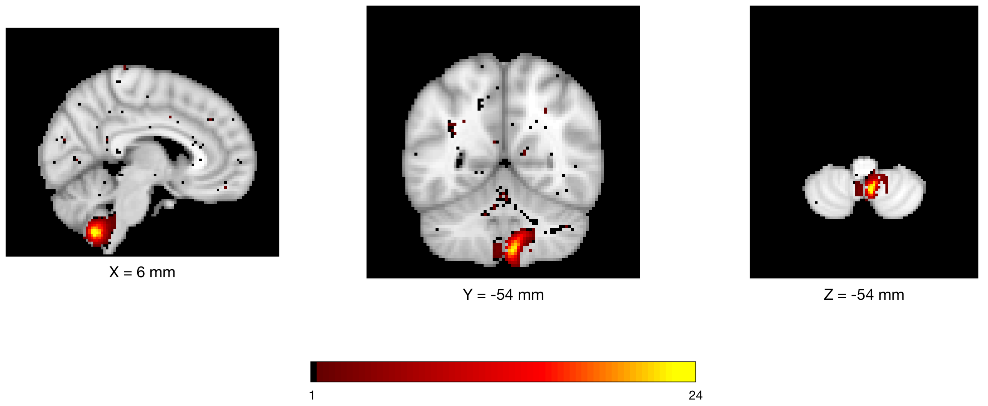


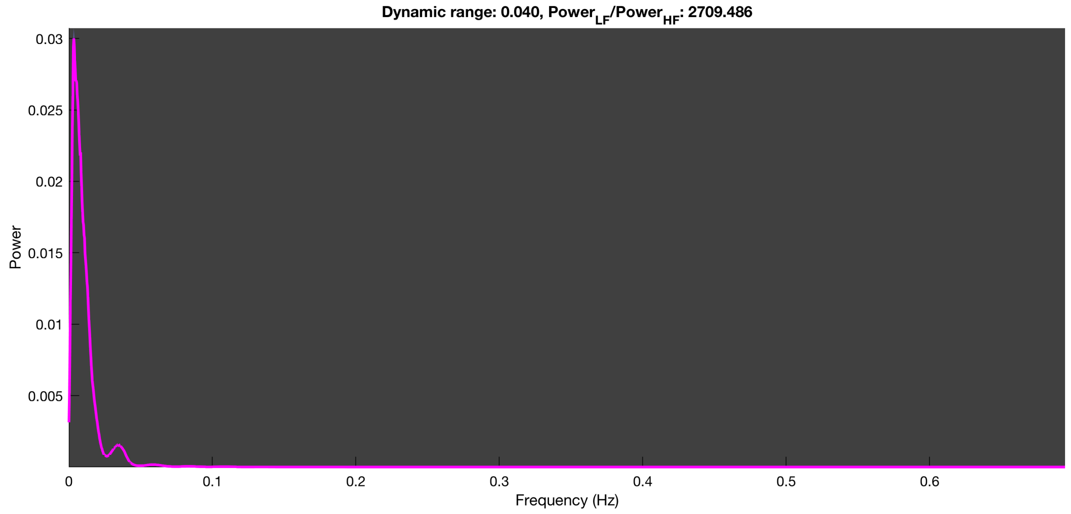

3-1 100n_tmap_component_ica_s1 003

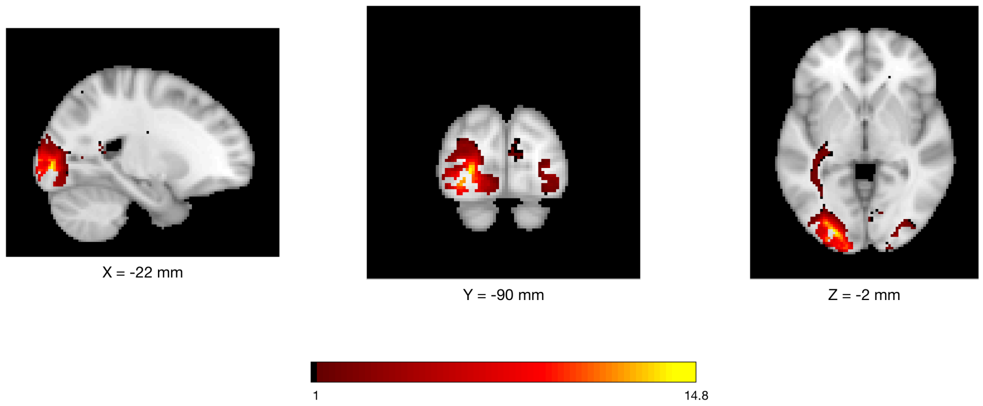


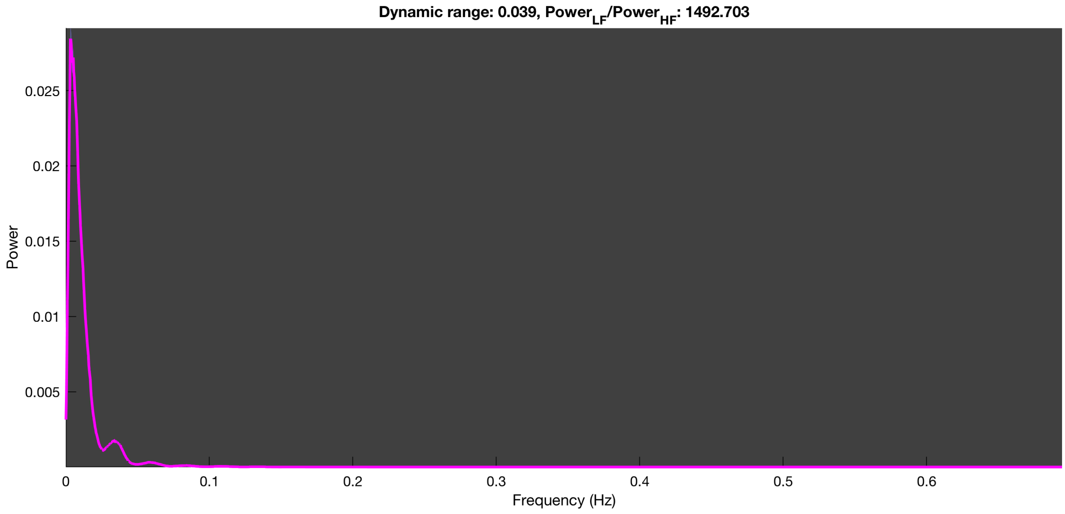

#### 3-1 100n_tmap_component_ica_s1 004

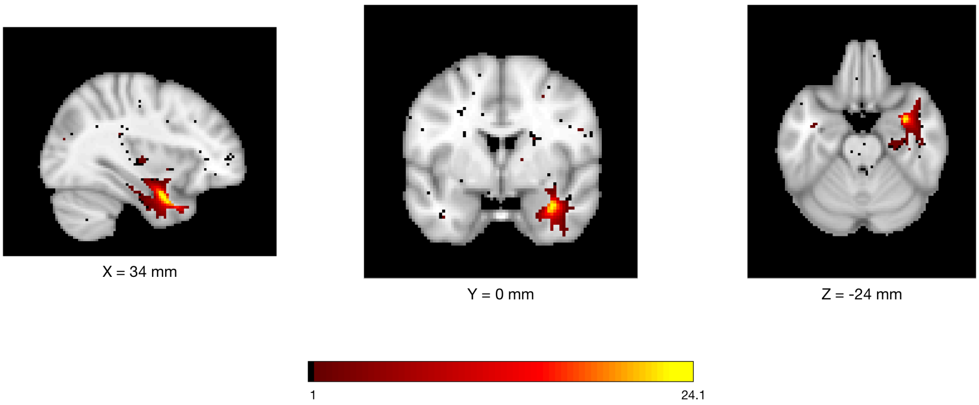


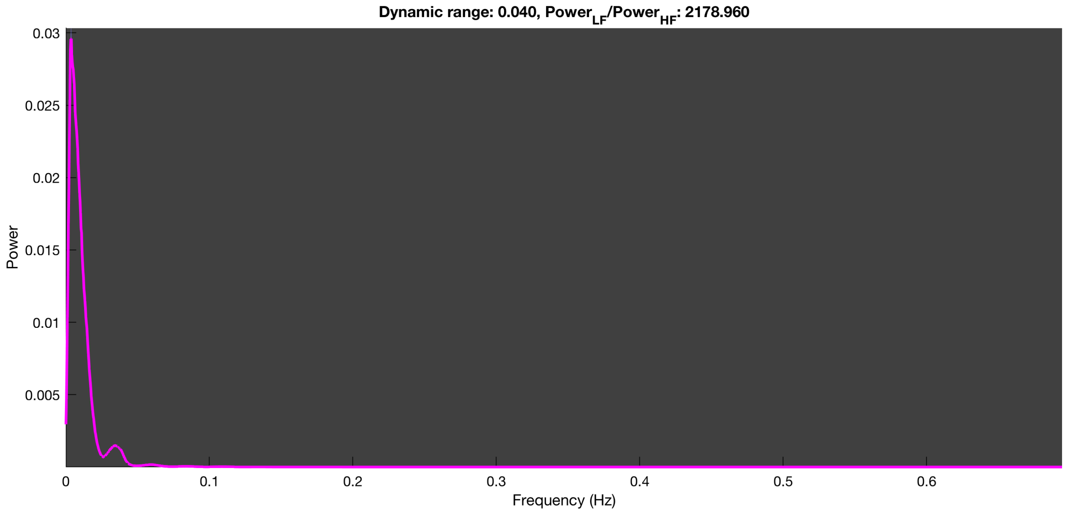

3-1 100n_tmap_component_ica_s1 005

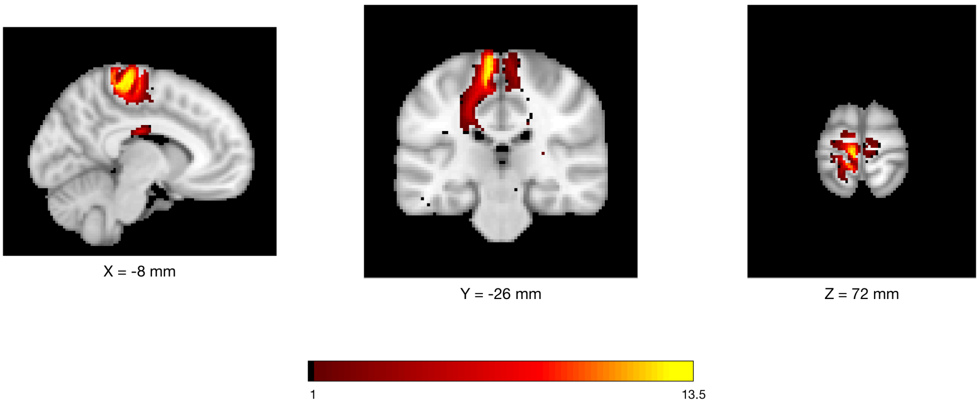


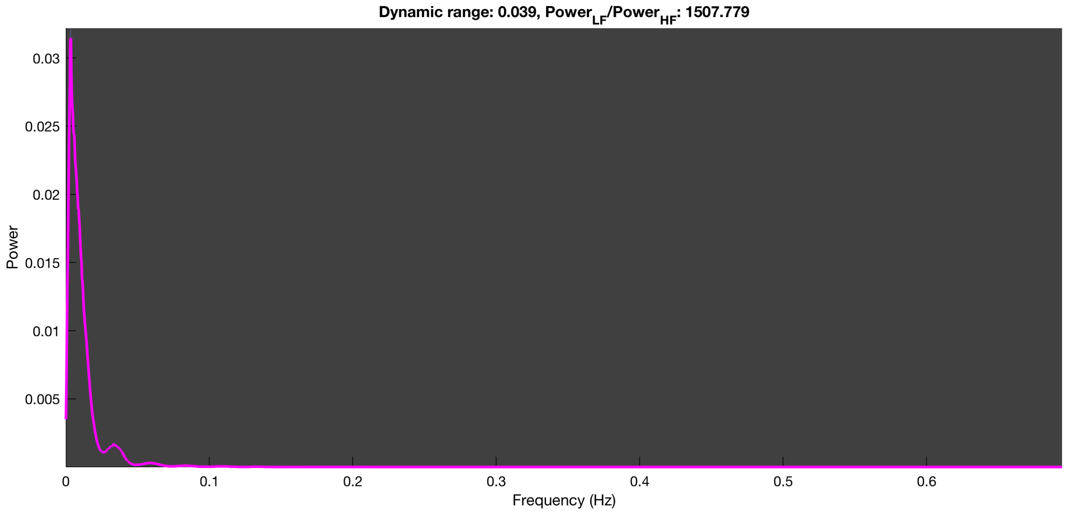

3-1 100n_tmap_component_ica_s1 006

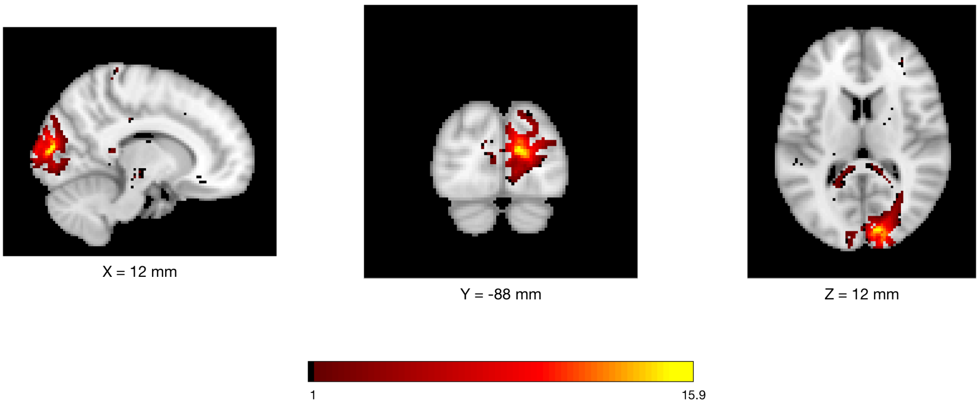


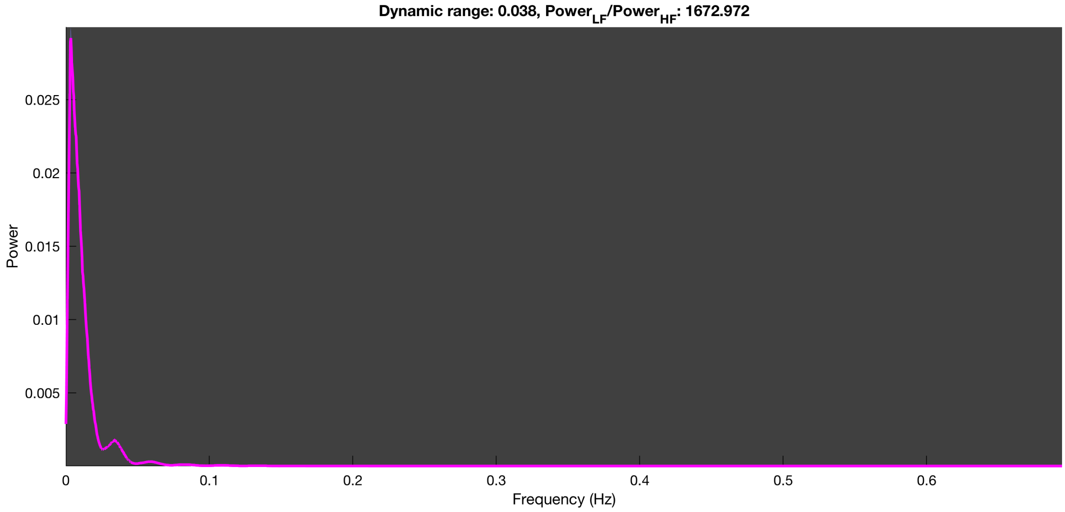

3-1 100n_tmap_component_ica_s1 007

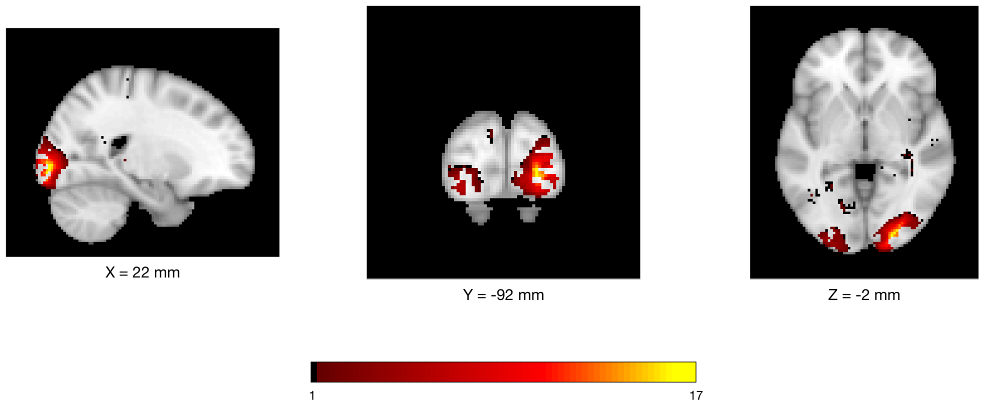


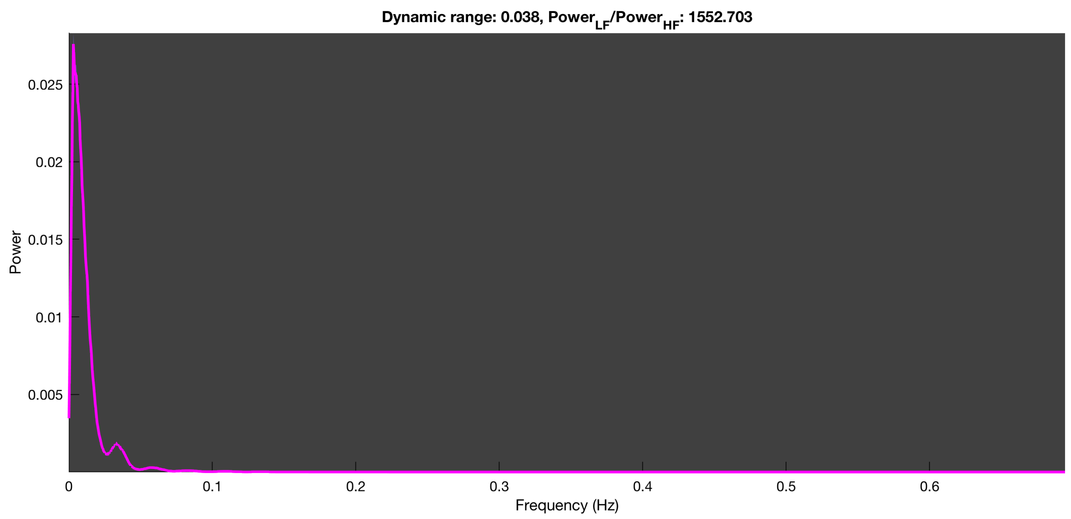

3-1 100n_tmap_component_ica_s1 008

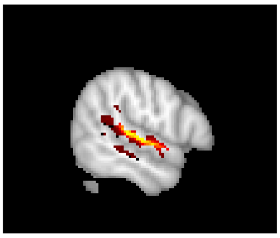

X = 56 mm

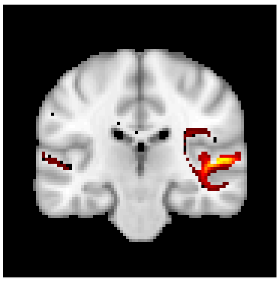

Y = -24 mm

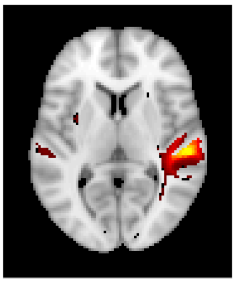

Z = 6 mm

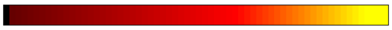


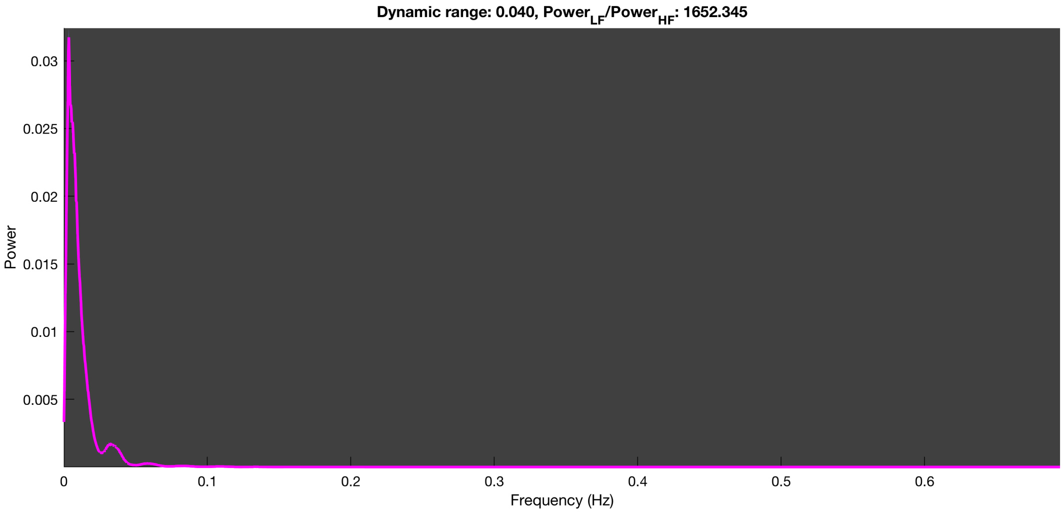

**3-1 100n_tmap_component_ica_s1 009**

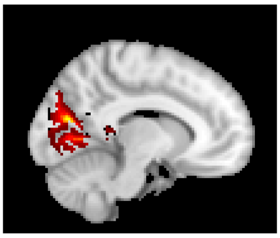

X = -10 mm

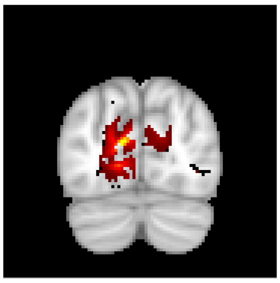

Y = -76 mm

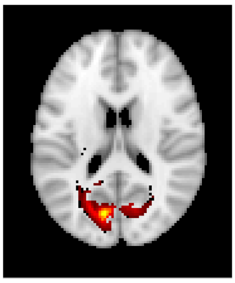

Z = 18 mm

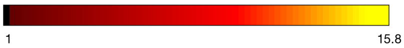


**Dynamic range: 0.040, $Power_{LF}/Power_{HF}$: 1338.582**

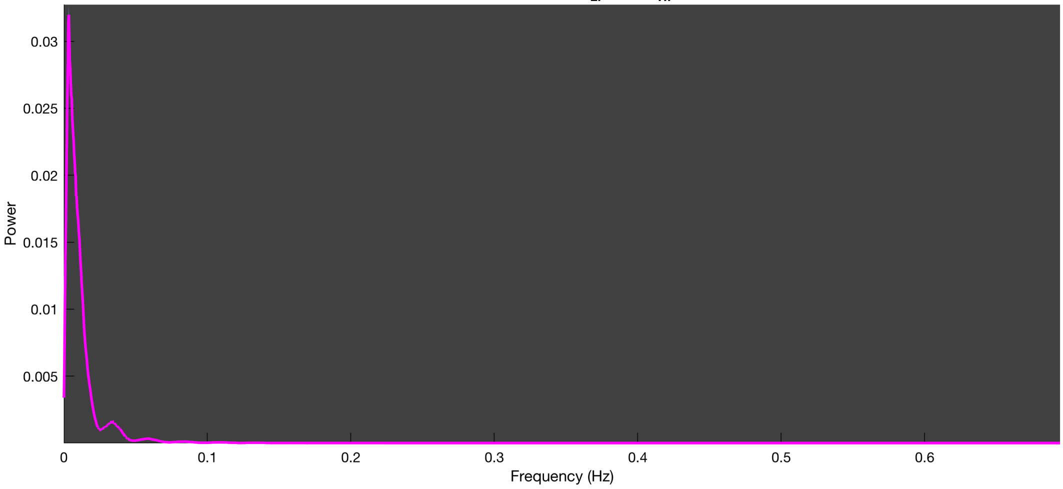

3-1 100n_tmap_component_ica_s1 010

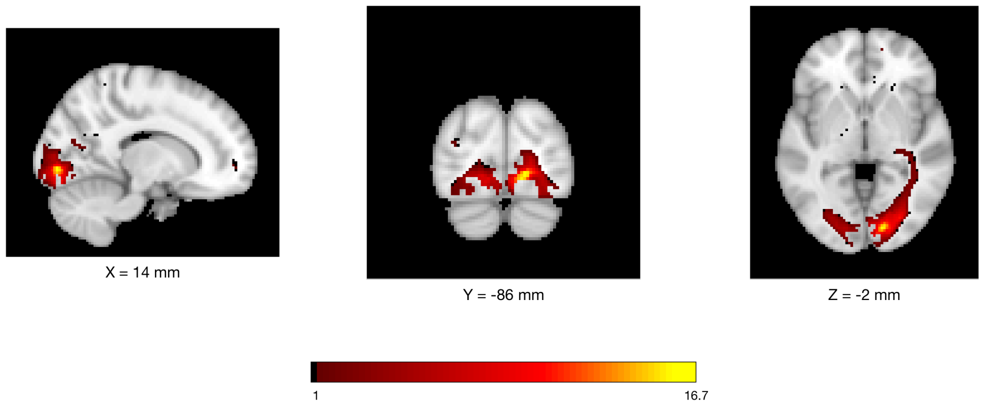


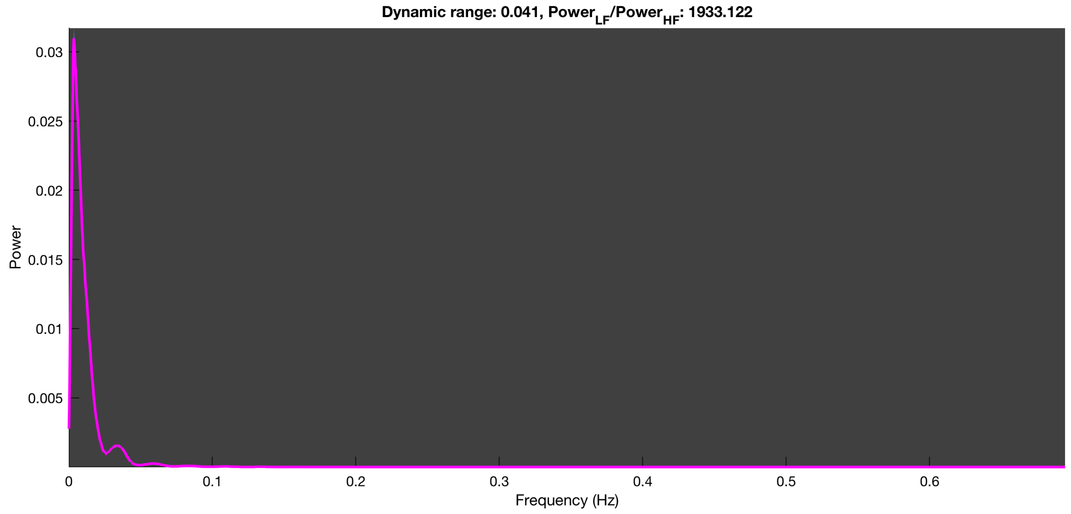

3-1 100n_tmap_component_ica_s1 011

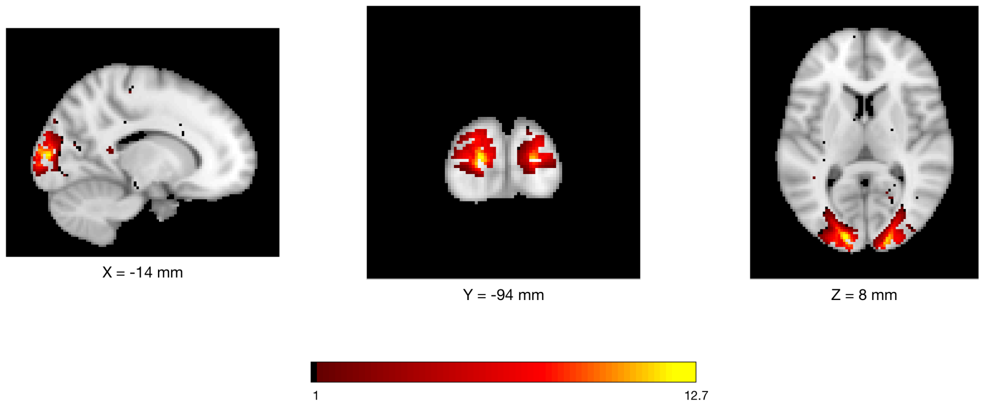


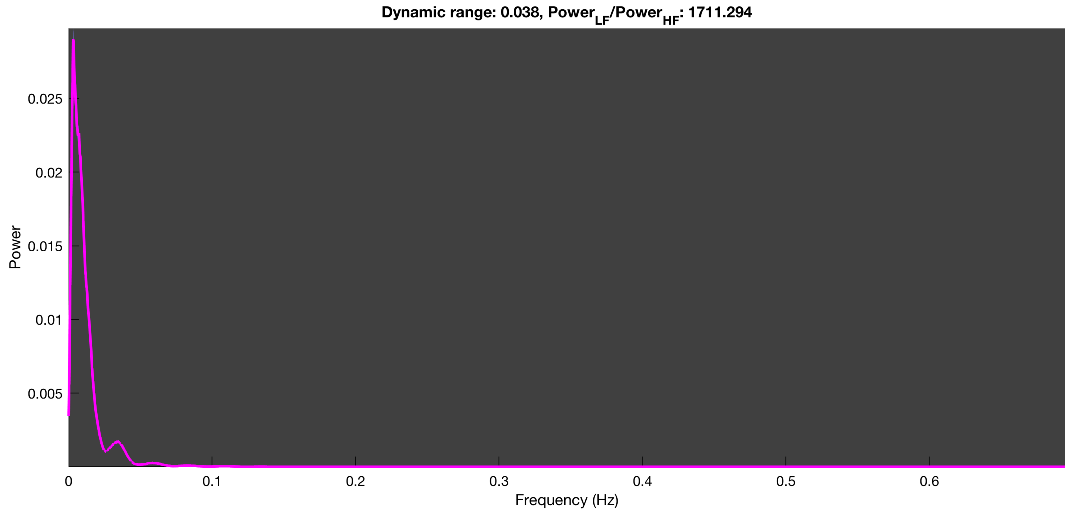

3-1 100n_tmap_component_ica_s1 012

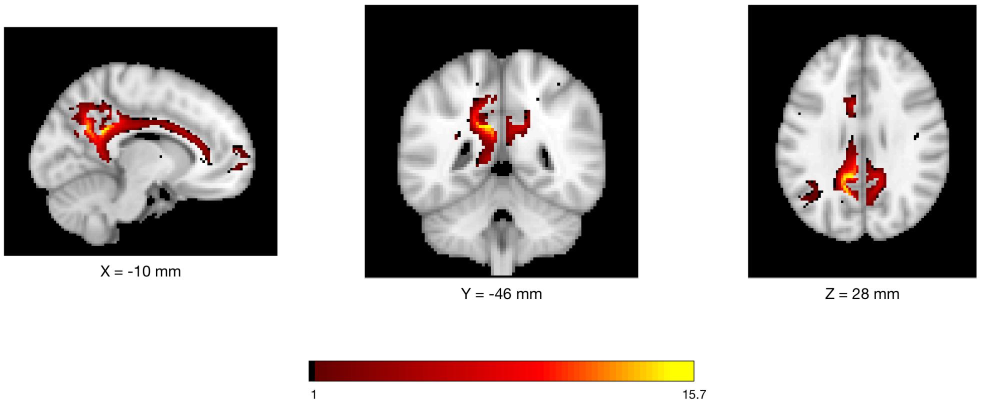


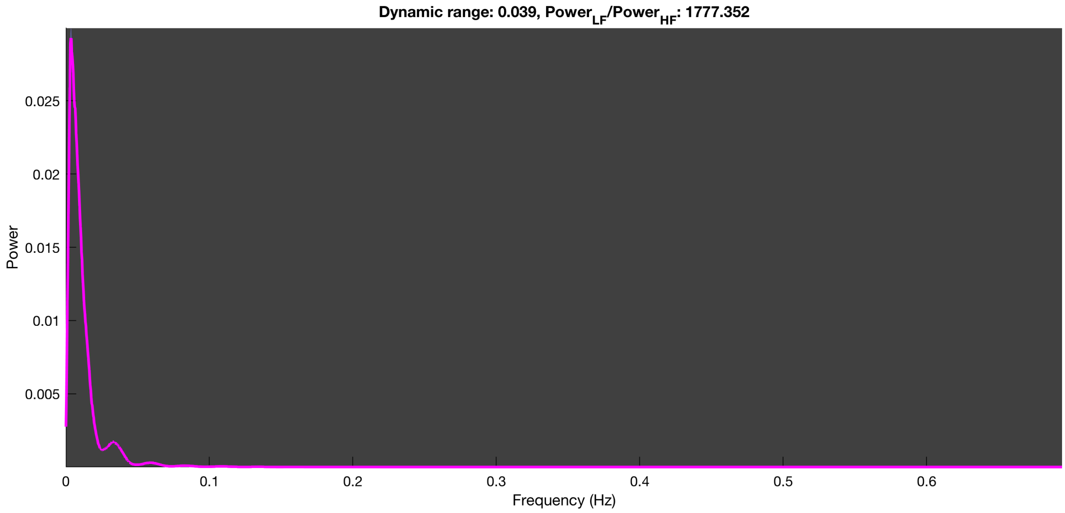

#### 3-1 100n_tmap_component_ica_s1 013

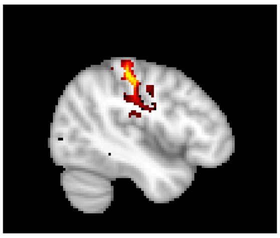

X = -44 mm

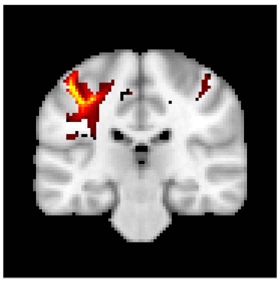

Y = -26 mm

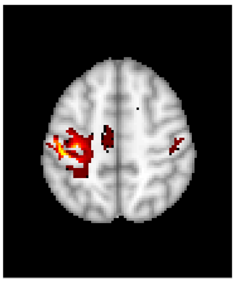

Z = 52 mm

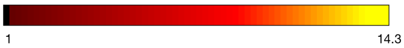


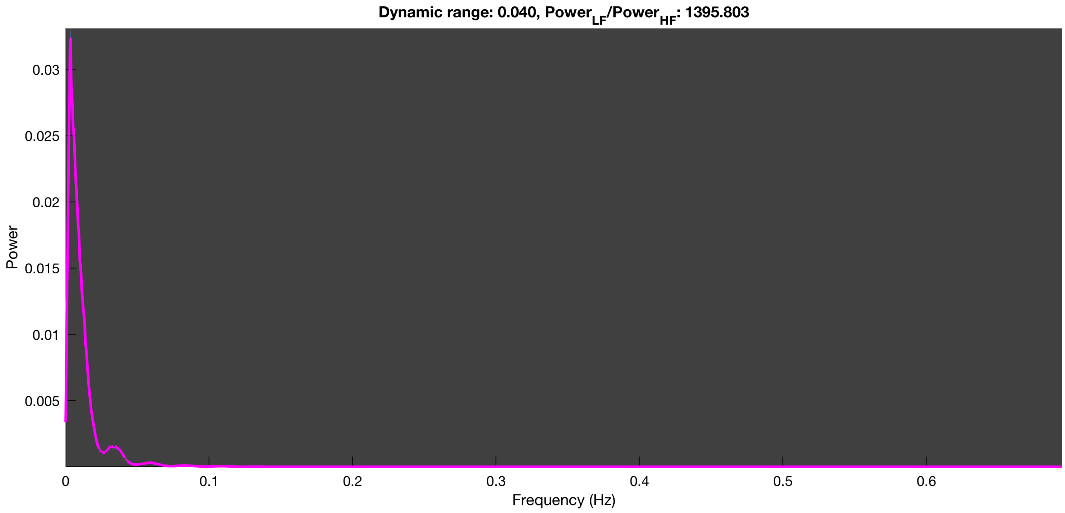

3-1 100n_tmap_component_ica_s1 014

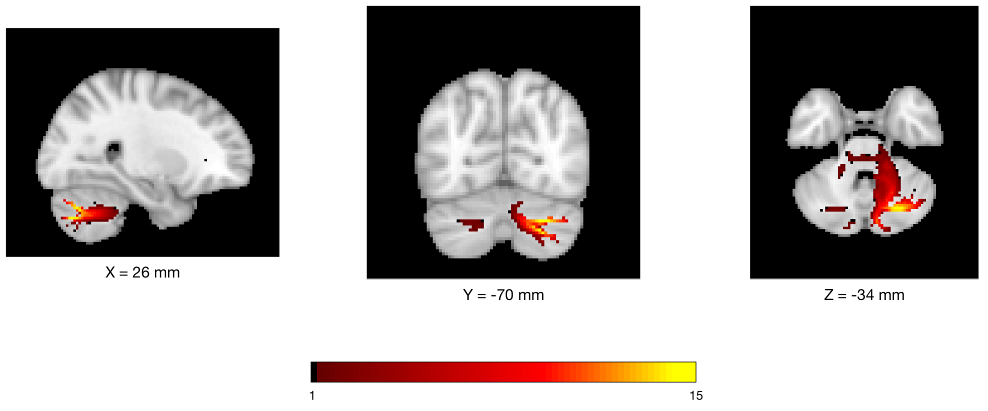


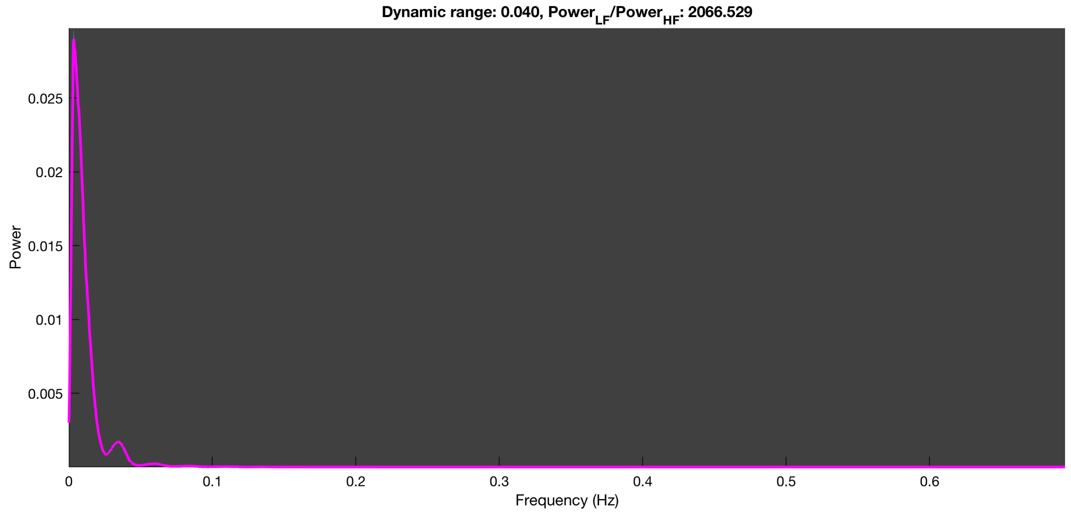

3-1 100n_tmap_component_ica_s1 015

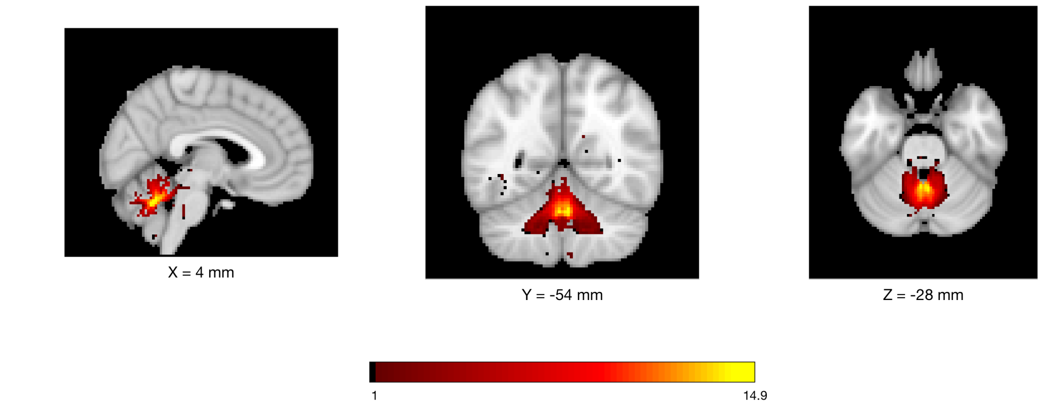


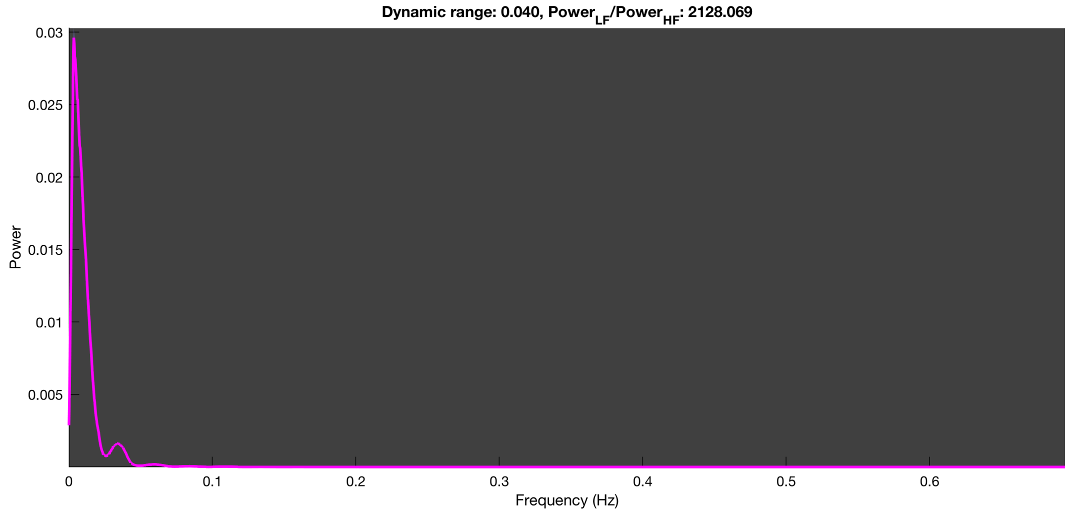

3-1 100n_tmap_component_ica_s1 016

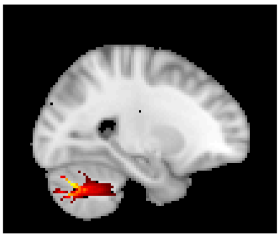

X = -24 mm

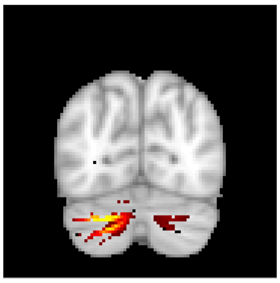

Y = -72 mm

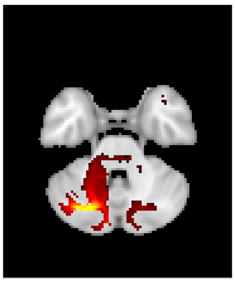

Z = -34 mm

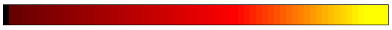


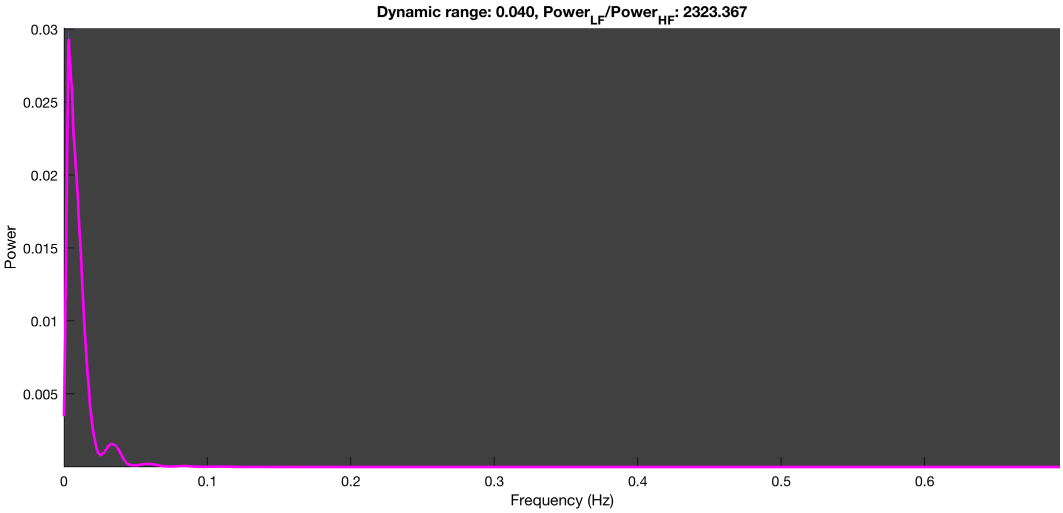

3-1 100n_tmap_component_ica_s1 017

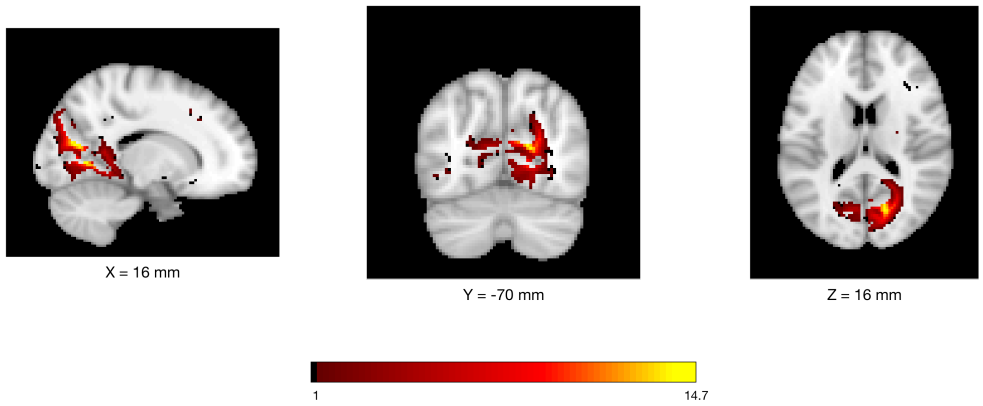


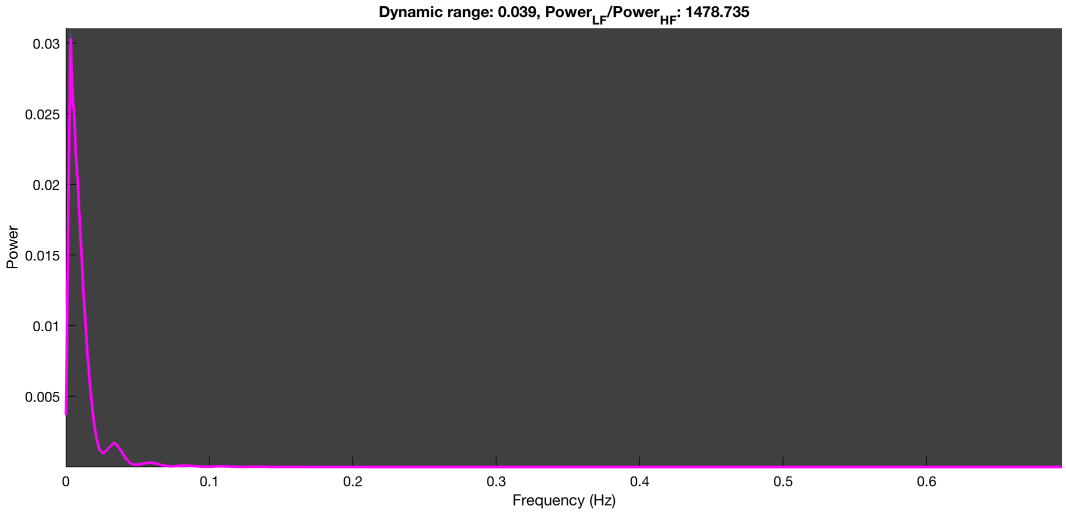

3-1 100n_tmap_component_ica_s1 018

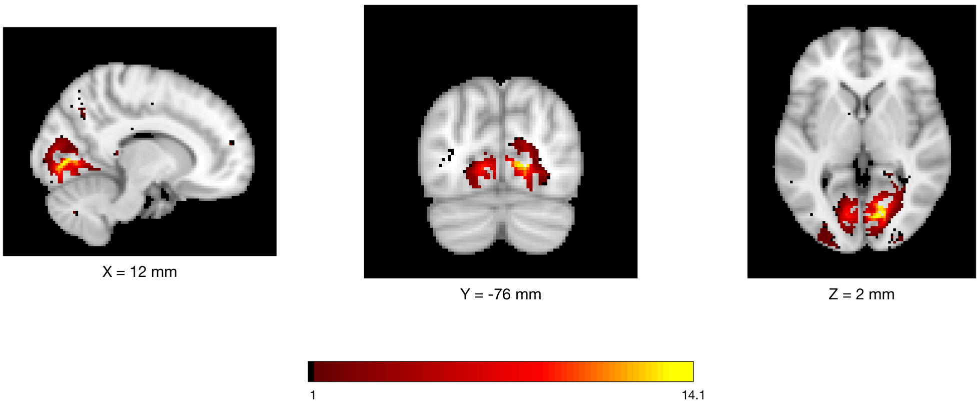


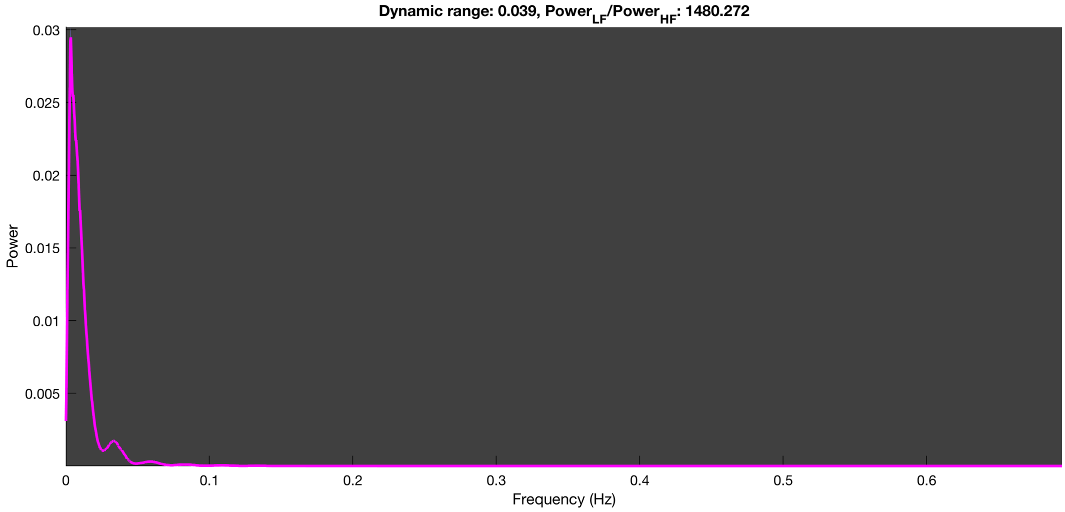

3-1 100n_tmap_component_ica_s1 019

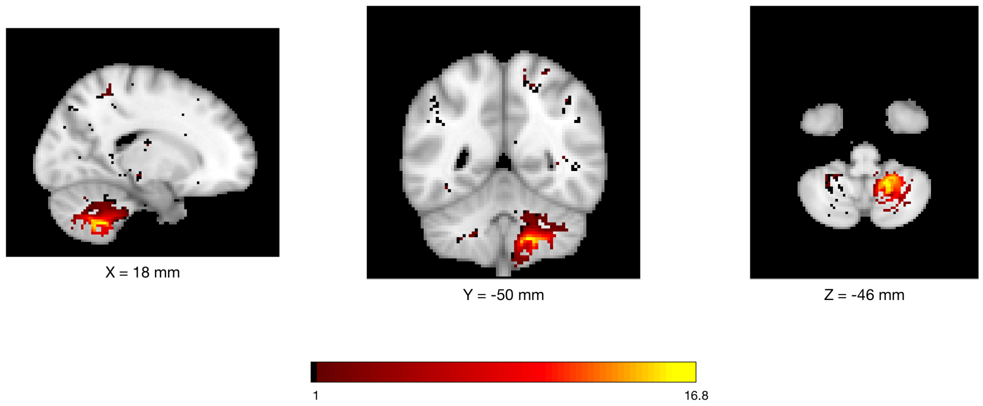


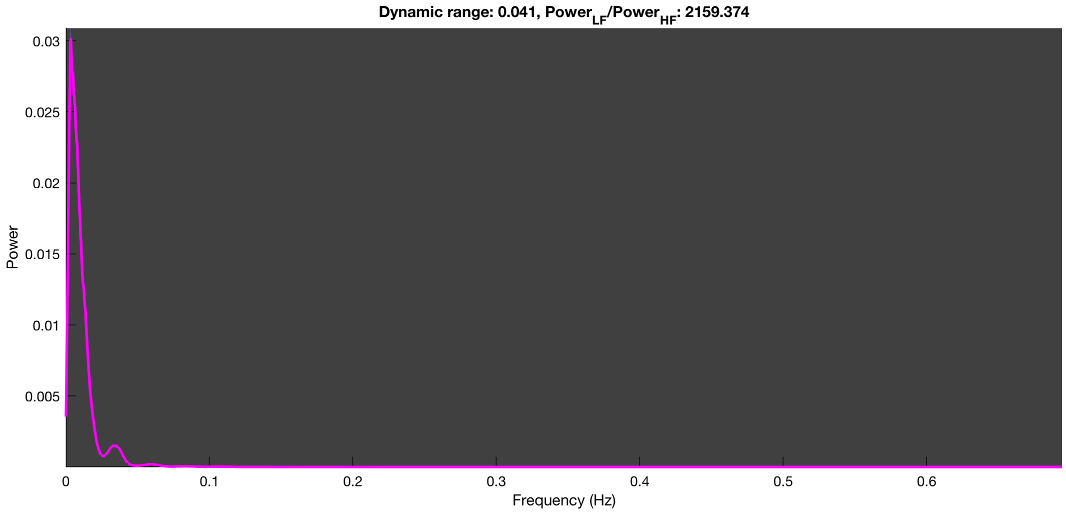

3-1 100n_tmap_component_ica_s1 020

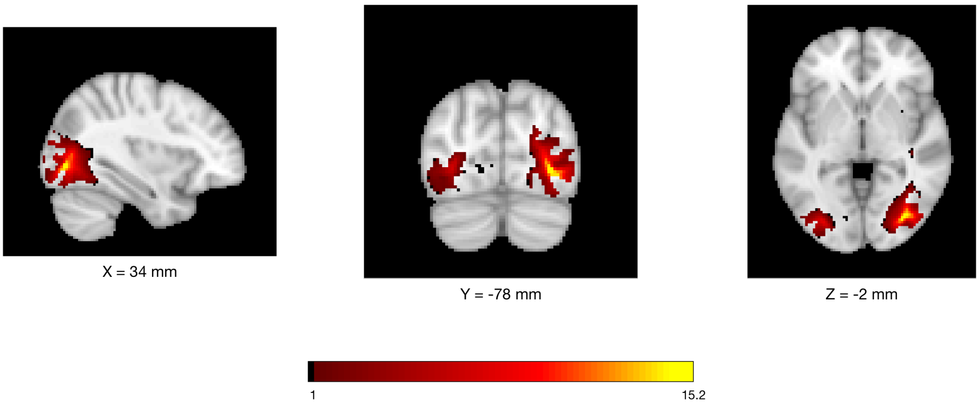


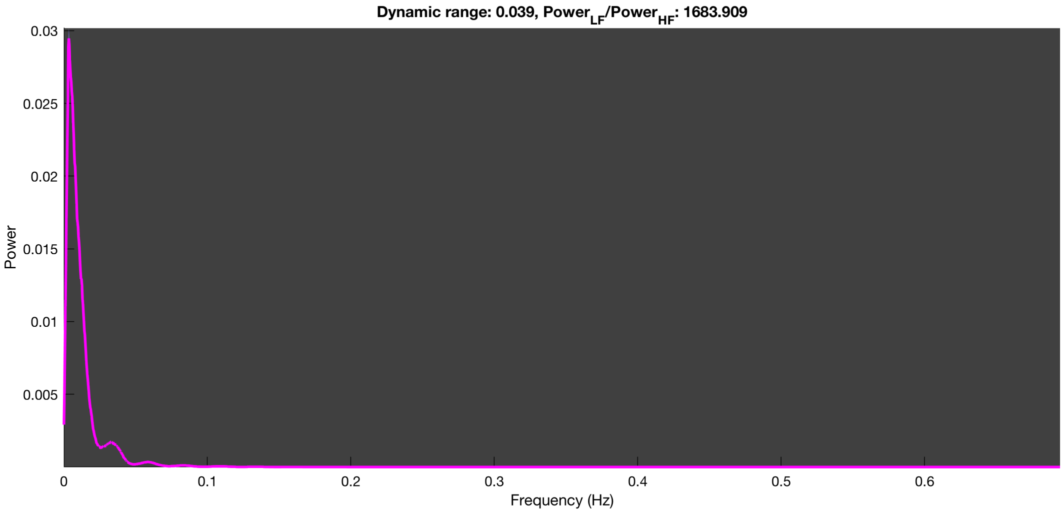

3-1 100n_tmap_component_ica_s1 021

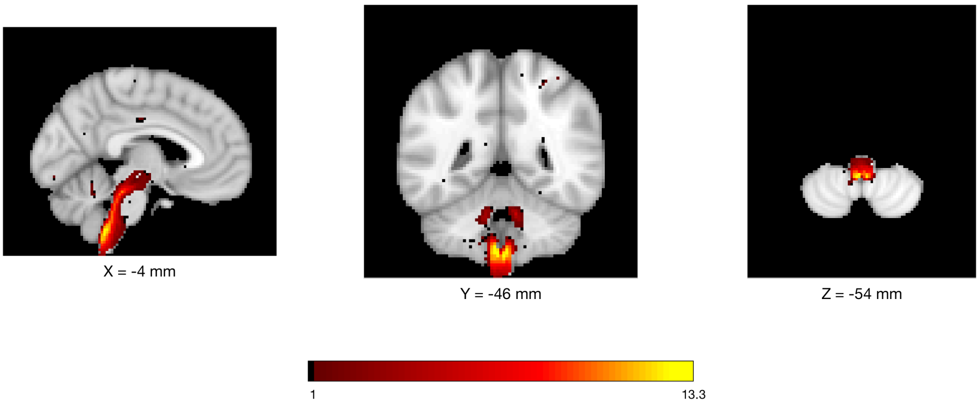


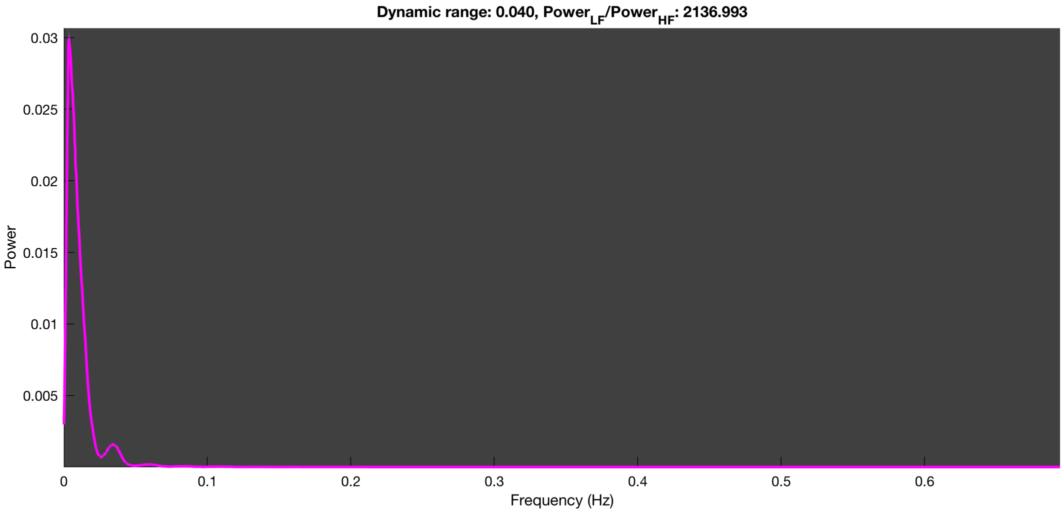

3-1 100n_tmap_component_ica_s1 022

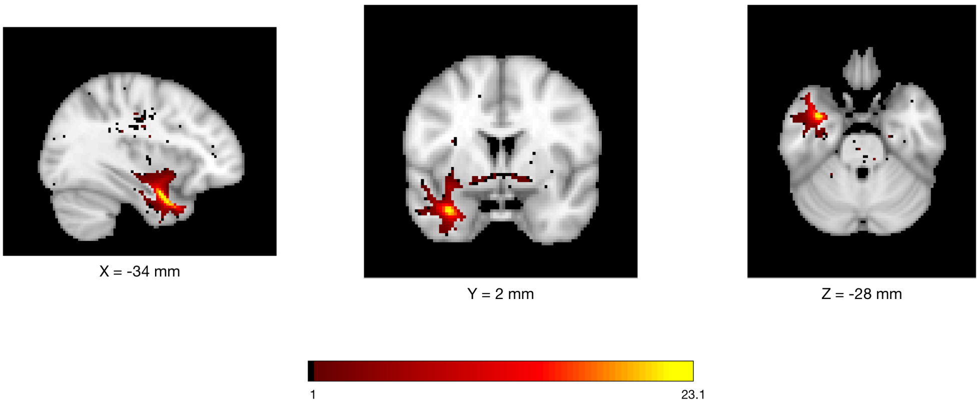


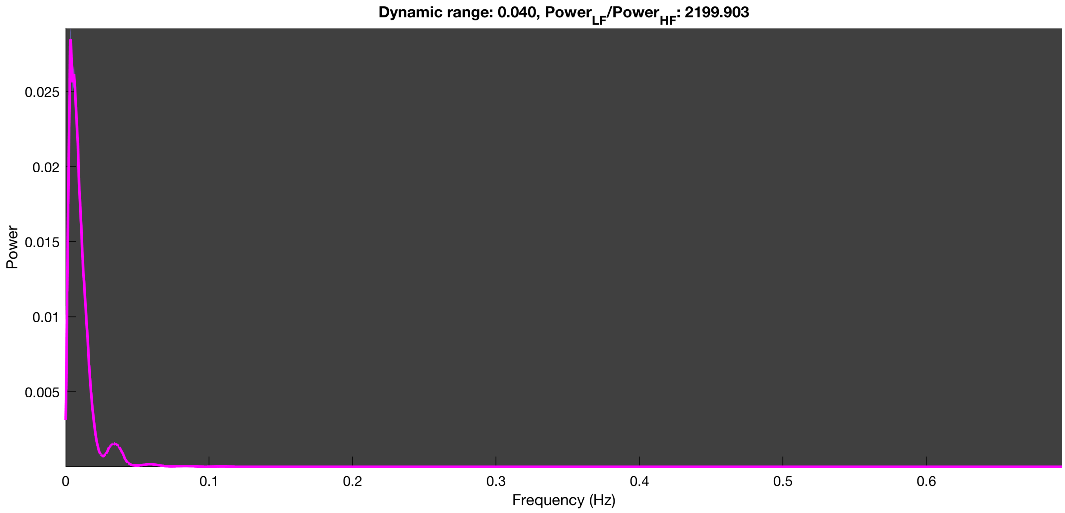

3-1 100n_tmap_component_ica_s1 023

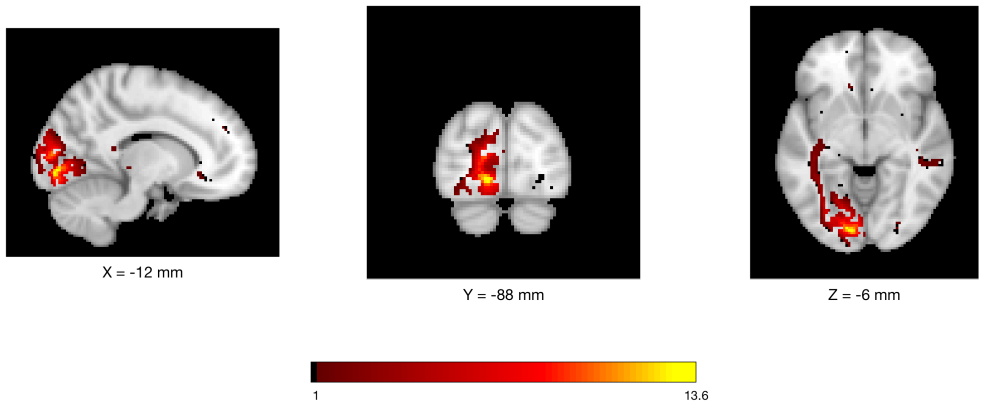


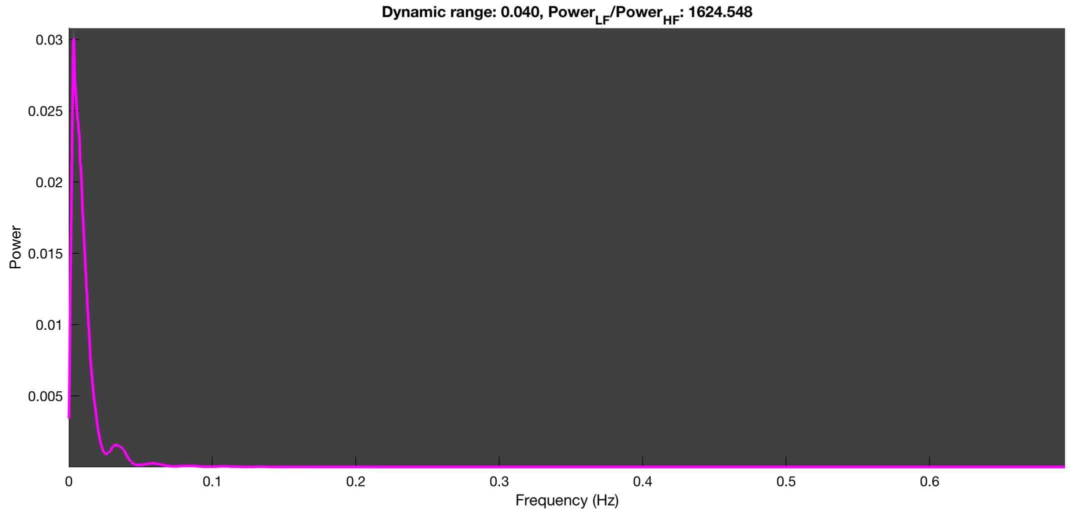

3-1 100n_tmap_component_ica_s1 024

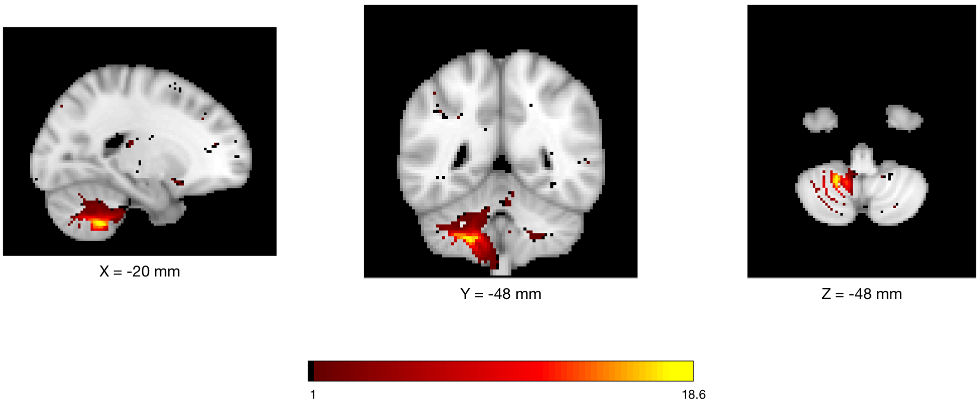


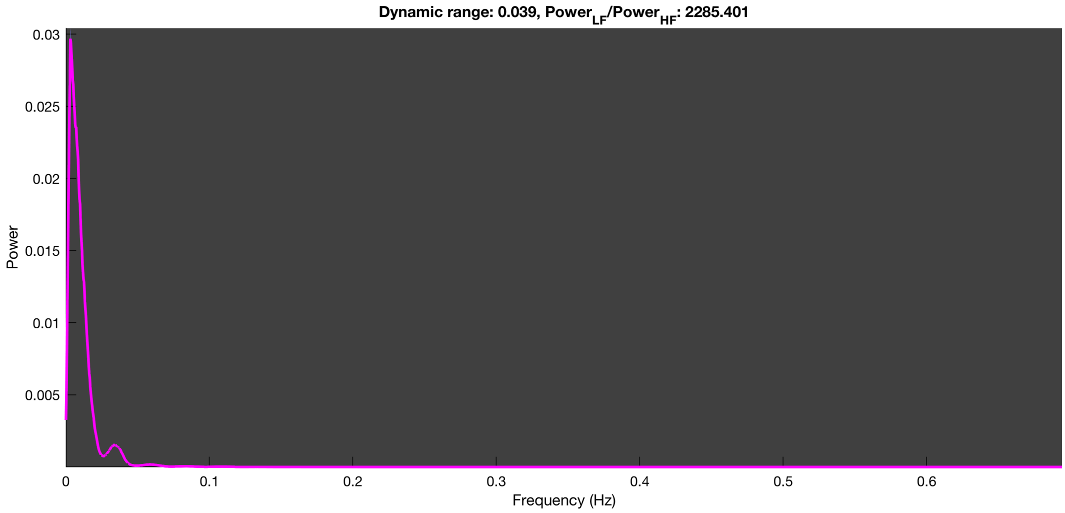

3-1 100n_tmap_component_ica_s1 025

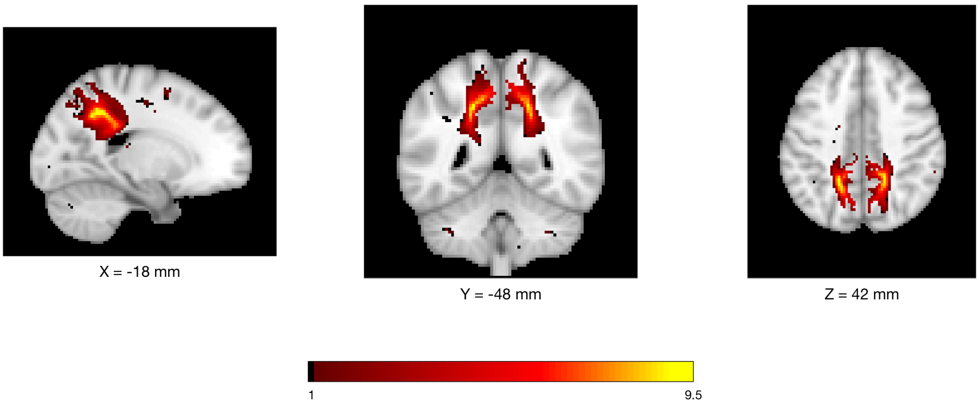


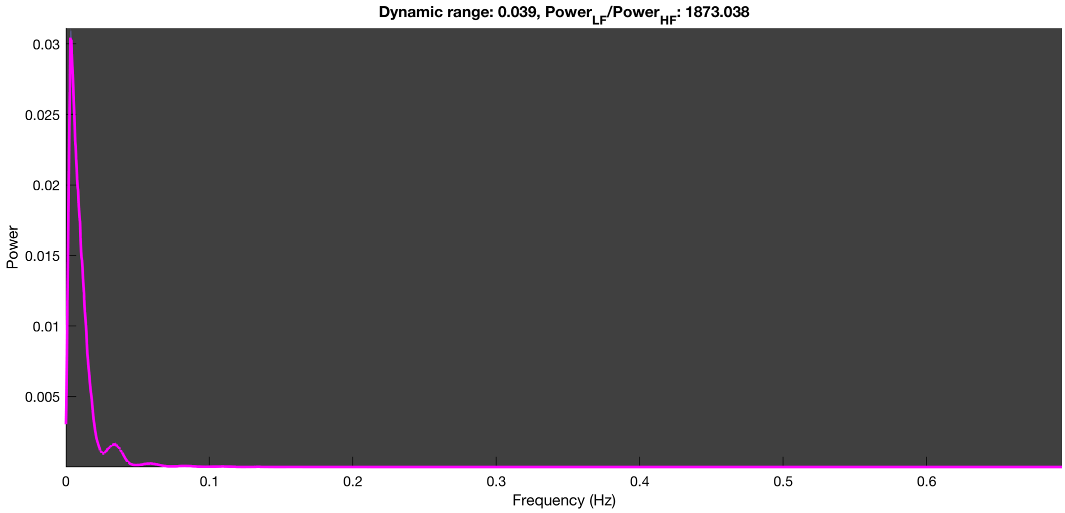

3-1 100n_tmap_component_ica_s1 026

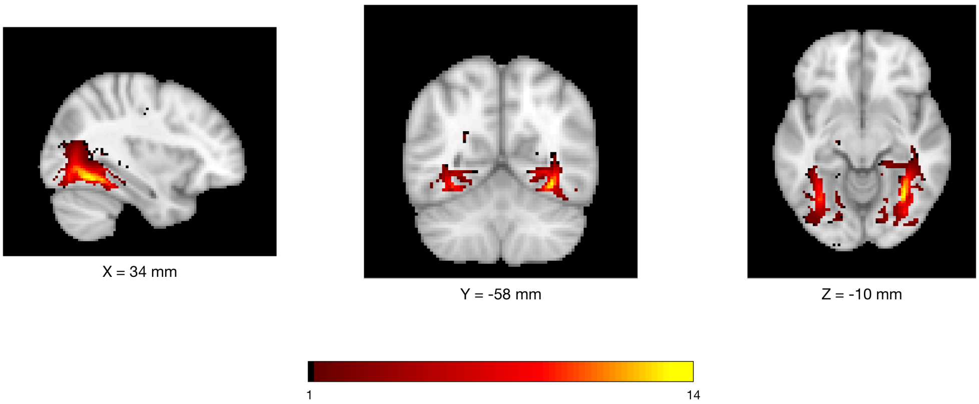


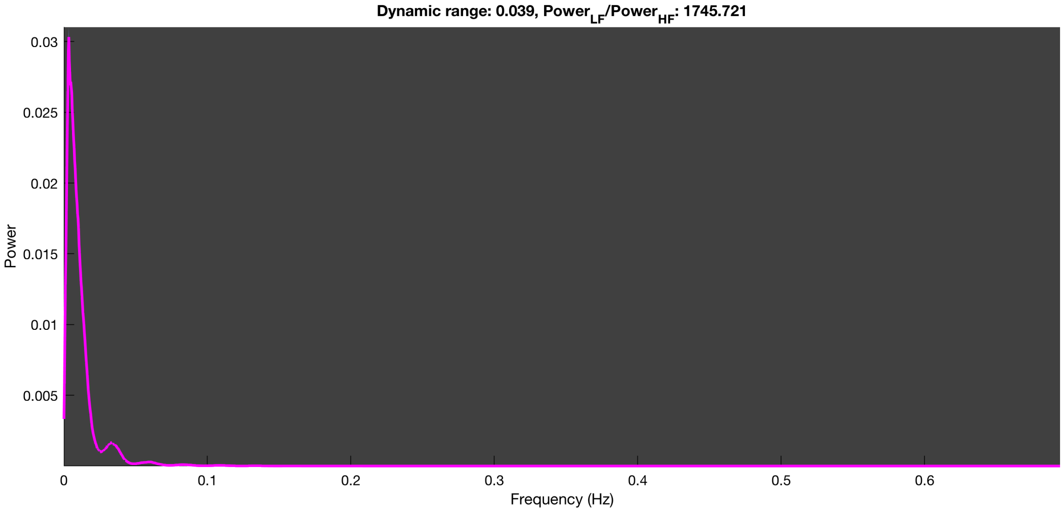

3-1 100n_tmap_component_ica_s1 027

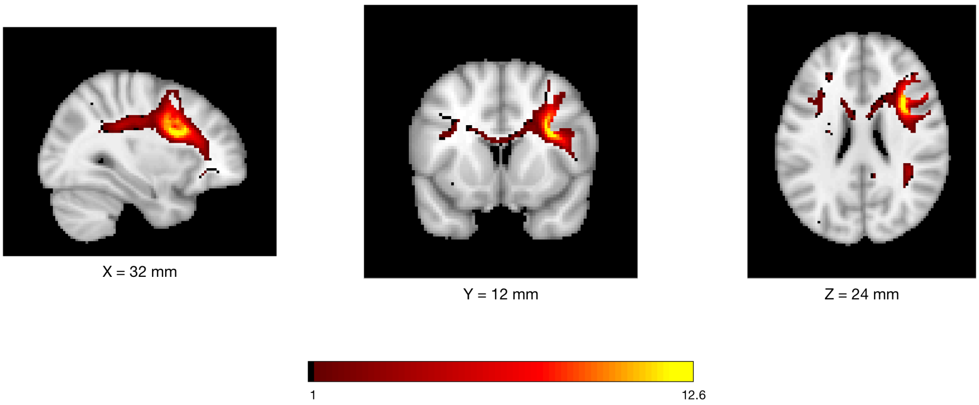


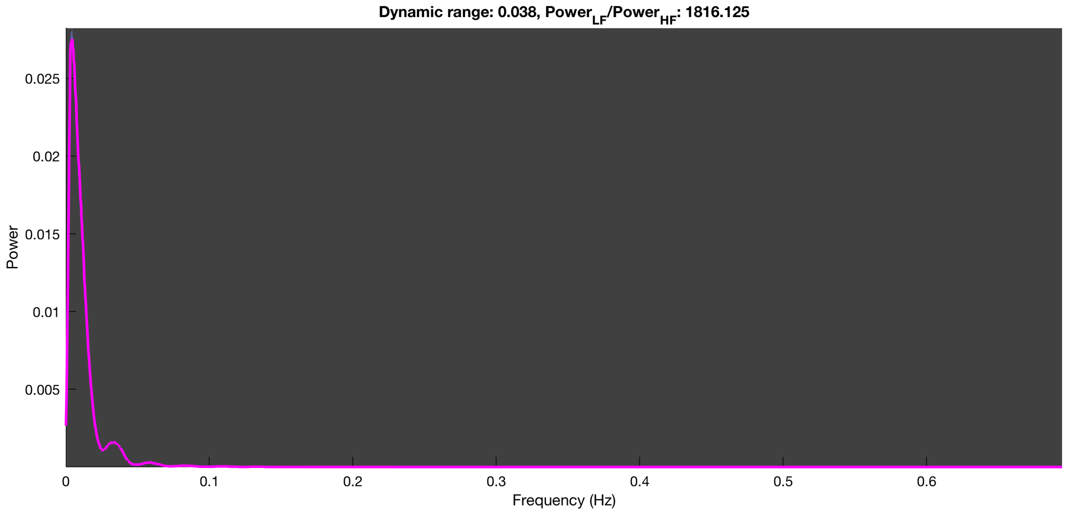

3-1 100n_tmap_component_ica_s1 028

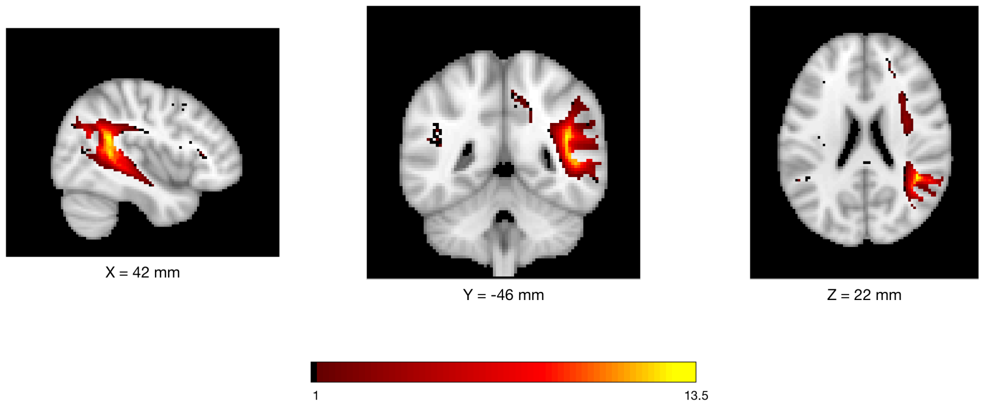


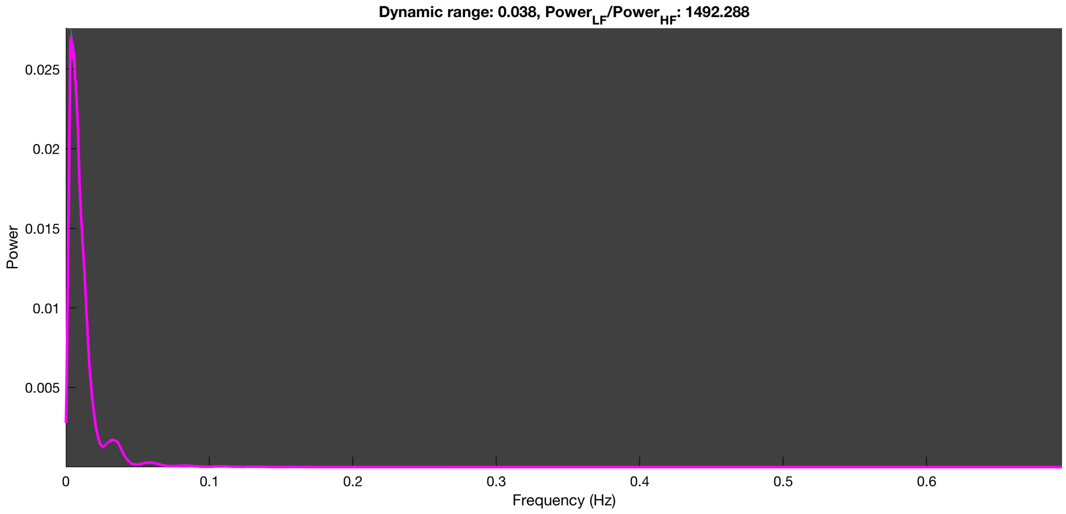

3-1 100n_tmap_component_ica_s1 029

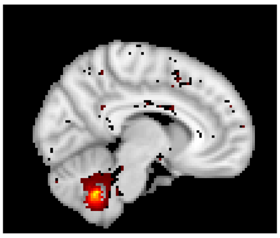

X = 8 mm

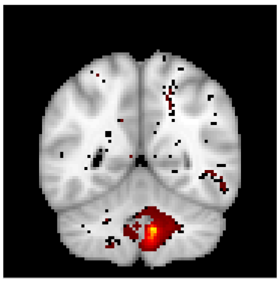

Y = -52 mm

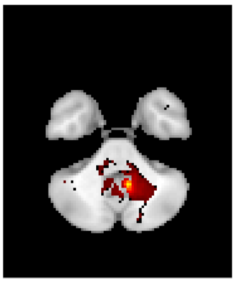

Z = -40 mm

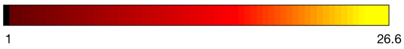


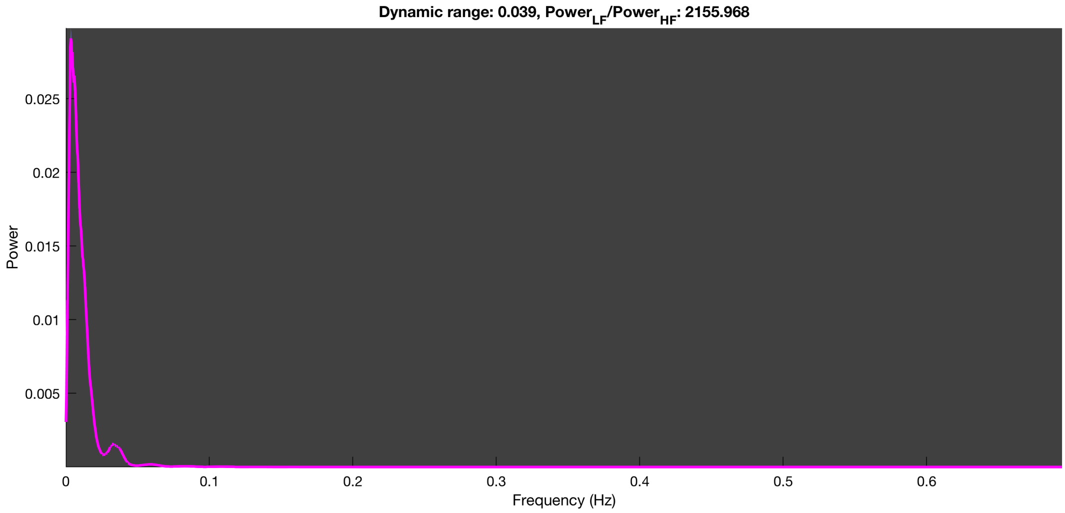

3-1 100n_tmap_component_ica_s1 030

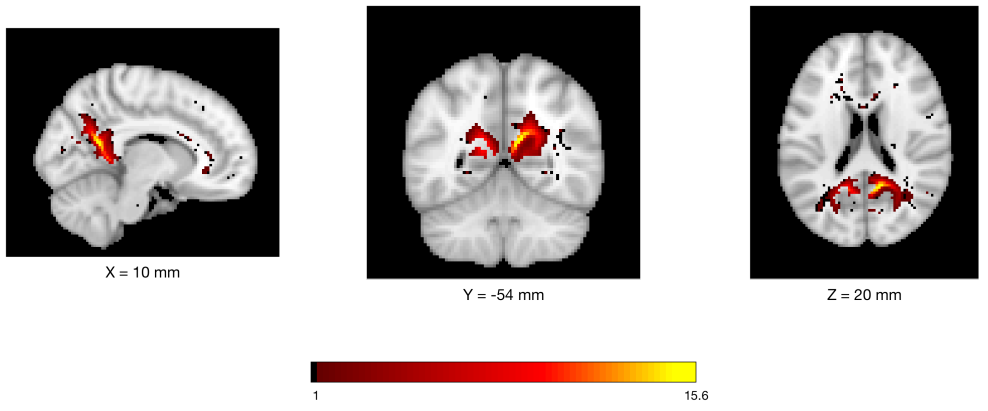


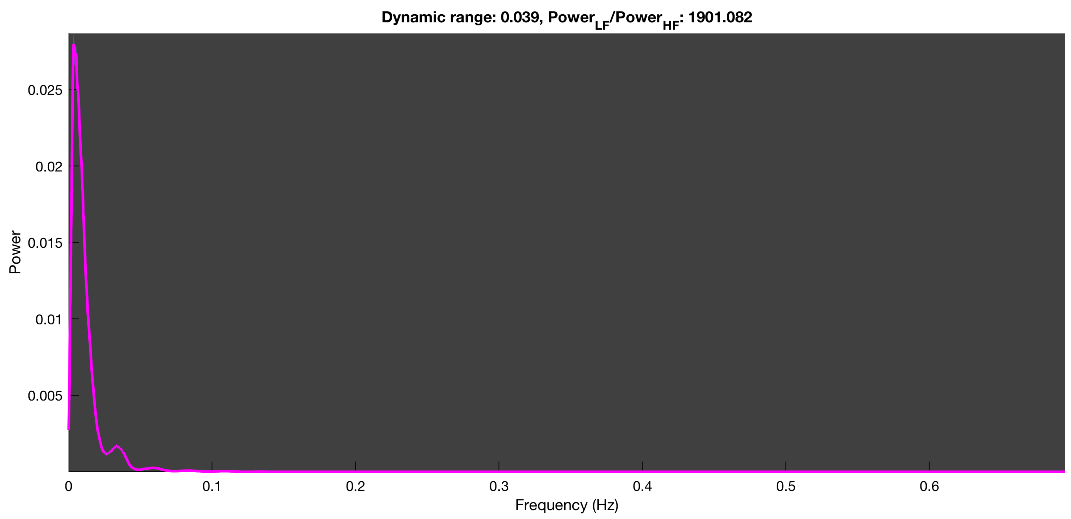

3-1 100n_tmap_component_ica_s1 031

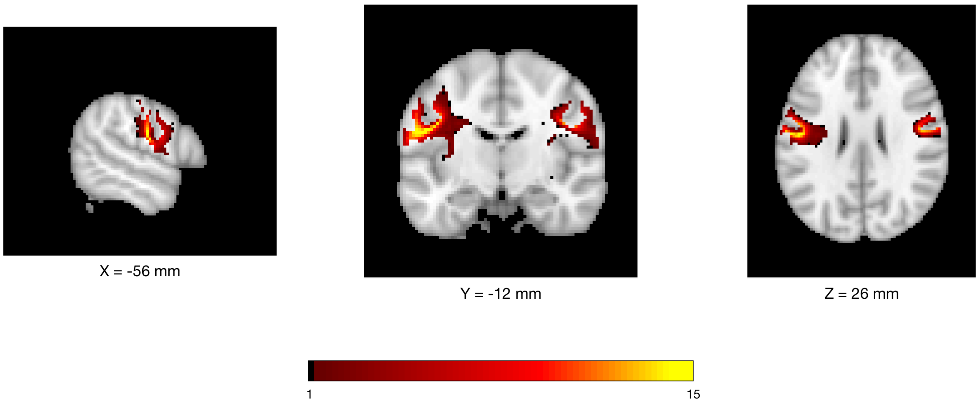


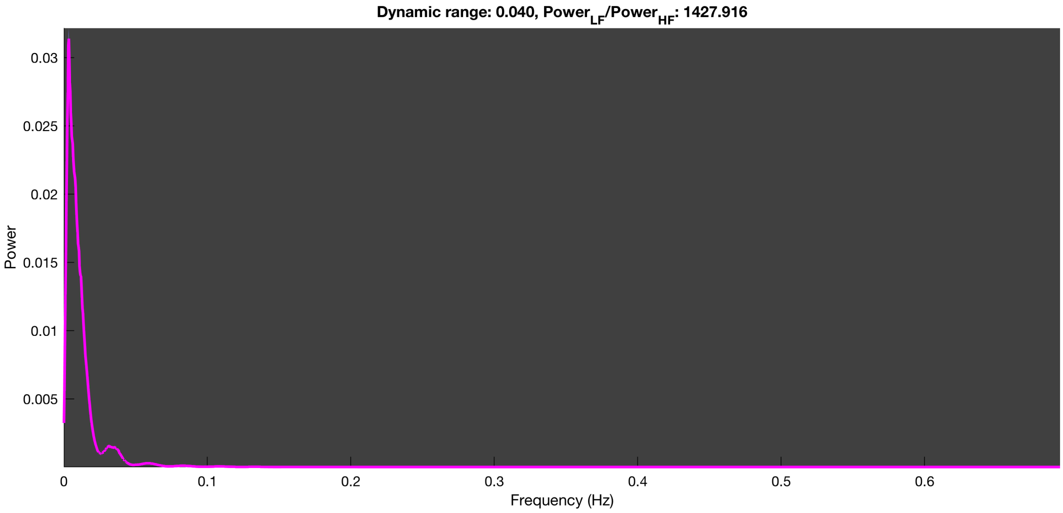

3-1 100n_tmap_component_ica_s1 032

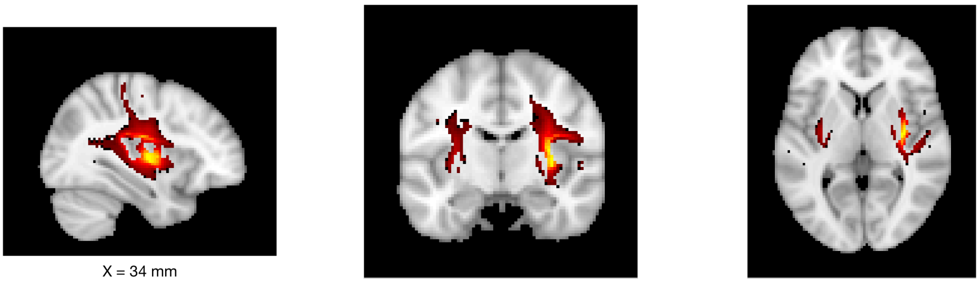

X = 34 mm    Y = -10 mm    Z = 4 mm

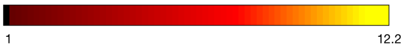


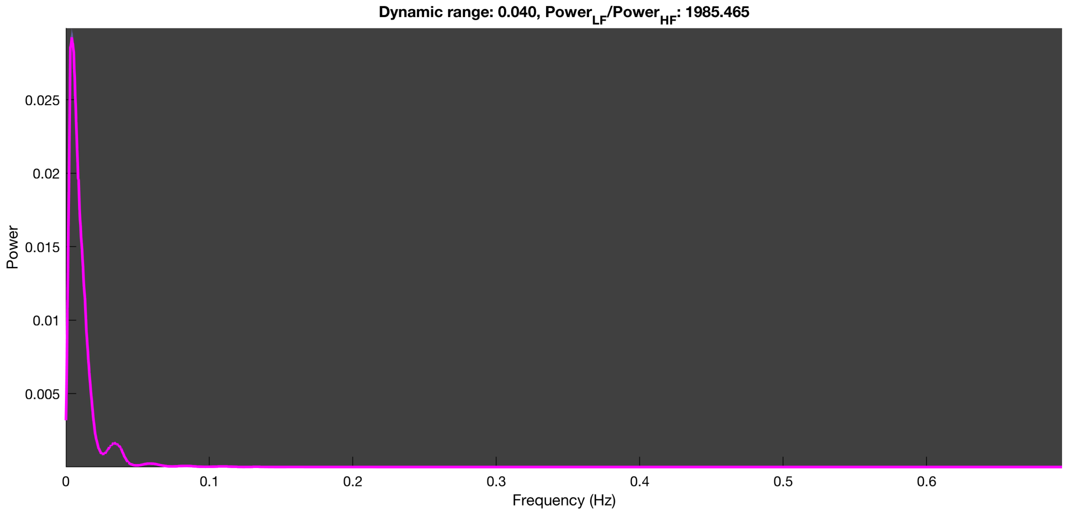

3-1 100n_tmap_component_ica_s1 033

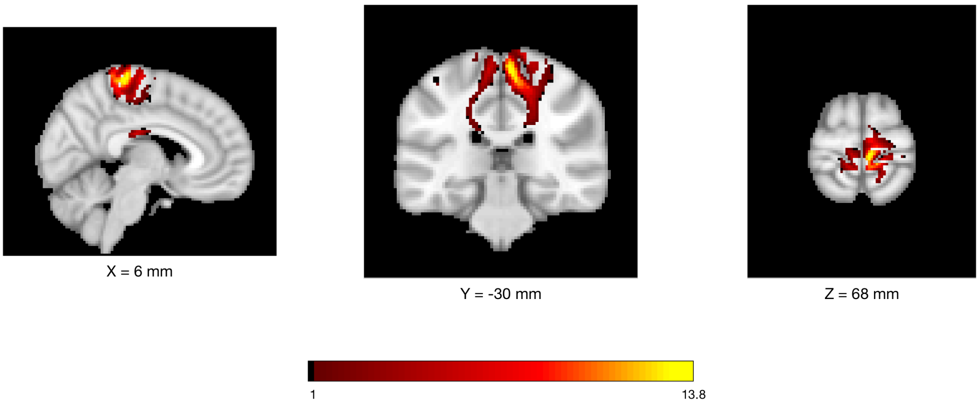


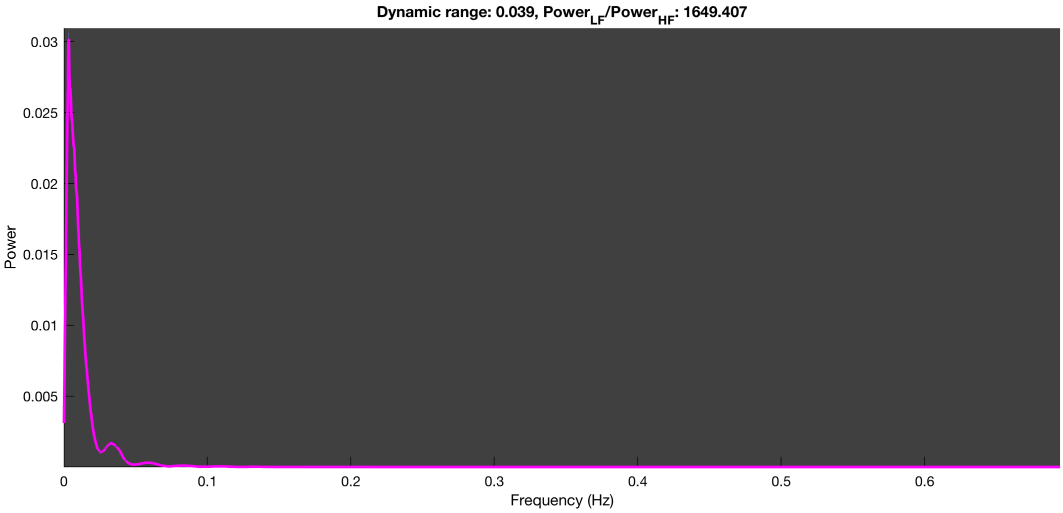

3-1 100n_tmap_component_ica_s1 034

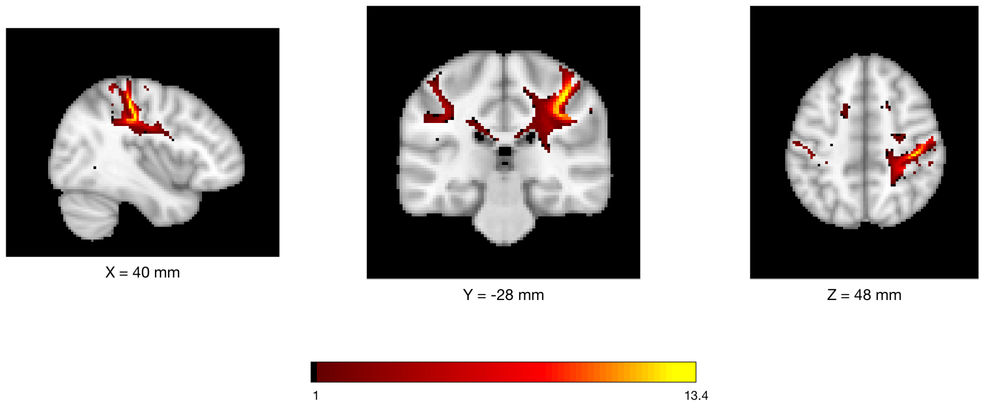


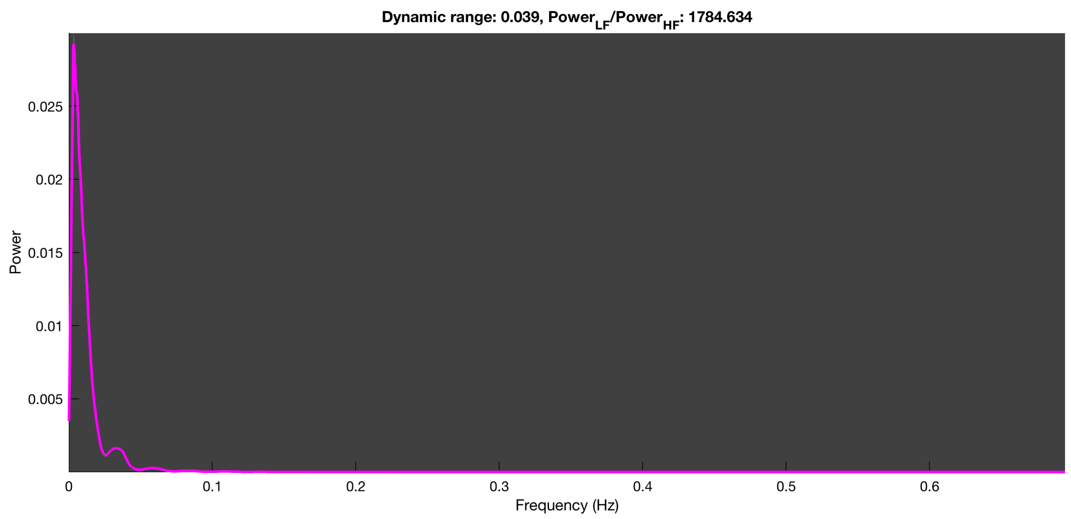

3-1 100n_tmap_component_ica_s1 035

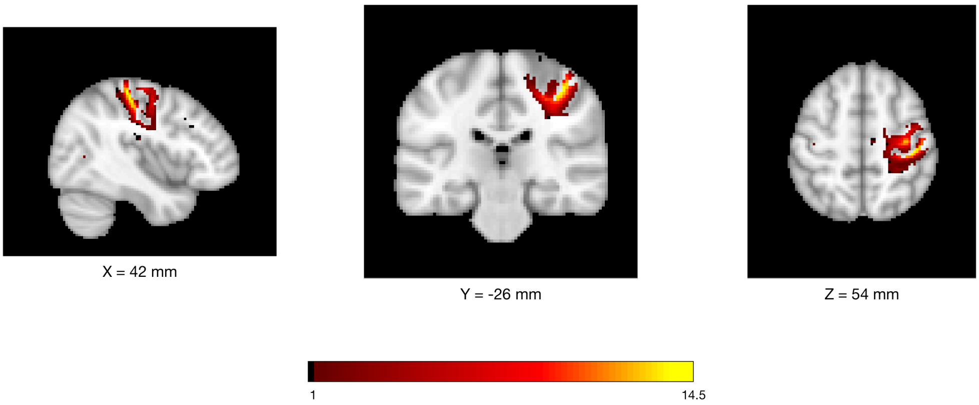


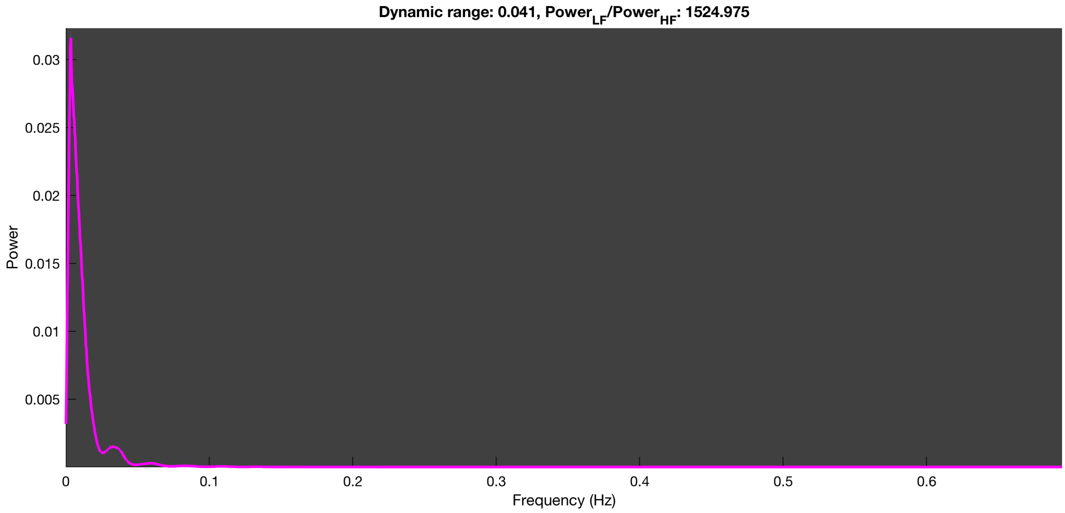

3-1 100n_tmap_component_ica_s1 036

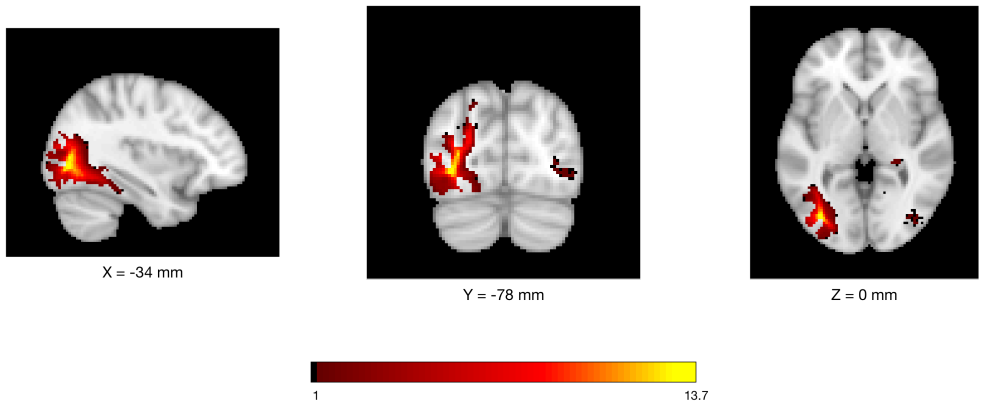


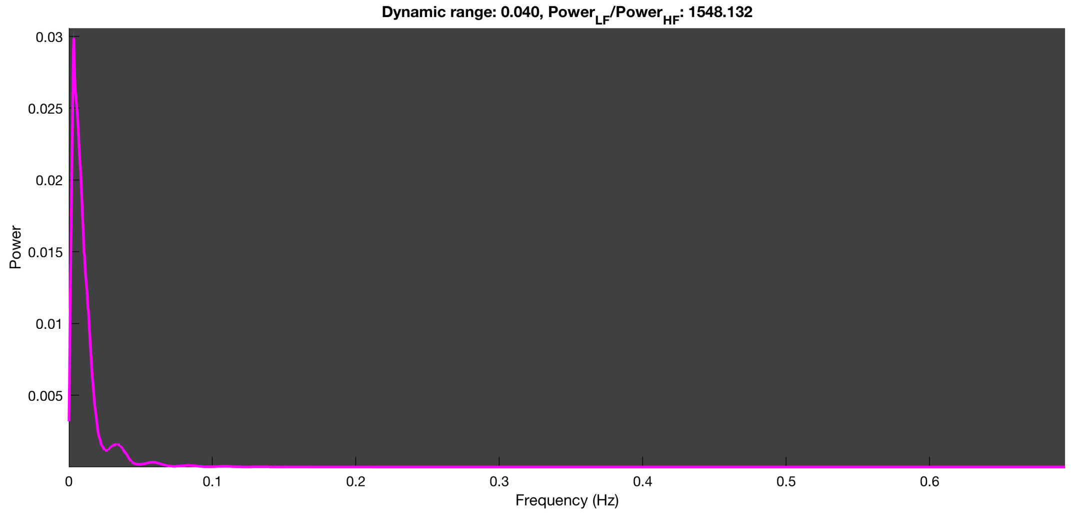

3-1 100n_tmap_component_ica_s1 037

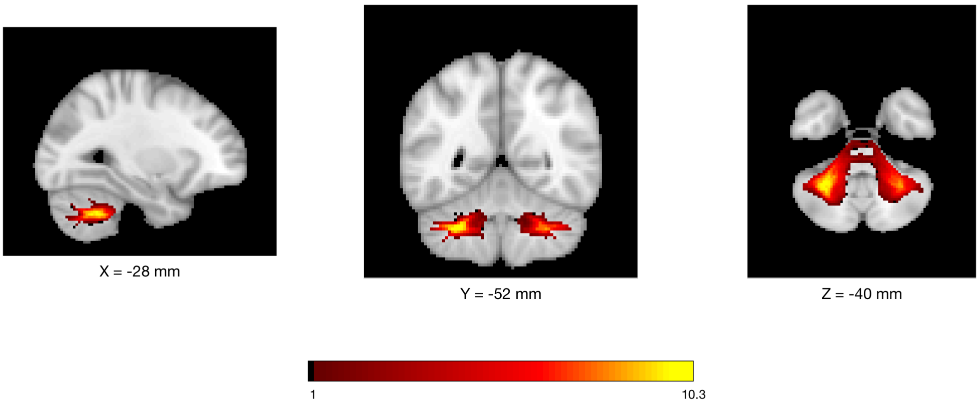


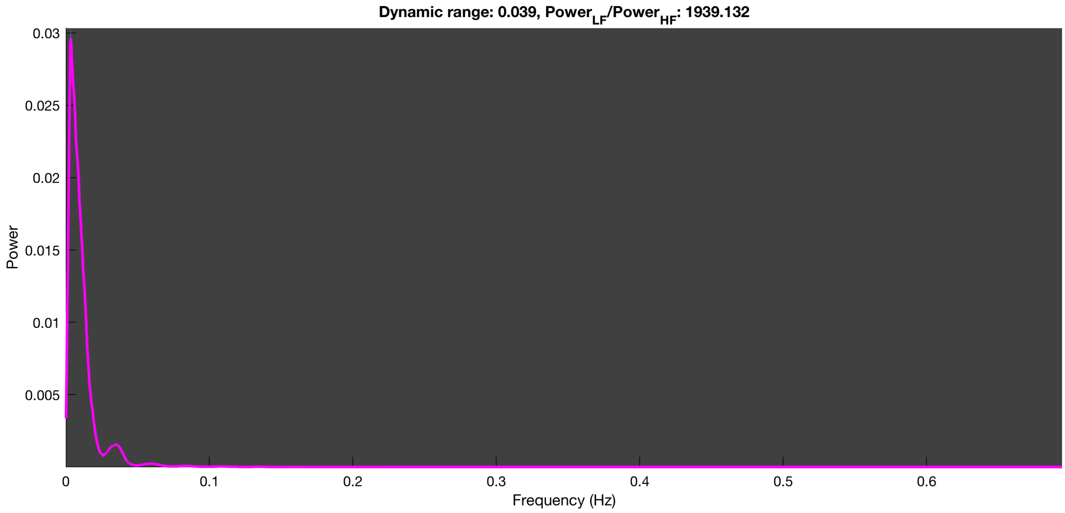

3-1 100n_tmap_component_ica_s1 038

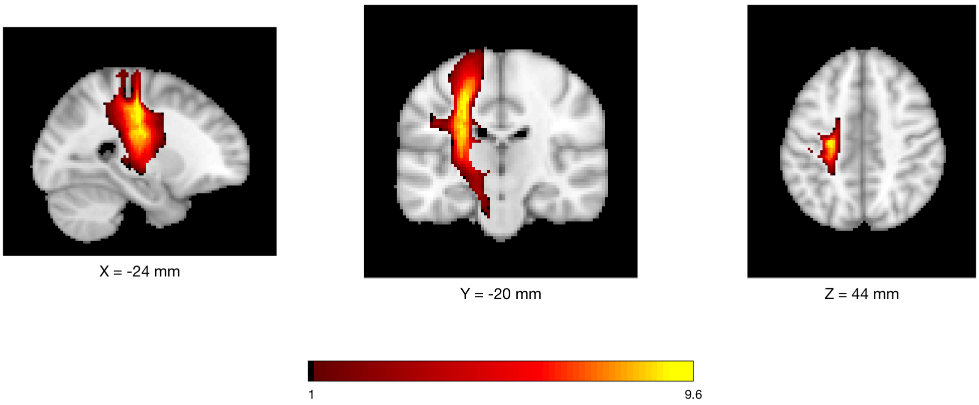


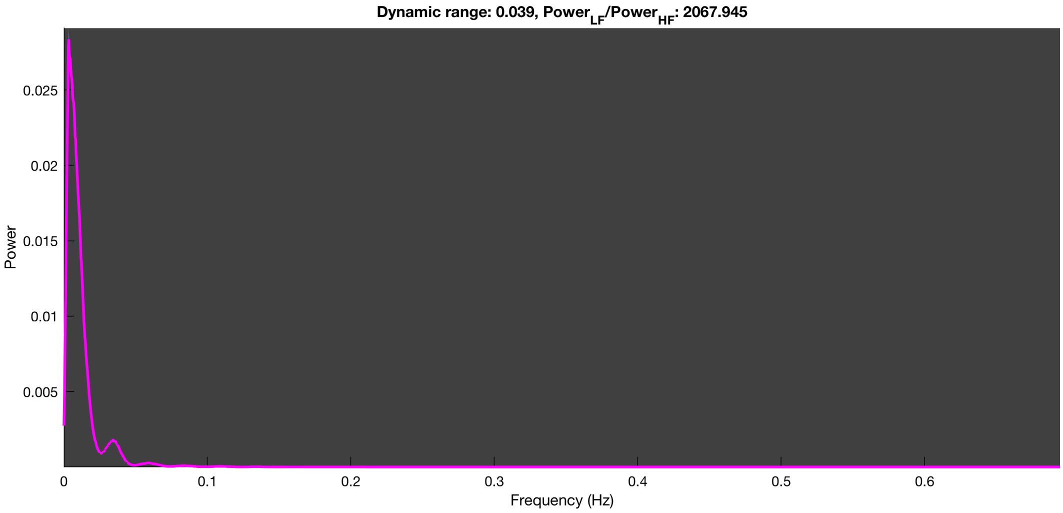

3-1 100n_tmap_component_ica_s1 039

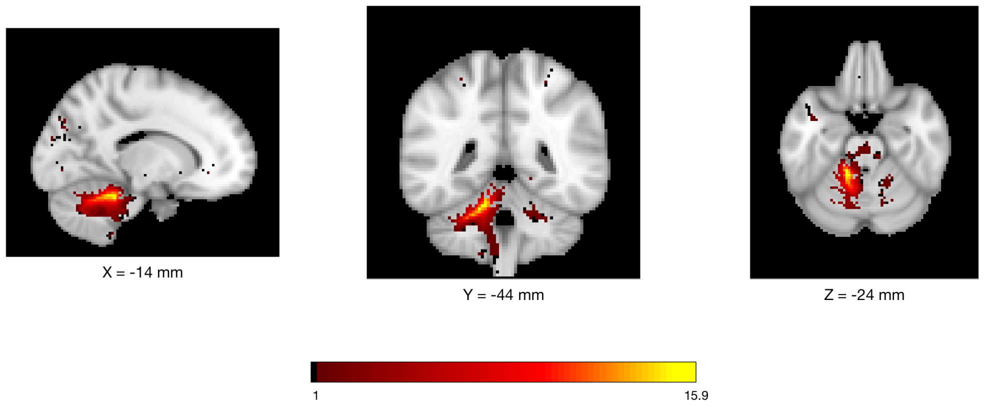


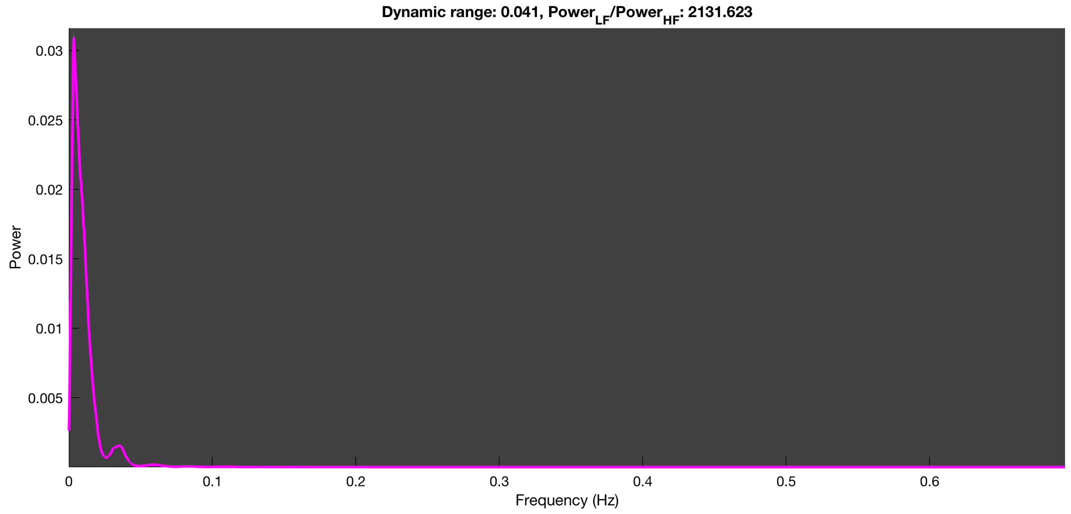

3-1 100n_tmap_component_ica_s1 040

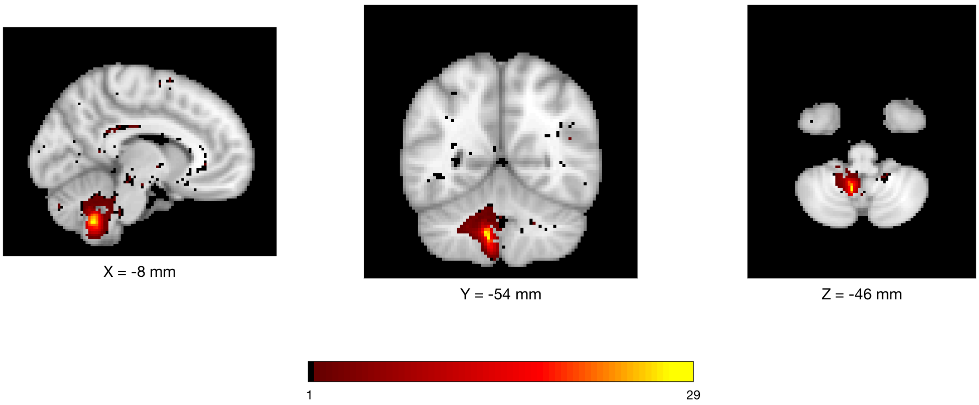


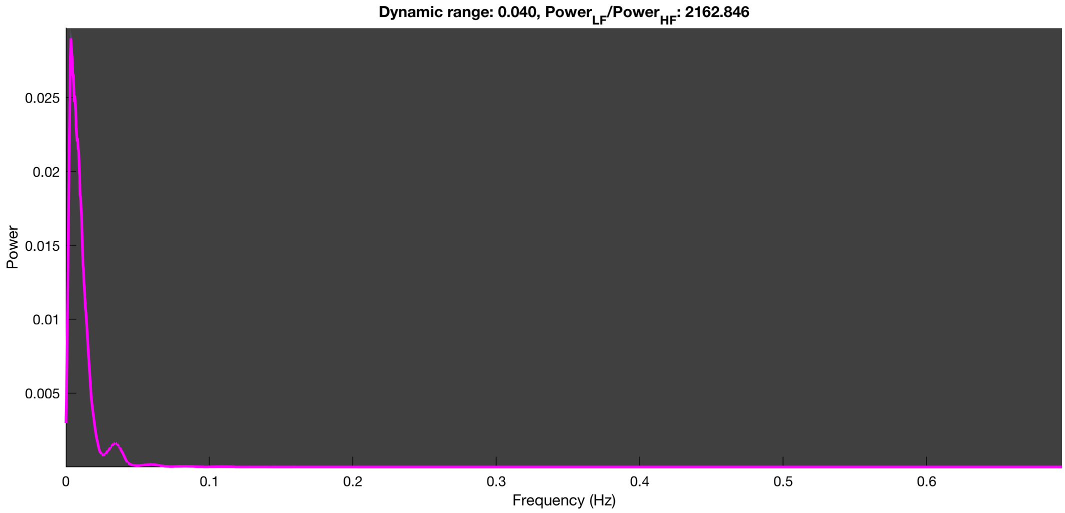

##### 3-1 100n_tmap_component_ica_s1 041

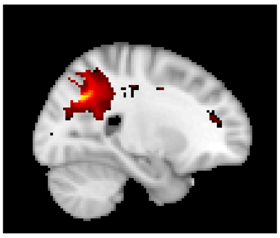

X = 24 mm

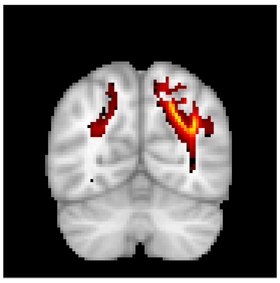

Y = -64 mm

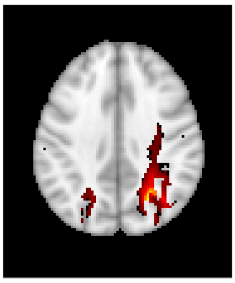

Z = 36 mm

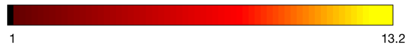


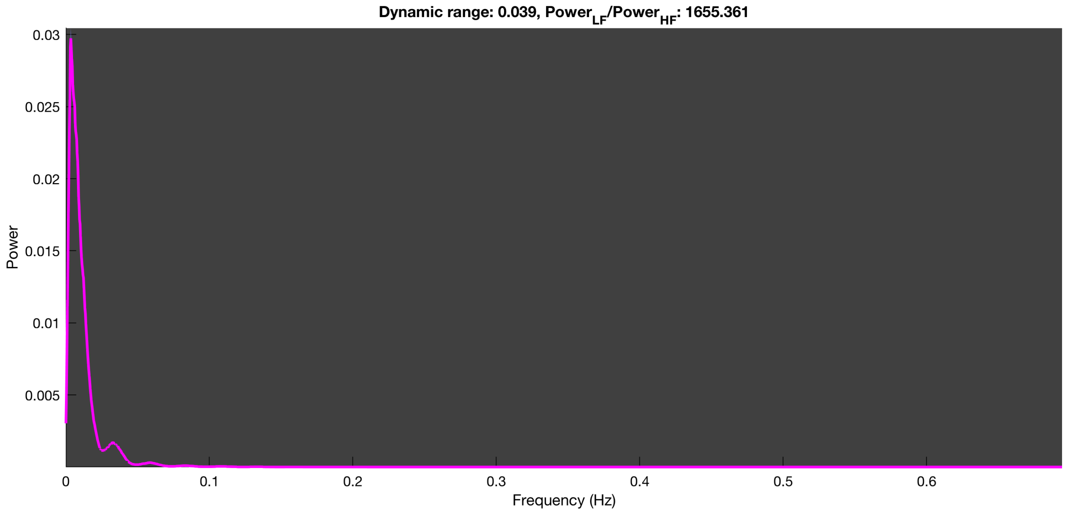

3-1 100n_tmap_component_ica_s1 042

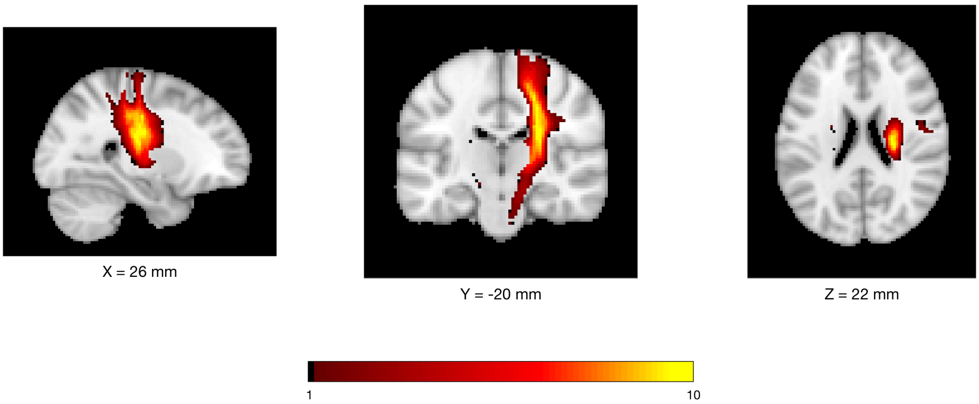


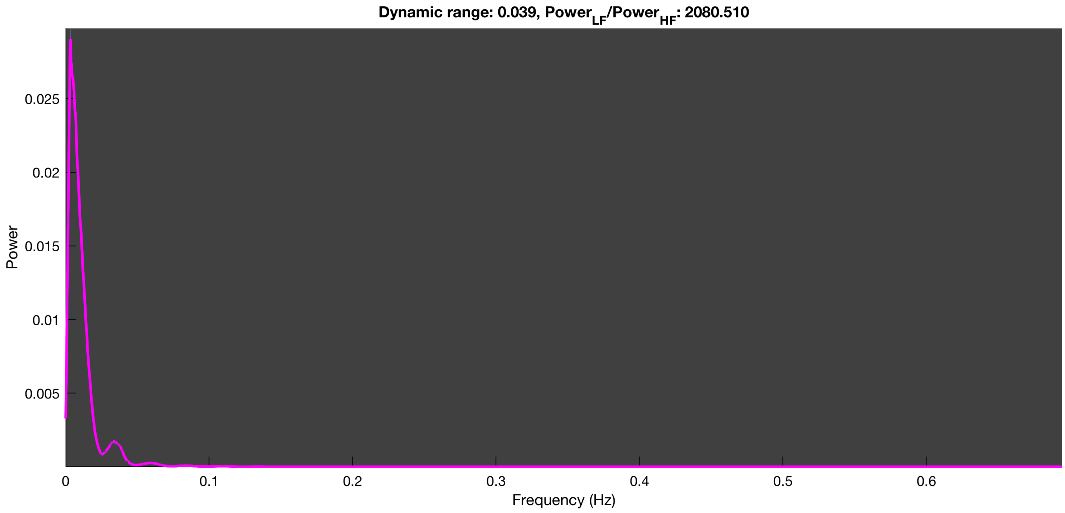

3-1 100n_tmap_component_ica_s1 043

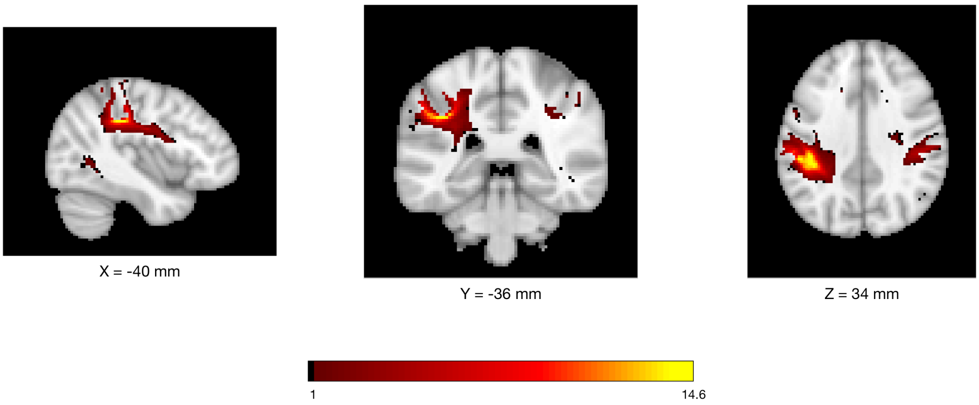


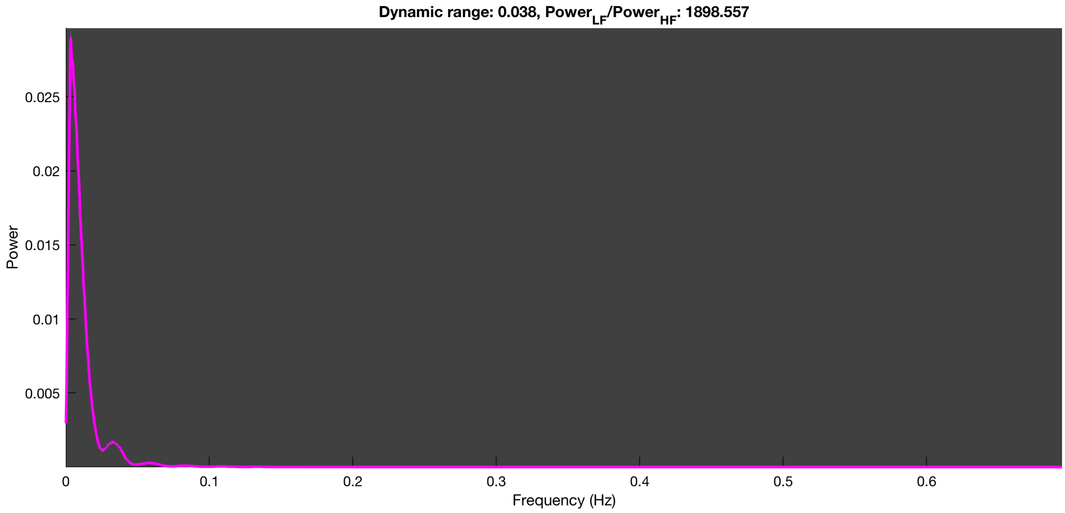

3-1 100n_tmap_component_ica_s1 044

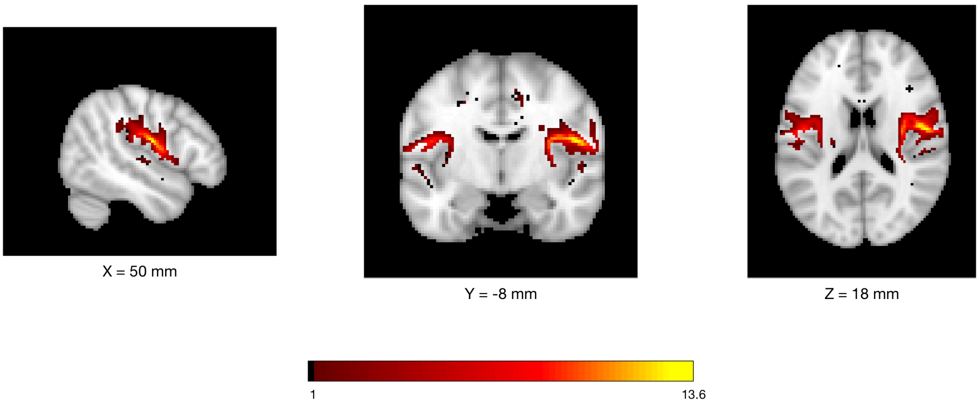


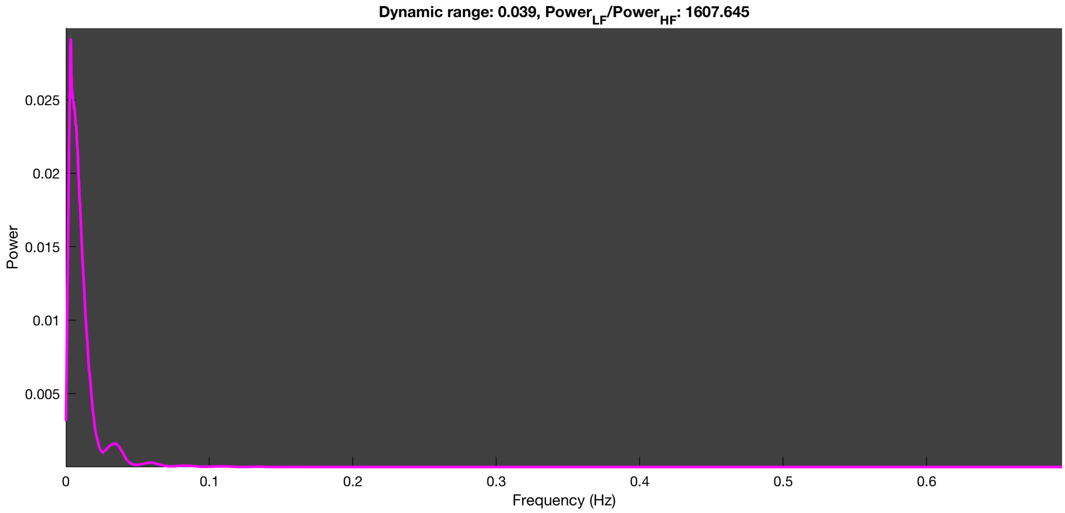

#### 3-1 100n_tmap_component_ica_s1 045

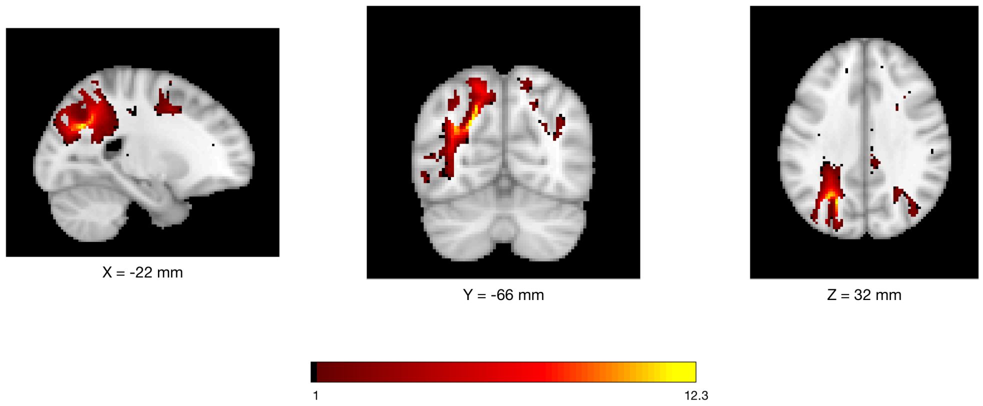


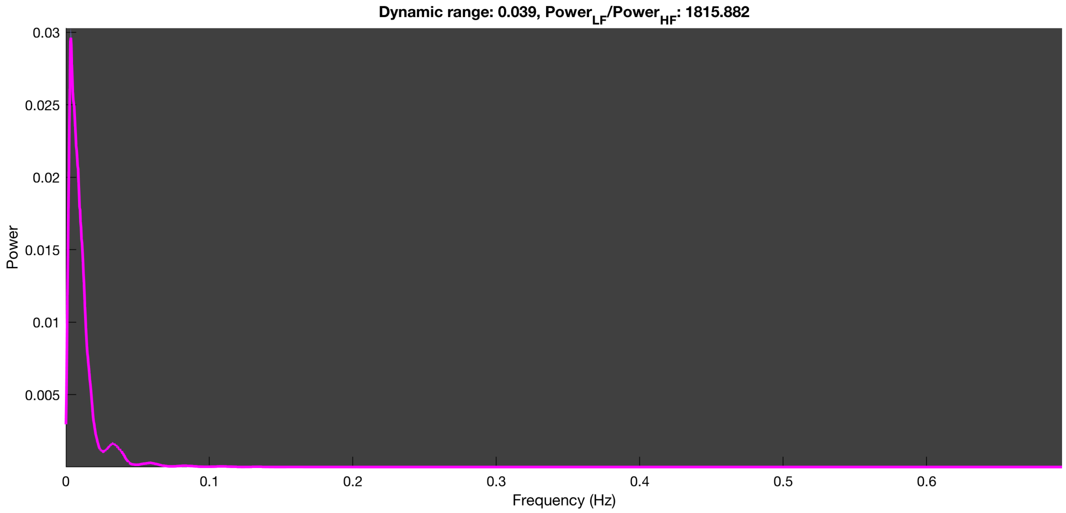

3-1 100n_tmap_component_ica_s1 046

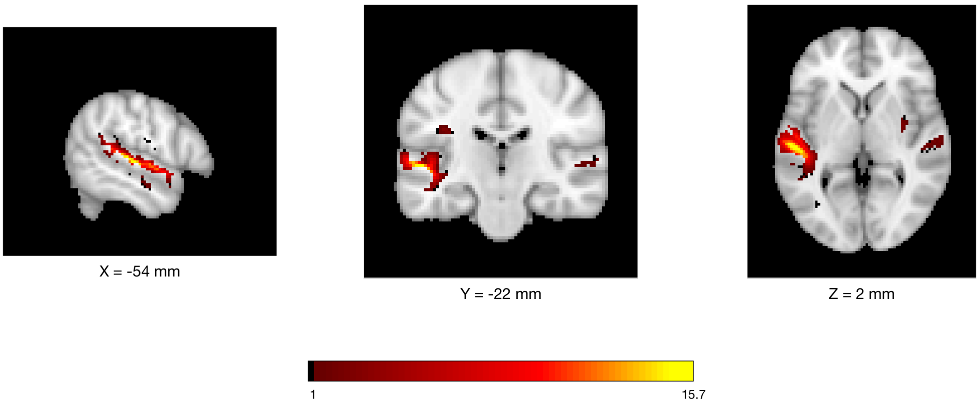


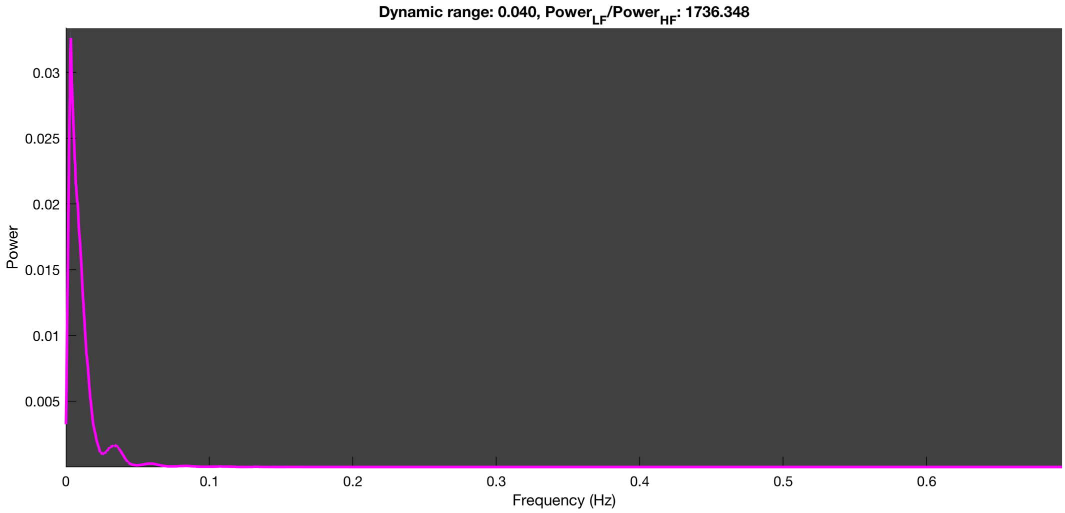

3-1 100n_tmap_component_ica_s1 047

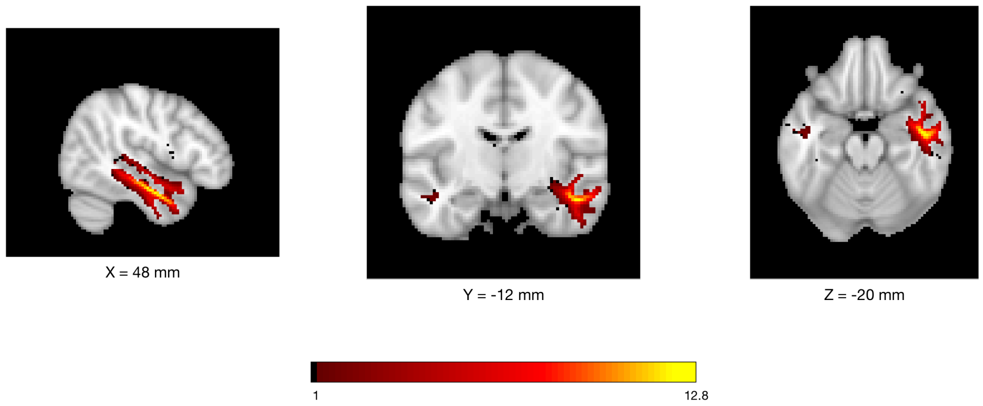


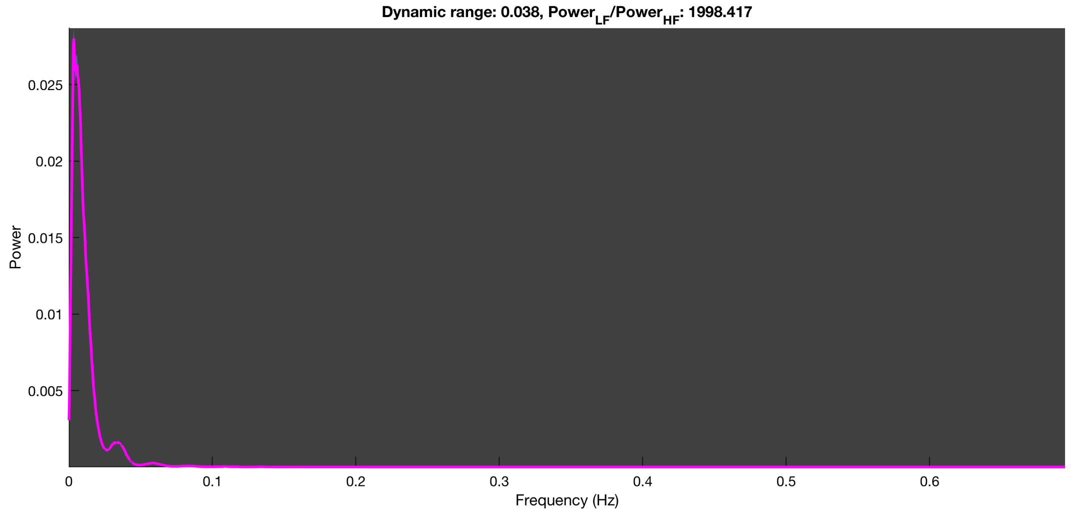

3-1 100n_tmap_component_ica_s1 048

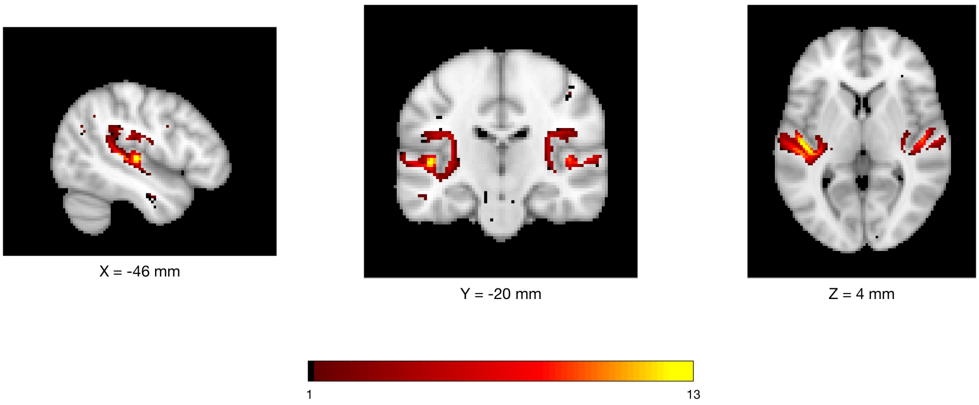


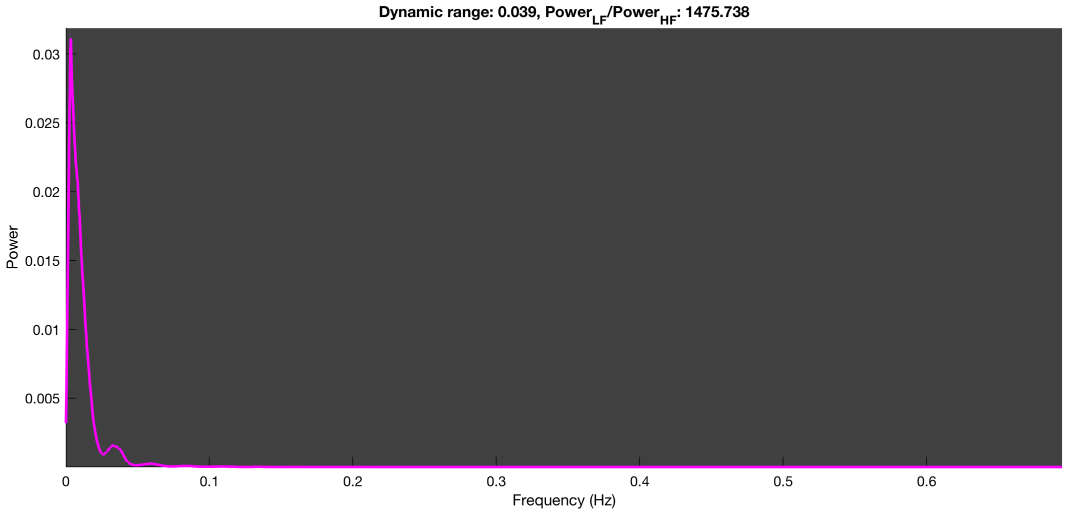

3-1 100n_tmap_component_ica_s1 049

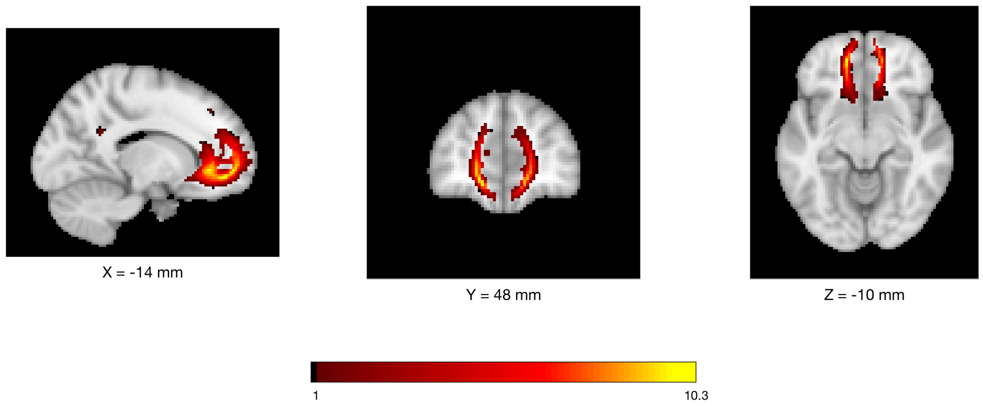


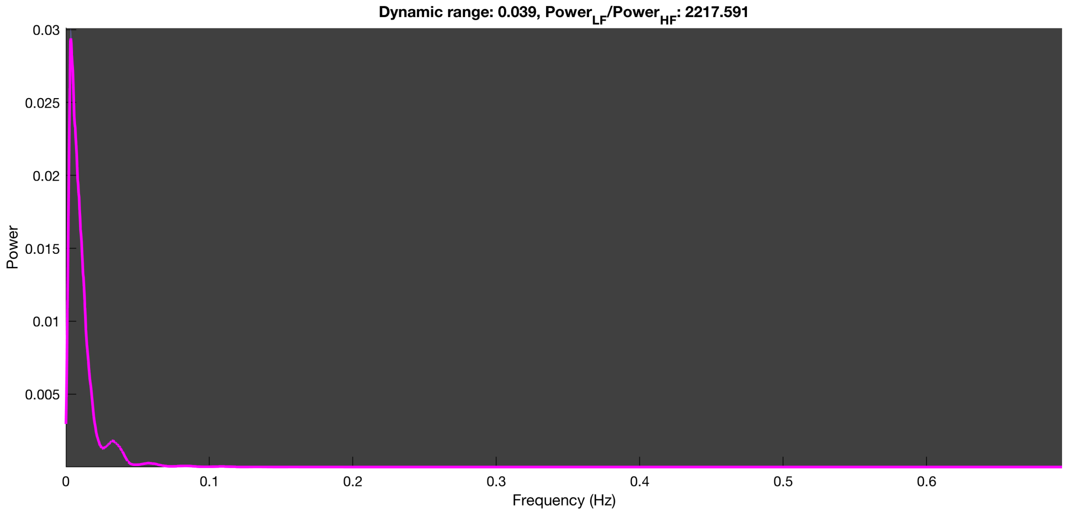

3-1 100n_tmap_component_ica_s1 050

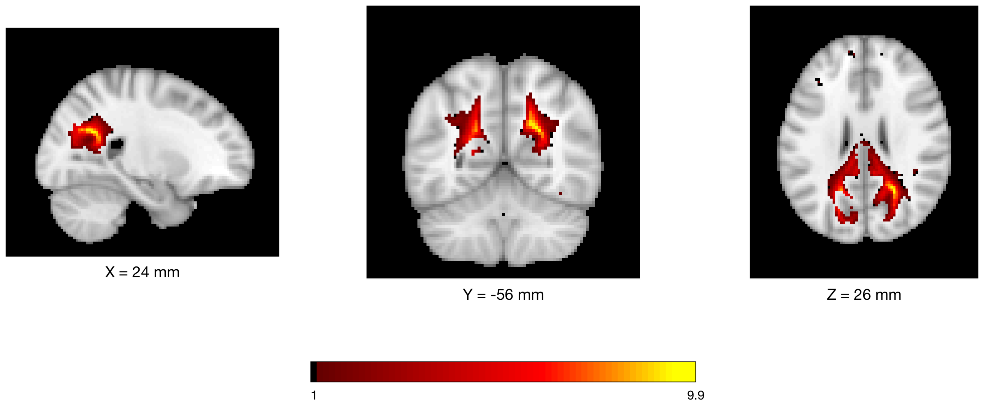


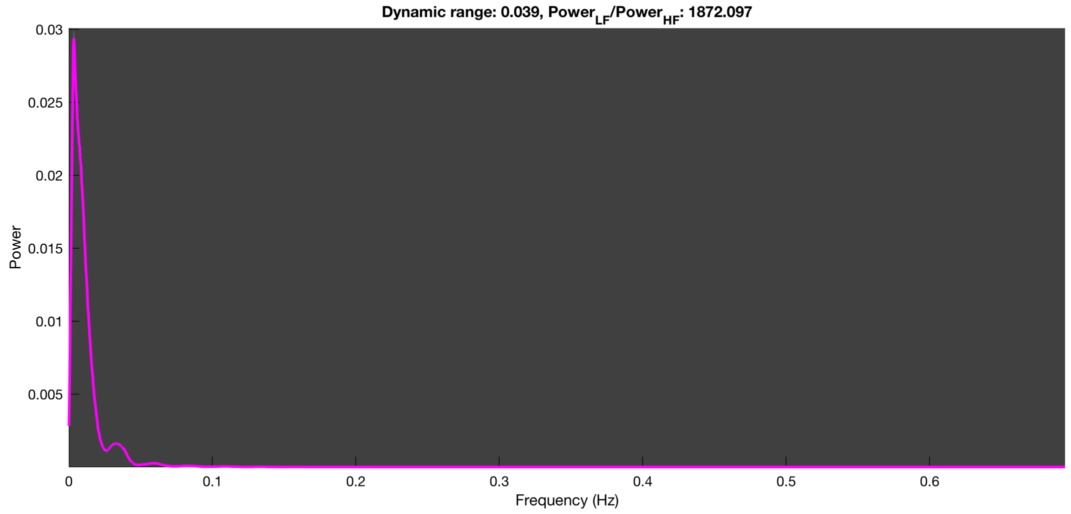

3-1 100n_tmap_component_ica_s1 051

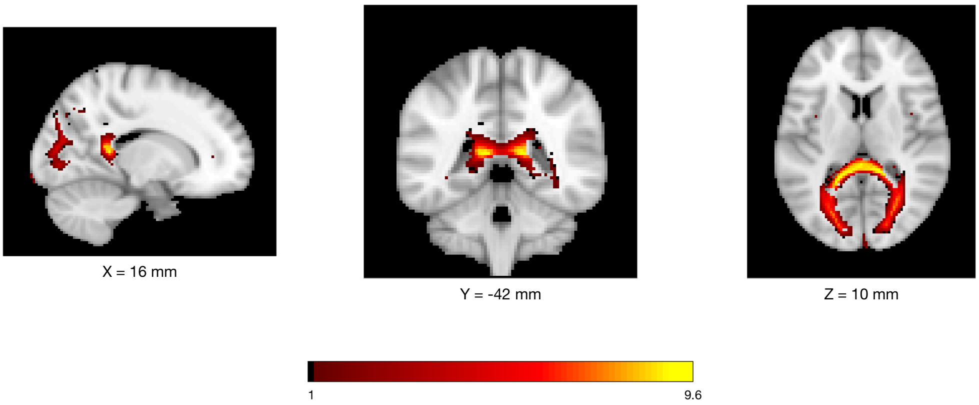


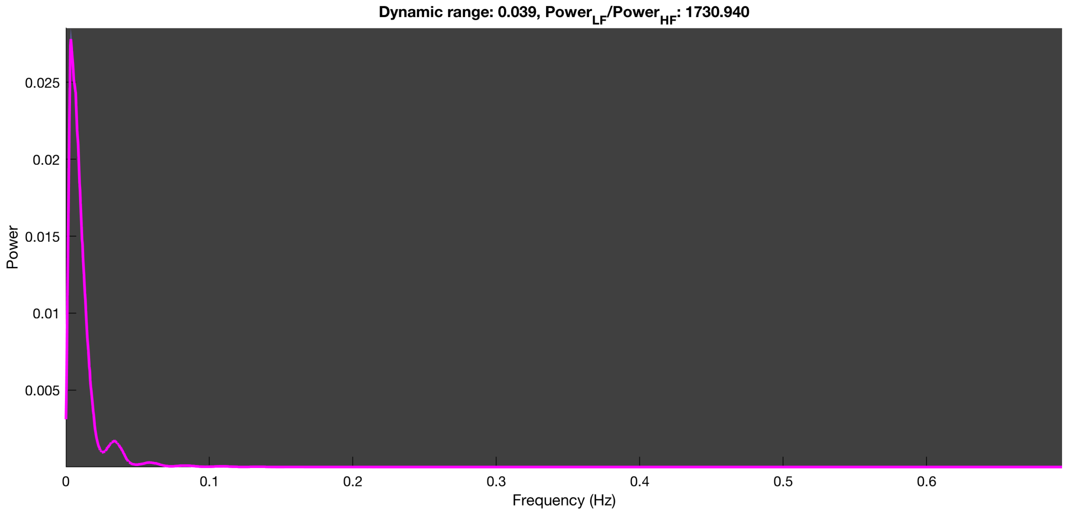

3-1 100n_tmap_component_ica_s1 052

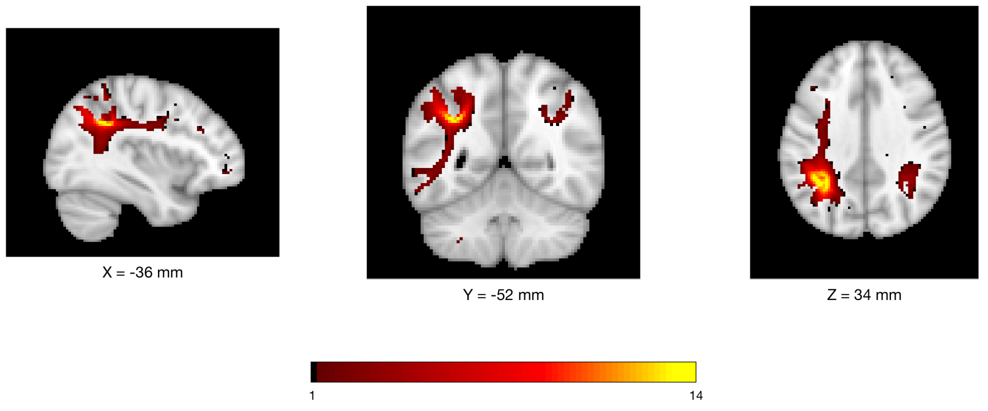


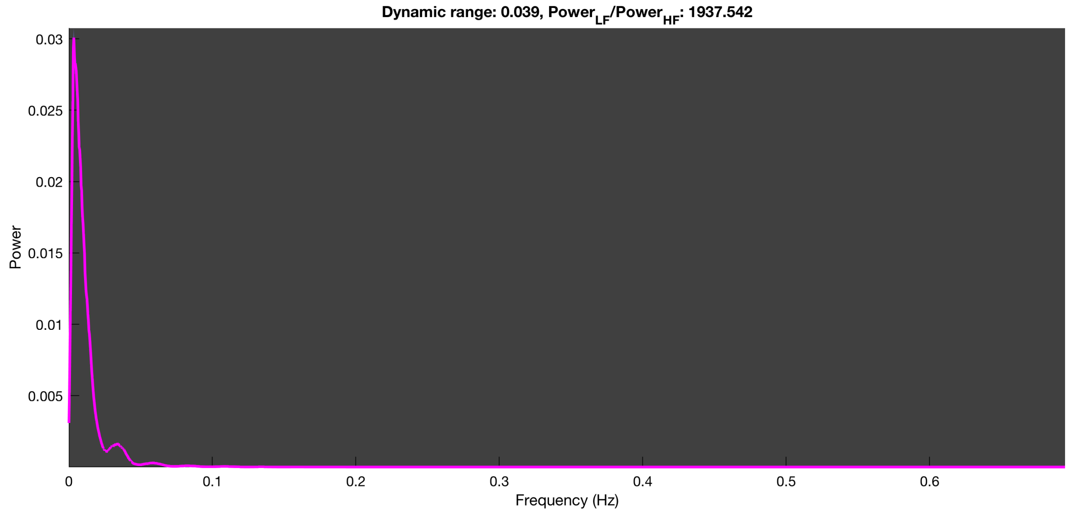

#### 3-1 100n_tmap_component_ica_s1 053

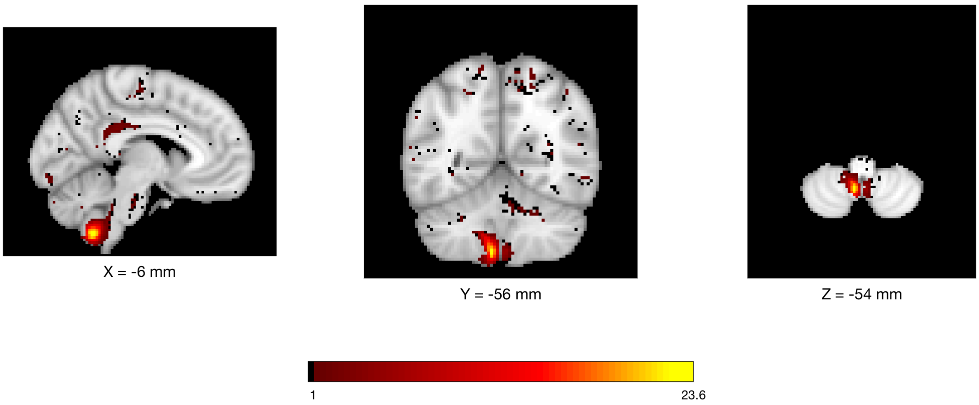


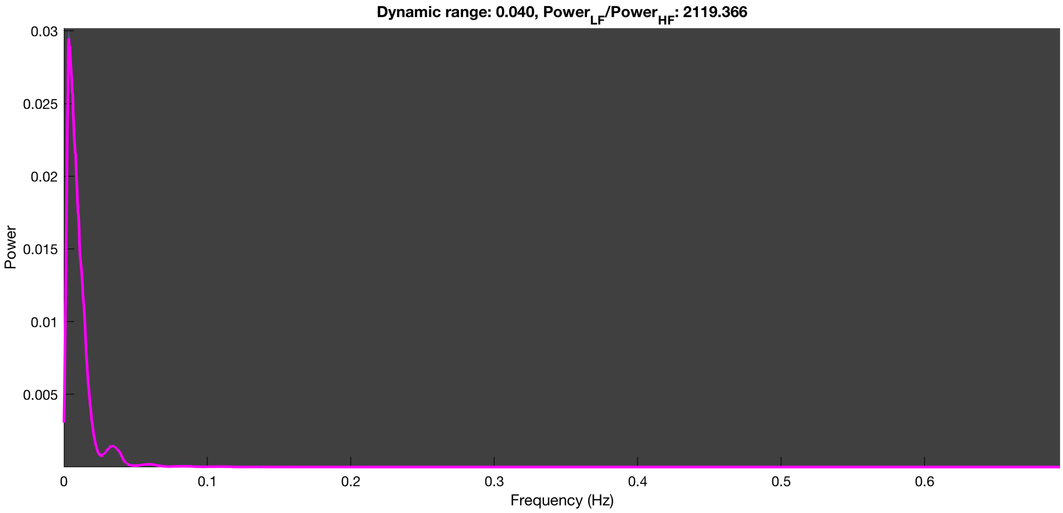

3-1 100n_tmap_component_ica_s1 054

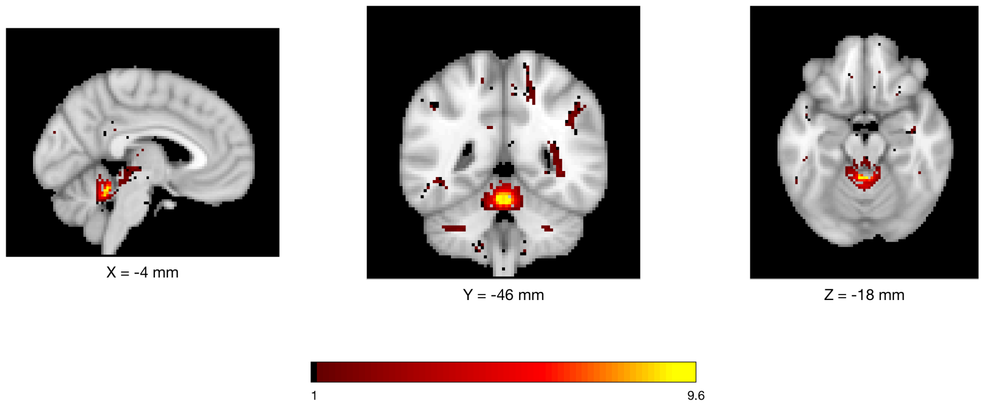


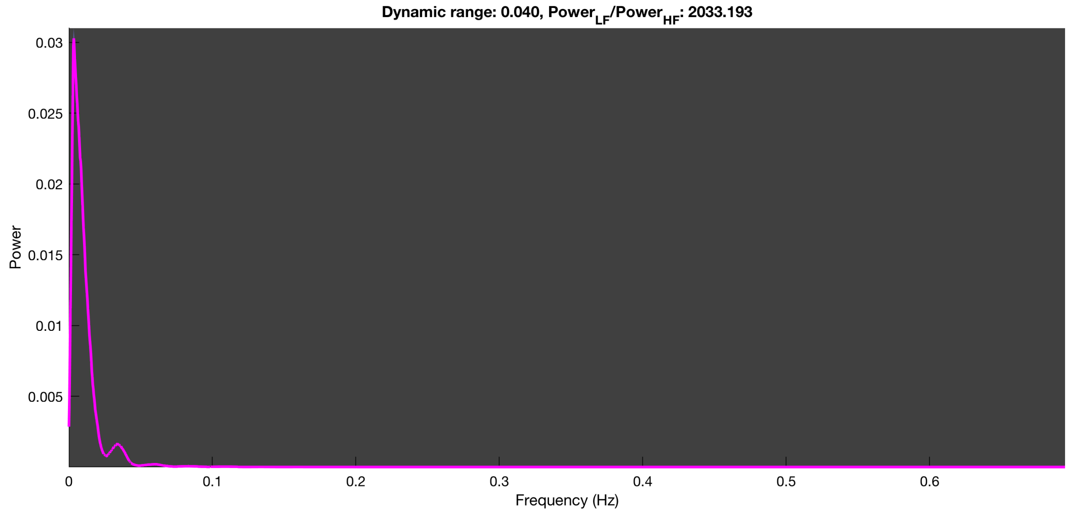

3-1 100n_tmap_component_ica_s1 055

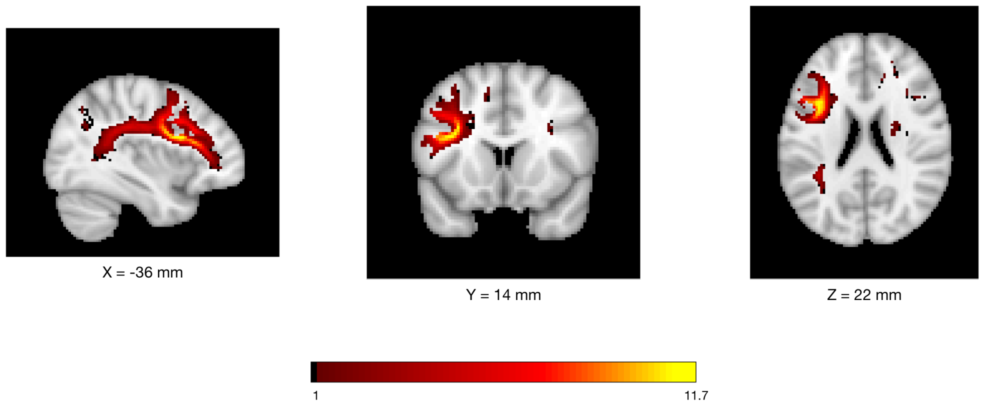


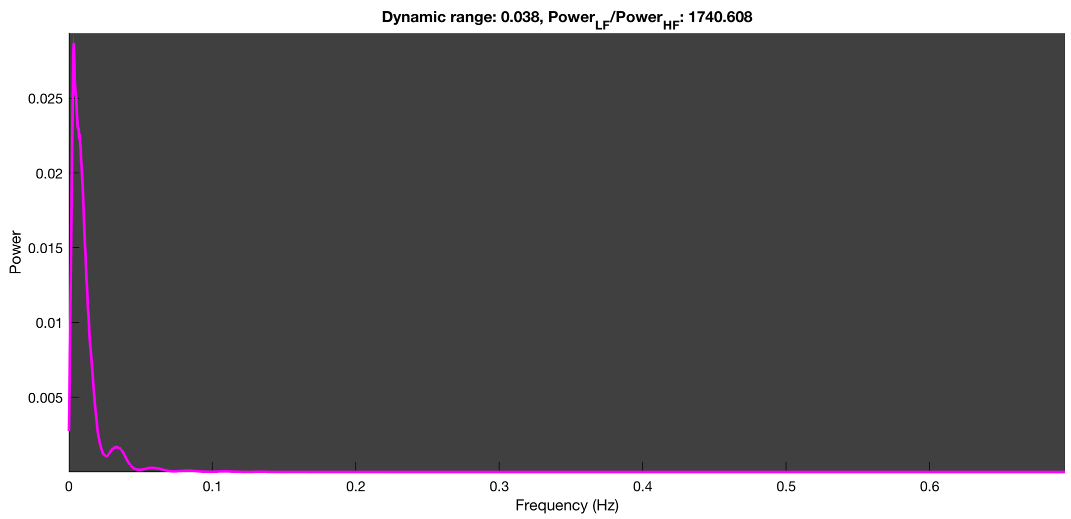

3-1 100n_tmap_component_ica_s1 056

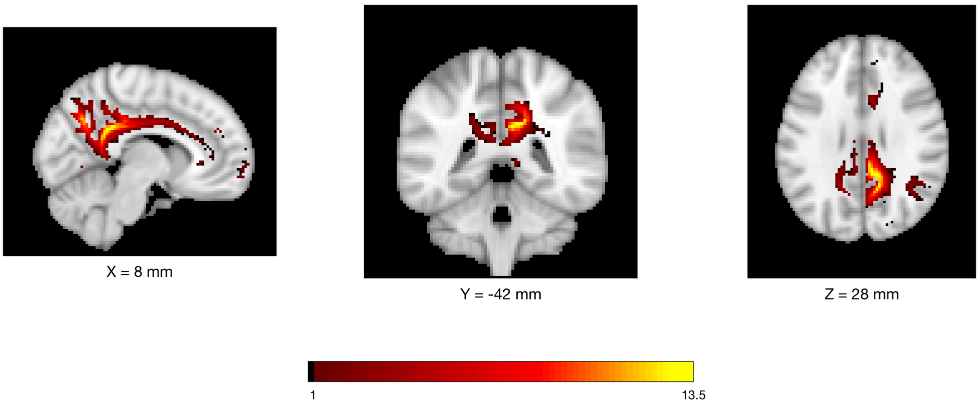


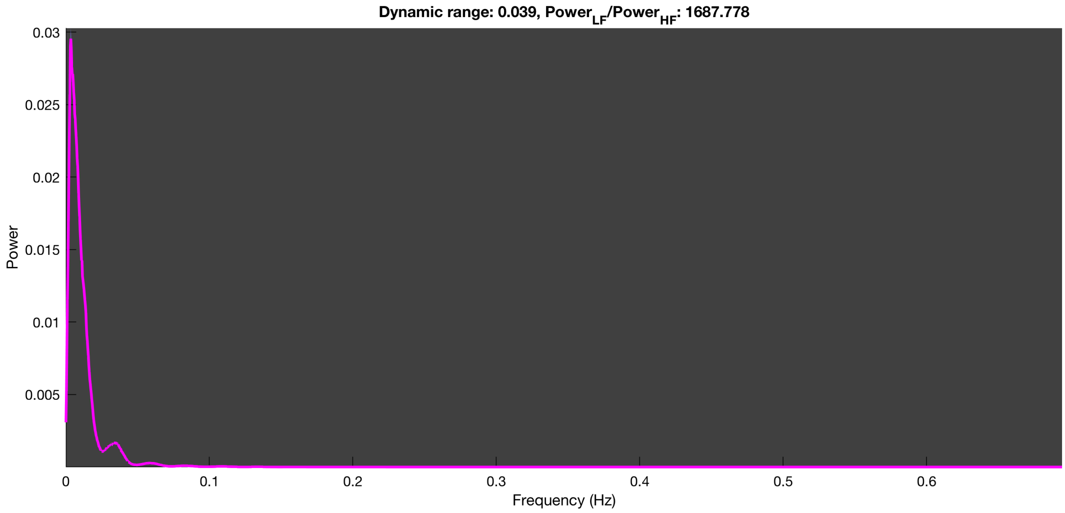

3-1 100n_tmap_component_ica_s1 057

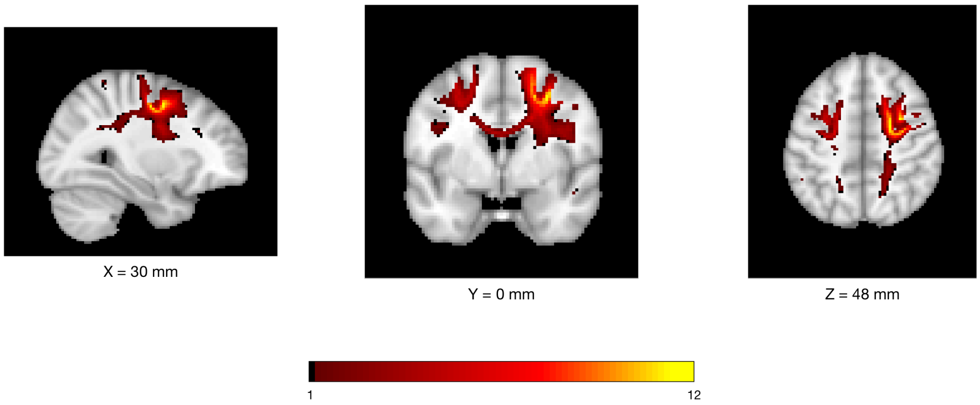


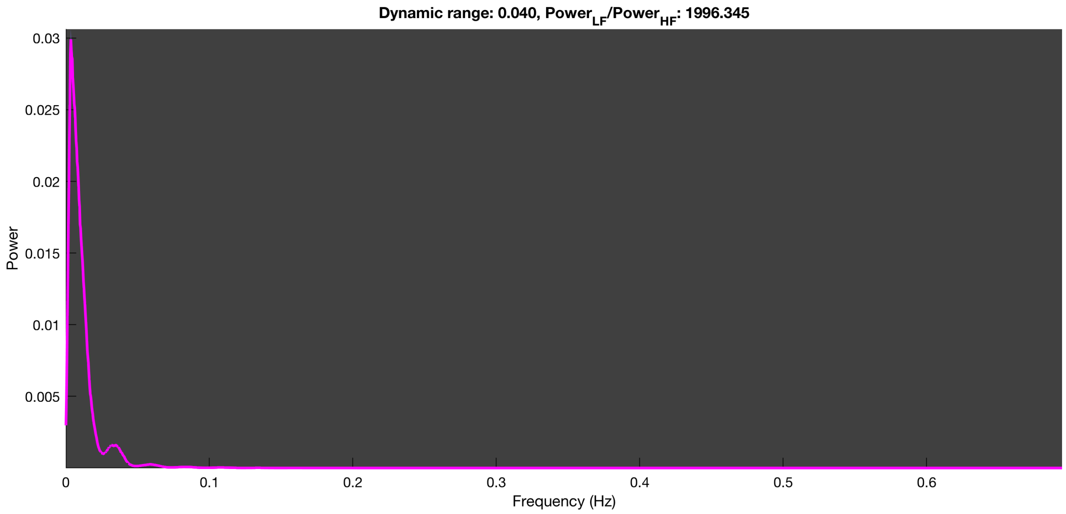

3-1 100n_tmap_component_ica_s1 058

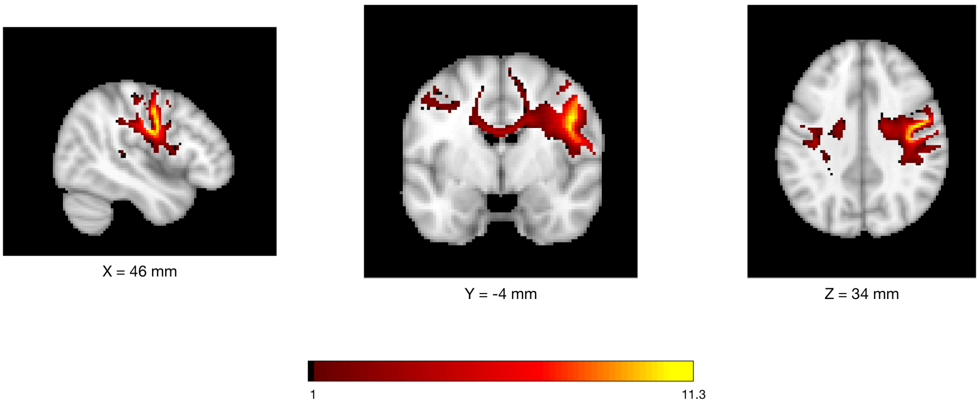


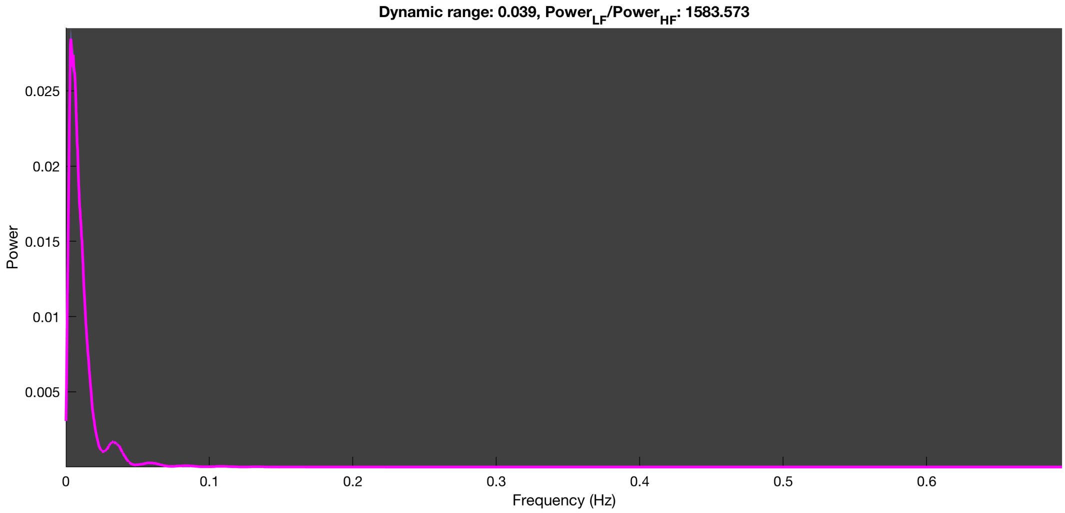

3-1 100n_tmap_component_ica_s1 059

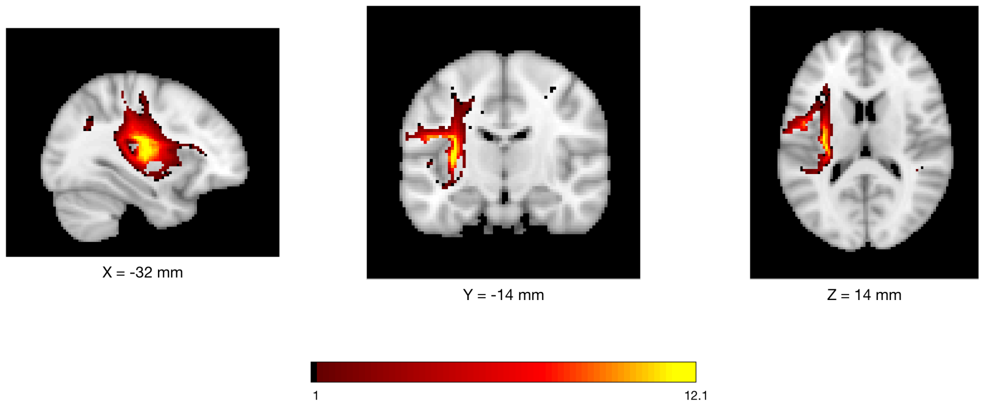


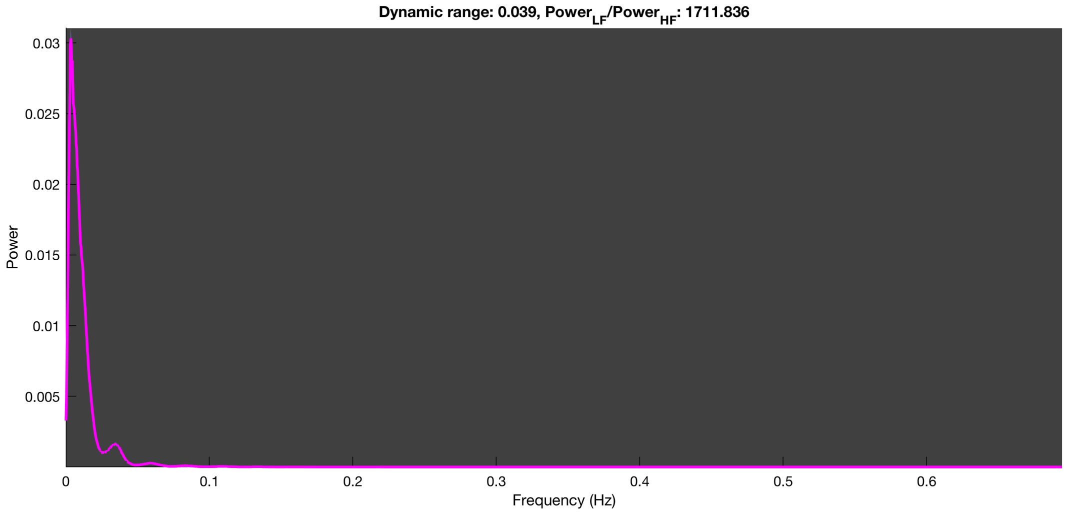

3-1 100n_tmap_component_ica_s1 060

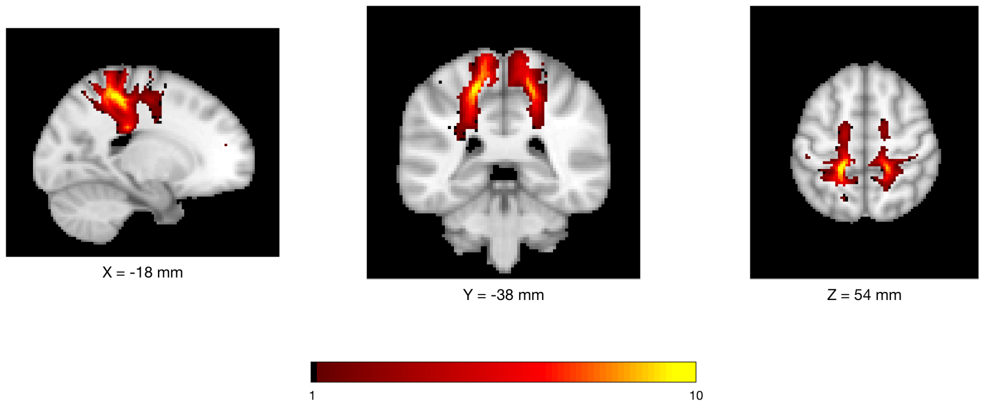


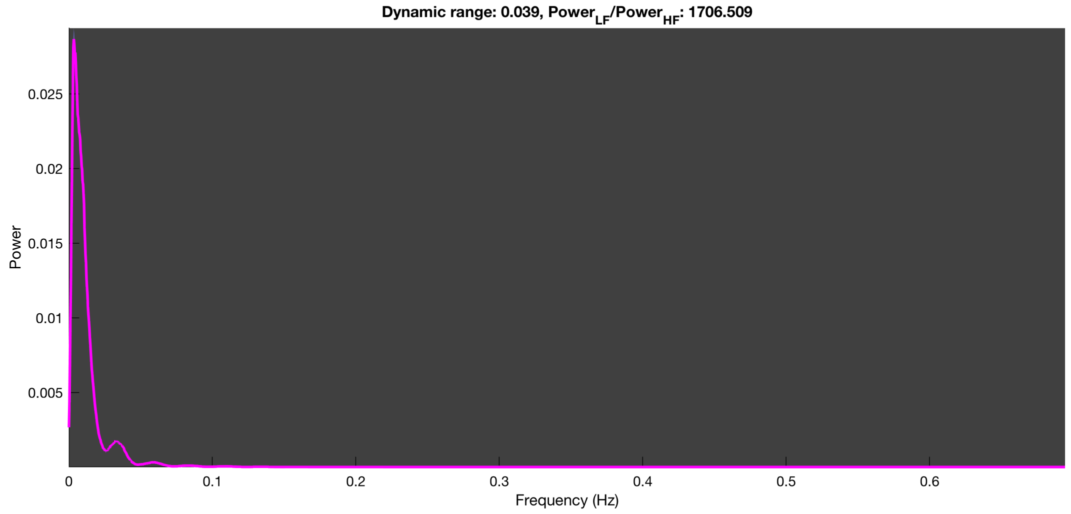

3-1 100n_tmap_component_ica_s1 061

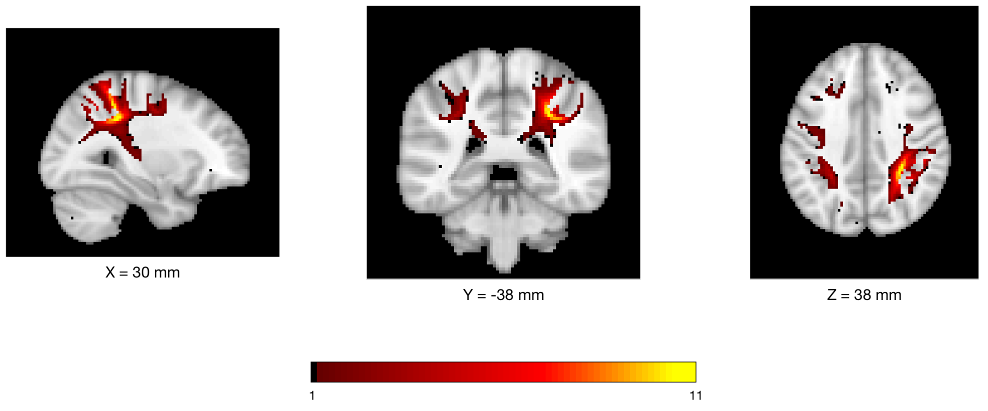


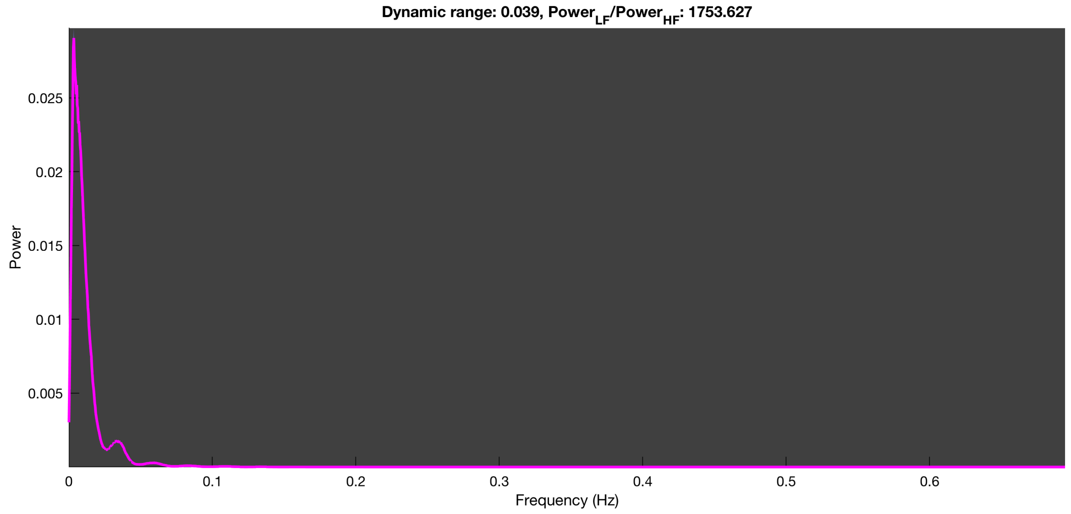

3-1 100n_tmap_component_ica_s1 062

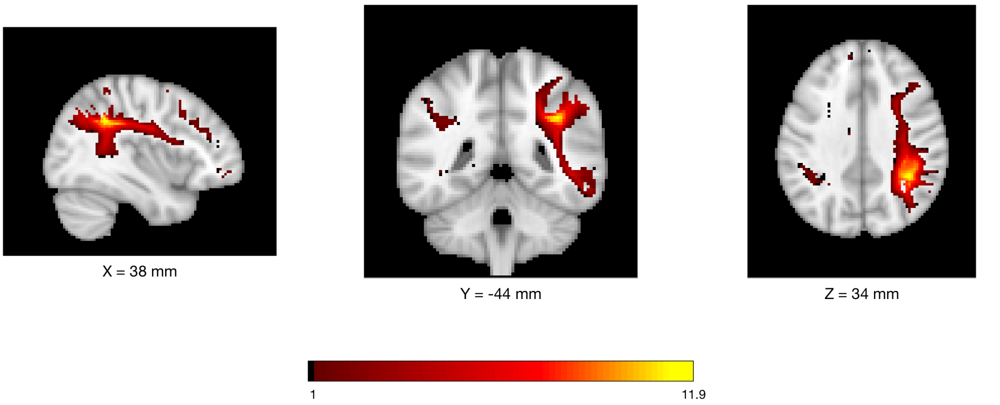


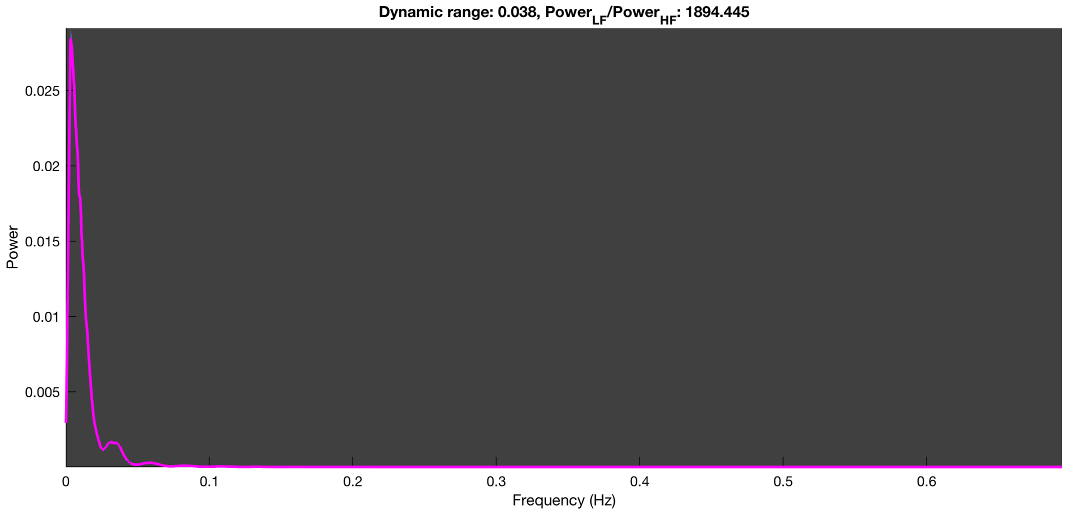

3-1 100n_tmap_component_ica_s1 063

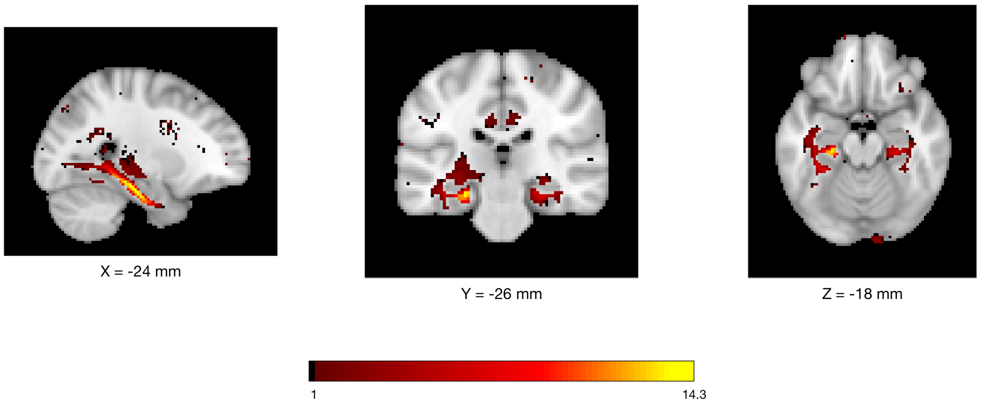


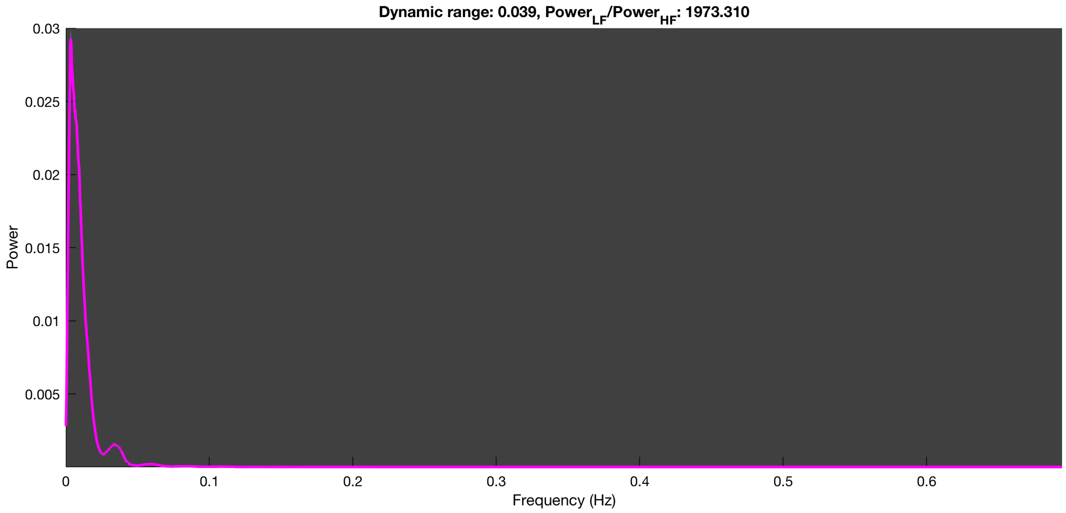

**3-1 100n_tmap_component_ica_s1 064**

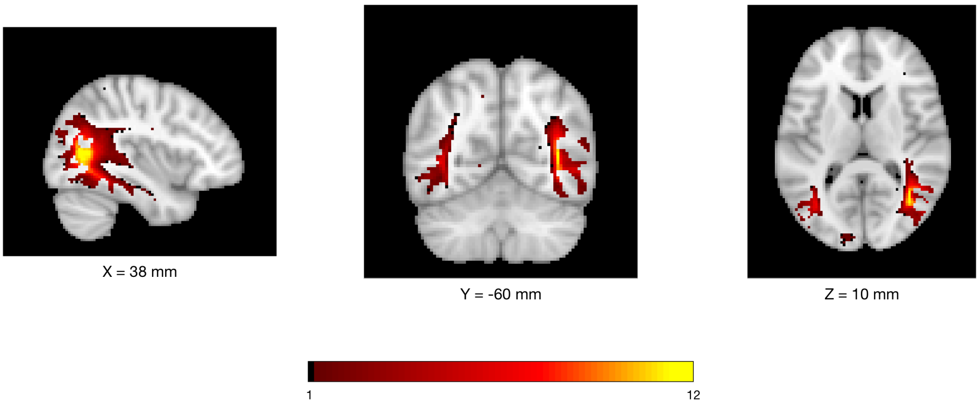


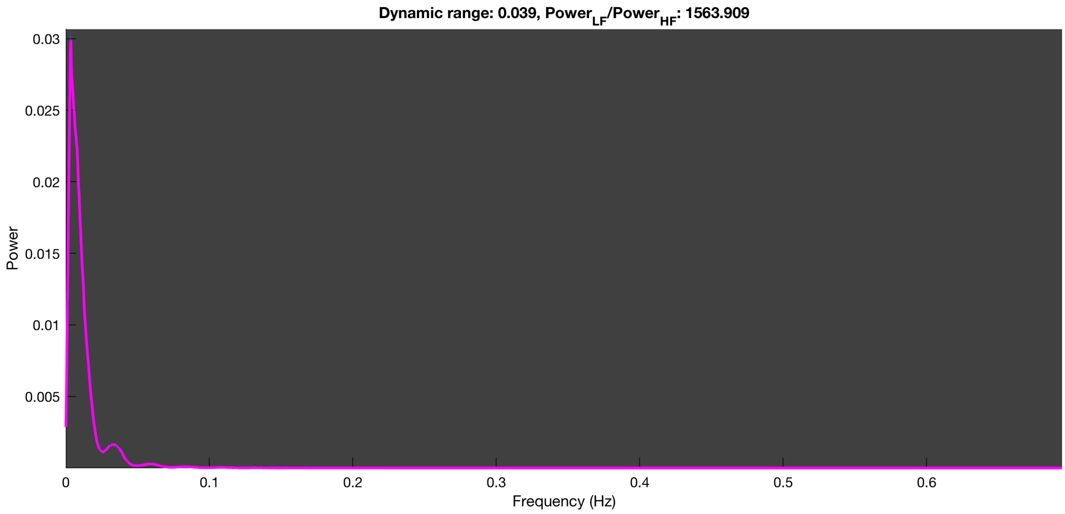

3-1 100n_tmap_component_ica_s1 065

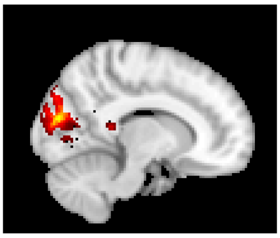

X = 12 mm

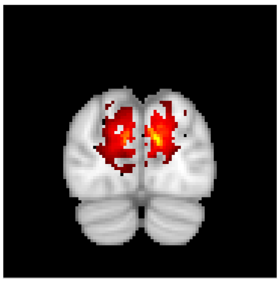

Y = -82 mm

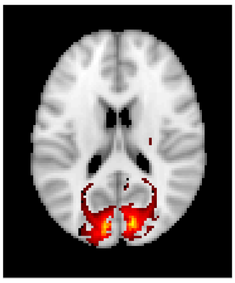

Z = 18 mm

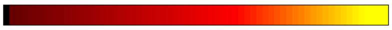


Dynamic range: 0.038, $Power_{LF}/Power_{HF}$: 1404.694

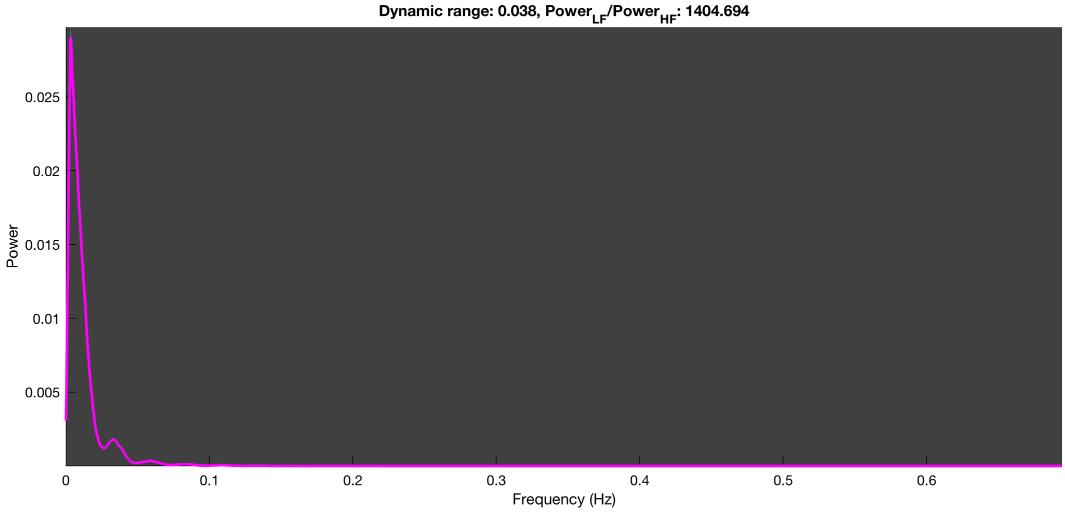

#### 3-1 100n_tmap_component_ica_s1 066

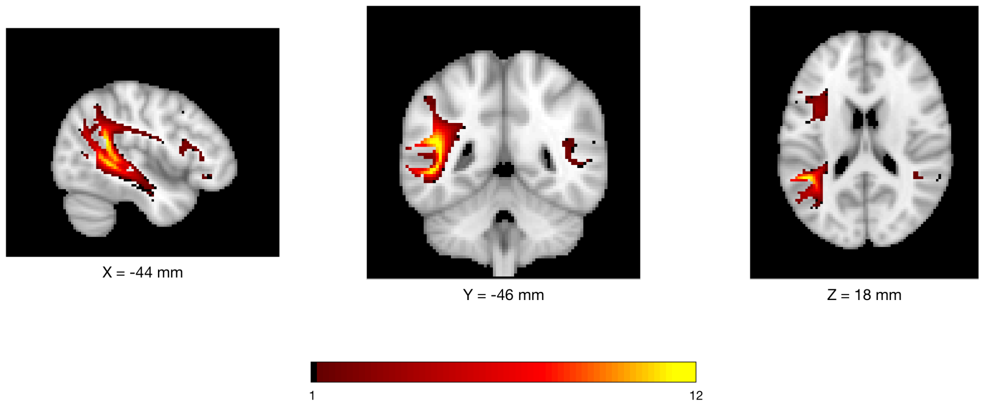


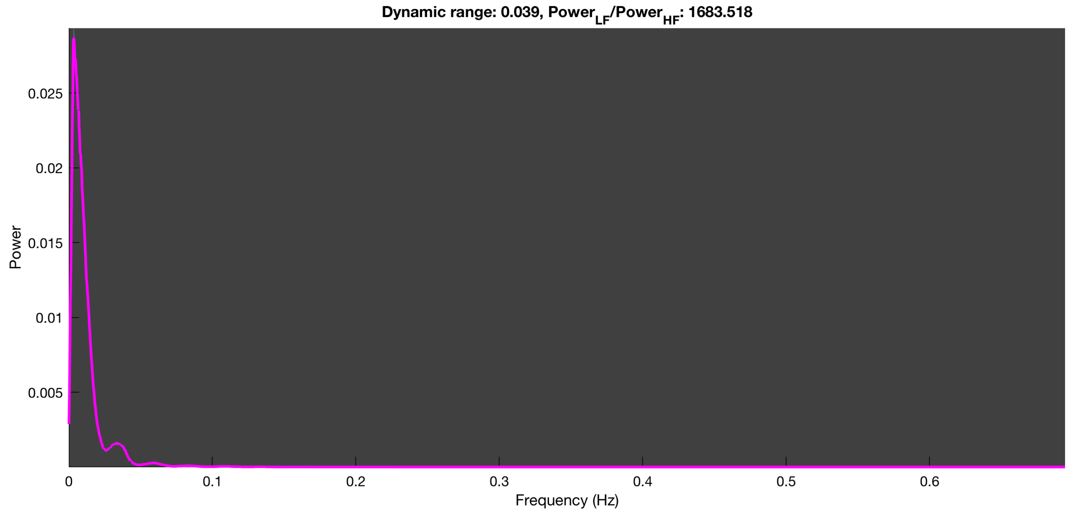

3-1 100n_tmap_component_ica_s1 067

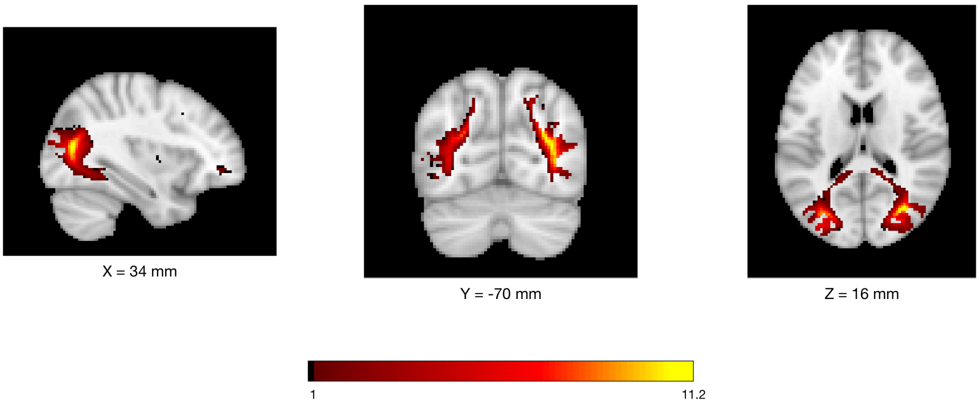


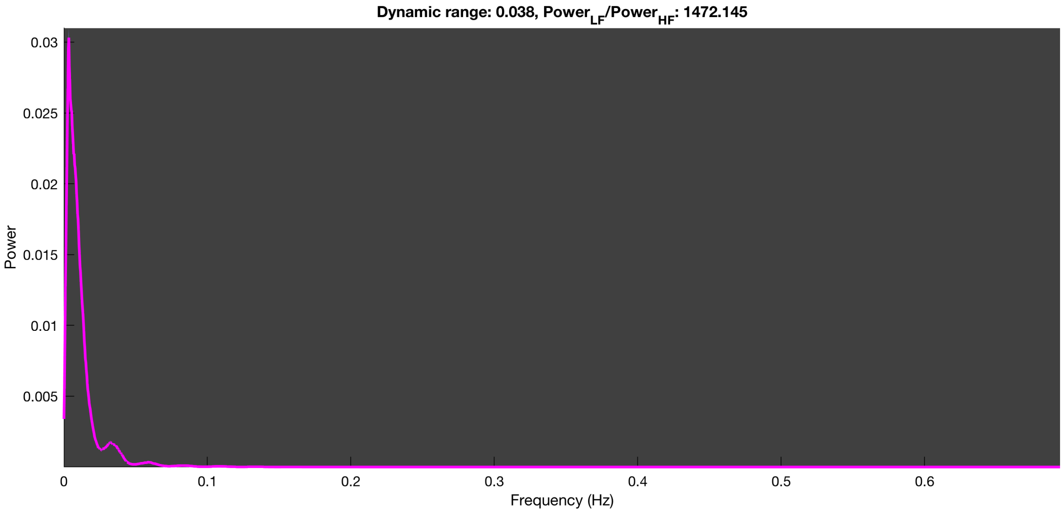

3-1 100n_tmap_component_ica_s1 068

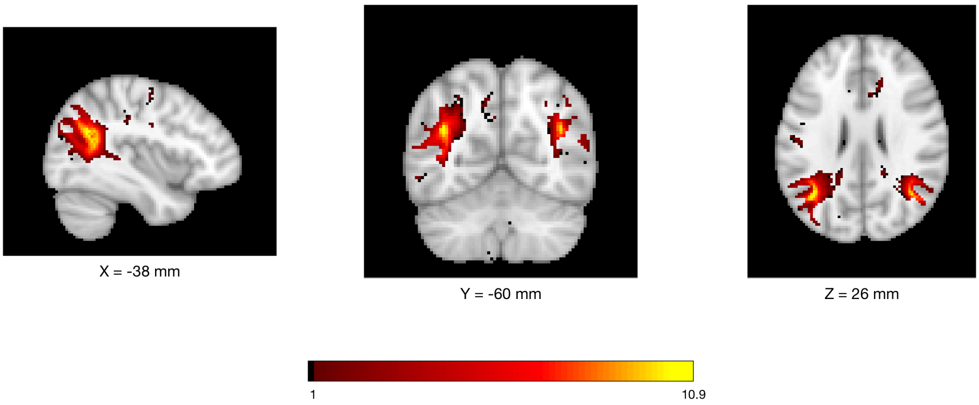


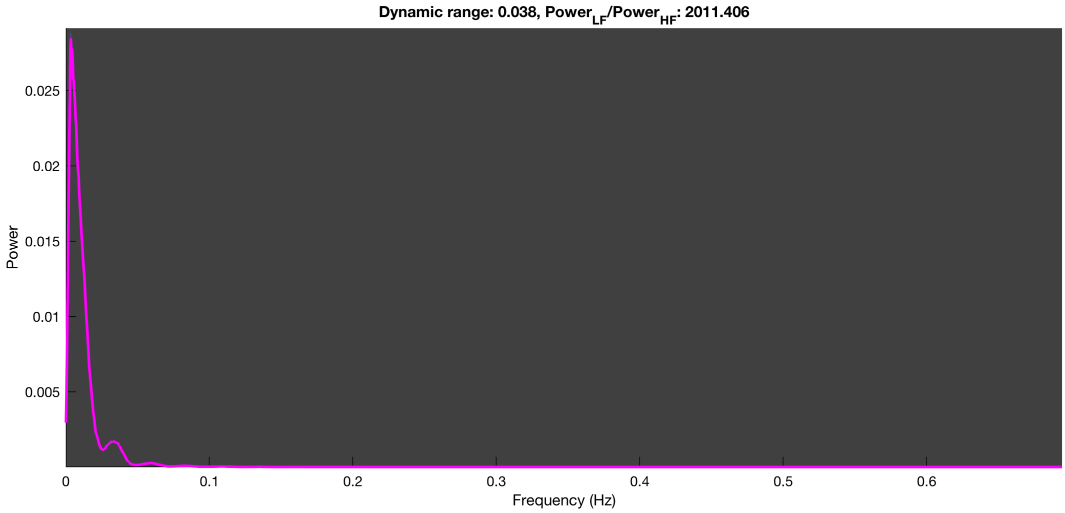

#### 3-1 100n_tmap_component_ica_s1 069

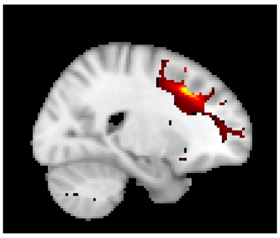

X = 22 mm

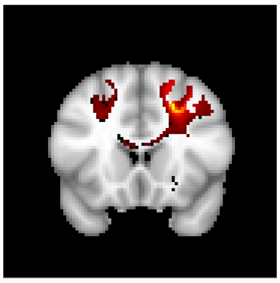

Y = 18 mm

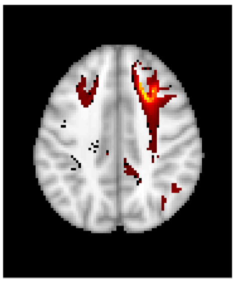

Z = 40 mm

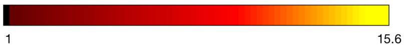


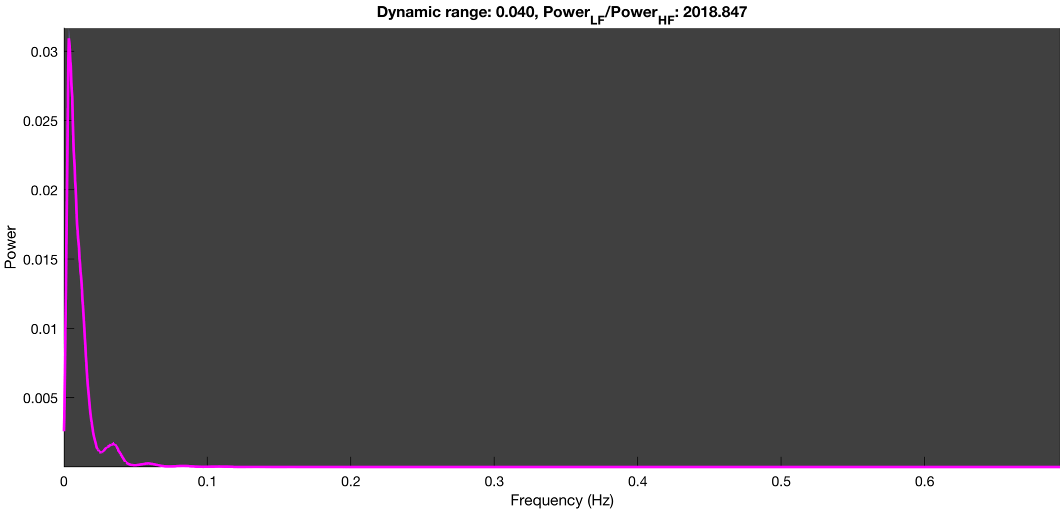

3-1 100n_tmap_component_ica_s1 070

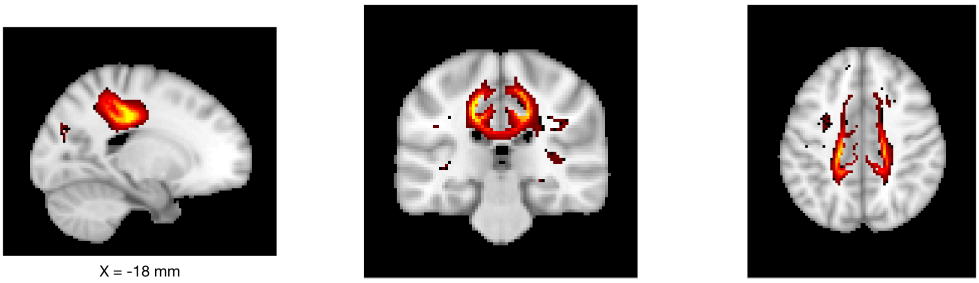

X = -18 mm

Y = -28 mm

Z = 42 mm

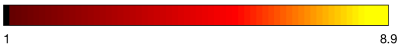


Dynamic range: 0.039, $Power_{LF}/Power_{HF}$: 1740.251

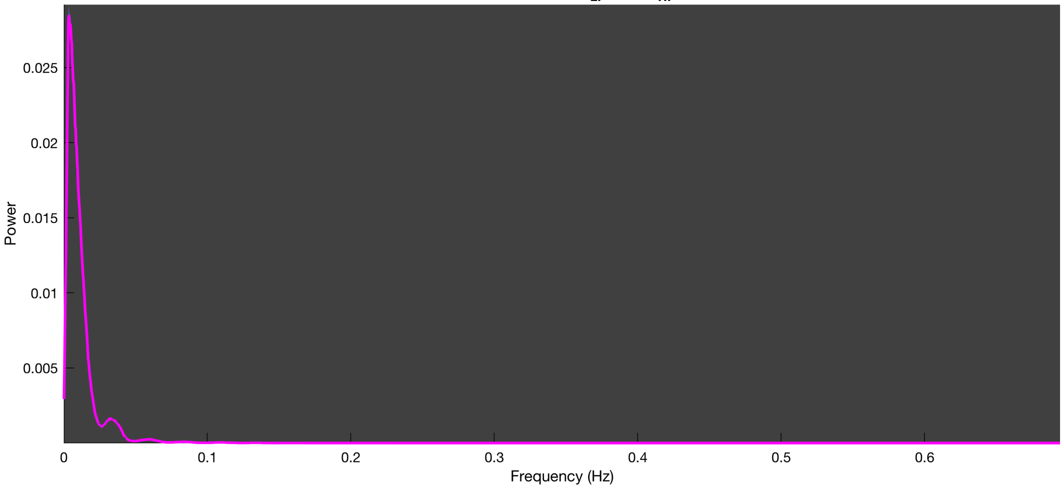

3-1 100n_tmap_component_ica_s1 071

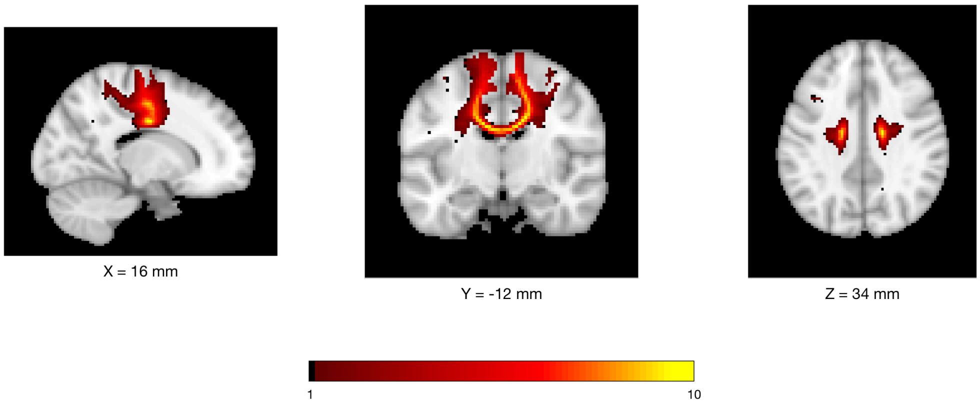


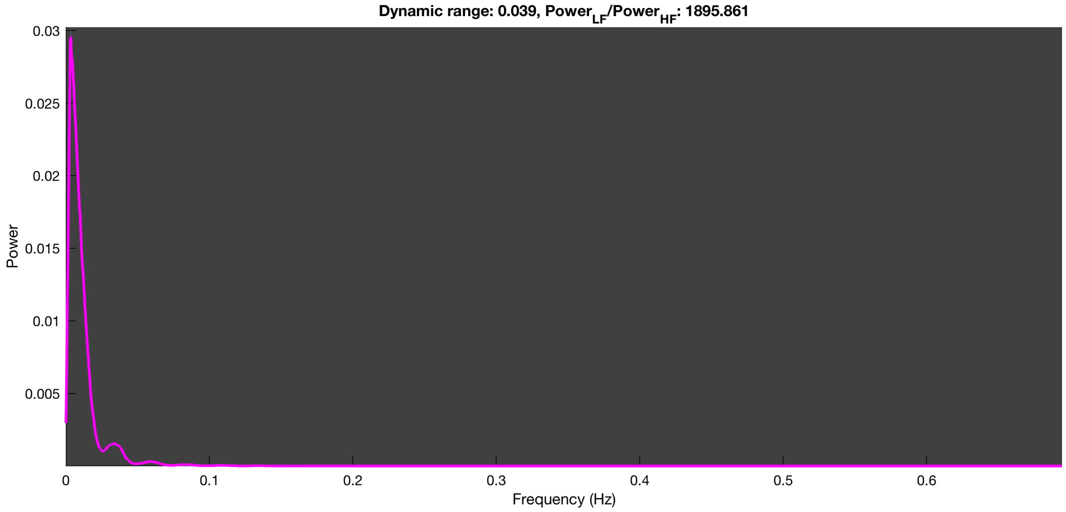

3-1 100n_tmap_component_ica_s1 072

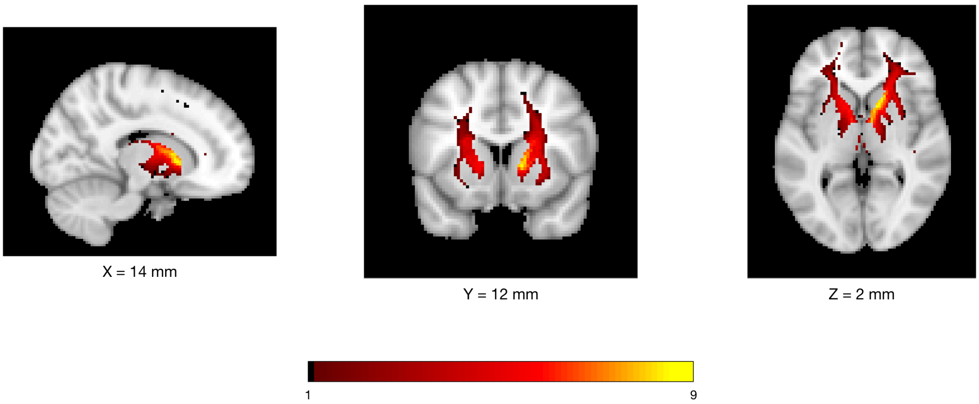


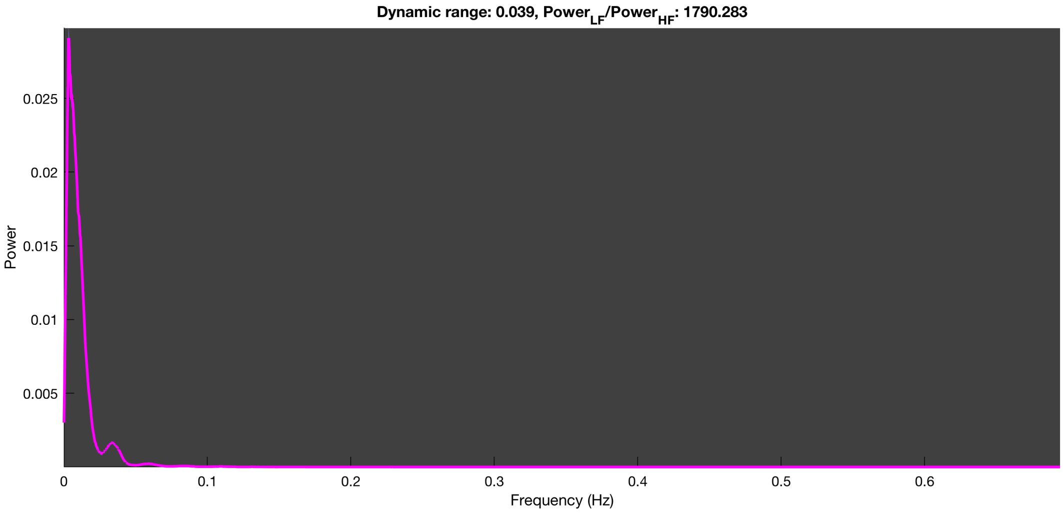

3-1 100n_tmap_component_ica_s1 073

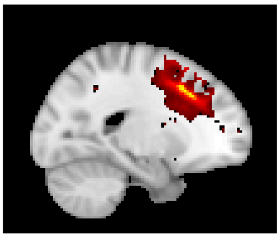

X = -20 mm

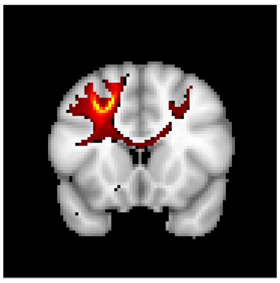

Y = 16 mm

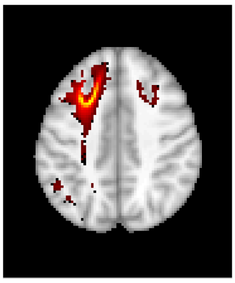

Z = 42 mm

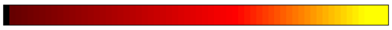


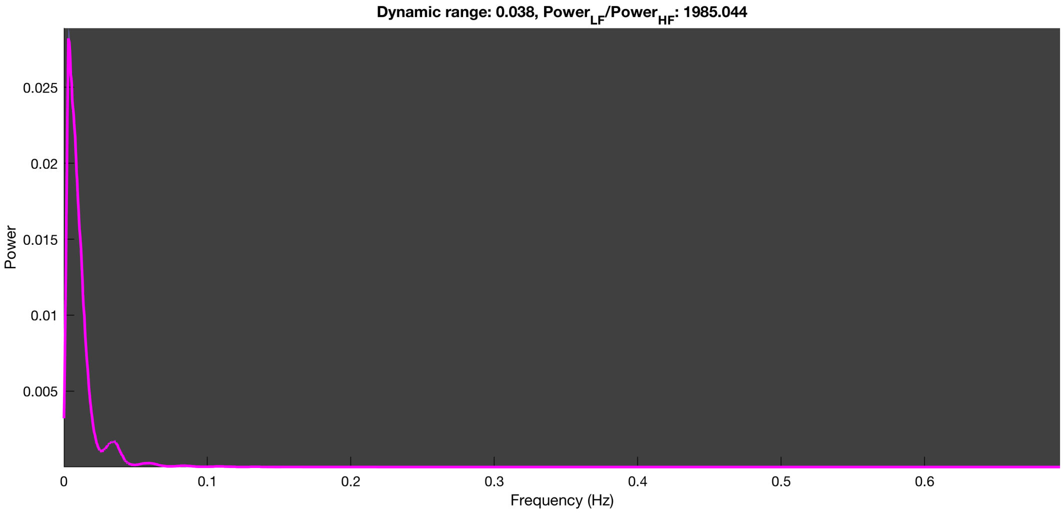

3-1 100n_tmap_component_ica_s1 074

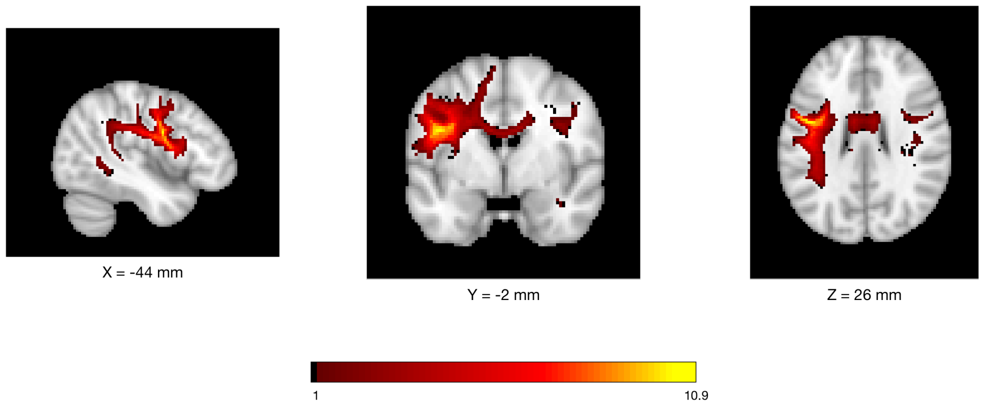


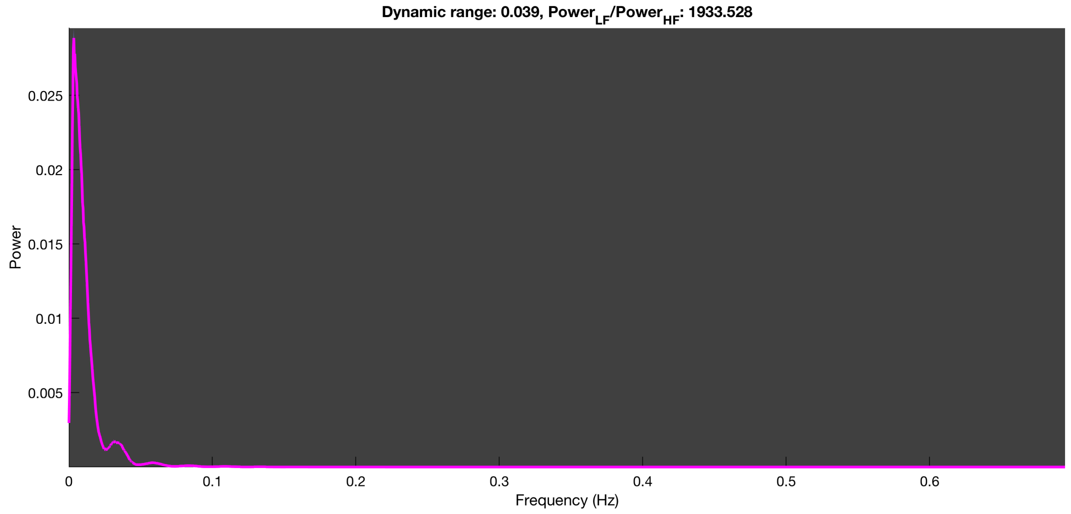

3-1 100n_tmap_component_ica_s1 075

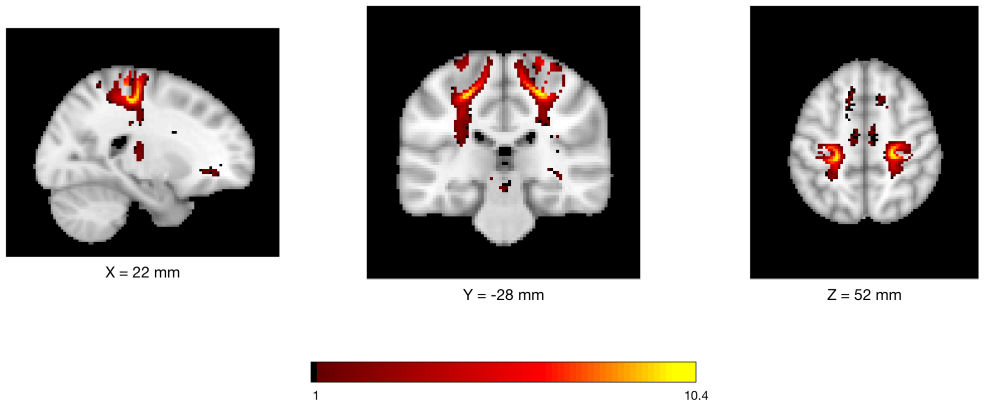


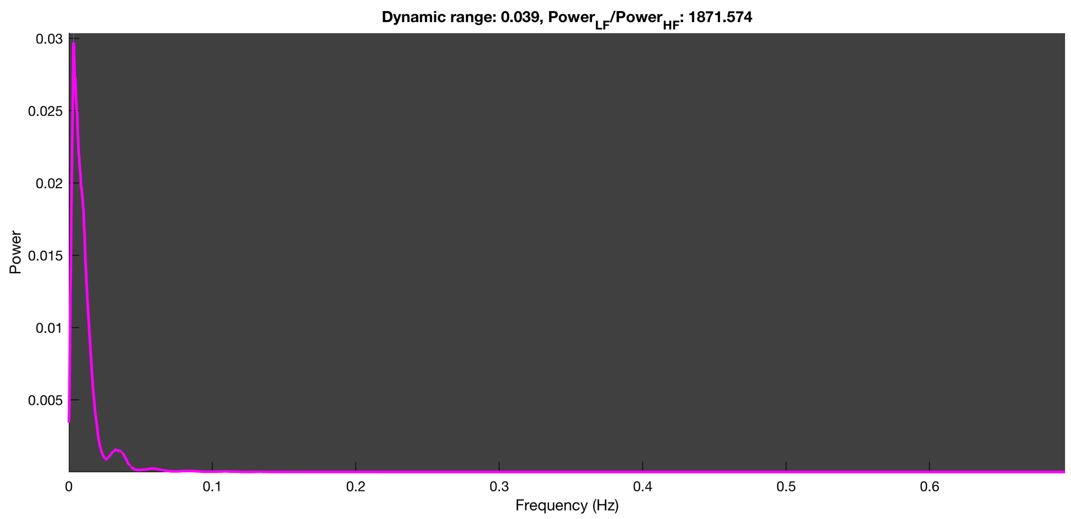

3-1 100n_tmap_component_ica_s1 076

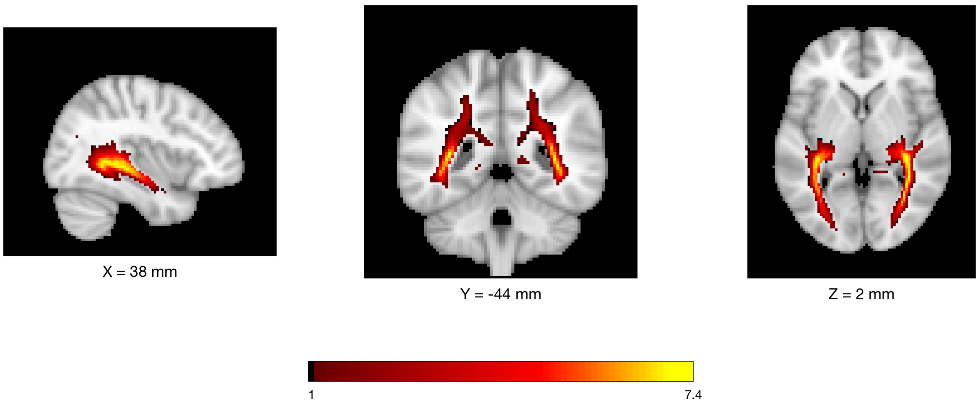


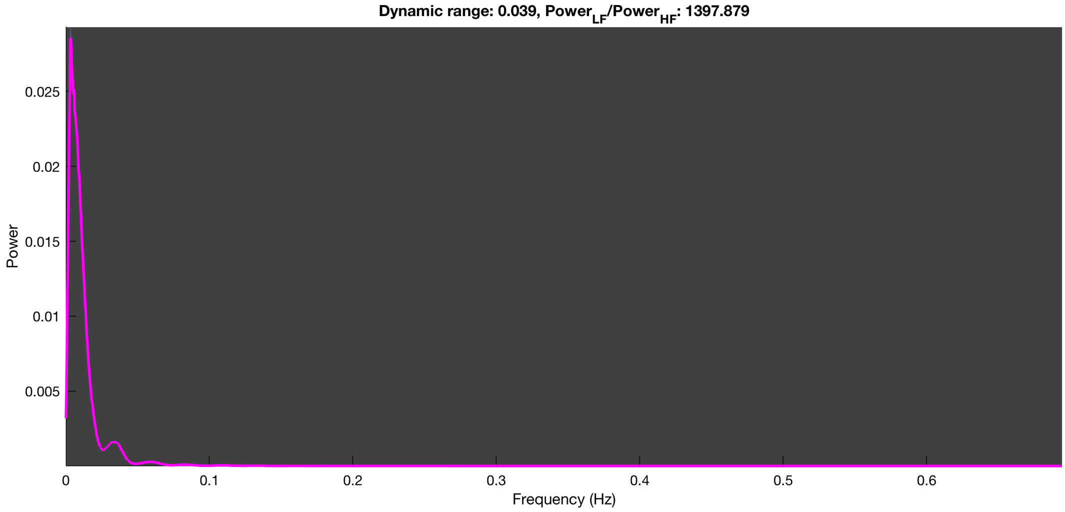

3-1 100n_tmap_component_ica_s1 077

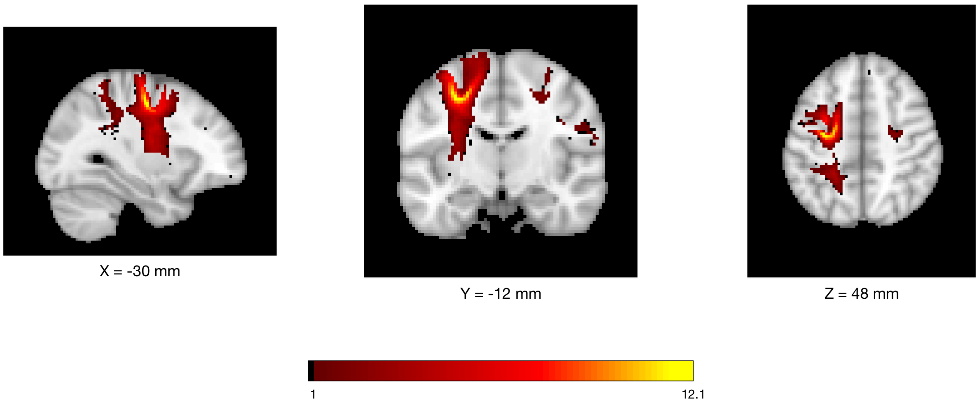


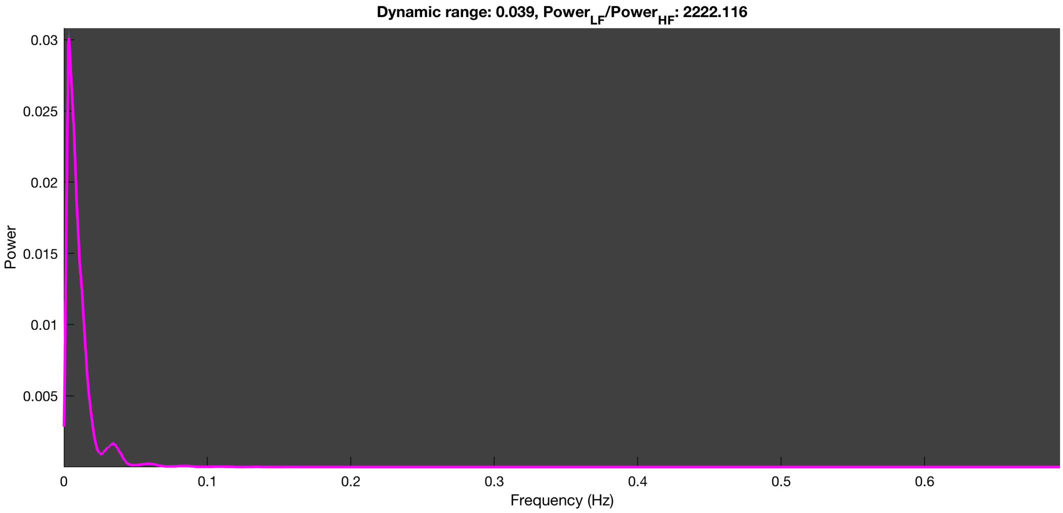

3-1 100n_tmap_component_ica_s1 078

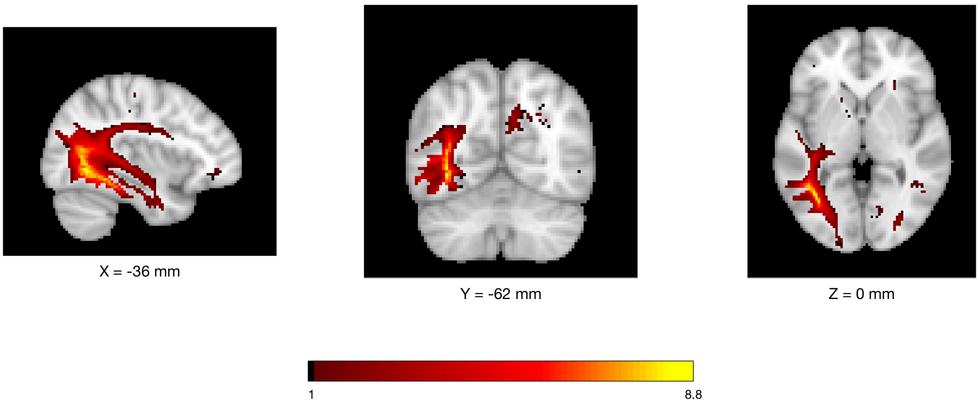


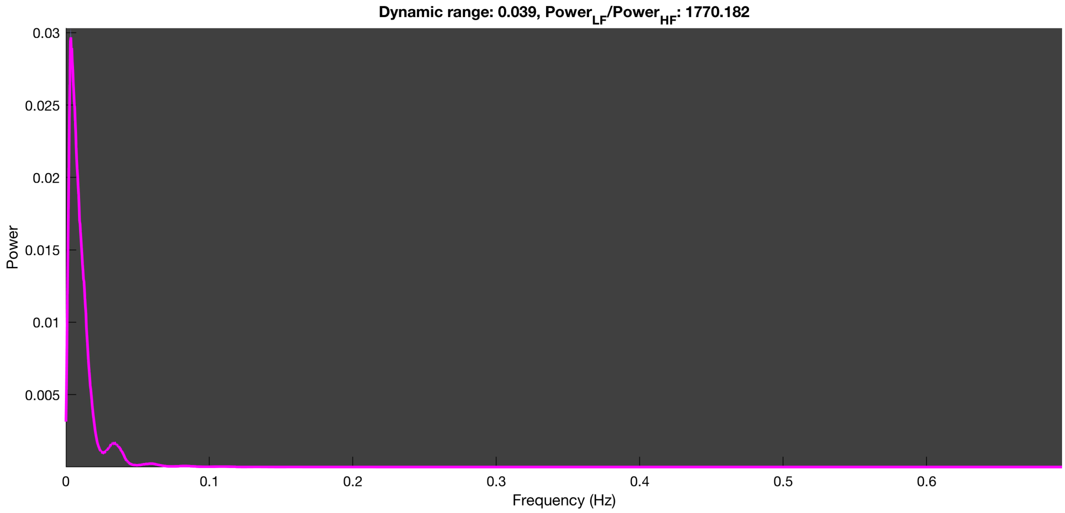

3-1 100n_tmap_component_ica_s1 079

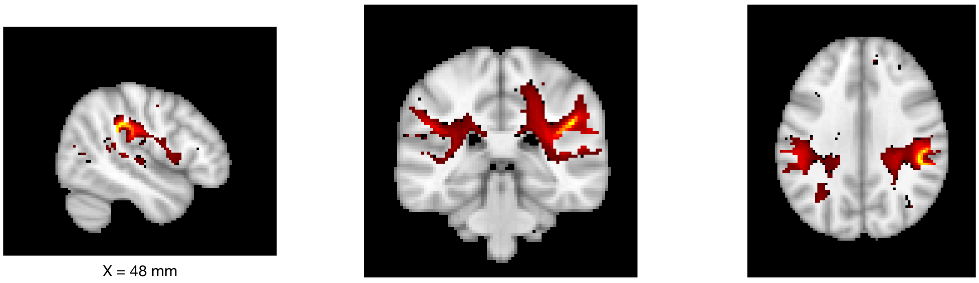

X = 48 mm    Y = -34 mm    Z = 30 mm

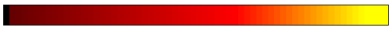


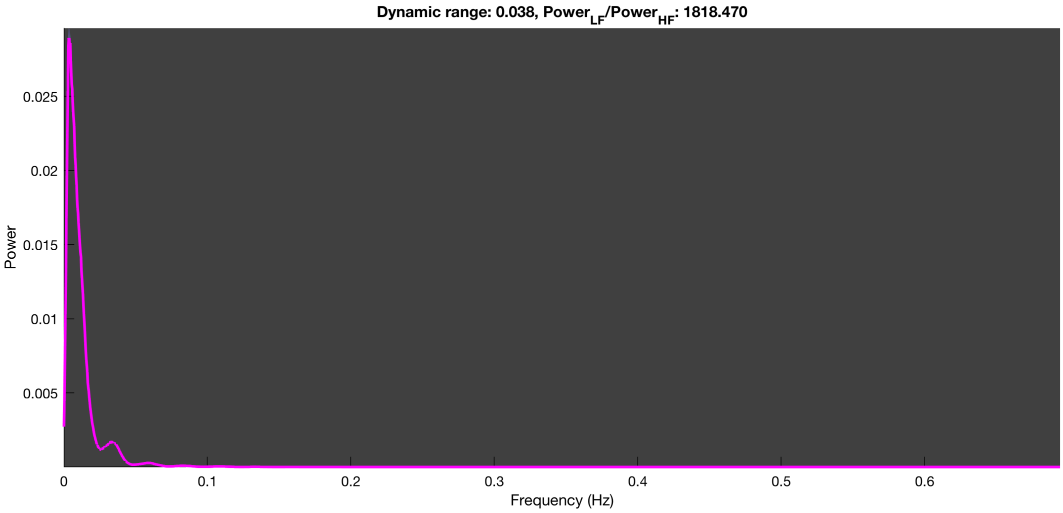

#### 3-1 100n_tmap_component_ica_s1 080

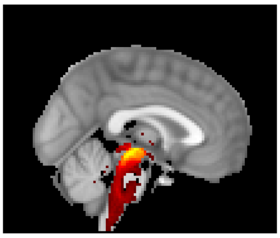

X = 0 mm

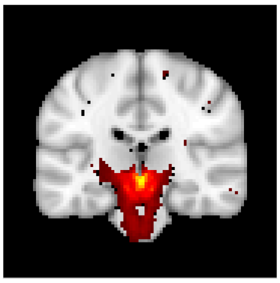

Y = -24 mm

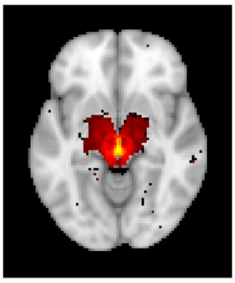

Z = -8 mm

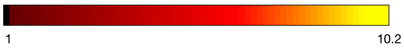


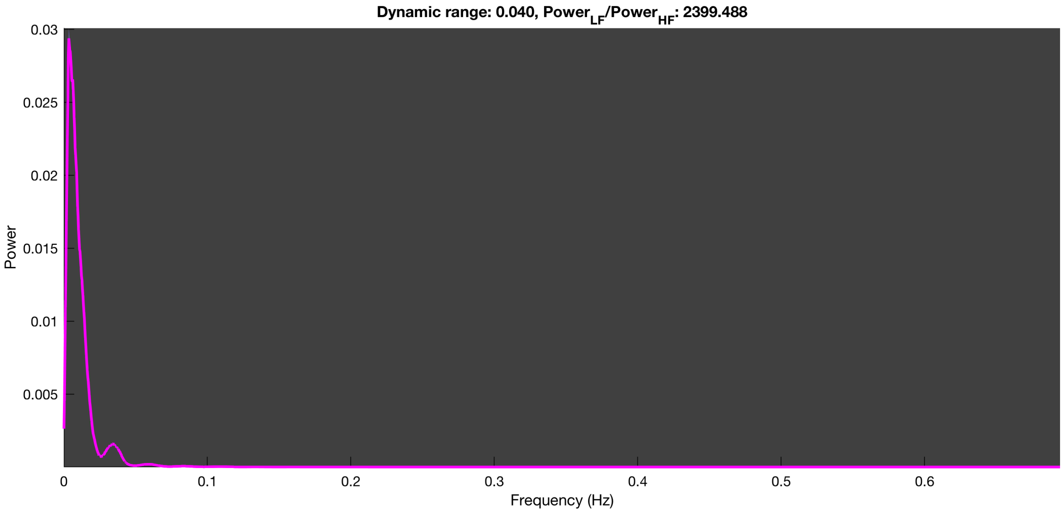

3-1 100n_tmap_component_ica_s1 081

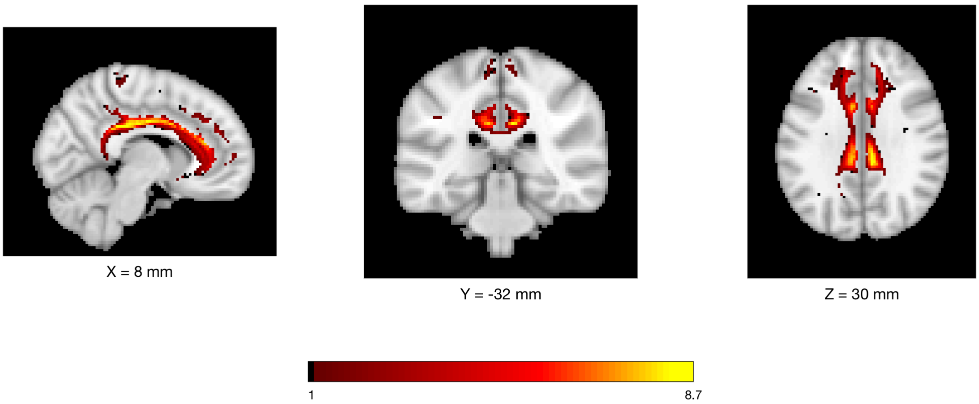


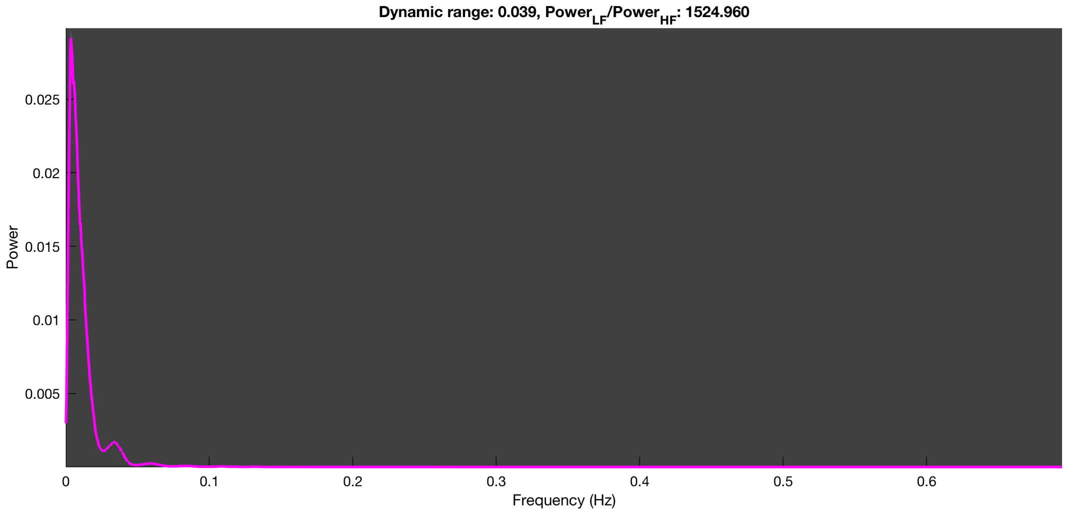

3-1 100n_tmap_component_ica_s1 082

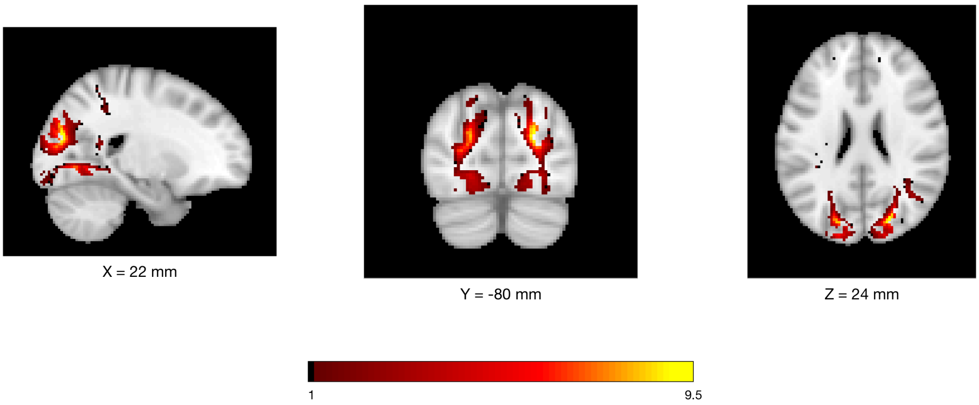


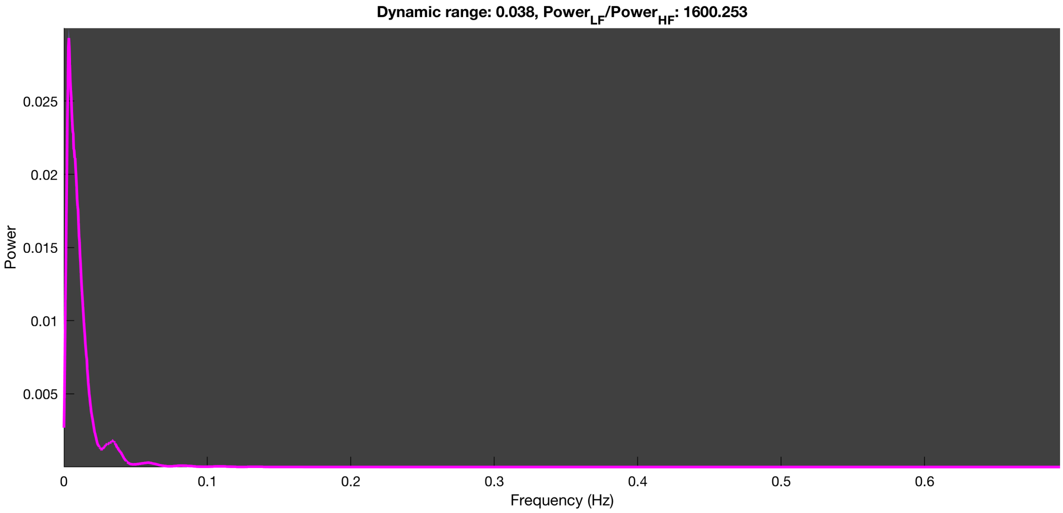

3-1 100n_tmap_component_ica_s1 083

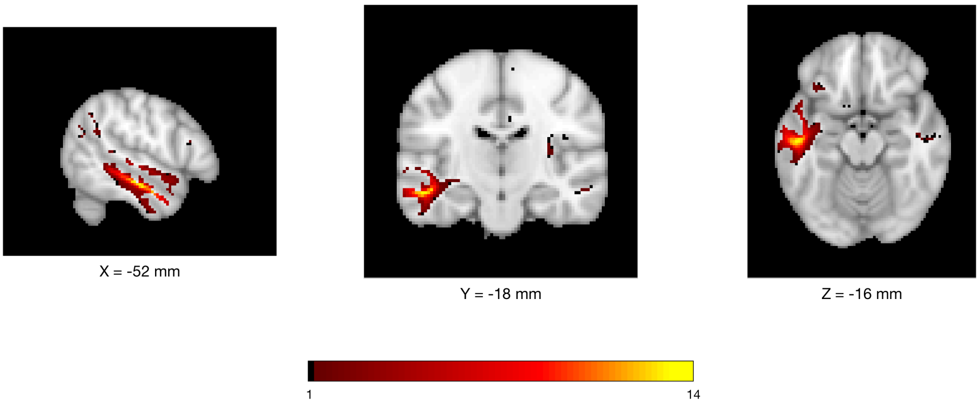


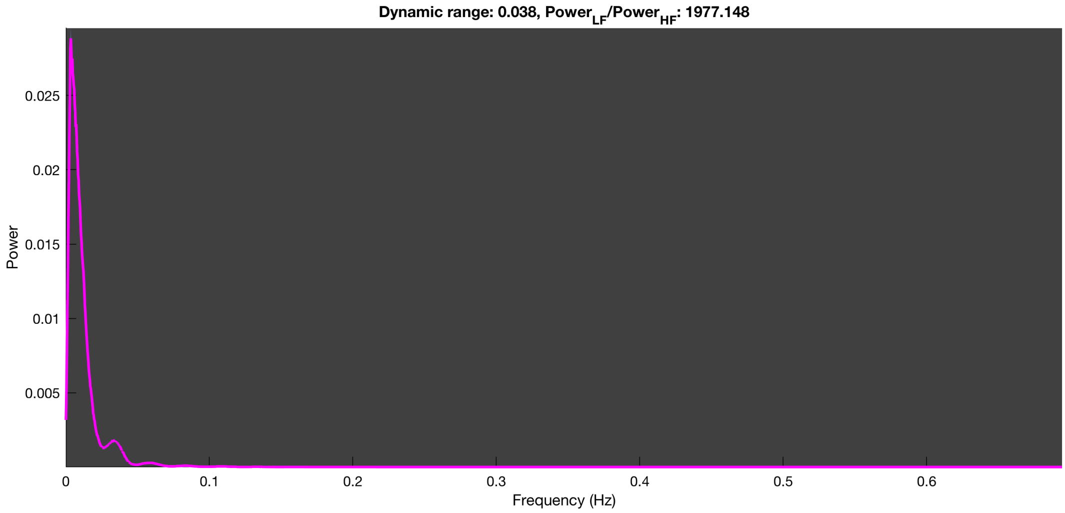

3-1 100n_tmap_component_ica_s1 084

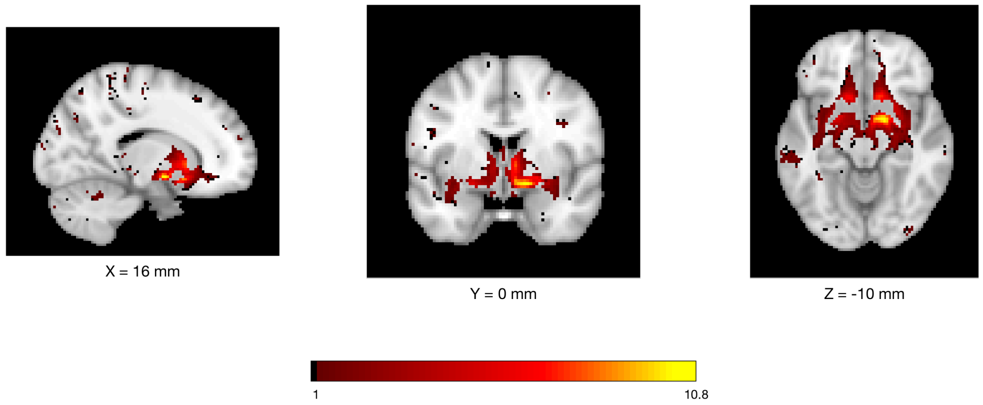


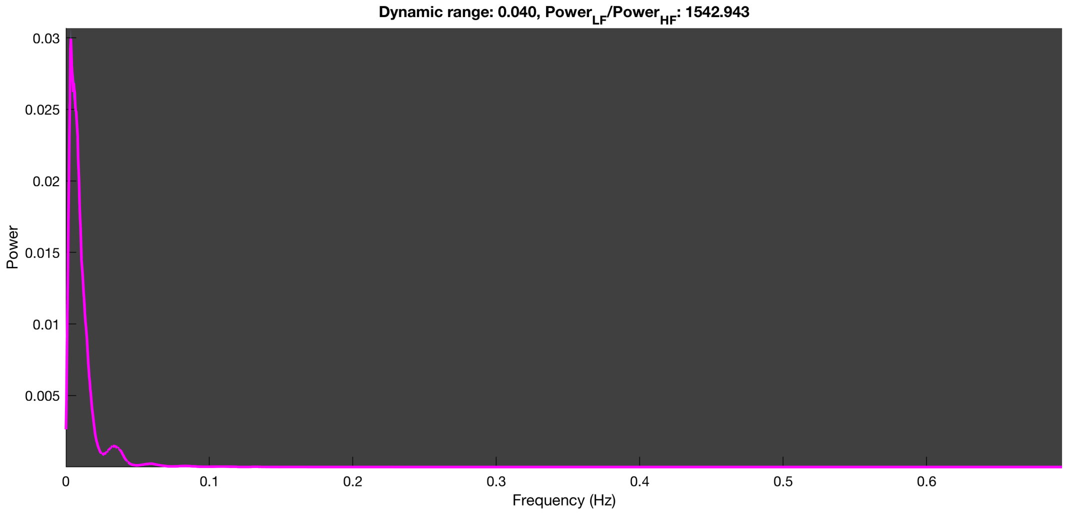

3-1 100n_tmap_component_ica_s1 085

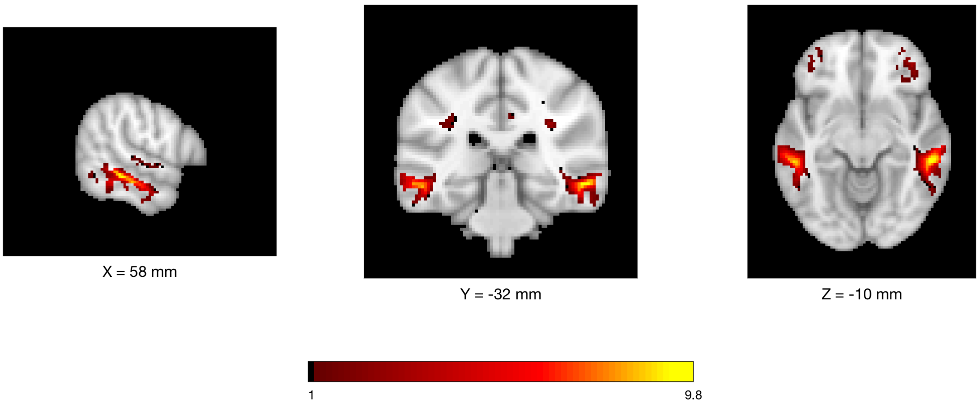


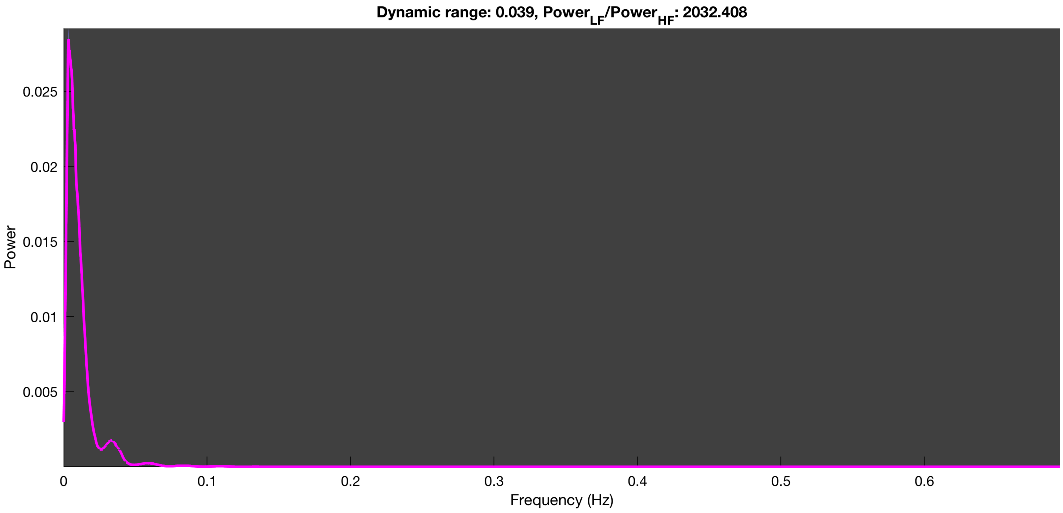

3-1 100n_tmap_component_ica_s1 086

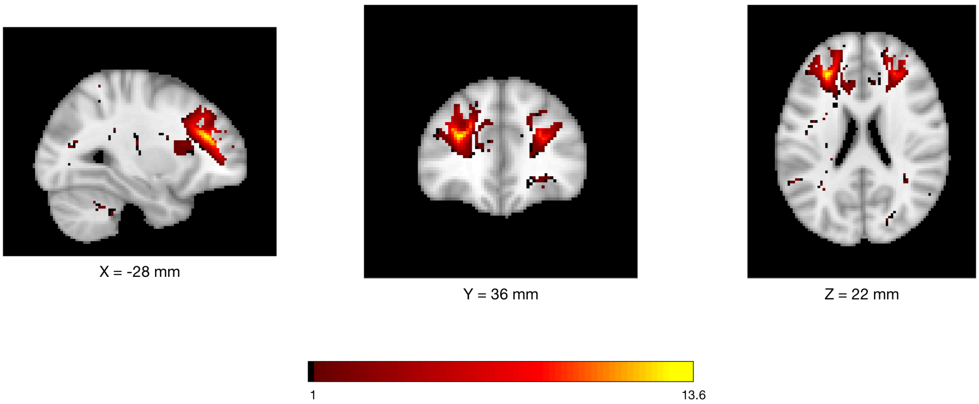


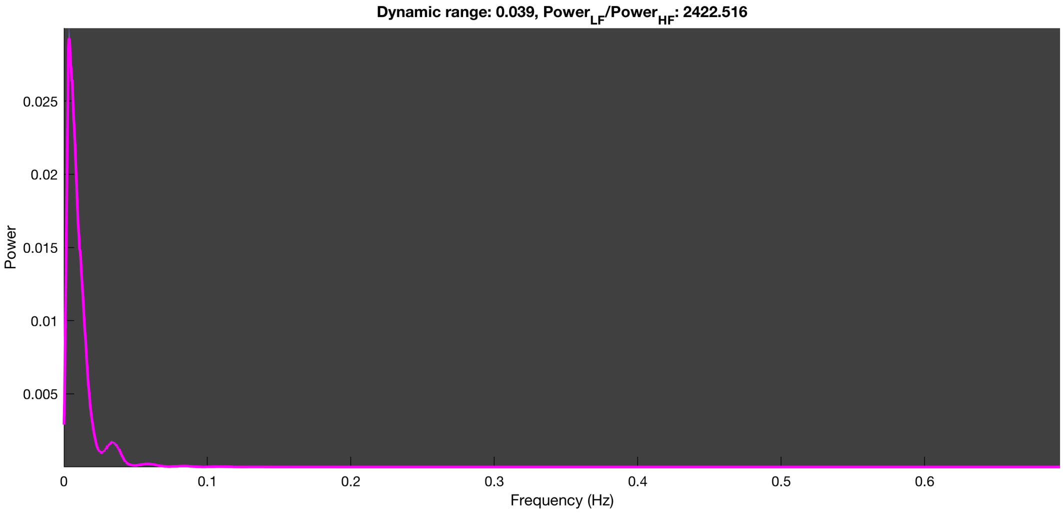

3-1 100n_tmap_component_ica_s1 087

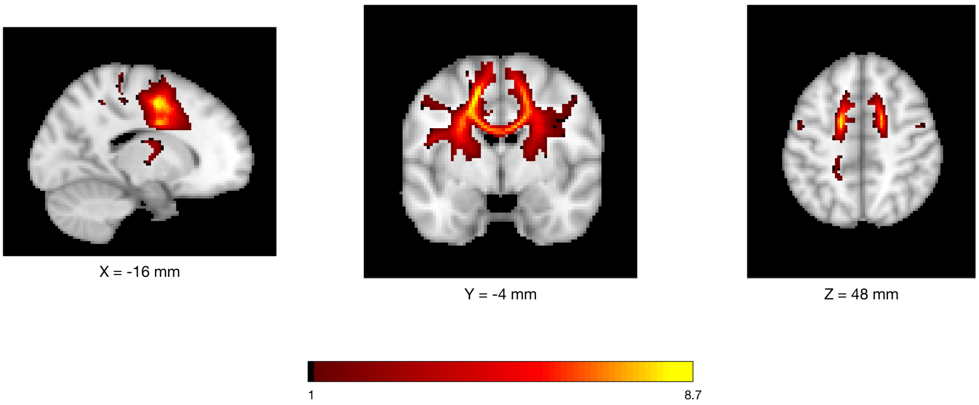


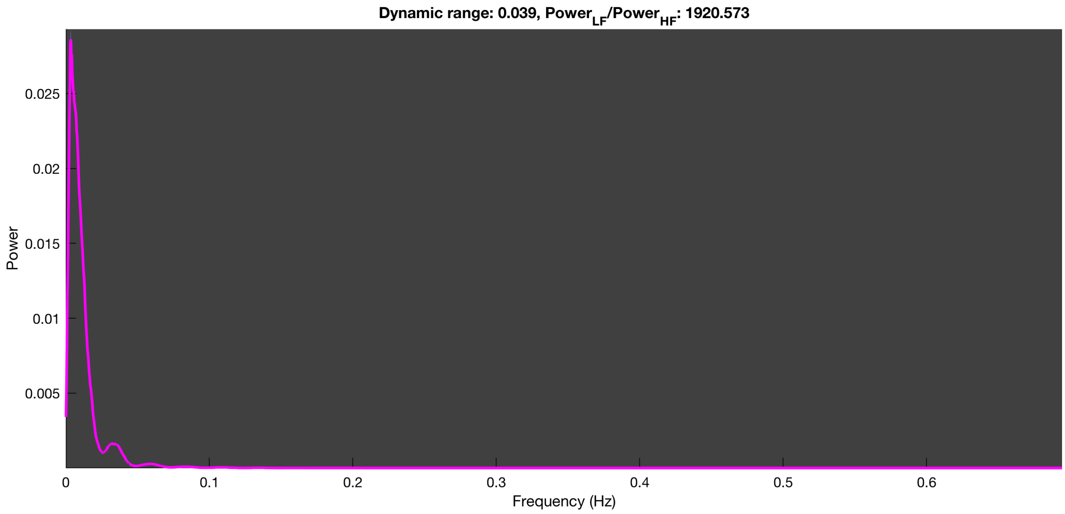

3-1 100n_tmap_component_ica_s1 088

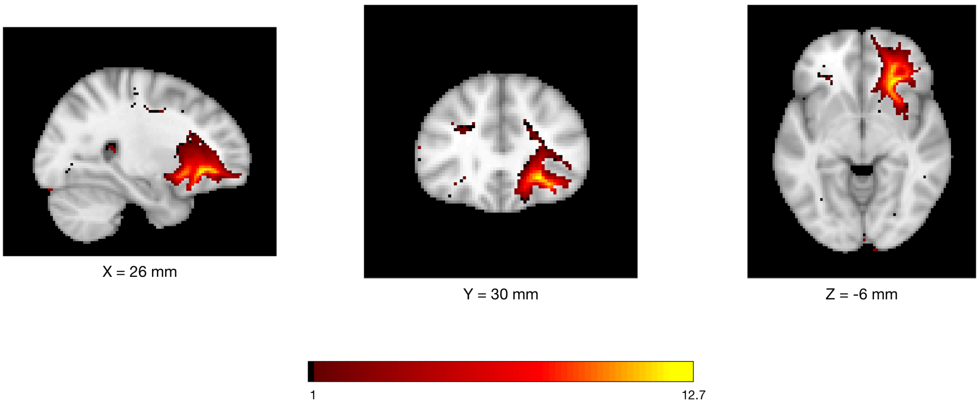


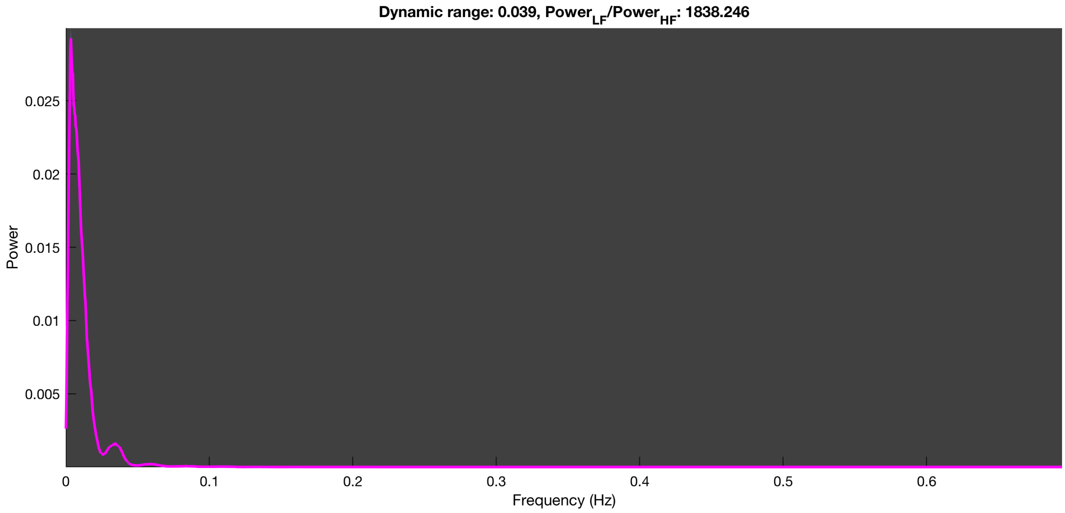

3-1 100n_tmap_component_ica_s1 089

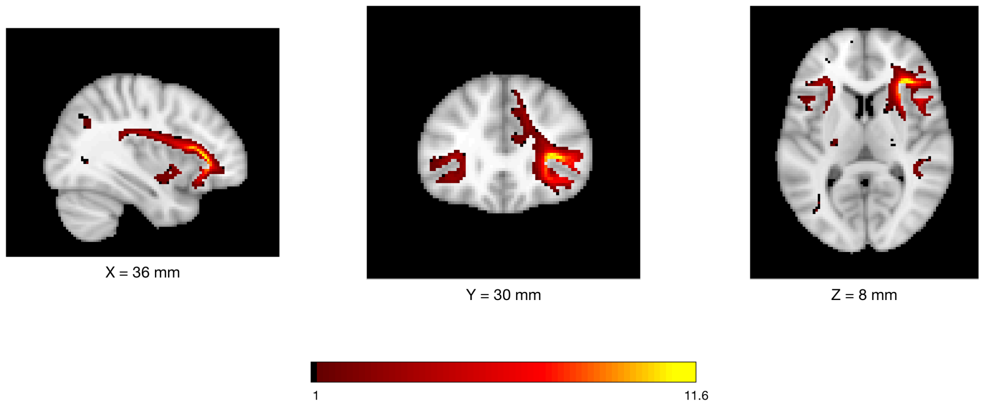


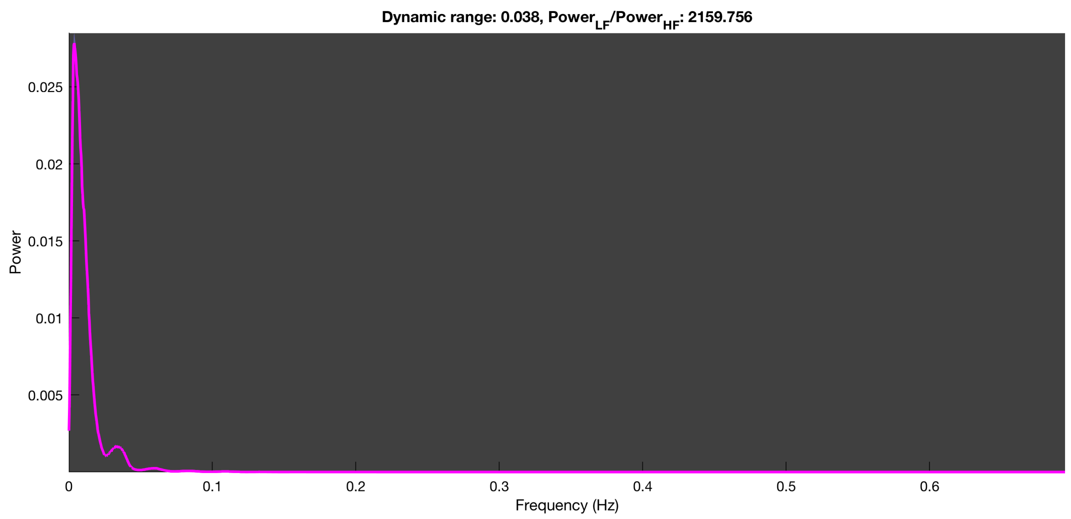

3-1 100n_tmap_component_ica_s1 090

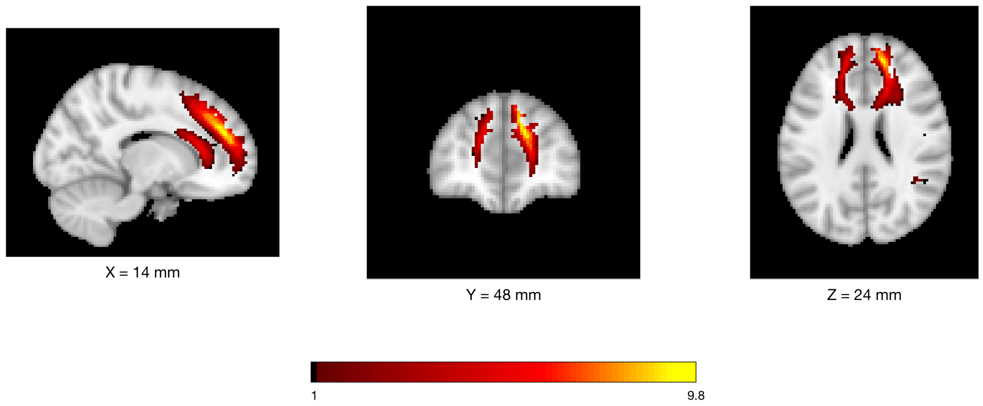


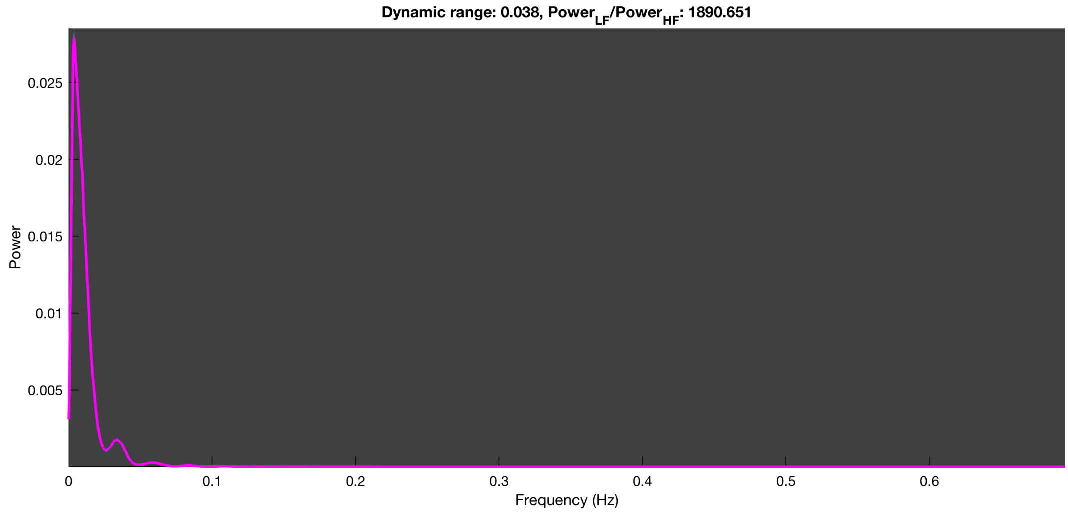

##### 3-1 100n_tmap_component_ica_s1 091

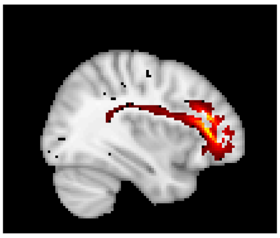

X = 36 mm

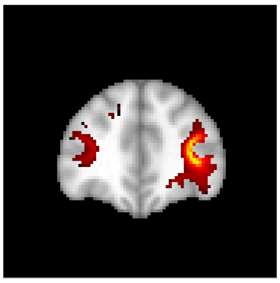

Y = 38 mm

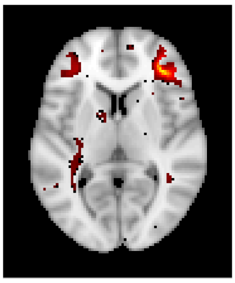

Z = 6 mm

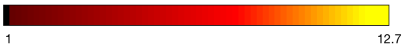


Dynamic range: 0.039, $Power_{LF}/Power_{HF}$: 1916.178

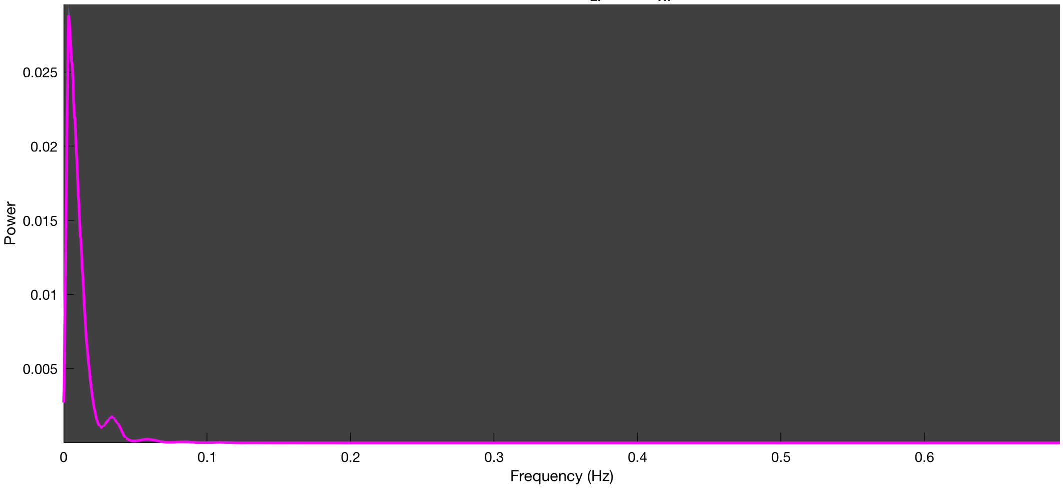

3-1 100n_tmap_component_ica_s1 092

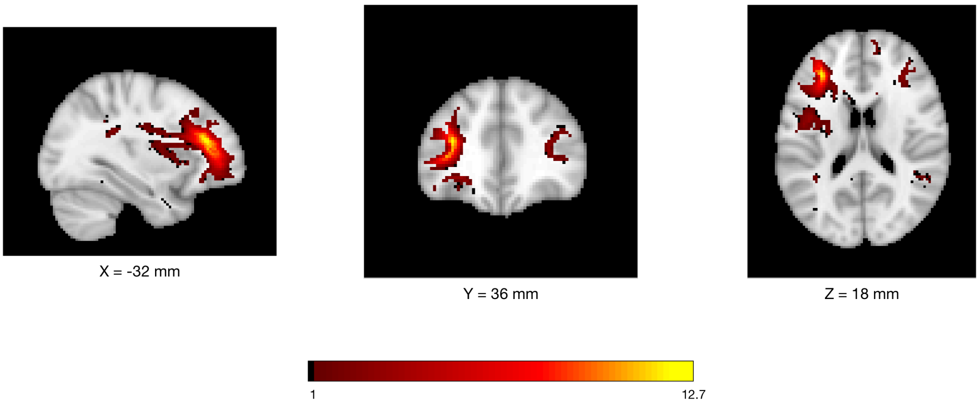


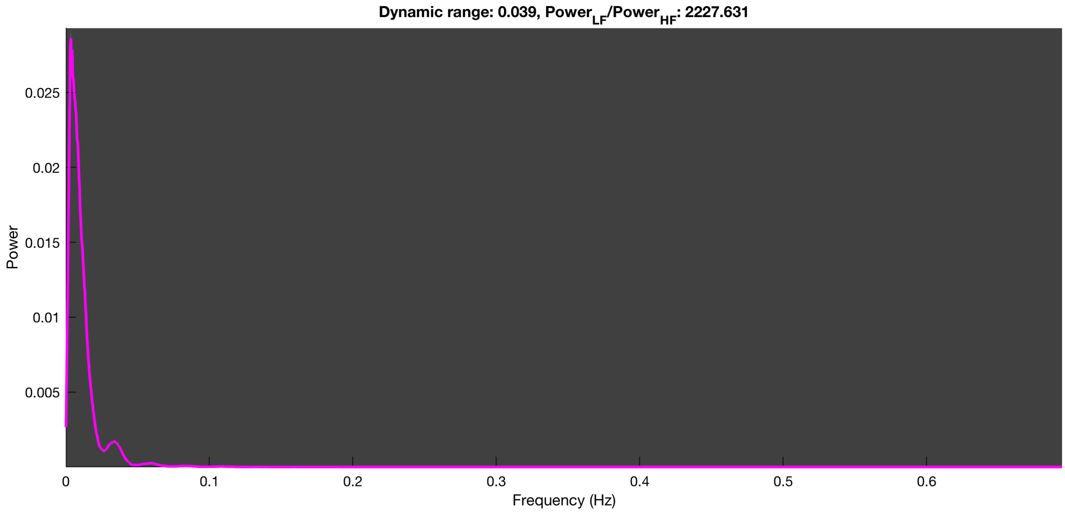

3-1 100n_tmap_component_ica_s1 093

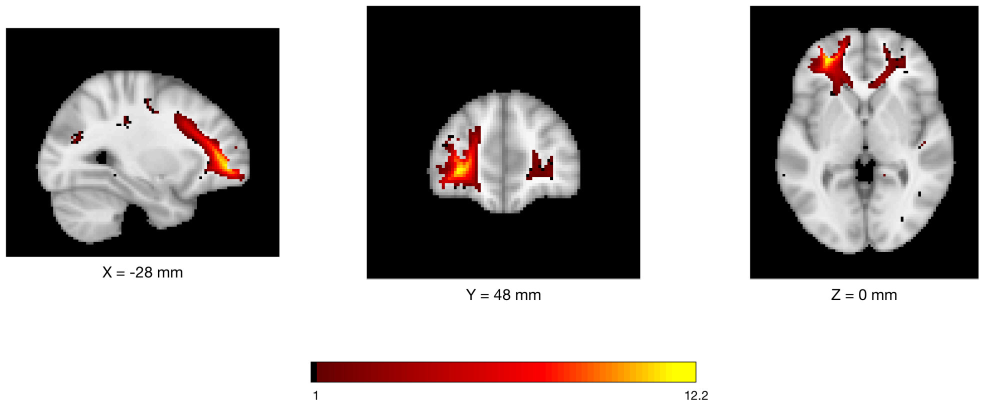


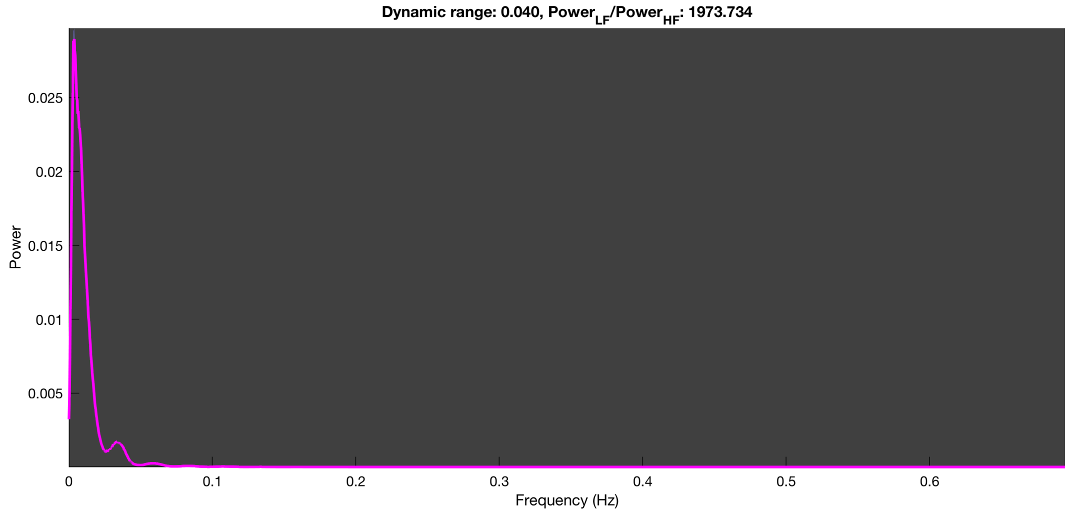

3-1 100n_tmap_component_ica_s1 094

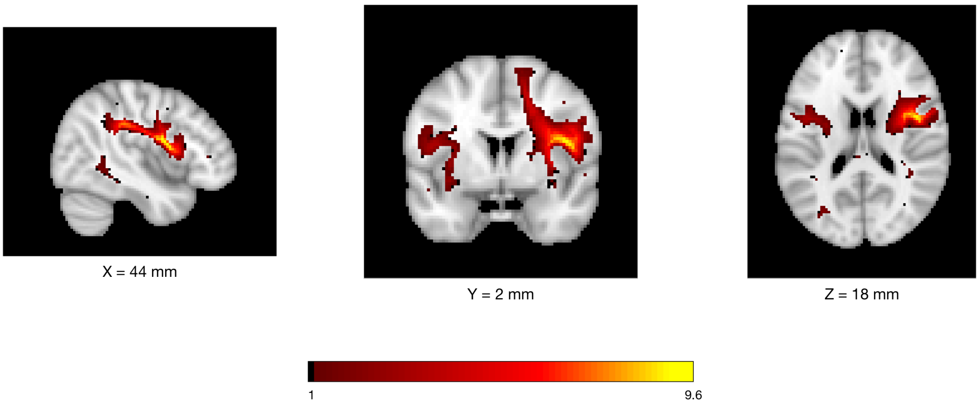


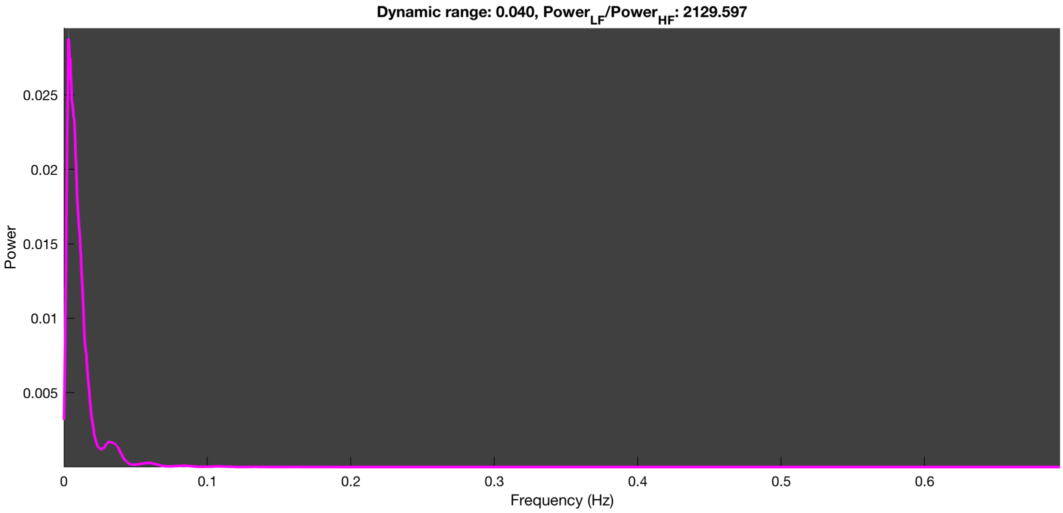

3-1 100n_tmap_component_ica_s1 095

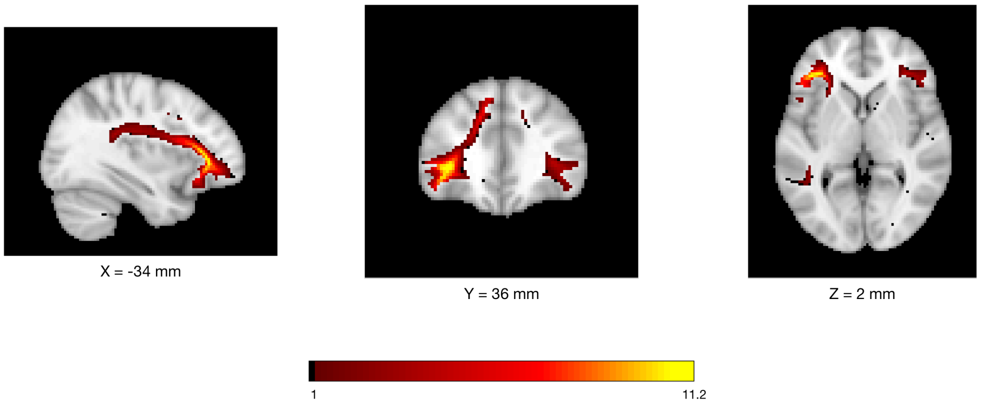


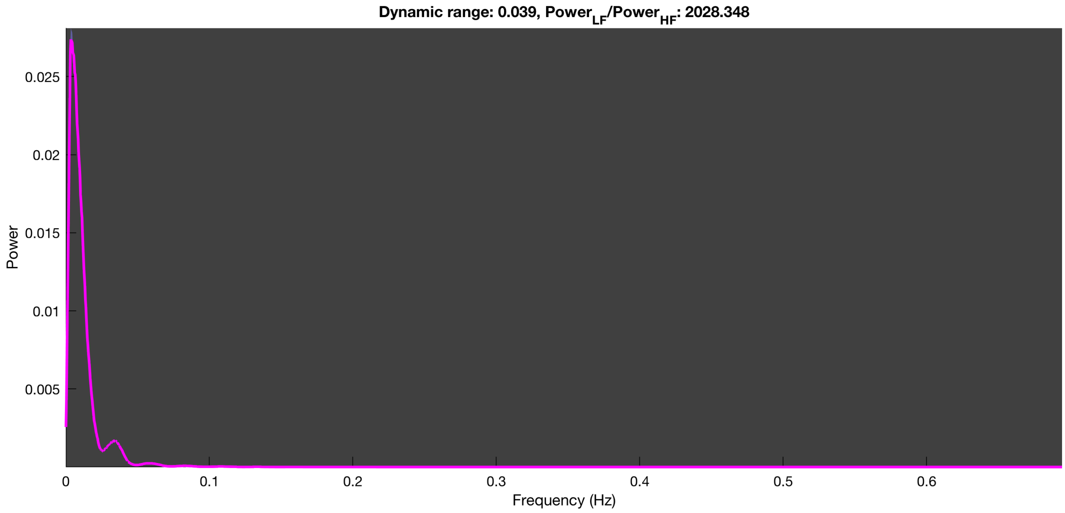

3-1 100n_tmap_component_ica_s1 096

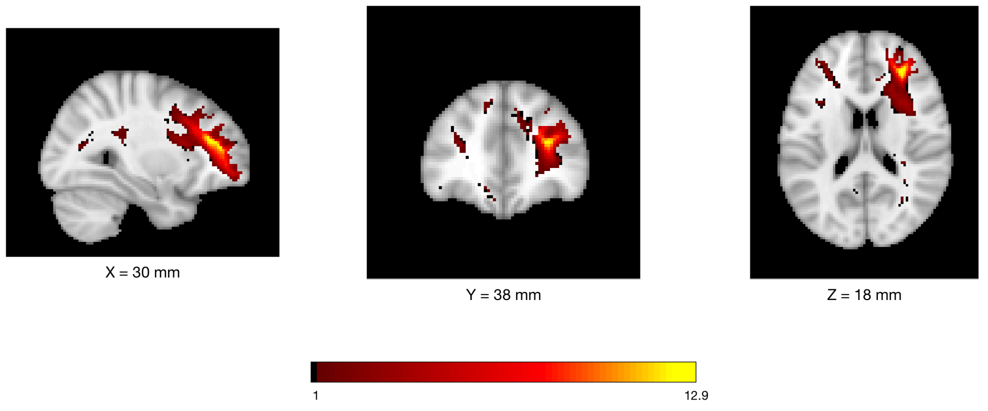


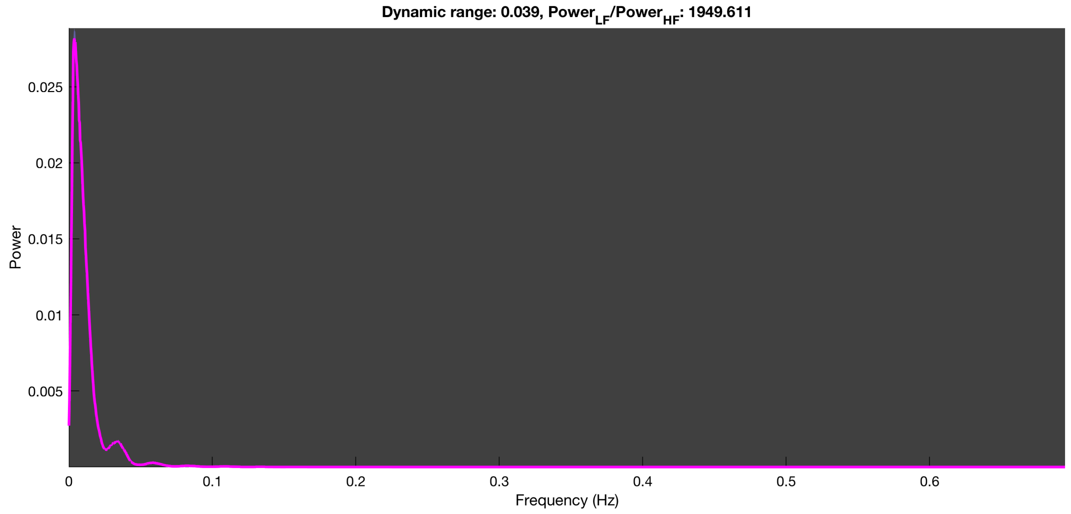

3-1 100n_tmap_component_ica_s1 097

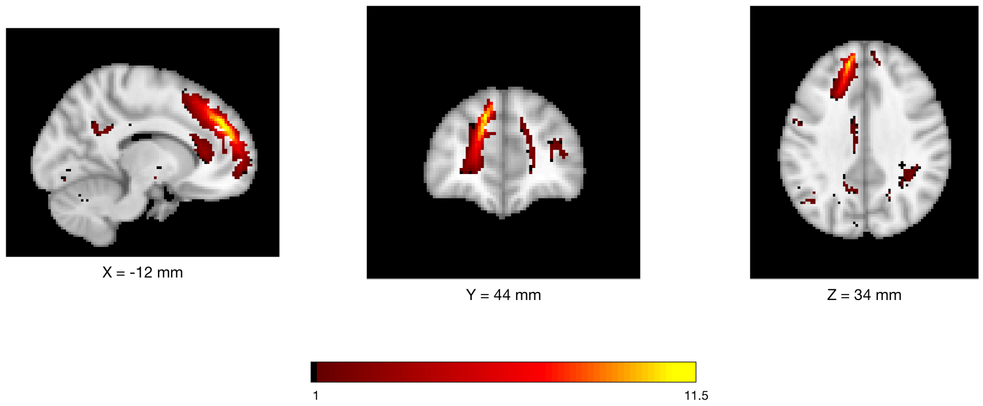


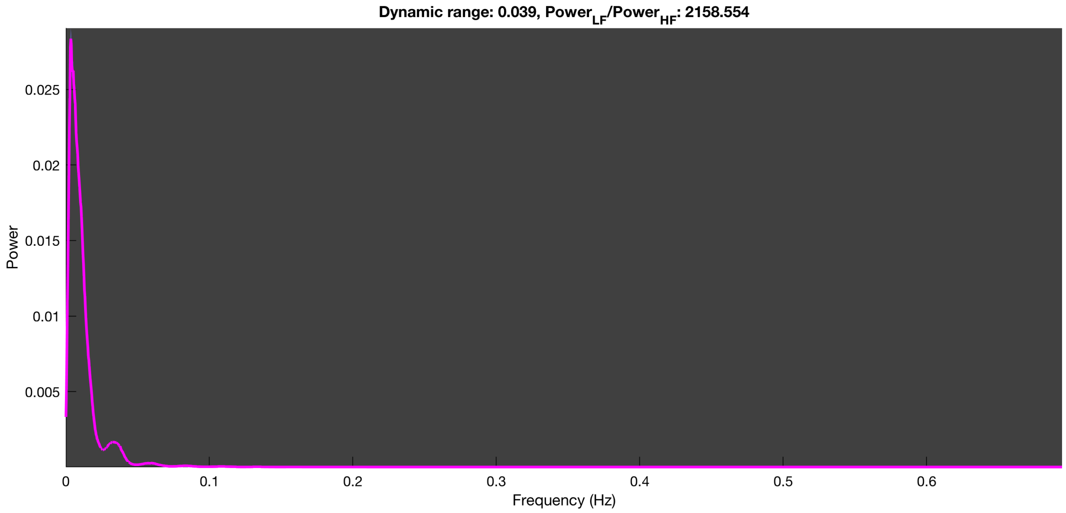

3-1 100n_tmap_component_ica_s1 098

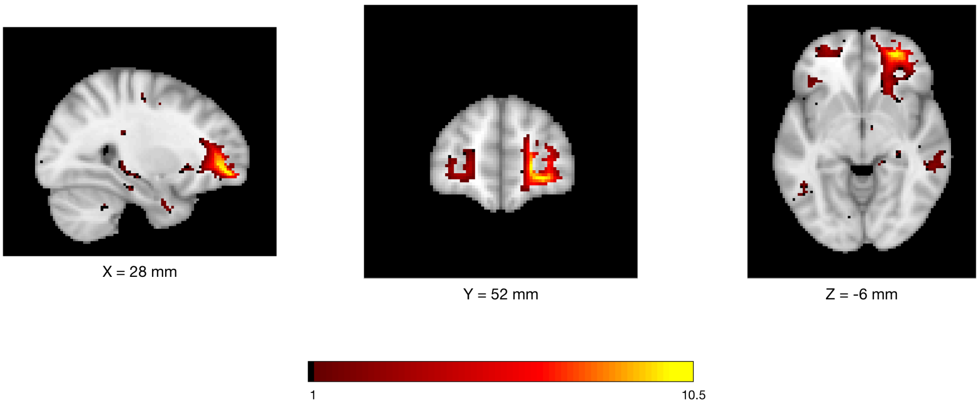


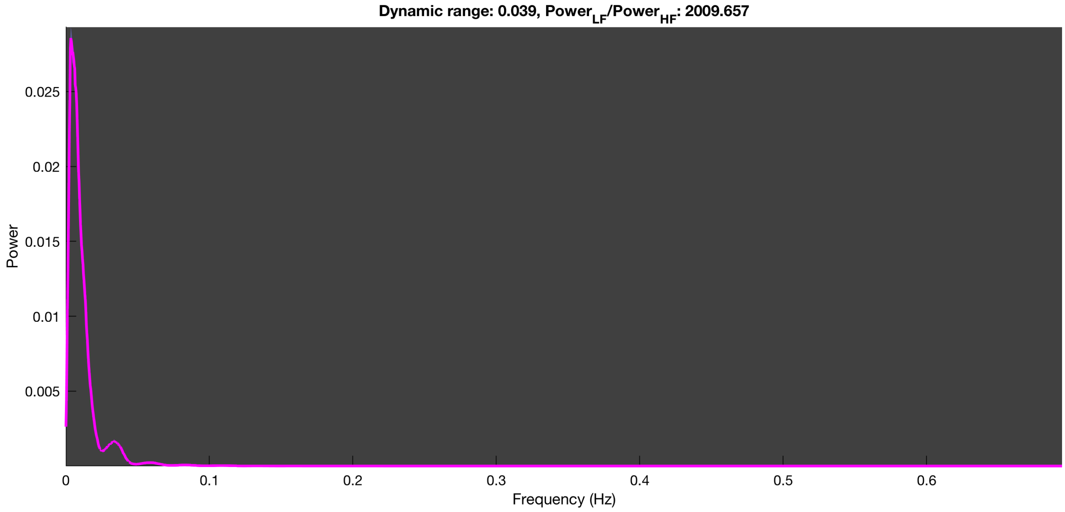

3-1 100n_tmap_component_ica_s1 099

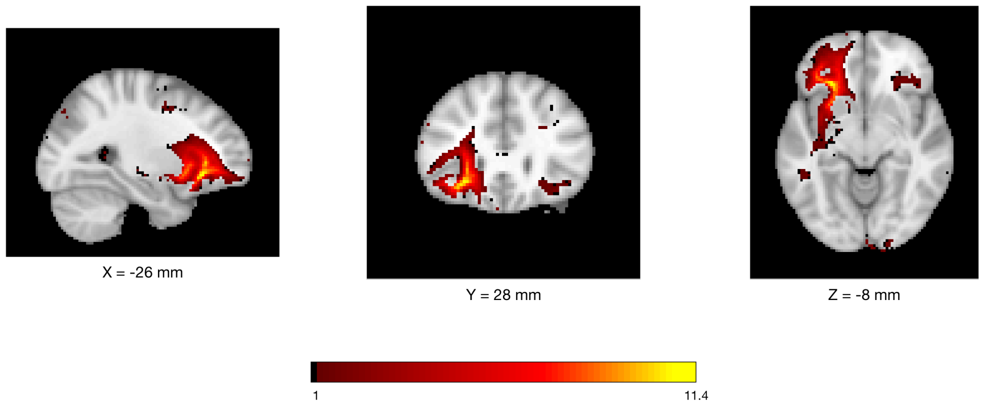


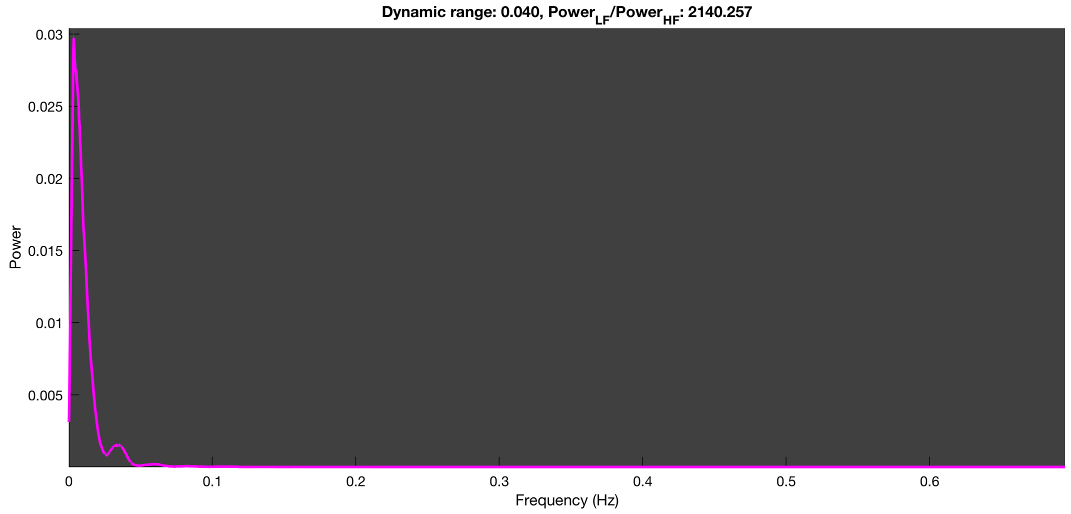

3-1 100n_tmap_component_ica_s1 100

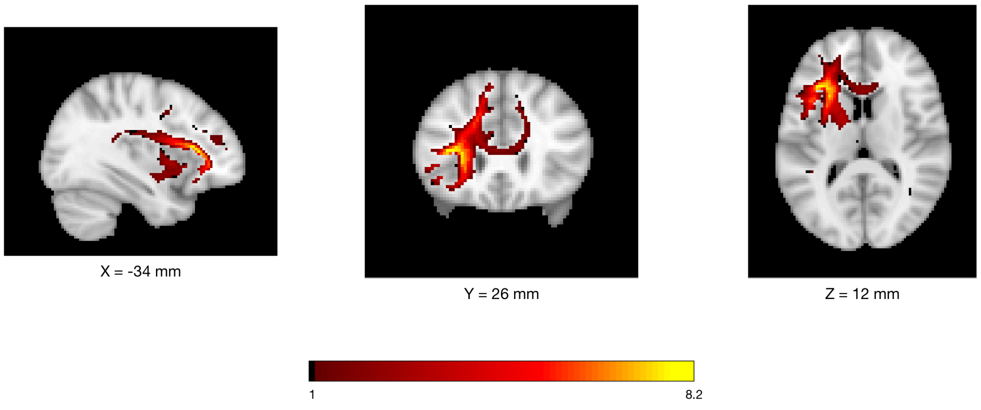


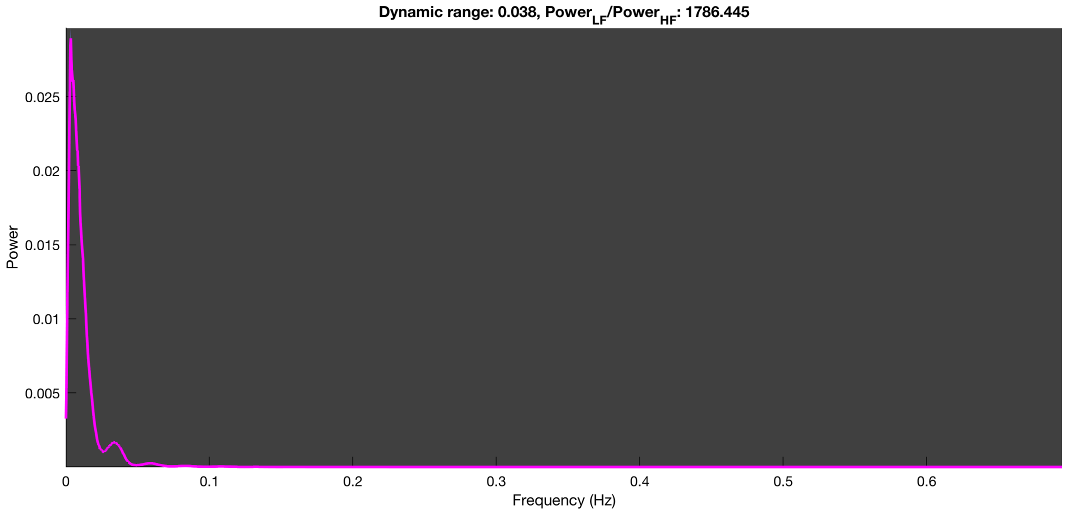

### LEMON dataset

*n=10 run*

3-1 10n_tmap_component_ica_s1 001

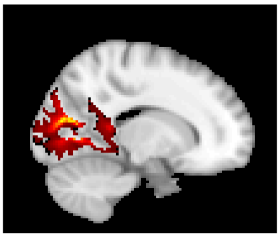

X = 16 mm

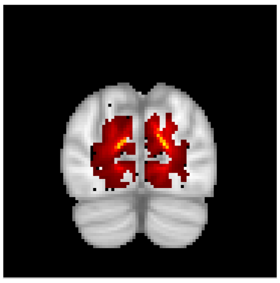

Y = -80 mm

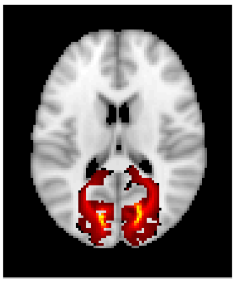

Z = 16 mm

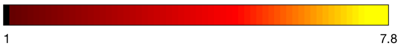


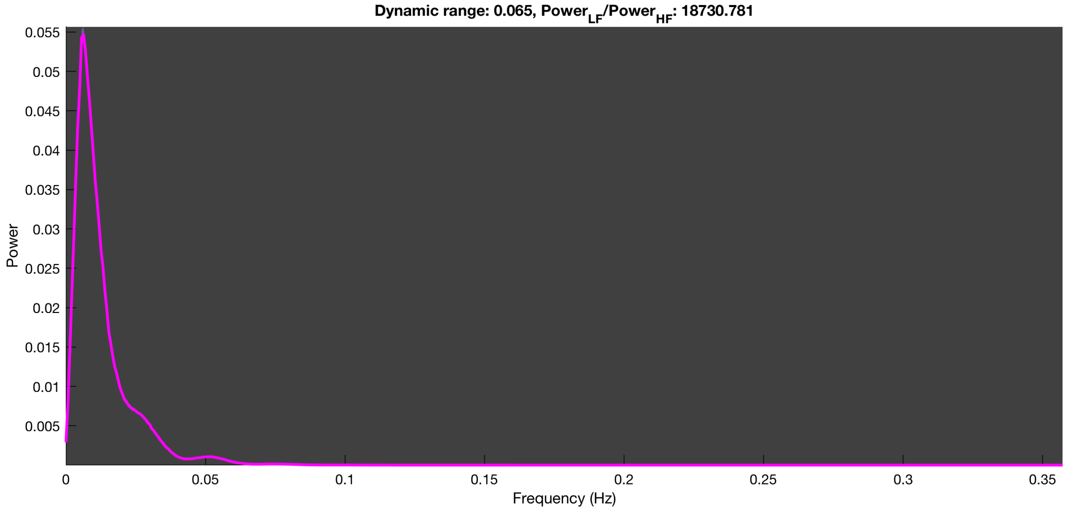

3-1 10n_tmap_component_ica_s1 002

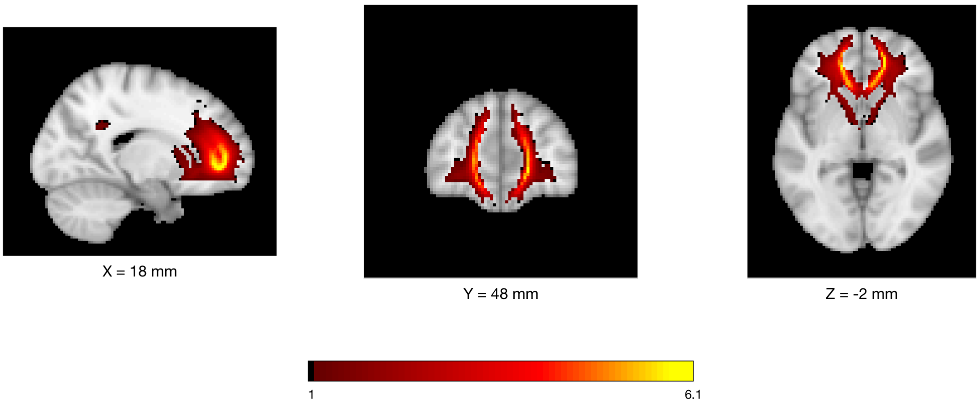


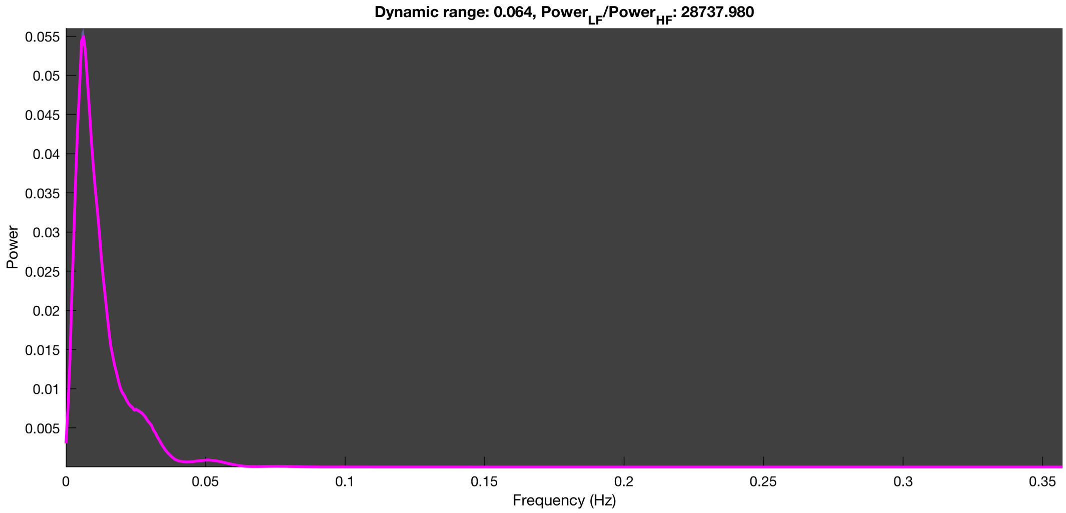

#### 3-1 10n_tmap_component_ica_s1 003

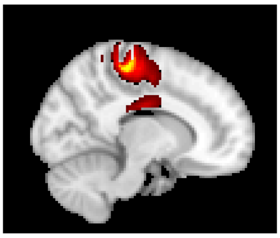

X = 12 mm

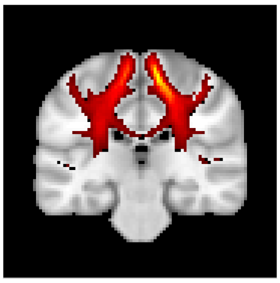

Y = -26 mm

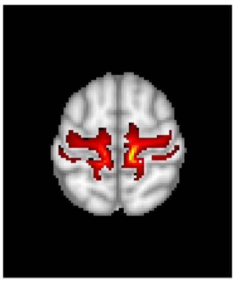

Z = 60 mm

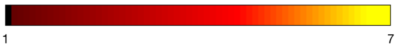


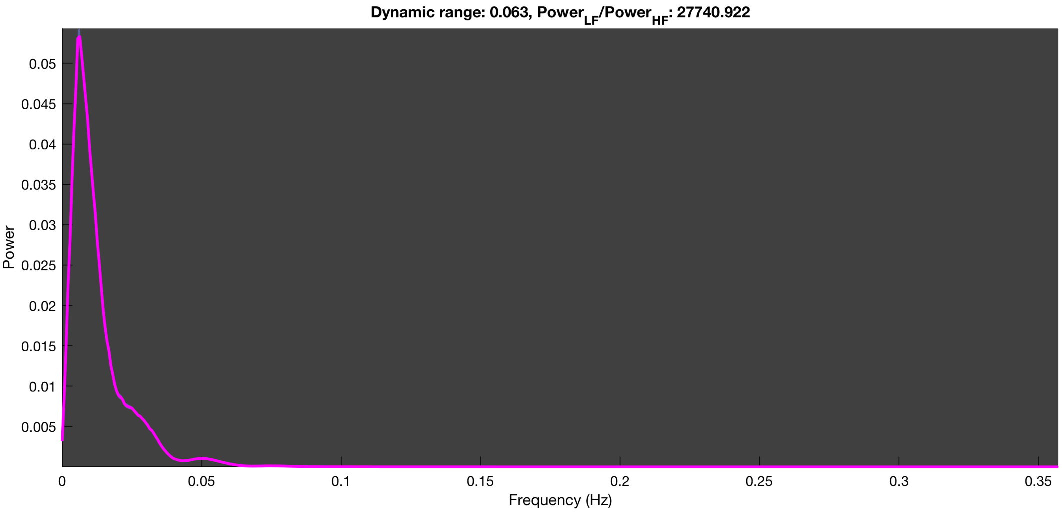

3-1 10n_tmap_component_ica_s1 004

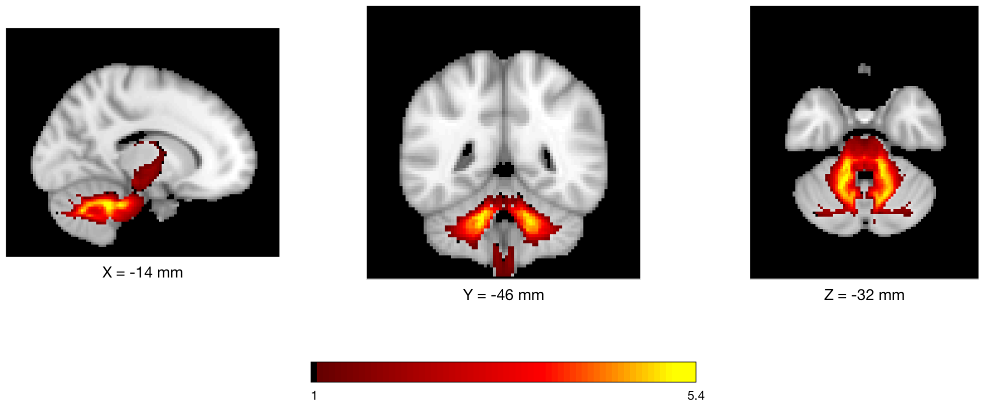


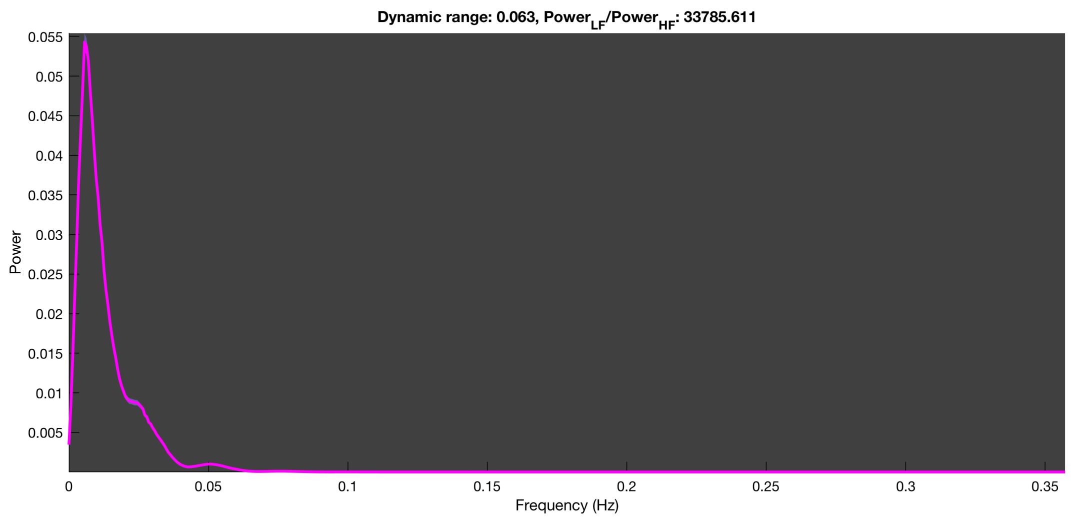

3-1 10n_tmap_component_ica_s1 005

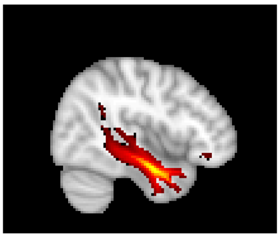

X = 44 mm

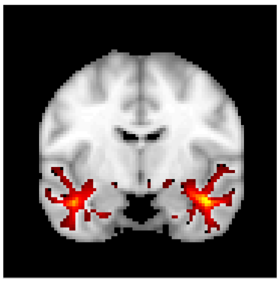

Y = -8 mm

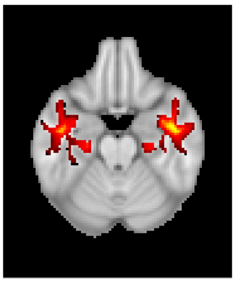

Z = -22 mm

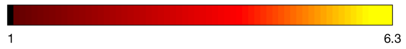


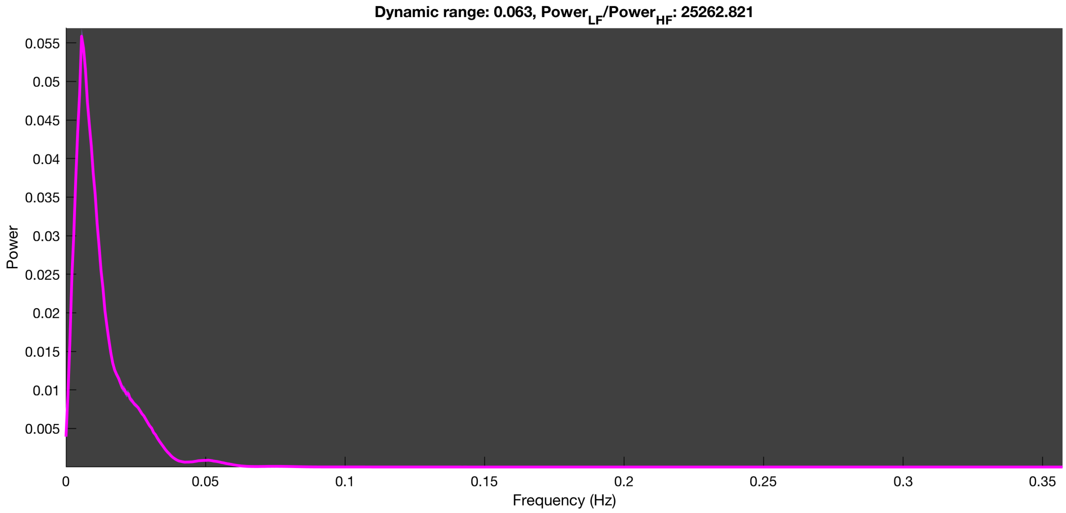

3-1 10n_tmap_component_ica_s1 006

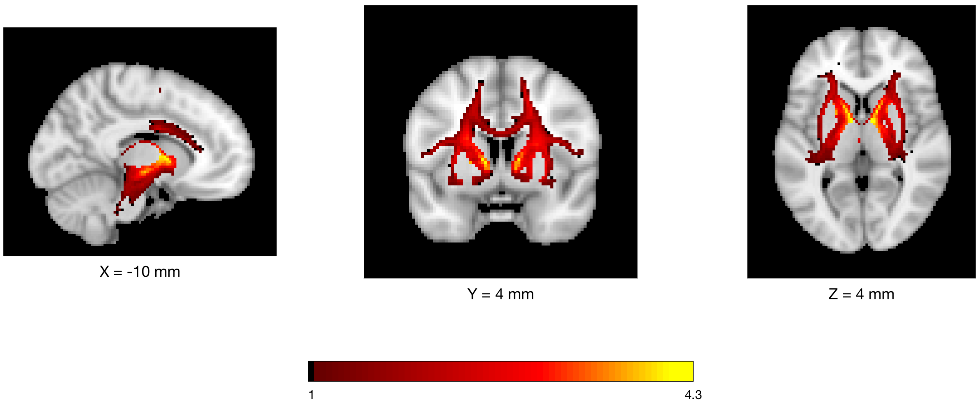


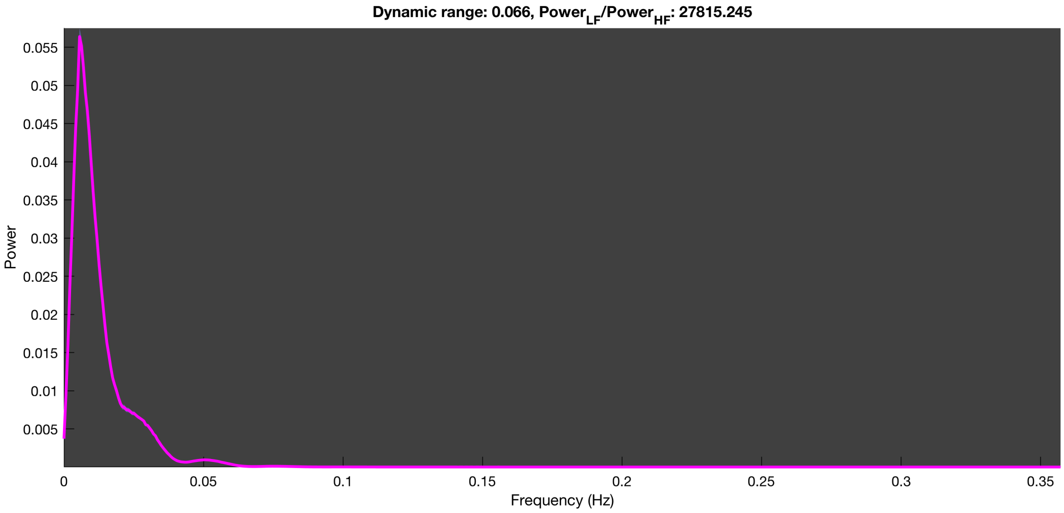

3-1 10n_tmap_component_ica_s1 007

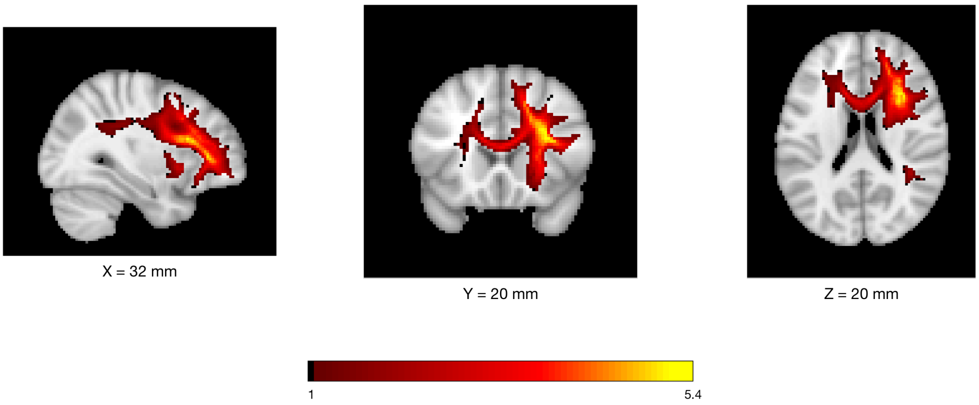


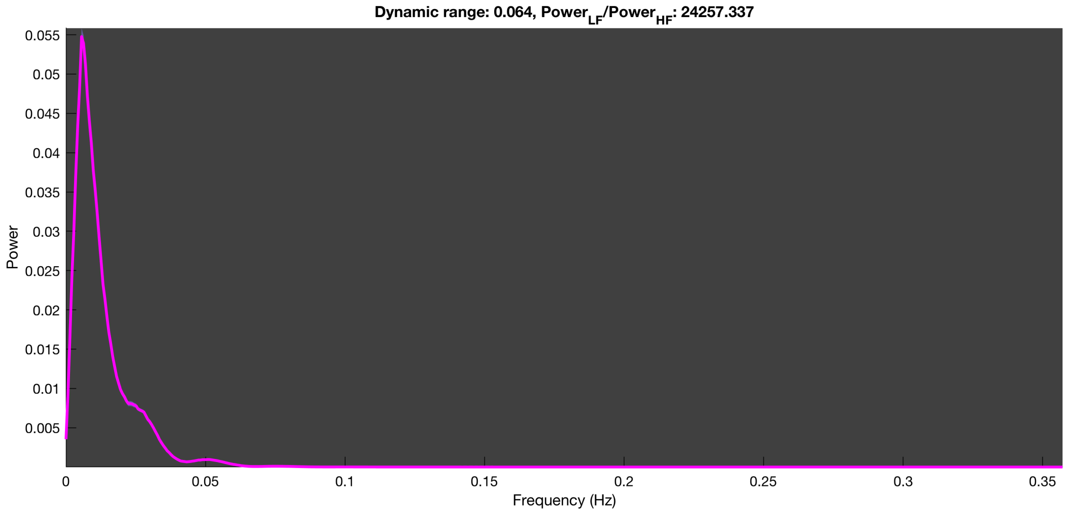

3-1 10n_tmap_component_ica_s1 008

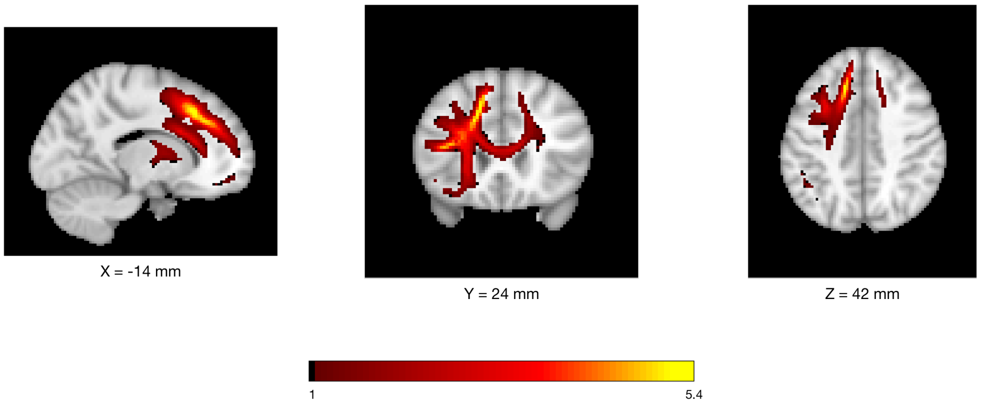


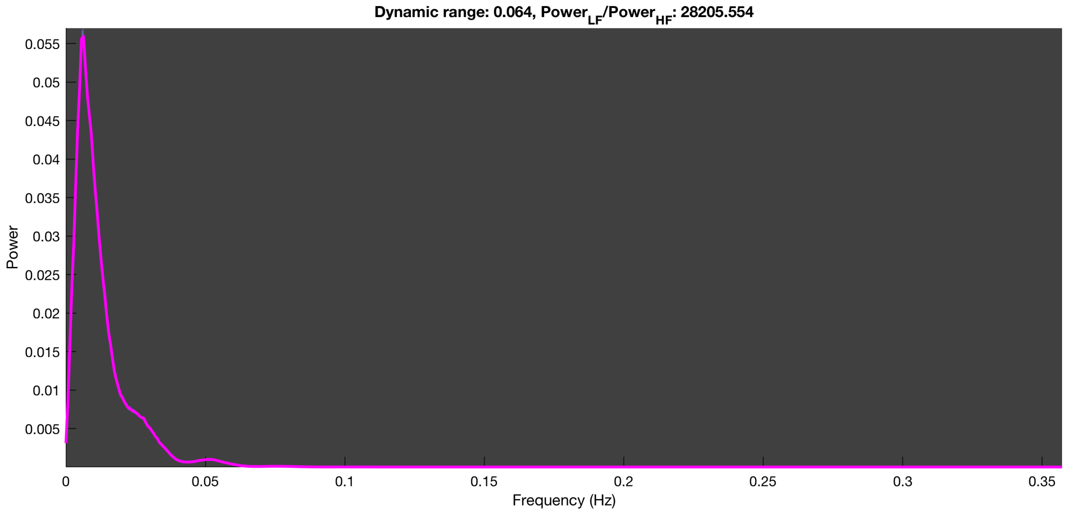

3-1 10n_tmap_component_ica_s1 009

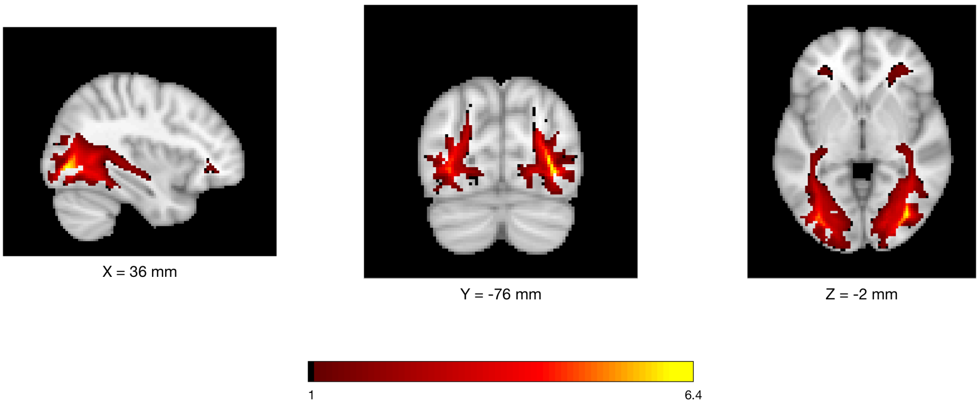


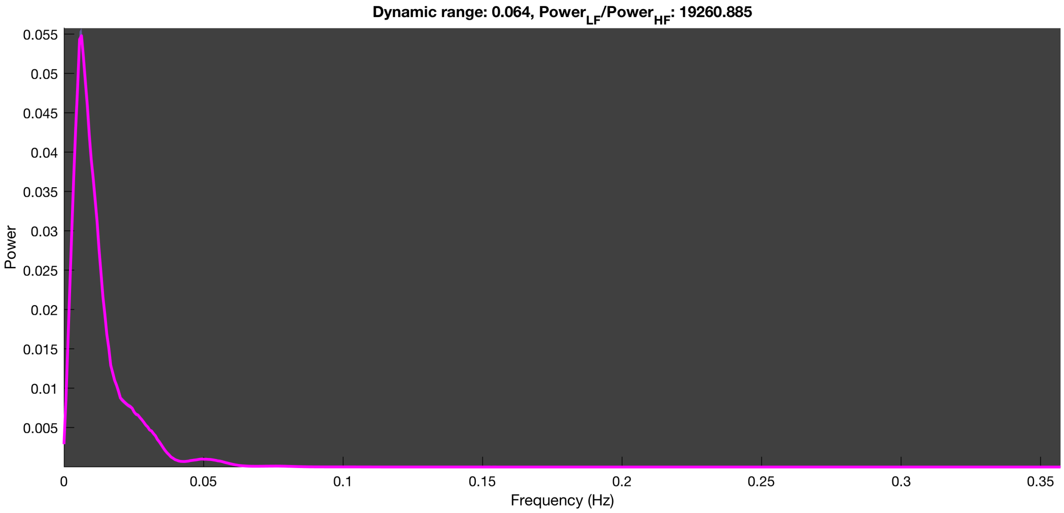

3-1 10n_tmap_component_ica_s1 010

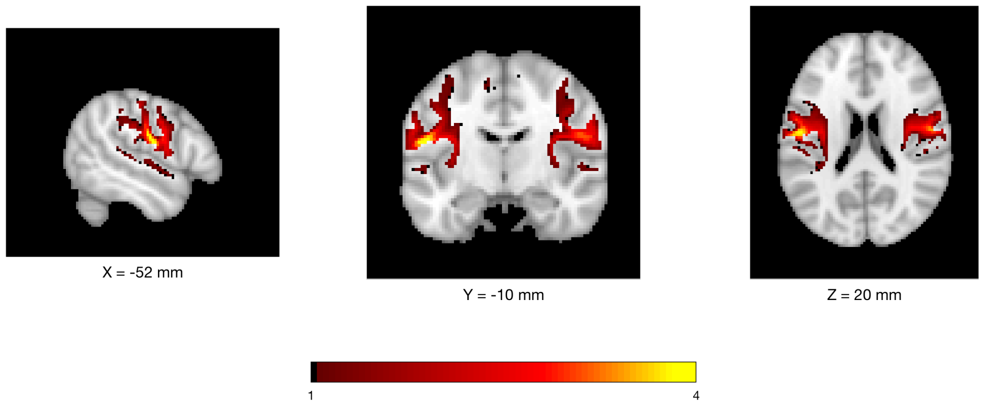


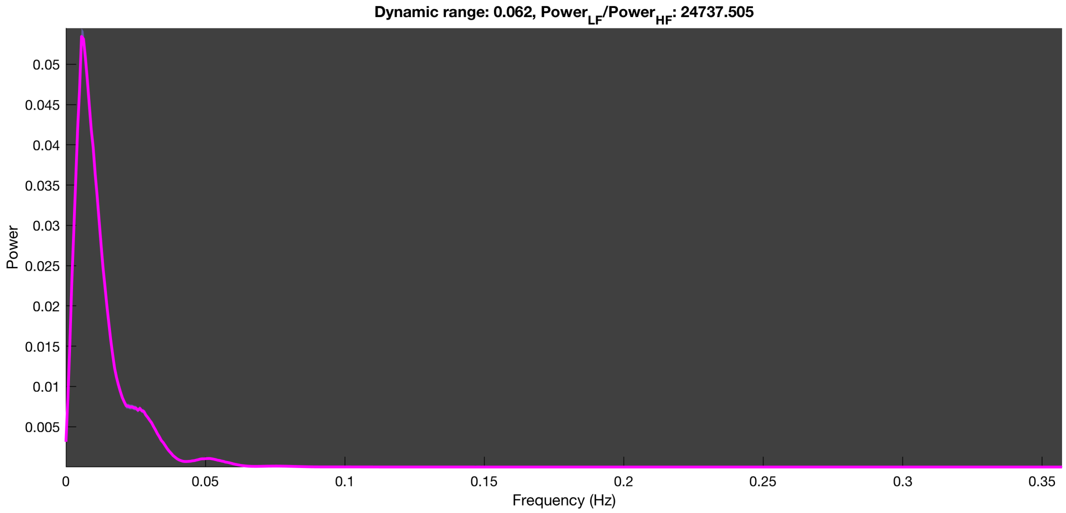

### LEMON dataset

*n=20 run*

3-1 20n_tmap_component_ica_s1 001

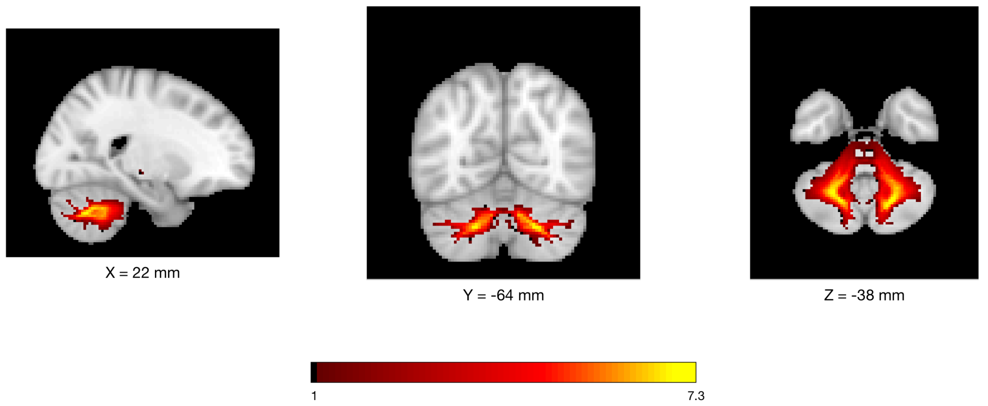


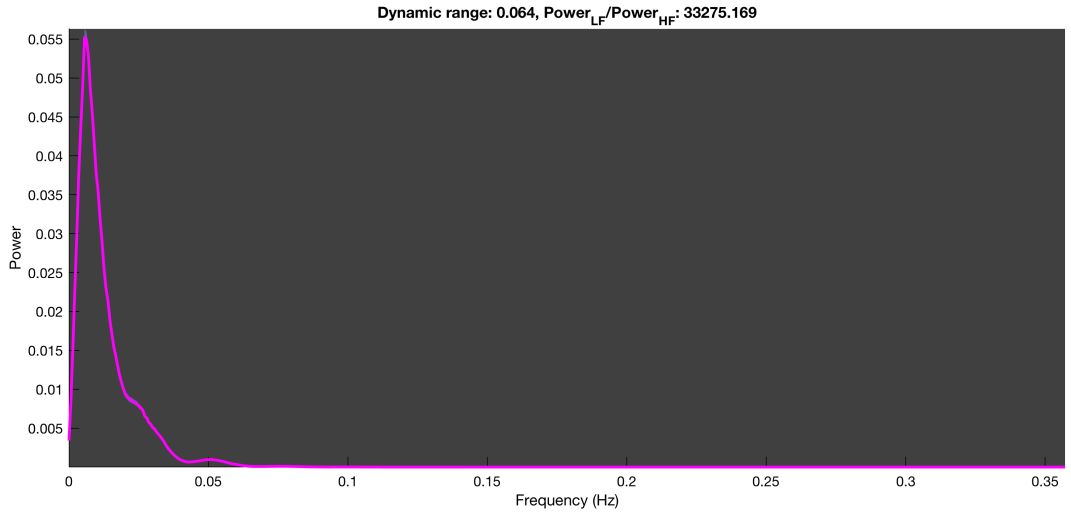

3-1 20n_tmap_component_ica_s1 002

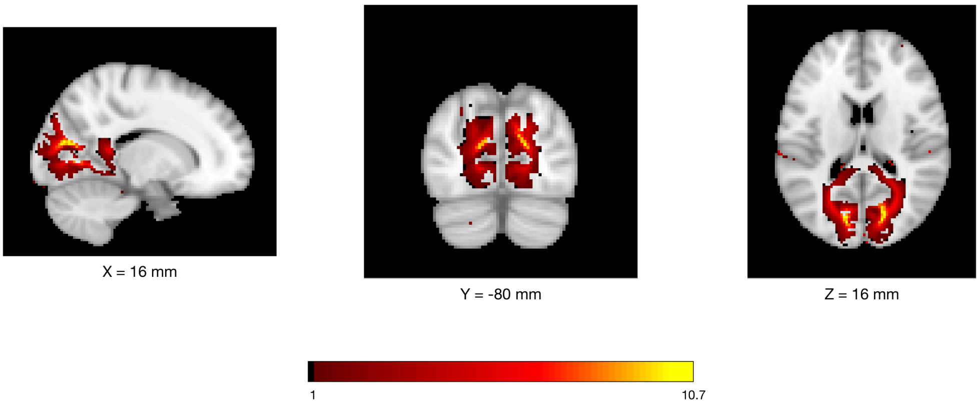


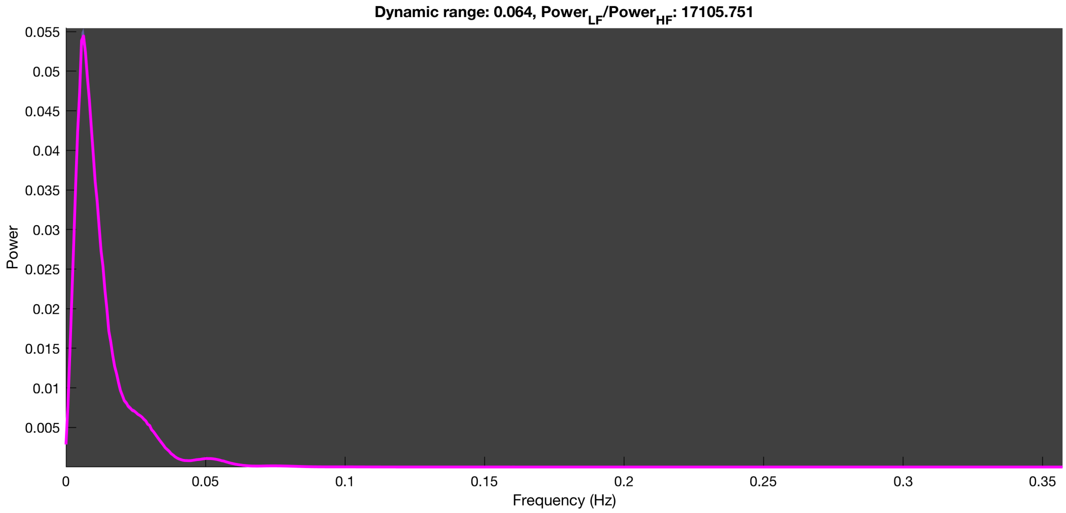

#### 3-1 20n_tmap_component_ica_s1 003

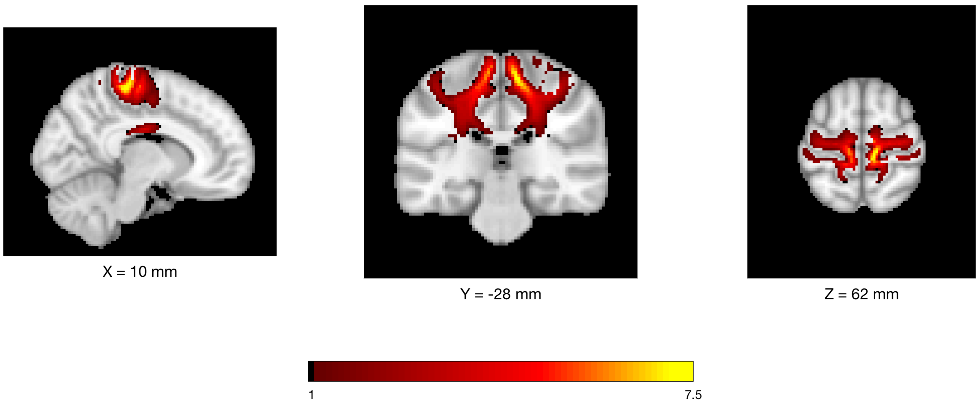


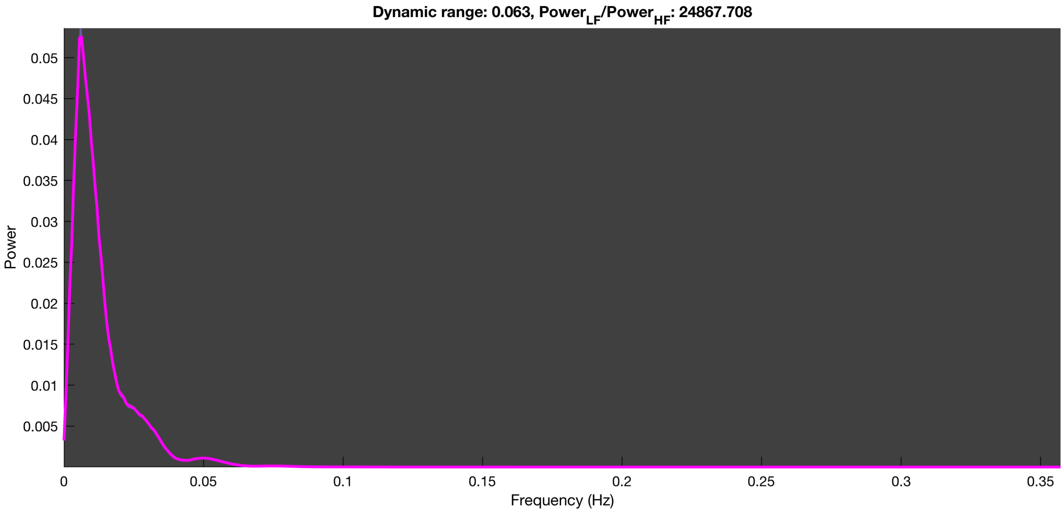

3-1 20n_tmap_component_ica_s1 004

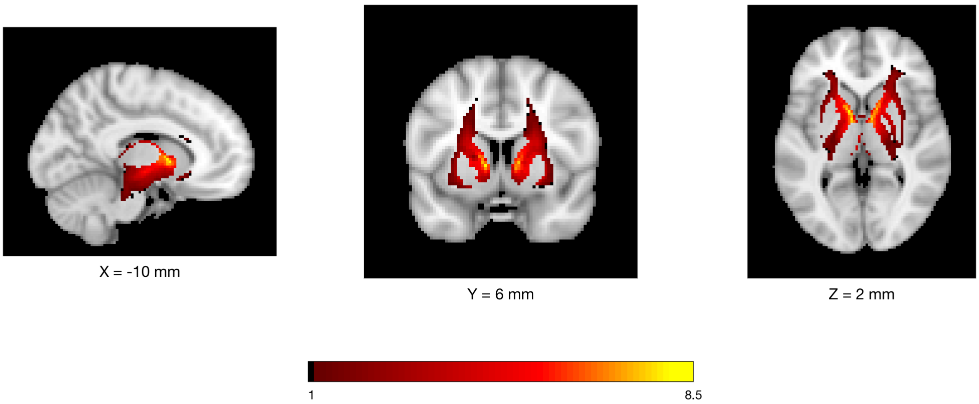


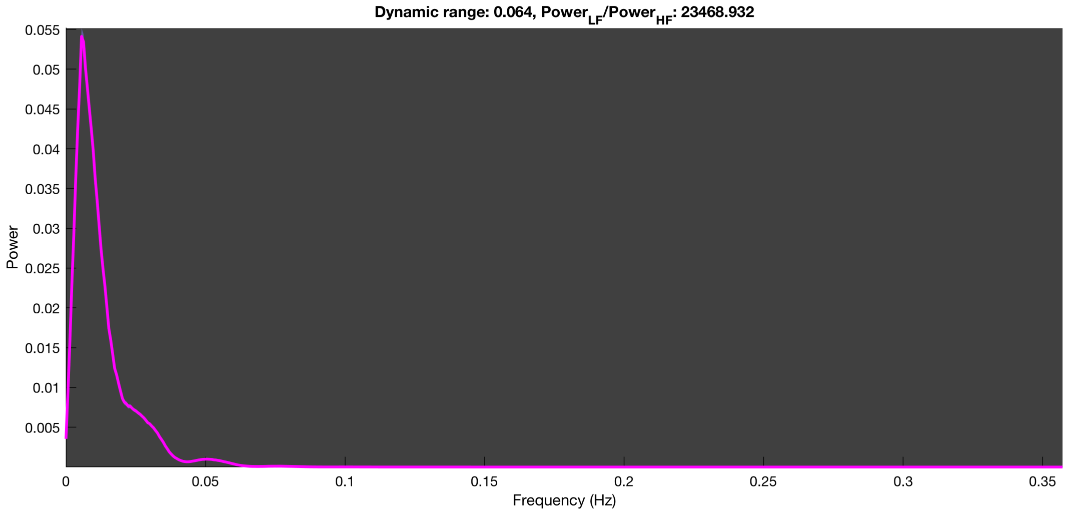

3-1 20n_tmap_component_ica_s1 005

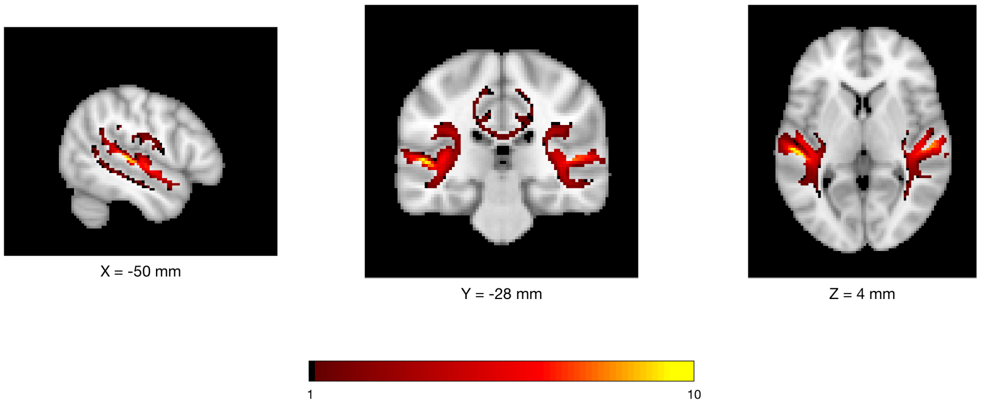


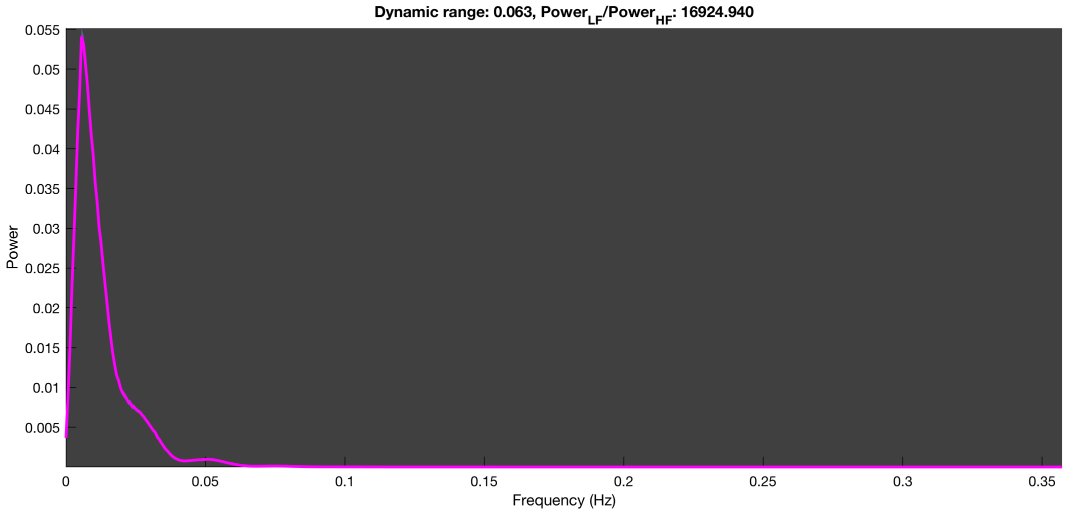

3-1 20n_tmap_component_ica_s1 006

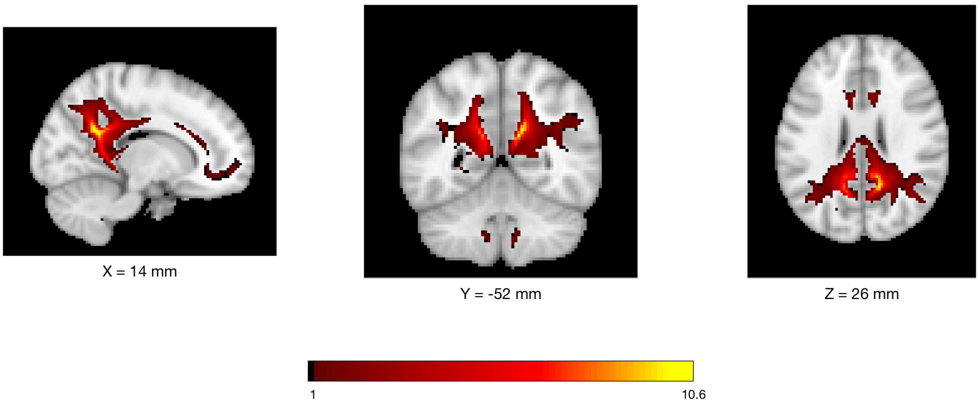


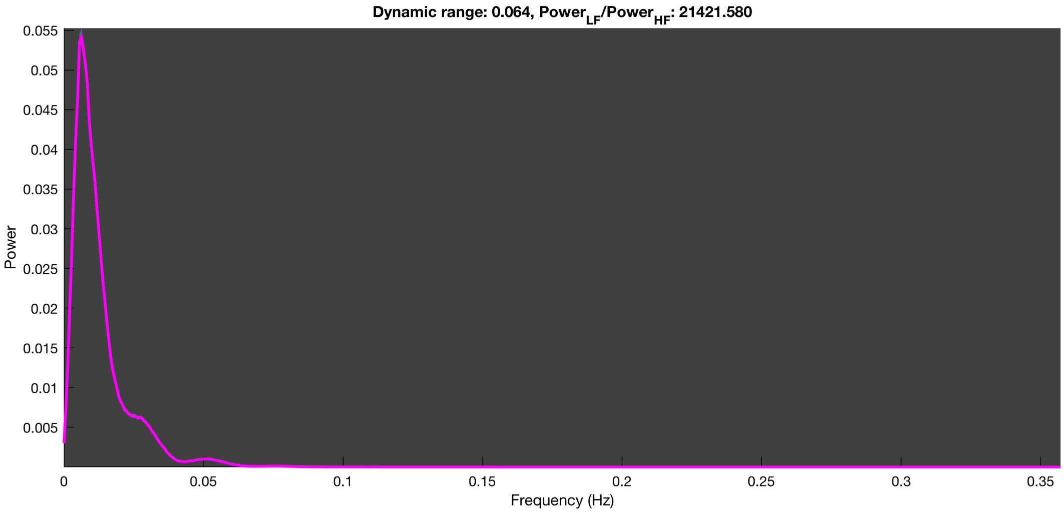

3-1 20n_tmap_component_ica_s1 007

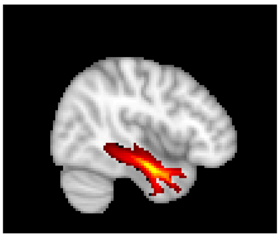

X = 44 mm

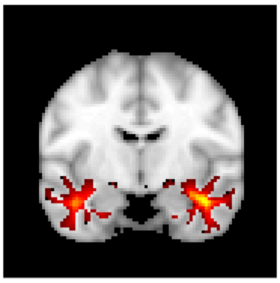

Y = -8 mm

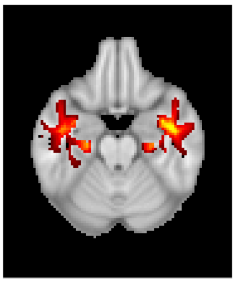

Z = -22 mm

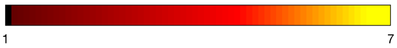


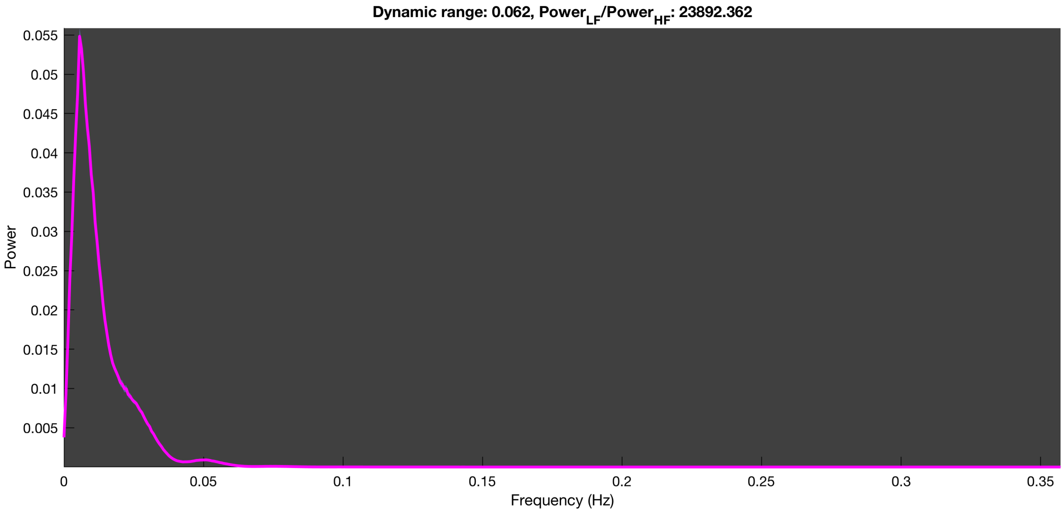

#### 3-1 20n_tmap_component_ica_s1 008

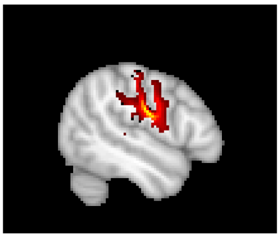

X = -50 mm

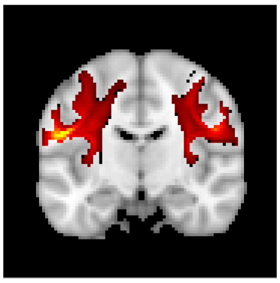

Y = -12 mm

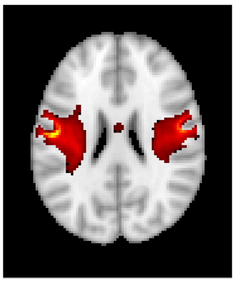

Z = 24 mm

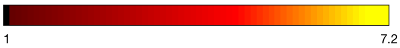


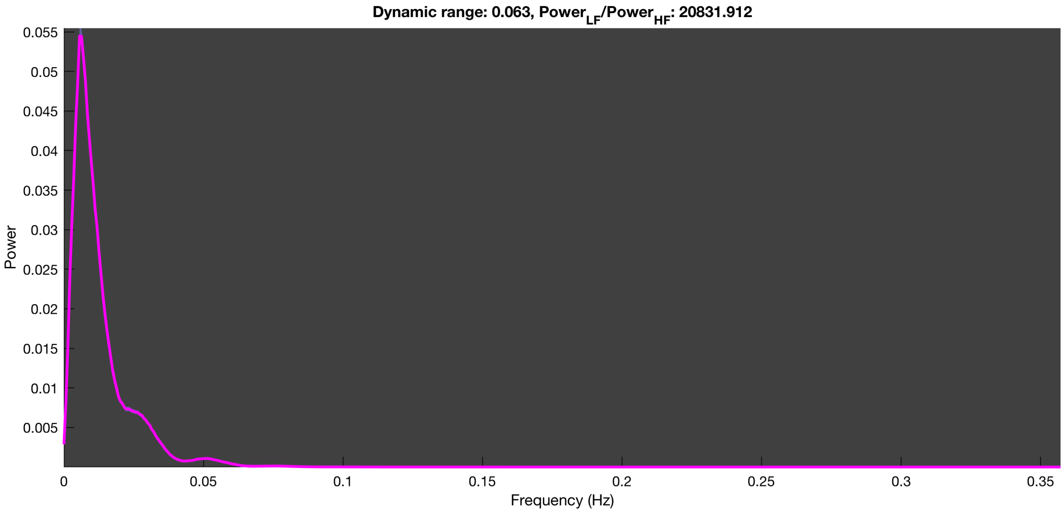

### 3-1 20n_tmap_component_ica_s1 009

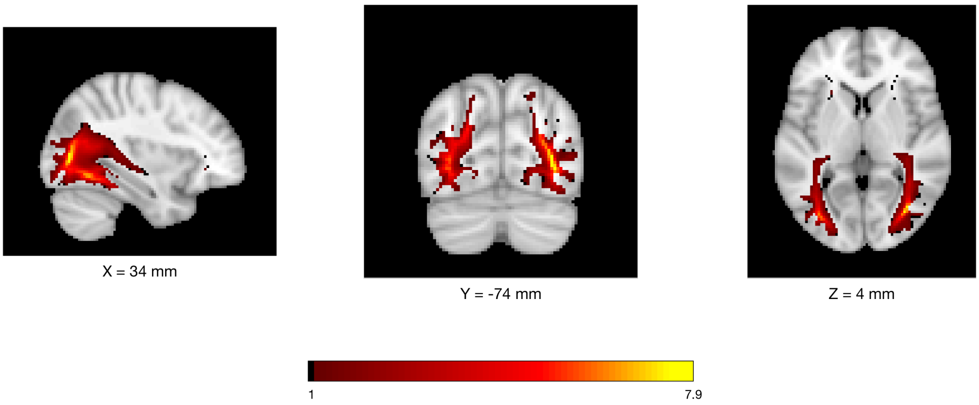


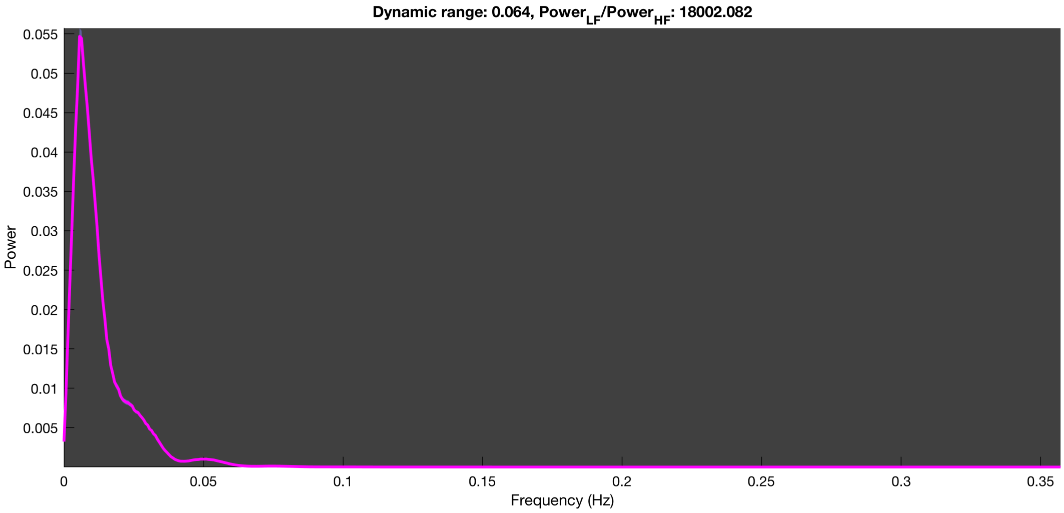

3-1 20n_tmap_component_ica_s1 010

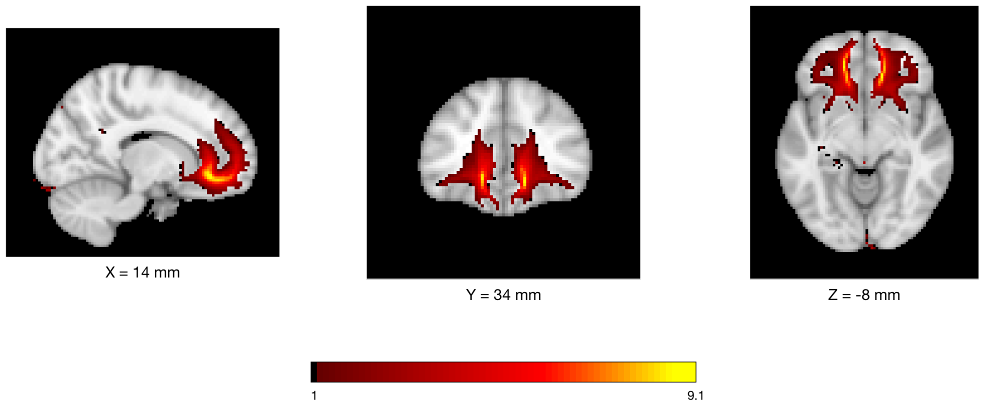


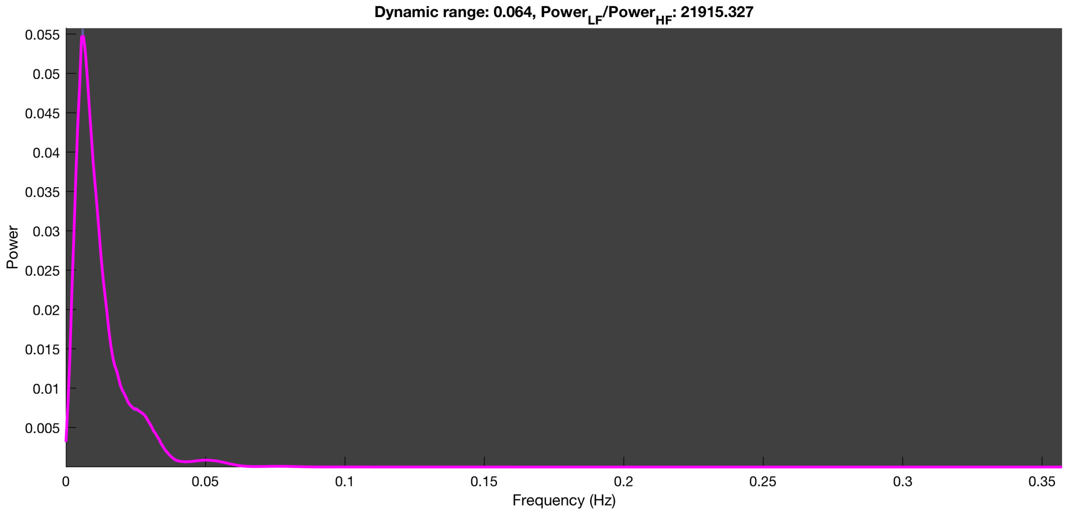

3-1 20n_tmap_component_ica_s1 011

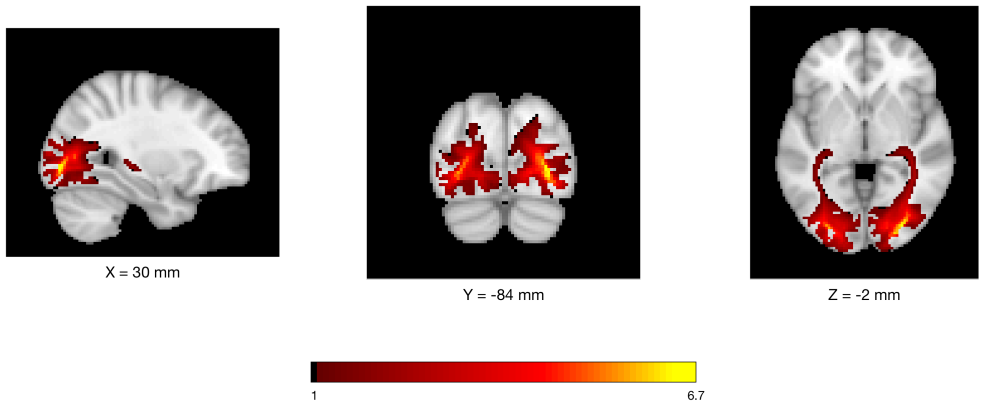


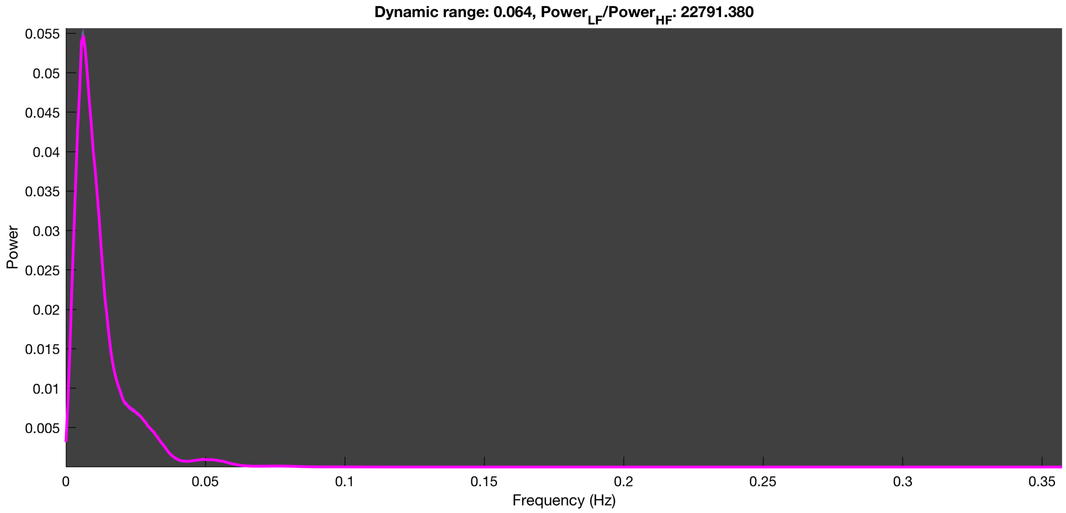

3-1 20n_tmap_component_ica_s1 012

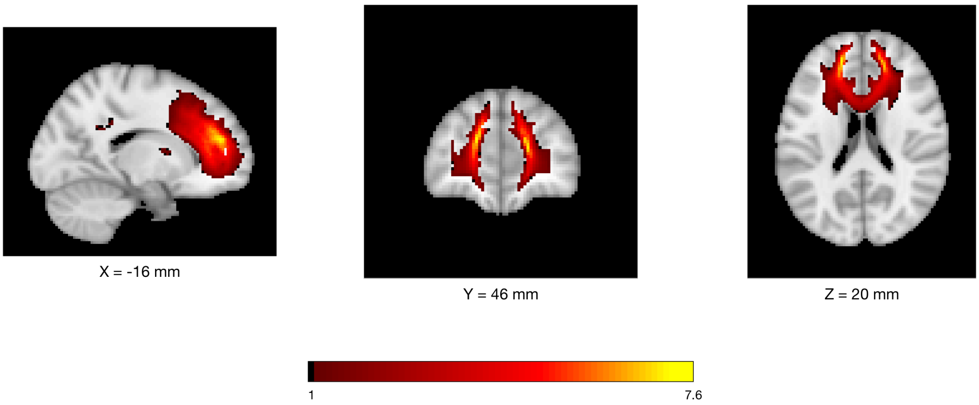


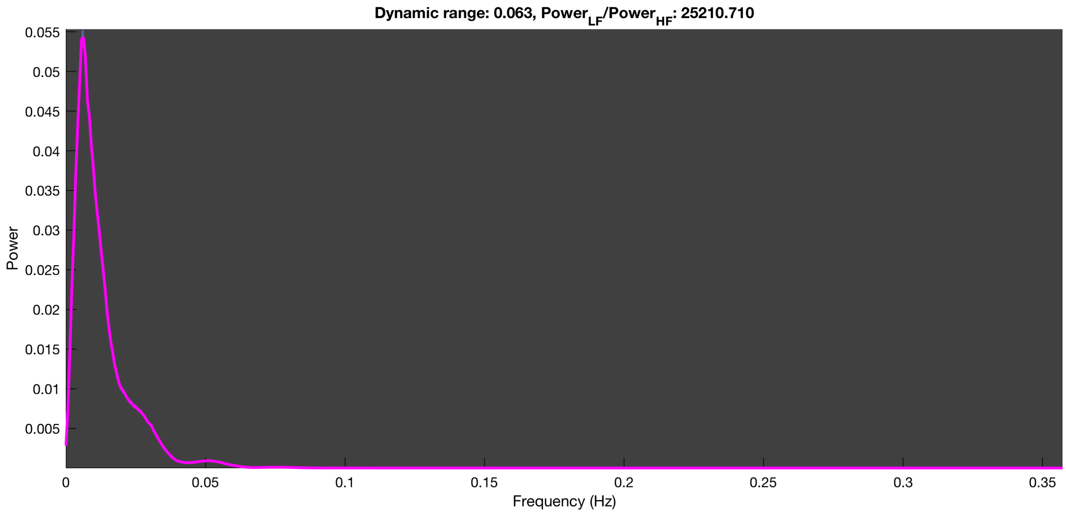

3-1 20n_tmap_component_ica_s1 013

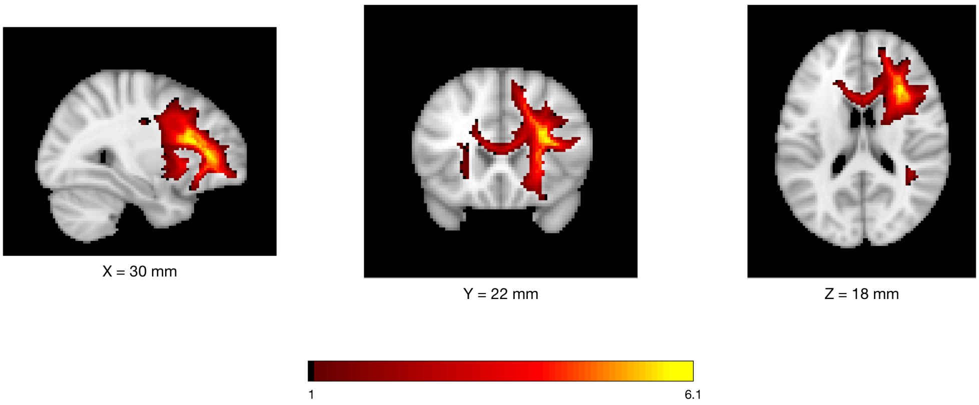


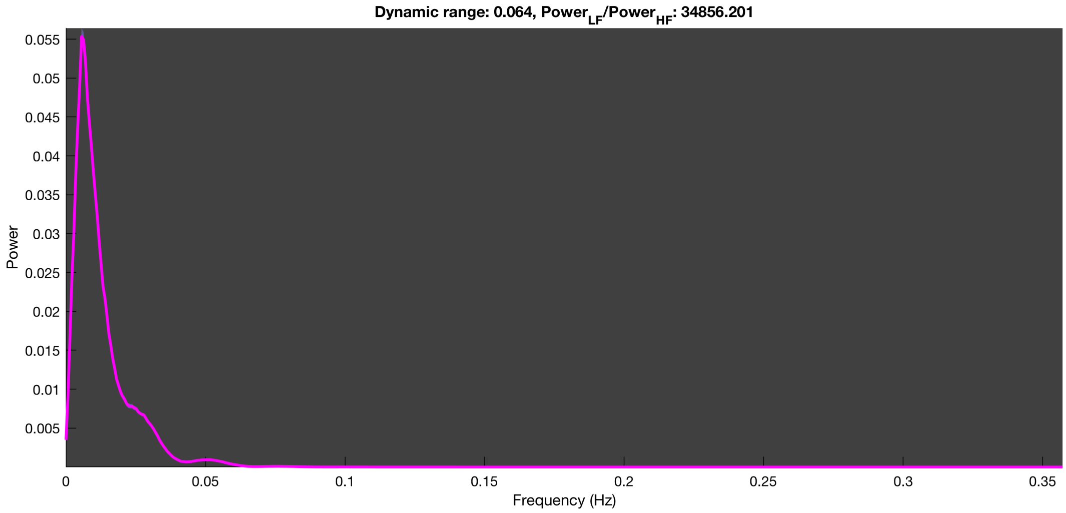

3-1 20n_tmap_component_ica_s1 014

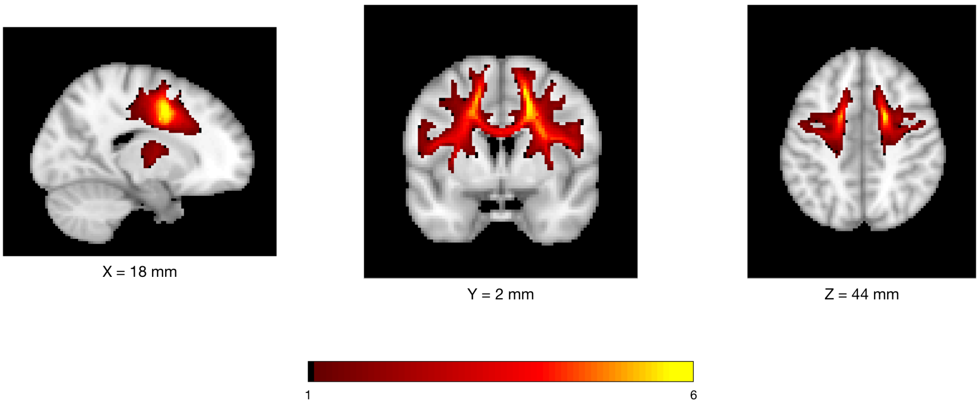


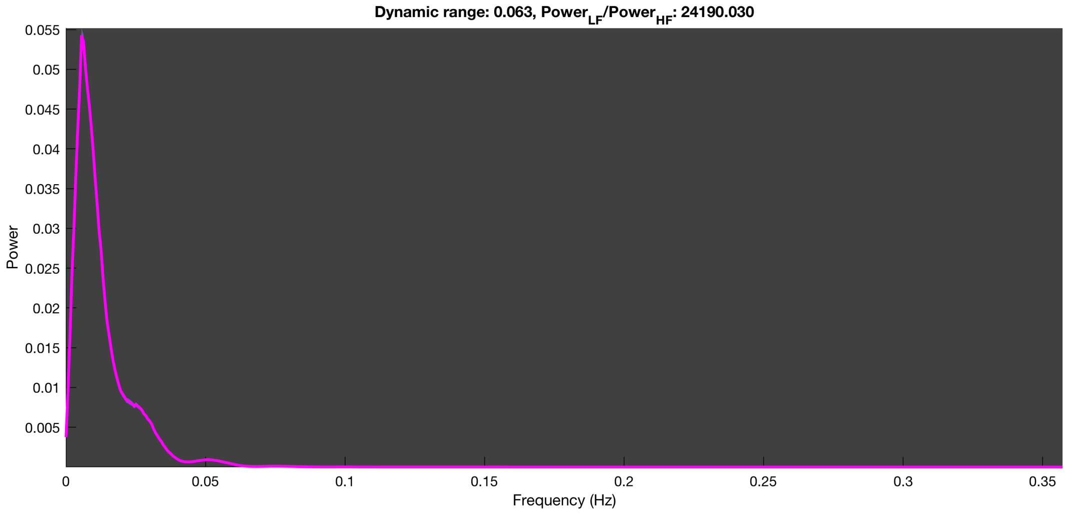

3-1 20n_tmap_component_ica_s1 015

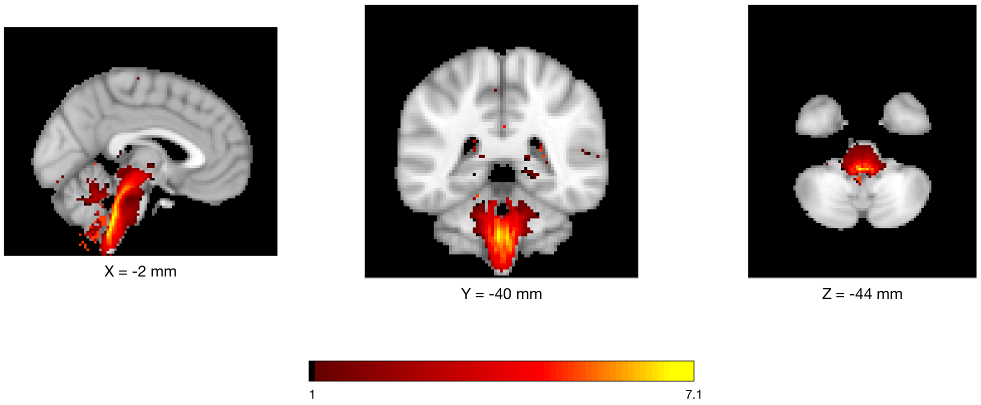


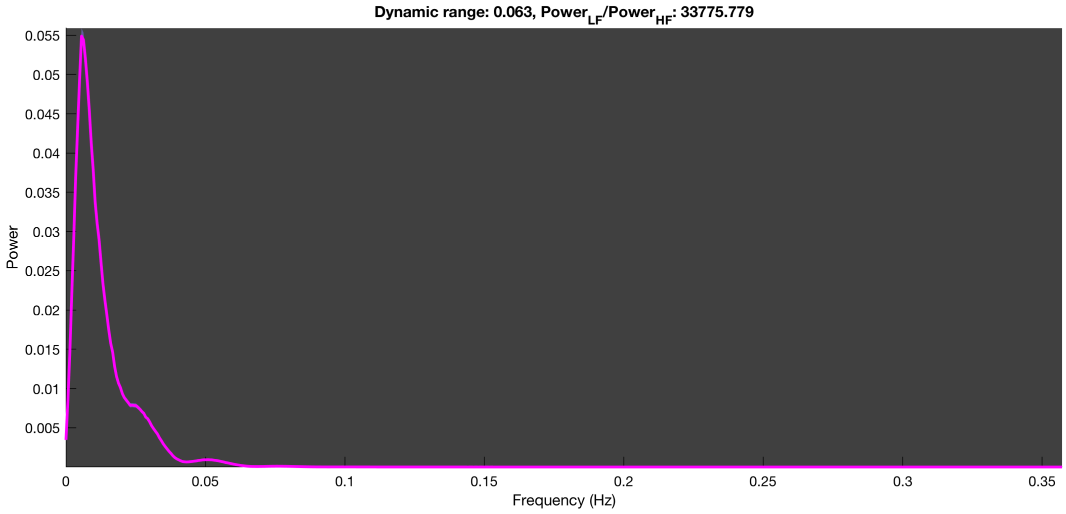

#### 3-1 20n_tmap_component_ica_s1 016

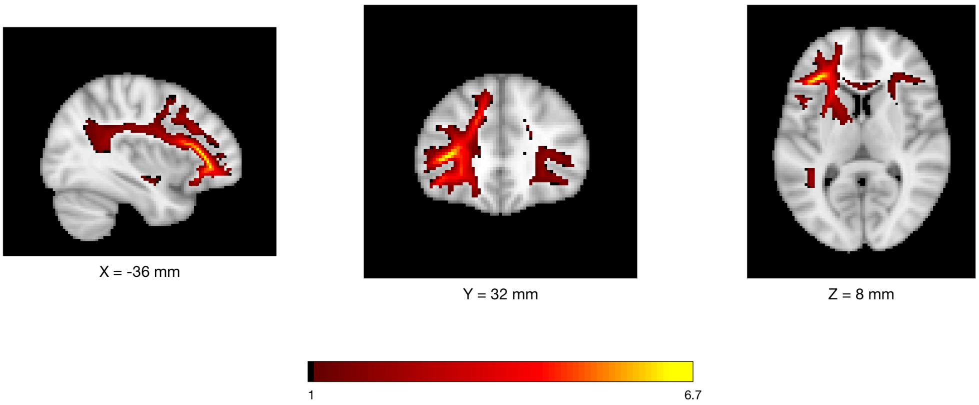


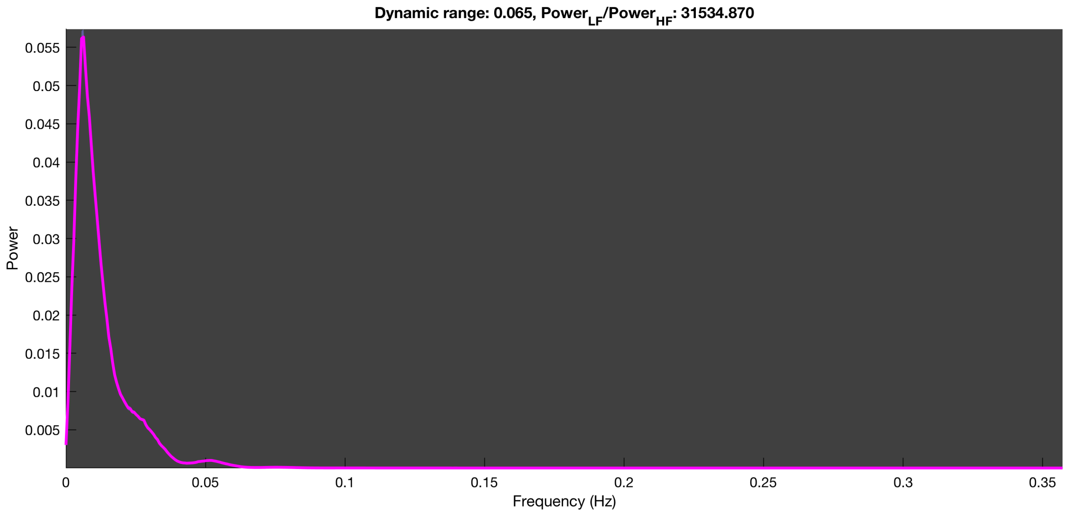

#### 3-1 20n_tmap_component_ica_s1 017

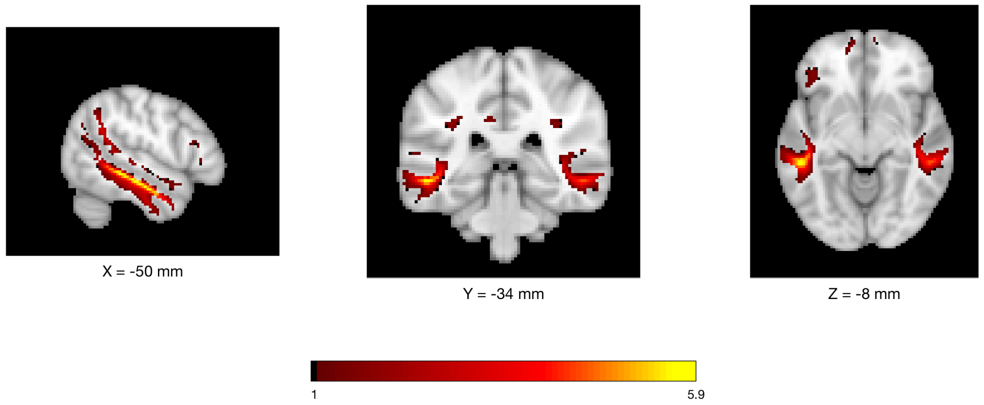


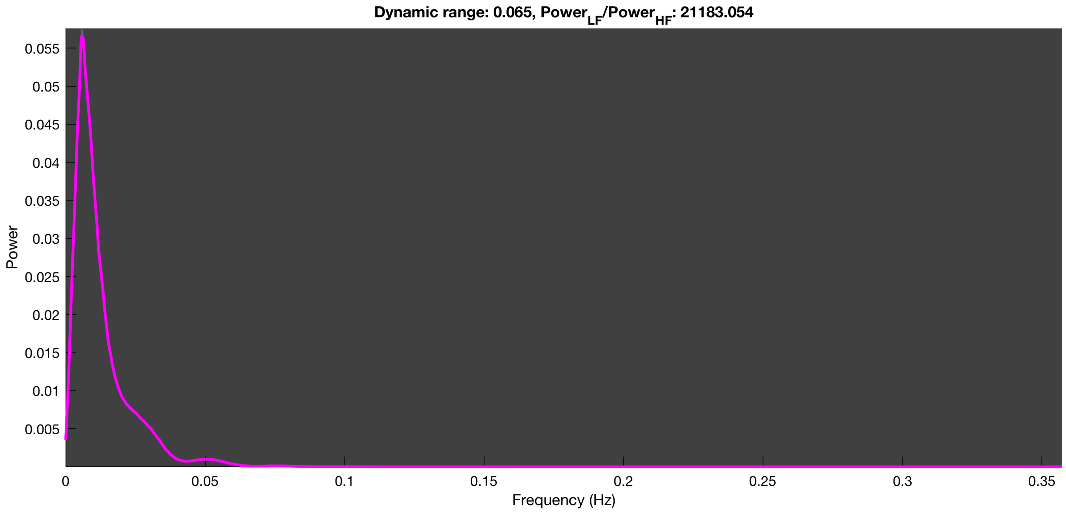

3-1 20n_tmap_component_ica_s1 018

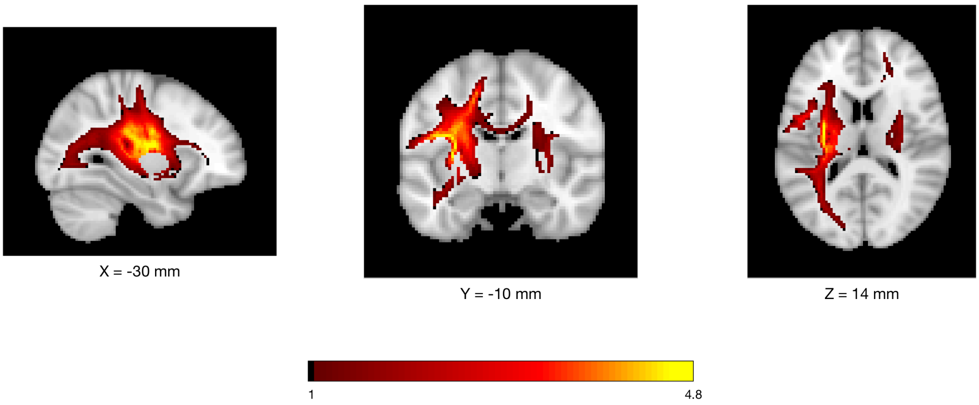


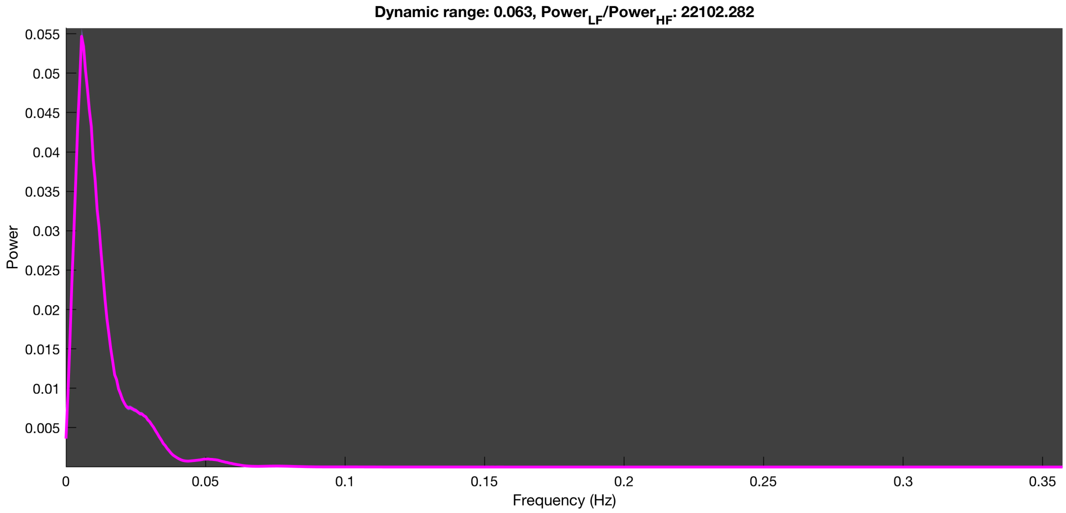

3-1 20n_tmap_component_ica_s1 019

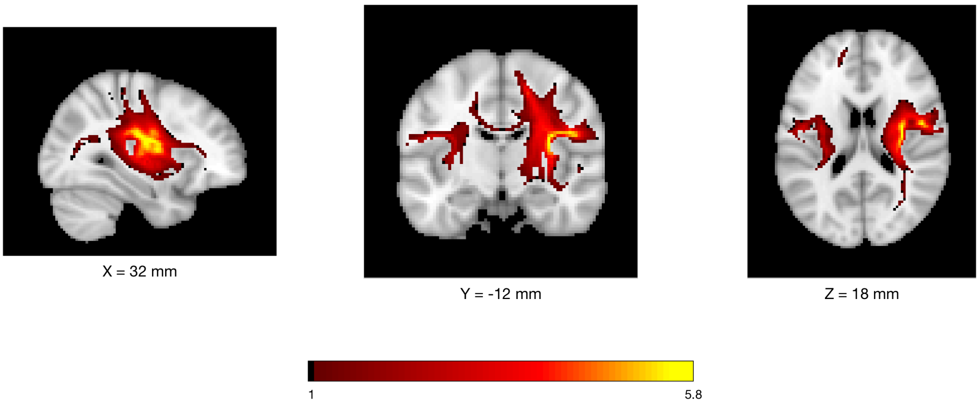


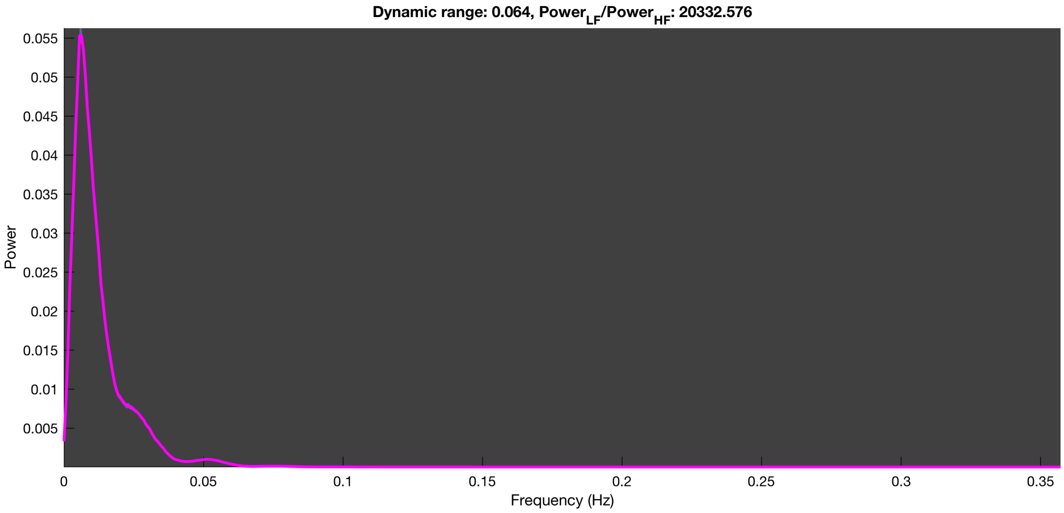

3-1 20n_tmap_component_ica_s1 020

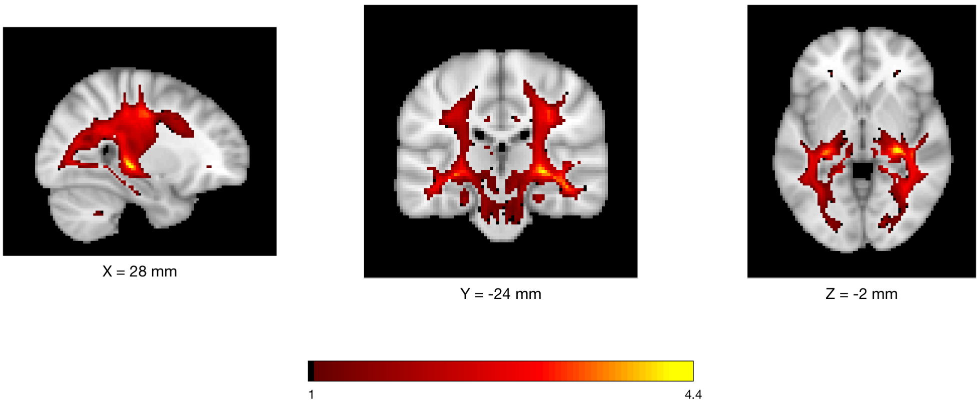


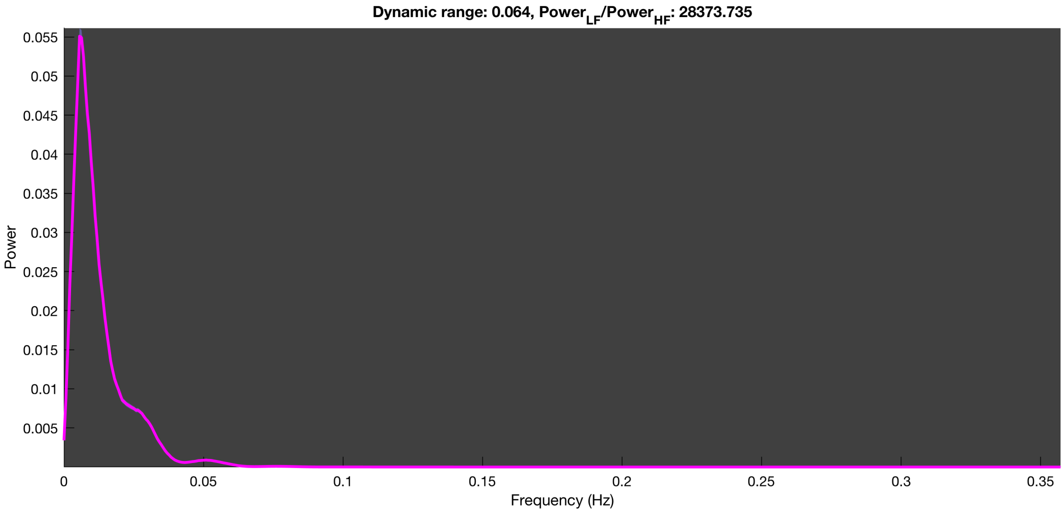

### LEMON dataset
*n=100 run*

3-1 100n_tmap_component_ica_s1 001

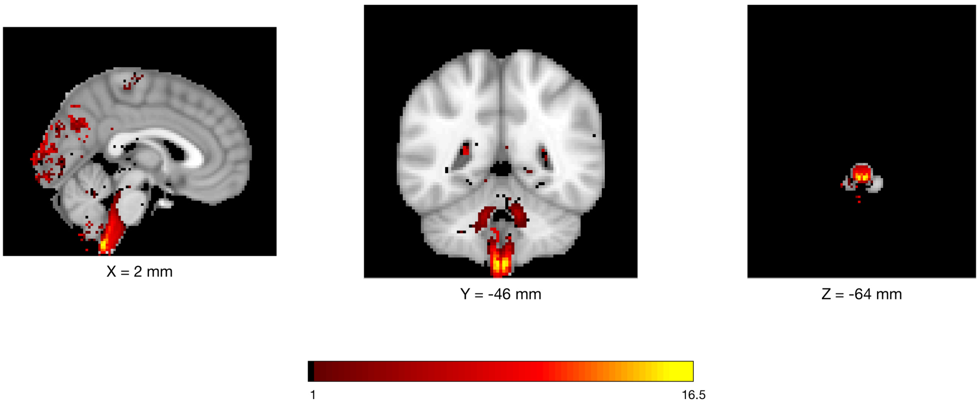


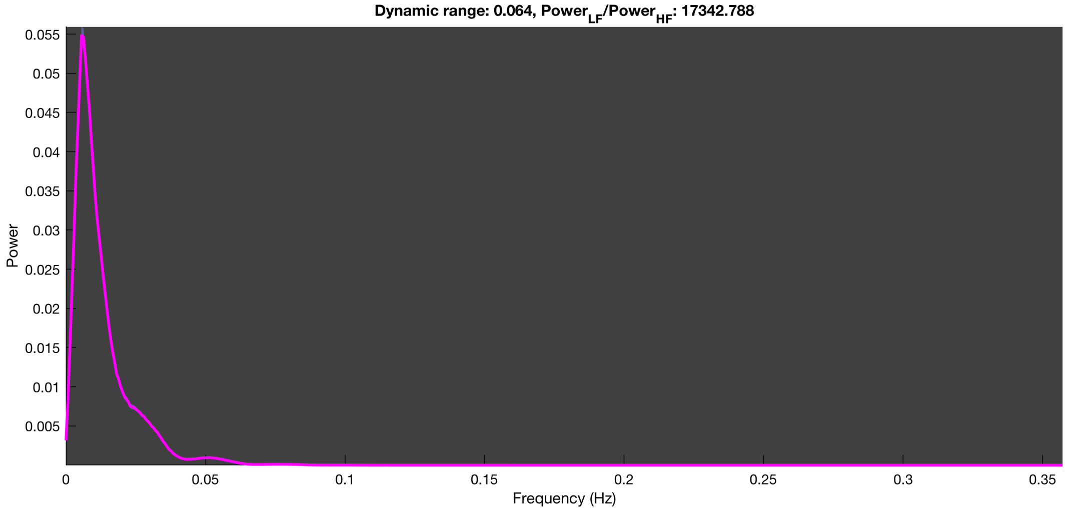

3-1 100n_tmap_component_ica_s1 002

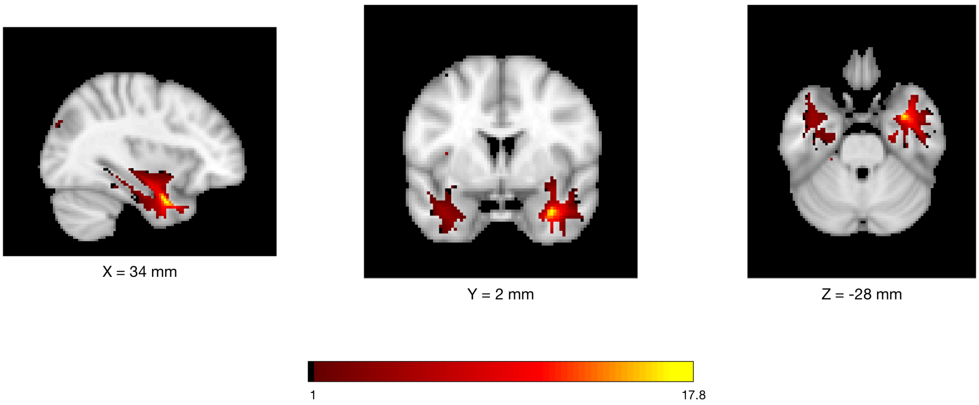


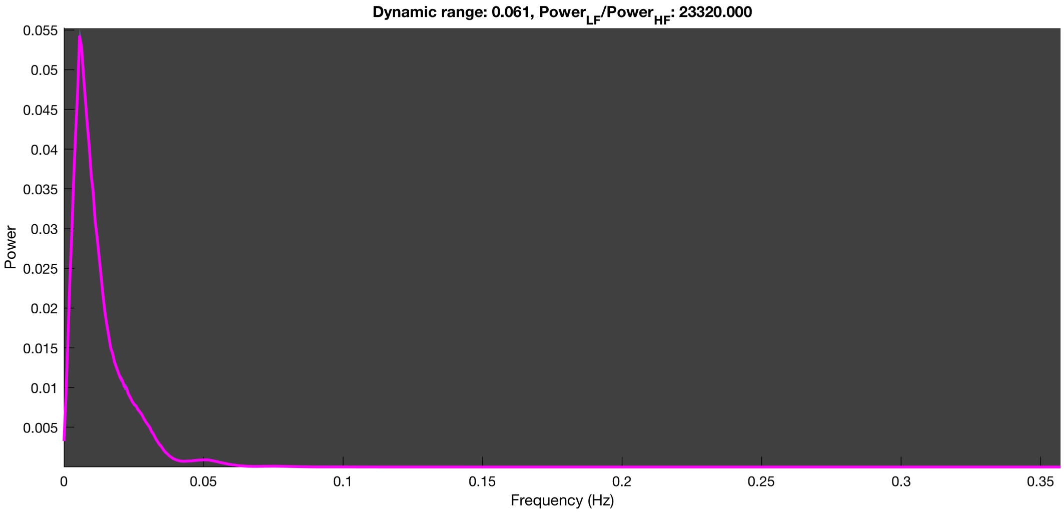

3-1 100n_tmap_component_ica_s1 003

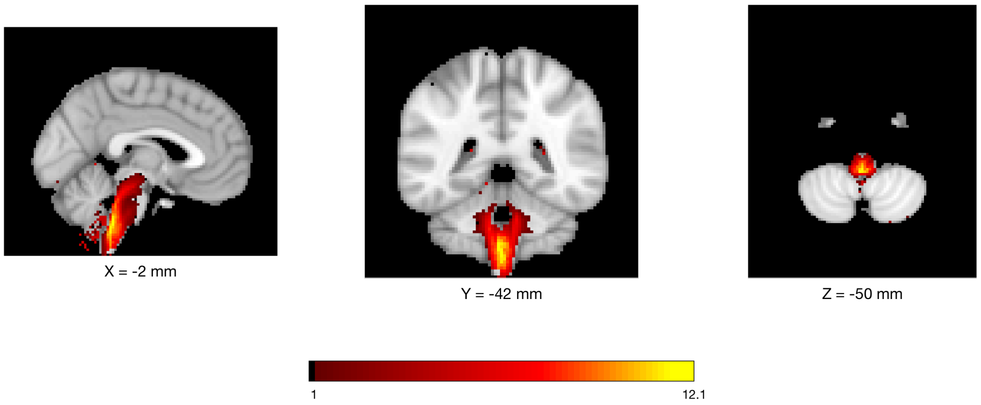


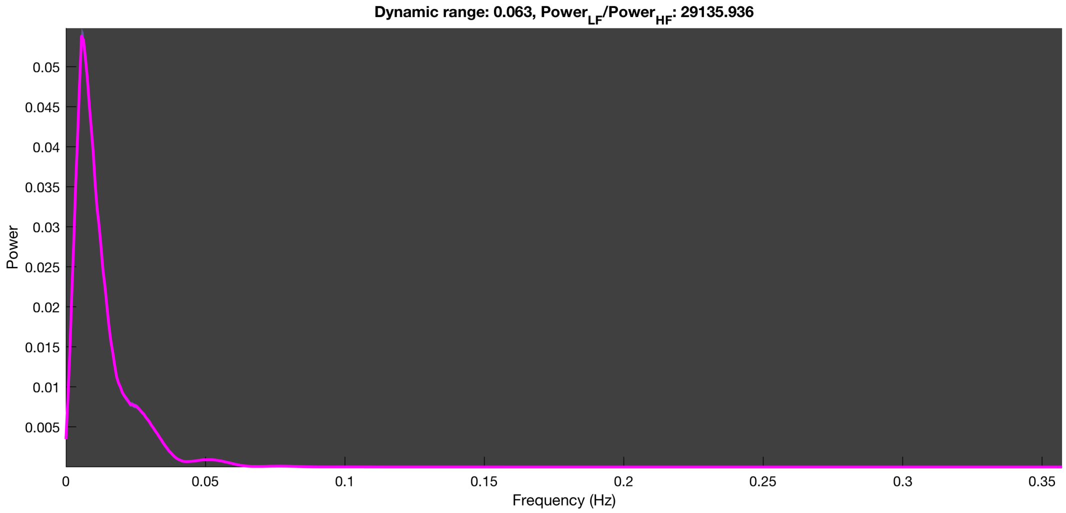

3-1 100n_tmap_component_ica_s1 004

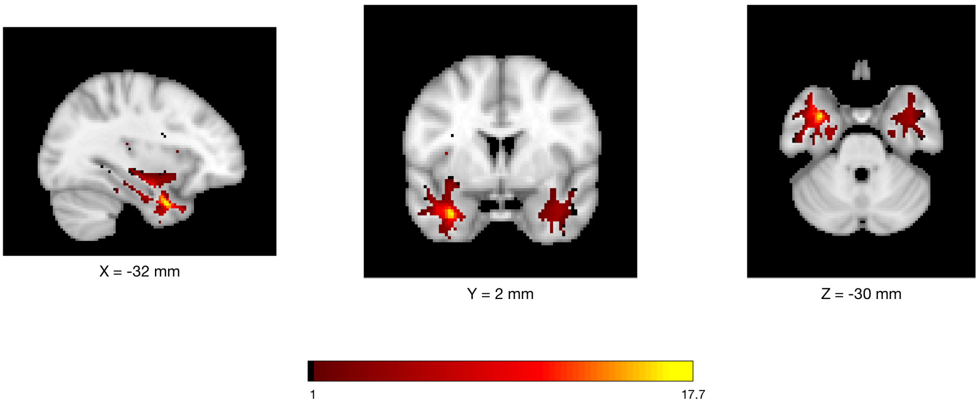


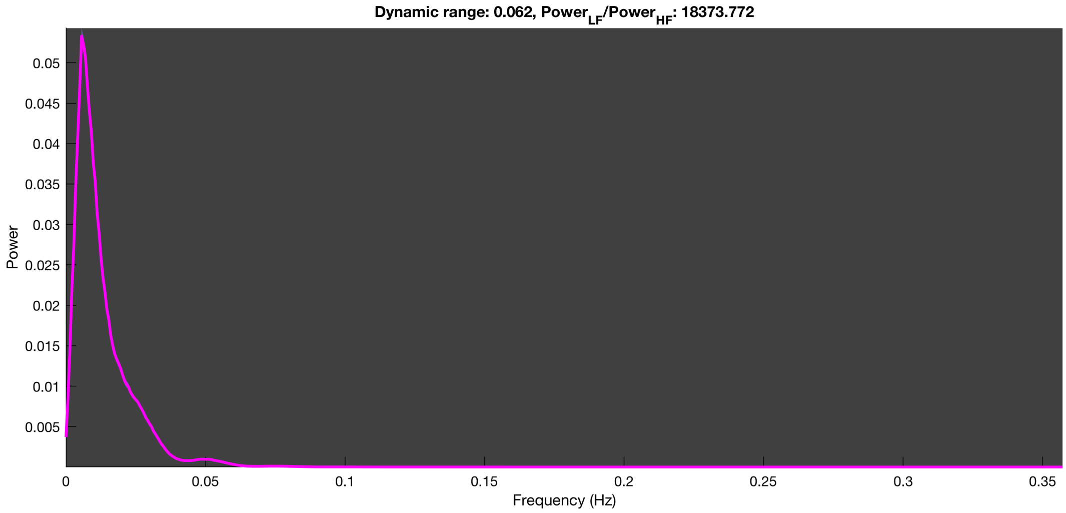

3-1 100n_tmap_component_ica_s1 005

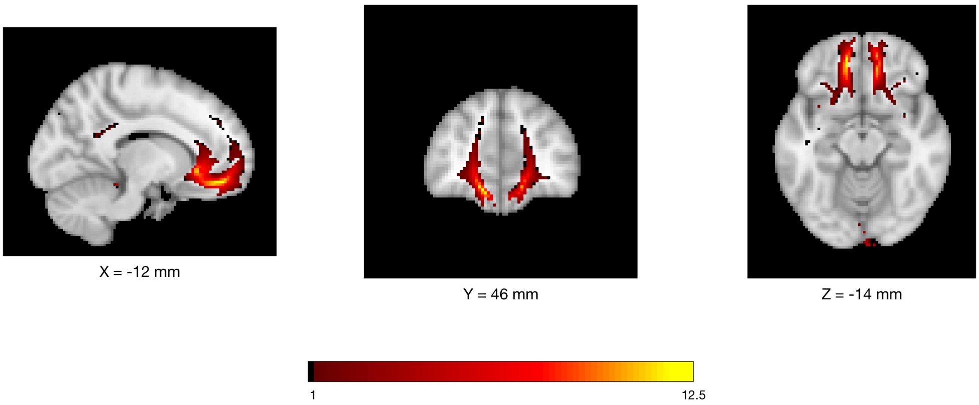


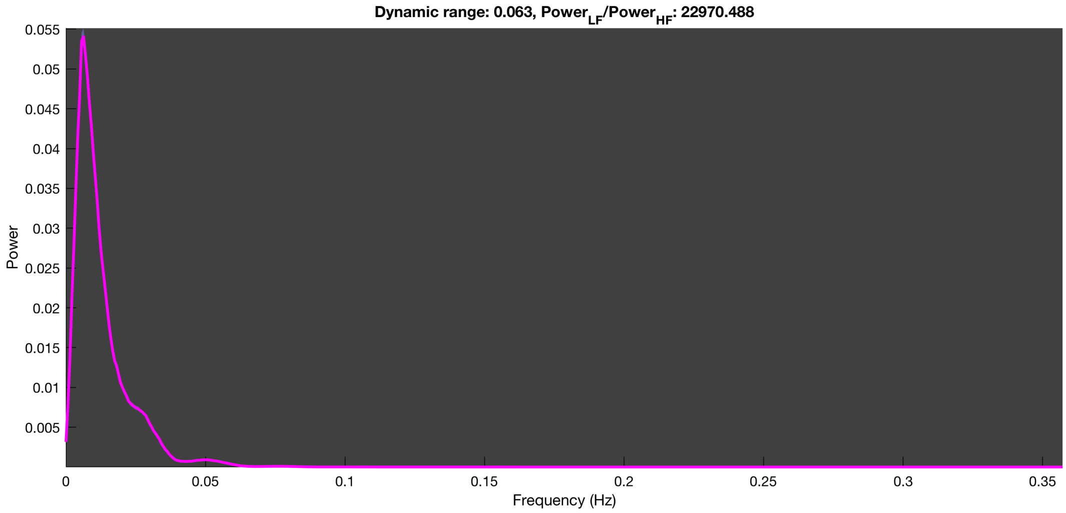

3-1 100n_tmap_component_ica_s1 006

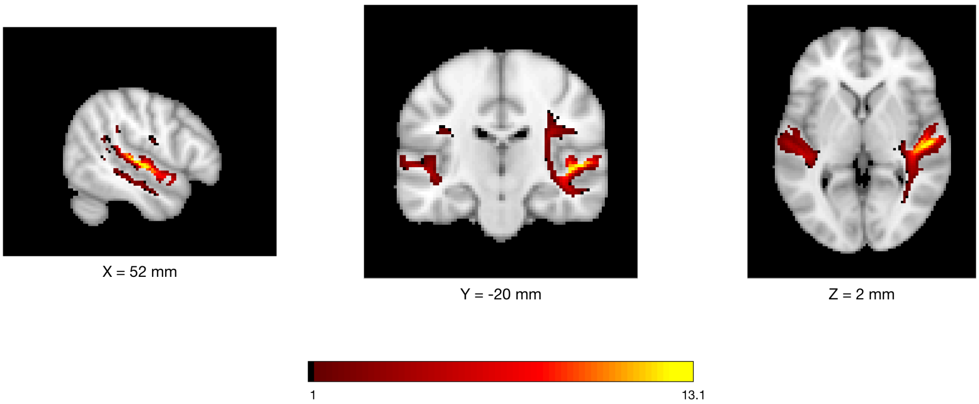


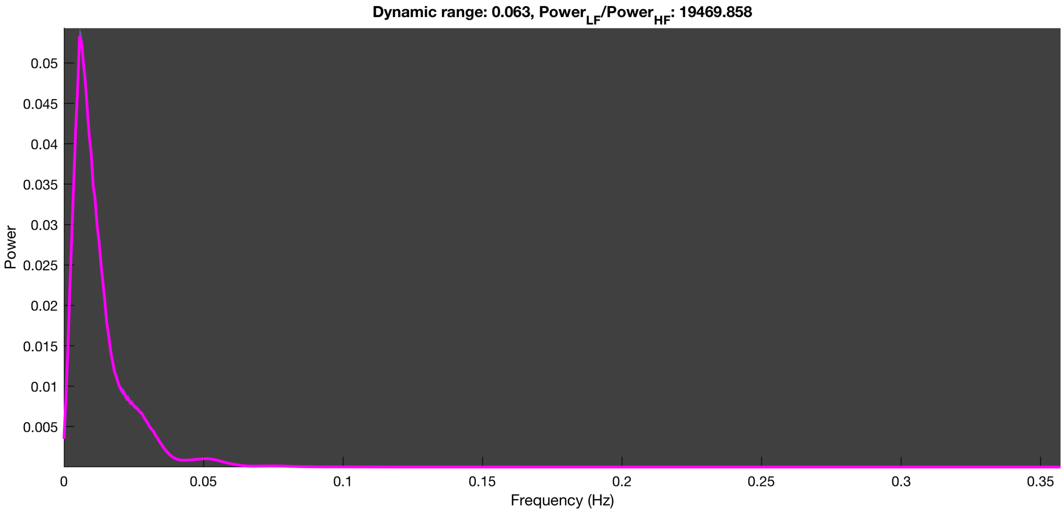

3-1 100n_tmap_component_ica_s1 007

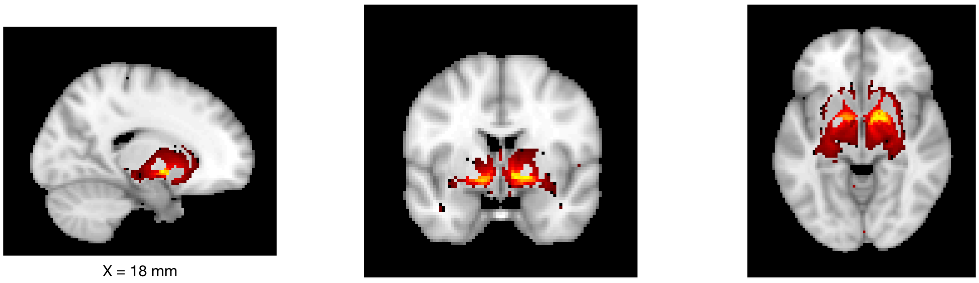

X = 18 mm Y = 0 mm Z = -6 mm

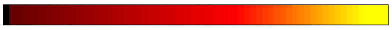


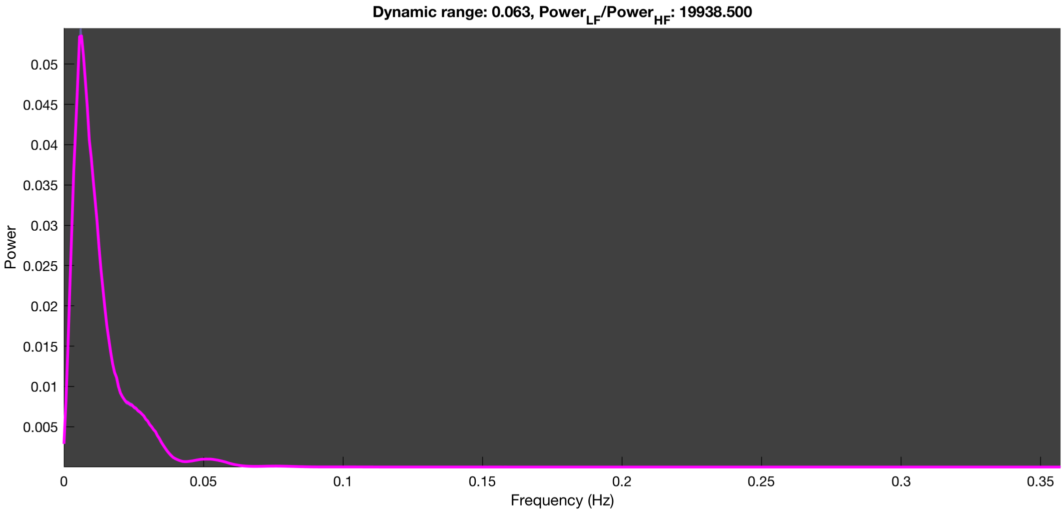

3-1 100n_tmap_component_ica_s1 008

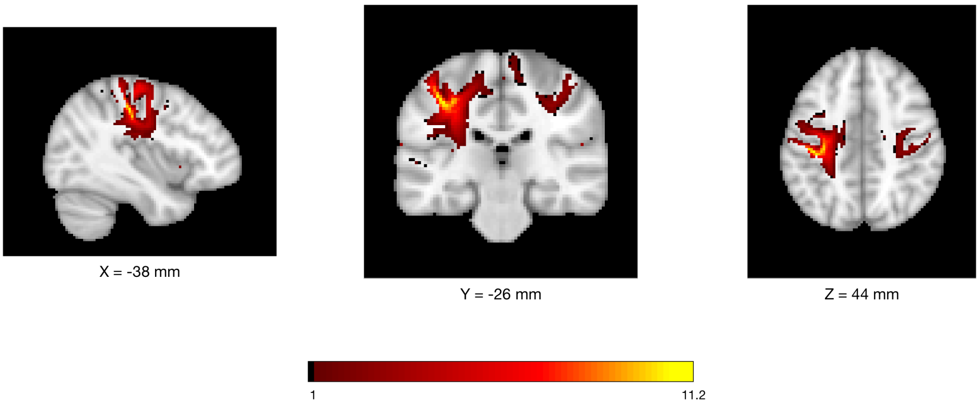


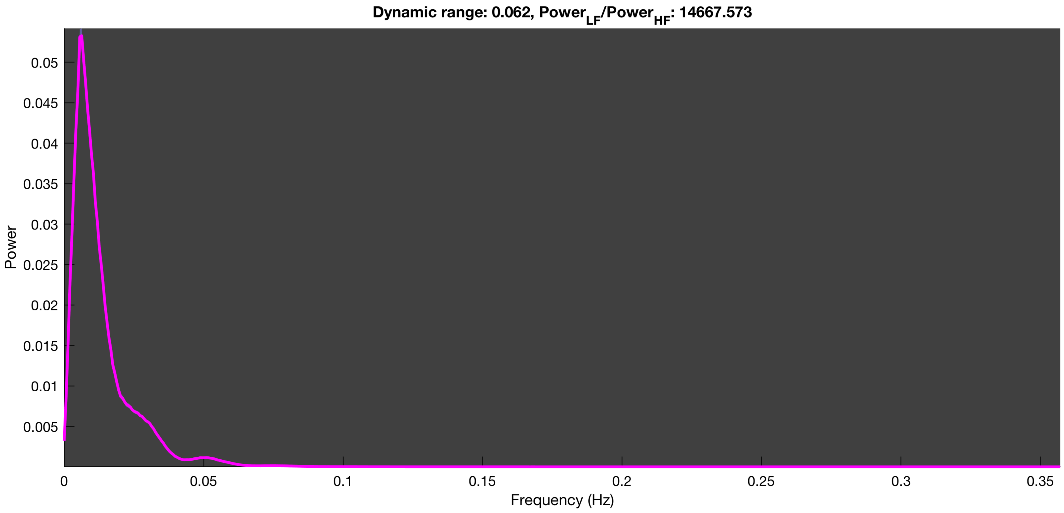

3-1 100n_tmap_component_ica_s1 009

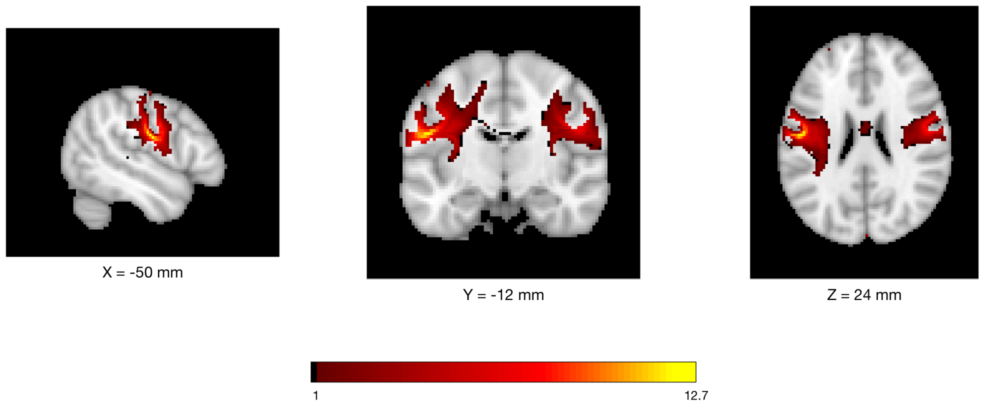


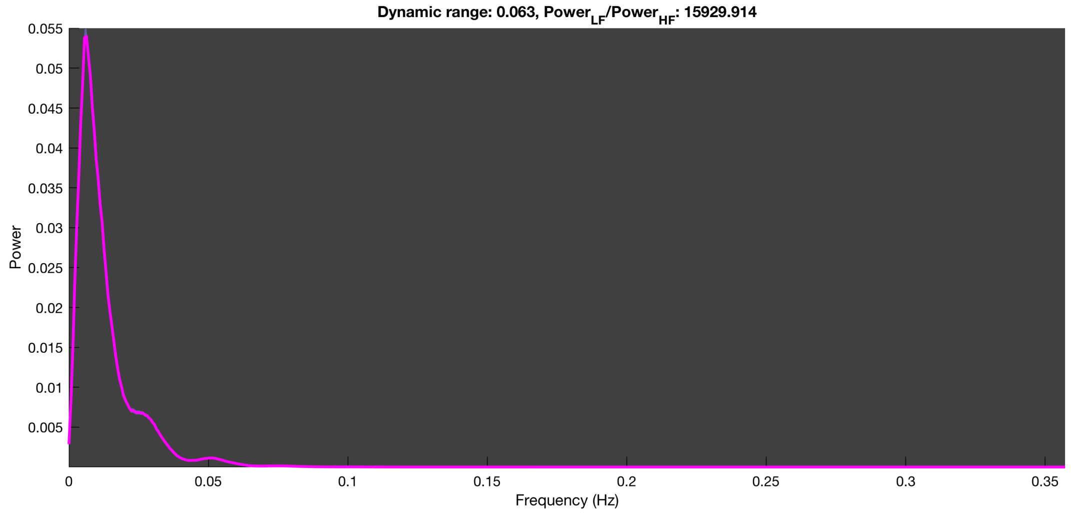

3-1 100n_tmap_component_ica_s1 010

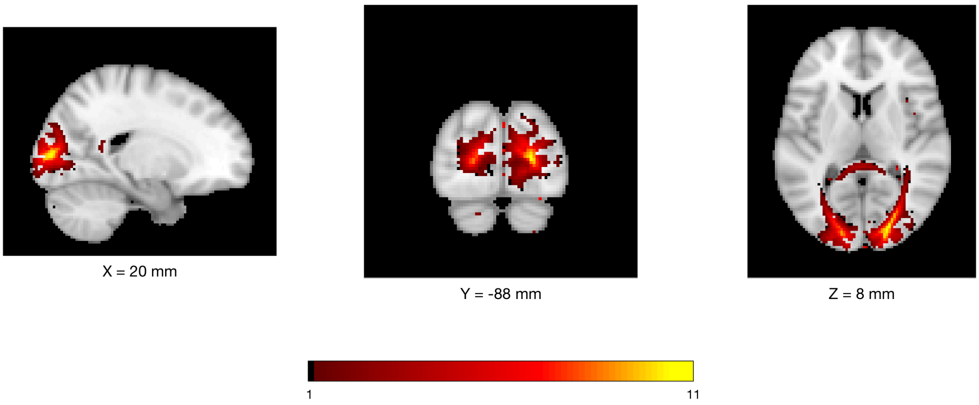


Dynamic range: 0.063, $Power_{LF}/Power_{HF}$: 14682.681

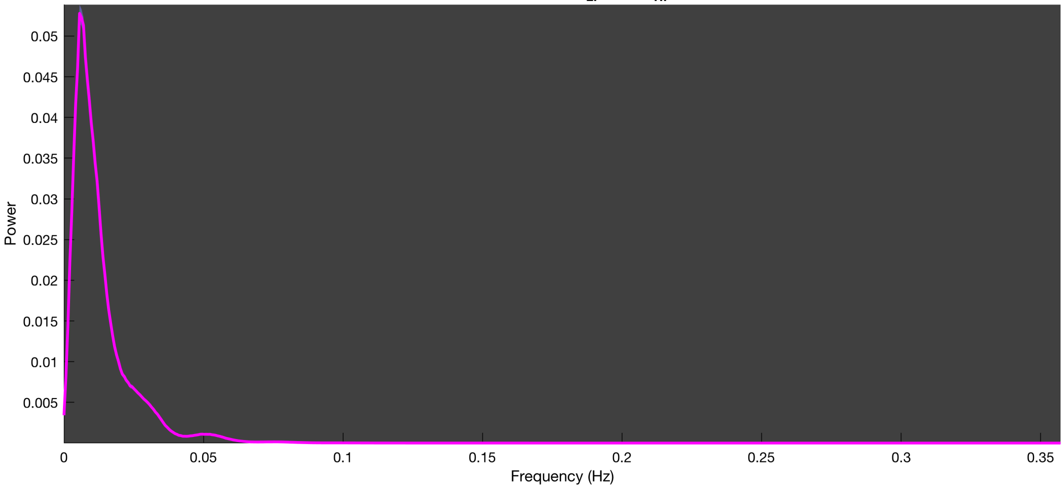

3-1 100n_tmap_component_ica_s1 011

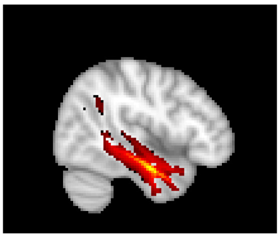

X = 46 mm

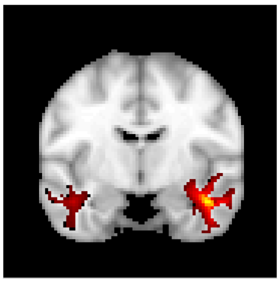

Y = -8 mm

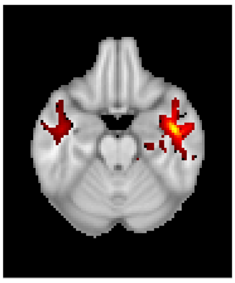

Z = -22 mm

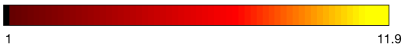


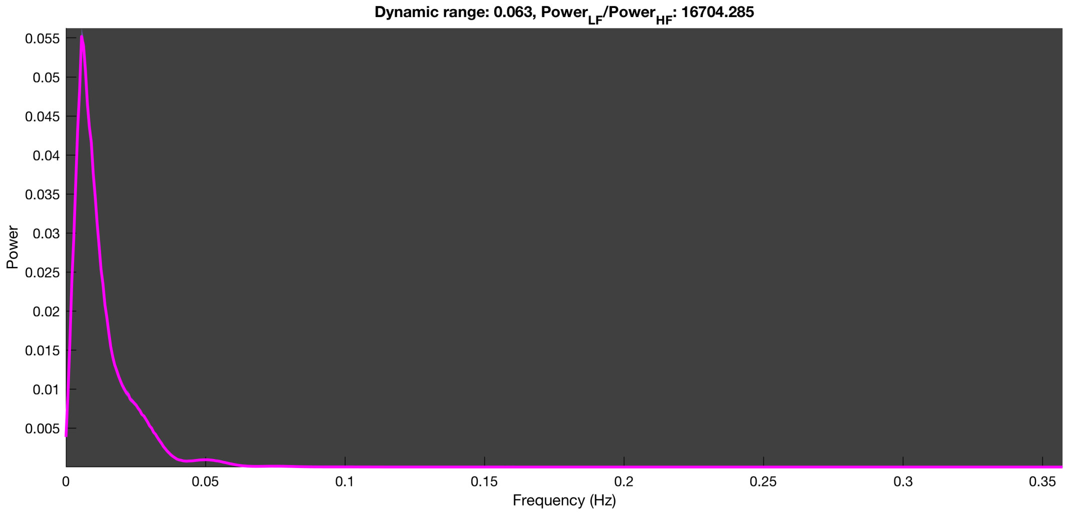

3-1 100n_tmap_component_ica_s1 012

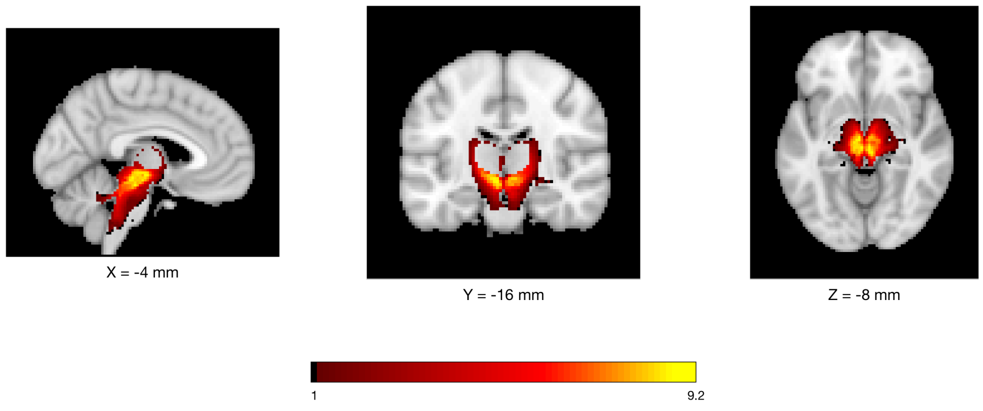


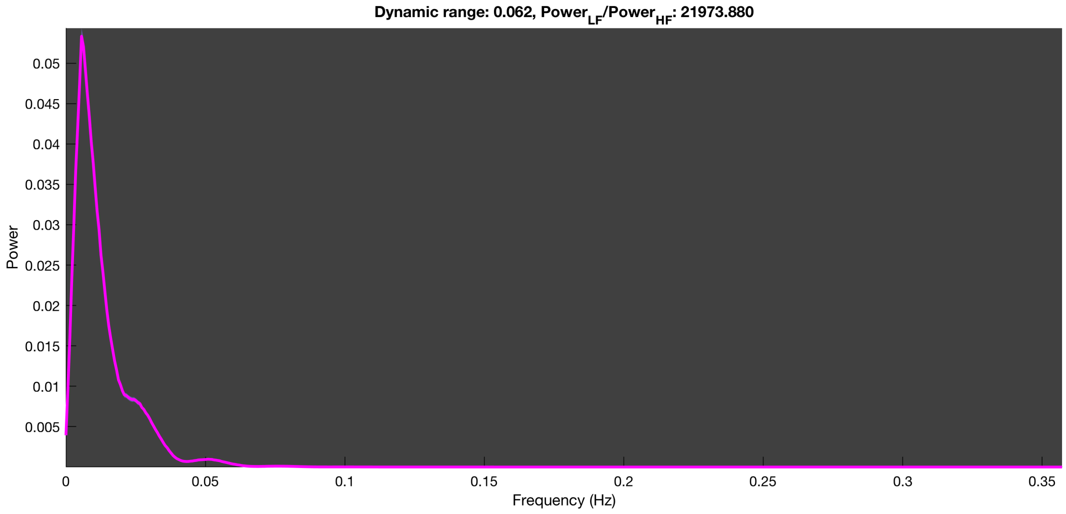

3-1 100n_tmap_component_ica_s1 013

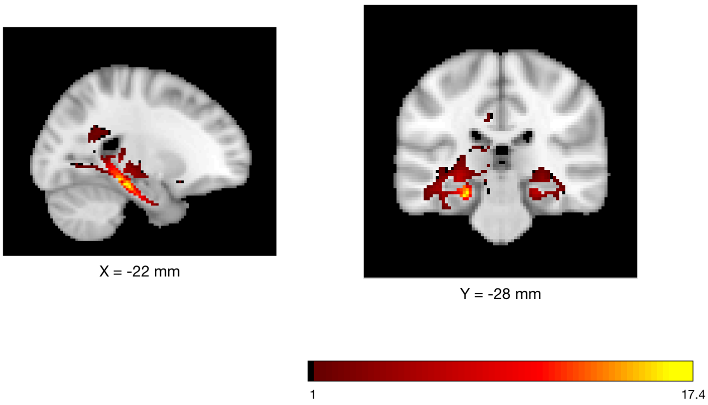


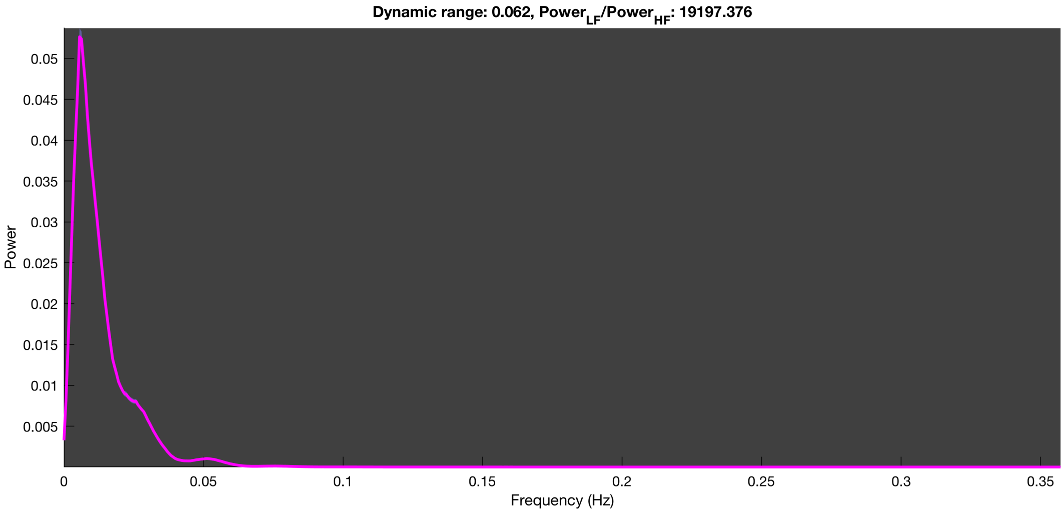

3-1 100n_tmap_component_ica_s1 014

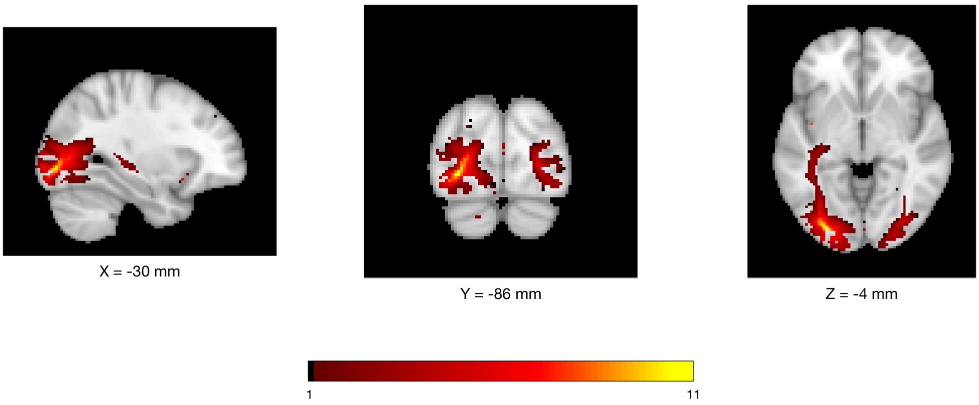


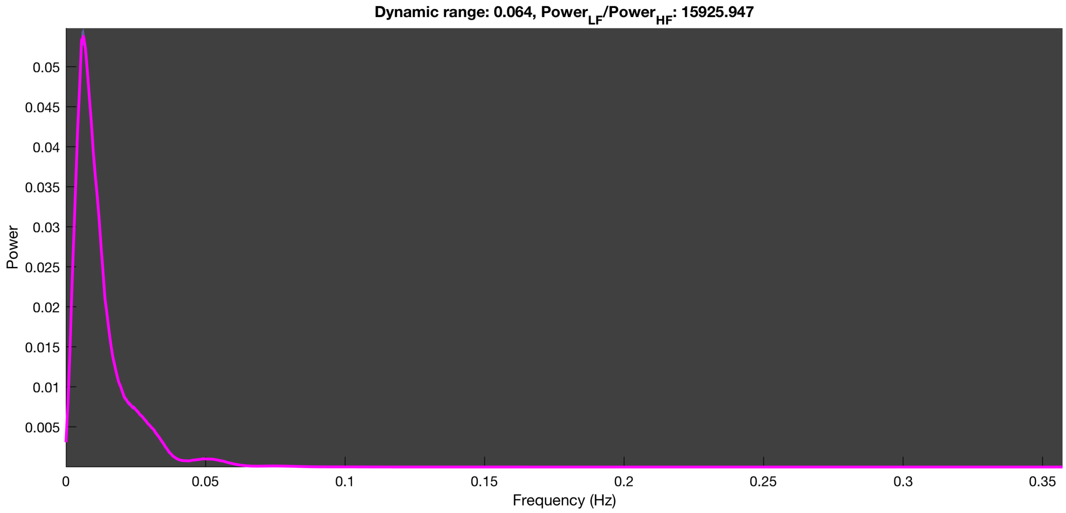

3-1 100n_tmap_component_ica_s1 015

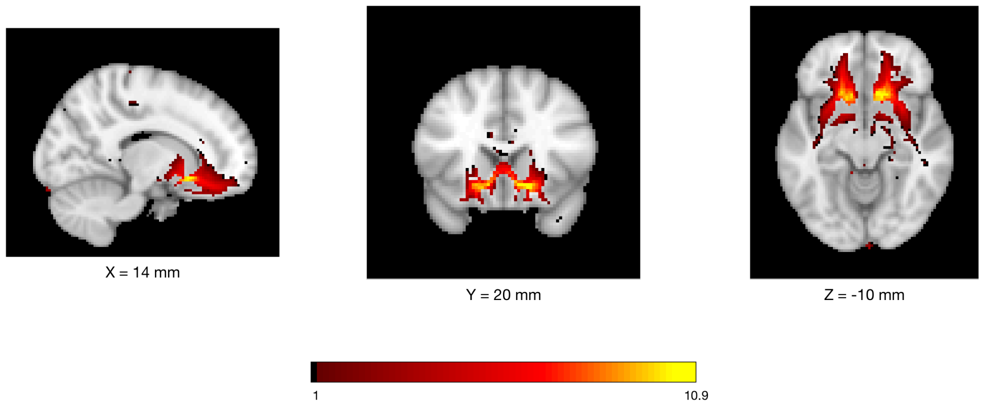


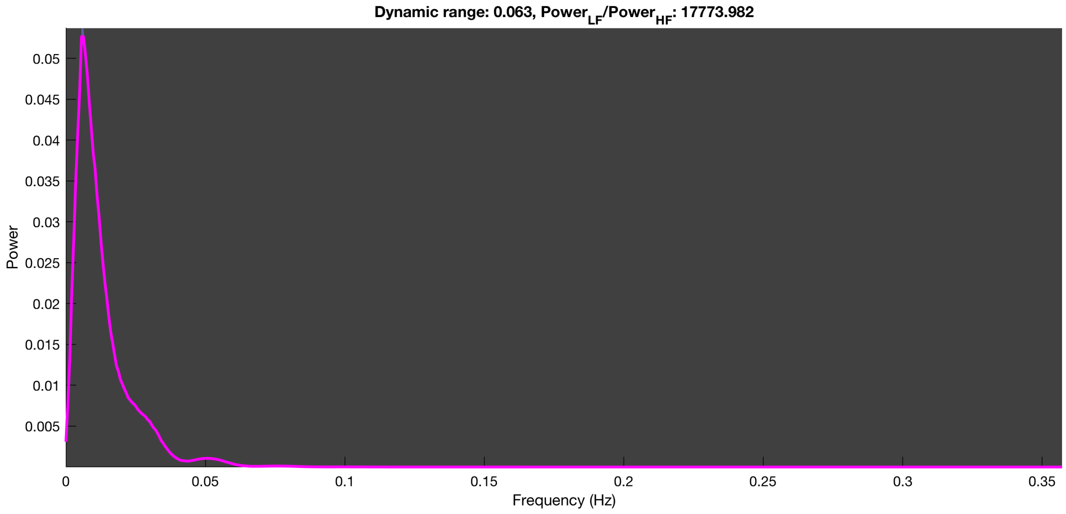

3-1 100n_tmap_component_ica_s1 016

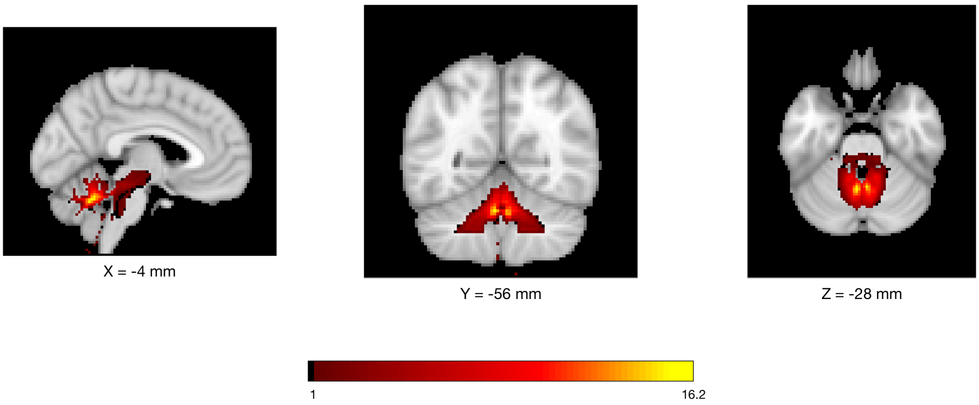


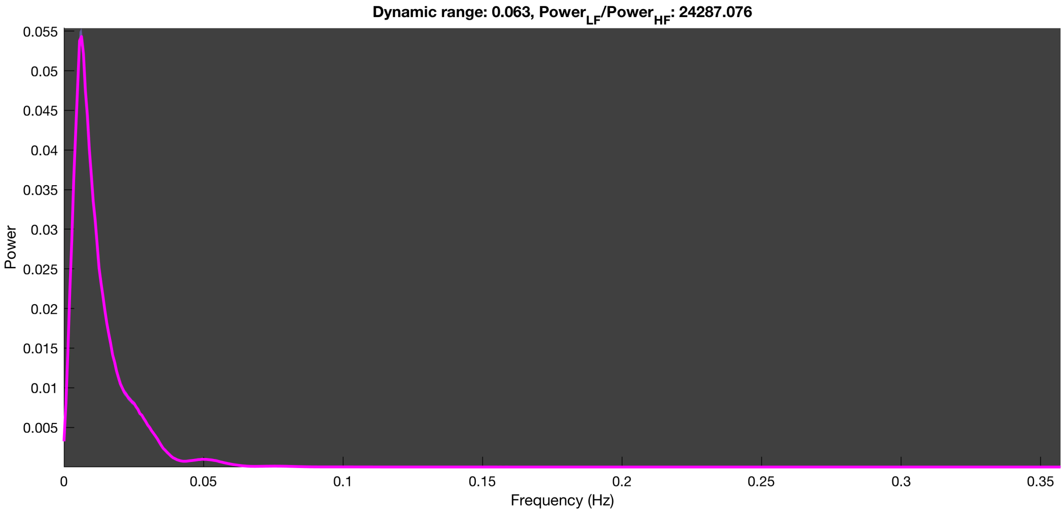

3-1 100n_tmap_component_ica_s1 017

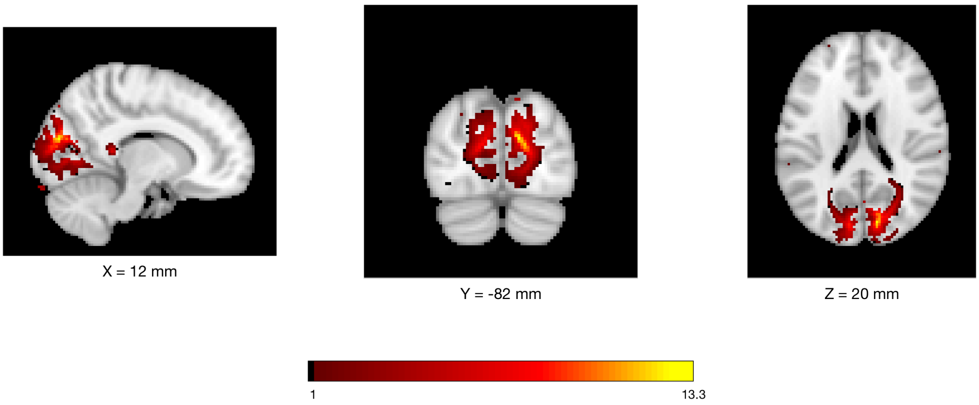


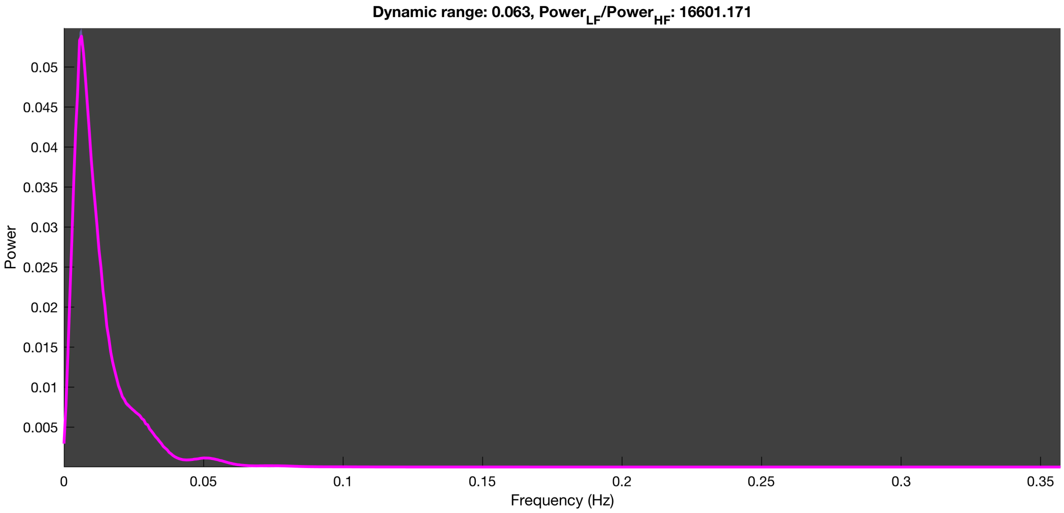

3-1 100n_tmap_component_ica_s1 018

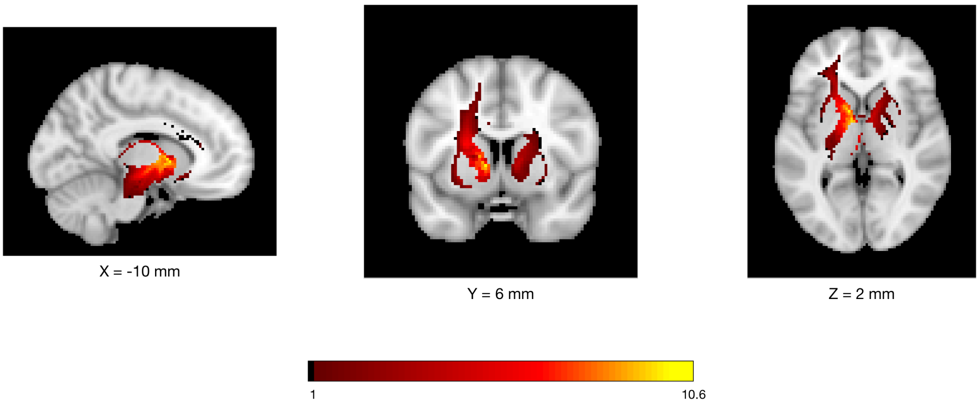


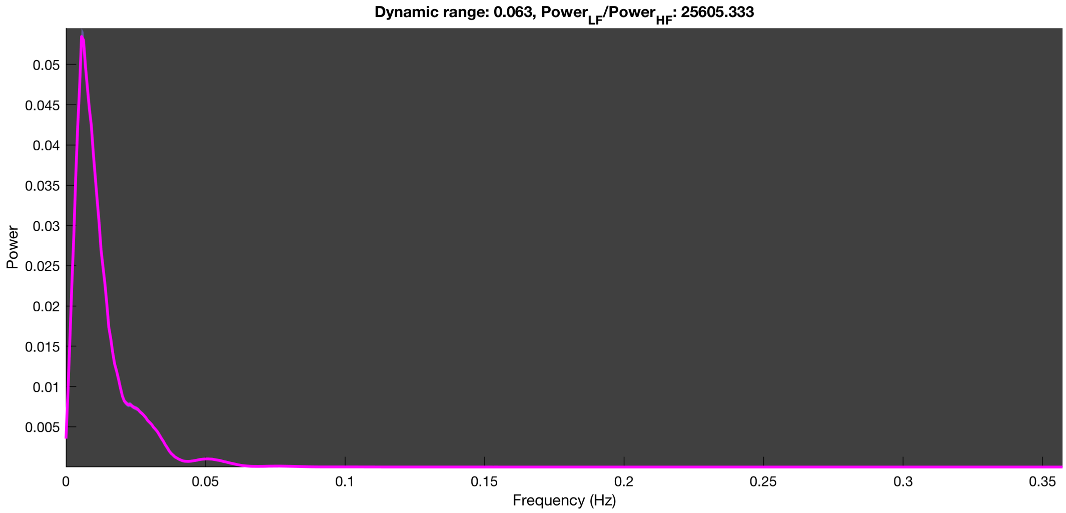

3-1 100n_tmap_component_ica_s1 019

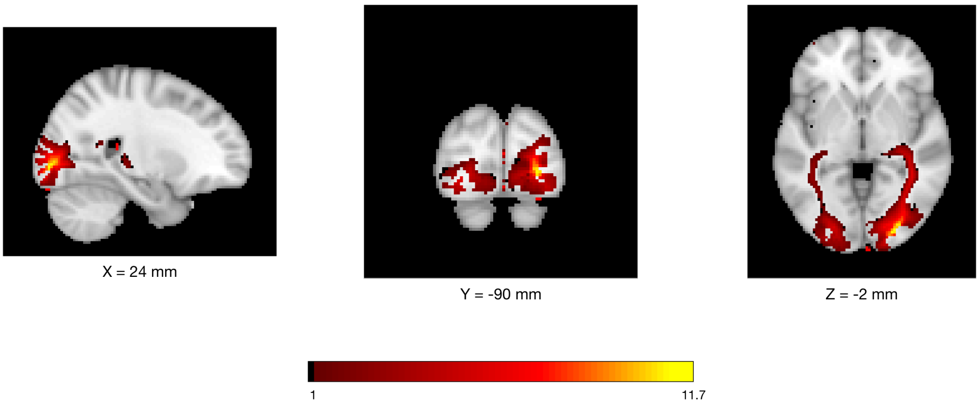


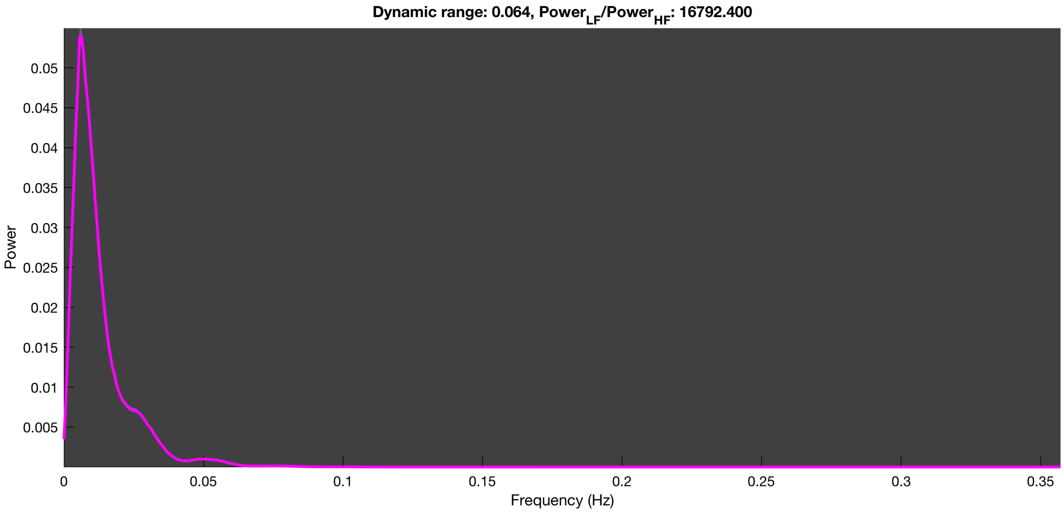

3-1 100n_tmap_component_ica_s1 020

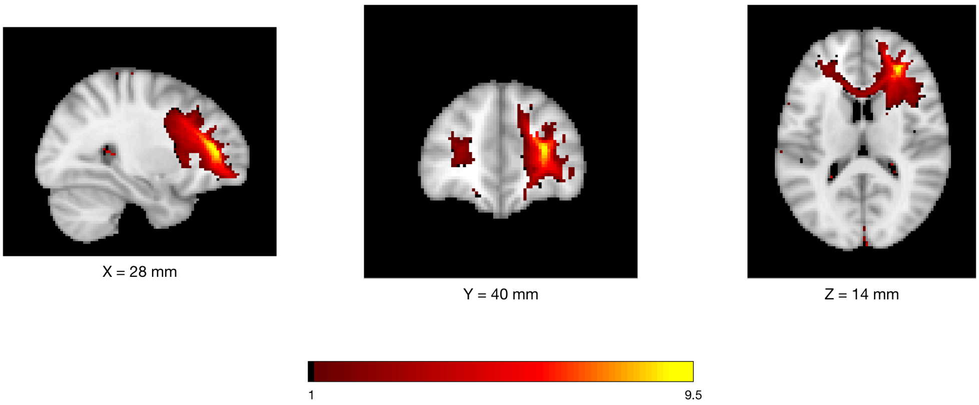


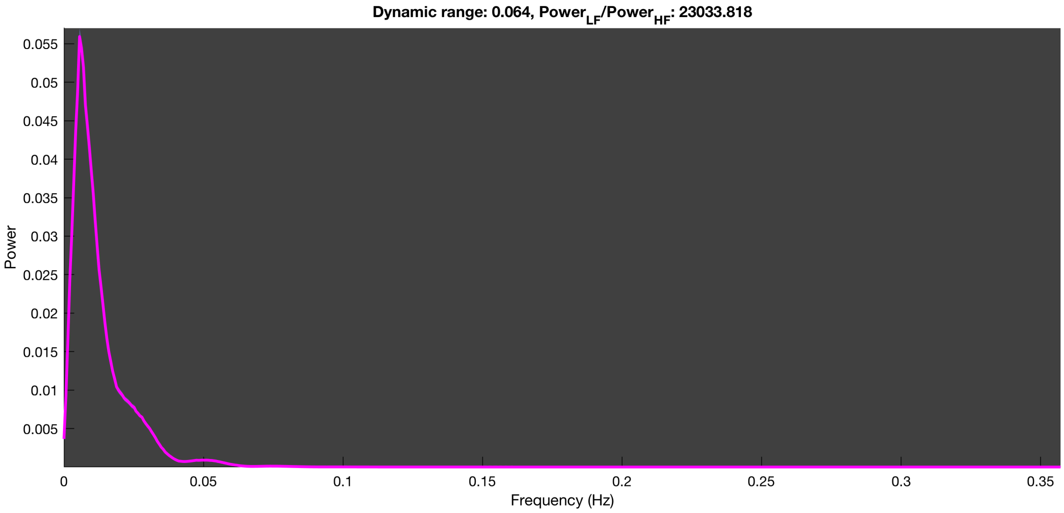

3-1 100n_tmap_component_ica_s1 021

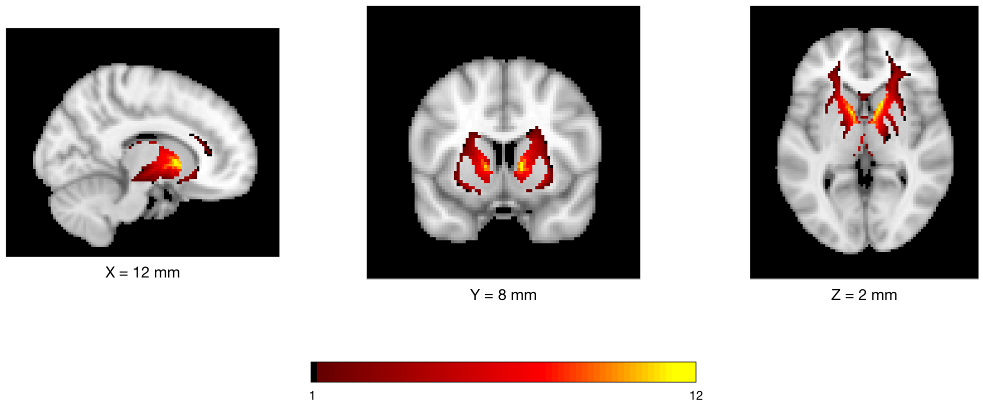


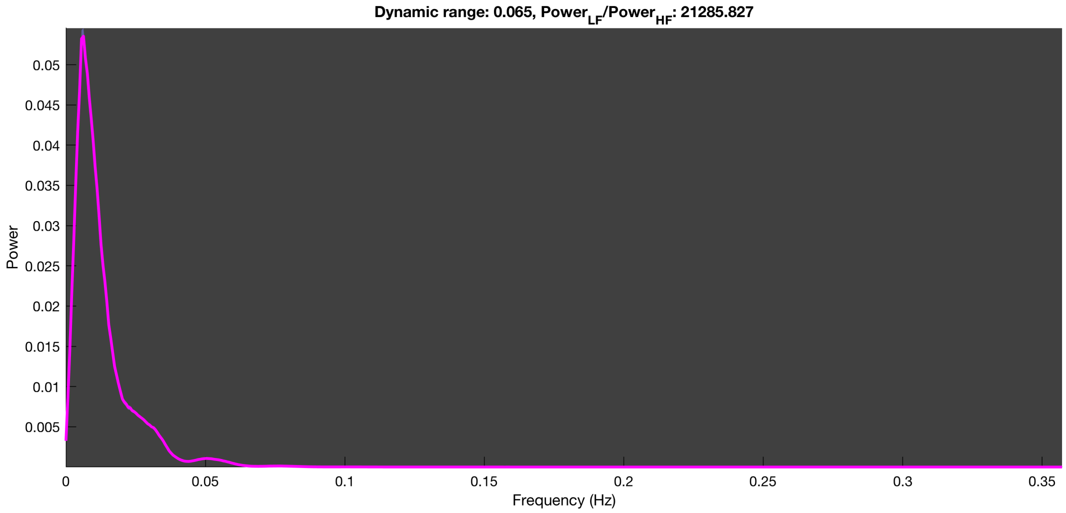

3-1 100n_tmap_component_ica_s1 022

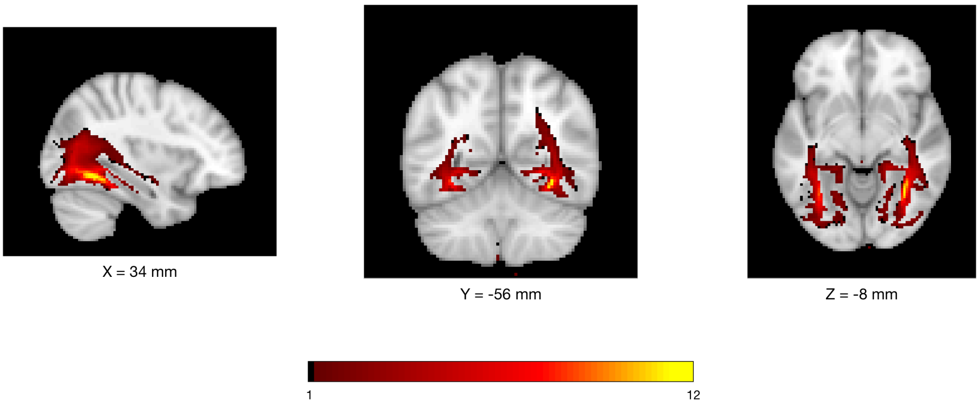


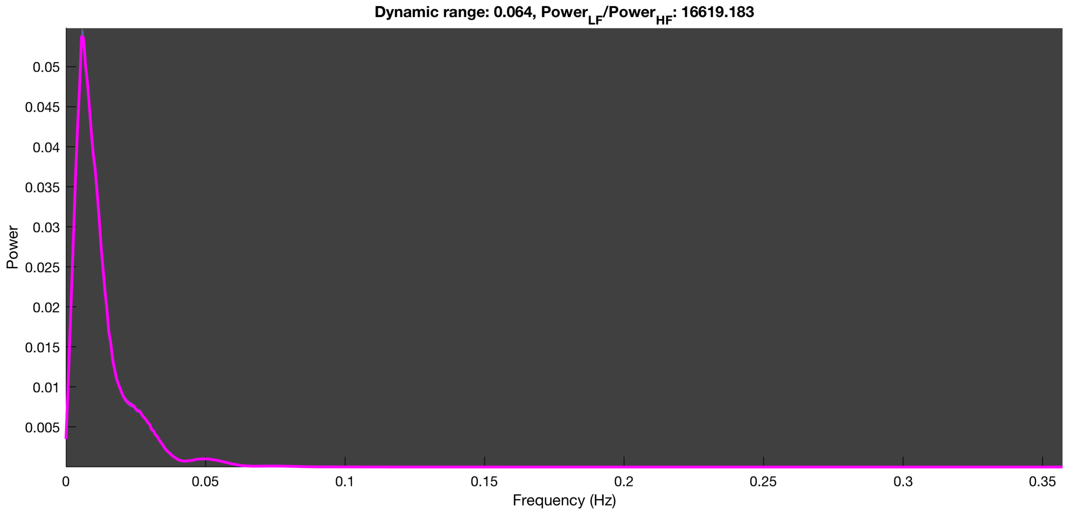

3-1 100n_tmap_component_ica_s1 023

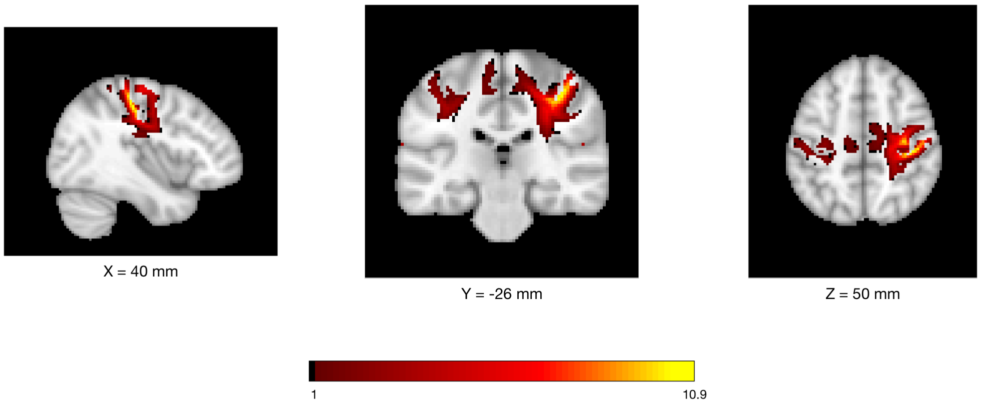


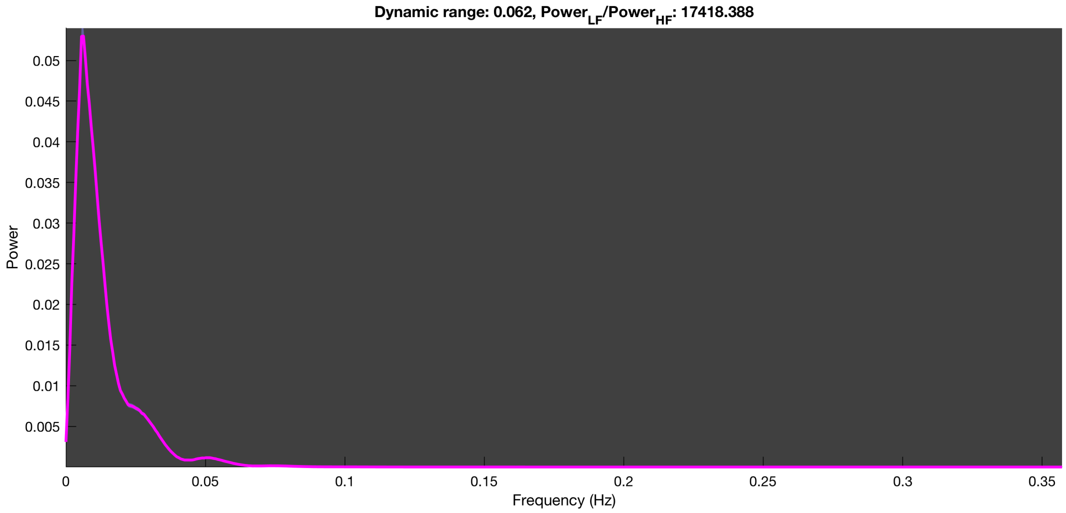

3-1 100n_tmap_component_ica_s1 024

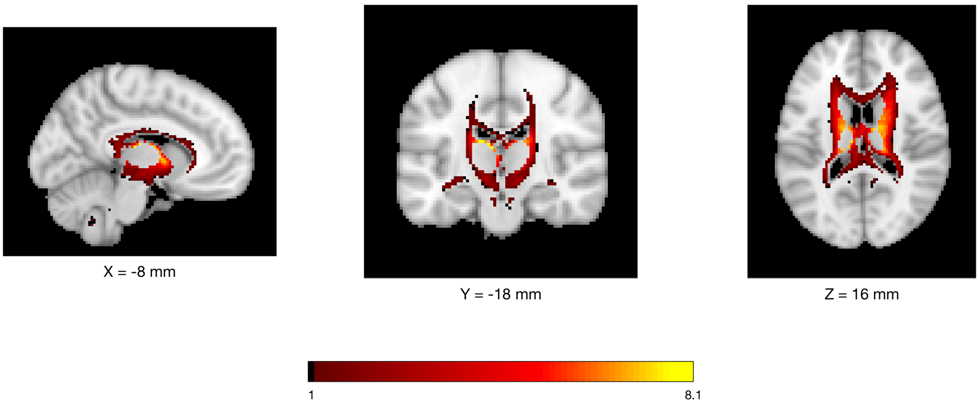


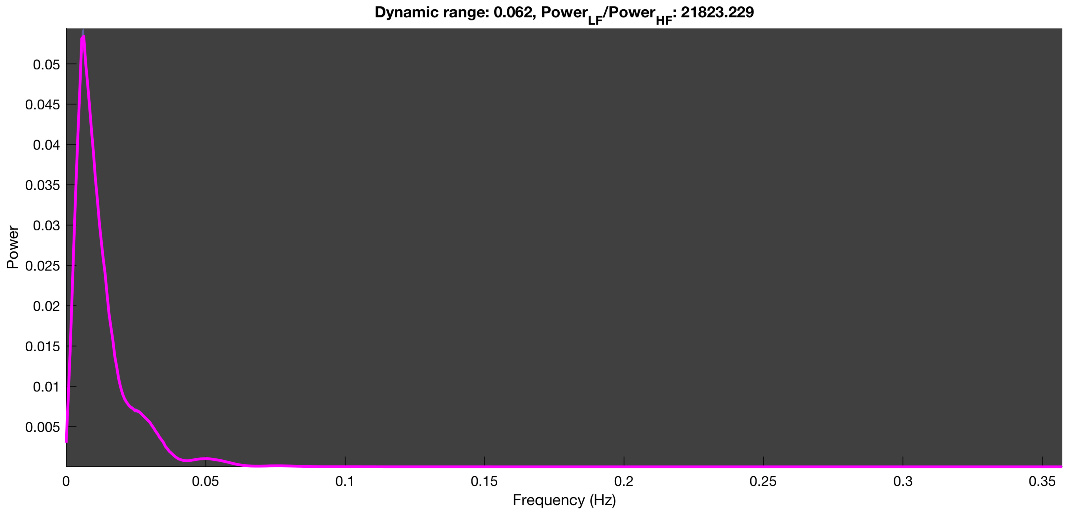

3-1 100n_tmap_component_ica_s1 025

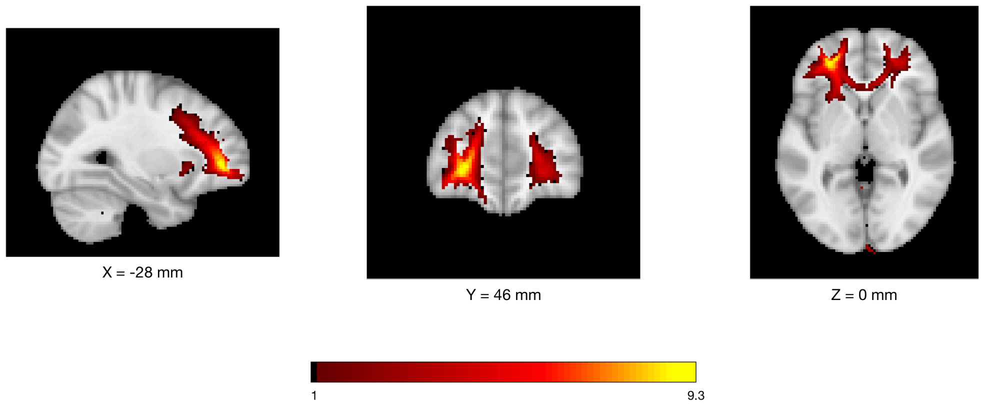


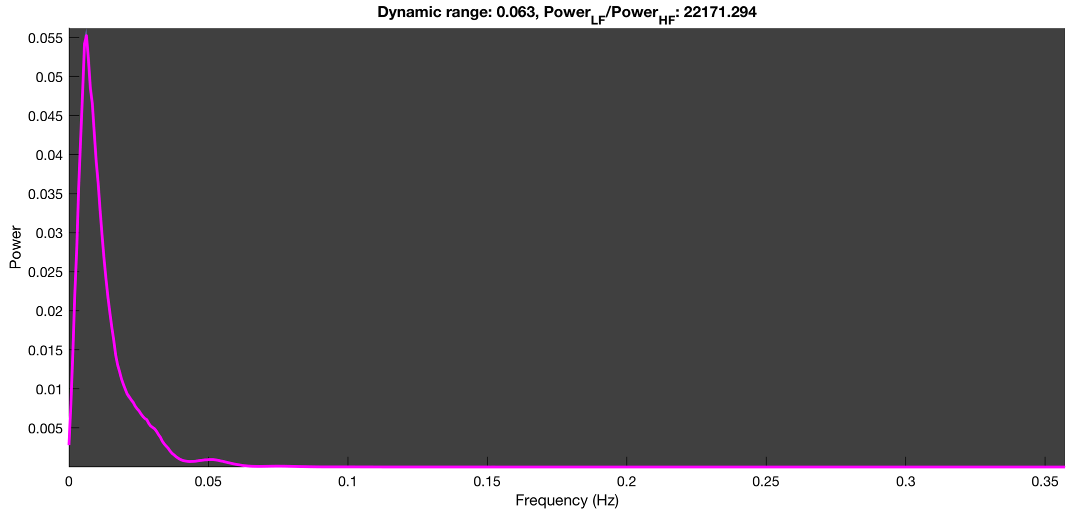

3-1 100n_tmap_component_ica_s1 026

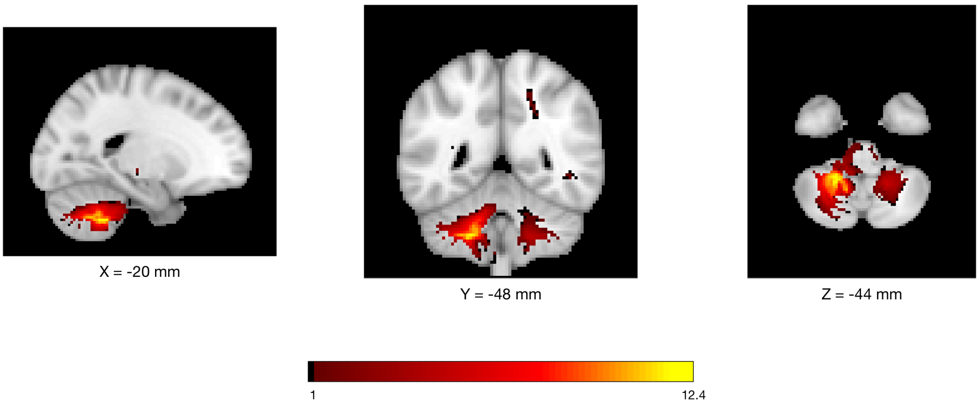


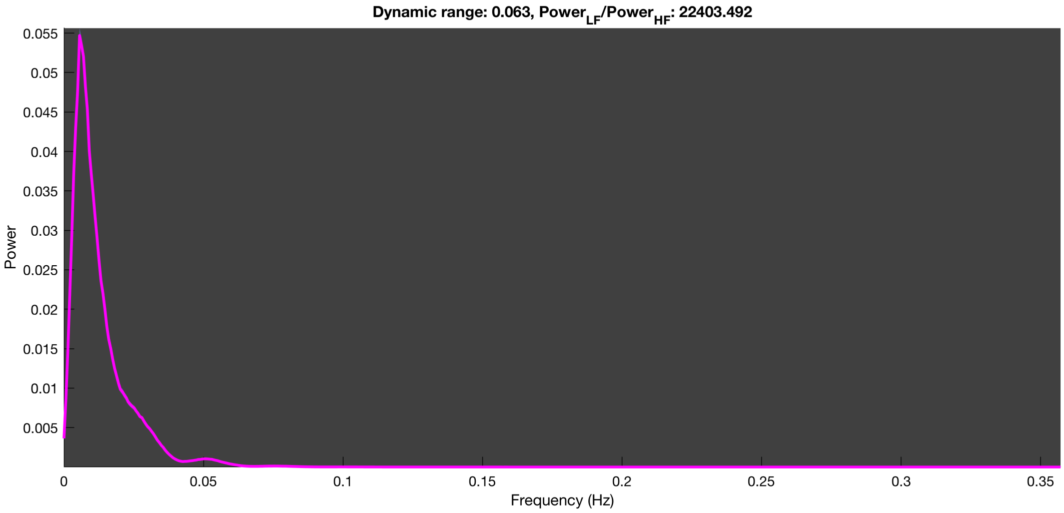

3-1 100n_tmap_component_ica_s1 027

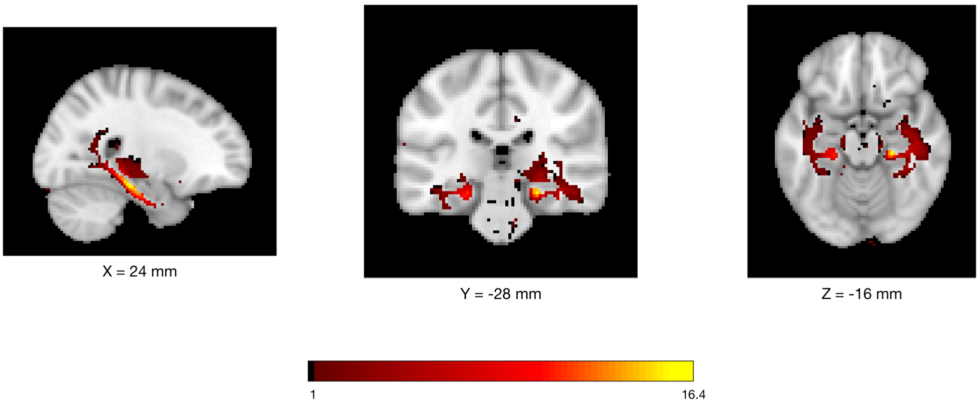


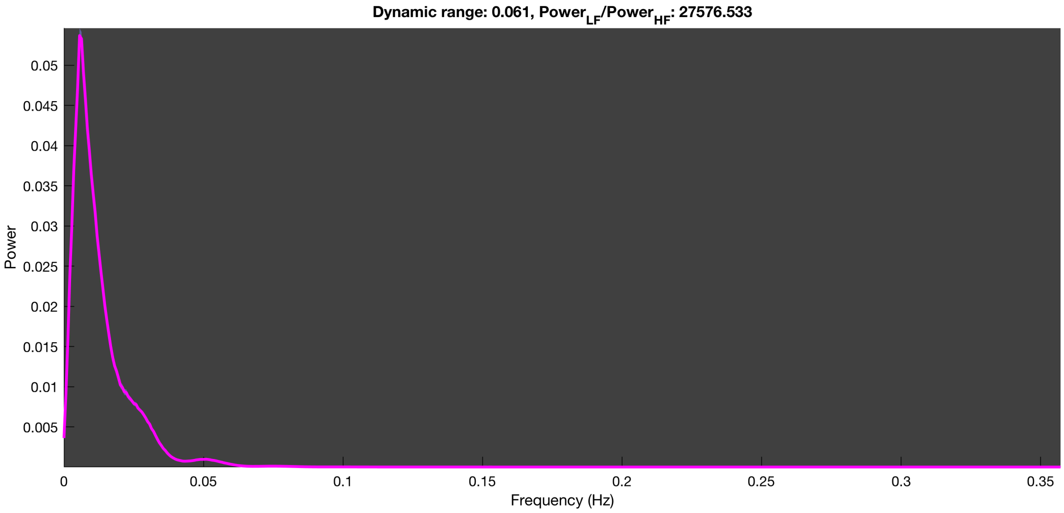

3-1 100n_tmap_component_ica_s1 028

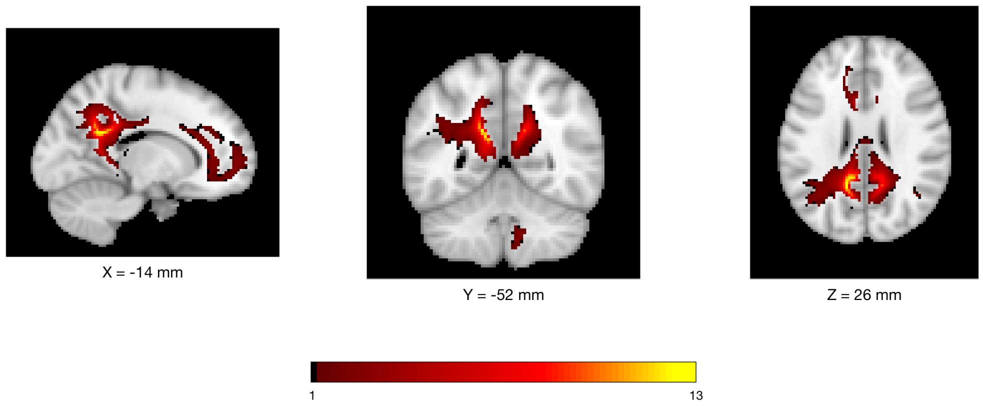


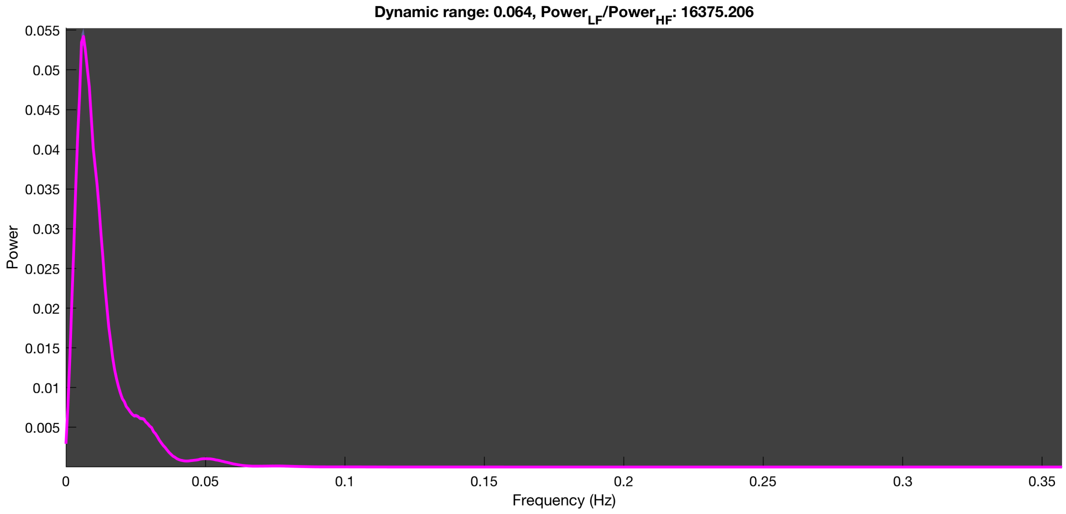

3-1 100n_tmap_component_ica_s1 029

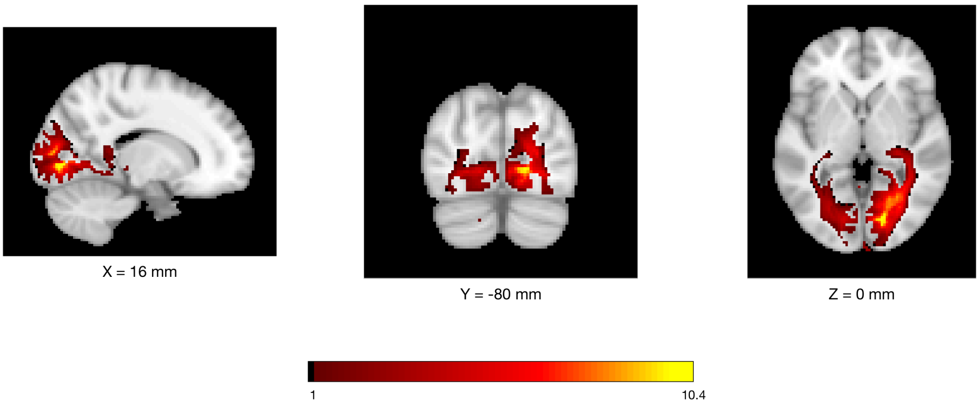


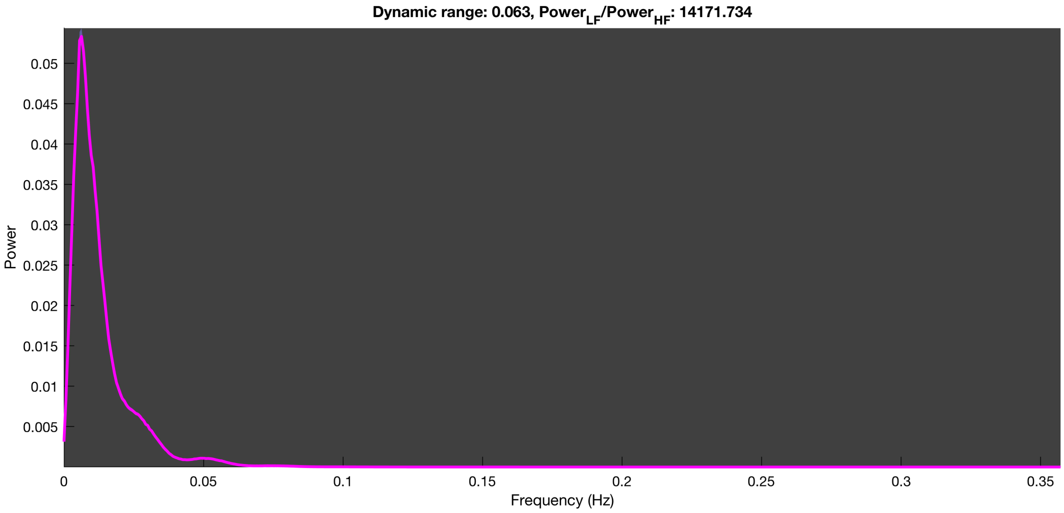

3-1 100n_tmap_component_ica_s1 030

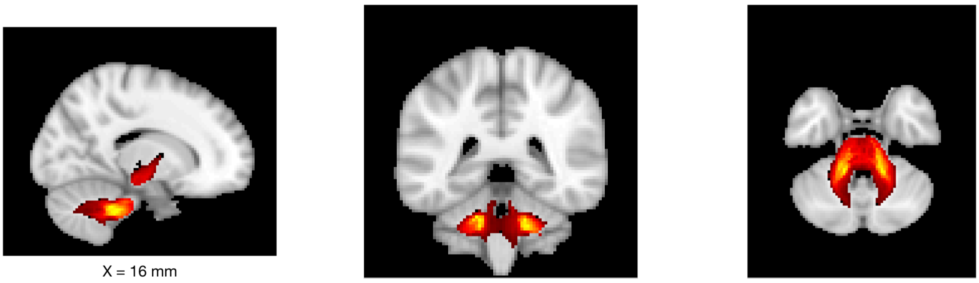

X = 16 mm

Y = -40 mm

Z = -34 mm

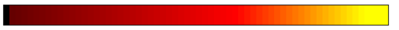


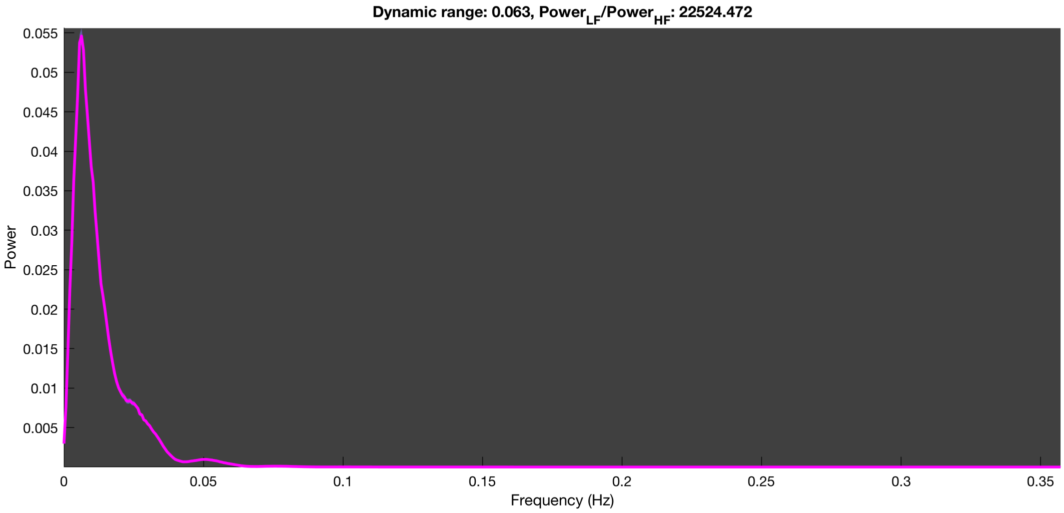

3-1 100n_tmap_component_ica_s1 031

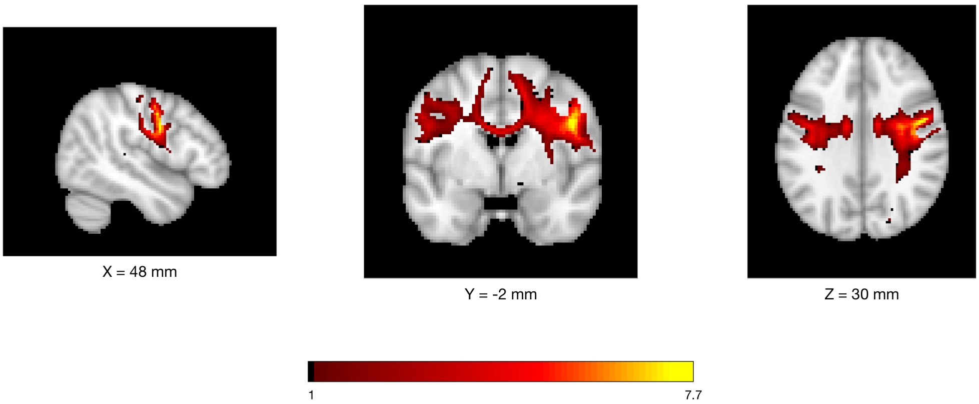


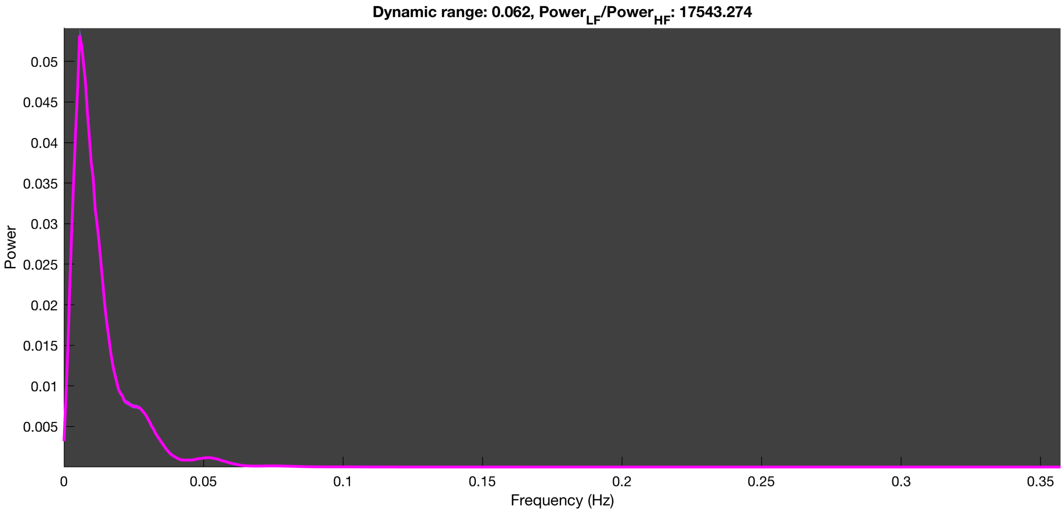

3-1 100n_tmap_component_ica_s1 032

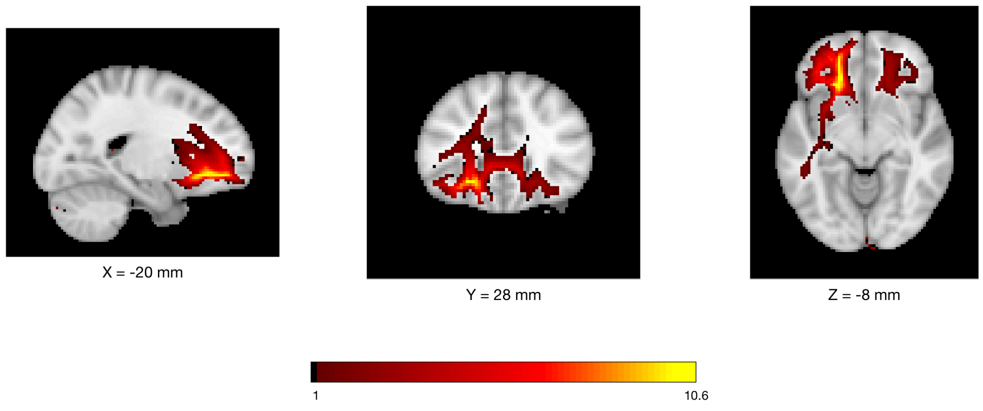


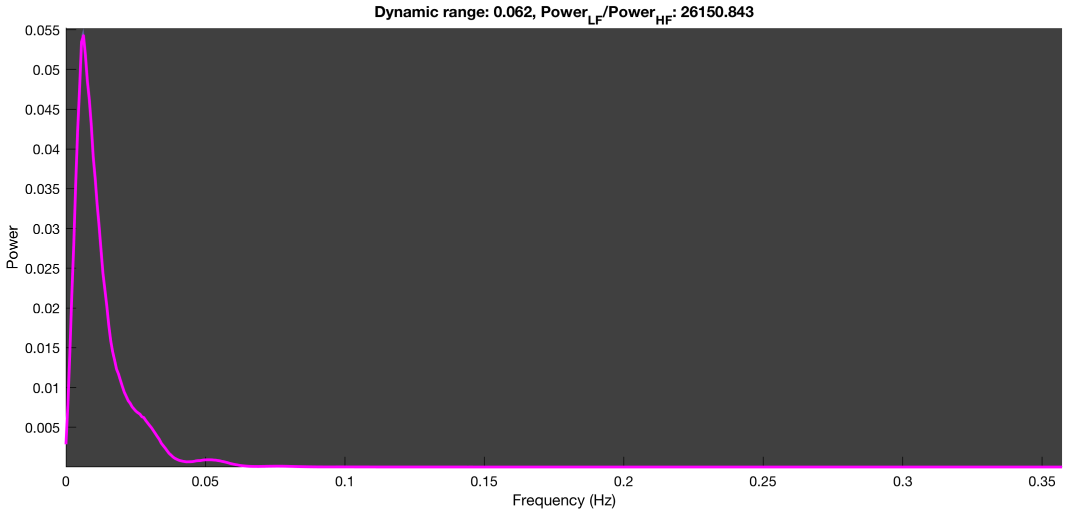

3-1 100n_tmap_component_ica_s1 033

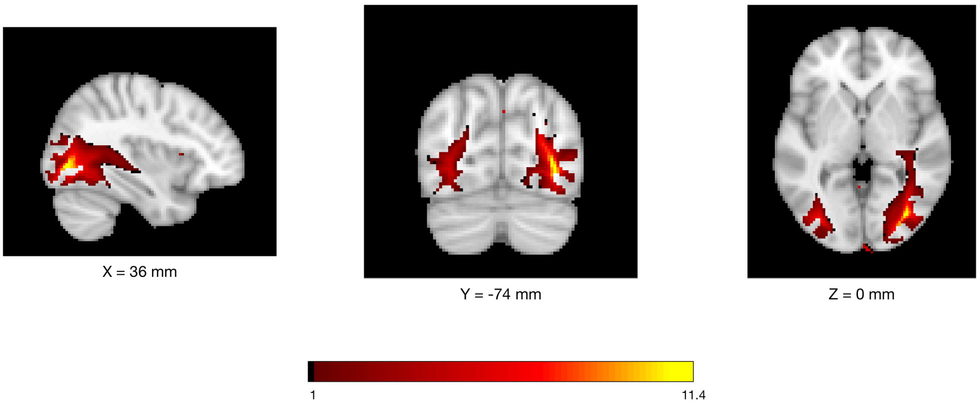


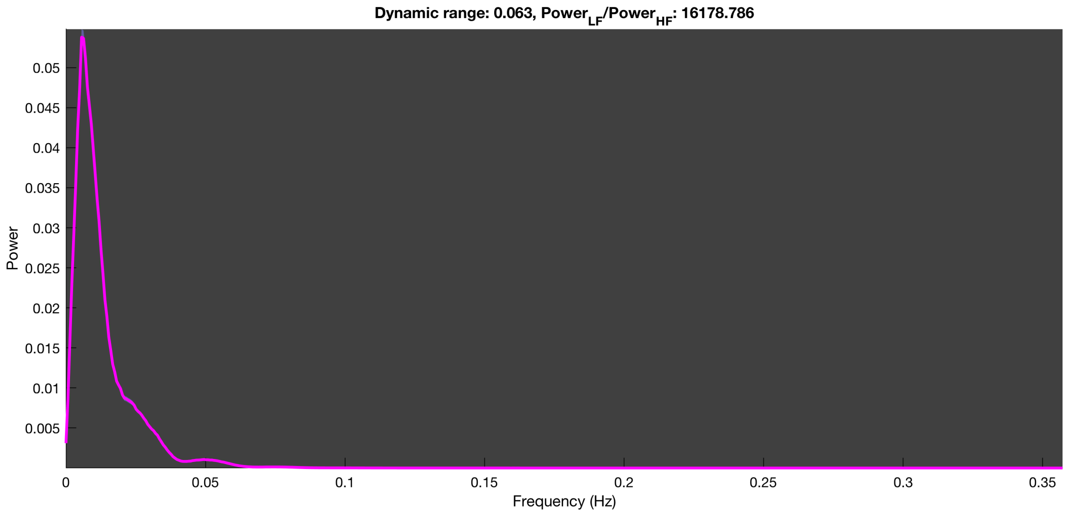

3-1 100n_tmap_component_ica_s1 034

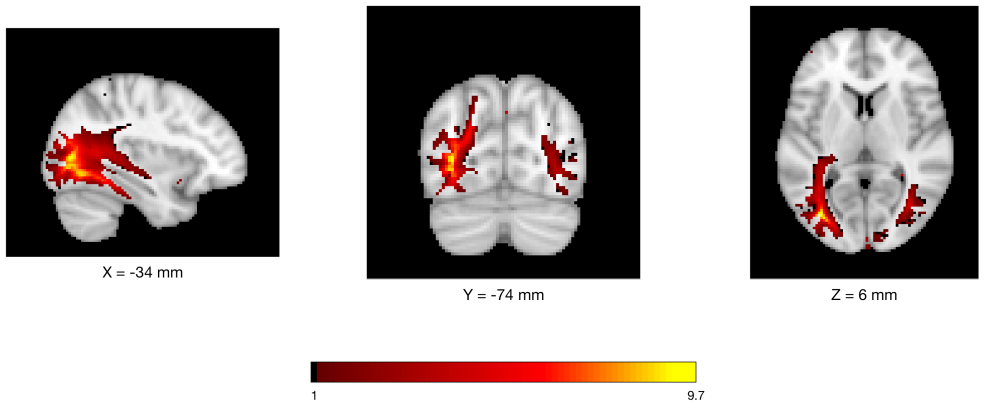


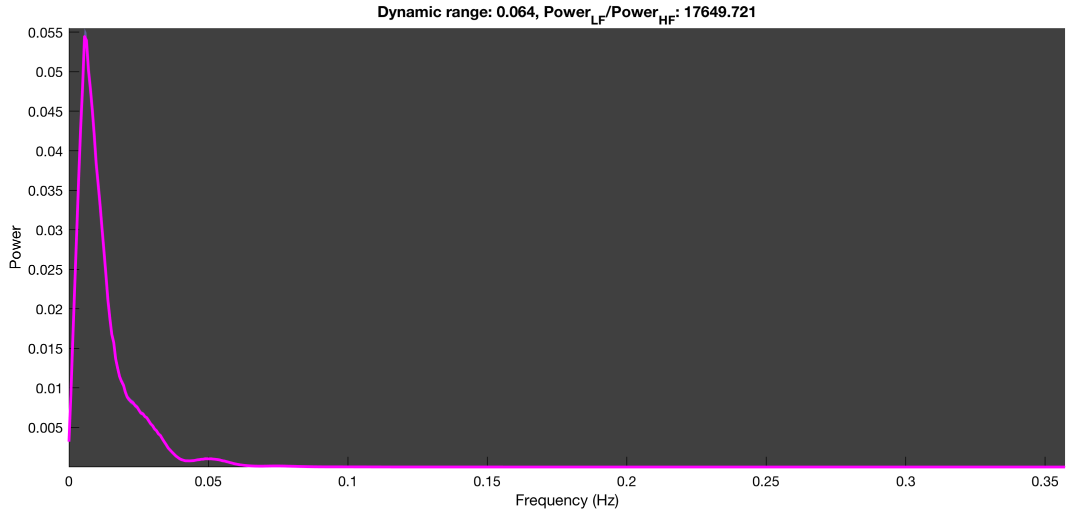

3-1 100n_tmap_component_ica_s1 035

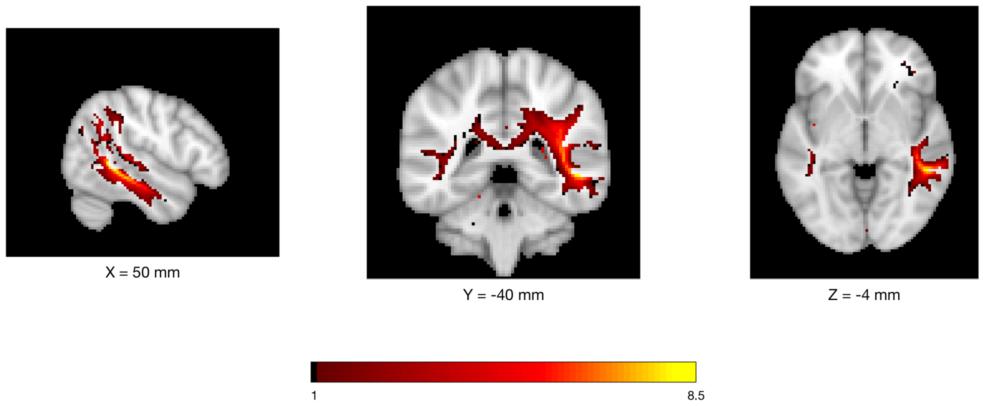


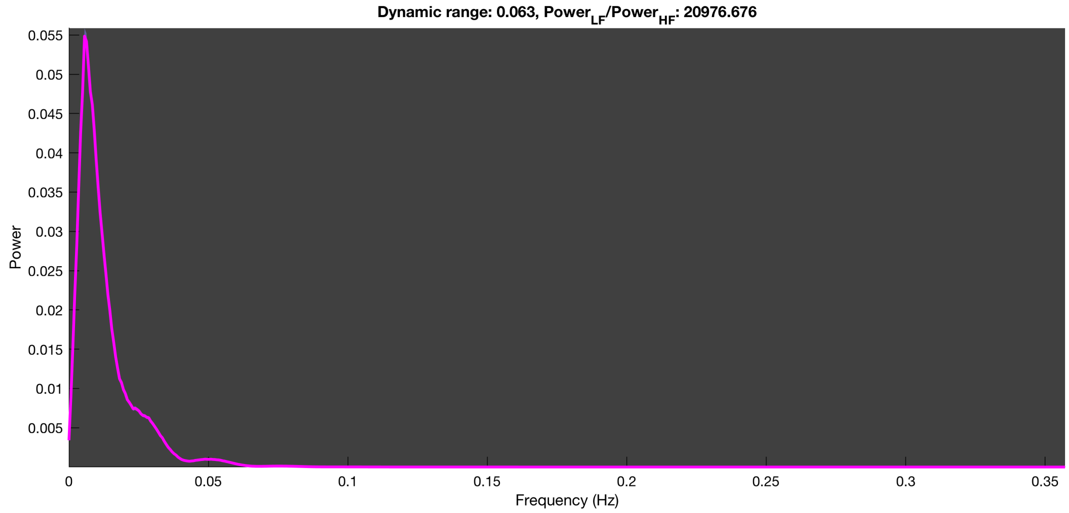

3-1 100n_tmap_component_ica_s1 036

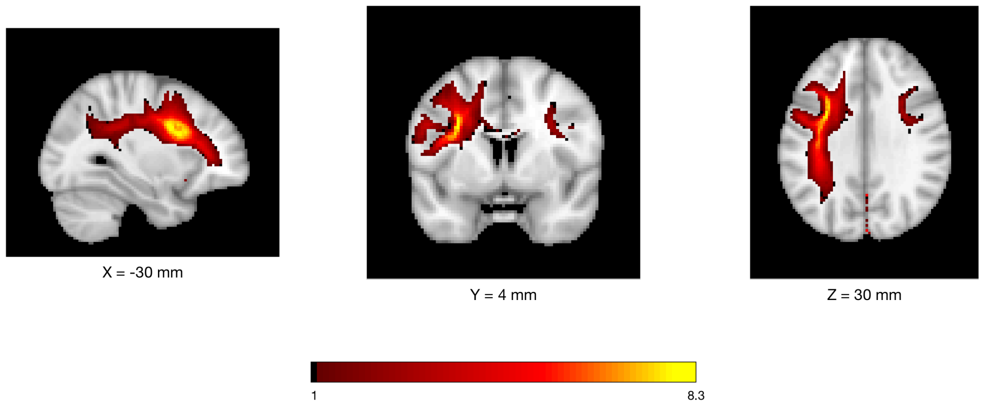


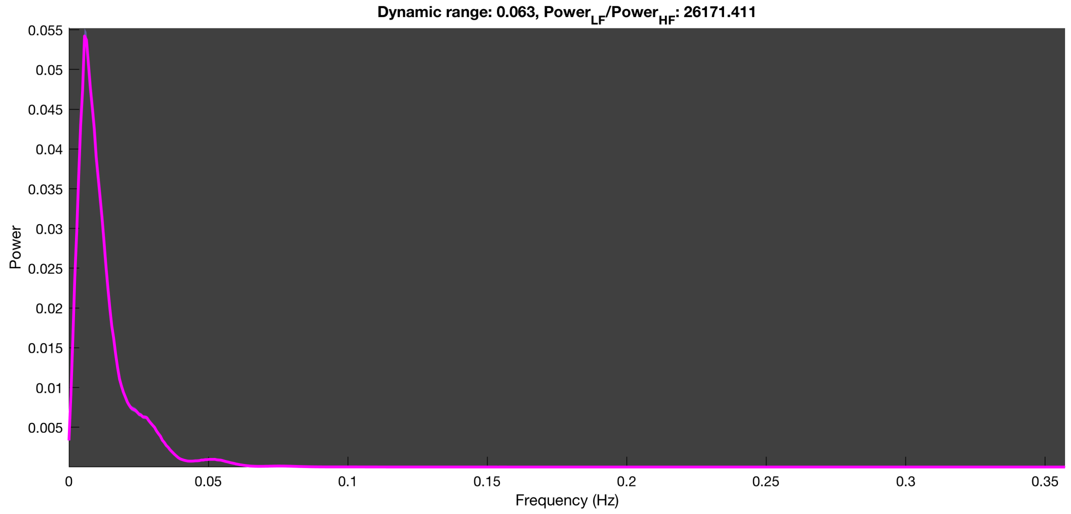

3-1 100n_tmap_component_ica_s1 037

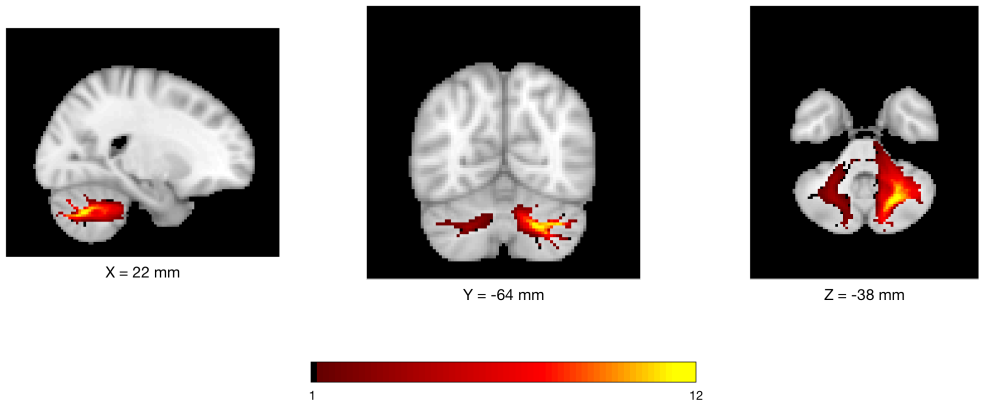


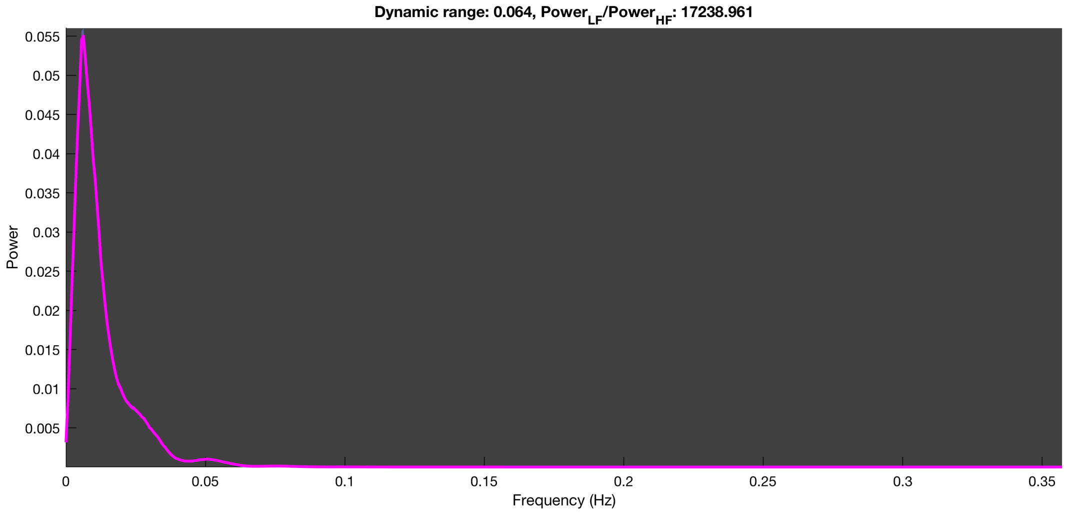

3-1 100n_tmap_component_ica_s1 038

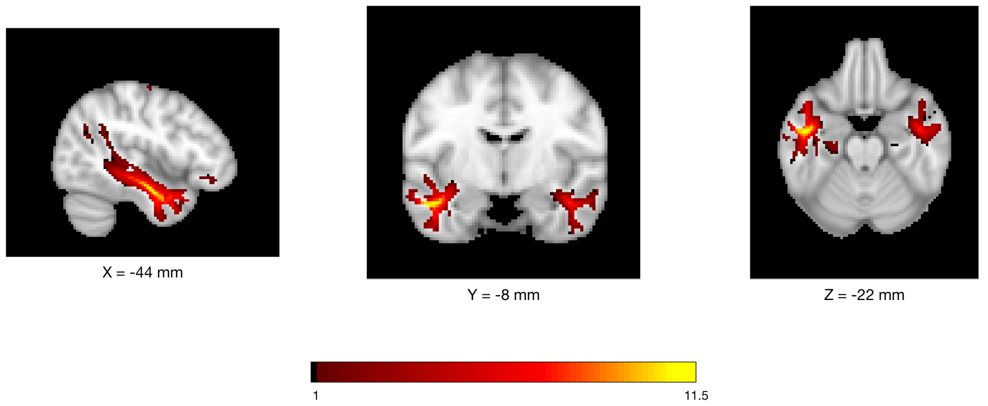


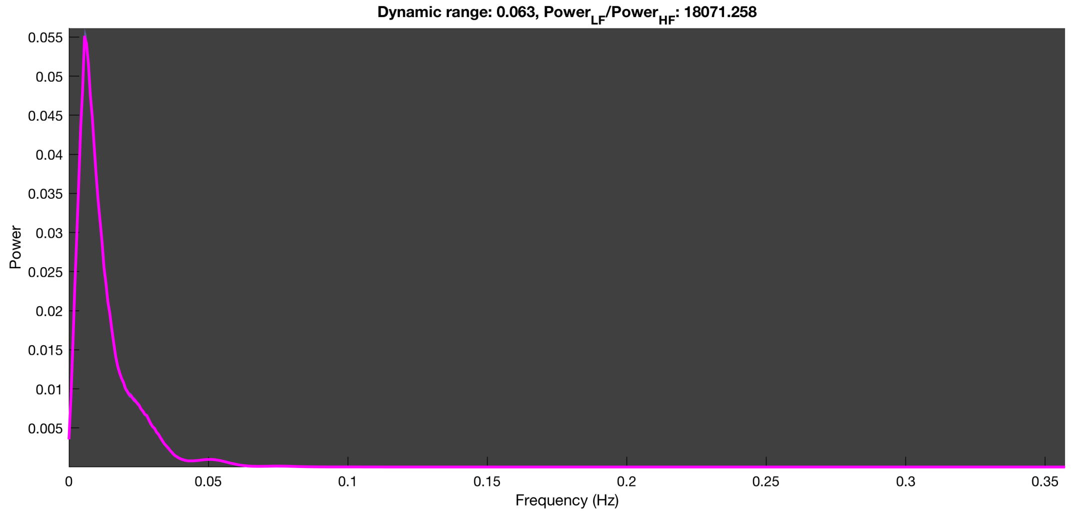

3-1 100n_tmap_component_ica_s1 039

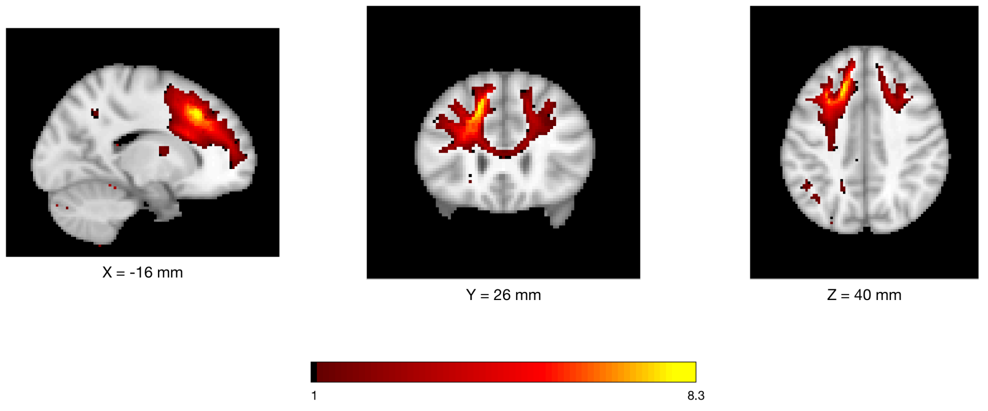


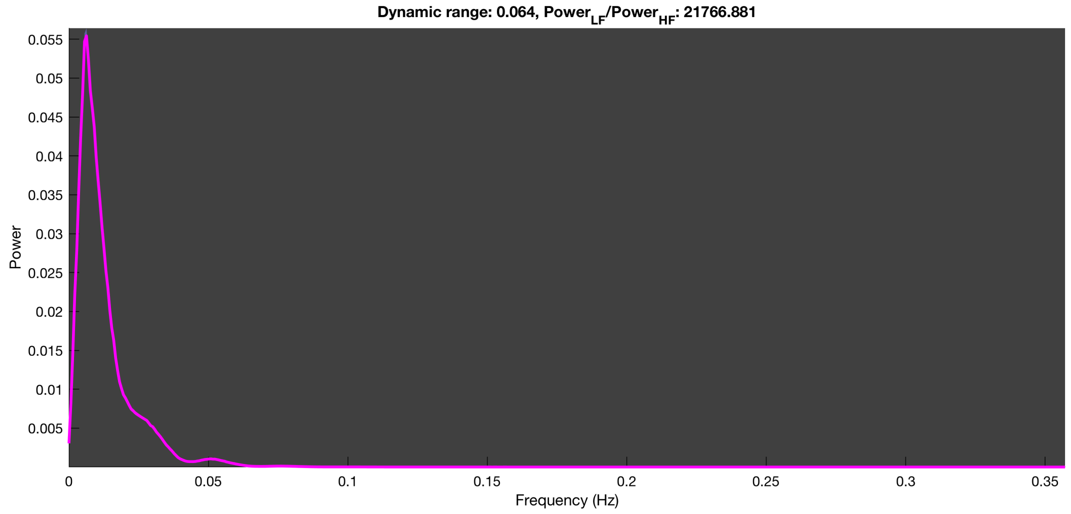

3-1 100n_tmap_component_ica_s1 040

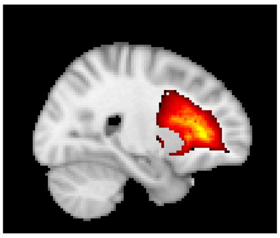


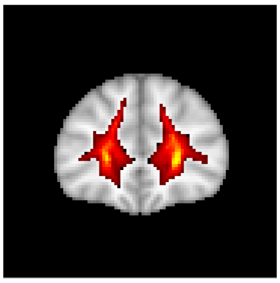


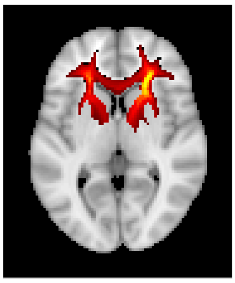


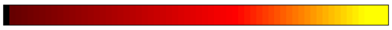


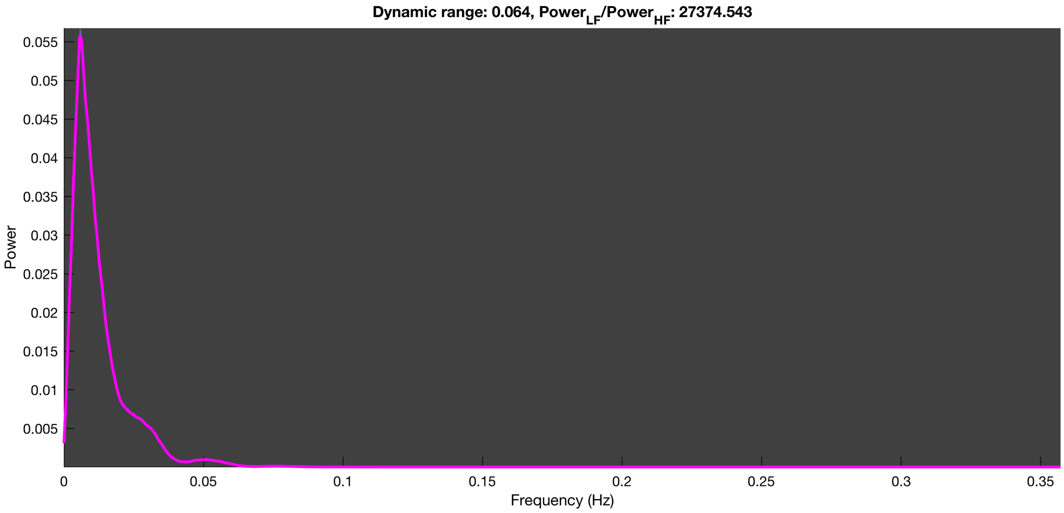

3-1 100n_tmap_component_ica_s1 041

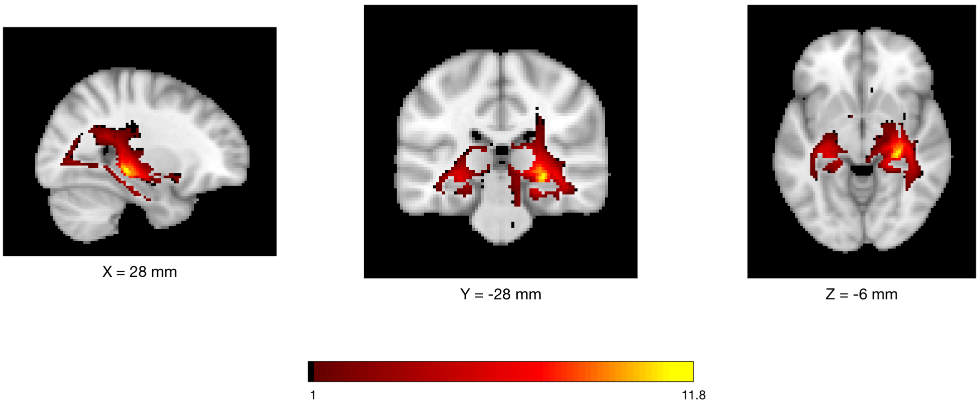


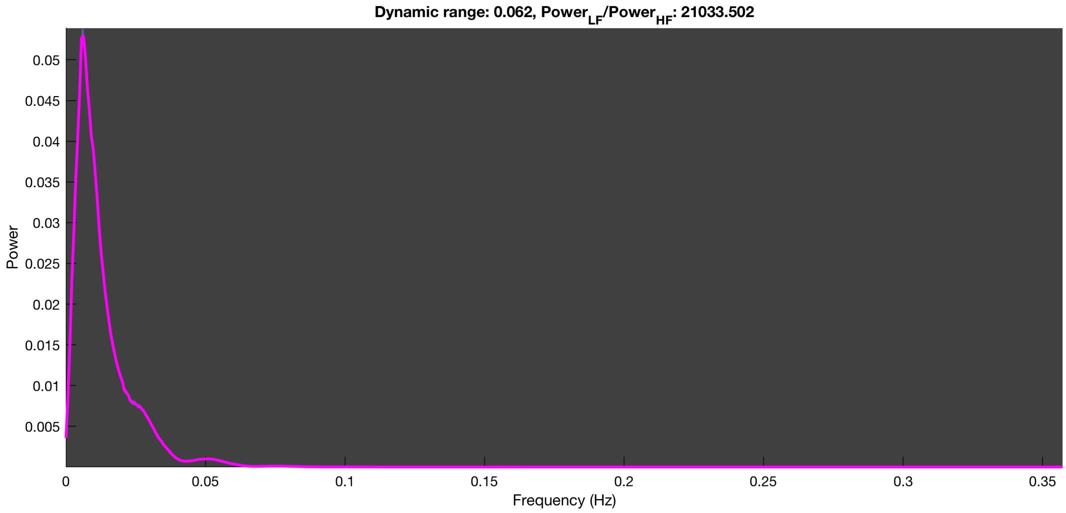

3-1 100n_tmap_component_ica_s1 042

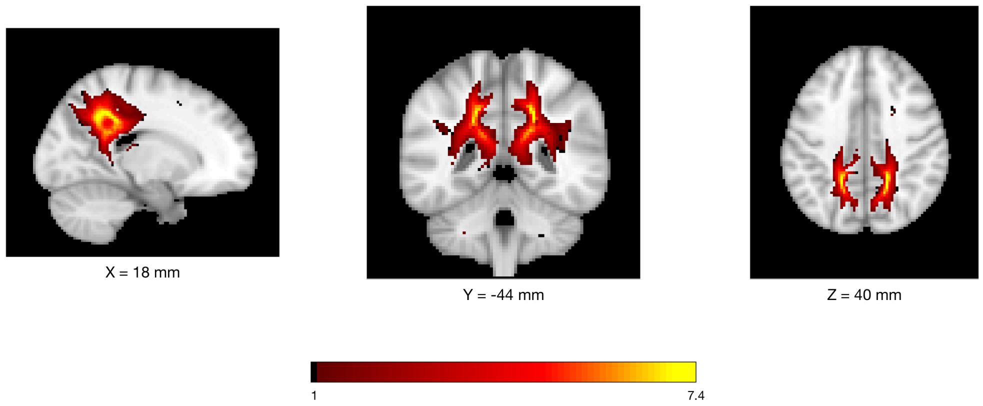


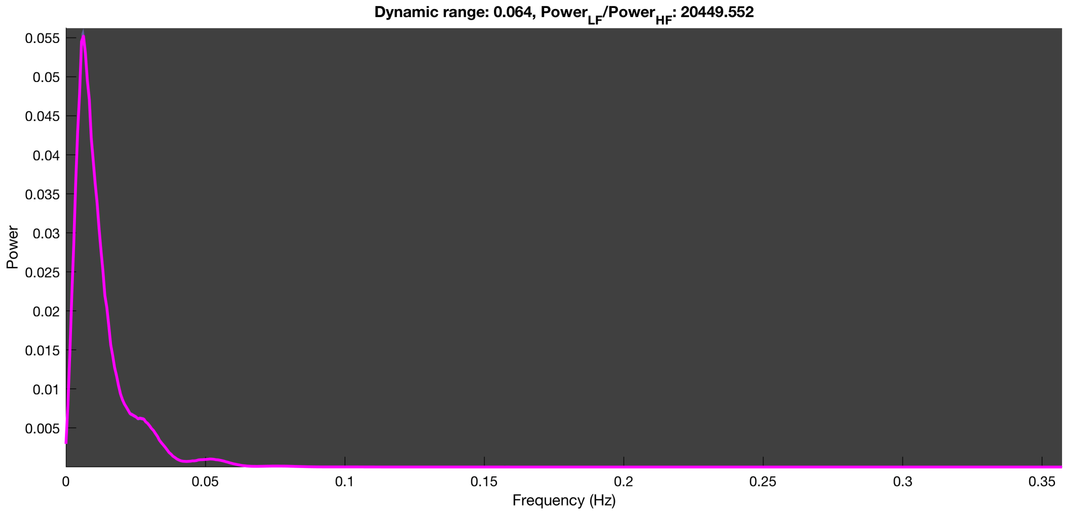

3-1 100n_tmap_component_ica_s1 043

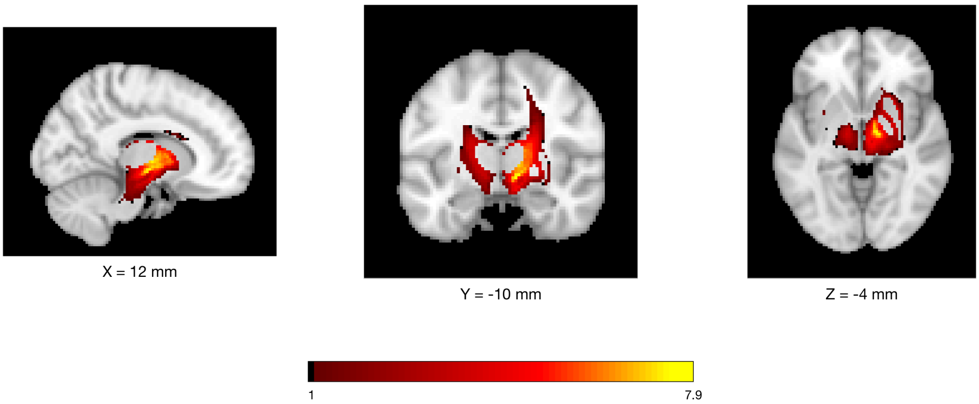


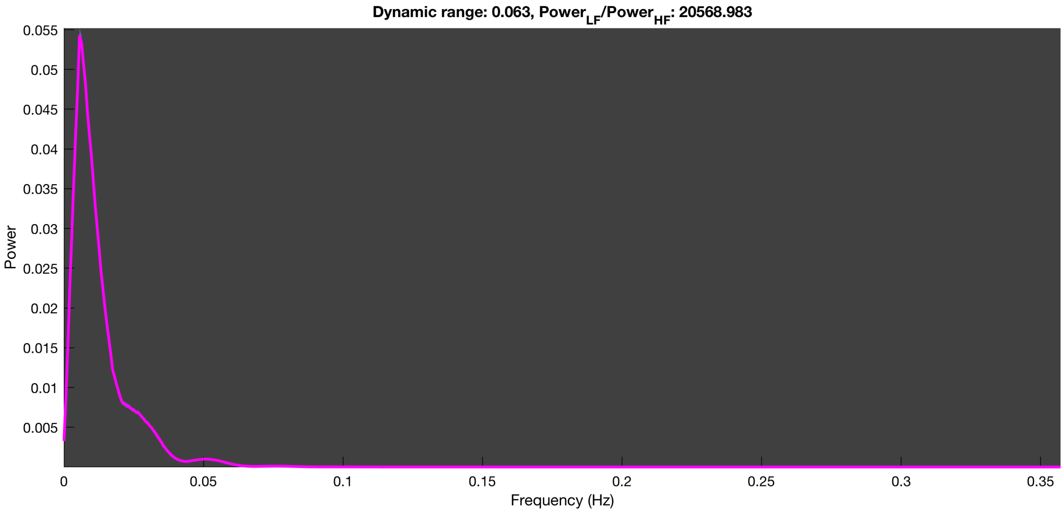

#### 3-1 100n_tmap_component_ica_s1 044

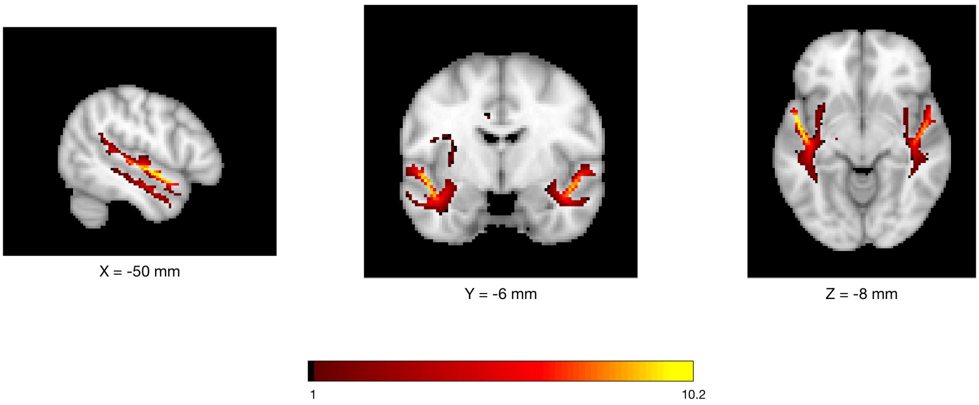


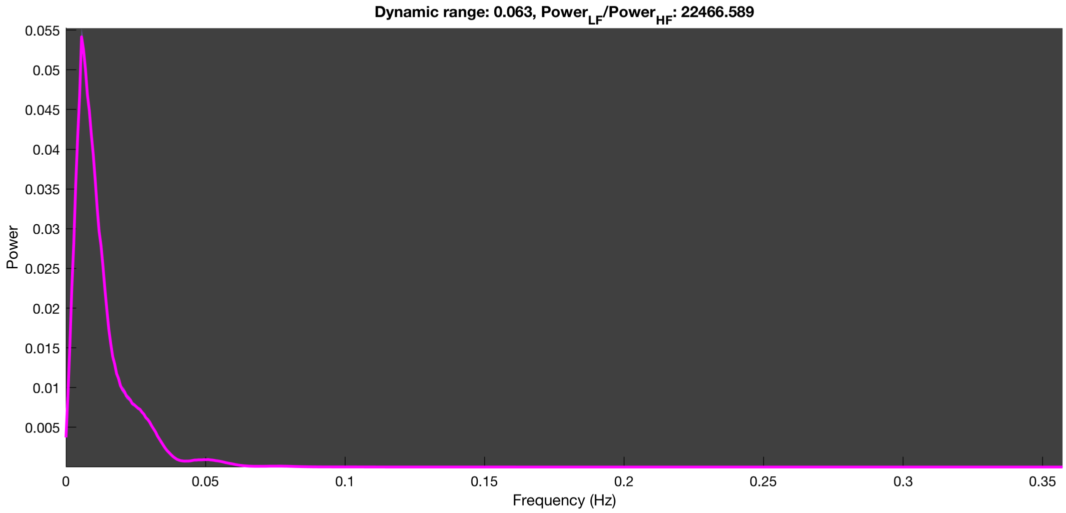

3-1 100n_tmap_component_ica_s1 045

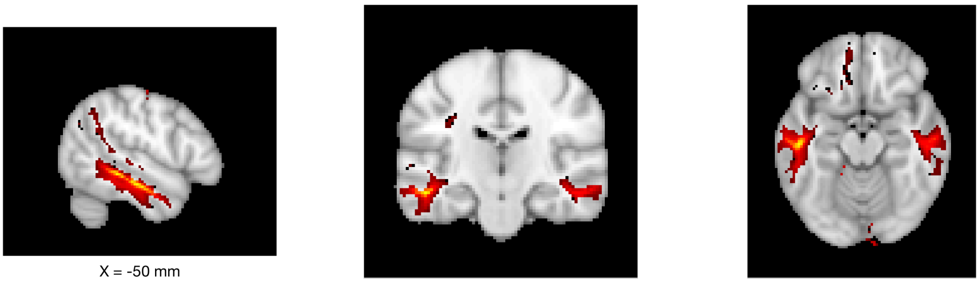

X = -50 mm

Y = -20 mm

Z = -16 mm

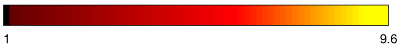


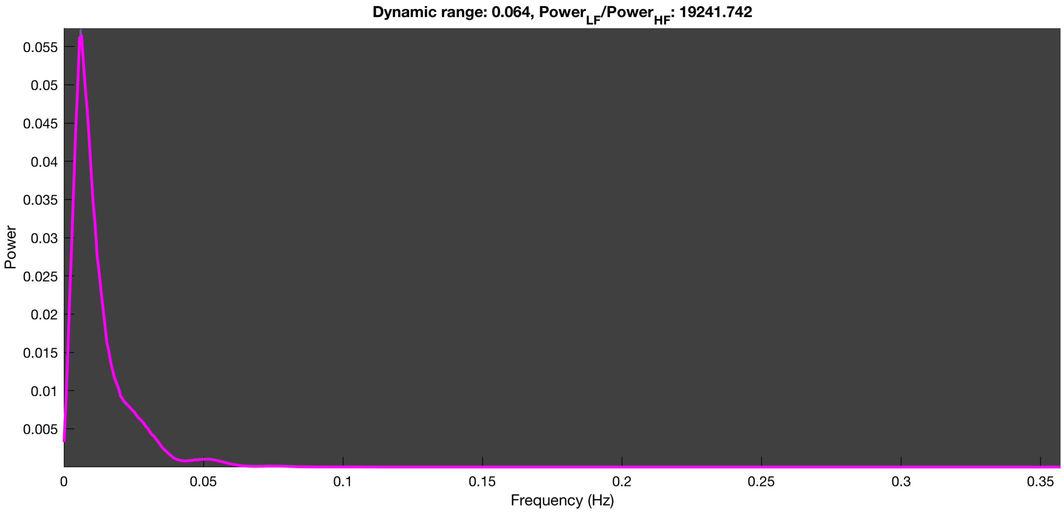

3-1 100n_tmap_component_ica_s1 046

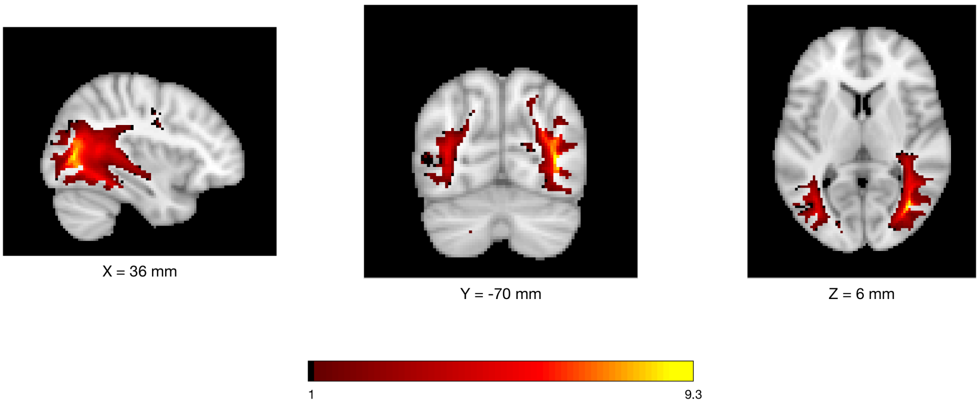


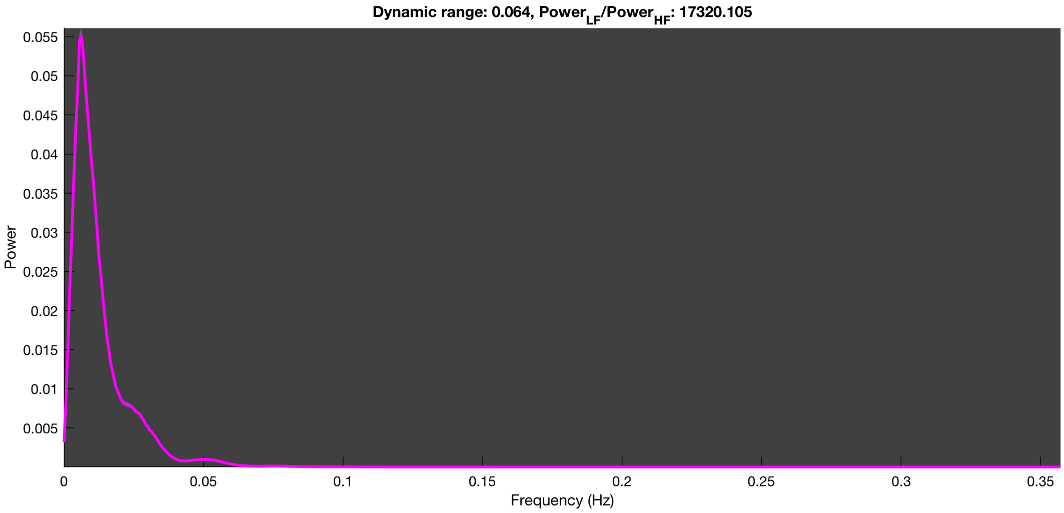

3-1 100n_tmap_component_ica_s1 047

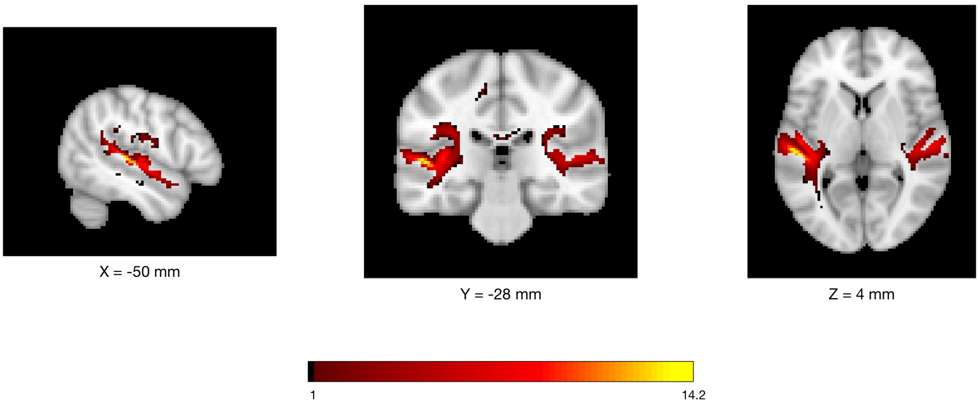


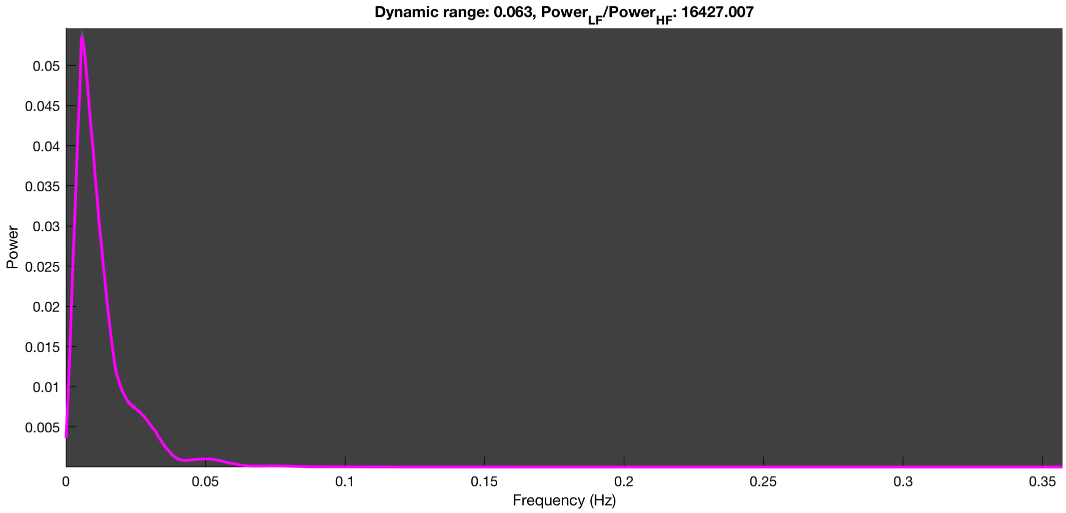

#### 3-1 100n_tmap_component_ica_s1 048

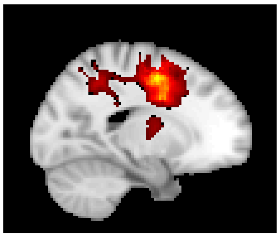

X = -18 mm

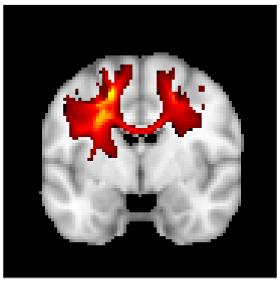

Y = -4 mm

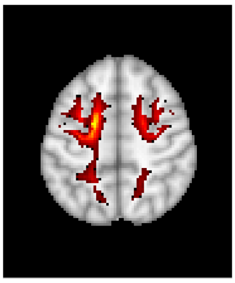

Z = 50 mm

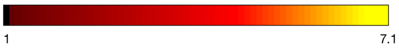


**Dynamic range: 0.063, $Power_{LF}/Power_{HF}$: 24301.131**

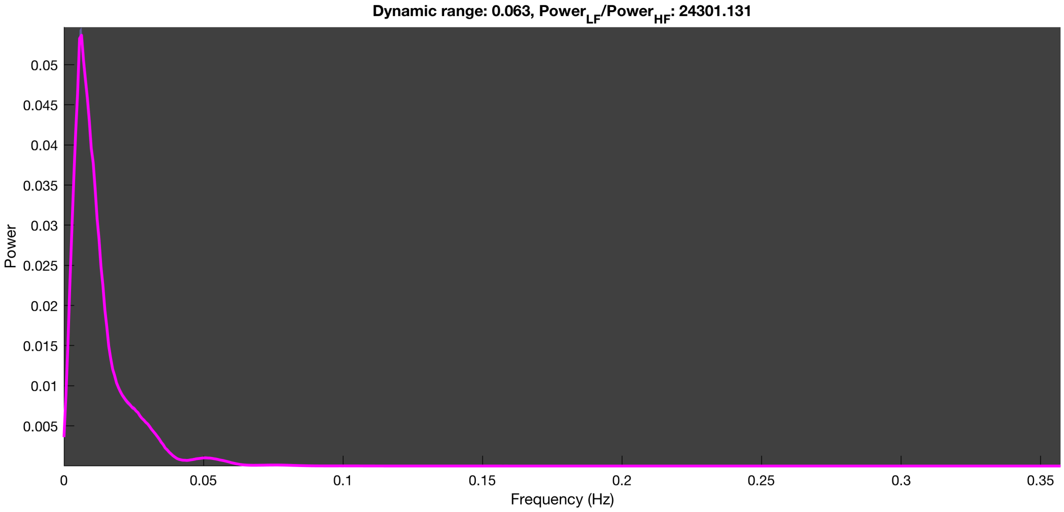

3-1 100n_tmap_component_ica_s1 049

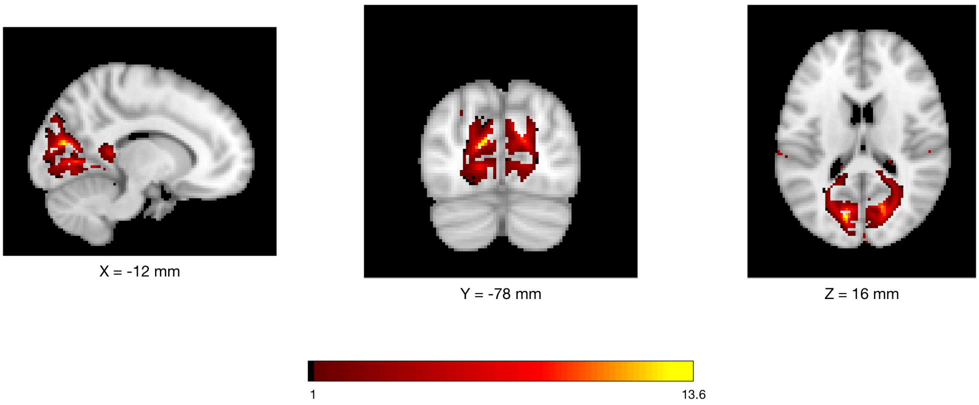


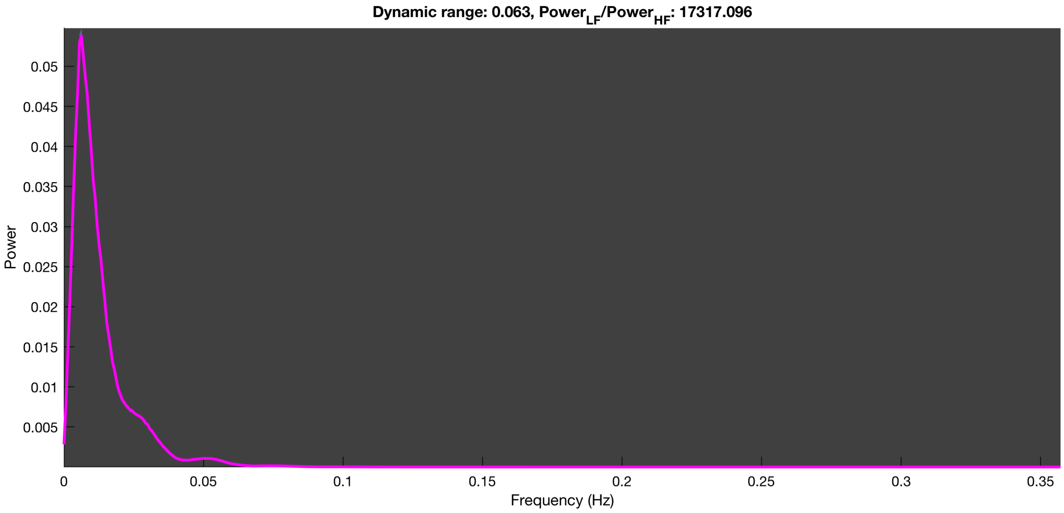

3-1 100n_tmap_component_ica_s1 050

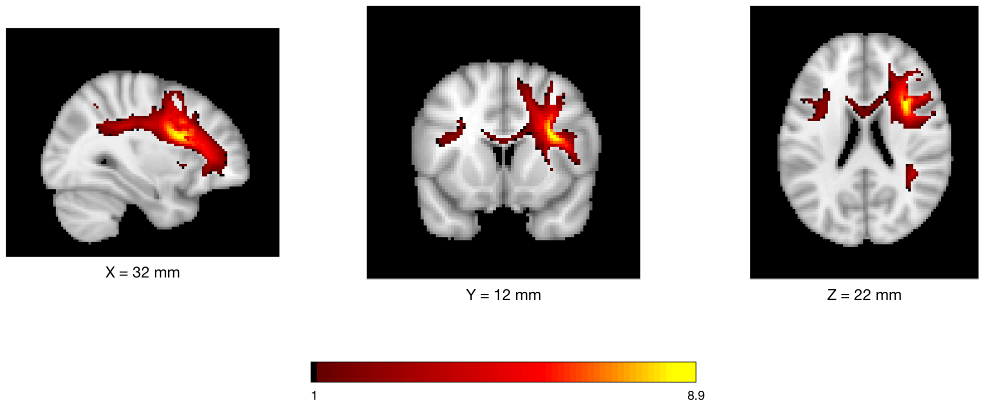


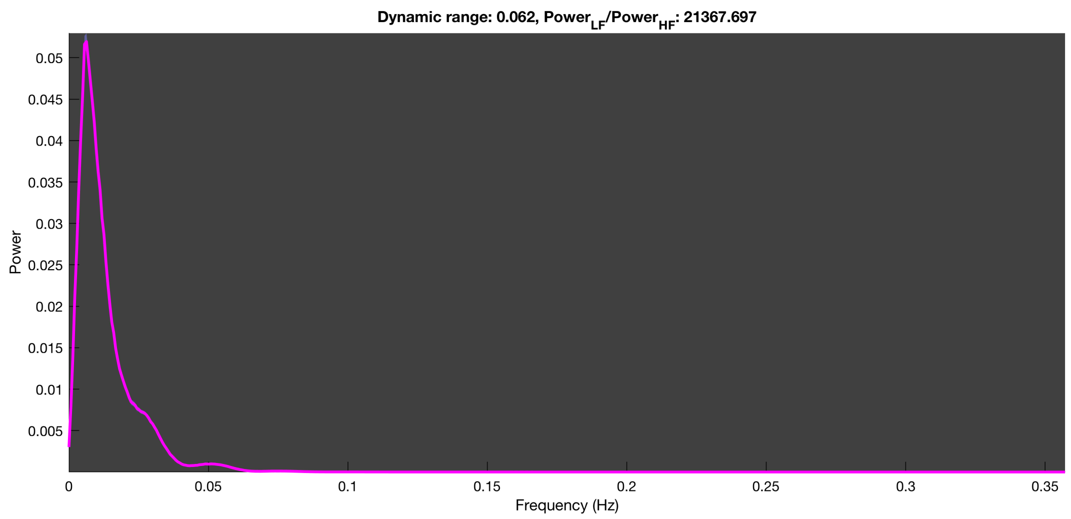

3-1 100n_tmap_component_ica_s1 051

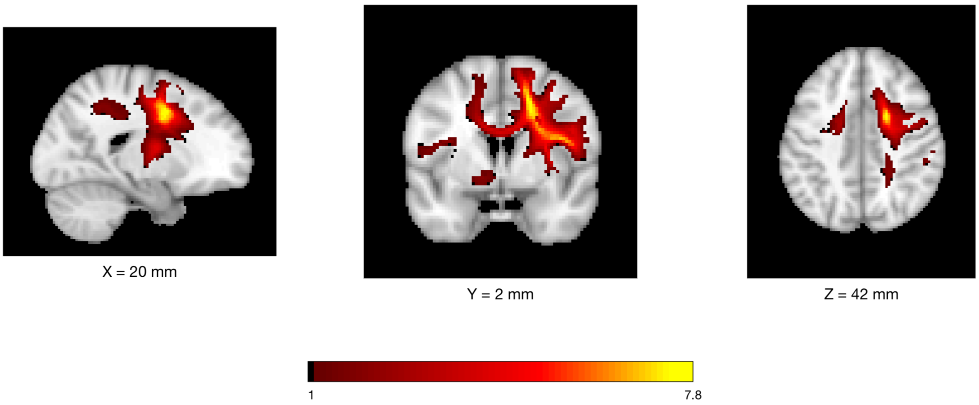


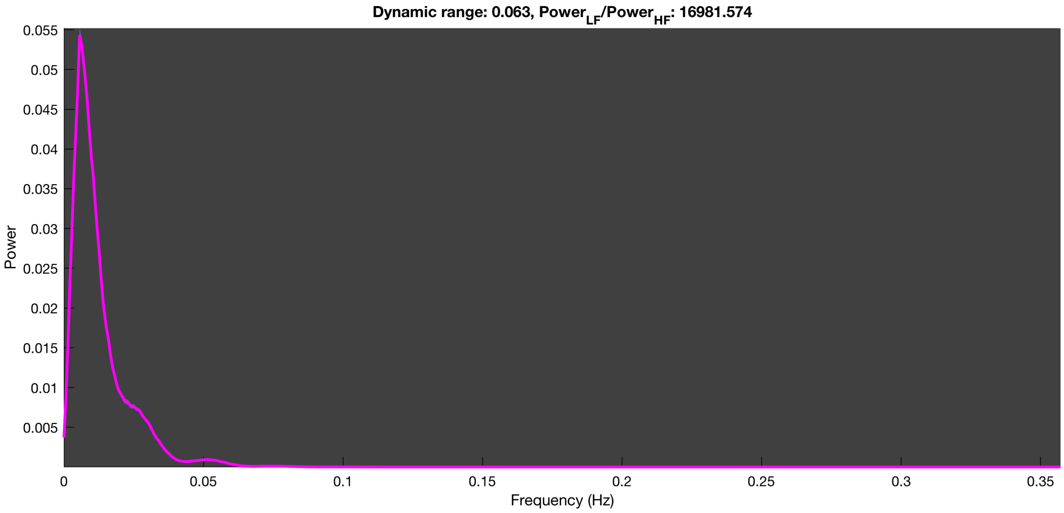

3-1 100n_tmap_component_ica_s1 052

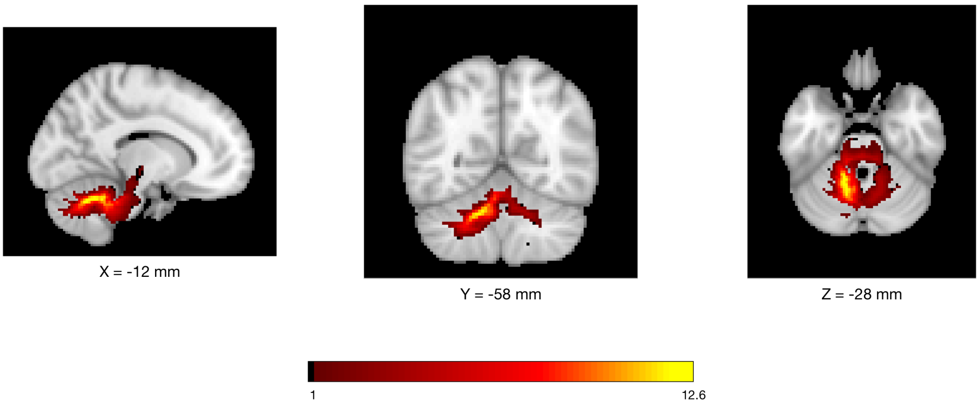


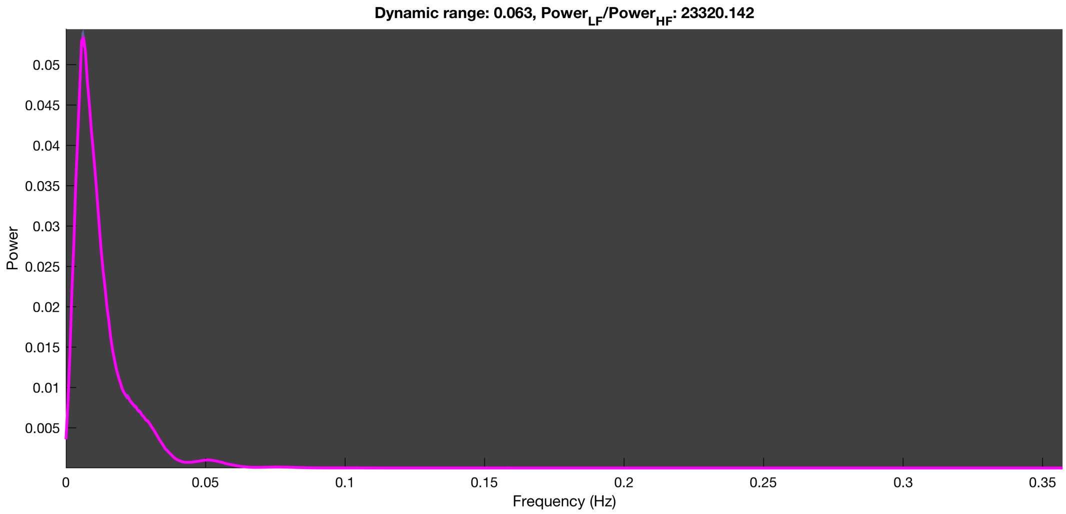

3-1 100n_tmap_component_ica_s1 053

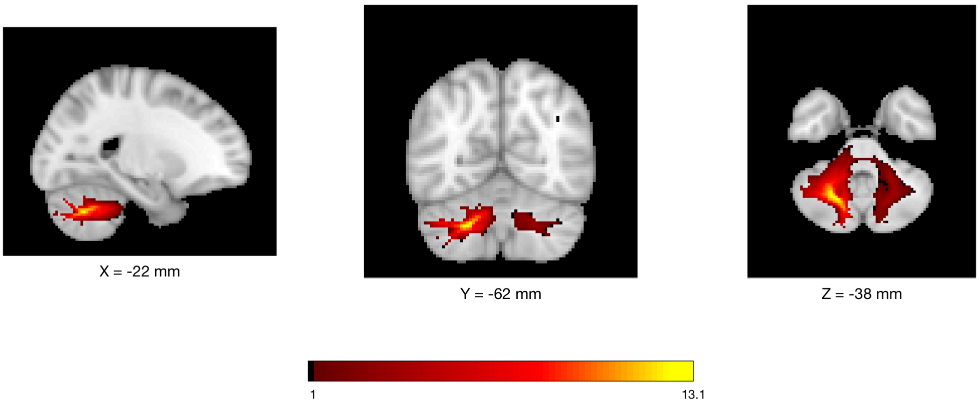


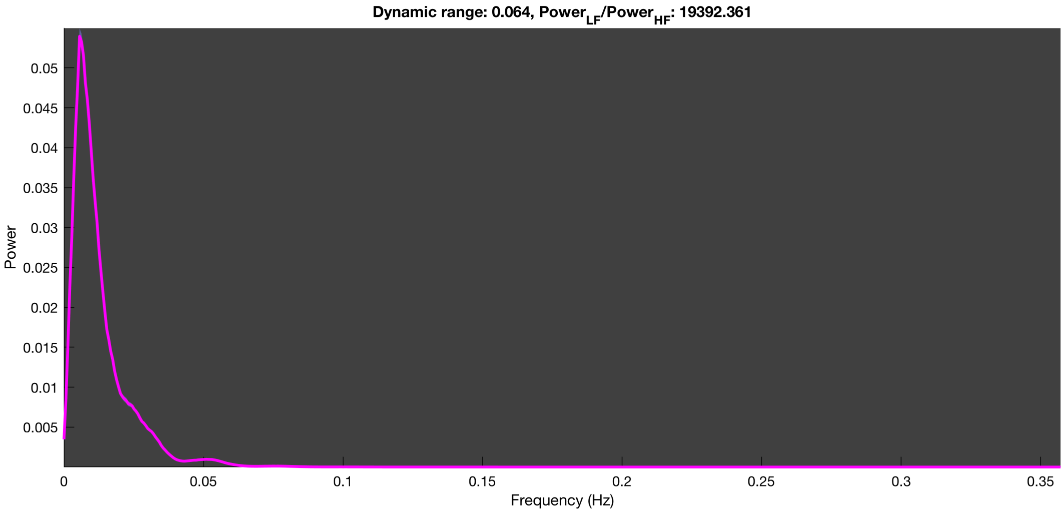

3-1 100n_tmap_component_ica_s1 054

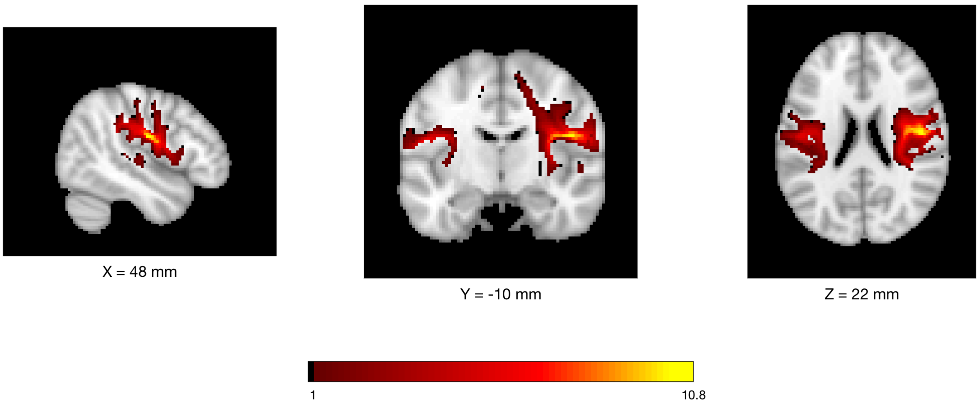


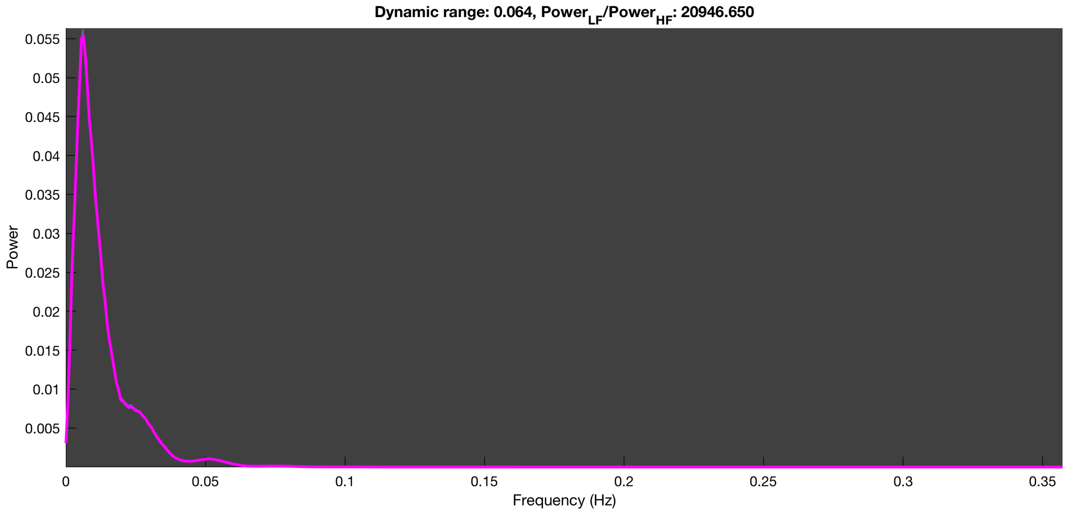

3-1 100n_tmap_component_ica_s1 055

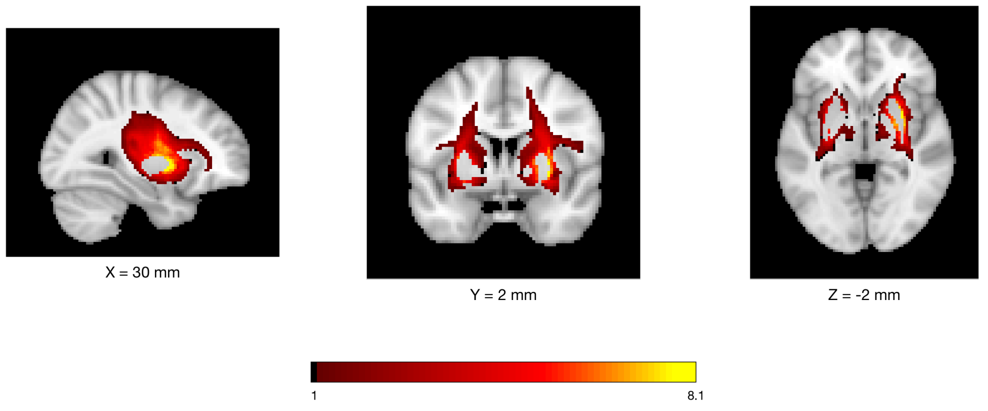


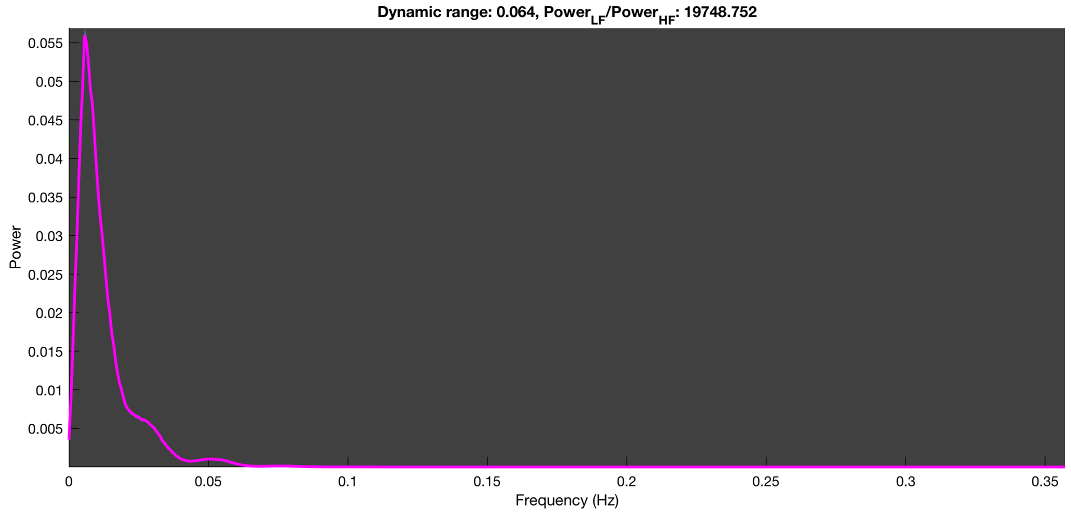

3-1 100n_tmap_component_ica_s1 056

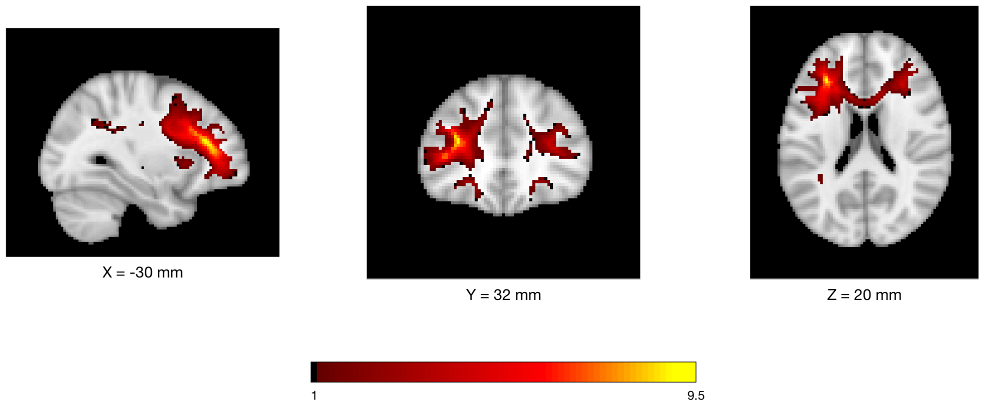


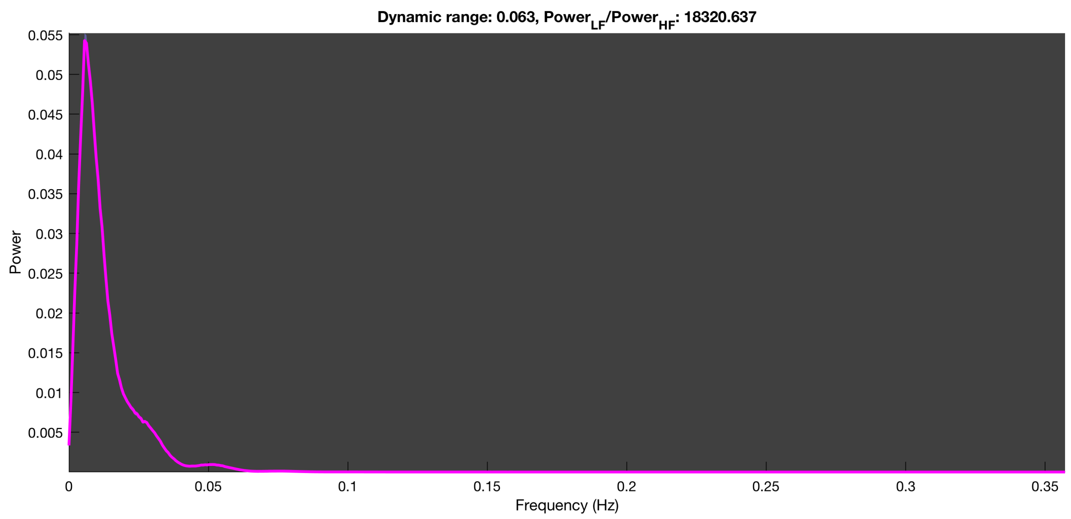

3-1 100n_tmap_component_ica_s1 057

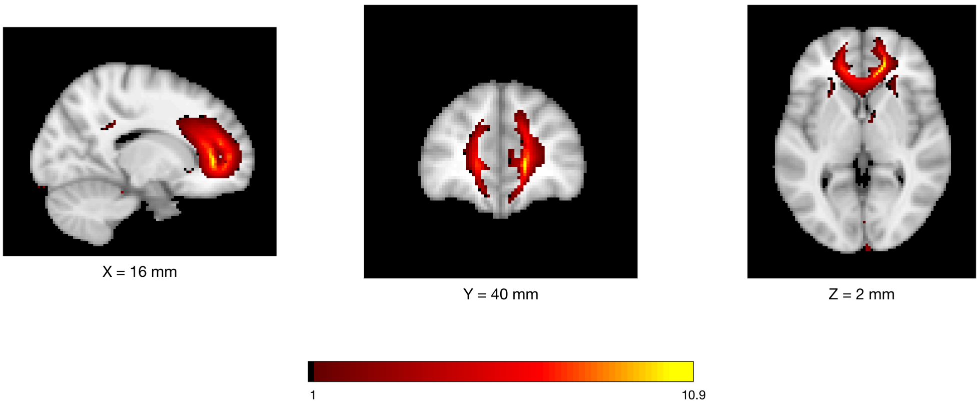


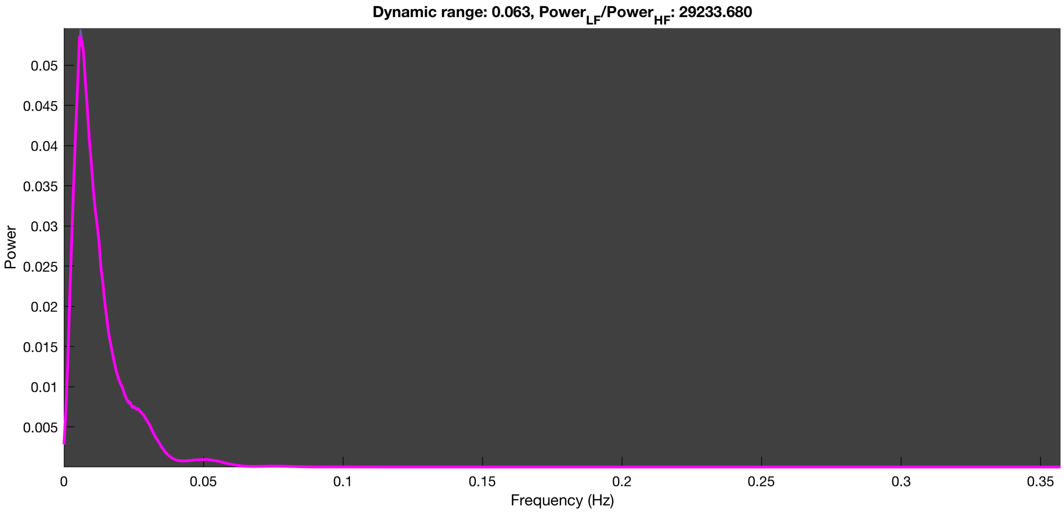

3-1 100n_tmap_component_ica_s1 058

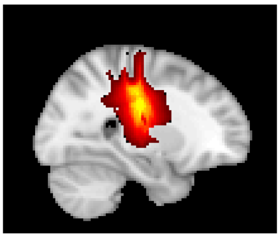

X = 26 mm

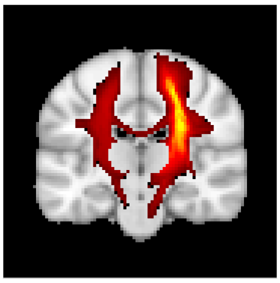

Y = -20 mm

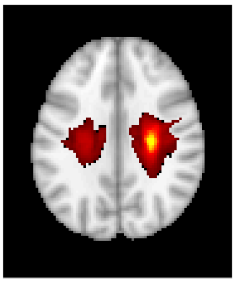

Z = 30 mm

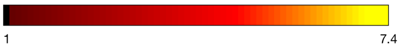


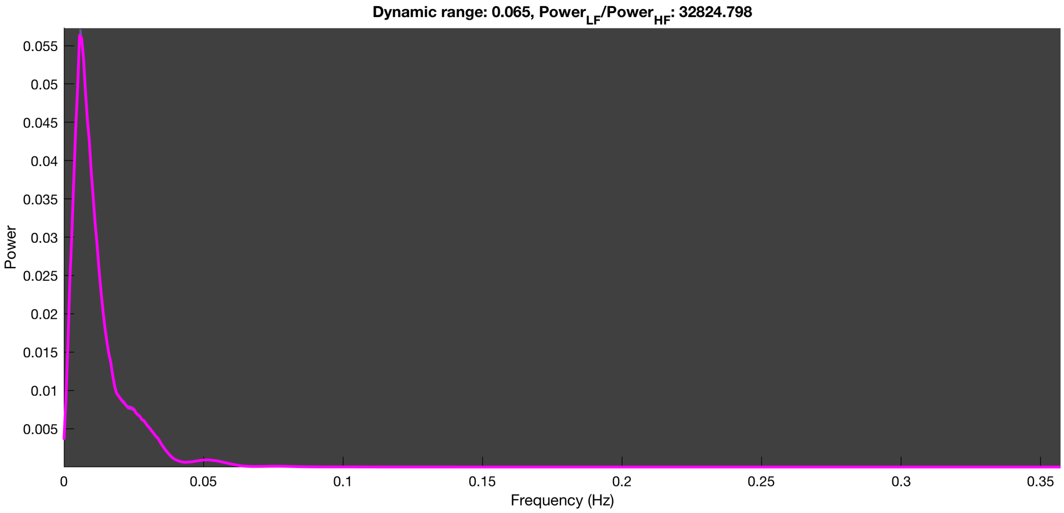

3-1 100n_tmap_component_ica_s1 059

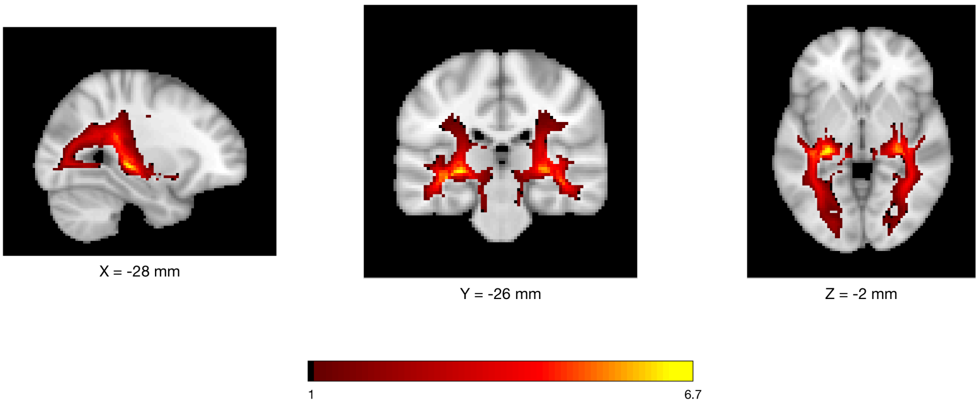


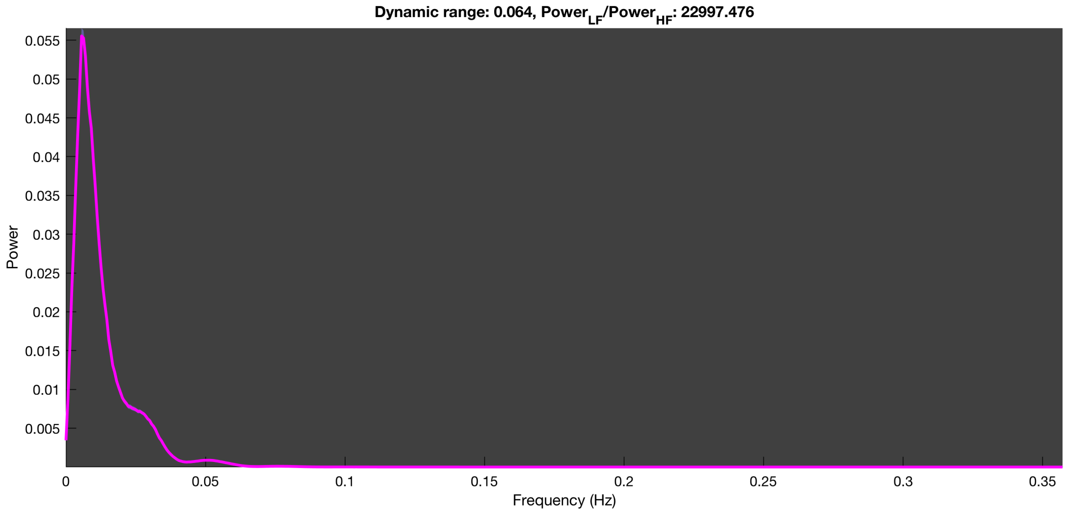

3-1 100n_tmap_component_ica_s1 060

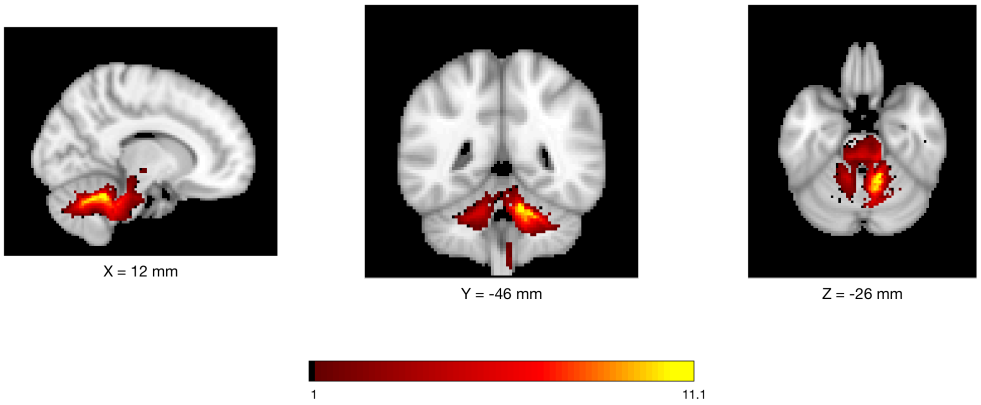


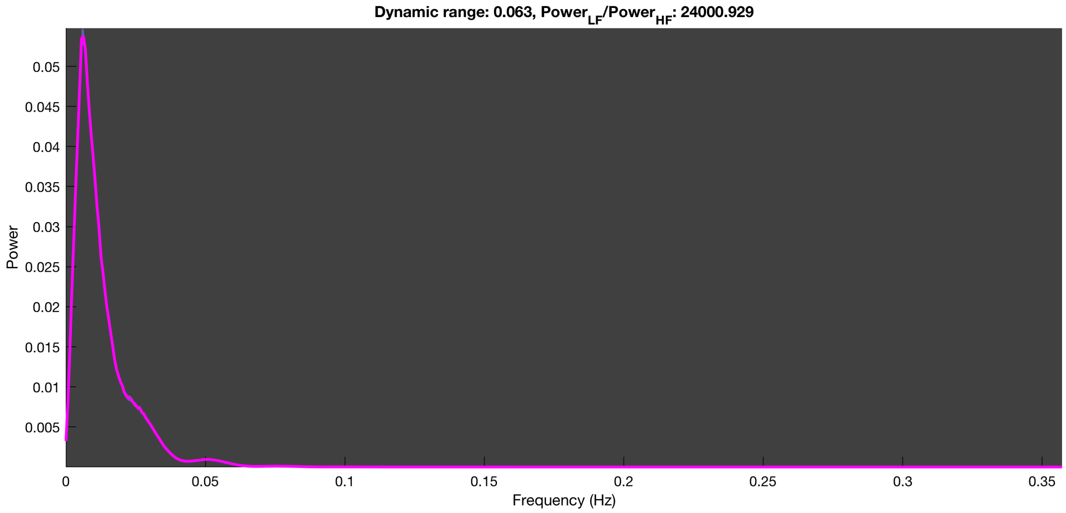

3-1 100n_tmap_component_ica_s1 061

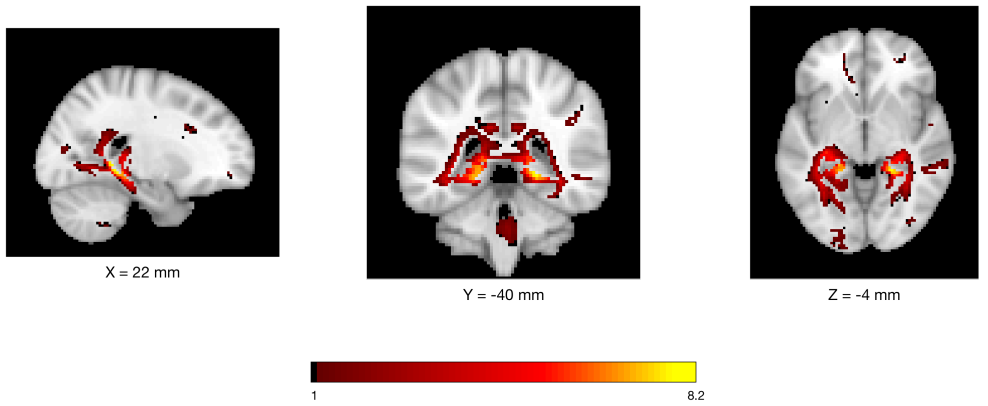


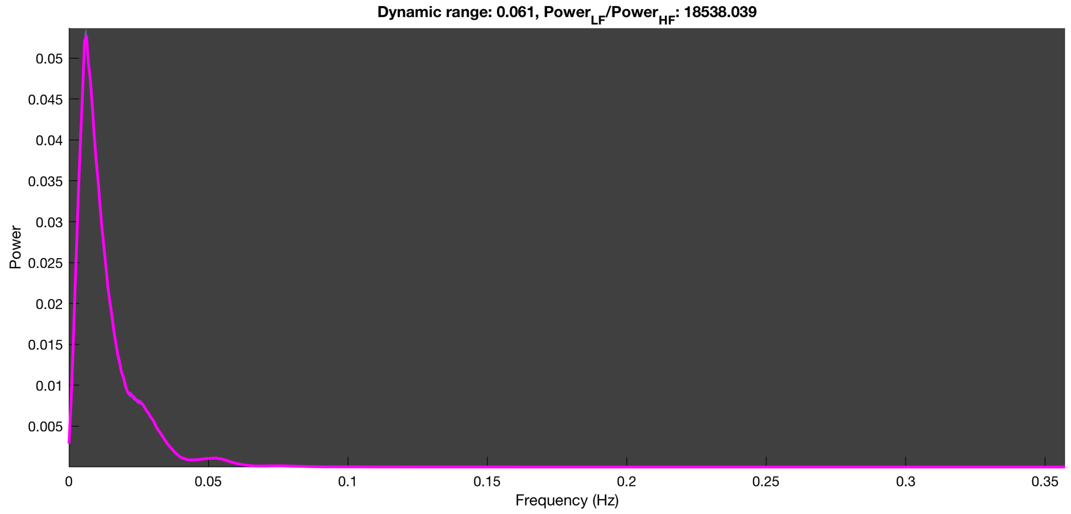

3-1 100n_tmap_component_ica_s1 062

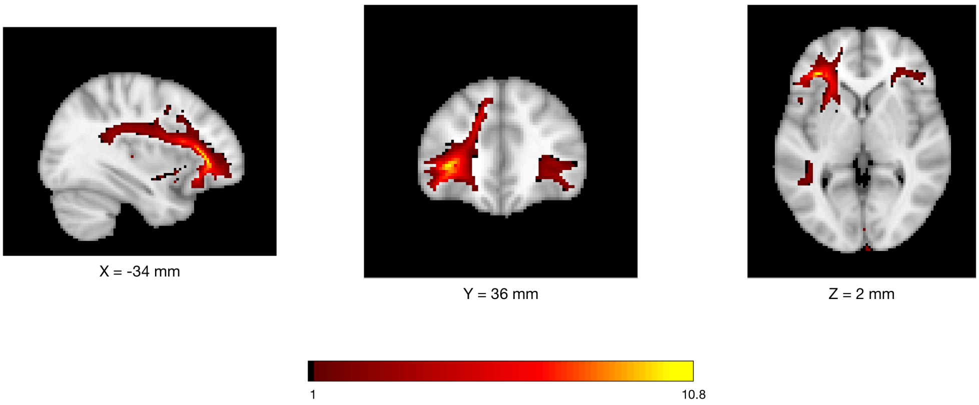


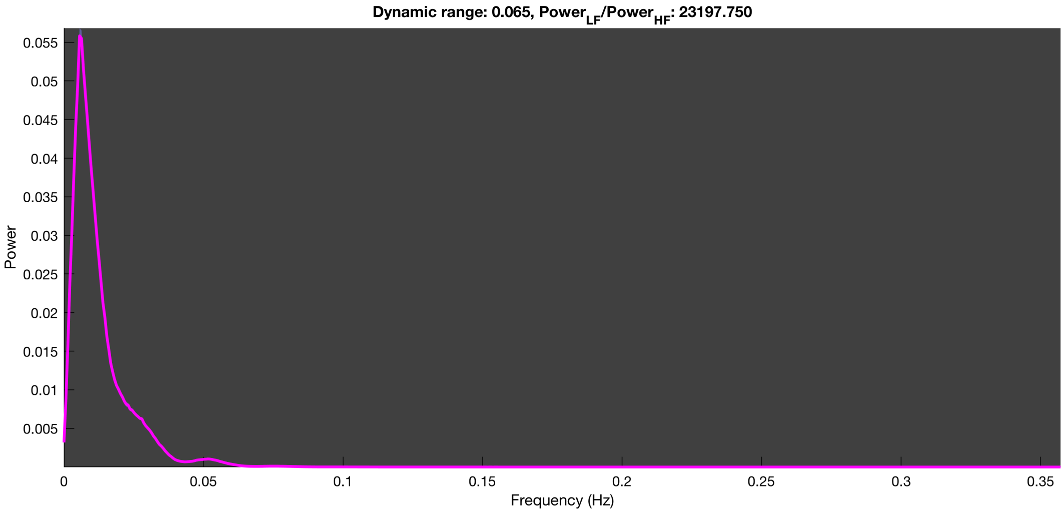

3-1 100n_tmap_component_ica_s1 063

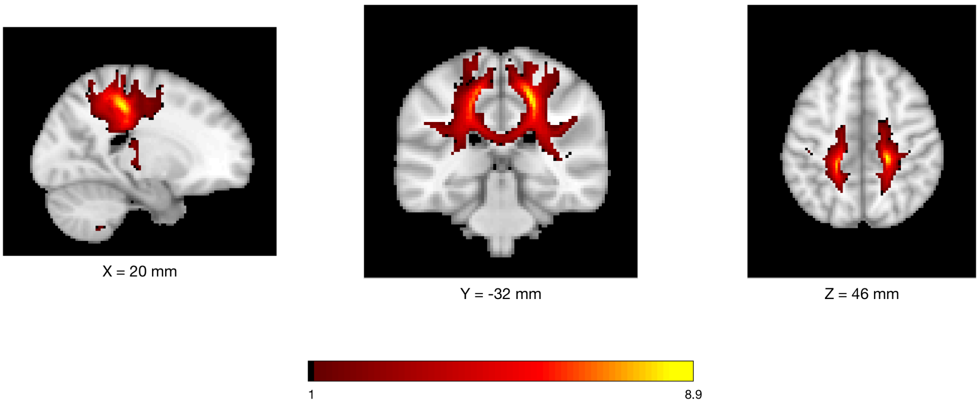


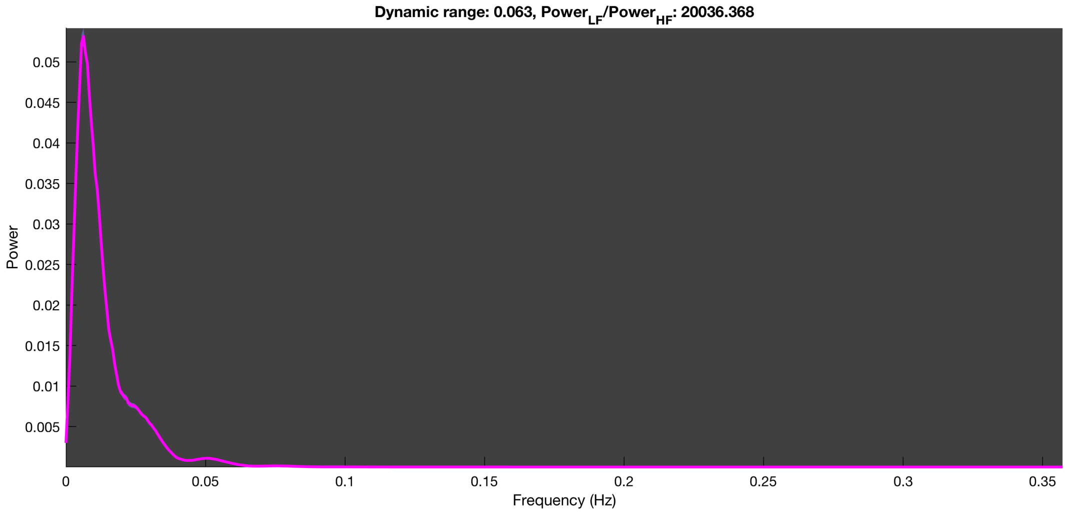

3-1 100n_tmap_component_ica_s1 064

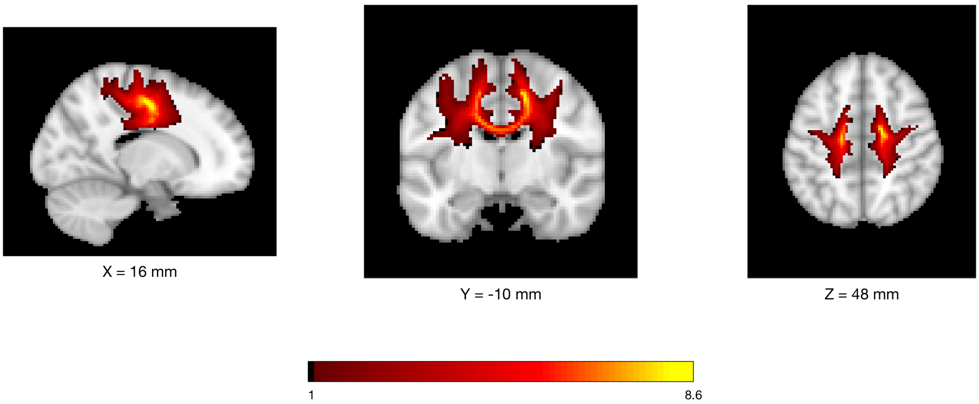


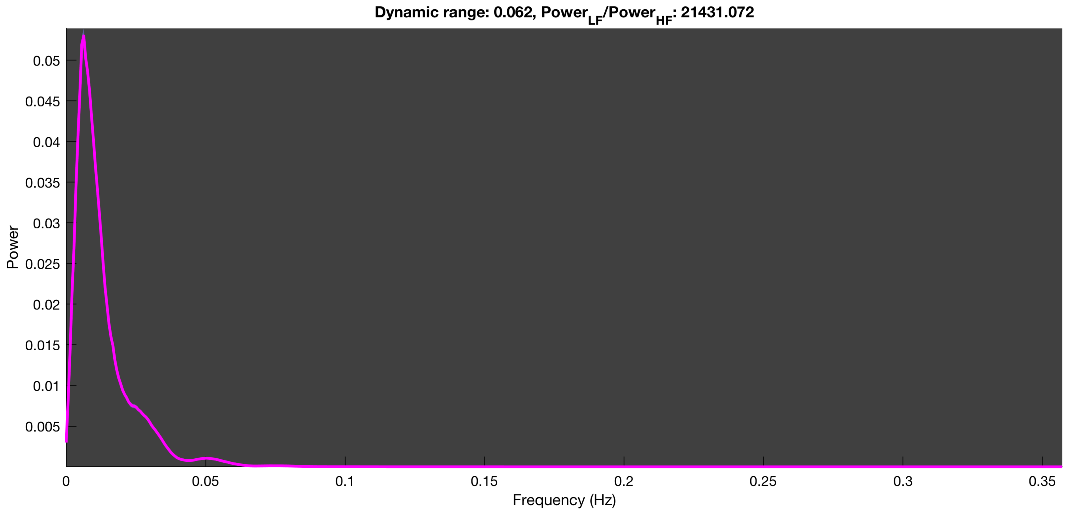

3-1 100n_tmap_component_ica_s1 065

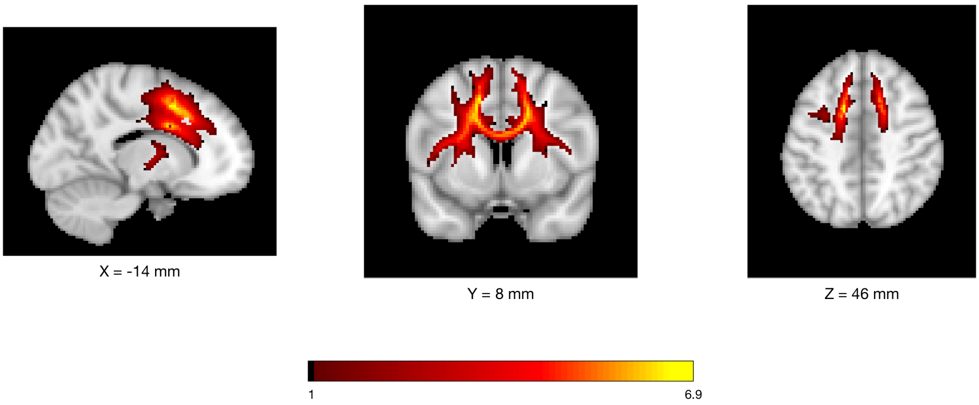


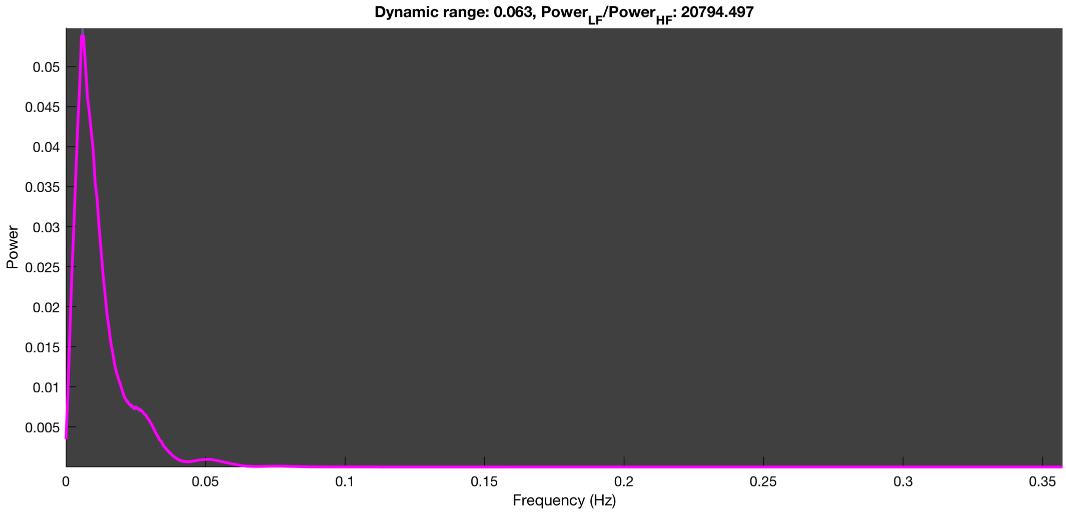

3-1 100n_tmap_component_ica_s1 066

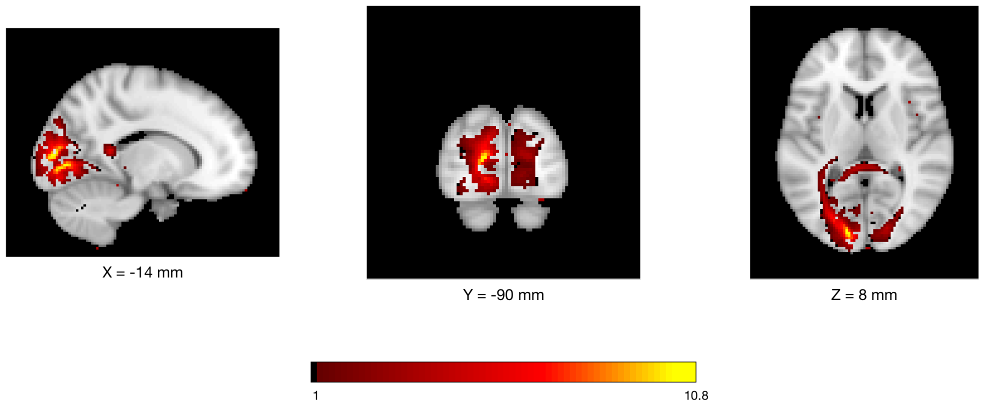


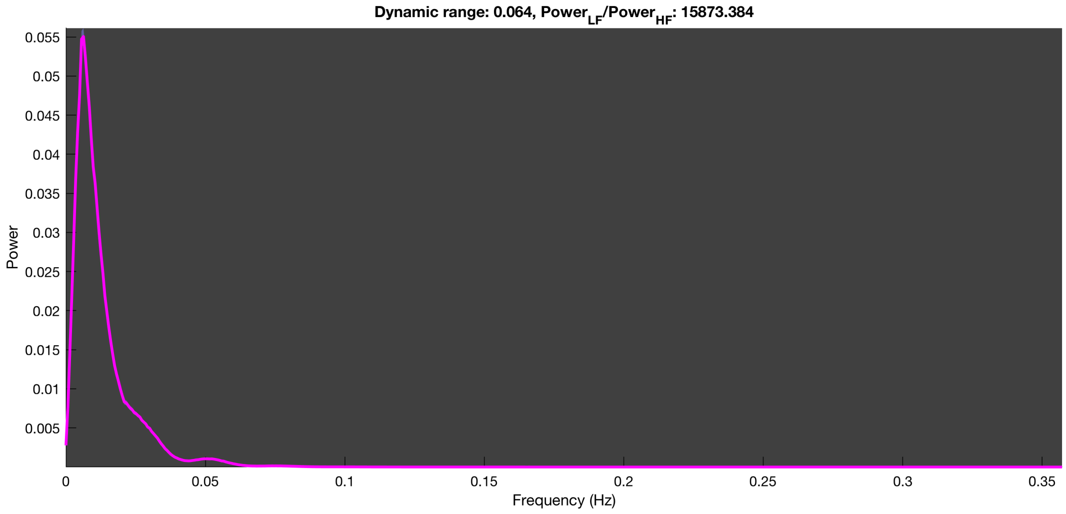

3-1 100n_tmap_component_ica_s1 067

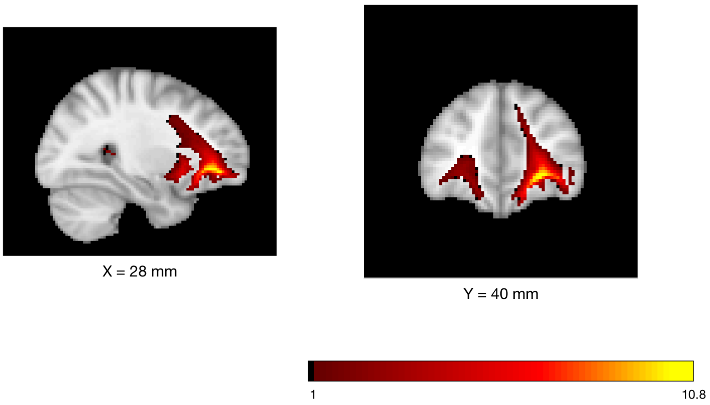


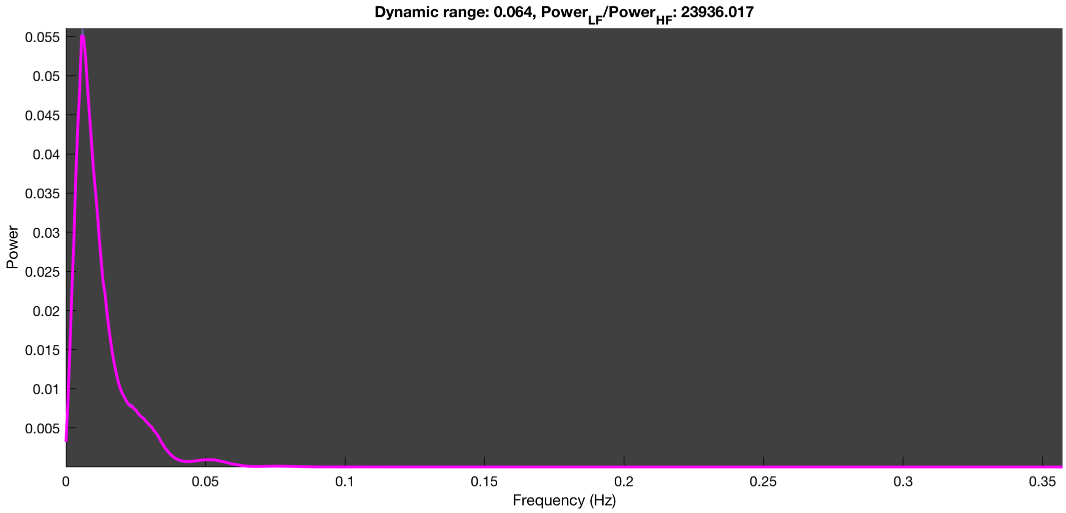

3-1 100n_tmap_component_ica_s1 068

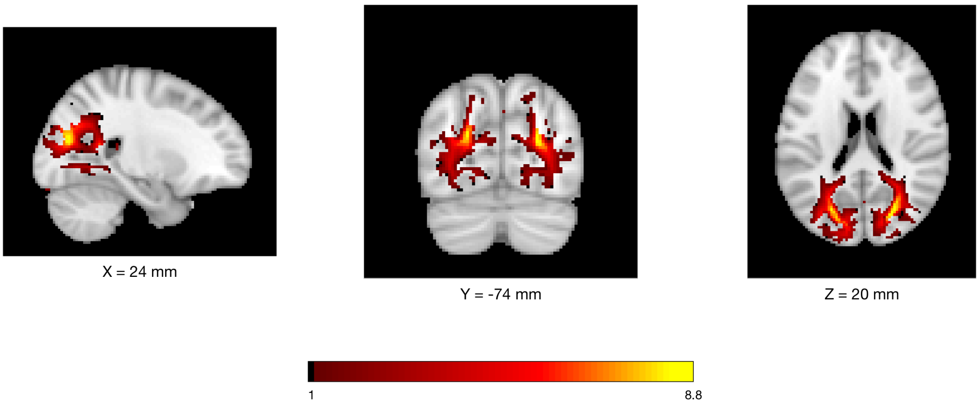


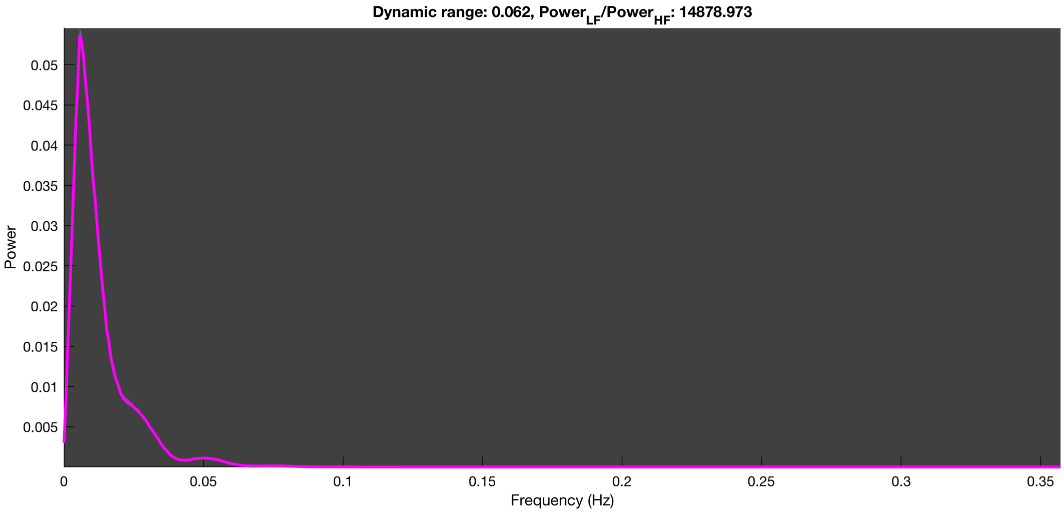

3-1 100n_tmap_component_ica_s1 069

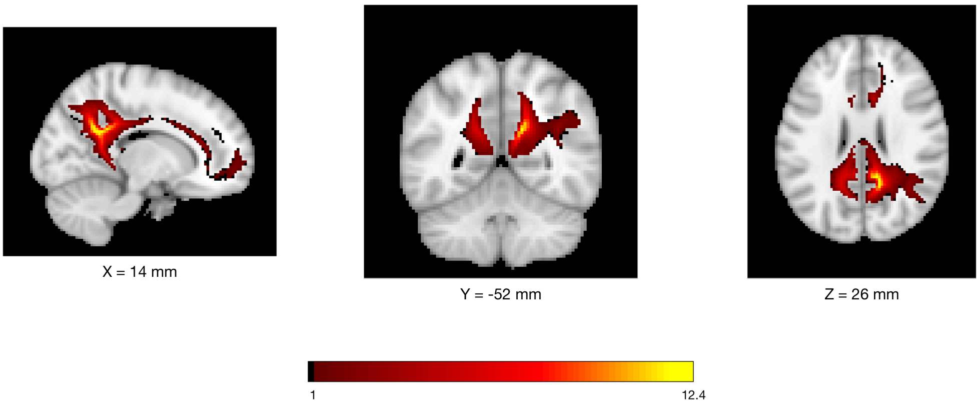


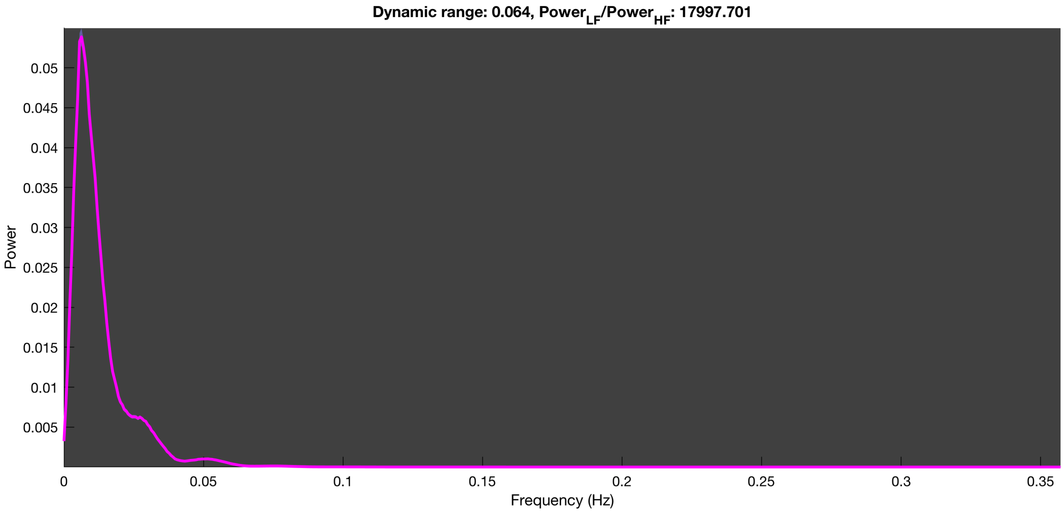

3-1 100n_tmap_component_ica_s1 070

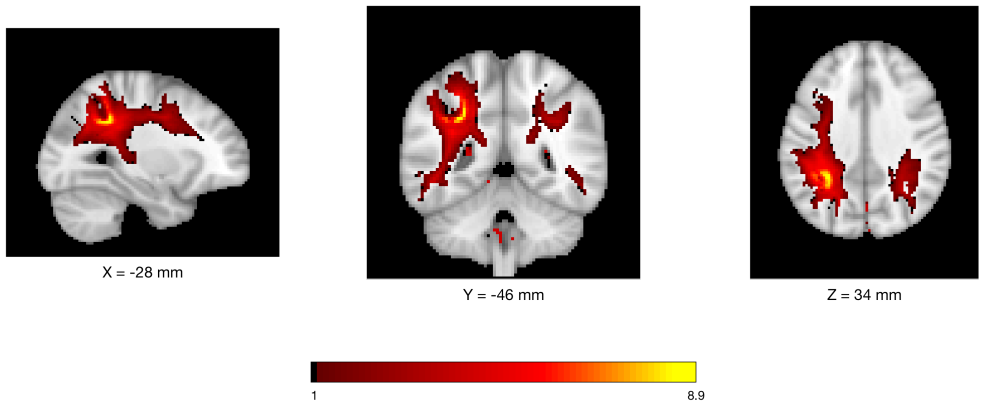


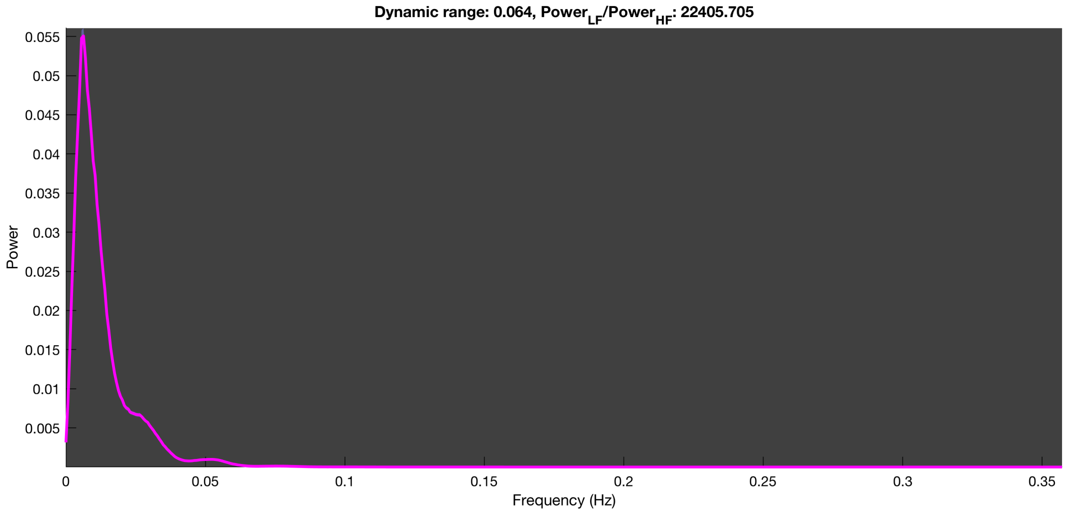

3-1 100n_tmap_component_ica_s1 071

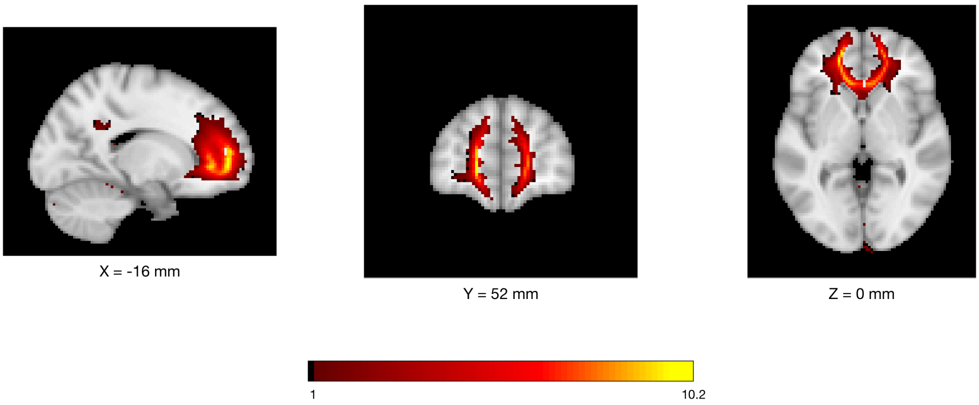


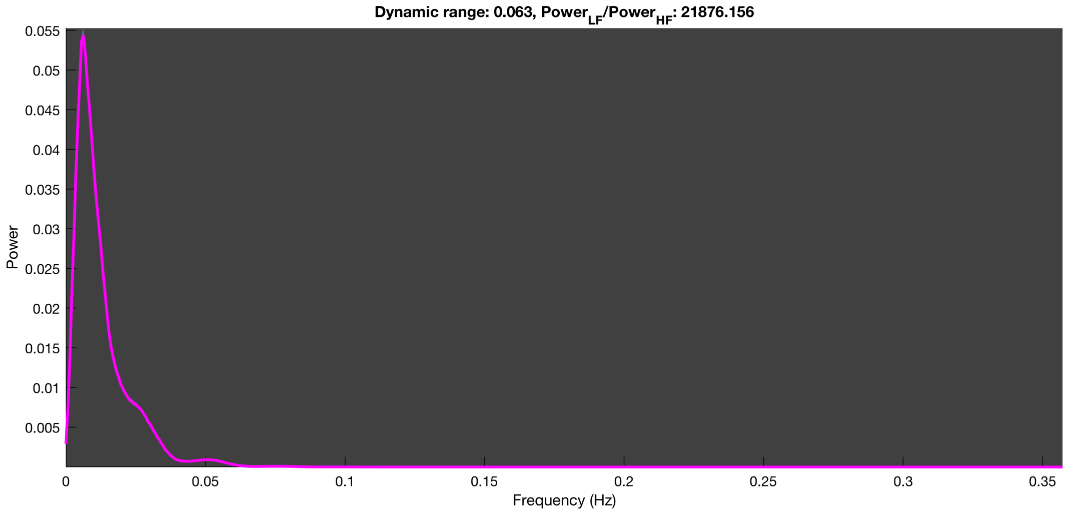

3-1 100n_tmap_component_ica_s1 072

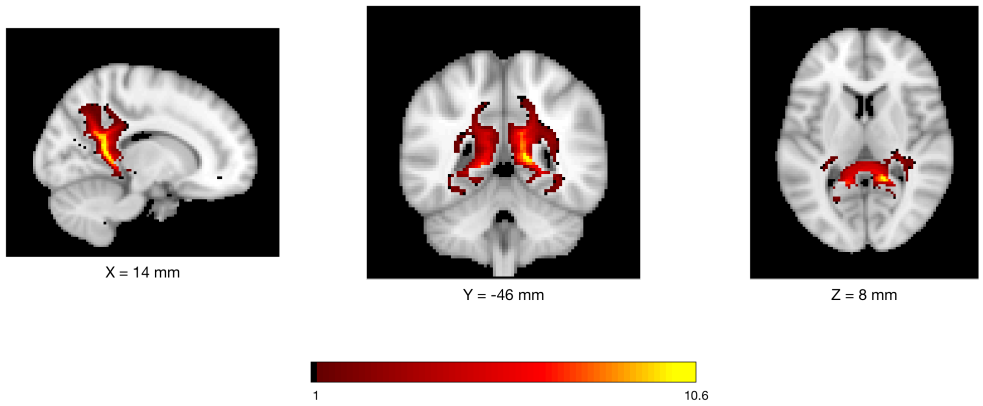


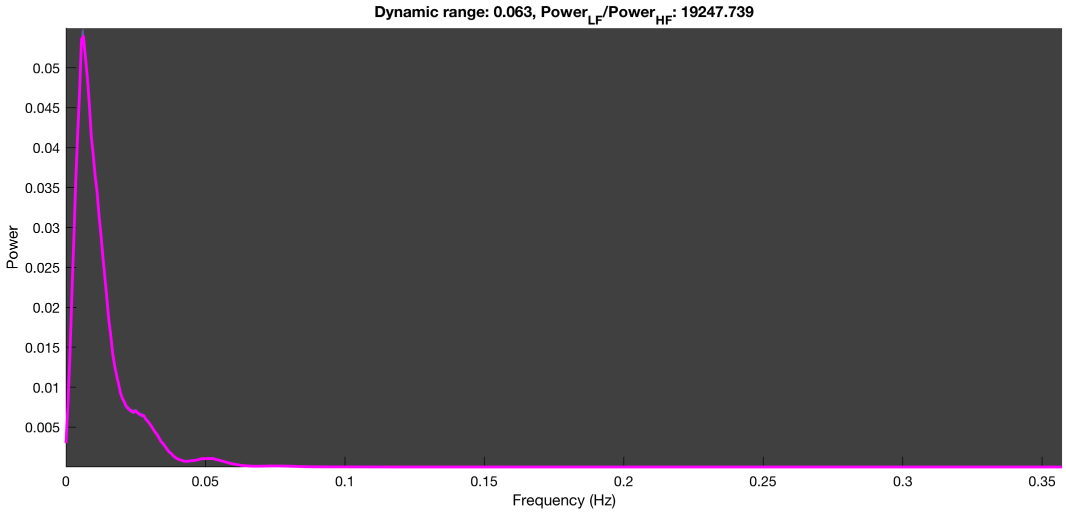

3-1 100n_tmap_component_ica_s1 073

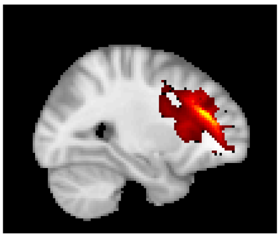


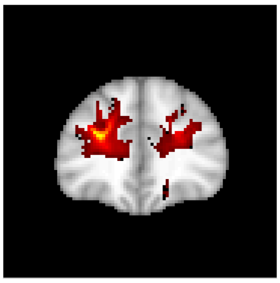


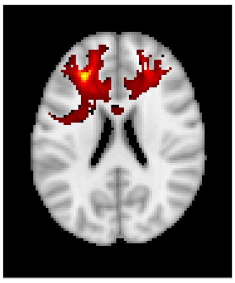


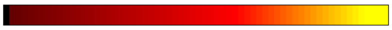


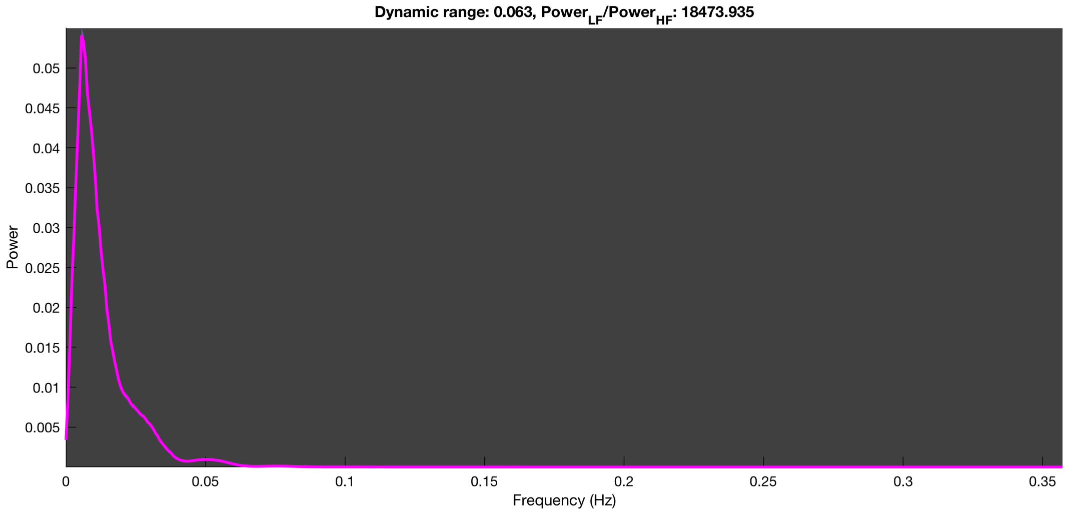

3-1 100n_tmap_component_ica_s1 074

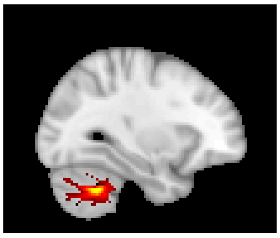

X = -30 mm

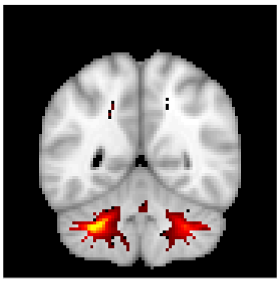

Y = -52 mm

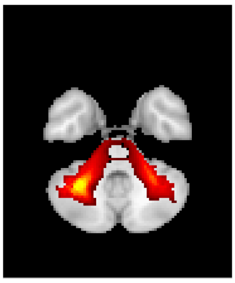

Z = -38 mm

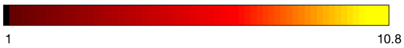


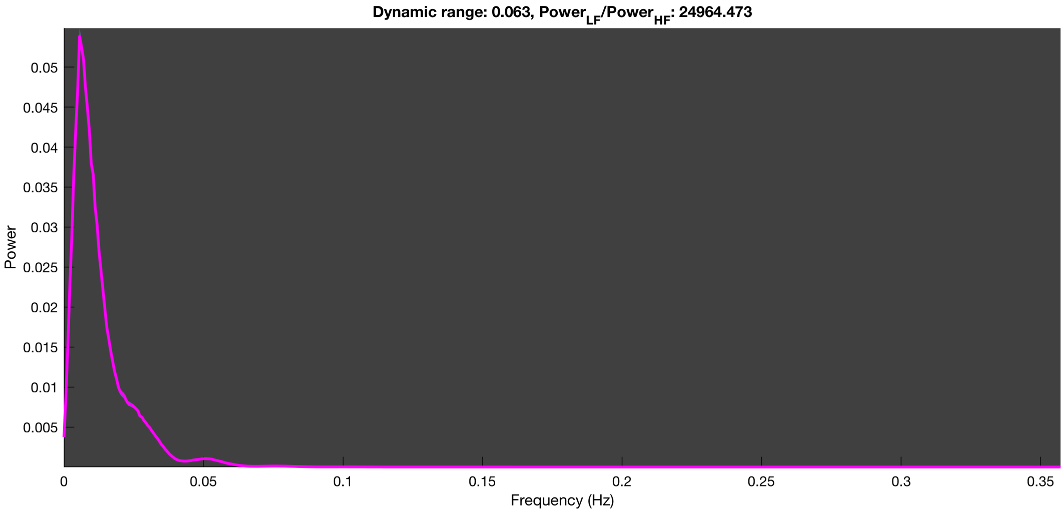

3-1 100n_tmap_component_ica_s1 075

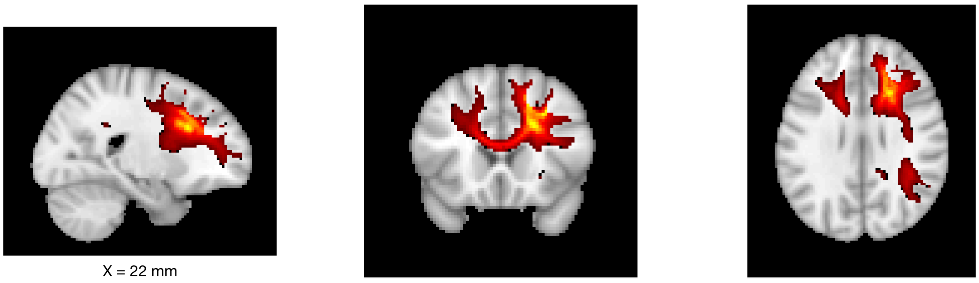

X = 22 mm Y = 20 mm Z = 28 mm

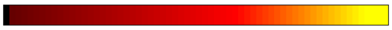


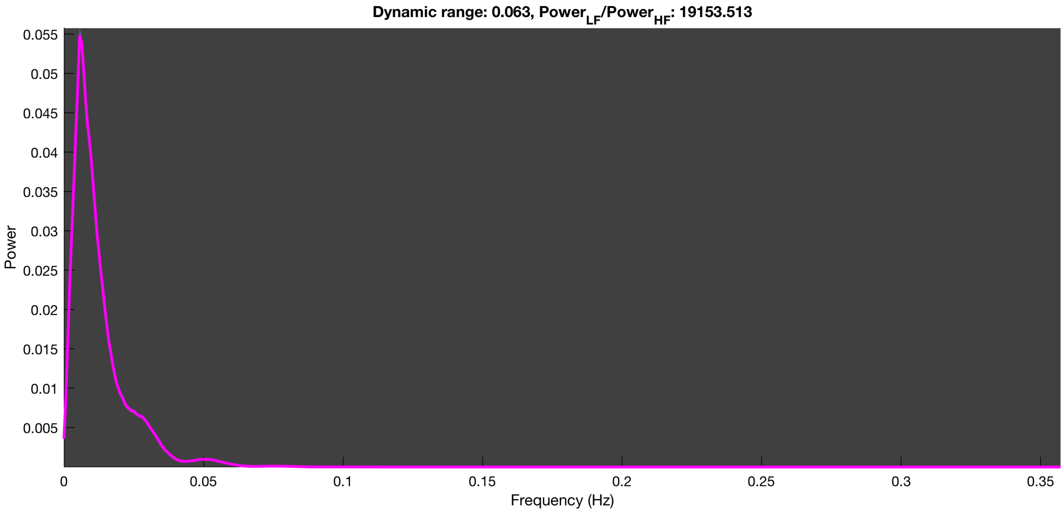

3-1 100n_tmap_component_ica_s1 076

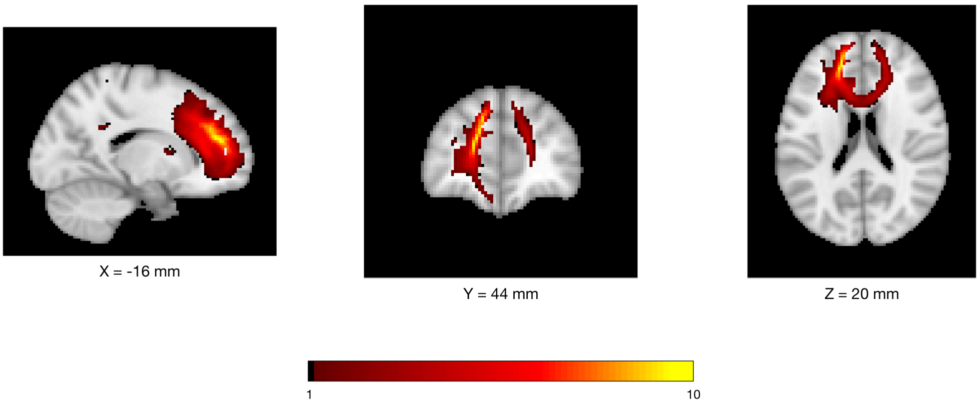


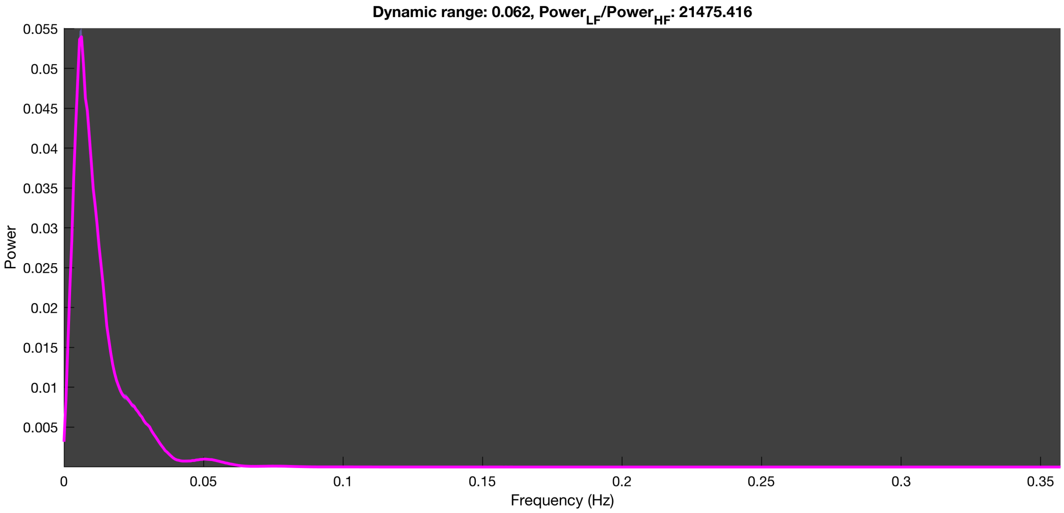

3-1 100n_tmap_component_ica_s1 077

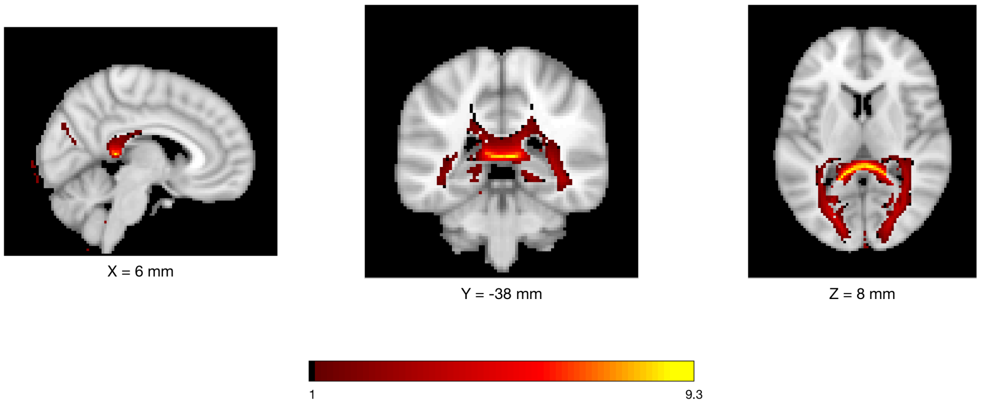


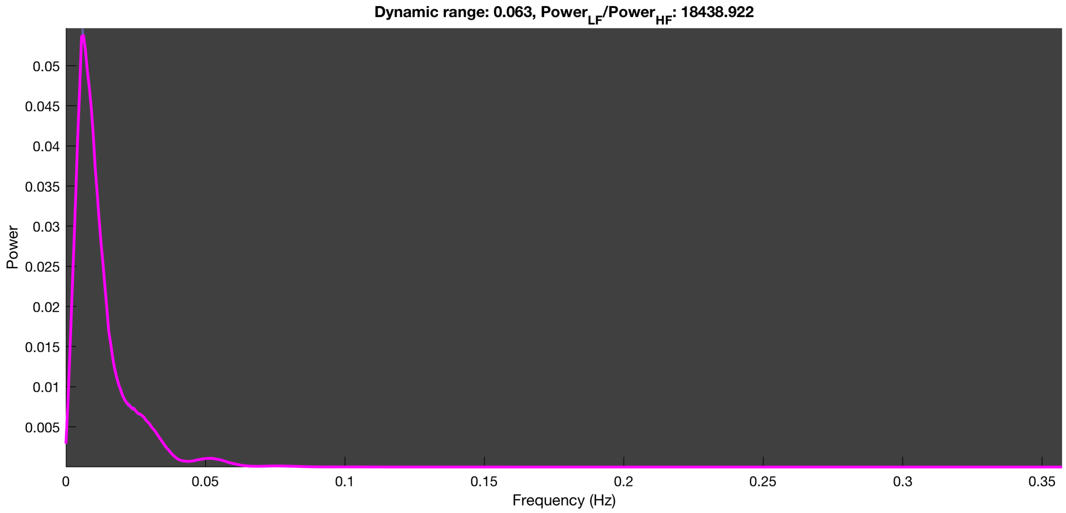

3-1 100n_tmap_component_ica_s1 078

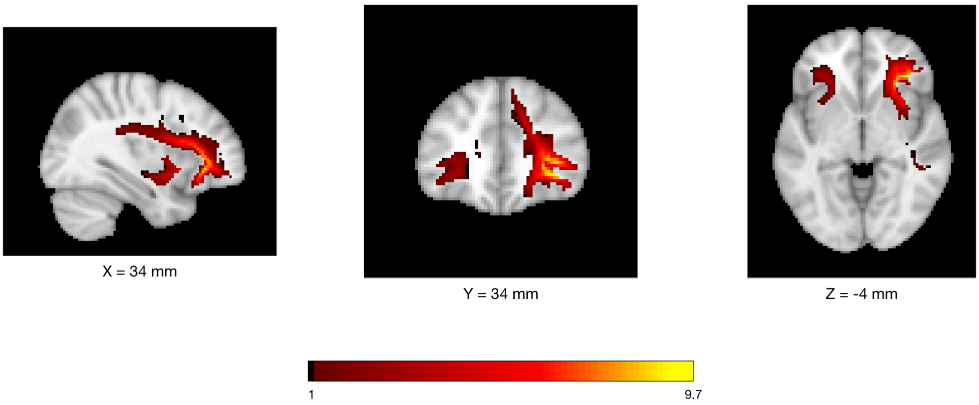


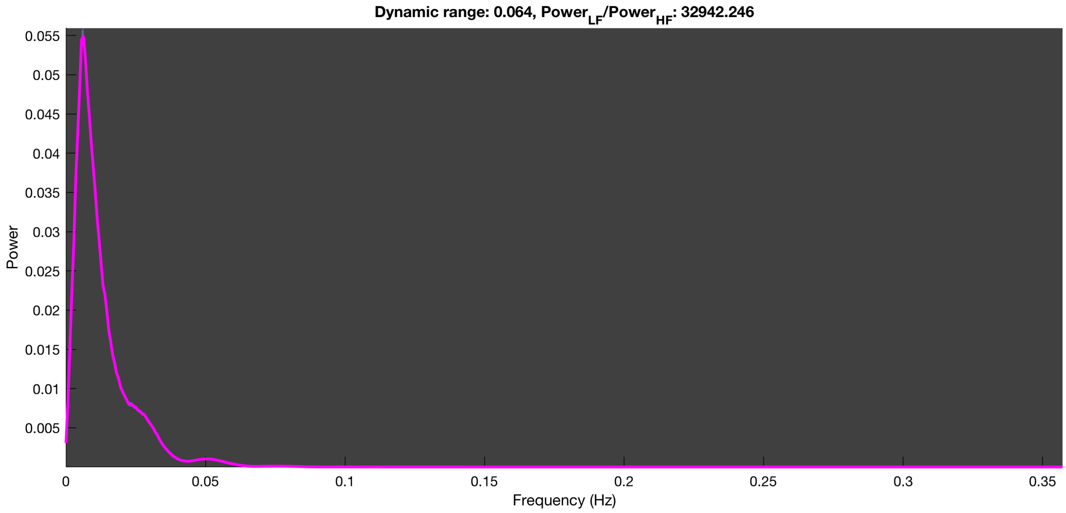

3-1 100n_tmap_component_ica_s1 079

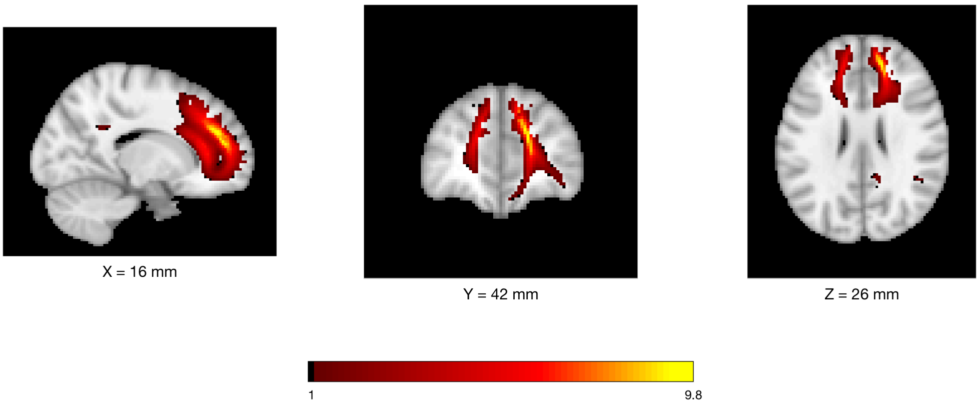


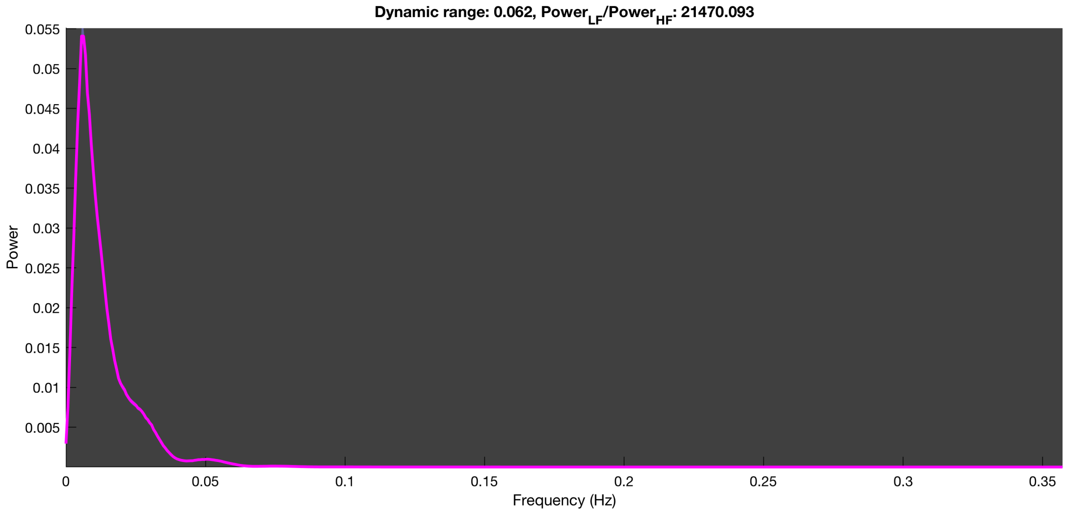

##### 3-1 100n_tmap_component_ica_s1 080

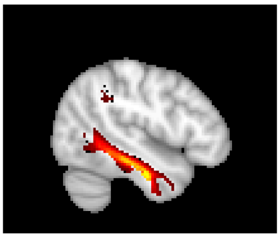

X = -46 mm

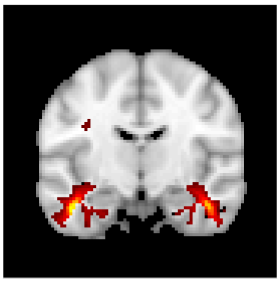

Y = -12 mm

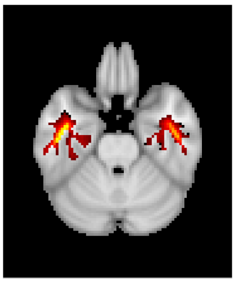

Z = -26 mm

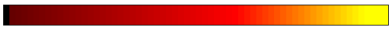


Dynamic range: 0.062, $Power_{LF}/Power_{HF}$: 21508.921

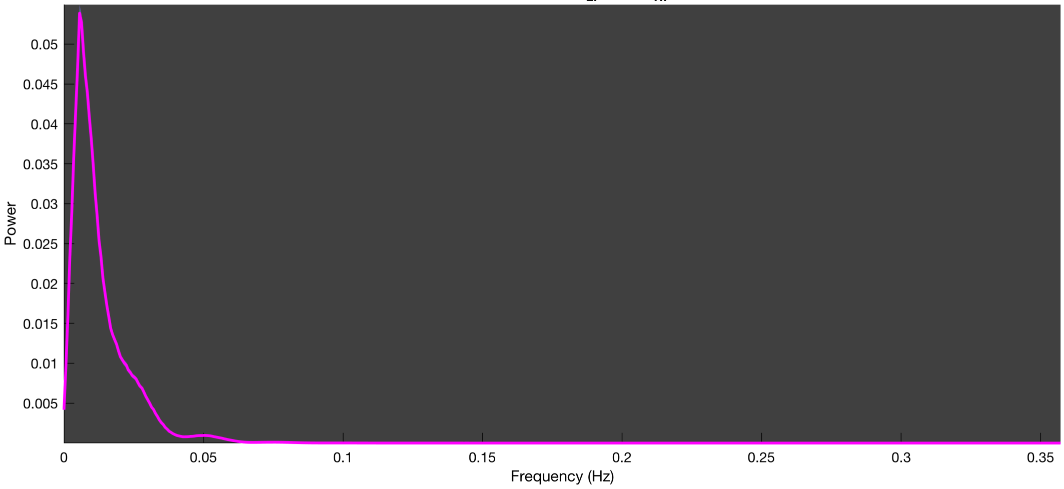

3-1 100n_tmap_component_ica_s1 081

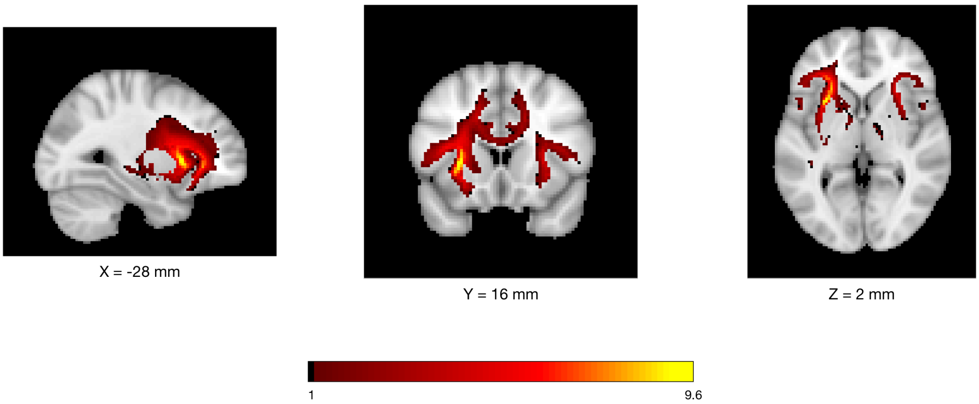


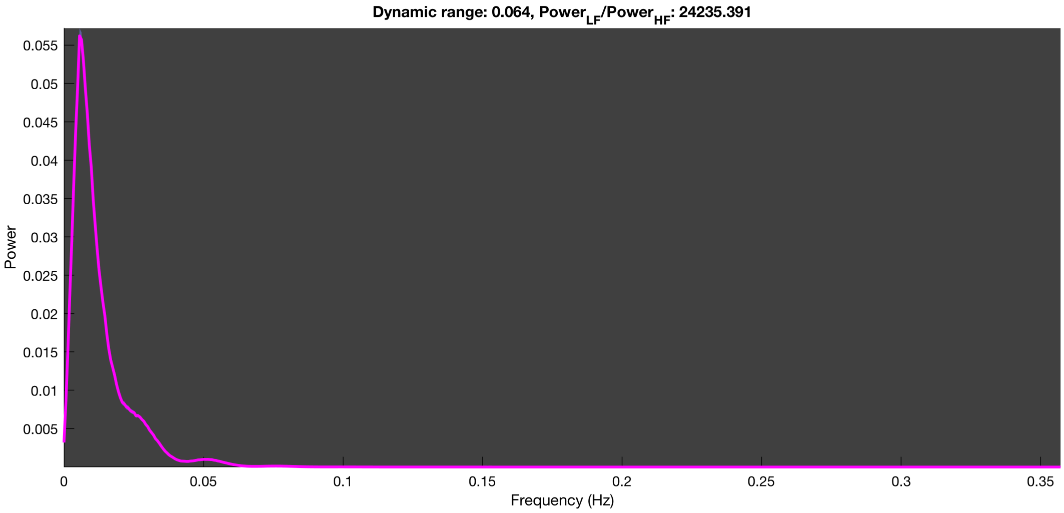

3-1 100n_tmap_component_ica_s1 082

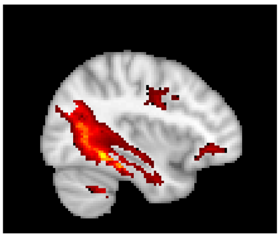

X = -36 mm

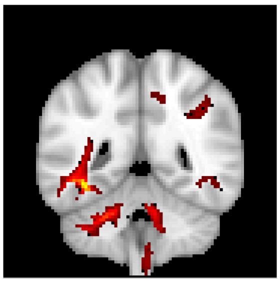

Y = -46 mm

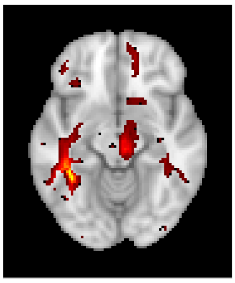

Z = -12 mm

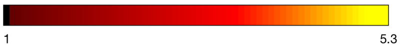


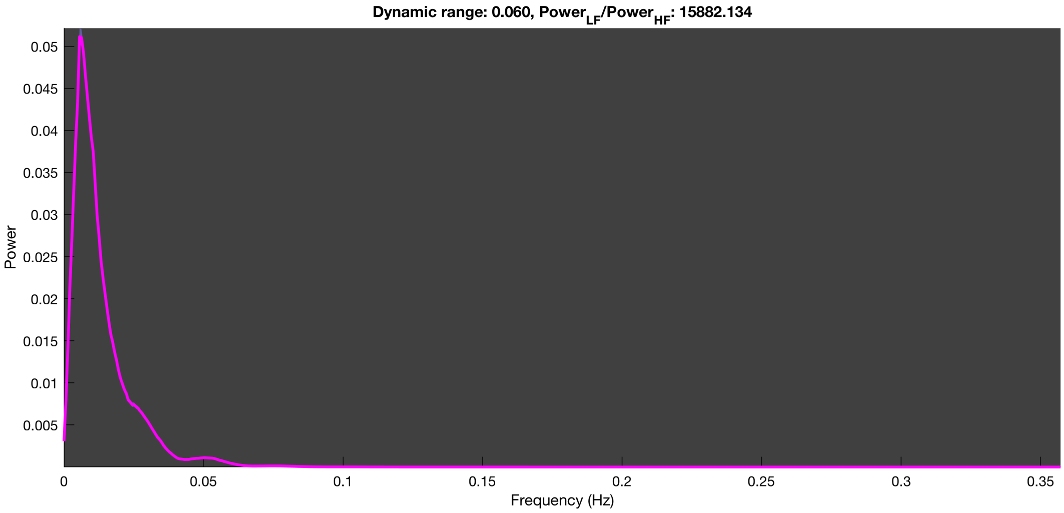

##### 3-1 100n_tmap_component_ica_s1 083

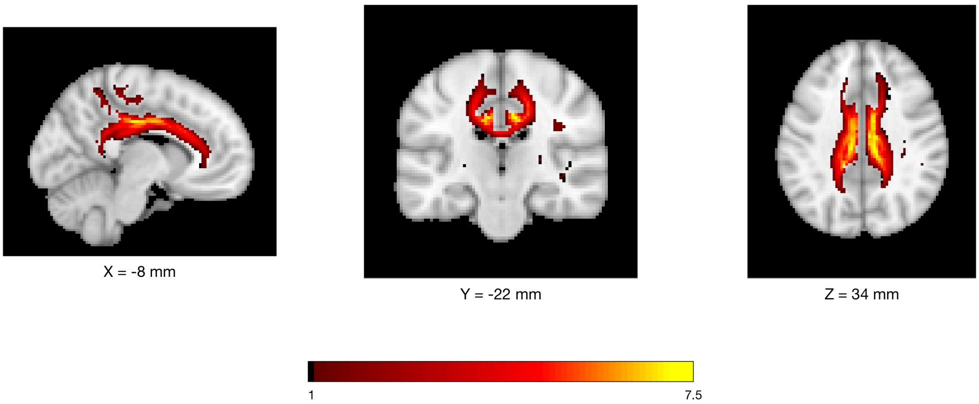


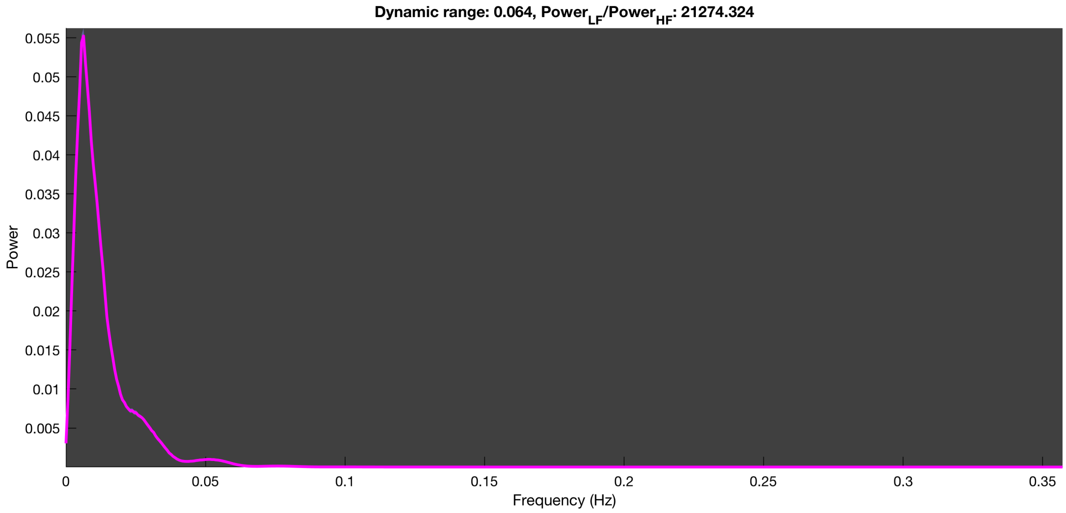

3-1 100n_tmap_component_ica_s1 084

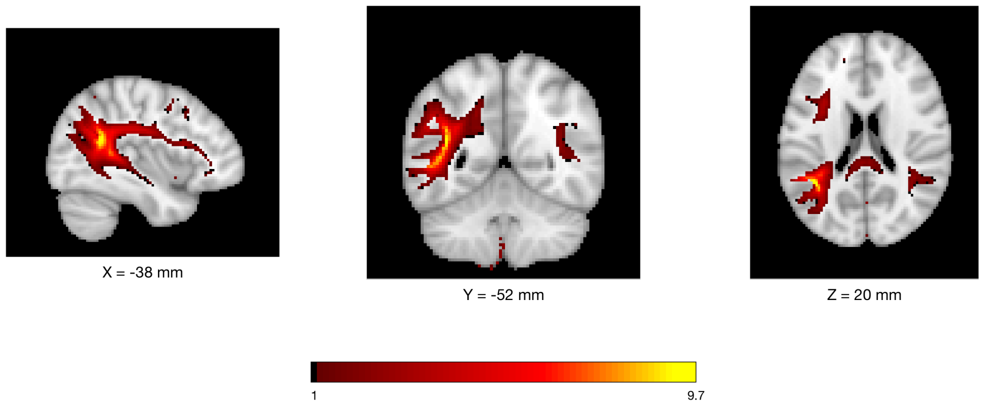


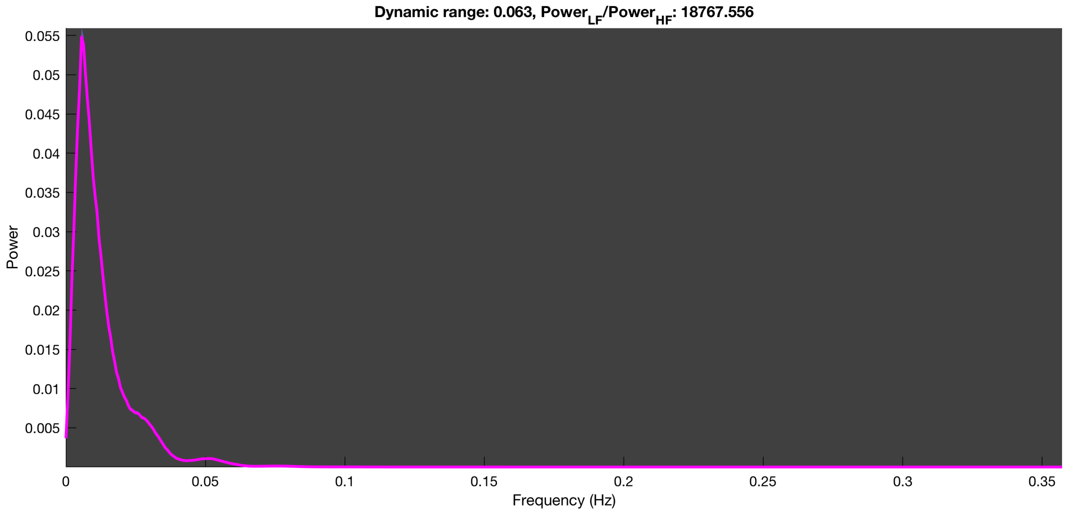

3-1 100n_tmap_component_ica_s1 085

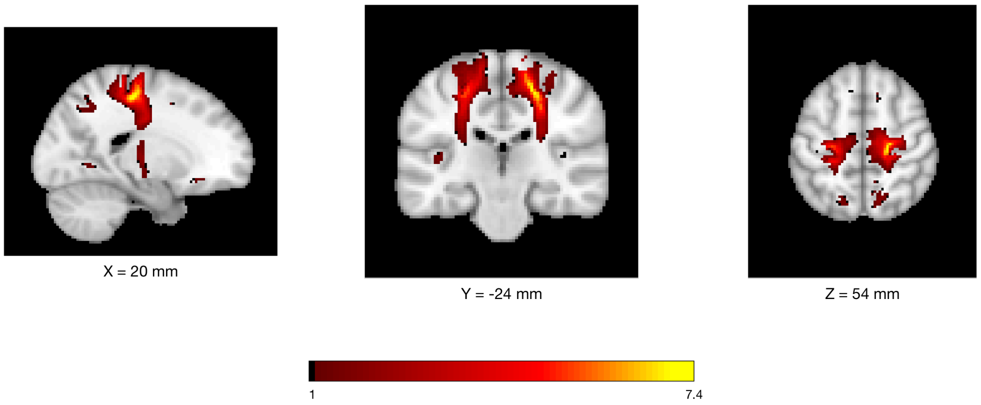


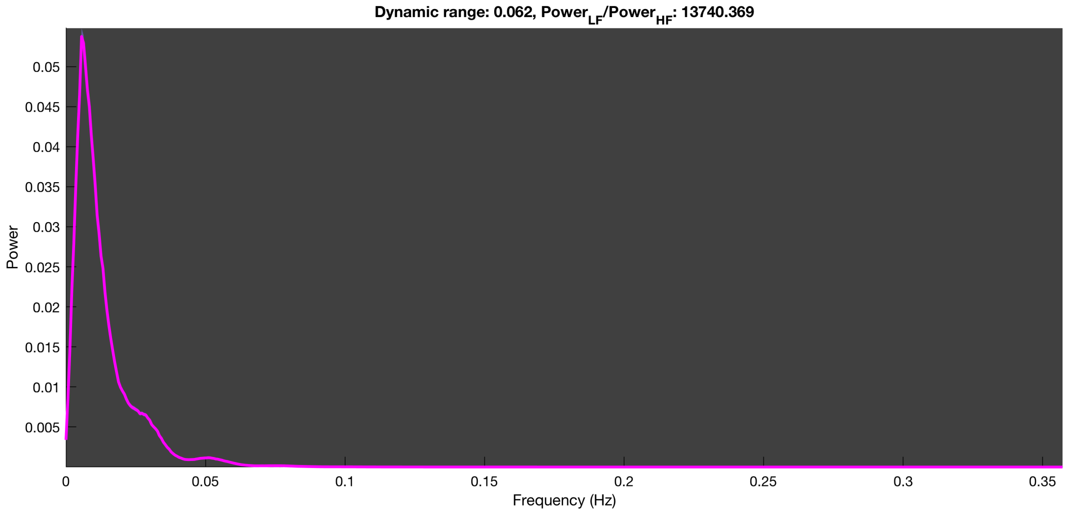

3-1 100n_tmap_component_ica_s1 086

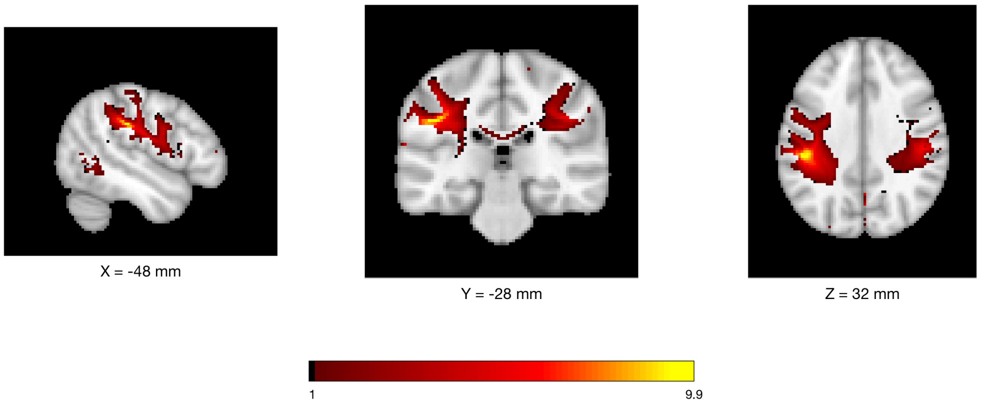


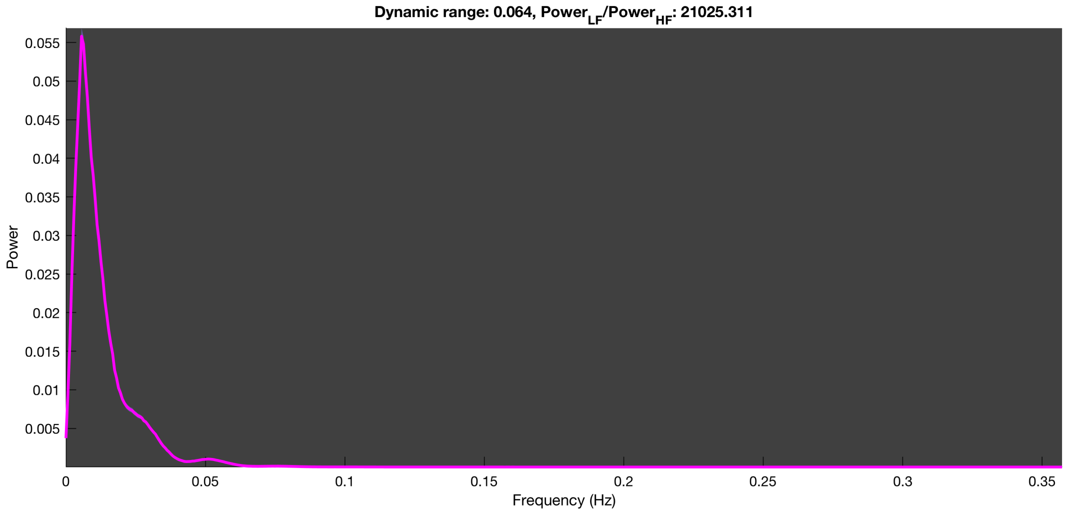

3-1 100n_tmap_component_ica_s1 087

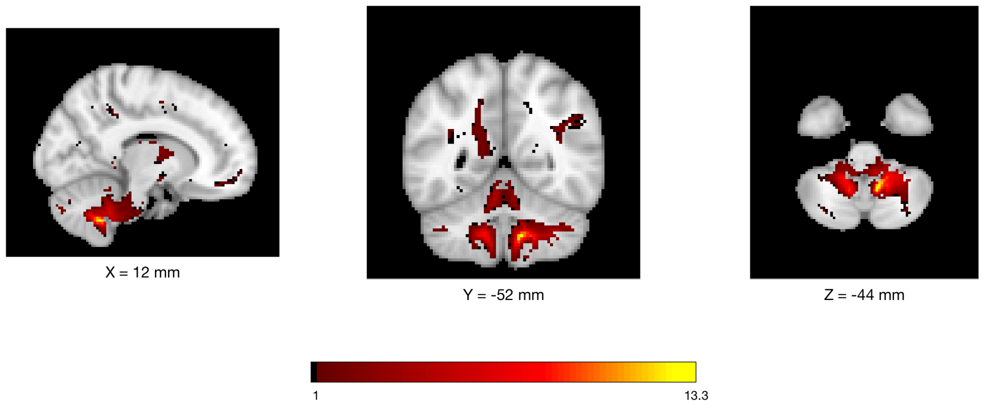


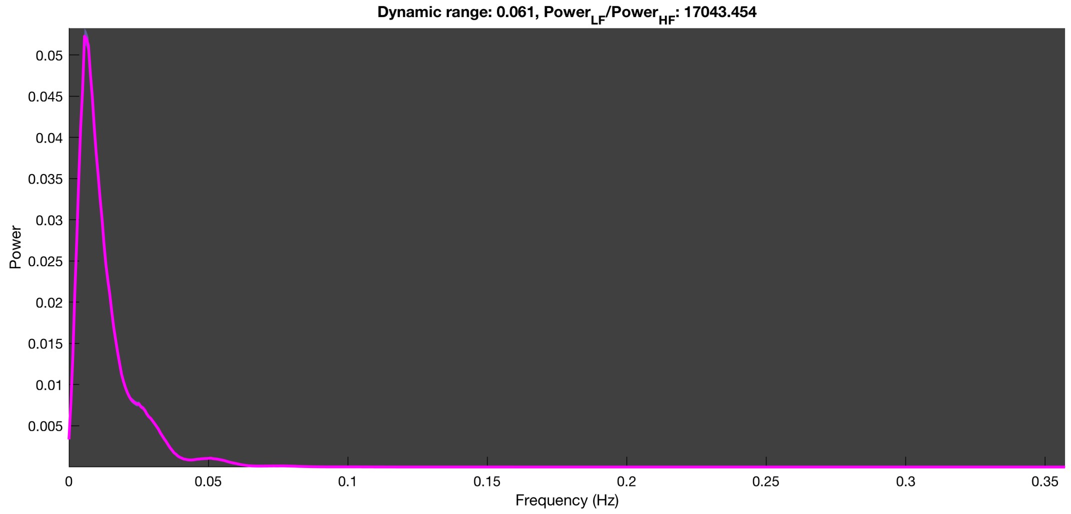

3-1 100n_tmap_component_ica_s1 088

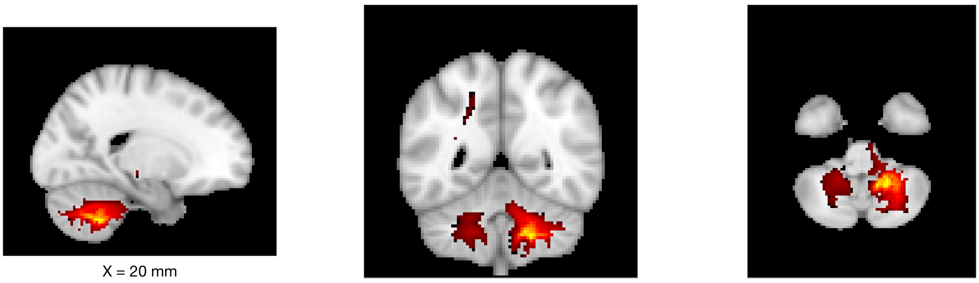

X = 20 mm

Y = -50 mm

Z = -44 mm

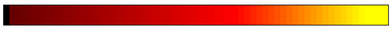


Dynamic range: 0.064, $Power_{LF}/Power_{HF}$: 19040.646

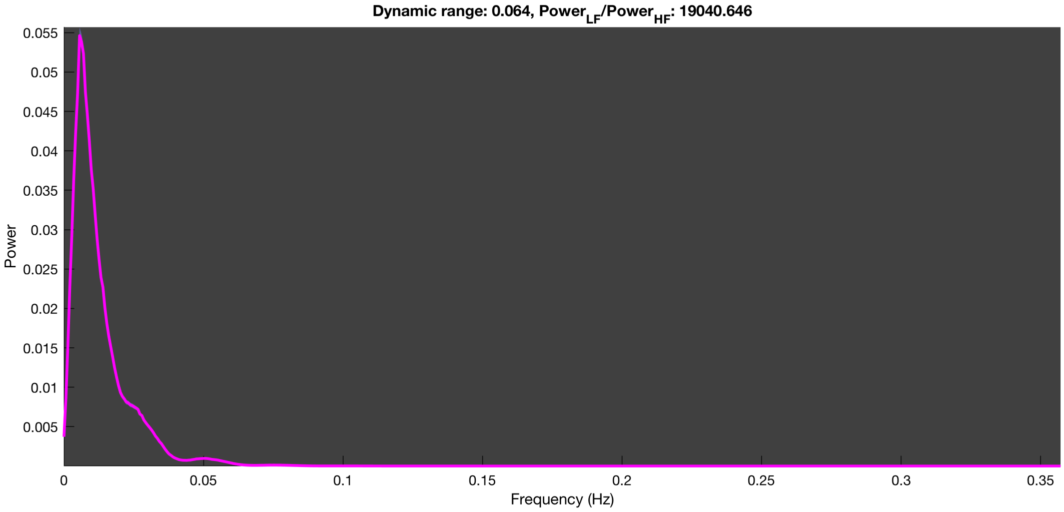

3-1 100n_tmap_component_ica_s1 089

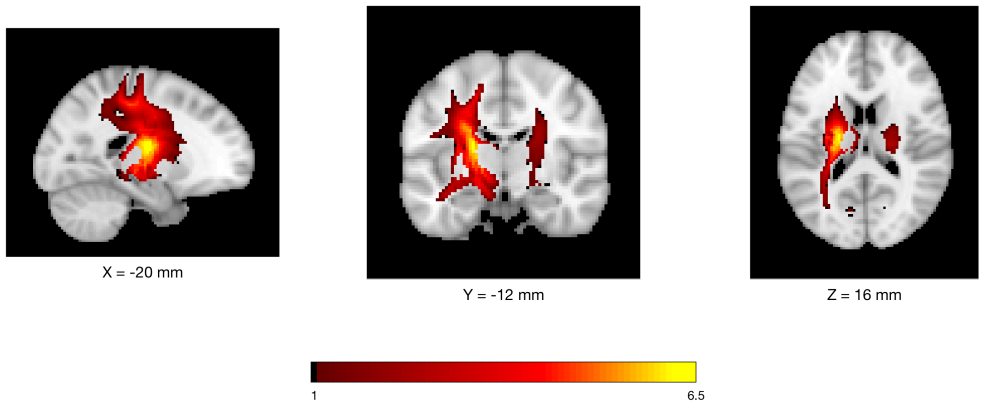


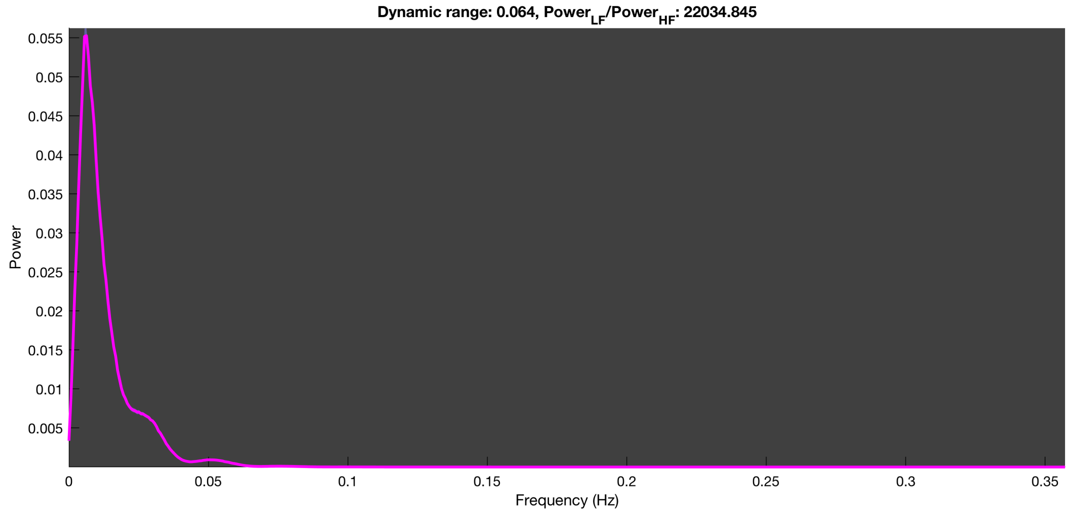

3-1 100n_tmap_component_ica_s1 090

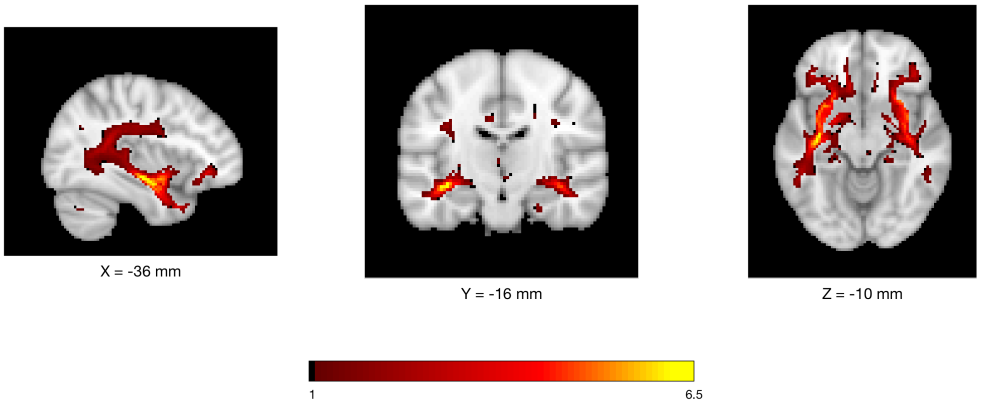


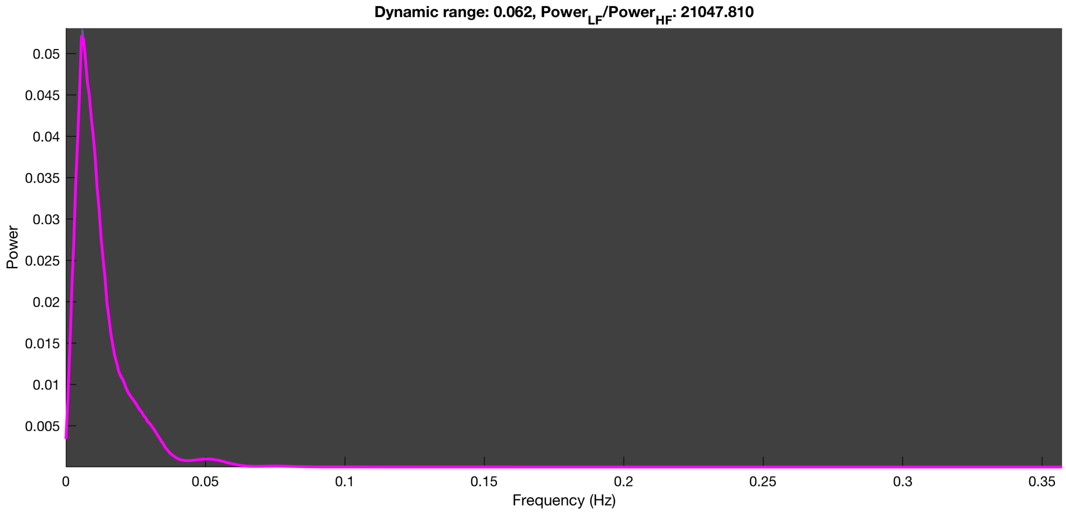

3-1 100n_tmap_component_ica_s1 091

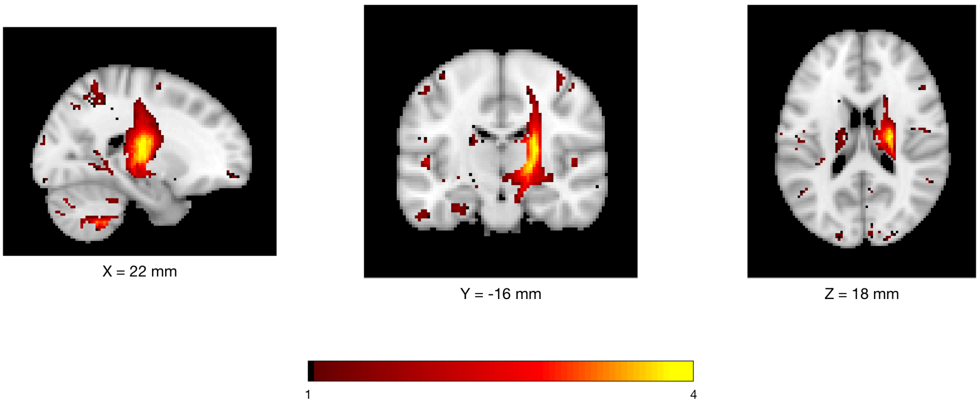


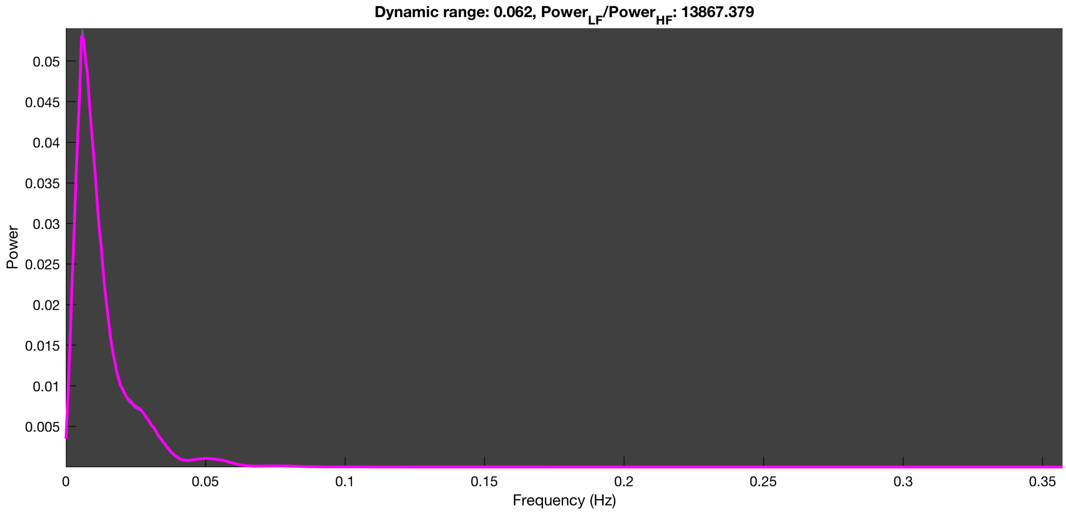

3-1 100n_tmap_component_ica_s1 092

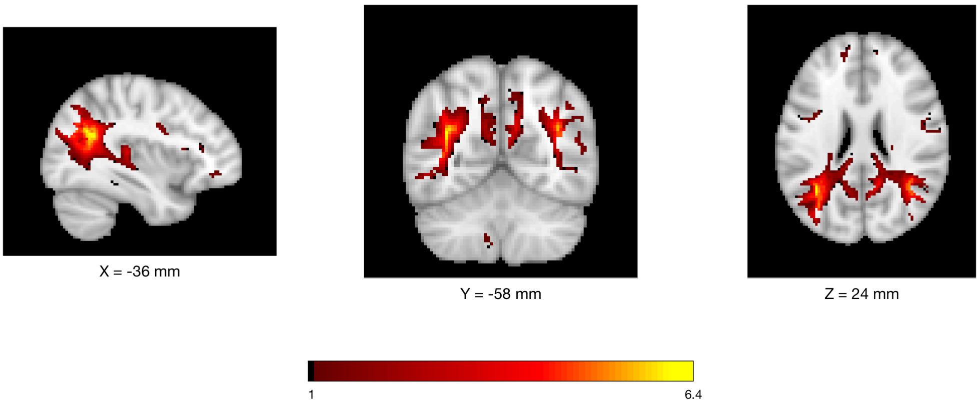


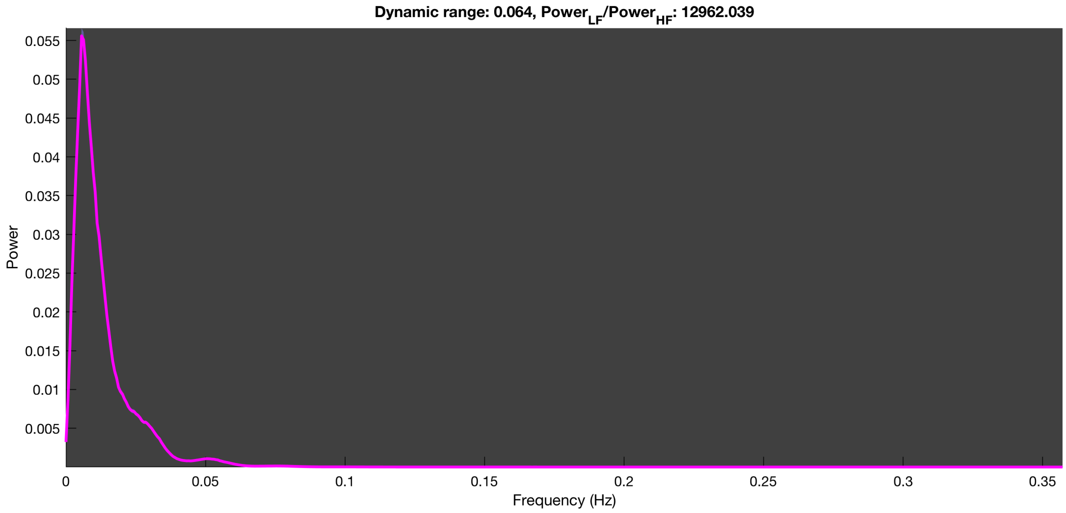

**3-1 100n_tmap_component_ica_s1 093**

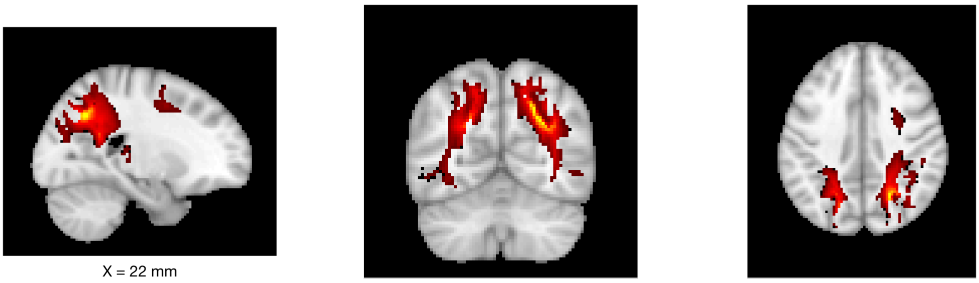

X = 22 mm Y = -62 mm Z = 38 mm

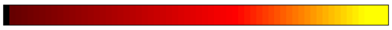


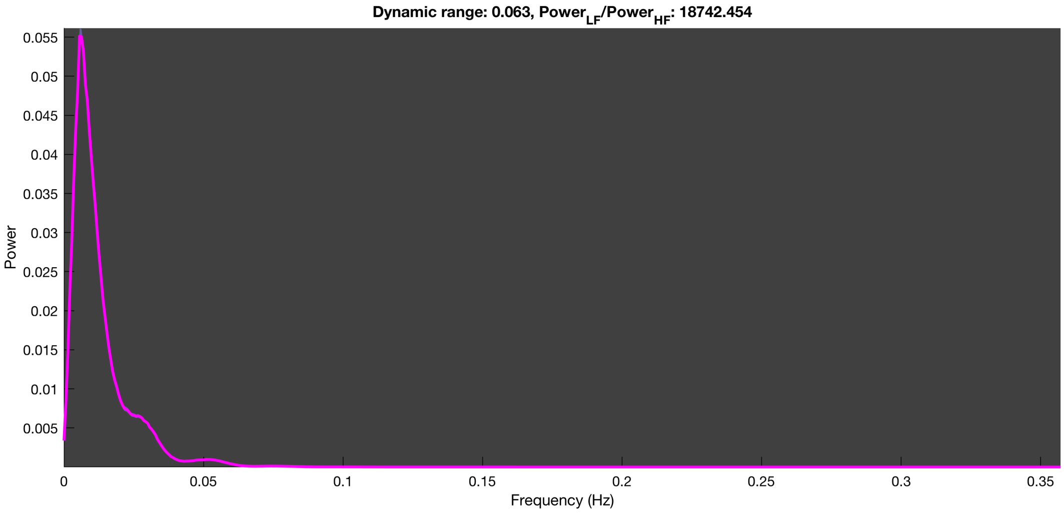

3-1 100n_tmap_component_ica_s1 094

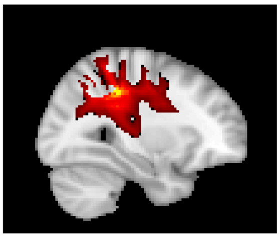

X = 30 mm

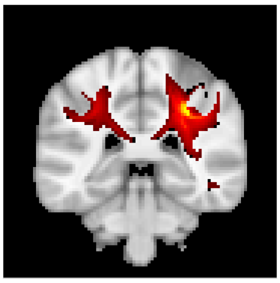

Y = -36 mm

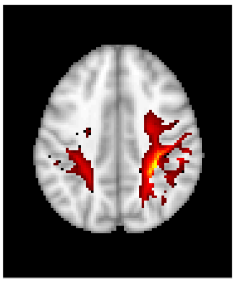

Z = 40 mm

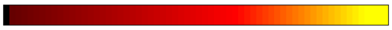


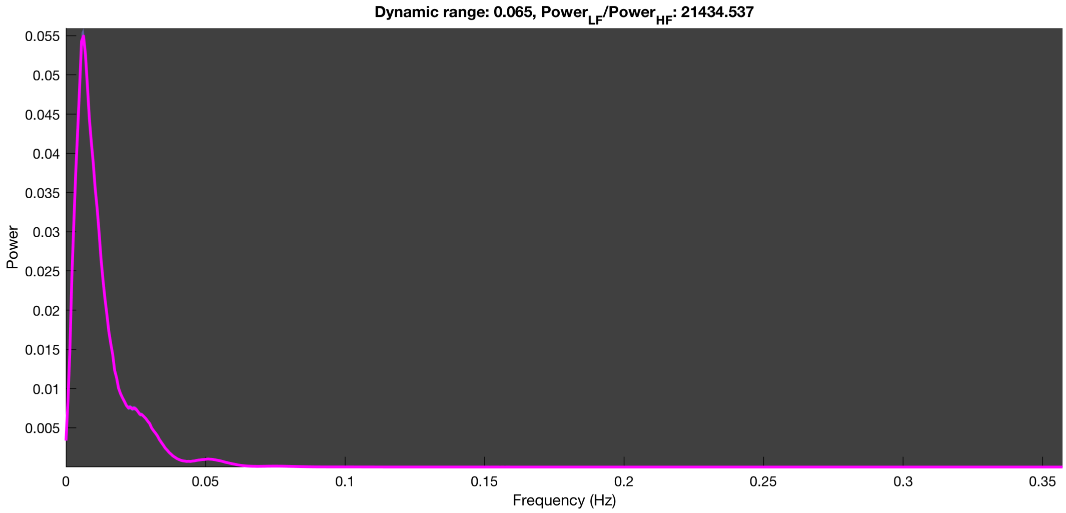

3-1 100n_tmap_component_ica_s1 095

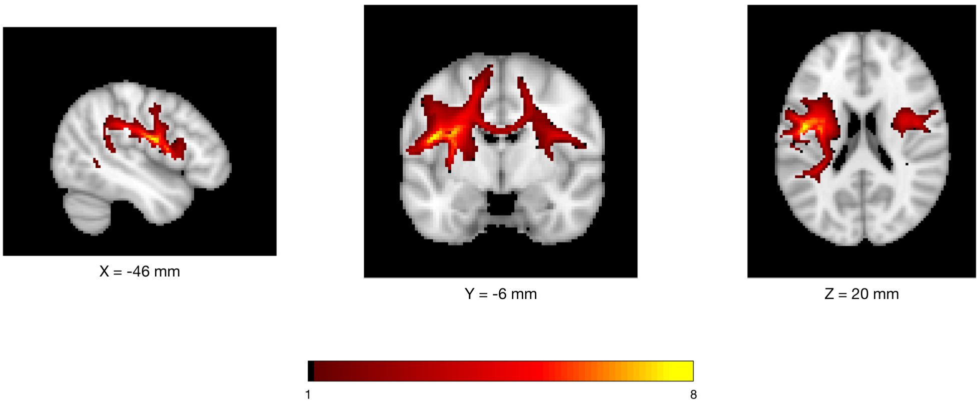


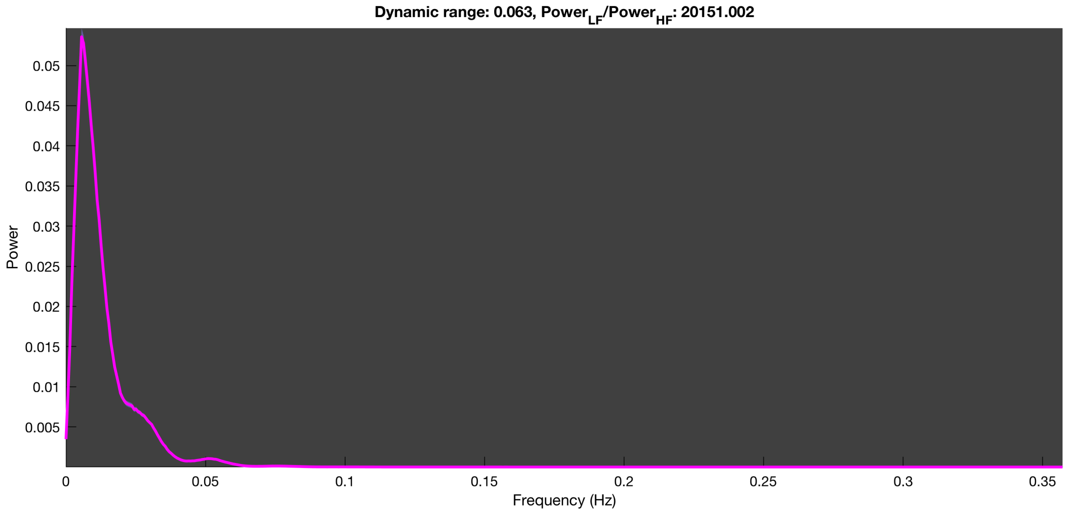

3-1 100n_tmap_component_ica_s1 096

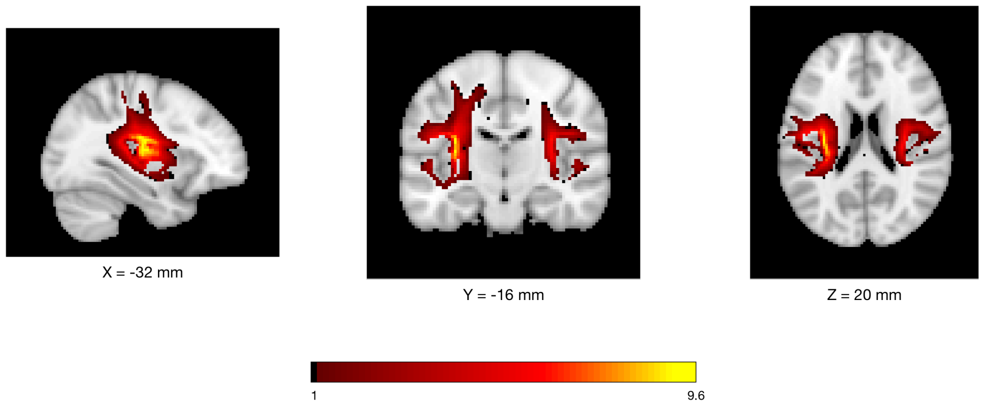


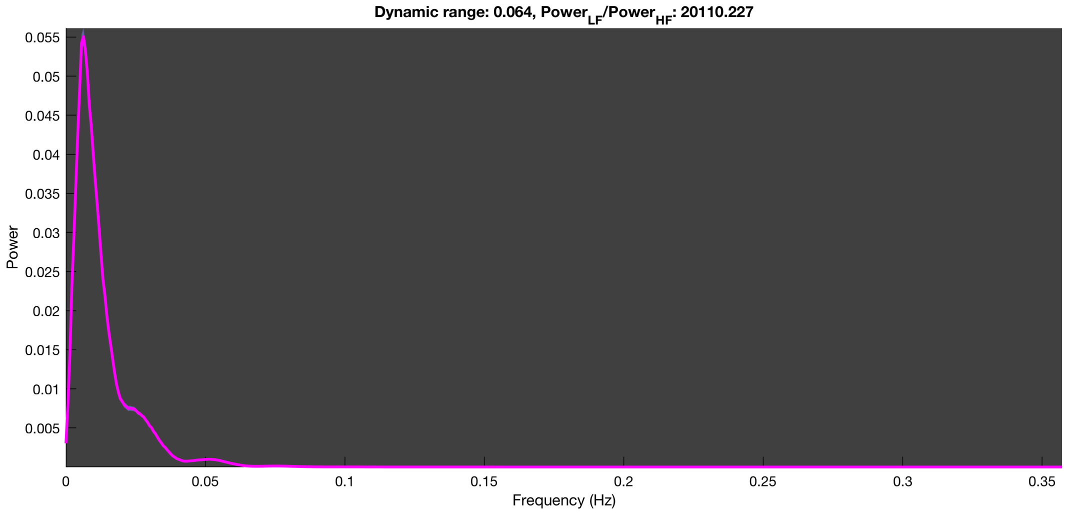

3-1 100n_tmap_component_ica_s1 097

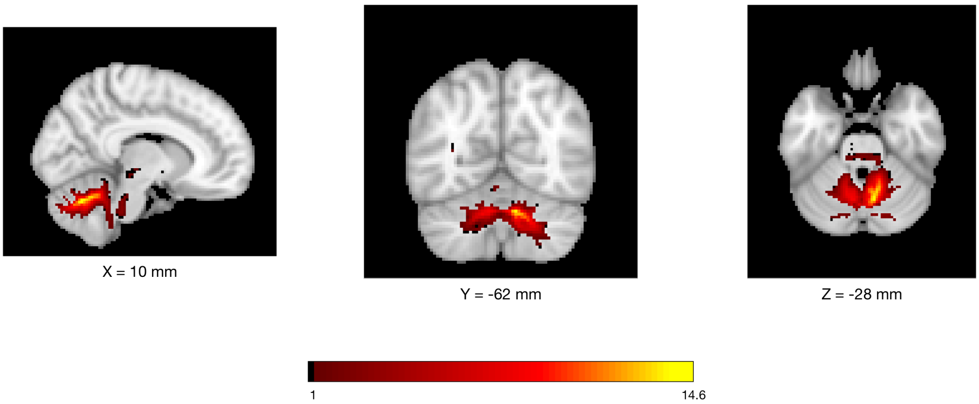


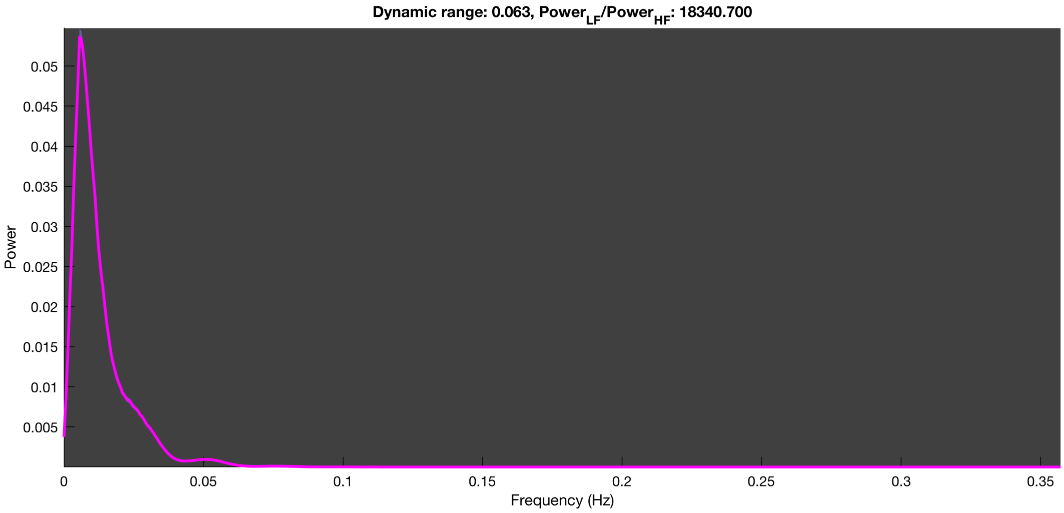

3-1 100n_tmap_component_ica_s1 098

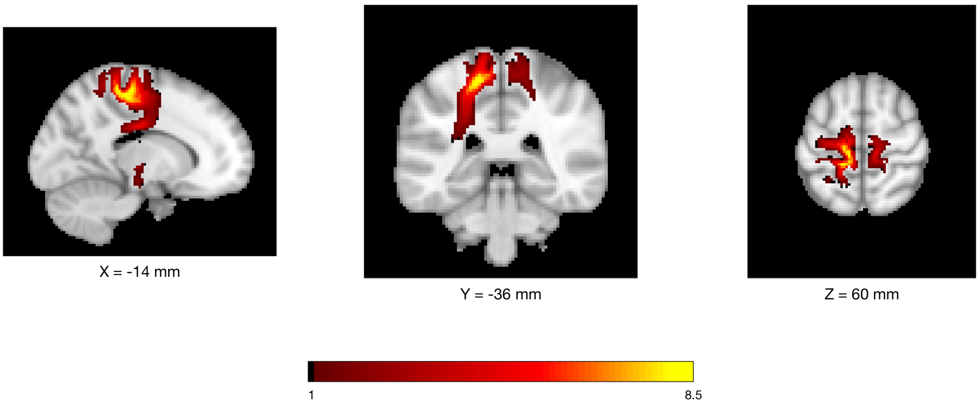


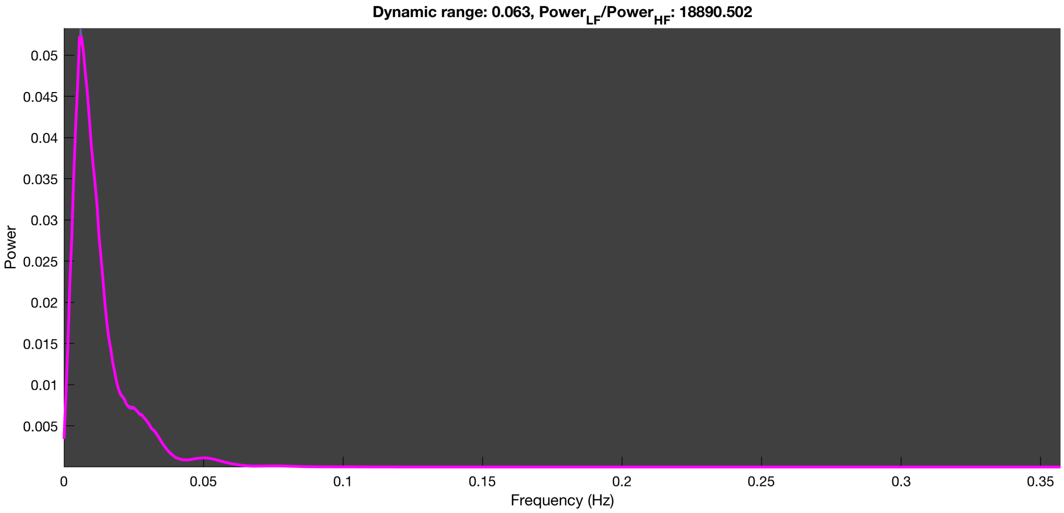

3-1 100n_tmap_component_ica_s1 099

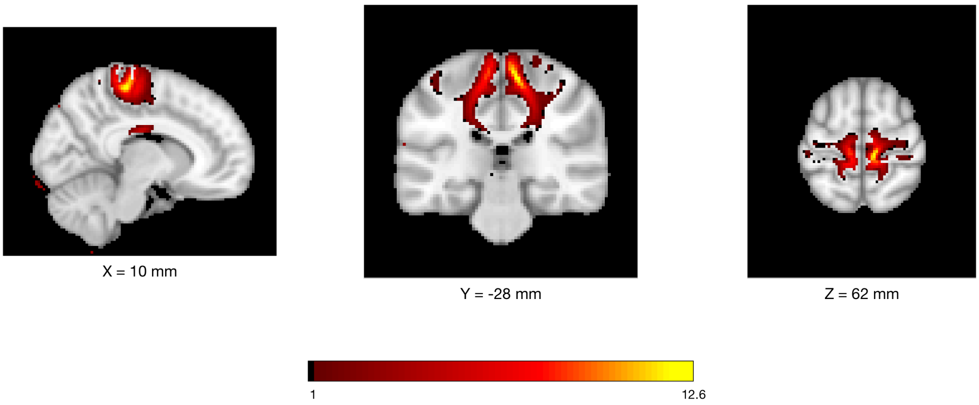


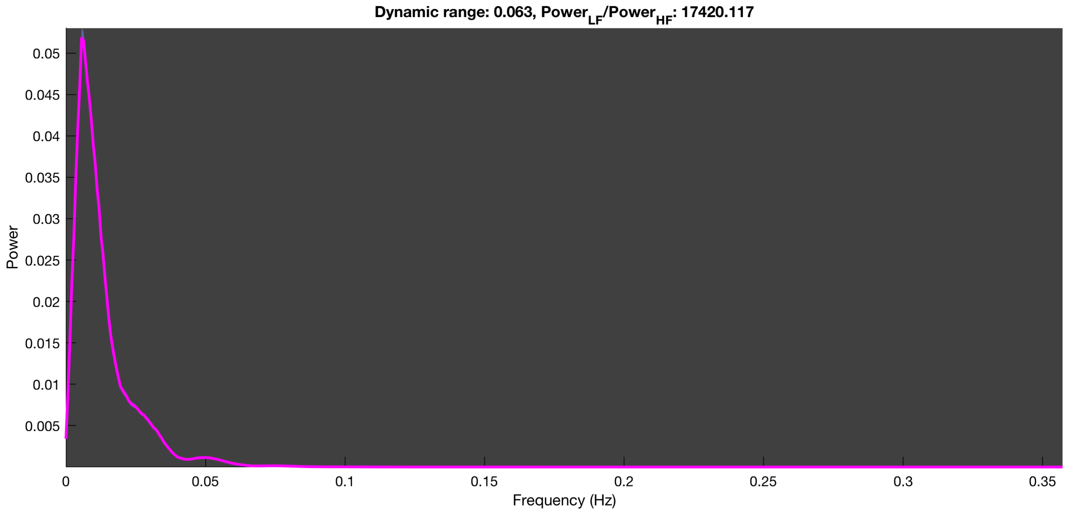

3-1 100n_tmap_component_ica_s1 100

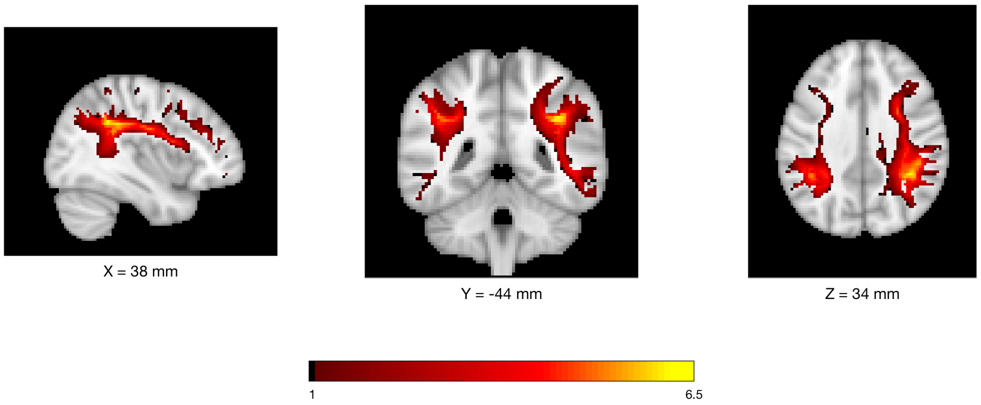


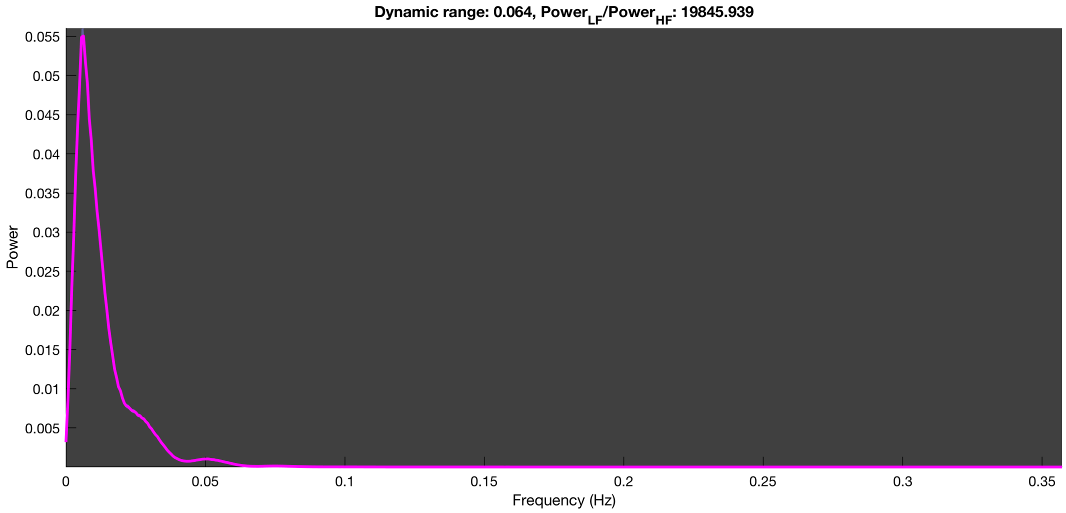
